# Histone methylation has a direct metabolic role in human cells

**DOI:** 10.1101/2023.04.22.537846

**Authors:** Marcos Francisco Perez, Peter Sarkies

## Abstract

The N-terminal tails of eukaryotic histones are frequently post-translationally modified. The role of these modifications in transcriptional regulation is well-documented. However, the extent to which the enzymatic process of histone post-translational modification itself contributes to metabolic regulation is less clear. Here we investigated the metabolic role of histone methylation using metabolomics, proteomics and RNA-seq data from cancer cell lines, primary tumour samples and healthy tissue samples. In cancer the transcription of histone methyltransferases was inversely correlated to the activity of NNMT, an enzyme previously characterised as a methyl sink that disposes of excess methyl groups carried by the universal methyl donor S-adenosyl methionine (SAM or AdoMet). In healthy tissues histone methylation was inversely correlated to the levels of an alternative methyl sink, PEMT. These associations affected the levels of multiple histone marks on chromatin genome-wide but had no detectable impact on transcriptional regulation. We show that histone methyltransferases with a variety of different associations to transcription are co-regulated by the Retinoblastoma (Rb) tumour suppressor in human cells. Total HMT expression is increased in Rb-mutant cancers, and this leads to *NNMT* downregulation. Together, our results suggest a direct metabolic role for histone methylation in SAM homeostasis, independent of transcriptional regulation.

## INTRODUCTION

The discovery of strong associations between the transcriptional states of genes and methylation of histones at these loci was a seminal moment in the study of gene regulation (*1*). Soon the concept of the histone code was born (*2*), whereby it was imagined that in the near future knowledge of the particular combination of epigenetic marks in chromatin would allow a deterministic prediction of gene expression, just as the deciphering of the genetic code had allowed precise prediction of gene products. However, two decades later, despite strong correlations of some histone marks with specific transcriptional states (*3*), these associations can be ambiguous (*4, 5*), while evidence of causal links between histone marks and transcriptional activation or repression remains equivocal (*6*).

In that time, it has become widely appreciated that histone modifications can be influenced by cellular metabolism (*7*). Histone methylation is influenced by the availability of S-adenosyl methionine (SAM, also abbreviated as AdoMet or SAMe), the universal methyl donor that is required for cellular methylation of lipids, proteins, nucleic acids and metabolites, and which can be modulated by dietary methionine supplementation (*8*). However, the abundance of histones in the cell offers the potential for histone modifications to impact metabolism, by acting as sinks for metabolic intermediates such as SAM (*9*). Methyl sinks may play key metabolic roles, acting to buffer the ratio of SAM to S-adenosyl homocysteine (SAH, also abbreviated as AdoHcy) and supporting the synthesis of important sulphur-containing metabolites such as cysteine and glutathione via the SAH-dependent transsulphuration pathway.

Here we discovered strong negative relationships between the total expression of histone methyltransferases and distinct established methyl sinks in cancer and healthy tissues. We show that these relationships affected genome-wide levels of histone post-translational modifications but did not have significant consequences for transcriptional regulation. We show that histone methyltransferases were co-expressed and regulated by E2F transcription factors in cancer. Our data indicates that histone methylation has an important metabolic role in SAM homeostasis in healthy human tissues and tumours, independent of a role in transcriptional regulation.

## RESULTS

### Histone methyltransferase expression correlates to cellular metabolite levels

We set out to investigate a potential link between metabolism and histone methylation. We reasoned that effects of histone methyltransferase (HMT) activity on metabolism might result in correlations between HMT levels and cellular metabolite concentrations. To investigate this possibility, we used a publicly-available metabolomics dataset consisting of 225 metabolites profiled by liquid chromatography-mass spectrometry (LC-MS) across 911 cell lines from the Cancer Cell Line Encyclopedia (CCLE), representing more 23 distinct cancer types (*10*). We related metabolite levels to the normalized expression of HMTs in the same cell line. We curated a list of 38 HMTs (**Table S1**) and examined the correlation of each HMT to all metabolites. Across this set 1-methylnicotinamide (1MNA) consistently emerged as the metabolite most strongly associated to histone methyltransferases, with an FDR < 0.05 for 18 individual HMTs, a geometric mean FDR of 0.001 (**Table S2**) and an average Pearson’s correlation of-0.090 (range-0.261 to 0.160). Indeed, 1MNA was by far the metabolite that (anti-)correlated most strongly with the total level of HMTs obtained by summing the expression of the 38 individual enzymes (**Fig 1A**; Pearson’s correlation =-0.274, FDR = 1.77 × 10^-14^).

**Fig 1.**
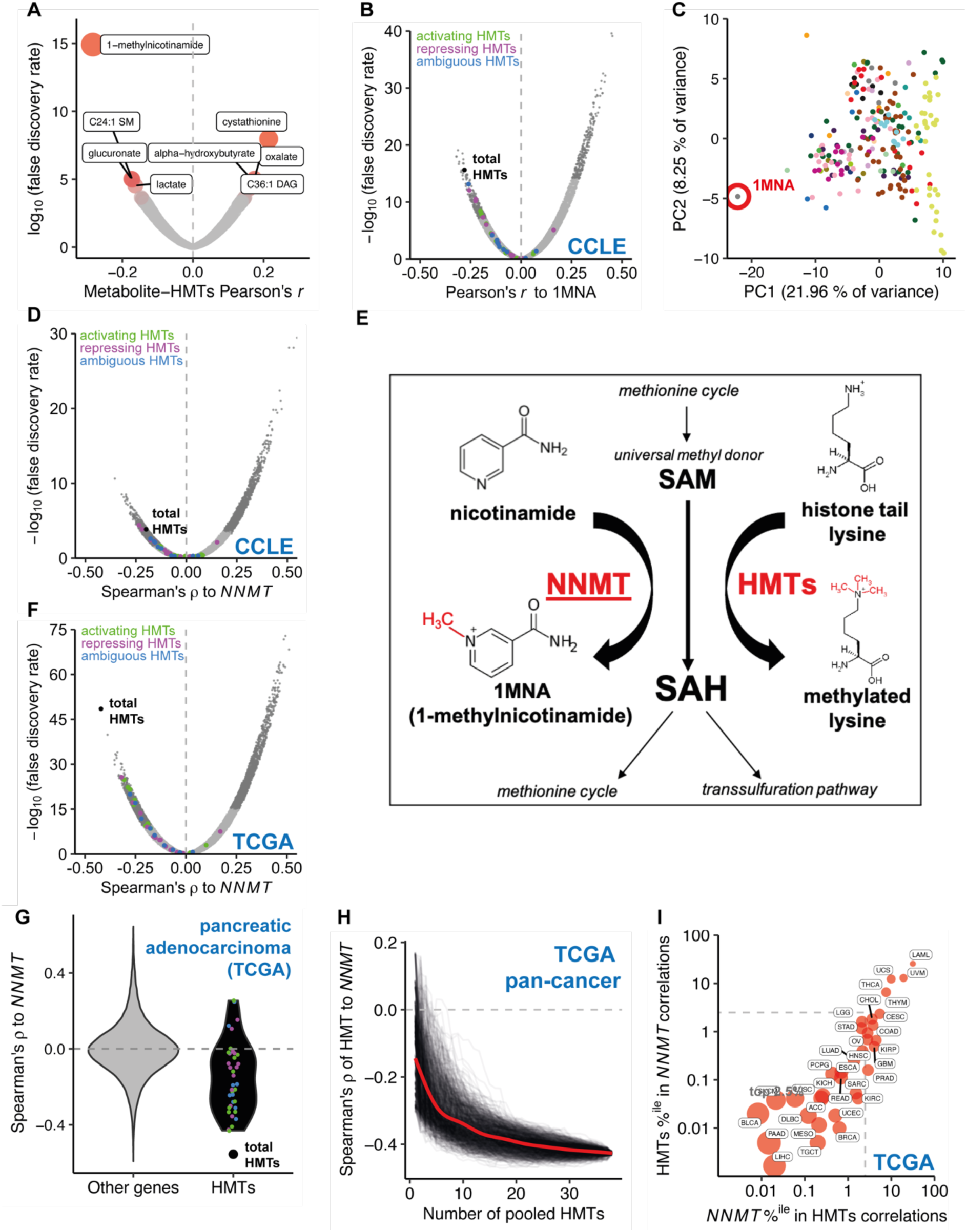
Total histone methyltransferase expression is strongly anticorrelated with the activity of NNMT in cancers. A) Volcano plot showing Pearson’s correlation and FDR for 225 metabolites to total histone methyltransferase (HMT) expression (total RNA-seq median of ratios-normalised pseudocounts) across 927 cancer cell lines from the CCLE. B) Volcano plot showing Pearson’s correlation and FDR for expression of 10275 expressed genes to levels of 1-methylnicotinamide (1MNA) across 927 CCLE cancer cell lines. The top and bottom 2.5% of points are shown in darker grey. HMT-encoding genes are shown as points coloured according to their association with transcriptional activation (green), repression (magenta) or an unclear relationship (blue). Pearson’s r for total HMT expression is shown as a black point. C) Principal component analysis of metabolite levels across 927 cancer cell lines from the CCLE. 1MNA is highlighted with a red circle. D) Volcano plot showing Spearman’s correlation and false discovery rate (FDR) for expression of *NNMT* vs 52440 genes in a pan-cancer analysis of 927 CCLE cell lines across 23 cancer types. HMT-encoding genes are shown as points as in panel **1B**. E) NNMT and HMTs both convert SAM to SAH and so can affect cellular methylation potential by acting as a ‘sink’. F) Volcano plot showing Spearman’s correlation and FDR for expression of *NNMT* vs 60489 genes in a pan-cancer analysis of TCGA primary tumours across 33 cancer types. HMT-encoding genes are shown as points as in panel **1B**. G) Violin plot showing Spearman’s correlation to *NNMT* for HMTs (black, right) or other genes (left, grey) in 79 primary adrenocortical carcinoma (ACC) tumours from the TCGA. Individual HMT-encoding genes are shown as points as in panel **1B**. H) Spearman’s correlation vs *NNMT* expression of total expression of pooled HMTs added to the pool in a random order. 1000 individual iterations are shown as black lines, with the locally estimated smoothing (Loess fit) trendline shown in red. I) TCGA pan-cancer analysis showing rank percentile position of total HMTs among correlations of *NNMT* expression to 60489 genes and vice versa in 33 distinct cancer types. Bubble size is inversely proportional to the log of the ‘relative reciprocal score’, the sum of squares of the ranks of total HMTs/*NNMT* in the reciprocal distribution. The dashed grey box indicates correlations in the strongest 2.5% of anticorrelated genes.

### HMT expression varies reciprocally with the activity of the 1MNA/NNMT methyl sink

The strong relationship between 1MNA and HMT levels indicated that HMT expression variation across cancer cell lines was associated with changes in metabolism. To investigate which metabolic pathways might be responsible we performed principal component analysis (PCA) and clustering analysis on all 225 metabolites. Related metabolites from known biochemical pathways tended to cluster together. 1MNA was a clear outlier in both analyses (**Fig 1C, Fig S1**), indicating that 1MNA synthesis reflects a discrete metabolic process. Indeed, 1MNA is known to be a stable metabolic end-product that has no downstream metabolites in cancer (*11*) and is excreted from cells in healthy tissues (*12*).

1MNA is the product of methylation of nicotinamide by the enzyme nicotinamide N-methyltransferase (NNMT). 1MNA levels were strongly correlated with *NNMT* expression as measured by RNA-seq (**Fig S2A, Table S2**) and with NNMT protein levels as measured by quantitative mass spectrometry (**Fig S2B, Table S3**; (*13*)), while nicotinamide levels were anticorrelated with *NNMT* (**Table S2, Table S3**). We conclude that 1MNA levels reflect the activity of the metabolic pathway that converts nicotinamide to 1MNA, catalysed by the enzyme NNMT. Moreover, this pathway is not tightly coupled to the activity of other pathways of core metabolism. We therefore decided to investigate possible explanations for relationship between HMT levels and 1MNA pathway activity.

### HMT and NNMT act as alternative sink pathways to dispose of methyl groups

The reaction catalysed by NNMT uses SAM as a cofactor, transferring a methyl group from SAM to nicotinamide to form 1MNA. NNMT has been proposed function as a “sink” for methyl groups, allowing excess SAM to be converted to SAH. High NNMT activity can act as a methyl sink by reducing the SAM:SAH ratio (*11, 14–16*).

To further investigate the relationship between the 1MNA synthesis pathway and HMT activity we investigated the relationship between *NNMT* expression and total HMT expression. NNMT protein levels in two cancer cell line panels, the CCLE and NCI60 (*17*), correlated strongly to *NNM*T expression (**Fig S2C-D** respectively). *NNMT* expression is therefore a reliable indicator of NNMT protein levels and catalytic activity. *NNMT* expression and total HMT expression in the CCLE were negatively correlated (**Fig 1D**). HMT protein levels were negatively correlated with NNMT protein levels both for individual HMTs (for 20 HMTs detected in >90% of samples, mean Pearson’s *r* to NNMT protein levels =-0.108, range-0.306 – 0.199, FDR < 0.05 negative correlation for 12/20 and positive correlation for 1/20) and collectively (**Fig S2E**; mean Pearson’s *r* with sample mean HMT protein Z-score =-0.244, p-value = 2.62 × 10^-5^). HMT protein levels are also negatively correlated with 1MNA levels (**Table S3**; mean Pearson’s *r* with sample mean HMT protein Z-score =-0.102, p-value = 7.94 × 10^-2^). Altogether, this suggested that elevated 1MNA synthesis is associated with reduced HMT activity.

One possible explanation for the negative association between HMT levels and 1MNA synthesis is that 1MNA directly represses HMT transcription. If so, this would predict that the relationship between HMT and 1MNA should persist even if the correlation of *NNMT* expression on HMT expression was controlled. However, partial correlation analysis indicated that the correlation between HMTs and 1MNA was greatly weakened from-0.280 to-0.073 after taking into account *NNMT* expression (**Fig S2F**). We conclude that 1MNA itself is unlikely to regulate HMT expression. Instead, HMT levels are primarily associated with *NNMT* expression, and thus with the rate of 1MNA synthesis, rather than 1MNA levels themselves. These findings are consistent with the proposal that HMT activity and NNMT activity are parallel pathways capable of acting as methyl sinks (*9*). This implies that the total activity of these two pathways is balanced, which would result in a reciprocal relationship between the activity of HMTs and NNMT. Histone methylation may be used as an alternative methyl sink when NNMT activity is low and vice versa (**Fig 1E**)

### HMT and NNMT expression are tightly coupled in primary tumours

To test if the relationship we uncovered in cancer cell lines was also seen in tumours we used RNA-seq data from primary tumours from 33 distinct cancer types found in the Cancer Genome Atlas (TCGA) database to interrogate the correlation between HMTs and *NNMT* expression. The HMT-*NNMT* expression relationship was much stronger than that observed in the CCLE cell lines, both in a pan-cancer analysis (**Fig 1E-F**) and within individual cancer types (**Supp. Files 1**-**2**). The relationship was strengthened by pooling the expression of HMTs together (**Fig 1F, Supp. Files 1-2**). As an example, among the 177 pancreatic adenocarcinoma (PAAD) primary tumour samples, the relationship of *NNMT* expression with expression of individual HMTs was for the most part not exceptionally strong. However, when their expression was pooled, HMT expression anticorrelated with *NNMT* better than almost any single gene (rank 3, top 0.00496%; **Fig 1G**). Conversely, *NNMT* was one of the most negatively correlated genes to pooled HMT expression (rank 9, top 0.0149% of genes overall).

Raw correlation statistics (such as *ρ*) do not necessarily provide a reliable comparator of the strength of the relationship, as the distribution of gene correlations can differ between cancer or tissue types (*18*). To overcome this, we computed a relative reciprocal relationship score for HMTs and *NNMT* (see Methods). The relative reciprocal relationship scores were much stronger in the TCGA primary tumours than in the CCLE cell lines (**Fig 1I, Fig S2G**). The HMT-*NNMT* relationship was found across most cancer types and was particularly strong in liver, pancreatic and bladder cancers. The only cancer in which this relationship was not evident at all was acute myeloid leukaemia (LAML). This may be a result of the low expression of *NNMT* in this cancer type (**Fig S2H**). Indeed, across all cancers there was a significant correlation between the strength of the relationship between *NNMT* and HMT levels and the expression of *NNMT* (**Fig S2I**). Importantly, the HMT-*NNMT* relationship remained robust when controlling for immune cell infiltration, as estimated by two distinct gene expression deconvolution tools (**Fig S3**).

To visualise how the correlation strengthened as HMT expression was combined, we performed a pan-cancer analysis in which we added successively more HMTs and calculated the correlation between these HMTs and *NNMT*. The relationship between HMTs and *NNMT* became stronger as more HMTs were added, regardless of the order with which the HMTs were combined (**Fig 1H**). This relationship indicates that the anticorrelation between HMTs and *NNMT* is distributed across HMTs and not due to one or two HMTs with unusually strong anticorrelations. Using a simple stochastic modelling approach, we determined that the reinforcement of this relationship when more HMTs are added is consistent with co-regulation of the HMTs at the transcriptional level (**Fig S2J**).

### Among cellular methyltransferases, HMTs have the strongest relationship with NNMT

To test whether the strong anticorrelation to *NNMT* is specific to histone lysine methyltransferases, we calculated the relative reciprocal relationship scores for protein arginine methyltransferases (PRMTs), DNA methyltransferases and groups of RNA methyltransferases (**Fig S4**; gene sets in **Table S1**). The relationship of *NNMT* to histone lysine methyltransferases was the strongest and most widespread. There was no significant relationship between *NNMT* and protein arginine methyltransferases or tRNA/rRNA methyltransferases. In breast cancer (BRCA) and 16 other cancer types *NNMT* had a strong negative relationship to mRNA methyltransferases. In lung squamous cell carcinoma (LUSC) and 4 other cancer types there was a strong negative relationship between *NNMT* and small RNA methyltransferases. More notably, 9 out of 33 cancer types displayed a strong relationship between *NNMT* and methyltransferases with unknown substrates, with very strong relationships evident in LUSC and stomach and colon adenocarcinomas (STAD & COAD). We conclude that other cellular methyltransferases act in parallel to lysine histone methyltransferases and *NNMT* in disposing of excess methyl groups. However, this function is most evident for histone lysine methyltransferases across cancer.

### In healthy tissues, HMTs correlate with PEMT, an alternative methyl sink

We wondered whether the relationship between HMTs and *NNMT* was specific to cancer. We therefore investigated the HMT-*NNMT* relationship in RNA-seq data from healthy tissue samples available through the Genotype-Tissue Expression (GTEx) project. A strong relationship between *NNMT* expression and HMT expression was observed in only one individual tissue type (muscle) (**Fig 2A-B, Supp. File 3**). Moreover, in 10 out of the 12 TCGA cancer types for which matched tumour and normal tissue samples from at least 30 patients were available, the negative relationship of *NNMT* and HMT was stronger in the cancer samples relative to the matched normal samples (paired Wilcoxon test on reciprocal scores, p-value = 1.47 × 10^-3^; **Fig S1K**).

**Fig 2.**
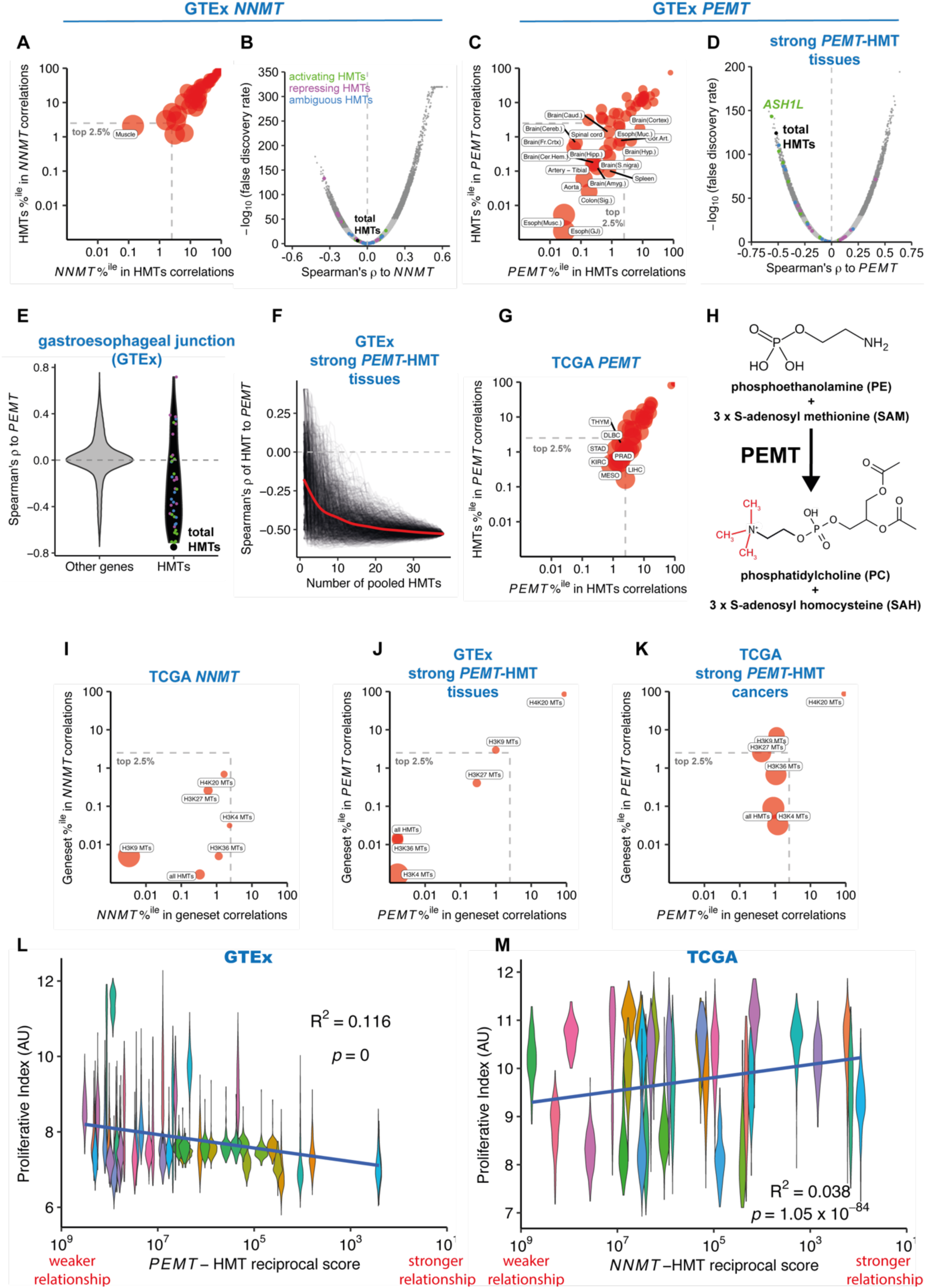
Total histone methyltransferase expression is strongly anticorrelated with the expression of *PEMT* in healthy tissues. A) Analysis showing rank percentile position of total HMTs among correlations of *NNMT* expression to 56200 genes and vice versa in 48 distinct healthy tissue types from the GTEx project. Bubble size is inversely proportional to the log of the ‘relative reciprocal score’, the sum of squares of the ranks of total HMTs/*NNMT* in the reciprocal distribution (see methods). The dashed grey box indicates correlations in the strongest 2.5% of anticorrelated genes, with tissues labelled. B) Volcano plot showing Spearman’s correlation and FDR for expression of *NNMT* vs 56200 genes in a pan-cancer analysis of GTEx primary tumours across 48 tissue types. HMT-encoding genes are shown as points coloured according to association with transcriptional regulation; correlation for total HMT expression is shown as a black point. C) Analysis showing rank percentile position of total HMTs among correlations of *PEMT* expression and vice versa in healthy tissue types from the GTEx project. Bubble size and dashed grey box as in panel **2A**. D) Volcano plot showing Spearman’s correlation and FDR for expression of *PEMT* vs 56200 genes in a cross-tissue analysis of 18 tissue types tissues with a strong HMT-*PEMT* relationship (within the grey box in panel **2C**). HMT-encoding genes are shown as points as in panel **2B**. E) Violin plot showing Spearman’s correlation to *PEMT* of HMTs (black, right) or other genes (left, grey) in 375 patient samples from the gastroesophageal junction. HMT-encoding genes are shown as points as in panel **2B**. F) Spearman’s correlation vs *PEMT* expression of total expression of pooled HMTs added to the pool in a random order in a cross-tissue analysis of tissues with a strong HMT-*PEMT* relationship (within the grey box in panel **2C**). 1000 individual iterations are shown as black lines, with Loess fit trendline in red. G) Analysis showing rank percentile position of total HMTs among correlations of *PEMT* expression to 60489 genes and vice versa in 33 cancer types from the TCGA. Bubble size and dashed grey box as in panel **2A**. H) PEMT sequentially methylates phosphoethanolamine to produce phosphatidylcholine, converting 3 molecules of SAM to SAH. I) Analysis showing rank percentile position of HMTs classified by their substrate histone lysine residues among correlations of *NNMT* expression and vice versa in cancer types from the TCGA. Bubble size and dashed grey box as in panel **2A**. J) Analysis showing rank percentile position of HMT sets methylating distinct histone lysine residues among correlations of *PEMT* expression to 56200 genes and vice versa in a pan-tissue analysis of 18 tissue types from the GTEx with a strong HMT-*PEMT* relationship (within the grey box in panel **2C**). Bubble size and dashed grey box as in panel **2A**. K) Analysis showing rank percentile position of HMT sets methylating distinct histone lysine residues among correlations of *PEMT* expression and vice versa in a pan-cancer analysis of 7 cancer types from the TCGA with a strong HMT-*PEMT* relationship (within the grey box in panel **2G**). Bubble size and dashed grey box as in panel **2J**. L) Violin plot showing healthy tissue sample proliferative index (PI), a measure of proliferation inferred from sample RNA-seq gene expression data, for 48 tissue types of the GTEx arranged by the strength of the anticorrelating relationship between *PEMT* and total HMTs. Note the x axis is inverted as a lower relative reciprocal score indicates a stronger relationship. M) Violin plot showing tumour PI for 31 cancer types of the TCGA arranged by the strength of the anticorrelating relationship between *NNMT* and total HMTs.

*NNMT* expression is generally low in healthy tissues, with high expression only evident in the liver, whereas it is often overexpressed in cancers (*19–26*). We tested whether total HMT expression might show an analogous relationship to another methyltransferase or class of methyltransferases operating within the cell. In 18/48 tissues total HMT expression anticorrelated strongly with *PEMT* (reciprocally in top 2.5% of genes; **Fig 2C, Supp. File 4**). Interestingly a particularly high proportion of brain tissues (10/13) showed a strong, significant relationship. Across the 18 tissues that showed strong associations between *PEMT* and HMT expression (**Fig 2D**), the relationship became stronger as more HMTs were pooled (**Fig 2E, F**). *PEMT* was strongly negatively correlated to HMTs in 7/33 cancers (**Fig 2G**).

PEMT is an enzyme that adds three methyl groups to the phospholipid phosphoethanolamine (PE) to produce phosphatidylcholine (PC) (**Fig 2H**). PC makes up around 40% of the lipid content of the plasma membrane in eukaryotic cells (*27*). PEMT activity contributes around 30% of cellular PC synthesis (*28*). The abundance of PC in the membrane suggested the possibility that PEMT could act as a sink for methyl groups, similar to NNMT; indeed, PEMT has been suggested to be the primary consumer of SAM in mammals (*29*).

We investigated whether other groups of methyltransferases also anticorrelated to *PEMT*. We found a significant negative relationship of *PEMT* to mRNA methyltransferases in 14 tissue types, including 9/13 brain tissues (**Fig S5**). Likewise, 7 tissues (5 from the brain) displayed an analogous relationship with DNA methyltransferases (**Fig S5**). Histone lysine methyltransferases had the strongest and most consistent anticorrelation to *PEMT*.

### Differential contributions of methylated residues to methyl sink activity

We investigated whether methyltransferases specific to particular lysine residues on histone tails contribute more strongly to the relationship with cellular methylation sinks. In cancer the HMTs that methylate the H3K9 residue showed a stronger relationship to *NNMT*, while the relationship to methyltransferases targeting other residues was weaker, albeit still highly significant (**Fig 2I**). In healthy tissues *PEMT* showed a strong reciprocal relationship to H3K4 and H3K36 methyltransferases, whereas the relationship with H3K9 and H3K27 methyltransferases, associated with transcriptional repression, was weaker and no strong relationship existed for H4K20 methyltransferases (**Fig 2J**). The difference between PEMT and NNMT was specific to the enzymes themselves, because cancers with a significant HMT-*PEMT* relationship showed similar residue specificity as healthy tissues (**Fig 2K**).

One factor that might be related to the specific HMTs that correlate with *PEMT* or *NNMT* could be the activity of the different pathways across the cell cycle. Transcription-associated methylation on H3K4me3 and H3K36me3 is enriched in quiescent cells relative to methylation of H3K9me3, which largely occurs in late S and G2 phase to restore H3K9 methylation to newly synthesised histones (*30*). *PEMT* expression is also reported to vary strongly across the cell cycle, peaking in G1 phase and declining in S phase (*31, 32*). We tested the correlation between the strength of HMT-*PEMT* relationship across the GTEx healthy tissues to the tissue sample’s Proliferative Index, a measure of proliferation inferred from RNA-seq data (*33*). The HMT-*PEMT* relationship tended to be stronger in tissues with a lower proliferative index (**Fig 2L**; R^2^ = 0.116, p-value ∼ 0). This may reflect the inability of proliferating cells to use PEMT as a methyl sink throughout the cell cycle. The HMT-*NNMT* relationship in cancers exhibited the opposite trend, albeit weakly (**Fig 2M**; R^2^ = 0.0384, p-value = 1.05 × 10^-84^). The aggressive proliferation of cancer cells and the lack of *NNMT* expression in most healthy tissues may explain why total HMT expression correlates to different methyl sinks in cancers and healthy tissues.

### Histone methylation in chromatin responds to methyl sink activity without affecting transcription

We investigated whether the relationship between HMT activity and *PEMT* or *NNMT* had consequences for histone methylation levels in chromatin genome-wide. We used ChIP-seq data from healthy tissues and cancer cell lines to assess histone methylation levels. In healthy tissues, we found that there was a global negative relationship of PEMT expression with H3K4me3, H3K9me3 and H3K27me3 at all classes of genomic regions examined. For instance, 99 % of 12355 gene promoters marked by H3K4me3 peaks showed an anticorrelated relationship between *PEMT* expression and total H3K4me3 signal or H3K4me3 peak width, an orthogonal measure of histone methylation levels (**Fig 3A-B, Fig S6A-B**). Similarly, 99 % of 2870 repetitive regions modelled showed an anticorrelated relationship for H3K9me3 signal and *PEMT* expression (**Fig 3A**). However, this was not the case for H3K36me3, which had a moderately positive relationship with *PEMT* expression at promoters and gene bodies. No negative relationships with *NNMT* expression were observed in healthy tissues (**Fig S6C**).

**Fig 3.**
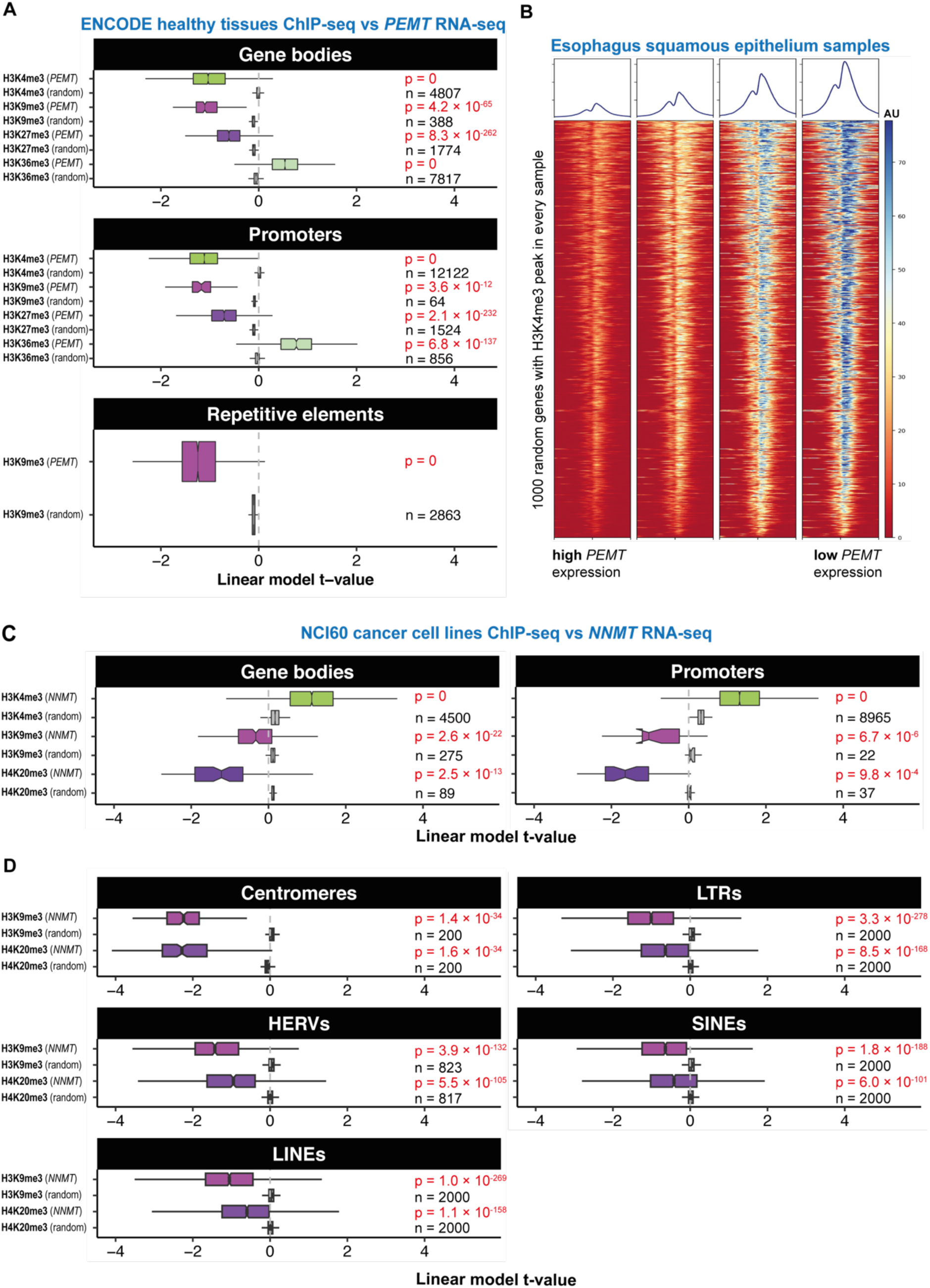
*PEMT* and *NNMT* expression anticorrelate globally with levels of specific histone marks genome-wide in healthy tissues and cancers respectively. A) Boxplot shows t-values from linear mixed effects model for sample *PEMT* expression predicting ChIP-seq signal for various histone marks (label left) on gene bodies, promoters or repetitive elements (sub-panel headers) in patient tissue samples collected as part of the ENCODE project. The number of individual sites is noted on the plot for each boxplot. p-values derive from paired Wilcoxon tests against a null distribution calculated by the mean t-value at each locus for 1000 random expressed genes. B) Heatmap showing H3K4me3 ChIP-seq signal (log_2_ fold change over input) over 1000 random genes for 4 samples from the squamous epithelium of the esophagus arranged in order of *PEMT* expression. C) Boxplot shows t-values from generalised linear models for *NNMT* expression (RNA-seq) predicting ChIP-seq signal for various histone marks (label left) on gene bodies and promoters in cell lines of the NCI60 cancer cell line panel. The number of individual sites is noted on the plot for each boxplot. p-values derive from paired Wilcoxon tests against a null distribution calculated by the mean t-value at each locus for 1000 random expressed genes. D) Boxplot shows t-values from generalised linear models for *NNMT* expression (RNA-seq) predicting ChIP-seq signal for various histone marks (label left) on different classes of repetitive elements in cell lines of the NCI60 cancer cell line panel. The number of individual sites is noted on the plot for each boxplot. p-values derive from paired Wilcoxon tests against a null distribution calculated by the mean t-value at each locus for 1000 random expressed genes. Sites shown are from bin with highest ChIP signal (cf **Fig S7D**). Abbreviations: LTRs, long terminal repeats; HERVs, human endogenous retroviruses; SINEs, short interspersed nuclear elements; LINEs, long interspersed nuclear elements.

Altered histone modification levels are often associated with changes in transcription of the genes at the corresponding loci. However, we found that despite low signal of H3K4me3 (associated with transcription) and H3K9me3 (associated with repression) in high-PEMT samples, expression from marked genes was not affected in either case (**Fig S6D**). Similarly, we found that variation across samples in total H3K4me3 signal (**Fig S6E**) or width (**Fig S6F**) at marked promoters, and H3K9me3 signal on H3K9me3-marked gene bodies, does not correlate with the expression of the corresponding genes. Thus in healthy tissues, PEMT varied with histone methylation levels independent of effects on transcription.

In cancer, H3K9me3 and H4K20me3 at both gene bodies and promoters were anticorrelated to cell line *NNMT* expression as measured by RNA-seq (**Fig 3C**), microarrays (**Fig S7A**) and proteomics (**Fig S7B**); however, H3K4me3 showed a positive relationship. No negative relationships were observed for *PEMT* expression (**Fig S7C**). Similarly, we found that both H3K9me3 and H4K20me3 levels at multiple classes of repetitive elements were negatively correlated with *NNMT* expression (**Fig 3D**). For all classes, the negative correlation was stronger at genomic sites with a higher average ChIP-seq signal across samples (**Fig S7D**). The anticorrelation with *NNMT* expression was particularly strong at centromeric satellites (**Fig 3D**), independent of their tendency to display higher signal of heterochromatic marks than other classes of repetitive element (**Fig S7E**).

While variation across samples in H4K20me3 signal at H4K20me3-marked gene bodies was negatively correlated with *NNMT* levels, expression from those genes displayed little relationship with *NNMT* (**Fig S7F**). We also estimated locus-specific expression of transposable elements, specifically human endogenous retroviruses (HERVs). Similarly, we found that despite reduced signal of this canonically repressive histone mark at HERVs in samples with high *NNMT* expression, HERV expression was not increased (**Fig S7F**). Indeed, variation in total signal of H3K9me3/H4K20me3 at marked sites was not associated with the level of transcription from either gene bodies or HERVs (**Fig S7G**).

### Histone methyltransferase genes are co-expressed

We wished to understand the structure and origin of the variation we observe in total HMT expression. We found that the expression of HMT genes was significantly more positively correlated to each other than to random genes (**Fig 4A-B**). The strongest co-expression was evident among the most highly expressed HMTs, which had a range of target lysine residues and canonical associations with transcription (see **Fig 4A** sidebars). This core of 14-16 highly expressed and highly correlated HMT genes was largely stable between healthy tissues and cancer (**Fig 4A**) and across individual tissue or cancer types (**Supp. Files 5-6**). We performed gene co-expression network analysis, showing a similar network architecture with a small number of modularity classes in both cancer and healthy tissue. The strength of network edges was highly concordant between pan-cancer and pan-tissue correlation analyses (**Fig 4C, Fig S8A**; Pearson’s correlation = 0.834; Jaccard Index on network with |edge strength| > 0.2 = 0.569). Whilst distinct modularity clusters showed some similarity to annotated associations with transcription (**Fig 4C-D, Fig S8A-B**), the overlap was not strong. Together, this suggested the possibility that HMTs might be co-ordinately regulated independent of their transcriptional functions.

**Fig 4.**
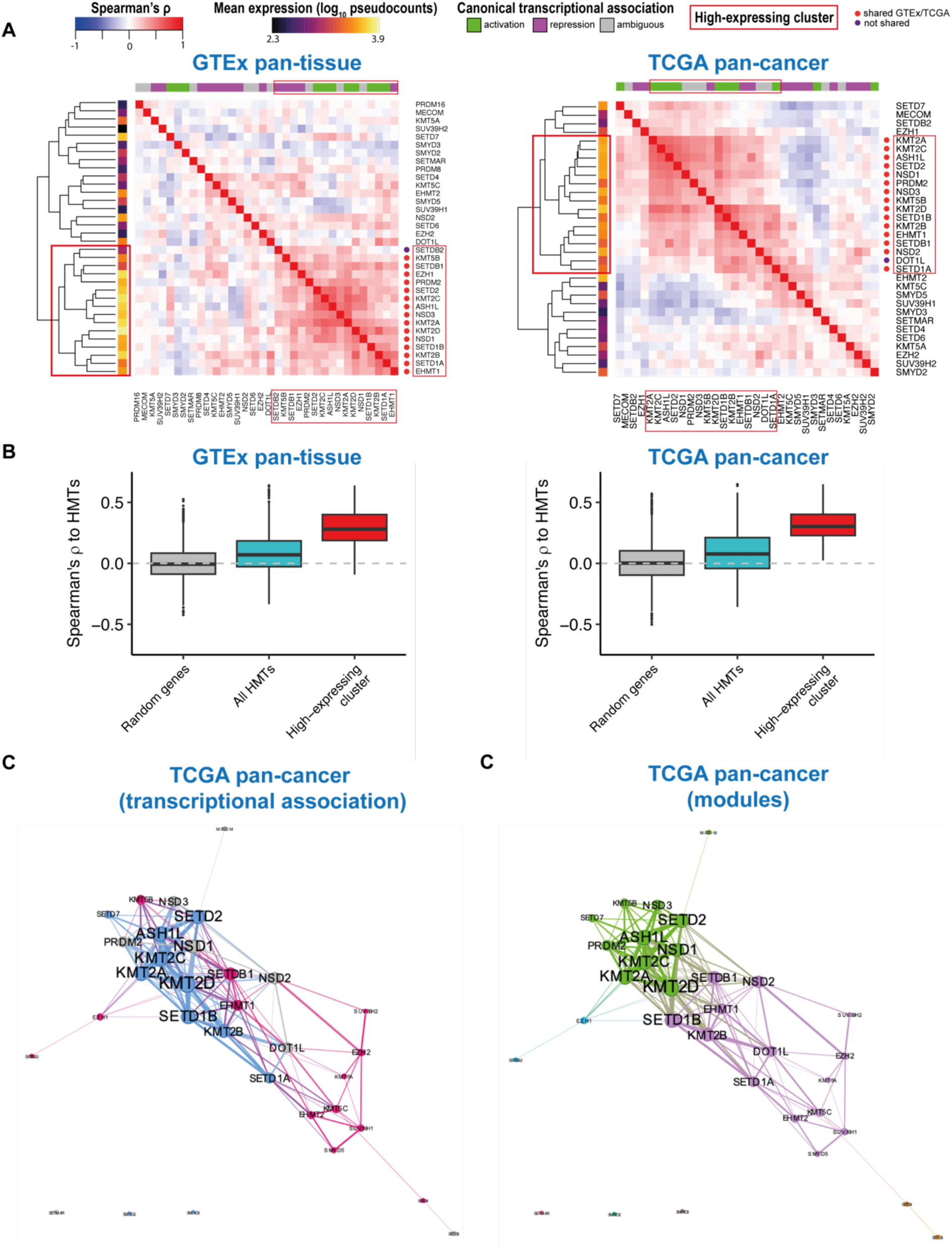
Highly-expressed histone methyltransferase genes are co-expressed. A) Heatmaps showing correlation of histone methyltransferase gene expression levels corrected for known confounders (GTEx left, TCGA right). Sidebars display expression level (left) and canonical transcriptional association (top). Dendrograms show hierarchical clustering of correlations. Red boxes indicate highly-expressed and correlated HMTs. Orange and purple dots indicate whether membership of the high-expressing cluster is shared across GTEx and TCGA analyses for each gene. B) Boxplots indicate distributions of Spearman’s correlations to HMT genes for random genes, within all HMTs and within the HE cluster. C) Network plot shows all correlations between HMTs with magnitude > 0.2 in a TCGA pan-cancer analysis. Edge weight is proportional to the correlation strength. Node size is proportional to the weighted degree. A ForceAtlas2 layout is displayed. Blue nodes indicate HMTs associated with transcriptional activation, magenta with transcriptional repression and grey with ambiguous association. Edge colours reflect the colours of the connected nodes. D) Network plot for TCGA pan-cancer analysis coloured to show distinct modules discovered using the Louvain method.

### Histone methyltransferase genes are regulated by E2F and Rb

Seeking to understand the possible basis for co-regulation of HMTs, we turned to *Caenorhabditis elegans*, a model organism with a simpler genetic regulatory architecture (*34*). We observed a positive correlation among HMTs across 206 diverse natural genetic backgrounds in *C. elegans*, with the strongest correlation in a core of 10 genes largely consisting of the most highly expressed HMTs (**Fig S9A**). Genes in this cluster were more likely to have orthologues in the human highly-expressed cluster (**Fig S9A**; odds ratio = 11.07, Fisher’s exact test p = 0.0108). Additionally, we observed very strong negative correlations between total HMT expression and expression of the *NNMT* orthologues *anmt-1/3* and the *PEMT* analogue, *pmt-1* (**Fig S9B**). This relationship was partially due to varying levels of *anmt-1/3* and *pmt-1* across development (**Fig S9C**) and was also evident when controlling for developmental age (**Fig S9D-E**).

We performed a *de novo* motif enrichment search on the upstream regions of coregulated *C. elegans* HMT genes. The most strongly enriched motif, present in 9/10 genes, resembled the binding motif of the E2F orthologue EFL-2 (**Fig 5A**). E2F transcription factors can be bound by the Retinoblastoma (Rb) protein, which represses transcription of E2F targets (*35*). Using previously published genome-wide ChIP-seq data (*36*) we observed an enrichment for binding of LIN-35, the *C. elegans* Rb orthologue, close to the transcription start site of HMT genes in the highly-expressed cluster relative to other HMTs or to random genes (**Fig 5B**). Indeed, in RNA-seq data from *lin-35* mutants (*37, 38*), we saw total HMT expression increased by 12-15% (1-way ANOVA, p = 0.053).

**Fig 5.**
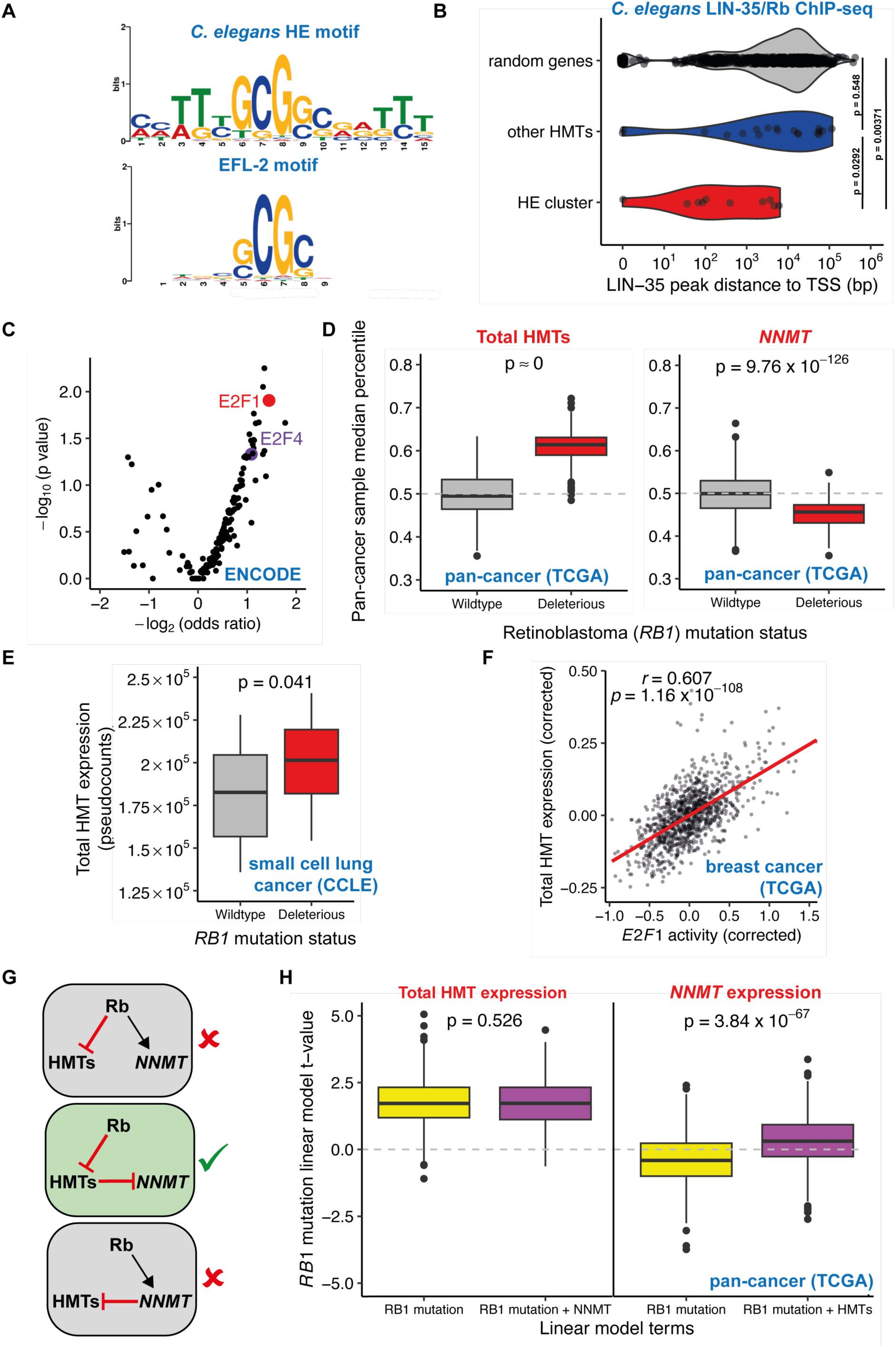
Histone methyltransferases are regulated by E2F and Retinoblastoma, with *NNMT* expression reduced downstream of HMTs in Rb-mutant cancers. A) Above: Sequence motif enriched in *C. elegans* highly-expressed cluster HMT promoters, relative to other HMTs promoters. Below: previously reported EFL-2 binding motif. B) Binding of the *C. elegans* Retinoblastoma orthologue LIN-35 upstream of the transcription start site (TSS) of the HE cluster, other HMT genes, and random genes. p-values from Wilcoxon test. C) Enrichment for transcription factor binding, from ENCODE ChIP-seq experiments, upstream of human HMT genes. Odds ratios and p-value derived from Fisher’s exact test. D) Boxplots show median total HMT or *NNMT* expression percentile drawn from 1000 iterations of pan-cancer sampling of tumours with wildtype *RB1* or potentially deleterious *RB1* mutations. p-value derived from t test. E) Total HMT expression in small cell lung cancer cell lines from the CCLE with wildtype *RB1* or deleterious *RB1* mutations. p-value derived from t test. F) Estimated E2F1 activity vs total HMT expression (both corrected for confounders) in breast cancer primary tumours from the TCGA. G) Potential architectures of the gene regulatory network linking *RB1*, *NNMT* and HMTs. H) Linear model t-values explaining total HMT and *NNMT* expression for *RB1* mutation status as the sole explanatory variable or jointly considered with *NNMT*/HMT expression respectively. p-value derived from t test.

To test whether E2F transcription factors also regulated histone methyltransferase expression in humans we performed an enrichment analysis for transcription factor binding sites upstream of human histone methyltransferase genes determined from ENCODE project ChIP-seq experiments for 181 distinct TFs. We observed that E2F1 was the second most significantly enriched TF (**Fig 5C**).

We identified 256 primary tumours from 10 cancer types in the TCGA that had potentially deleterious mutations in the Retinoblastoma-encoding *RB1* gene in at least 10 samples per cancer type. In all cancer types *RB1*-mutant tumours had higher mean total HMT expression; in a pan-cancer analysis we observed that this difference was highly significant (**Fig 5D**). Among 31 single HMTs with expression in all samples, 19 showed a marked upregulation in *RB1-*mutant tumours, with particularly notable upregulation of *EZH2*, *DOT1L* and *NSD2*, while only 5 displayed a clear downregulation (**Supp. File 7**).

We also identified cancer cell lines from the CCLE with deleterious Rb mutations. Around half of Rb mutations in CCLE cell lines were found in lung cancer cell lines, particularly small cell lung cancer lines. Total HMT expression was significantly upregulated in *RB1*-mutant lung cancer cell lines relative to wildtype-*RB1* cell lines (**Fig 5E**; 2-way ANOVA controlling for lung cancer subtype, p = 0.034).

To test whether variability in HMT expression across tumours was due to variable Rb/E2F activity, even if Rb was not mutated, we inferred the activity of 354 transcriptional regulators from expression of target genes in RNA-seq data from thousands of samples across the TCGA and GTEx (*39*). In the TCGA, inferred E2F1 activity was significantly (FDR < 0.1) and positively correlated with sample HMT expression in 30/33 cancer types, for example breast cancer (**Fig 5F**). E2F1 was the 10^th^ transcriptional regulator whose activity most strongly correlated with total HMT expression in a pan-cancer analysis (**Table S6**; Spearman’s rho = 0.457, 2.82 %^ile^). Interestingly, this relationship was much more notable in cancer than healthy tissue (**Table S7**). Altogether these results suggested that Rb activity represses HMT transcription and that this activity is conserved in *C. elegans* and humans.

### In Rb-mutant cancers NNMT is downregulated to compensate for increased HMT expression

The preceding results suggested that variation in HMT levels across cancers is associated to variability in Rb and E2F activity and their effects on transcription. *NNMT* expression was significantly reduced in *RB1*-mutant tumours (**Fig 5D**), consistent with their anticorrelation to HMTs. A key question is how HMT activity is co-ordinated with *NNMT*. One possible scenario is that HMTs and *NNMT* might be co-ordinated by a variable regulator (e.g. Rb) which has opposite effects on the transcription of HMTs and *NNMT*. Alternatively, *NNMT* expression might be regulated to compensate for existing variability in HMT levels, or vice versa (**Fig 5G**).

We tested this by modelling HMT expression and *NNMT* expression in response either to *RB1* mutation status alone, or in response to the combined effects of *RB1* and either HMTs or *NNMT*. Inclusion of *NNMT* in the model had no impact on the statistical relationship of HMT expression and *RB1* mutation status (**Fig 5H**). This is consistent with a direct transcriptional regulation of HMTs by Rb/E2F and demonstrates that expression of HMTs in these tumours is independent of *NNMT* expression. Conversely, including HMT expression when predicting *NNMT* expression abrogates the negative relationship between *NNMT* expression and *RB1* mutation status in tumours (**Fig 5H**). We conclude that in *RB1*-mutant cancers, *NNMT* is not directly downregulated. Instead, HMT expression is upregulated and *NNMT* expression is downregulated downstream of HMTs to compensate for their increased activity.

Altogether these analyses support the hypothesis that variability in E2F pathway activity drives variation in HMT expression in cancer and that this in turn affects the expression and activity of the *NNMT* methyl sink due to the important function that HMTs play in regulating SAM levels.

## DISCUSSION

Using data from tens of thousands of human samples, here we demonstrated that histone methyltransferase expression was strongly anticorrelated to the activity of two pathways known to consume excess methyl groups (known as methyl sinks): synthesis of 1-methylnicotinamide by the enzyme NNMT in cancers and production of phosphatidylcholine by PEMT in healthy tissues. This relationship implied that HMTs also acted as a methyl sink. Variation in HMT activity thus correlates to the extent to which alternative methyl sink pathways operate: high HMT activity is associated with low NNMT/PEMT activity and vice versa. However, we found no evidence that this variation in HMT activity had an effect on transcription. Below we discuss the implications of these results for understanding the roles of histone post-translational modifications.

We have shown a strong anticorrelation between HMT levels and *NNMT* activity in cancer. These results fit with earlier findings that changes in *NNMT* expression could modulate histone methylation (*11, 14–16*). This was previously argued to be due to a passive effect of NNMT activity on cellular methylation potential via the SAM/SAH ratio. However, we showed that this relationship corresponds to differences in expression of HMTs and *NNMT*, which is better explained by the hypothesis that histone methylation is playing a direct metabolic role in the cell. In this model, changes in HMT activity would lead to changes in NNMT activity because both pathways have an important role in maintaining SAM/SAH levels. Our discovery that HMT genes are co-expressed and co-regulated despite divergent associations to transcriptional regulation supports this hypothesis.

The histone code hypothesis proposed that specific histone modifications have direct and instructive effects on transcription (*2*). This would imply that changes in the levels of individual HMTs function to regulate transcriptional outputs. In contrast, we showed that much observed variation in histone methylation is associated with metabolic enzymes with functions far removed from gene regulation. Importantly, this variation does not have any detectable impact on transcription, even as a by-product. These results do not necessarily contradict a role for HMTs in instructing transcriptional regulation. For example, it may be that the variation we observe occurs within a range that does not affect transcription. Alternatively, the changes that occur in histone methylation levels at particular genes may require other changes, such as combinations of histone marks or specific transcription factors, in order to bring about transcriptional responses.

Our results demonstrate that co-ordinated histone methyltransferase expression is controlled transcriptionally by the activity of the Rb/E2F pathway, such that E2F1 activates multiple HMTs. What might the function of this regulation be? It is possible that HMTs are co-regulated in response to cellular metabolic requirements for methyl sinks. However, our results place *NNMT* expression downstream of HMT expression, which suggests instead that co-ordinated changes in HMTs require compensation through changes in methyl sink activity in order to maintain SAM levels. One possible function of coordinated change in HMT activity might be a requirement for multiple histone methylation marks to be reintroduced following DNA replication (*40*). It would be interesting to test whether E2F activity is required for correct maintenance of histone modification landscapes through cell division.

The relationship that we have discovered between total HMT activity and the activity of cellular methyl group sinks indicates that maintaining a consistent activity of methyl sink pathways is vital for cellular homeostasis. The importance of this activity might be in buffering cellular methylation potential by consuming SAM to maintain the SAM/SAH ratio. Additionally, SAH is required to support the transsulphuration pathway, which is the cell’s only pathway to *de novo* synthesise cysteine and downstream metabolites (e.g. glutathione). In primary tumours access to cysteine is limited and cells may be forced to rely on transsulphuration (*41, 42*). However, cultured cells enjoy abundant cysteine supplied in frequently-replenished culture medium. In support of this notion, we observed that the HMT/*NNMT* relationship is much stronger in primary tumours than cultured cells. Even so, the relationship is still evident in cancer cell lines and we note that in the CCLE metabolomics data the metabolite whose levels most strongly correlate positively to total HMT expression is cystathionine (**Fig 1A**), a characteristic marker of transsulphuration.

Taken together, our results indicate histone methylation participates directly in core, conserved pathways of cellular metabolism. This role is independent of the role of histone methylation marks in regulating transcription. Histone proteins evolved in archaea, where they have a limited role in transcriptional regulation and there is little evidence of post-translational modifications such as methylation (*43*). It is interesting to speculate whether the metabolic roles of histone post-translational modifications may predate their more familiar role in transcription.

## FIGURES AND TABLES

**Table S1** Methyltransferase gene sets used in this study

**Table S2** CCLE metabolite-RNA-seq correlations for HMTs and NNMT

**Table S3** CCLE metabolite-proteomics correlations for HMTs and NNMT

**Table S4** ENCODE ChIP-seq and RNA-seq files

**Table S5** NCI60 ChIP-seq and RNA-seq files

**Table S6** HMT correlations to transcription factor activity in TCGA

**Table S7** HMT correlations to transcription factor activity in GTEx

**Supp File 1** CCLE NNMT-HMT correlation all cancers full labelled plots

**Supp File 2** TCGA NNMT-HMT correlation all cancers full labelled plots

**Supp File 3** GTEx NNMT-HMT correlation all tissues full labelled plots

**Supp File 4** GTEx PEMT-HMT correlation all tissues full labelled plots

**Supp File 5** GTEx individual tissue HMT correlation matrices

**Supp File 6** TCGA individual tissue HMT correlation matrices

**Supp File 7** Individual HMT levels in Rb-mutant tumours in TCGA

## METHODS

### RNA-seq data

RNA-Seq data were downloaded from the GTEx data portal for GTEx v8. Data were downloaded as raw counts. “Harmonised” (hg38) RNA-seq data were downloaded for TCGA projects using the ‘TCGAbiolinks’ package in ‘R’ as raw counts. CCLE RNA-seq read counts were downloaded from the DepMap download portal in August 2021 (version: DepMap Public 21Q3).

Raw counts were subjected to a median-ratio normalisation (MRN) prior to all analyses. The MRN was performed using the ‘DESeq2’ package in ‘R’ (*44*). Normalizations were applied both individually for each tissue or cancer type cohort and across all samples within each database. Normalized pseudocounts were obtained by converting raw counts data to a *DEseq2DataSet* object using the *DESeqDataSetFromMatrix()* function, applying the *estimateSizeFactors()* function to the resulting *dds* object, and then retrieving the normalized pseudocounts with the function *counts()* with *normalized = TRUE*. All correlations presented are based on these MRN-normalised pseudocounts.

For TCGA cancer type analyses, we only considered samples annotated as Primary Tumours, except where we explicitly note otherwise (e.g. adjacent normal tissue samples).

For the CCLE, we excluded samples from Primary Disease types with fewer than 20 cell lines.

We restricted our analyses to GTEx tissues with at least 100 samples or TCGA cancer types with at least 35 Primary Tumour samples.

### Metabolomics and proteomics data

CCLE metabolomics data file ‘CCLE_metabolomics_20190502.csv’ was downloaded from the DepMap download portal. Quantitative proteomics data derived from mass spectrometry for the CCLE was obtained from (*13*), Table S2.

Metabolites were manually annotated to KEGG pathways. As metabolites can often be attributed to the function of multiple pathways, we chose appropriate pathways for each metabolite in a heuristic manner aiming to cover a maximum number of metabolites with as few pathways as possible.

### Principal component analysis and hierarchical clustering

Hierarchical clustering of metabolite abundances was performed by running the *hclust()* function in ‘R’ on a distance matrix produced by the *dist()* function on the transposed matrix containing metabolomics data. Principal component analysis of metabolite abundances was performed by running the *prcomp()* function in ‘R’ on the transposed matrix containing metabolomics data.

### Correlating metabolites to gene expression

In order to account for biases in cell lines deriving from particular disease types, both metabolite abundances and gene expression were converted to Z-scores for each Primary Disease type prior to correlating metabolite levels to gene expression in the CCLE data. This was done by subtracting the disease type mean abundance from the sample abundance and dividing by the disease type standard deviation for abundance. For RNA-seq pseudocounts, the same approach was taken but using log_10_-transformed values. These Z-scores were then pooled to perform the correlation analysis. The same approach was taken for correlations of metabolites or gene expression with protein levels measured by proteomics.

For the volcano plot in **Fig 1B**, the genes correlated to metabolites were limited to a list of 10275 gold-standard genes which are universally expressed across samples (TPM > 5 across all samples in the GTEx data) before calculation of Z-scores. This was done to exclude genes likely to contain samples with 0 values, which would hamper the viable calculation of Z-scores.

Partial correlation analysis was performed on Z-scores as above, using the *pcor()* function in ‘R’.

### Correlation distributions

To ensure equal representation of each tissue or cancer type when combining types across a database, we randomly sampled 100 (GTEx), 36 (TCGA) or 20 (CCLE) from each tissue. Combining raw gene expression data for tissues or cancer types may introduce artifacts even when correcting for average tissue/cancer gene expression, as high-expressing tissues/cancers may still have greater variance in the absolute value of the residuals. To account for this, we ranked the sample gene expression pseudocounts for each gene within the sample chosen for each tissue or cancer type. We then combined the ranks for the chosen samples across tissue types, using the ranks in place of the raw residuals; this gave us 4800 samples for GTEx, 1188 samples for TCGA or 460 samples for the CCLE. The Spearman’s correlation was then computed across these aggregated ranks. As the result varies slightly depending on the random sampling within each tissue, we repeated this process 100 times, and plotted the median correlation for each gene.

For analyses within a single tissue type, such a normalisation was not required and we simply correlated the uncorrected pseudocounts for our chosen gene against all others, using all samples available in the cohort.

For gene sets such as histone methyltransferases, we added together pseudocounts for each sample for all of the genes before conducting the analysis above.

When computing genome-wide correlations we did not correct for any potential confounders. Where explicitly noted that values were corrected for confounding variables in the text, we corrected for the following variables in GTEx: age, sex, speed of death, ischemic time and sequencing batch. For TCGA we corrected for age, race, sex, tumour stage and sequencing centre.

### Matched cancer and normal samples

Matched primary tumour and adjacent normal tissue samples were identified using TCGA metadata and barcodes. Tissues were identified with at least 30 normal tissue samples. Using the donor portion of the TCGA barcode, matching primary tumour samples were identified. If multiple primary tumour samples matched the adjacent normal tissue sample, one was retained at random and the remainder were discarded. Additionally, normal tissue samples without lacking identifiable primary tumour samples in the expression data were discarded, such that all normal tissue samples had one matching primary tumour sample and vice versa.

### Relative reciprocal relationship scores

To calculate relative reciprocal relationship scores in order to compare the strength of gene anticorrelation across different tissue/cancer types, we calculated the genome-wide correlation distribution for both of the interrogated gene (set) pair. We then extracted the genome-wide rank of each of the interrogated gene pair (i.e. a rank of 1 for the most anticorrelated gene), squared these ranks in order to penalise weak reciprocity (i.e. a rank of 1 and 200 in the respective distributions yields a weaker score than ranks of 10 and 10) and added them together to yield the relative reciprocal score.

### Proliferative Index

The Proliferative Index (PI) was calculated with the ‘ProliferativeIndex’ package in ‘R’ (*33*). Briefly, the entire dataset across all tissues or cancer types was normalized by MRN and variance stabilising transformation using the *varianceStabilizingTransformation()* function of ‘DESeq2’. Following the normalization, the PI was calculated by applying the *readDataForPI()* function with a randomly selected gene specified in the *modelIDs* argument, then running *calculatePI()* on the resulting object.

### Simulation of correlations among co-regulated genes

We constructed a toy model whereby a theoretical co-regulator positively regulates forty genes (A_1_, A_2_.A_n_; analogous to HMT genes) and negatively regulates another gene, B. We simulated different concentrations of the co-regulator in 500 different samples, with its influence on A_1,2.n_ and B subject to random noise. We then correlated the simulated concentrations of B to A genes as more A genes are pooled (analogous to our practice of pooling reads for HMT genes). We repeated this simulation 1000 times.

### Estimations of total immune fraction

The estimates for the immune cell infiltration of TCGA samples using both the TIMER and EPIC RNA-seq deconvolution algorithms were downloaded directly from http://timer.cistrome.org/.

### ENCODE ChIP-seq data processing

We identified publicly-available histone methylation ChIP-seq data from adult human patient samples from the ENCODE project for tissues with a strong HMT-*PEMT* correlation in the GTEx data (top 2.5% in both reciprocal correlation distributions) which had at least three samples with available RNA-seq data per histone mark. This gave us 19 to 23 samples (depending on the histone mark) from 5 tissues: the esophagus muscularis, the gastroesophageal sphincter, the esophagus squamous epithelium, the sigmoid colon and the spleen. Data for these samples were available for 4 distinct histone methylation marks: H3K4me3, H3K9me3, H3K27me3 and H3K36me3. For the corresponding ENCODE samples, the following processed data files were downloaded from the ENCODE data portal: ChIP signal fold change over control (as bigwig file) and pseudoreplicated peaks (in bed narrowPeak format). RNA-seq files were downloaded as raw counts for each sample. RNA-seq counts for all samples from all tissues were pooled and MRN-normalised as described above to yield pseudocounts. File names and experiment DOIs are listed in **Table S4**.

Fold change over control ChIP-seq signal files downloaded from ENCODE were converted to log_2_ fold change over control. This was done by using the *bigwigCompare* function of the command line package ‘deepTools’ (v3.5.0) (*45*) to compare the fold change file against an artificial bigWig file with a flat signal of 1 across all chromosomes, using the argument *--operation log2*.

To select genomic regions in which to model ChIP-seq signal by gene expression, we looked for regions marked by a peak in at least 75% of samples (for H3K9me3, H3K27me3, H3K36me3) or in 100% of samples (H3K4me3, due to greater reproducibility of peak overlaps across samples for this mark). We imported the pseudoreplicated peak files into ‘R’ and used the *countOverlaps()* function of the ‘GenomicRanges’ package against the co-ordinates of genomic regions of interest to determine the number of peaks which overlapped that region (e.g. a specific promoter) in each sample. We then excluded regions with 0 overlapping peaks in >25% of samples across tissues (or any samples for H3K4me3).

The co-ordinates of promoters and gene bodies were generated using the ‘TxDb.Hsapiens.UCSC.hg38.knownGene’ package in ‘R’ with the *genes()* and *promoter()* functions (we used the default settings by which the *promoter()* function returns windows from 2000bp upstream to 200bp downstream of each gene’s transcription start site). The co-ordinates of repeat regions were obtained using the hg38 ‘rmsk.txt’ file downloaded from http://hgdownload.cse.ucsc.edu/goldenpath/hg38/database/.

In order to quantify signal in our chosen genomic regions for each sample, first we exported the genomic co-ordinates of regions of interest from ‘R’ in bed file format. We then used the *computeMatrix* function of ‘deepTools’ with the sample log_2_ fold change bigwig file and the genomic regions bed file as inputs. We set the arguments *--binSize* and *--regionBodyLength* to be equal (usually at 100); this results in an output of a single number for average log_2_ fold change over control ChIP signal for each genomic region.

We went on to model this ChIP signal at each genomic range by expression of chosen gene. Prior to modelling we filtered out peaks with an average ChIP signal across samples that fell below the level of the input control. From any individual analysis we first excluded any tissue type that had fewer than 3 samples available for that particular combination of histone mark and tissue, as they could not be effectively modelled.

As multiple ENCODE samples from different tissues often came from the same individual (8 individual donors), we used linear mixed effects models using the ‘lmer’ package in ‘R’ to control for this lack of true sample independence as follows. We first corrected the sample chosen gene expression for tissue and donor of origin by fitting a linear mixed effects model explaining log_10_ RNA-seq pseudocounts with tissue as a fixed effect and donor as a random effect, extracting the gene expression residual from the model for each sample using the *residuals()* function. We then went on to fit a linear mixed effects model for each genomic region explaining the log_2_ fold change over control signal by corrected gene expression residual and tissue as fixed effects and donor as a random effect. From this model, we extracted the model t-value using the *summary()* function as a measure of the explanatory power of gene expression on ChIP signal at that locus and plotted boxplots of all of the t-values for each histone mark and type of genomic region examined.

To generate a null distribution to compare against, we repeated the same modelling procedure with expression of 1000 random genes from our gold standard set of ubiquitously-expressed genes (excluding histone methyltransferases and demethylases). For each genomic region we then took the mean t-value for the 1000 random genes as our null distribution that is plotted alongside that of the chosen gene on the boxplot. p-values were obtained from a two-tailed paired Wilcoxon test of the observed t-value for our chosen gene for each region against the mean t-value for that region across 1000 random genes.

To measure the ChIP-seq H3K4me3 peak width, we used the H3K4me3 pseudoreplicated peak narrowPeak bed files downloaded from ENCODE (**Table S4**). For each genomic region of interest we identified overlapping peaks using the *findOverlapPairs()* function of the ‘GenomicRanges’ package in ‘R’. We then calculated the width of these peaks from the start and end co-ordinates of peak calls from the bed file using the *width()* function. Note we calculated the entire width of the overlapping peak and not only the part which overlapped the region of interest. We then aggregated the peak widths for each genomic region in the case of multiple peaks, to provide a single figure for the sum total width of called peaks overlapping that region. We then went on to model peak width as described above for ChIP-seq signal.

When modelling or correlating gene expression from promoters, we restricted the analysis to genes with expression detected in every sample.

### NCI60 ChIP-seq data processing

To probe the relationship of histone methylation levels in chromatin to NNMT in cancer we used the NCI60 (*46*), a panel of 60 cancer cell lines with associated RNA-seq, microarray, proteomics and ChIP-seq data.

NCI60 ChIP-seq data were taken from (*46*). We downloaded raw sequencing reads from the NIH’s Sequence Read Archive, using the command line tool ‘SRAtoolkit’ (SRA identifier numbers found in **Table S5**). We then aligned the reads to the human genome (hg38) using the ‘bowtie2’ package (*47*). Each experiment was available as two replicates, in addition to an input control sample. Some sequencing replicates consisted of single-reads, while others were paired-end experiments. In order for all analyses to be comparable, we aligned only the forward reads from paired-end experiments. The output from ‘bowtie2’ was saved to a.sam file and converted to a.bam file using the ‘samtools’ package (version 1.16.1). bam files were then sorted by query name using the *sort* function from ‘samtools’.

We used the ‘Genrich’ package [https://github.com/jsh58/Genrich] to call ChIP-seq peaks on the sorted.bam files, as ‘Genrich’ can take two replicate experimental files, in addition to a control file, and perform an integrated peak call relative to the control. For all marks (H3K4me3, H3K9me3 and H4K20me3) we used the following settings, corresponding to ‘broad’ peaks:-q (max FDR) 0.1,-g (max distance between significant sites) 400. Additionally, for H3K4me3 we used the default settings, corresponding to “narrow” peaks. We then imported these peak call files into ‘R’ and used the *reduce()* function of the ‘GenomicRanges’ package to combine the peaks called under broad and narrow settings for H3K4me3.

For each experimental replicate, we used the bamCompare function of ‘deepTools’ with *--operation log2* to return a signal file in bigWig format for log_2_ fold change over control. We identified genomic regions of interest (promoters, gene bodies and repeats) as described above for ENCODE, limiting to genomic sites with a called peak in >2/3 of samples. We then ran the *computeMatrix* function of ‘deepTools’ with the sample log_2_ fold change bigwig file and the genomic regions.bed file as inputs, with equal bin size and body length as described above, to return a single figure for average log_2_ fold change for each region. At this point we took the average of the 2 replicates and carried that forward into our modelling approach.

We downloaded raw RNA sequencing reads using ‘SRAtoolkit’ (SRA identifier numbers in **Table S5**) and aligned them to the human genome (hg38) using ‘bowtie2’ using default settings. We calculated read counts per gene by counting reads overlapping exons using the *summarizeOverlaps* function of the ‘GenomicAlignments’ package in ‘R’. Read counts were then MRN-normalised before use in modelling. We downloaded NNMT / *NNMT* SWATH mass-spectrometry proteomics values and 5-microarray gene expression Z-scores for NCI60 cell lines from CellMinerCDB (https://discover.nci.nih.gov/rsconnect/cellminercdb/).

For each genomic region, we modelled the average log_2_ fold change over control signal using a generalised linear model with the *glm()* function of the ‘stats’ package in ‘R’. In the model we included the expression of our gene/protein of interest (sample RNA-seq pseudocounts, protein level or mRNA Z-score), in addition to the cell line tissue of origin and an interaction term between tissue and expression. We extracted and plotted the model t-values for expression as described above.

### Bayesian iterative reweighting analysis of multi-mapping ChIP-seq reads in NCI60

Both ChIP-seq signal and RNA-seq expression levels from repetitive elements are difficult to quantify accurately due to ambiguous multi-mapping of sequencing reads to highly similar genomic regions. These reads are typically discarded in data processing pipelines (as in the ENCODE pipeline). However, for the NCI60 we used a Bayesian iterative reweighting approach (‘SmartMap’) to apportion multi-mapping reads to individual genomic loci, providing more accurate estimates of ChIP-seq signal at repetitive elements (*48*). We performed this analysis for H3K9me3 and H4K20me3.

The two replicates in the NCI60 ChIP-seq data, as well as being paired and unpaired, have different read lengths. Additionally, the input controls are unpaired reads with shorter read lengths (150bp). In the case of multimapping reads, greater read length was likely to affect the likelihood of unique mapping and so affect the validity of comparisons to the input control. As such, we only made use of a single replicate for each histone mark and cell line, namely the unpaired replicate with shorter read lengths that matched the input control. We adapted the *SmartMapPrep* script from the ‘SmartMap’ package to process raw single-end reads, downloaded using ‘SRAtoolkit’ as described above, for treatment and input controls, before using the *SmartMap* function for a single reweighting iteration as default and as recommended by the authors. The output from ‘SmartMap’ is a bedgraph file. We converted the bedgraph files to bigwig files with the UCSC ‘bedGraphToBigWig’ utility. Log_2_ fold signal over input control was found using deepTools *bigwigCompare* with the default ‘log2’--operation choice.

For peak calling, we used ‘deepTools’ *bigwigCompare* with the *--operation subtract* setting to remove the signal from the appropriate input control track from each track. We then converted these bigWig files back to bedgraph files with the UCSC *bigWigToBedGraph* utility. We used MACS3 (v3.0.0) (*49*) for peak calling, as it can call peaks from a bedgraph file using the *bdgbroadcall* function.

As above we analysed individual repetitive elements that were marked by a peak in at least 40 of 60 cell lines. Repetitive elements were identified from the ‘rmsk.txt’ file as described above, with the exception of HERVs, which were taken from the annotation included in the Telescope package (see below). Custom bed files were created with the elements to be analysed and signal was quantified across the entire element using ‘deepTools’ *computeMatrix* as described above.

### Estimation of HERV expression in the NCI60

We also used a separate Bayesian reweighting approach (‘Telescope’) to estimate locus-specific expression estimates from a set of human endogenous retroviruses (HERVs) (*50*). We aligned raw RNA-sequencing reads (downloaded with ‘SRAtoolkit’ as above) with ‘bowtie2’ with options *--very-sensitive-local* and *-k 100* (allowing up to 100 alignments per read), as recommended by the ‘Telescope’ package authors. The resulting bam files were processed in ‘Telescope’ using the *telescope assign* function call. The HERVs annotation file ‘HERV_rmsk.hg38.v2.gtf’ was downloaded from the ‘telescope_annotation_db’ repository on GitHub (https://github.com/mlbendall/telescope_annotation_db). We analysed individual HERVs marked by ChIP peaks (identified using ‘SmartMap’) in 40 of 60 cell lines and which had expression detected in at least 30 of 60 cell lines. When using HERV expression as a response variable in a linear model, we used negative binomial generalised linear model (with the *glm.nb()* function)due to typical overdispersion of the data.

### Correlations within histone methyltransferase genes

To probe co-expression of histone methyltransferase genes, we first corrected the expression values for confounding variables as described above. Additionally for the GTEx pan-tissue analysis we corrected for donor ID as a random variable within a linear mixed-effects model, to account for the fact that when comparing across tissues multiple samples can originate from the same individual donor. Corrected residuals were rank-percentile transformed within each tissue or cancer type, before 100 (GTEx) or 36 (TCGA) samples were chosen from each and combined before Spearman’s correlations among the rank-percentile transformed values were computed across the grouped samples. The process of sampling was repeated 100 times and the median Spearman correlation from the 100 iterations was taken for plotting. We excluded very lowly expressed histone methyltransferase genes from plots by filtering according to a geometric mean expression across all samples of at least 100 pseudocounts; this accounts for slightly different numbers of samples in the GTEx/TCGA plots shown in **Fig 4**.

Correlations of histone methyltransferase genes to random genes were computed as above with 100 random genes from our gold standard set of ubiquitously expressed genes for each histone methyltransferase. Values computed were then pooled.

Network plots were prepared from pan-cancer or pan-tissue correlation matrices by first filtering out edges with correlations of magnitude less than 0.2. Node size was based on its degree and edges were weighted by the square of the magnitude of the correlation. Network analysis was performed in ‘Gephi’ (version 0.10.1), with the following visualisation properties: ForceAtlas2 layout, edge weight range 0.1-2.0 and attraction 30 for TCGA and 10 for GTEx.

### *de novo* transcription factor binding motif search

We used ‘MEME’ (version 5.5.1) in discriminative mode to find motif occurrences that were enriched in the *C. elegans* highly expressed cluster promoters relative to the remaining HMT promoters. We used 1000 bp upstream of the TSS as our promoter sequences. The previously published EFL-2 motif (*51*) was obtained from CisBP (*52*) version 1.02; motif identifier M0675_1.02).

### CeNDR *C. elegans* data and processing

We downloaded RNA-seq raw counts and TPM-normalised values for *C. elegans* strains from the *C. elegans* Natural Diversity Resource (CeNDR; (*53*)) from Gene expression Omnibus, accession number GSE186719. TPM values were used only for estimating sample ages using the ‘RAPToR’ tool (see below). We MRN-normalised the raw counts and used these counts for all other analyses. The raw counts were transcript-level counts; these were collapsed down to gene-level counts prior to all analyses.

We used the ‘RAPToR’ package in ‘R’ (*54*) to infer the age of the samples according to the author’s instructions. We used the *Cel_YA_2* reference series from the ‘wormRef’ package. To obtain age-corrected residuals for gene expression, we fitted a spline with 6 degrees of freedom using the *smooth.spline()* function of the ‘stats’ package in ‘R’ to predict log_10_ pseudocounts from inferred age, taking the residuals from the spline with the *residuals()* function.

*C. elegans* histone methyltransferase genes were selected according to their gene descriptions on WormBase (version WS287). The list of *C. elegans* HMTs can be found in **Table S1.** The vast majority of strains were represented by 3 independent RNA-seq samples. We took the mean of the age-corrected residuals for each strain to plot scatterplots and compute HMT correlations for the heatmap. *C. elegans* orthologues of human highly-expressed cluster genes were determined using OrthoList 2 (*55*); a gene was annotated as an orthologue if the orthology relationship was present in at least 3 of the 6 databases compiled in OrthoList 2.

### *C. elegans lin-35* mutants and ChIP-seq data

To identify RNA-seq datasets from *lin-35* mutants, we searched the Gene Expression Omnibus for *lin-35* and found two studies; (*37*) (GEO accession GSE62833) for *lin-35* mutant or wildtype L3 larvae and (*38*) (GEO accession GSE155190) for L1 larvae. We downloaded the raw data and aligned it to the *C. elegans* genome (version WS276) using ‘bowtie2’ with default settings, before obtaining gene level counts using *summarizeOverlaps()* from ‘GenomicRanges’ as described above for NCI60 and then MRN-normalising the resulting counts with ‘DESeq2’ as described above. In both studies total HMT counts were increased in *lin-35* mutants on average by 12-15%. To assess statistical significance we analysed the two studies together, performing a 2-way ANOVA for total HMT counts with genotype and developmental stage as explanatory variables.

LIN-35 ChIP-seq data was obtained from (*36*), Table S1. We used ‘GenomicRanges’ in ‘R’ to determine the distance from reported significant LIN-35 peaks to the TSS of HMT genes. For “other genes”, we excluded all genes which have 0 expression in any sample; we then randomly sampled 1000 of the remaining ∼11000 genes.

### Rb-mutant cancers in the TCGA and CCLE

Mutation calls for TCGA samples were downloaded as MAF files using the ‘TCGABiolinks’ package in ‘R’ (*56*). We identified samples with a reported mutation in *RB1* which was either a missense mutation, nonsense mutation, frame-shifting insertion, in-frame deletion or a frame-shift deletion. 10 cancer types had at least 10 *RB1-*mutant samples with available RNA-seq data; BLCA, BRCA, CESC, COAD, HNSC, LIHC, LUSC, LUAD, SARC and UCEC. We assumed samples had wildtype *RB1* if they had mutations called in other genes in the MAF files but none called in *RB1*.

We rank-percentile transformed all samples in these cancer types (*RB1* mutant and wildtype together) and then sampled 10 *RB1*-mutant and 10 wildtype cancers from each cancer type, taking the median rank-percentile of each random sample. We repeated this sampling process 1000 times and plotted the medians from these samples for each group, comparing the medians with a t test.

To model HMT or *NNMT* expression by *RB1* mutation status and/or counterpart expression, we rank-percentile transformed HMT and *NNMT* expression by cancer type after correction for confounding variables. We then performed pan-cancer sampling of *RB1*-mutant or wildtype cancers as above, combining all samples and fitting a linear model with HMT/*NNMT* expression as response variable and either *RB1*-mutation status alone as an explanatory variable or together with counterpart expression. We then extracted the t-values from the various linear models for the statistical association of *RB1*-mutation to either HMT or NNMT expression.

For the CCLE, we downloaded the mutation calls using the *depmap_mutationCalls()* function of the ‘depmap’ package in ‘R’. We then filtered *RB1* mutations by whether they were called as deleterious or not. We found 90/1236 cell lines had deleterious *RB1* mutations, of which 43 were annotated as lung cancer cell lines, 32 specifically small cell lung cancer. As small cell lung cancers had a higher expression of HMTs than other lung cancer subtypes, we performed a 2-way ANOVA with genotype and lung cancer subtype as explanatory variables.

### Transcriptional regulator activity estimation

To estimate transcriptional regulator (TR) activity in GTEx and TCGA samples, we used the ‘decoupleR’ package in ‘R’ (*39*) to infer TR activity from RNA-seq samples. ‘decoupleR’ requires a gene regulatory network (GRN) to use as a basis for TR activity inference; we used the ‘dorothea’ package in ‘R’ (*57*) previously developed by the same authors. The ‘dorothea’ package includes two different human GRNs; one general and one for cancers. We used the general GRN for estimating GTEx sample TR activity and the cancer GRN for TCGA sample TR activity. In order to increase our confidence in the estimates, we excluded the lowest confidence TR-target interactions that were the result only of *in silico* predictions. We then excluded any TRs that had fewer than 10 target genes remaining by which to infer their activity. This left us with 354 transcriptional regulators. Before running ‘decoupleR’, we also weighted targets by the confidence of the interaction, converting the confidence reported by ‘dorothea’ (letters A-D after low-confidence interactions had been eliminated) into an integer value (1–4) and using its inverse as an interaction weight.

### ENCODE transcription factor binding enrichment

We downloaded the ENCODE transcription factor binding site profiles from the ‘Harmonizome’ database web portal (*58*). For the genes bound by each transcription factor, we performed a Fisher’s exact test for enrichment of HMT genes among the bound genes. Odds ratios and p values are extracted from the Fisher’s exact test; p values reported in the volcano plot are raw and uncorrected.

---

**Fig S1.**
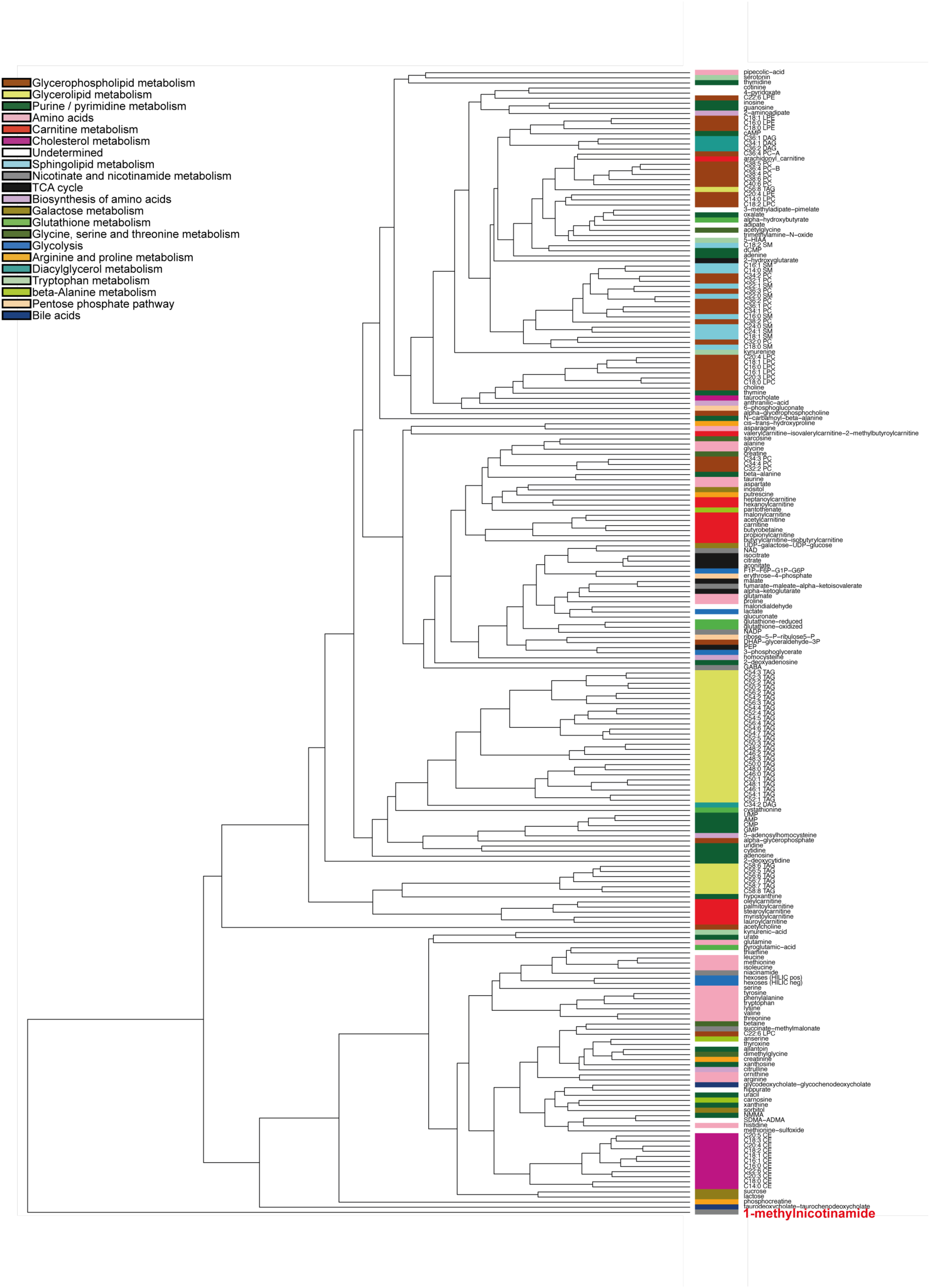
1-methylnicotinamide is an outlier when clustering metabolite levels across CCLE cell lines. Dendrogram showing unsupervised hierarchical clustering of 225 metabolites according to their levels in 927 CCLE cell lines. Sidebar is coloured according to representative metabolic pathways (see legend). 1-methylnicotinamide is classified as the outgroup and is highlighted in red.

**Fig S2.**
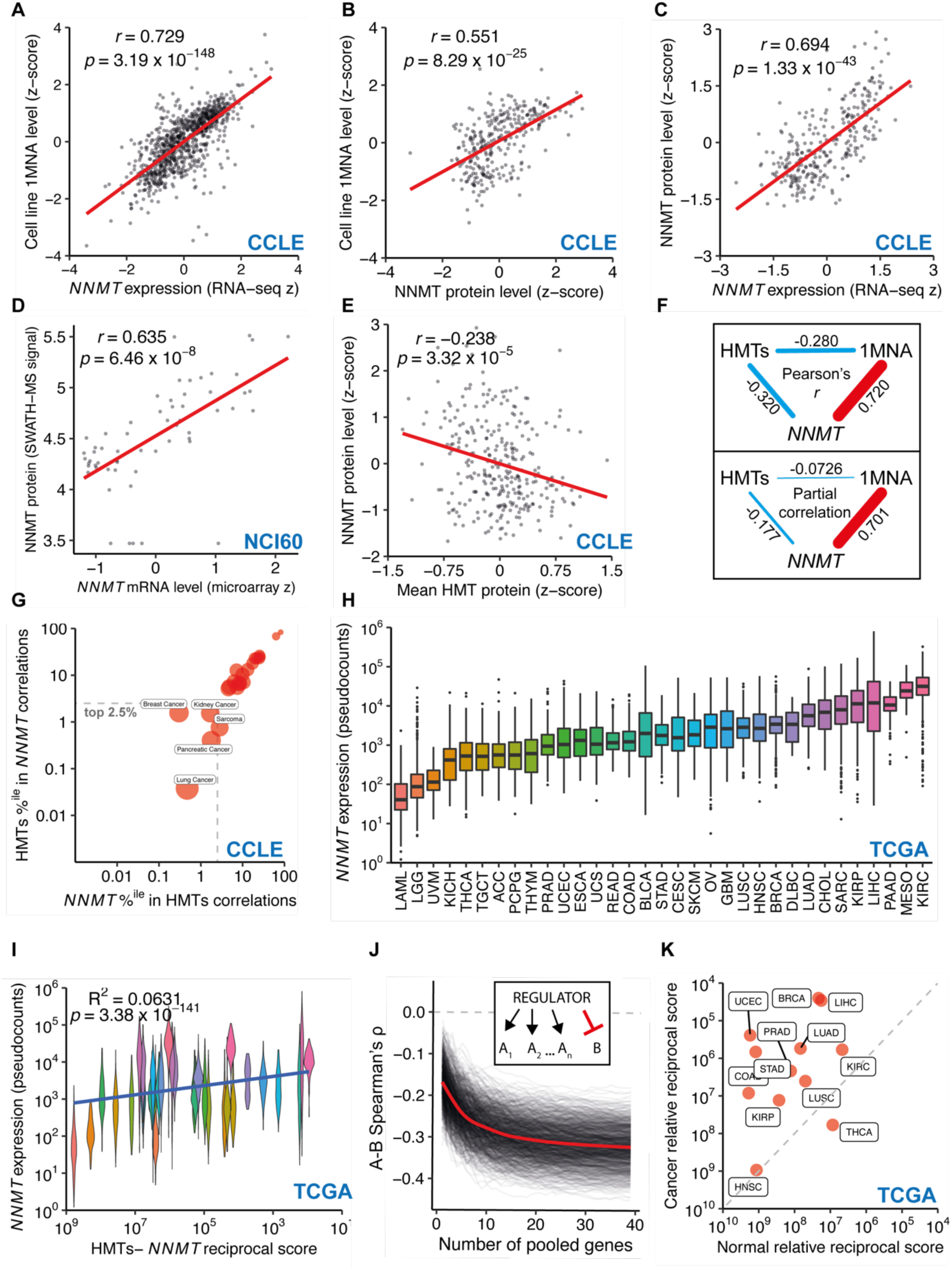
Total histone methyltransferase expression is strongly anticorrelated with the activity of NNMT in cancers (related to Fig 1) A) Scatterplot showing *NNMT* expression (RNA-seq) vs 1MNA levels (both normalised to cancer type z-scores) for 927 CCLE cell lines. B) Scatterplot showing NNMT protein levels (quantitative MS) vs 1MNA levels (both normalised to cancer type z-scores) for 298 CCLE cell lines. C) Scatterplot showing *NNMT* expression (RNA-seq) vs NNMT protein levels (both normalised to cancer type z-scores) for 927 CCLE cell lines. D) Scatterplot showing *NNMT* expression (average z-score across 5 distinct microarrays) vs NNMT protein levels (SWATH-MS signal) for 59 NCI60 cell lines. E) Scatterplot showing cell line mean cancer type-normalised z-score for protein levels of 20 reliably-detected HMTs vs NNMT protein levels (normalised to cancer type z-scores) for 298 CCLE cell lines. F) Pearsons’s correlations (blue) and partial corelations (orange) of *NNMT* expression, total HMT expression and 1MNA levels in 927 CCLE cell lines. G) CCLE pan-cancer analysis showing rank percentile position of total HMTs among correlations of *NNMT* expression to 52440 genes and vice versa in 23 distinct primary cancer types. Bubble size is inversely proportional to the log of the ‘relative reciprocal score’, the sum of squares of the ranks of total HMTs/NNMT in the reciprocal distribution. H) Boxplot showing *NNMT* expression levels among primary tumours of 33 distinct cancer types from the TCGA. I) Violin plot showing relationship of cancer type HMTs-NNMT relative reciprocal score to *NNMT* expression. Note the x-axis is inverted, because a low score indicates a strong relationship. J) Simulation of correlations between a pooled gene family (A_1,2…n_) and another gene (B) when they are conversely affected by a hypothetical transcriptional regulator. K) Scatterplot showing HMT-*NNMT* relative reciprocal relationship scores in primary tumour samples and matched normal samples. Note the axes are inverted as a lower score indicates a stronger relationship.

**Fig S3.**
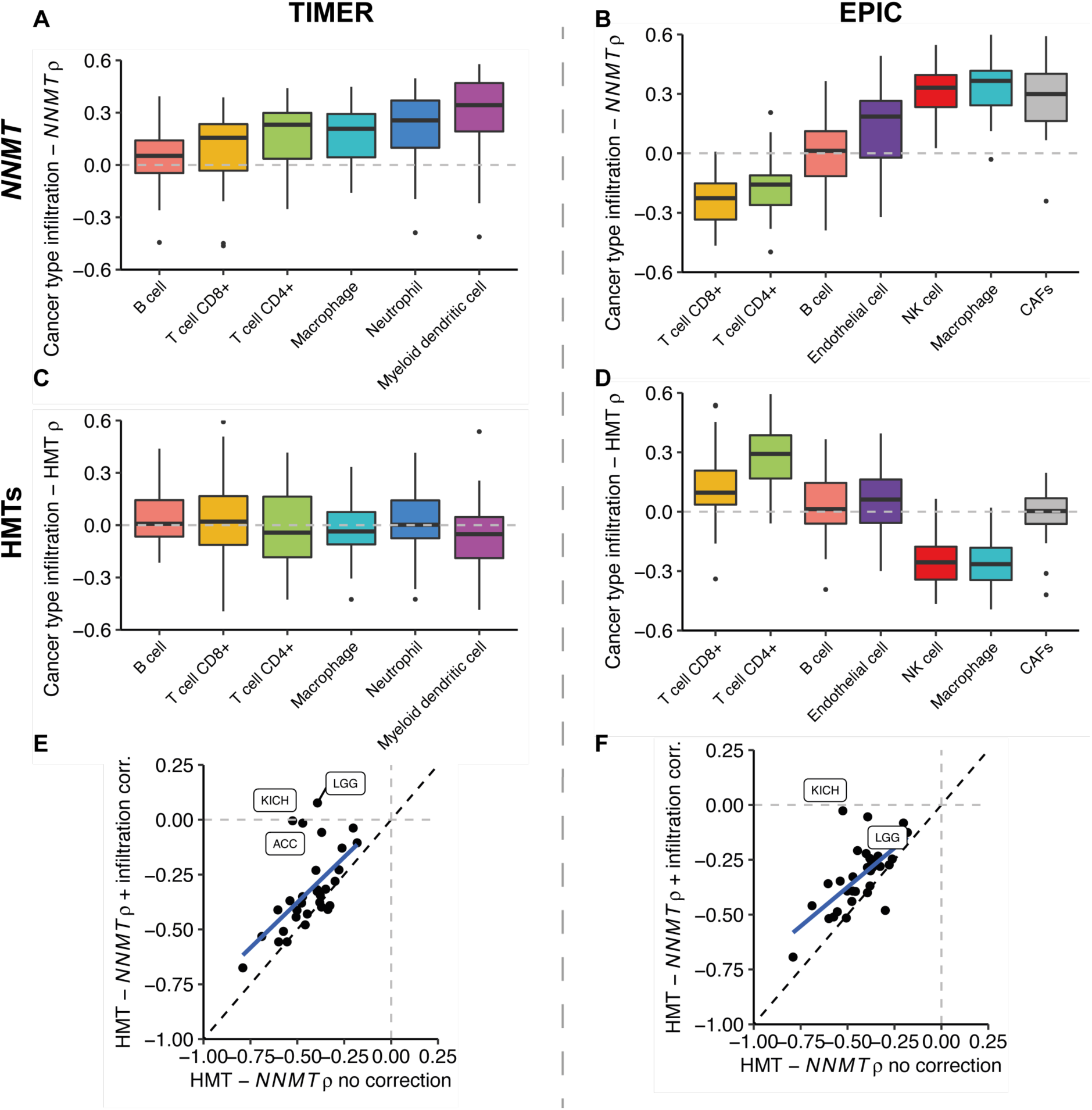
Infiltration of specific immune cell types into primary tumours correlates positively with *NNMT* expression but does not strongly confound the HMT-*NNMT* relationship. A) Boxplot showing the distribution of cancer-type correlations between *NNMT* expression and infiltration of specific immune cells as estimated by TIMER. B) Boxplot showing the distribution of cancer-type correlations between *NNMT* expression and infiltration of specific immune cells as estimated by EPIC. C) Boxplot showing the distribution of cancer-type correlations between total HMT expression and infiltration of specific immune cells as estimated by TIMER. D) Boxplot showing the distribution of cancer-type correlations between total HMT expression and infiltration of specific immune cells as estimated by EPIC. E) Scatterplot showing cancer-type correlations between total HMT expression and *NNMT* with or without correction for infiltration of specific immune cells as estimated by TIMER. Cancer types with a strong loss of HMT-*NNMT* relationship are labelled. F) Scatterplot showing cancer-type correlations between total HMT expression and *NNMT* with or without correction for infiltration of specific immune cells as estimated by EPIC. Cancer types with a strong loss of HMT-*NNMT* relationship are labelled.

**Fig S4.**
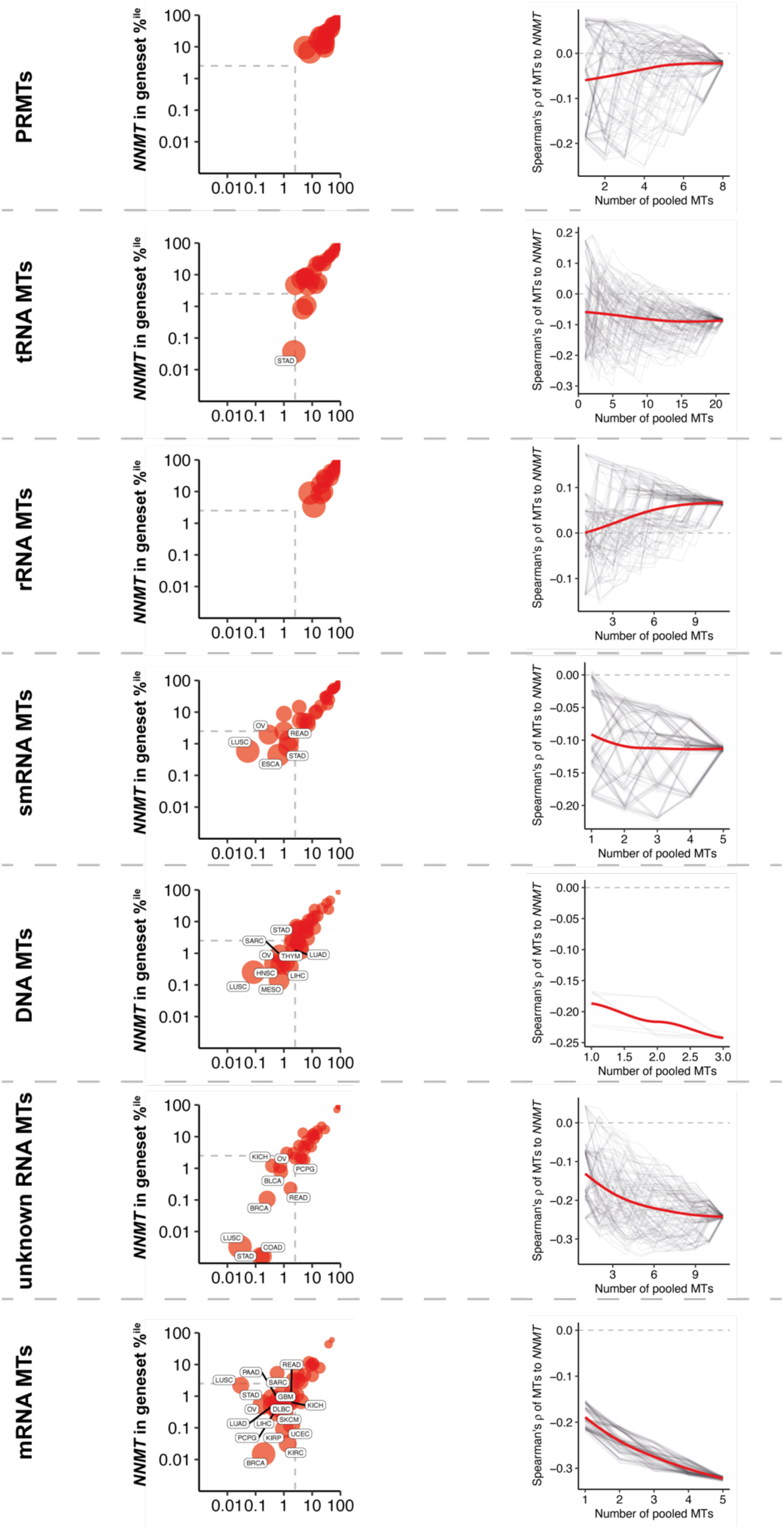
Relationship of other groups of methyltransferases to *NNMT* in cancers. For various functionally-related groups of methyltransferase genes (listed in Table S1) the left-hand plot shows the rank percentile position of total methyltransferase group expression among correlations of *NNMT* expression to 60489 genes and vice versa in 33 distinct cancer types. Bubble size is inversely proportional to the log of the ‘relative reciprocal score’, the sum of squares of the ranks of total MTs/*NNMT* in the reciprocal distribution. The dashed grey box indicates correlations in the strongest 2.5% of anticorrelated genes. Compare to **Fig 1I**. The right-hand plot shows the Spearman’s correlation vs *NNMT* of total expression of pooled HMTs added to the pool in a random order. 120 individual iterations, or the maximum number of permutations if below 120, are shown as black lines, with Loess trendline in red. Compare to **Fig 1H**. Abbreviations: PRMTs, protein arginine methyltransferases; tRNA MTs, transfer RNA methyltransferases; rRNA MTs, ribosomal RNA methyltransferases; smRNA, small RNA methyltransferases; unknown RNA MTs, unknown substrate RNA methyltransferases; mRNA MTs, messenger RNA methyltransferases.

**Fig S5.**
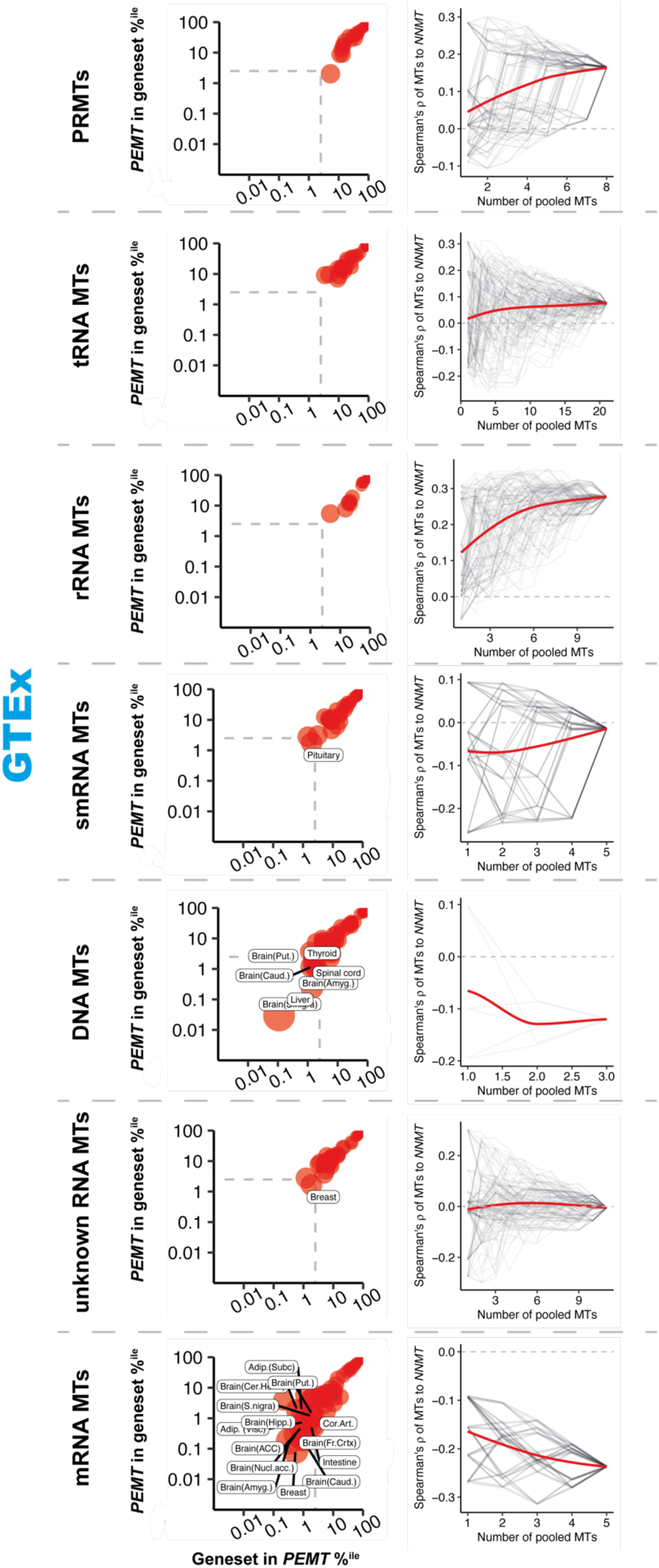
Relationship of other groups of methyltransferases to *PEMT* in healthy tissues. For various functionally-related groups of methyltransferase genes (listed in **Table S1**) the left-hand plot shows the rank percentile position of total methyltransferase group expression among correlations of *PEMT* expression to 56200 genes and vice versa in 48 distinct GTEx tissue types. Bubble size is inversely proportional to the log of the ‘relative reciprocal score’, the sum of squares of the ranks of total MTs/*PEMT* in the reciprocal distributions. The dashed grey box indicates correlations in the strongest 2.5% of anticorrelated genes. Compare to **Fig 2C**. The right-hand plot shows the Spearman’s correlation vs *PEMT* of total expression of pooled HMTs added to the pool in a random order. 120 individual iterations, or the maximum number of permutations if below 120, are shown as black lines, with Loess trendline in red. Compare to **Fig 2F**. Abbreviations: PRMTs, protein arginine methyltransferases; tRNA MTs, transfer RNA methyltransferases; rRNA MTs, ribosomal RNA methyltransferases; smRNA, small RNA methyltransferases; unknown RNA MTs, unknown substrate RNA methyltransferases; mRNA MTs, messenger RNA methyltransferases.

**Fig S6.**
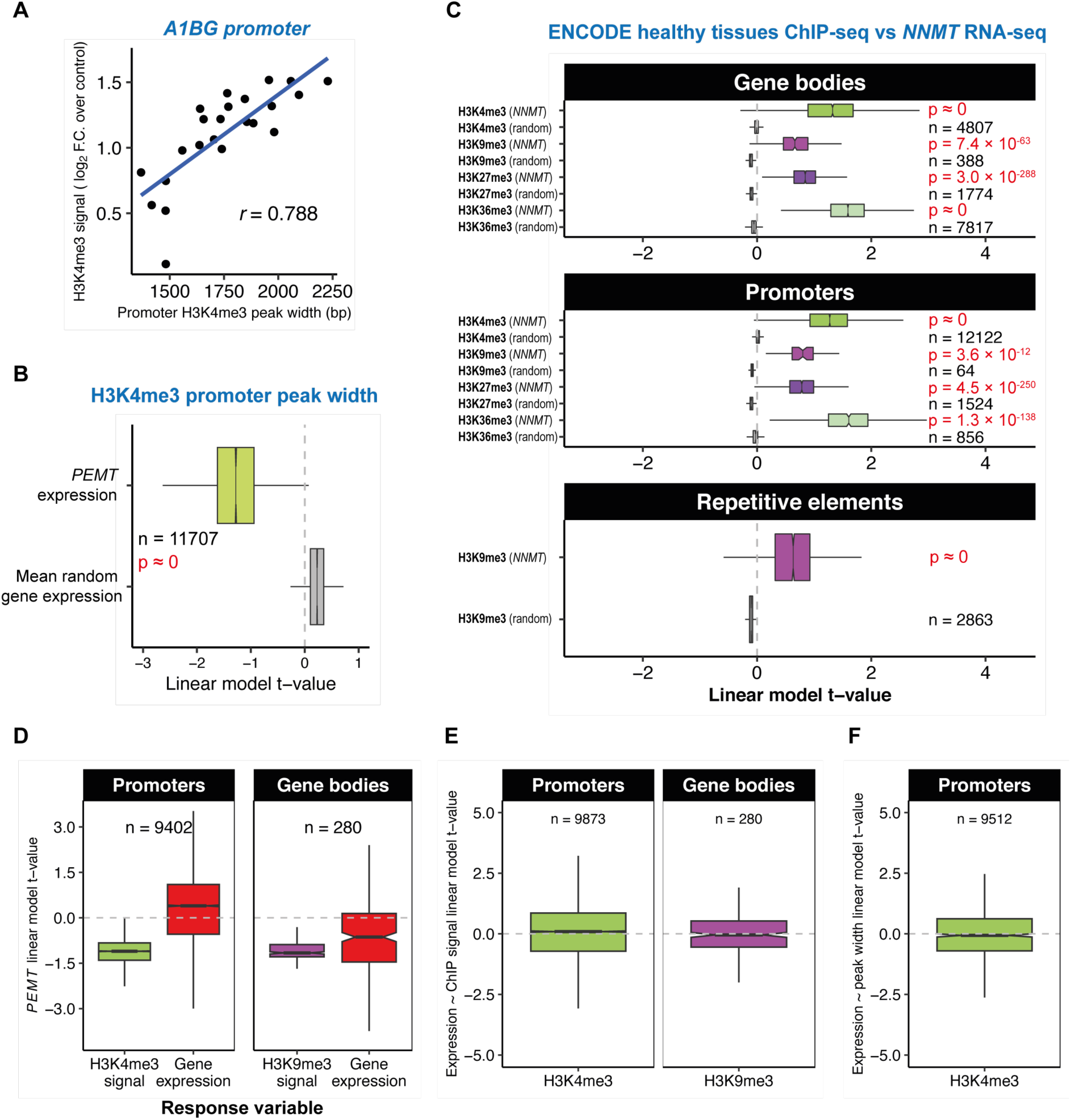
*PEMT* expression anticorrelate globally with levels of specific histone marks genome-wide in healthy tissues (related to Fig 3) A) Scatterplot showing correlation between H3K4me3 ChIP signal and H3K4me3 peak width at a representative promoter (*A1BG*). B) Boxplots showing linear model t-values for *PEMT* expression predicting H3K4me3 peak width at gene promoters and the mean t-values at promoters for 1000 random genes. C) Boxplot shows t-values from linear mixed effects model for sample *NNMT* expression predicting ChIP-seq signal for various histone marks (label left) on gene bodies, promoters or repetitive elements (sub-panel headers) in patient tissue samples collected as part of the ENCODE project. The number of individual sites is noted on the plot for each boxplot. p-values derive from paired Wilcoxon tests against a null distribution calculated by the mean t-value for 1000 random expressed genes. D) Boxplot shows linear mixed effects model t values for sample *PEMT* expression predicting H3K4me3 or H3K9me3 signal at promoters or gene bodies respectively, and predicting expression of those marked genes. E) Linear mixed effects model t-values for ChIP-seq signal of H3K4me3 at promoters or H3K9me3 at gene bodies predicting expression from the marked genes. F) Linear mixed effects model t-values for H3K4me3 peak width at promoters predicting expression from marked genes.

**Fig S7.**
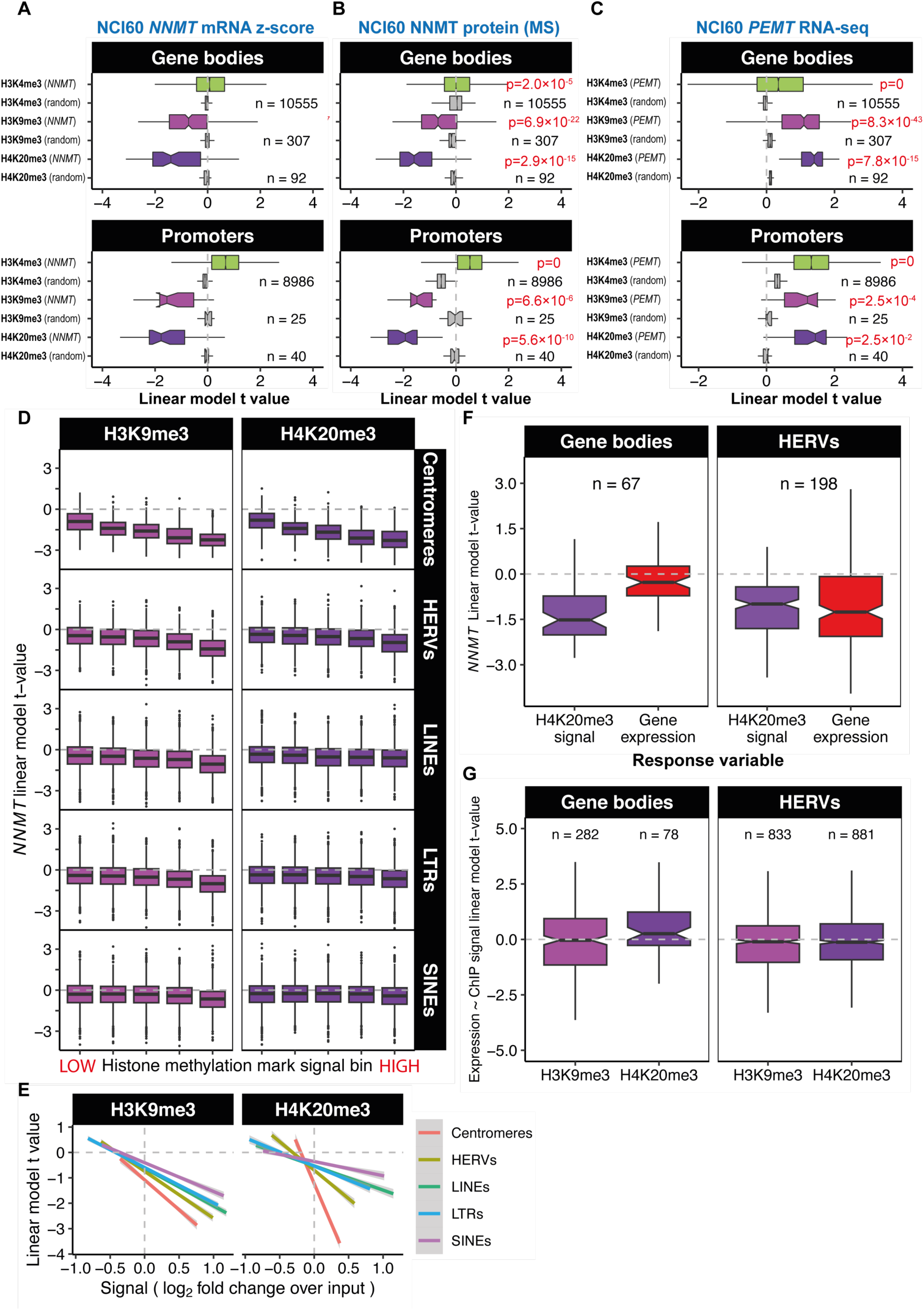
*NNMT* expression anticorrelate globally with levels of specific histone marks genome-wide in cancer cell lines (related to Fig 3) A) Boxplot shows t-values from generalised linear models for *NNMT* expression (Z-score pooled from expression profiling with 5 independent microarrays) predicting ChIP-seq signal for various histone marks on gene bodies or promoters in the NCI60 cancer cell line panel. B) Boxplot shows t-values from generalised linear models for NNMT protein levels (measured by SWATH mass spectrometry) predicting ChIP-seq signal for various histone marks on gene bodies, promoters or repetitive elements in the NCI60 cancer cell line panel. C) Boxplot shows t-values from generalised linear models for *PEMT* expression (RNA-seq) predicting ChIP-seq signal for various histone marks on gene bodies, promoters or repetitive elements in cell lines of the NCI60 cancer cell line panel. D) Boxplot shows linear model t values for *NNMT* predicting total ChIP-seq signal at repetitive elements, with loci binned by increasing average signal strength. E) Linear model t values for *NNMT* predicting total ChIP-seq signal at different classes of repetitive element by total average signal strength. F) Boxplot shows linear mixed effects model t values for sample *NNMT* expression predicting H4K20me3 at gene bodies and human endogenous retroviruses (HERVs), and predicting expression of those marked loci. G) Linear mixed effects model t-values for ChIP-seq signal of H3K9me3 and H4K20me3 at gene bodies and HERVs predicting expression from the marked loci.

**Fig S8.**
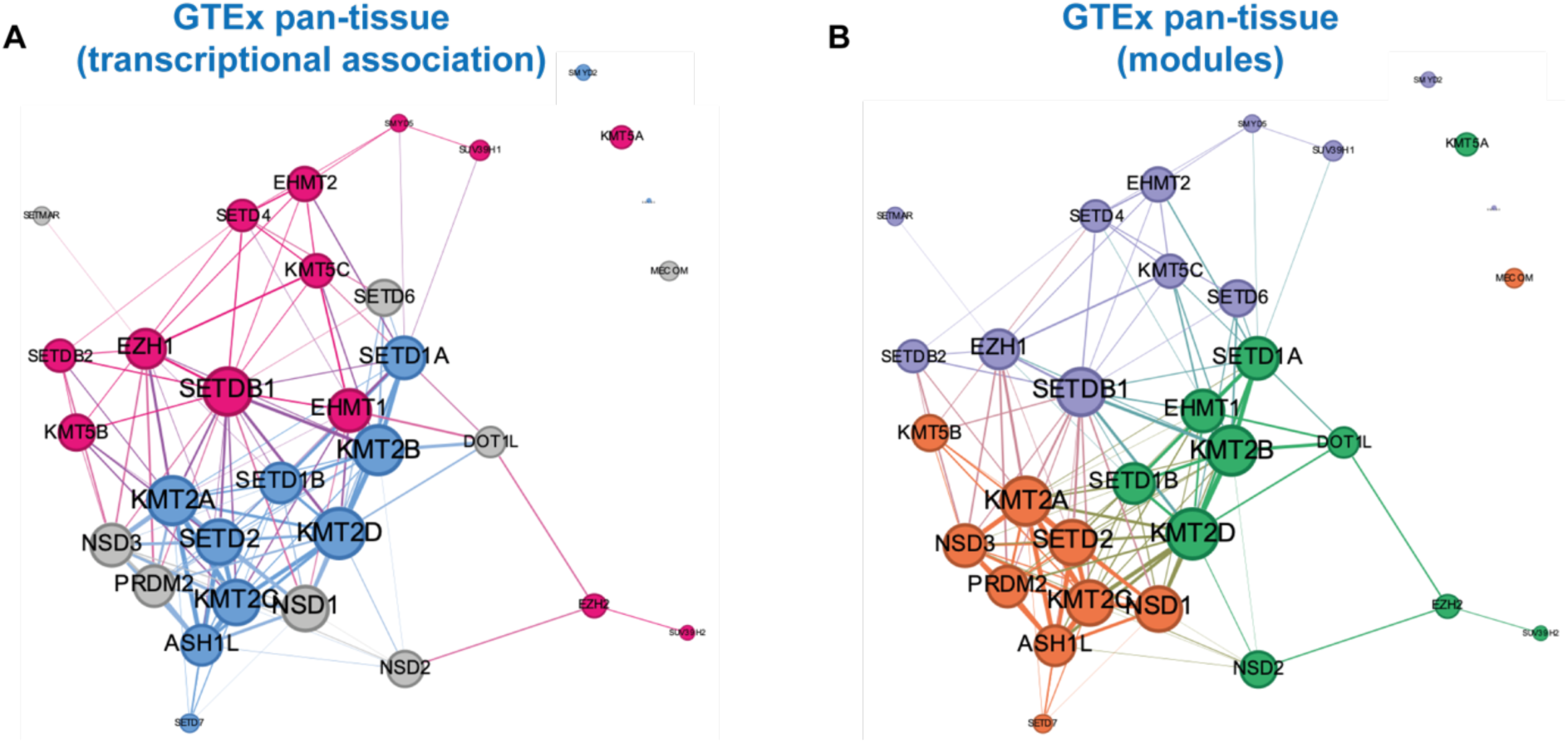
Histone methyltransferase correlation network in healthy tissues (related to Fig 4) A) Network plot shows all correlations between HMTs with magnitude > 0.2 in a GTEx pan-tissue analysis. Edge weight is proportional to the correlation strength. Node size is proportional to the weighted degree. Blue nodes indicate HMTs associated with transcriptional activation, magenta with transcriptional repression and grey with ambiguous association. Edge colours reflect the colours of the connected nodes. B) Network plot for GTEx pan-tissue analysis coloured to show distinct modules discovered using the Louvain method.

**Fig S9.**
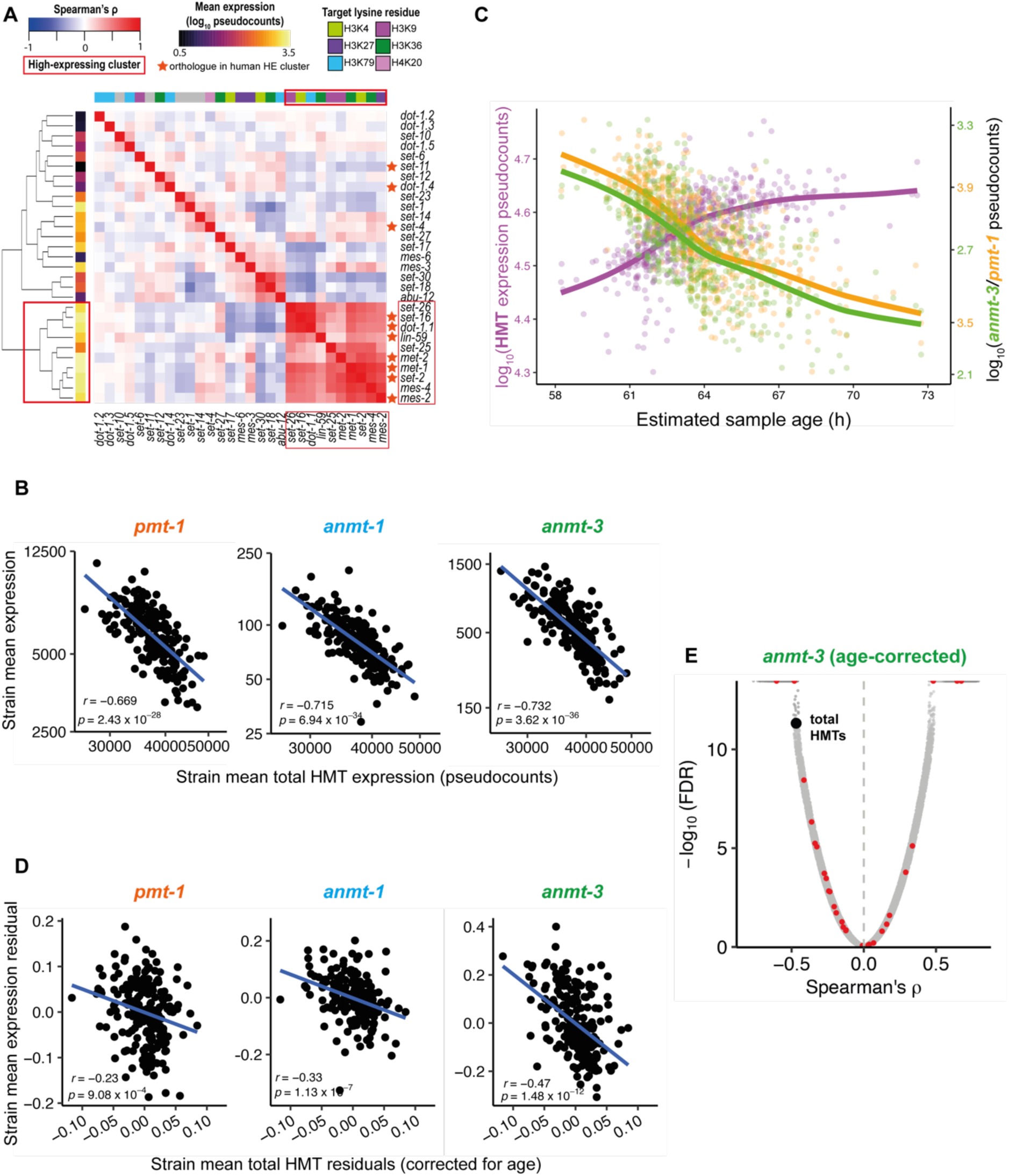
Histone methyltransferases are co-expressed and correlate with *NNMT/PEMT* orthologues/analogues in *Caenorhabditis elegans*. A) Heatmaps showing correlation of histone methyltransferase gene expression levels across 206 genetically diverse strains from the CeNDR resource. Sidebars display expression level (left) and target lysine residue (top – grey is unknown). Dendrogram shows hierarchical clustering of correlations. Red box indicates the cluster of highly-expressed and correlated HMTs. Orange stars indicate orthologues of human genes in the human highly-expressed cluster. B) Scatterplots of expression of the *PEMT* analogue *pmt-1* and the *NNMT* orthologues *anmt-1* and *anmt-3* against total HMT expression. C) Total HMT (purple, left y axis), *pmt-1* (orange, right y axis) and *anmt-3* (green, right y axis) expression against age of *C. elegans* sample inferred from RNA-seq data. D) Scatterplots of expression of the *PEMT* analogue *pmt-1* and the *NNMT* orthologues *anmt-1* and *anmt-3* against total HMT expression after correcting for inferred age. E) Volcano plot showing Spearman’s correlation and FDR for expression of *anmt-3* vs 29807 genes. HMT-encoding genes are shown as red points; correlation for total HMT expression is shown as a black point.

## REFERENCES

1. T. Kouzarides, Histone methylation in transcriptional control. Current Opinion in Genetics & Development 12, 198–209 (2002).

2. T. Jenuwein, C. D. Allis, Translating the histone code. Science 293, 1074–1080 (2001).

3. S. L. Berger, The complex language of chromatin regulation during transcription. Nature 447, 407–412 (2007).

4. E. L. Greer, Y. Shi, Histone methylation: a dynamic mark in health, disease and inheritance. Nature Reviews Genetics 13, 343–357 (2012).

5. E. J. Wagner, P. B. Carpenter, Understanding the language of Lys36 methylation at histone H3. Nature Reviews Molecular Cell Biology 13, 115–126 (2012).

6. F. S. Howe, H. Fischl, S. C. Murray, J. Mellor, Is H3K4me3 instructive for transcription activation? Bioessays 39, 1–12 (2017).

7. A. Kinnaird, S. Zhao, K. E. Wellen, E. D. Michelakis, Metabolic control of epigenetics in cancer. Nature Reviews Cancer 16, 694–707 (2016).

8. N. Zhang, Role of methionine on epigenetic modification of DNA methylation and gene expression in animals. Animal Nutrition 4, 11–16 (2018).

9. C. Ye, B. P. Tu, Sink into the epigenome: histones as repositories that influence cellular metabolism. Trends in Endocrinology & Metabolism 29, 626–637 (2018).

10. H. Li et al., The landscape of cancer cell line metabolism. Nature Medicine 25, 850–860 (2019).

11. O. A. Ulanovskaya, A. M. Zuhl, B. F. Cravatt, NNMT promotes epigenetic remodeling in cancer by creating a metabolic methylation sink. Nature Chemical Biology 9, 300–306 (2013).

12. P. Pissios, Nicotinamide N-methyltransferase: more than a vitamin B3 clearance enzyme. Trends in Endocrinology & Metabolism 28, 340–353 (2017).

13. D. P. Nusinow et al., Quantitative proteomics of the cancer cell line encyclopedia. Cell 180, 387–402. e316 (2020).

14. D. Kraus et al., Nicotinamide N-methyltransferase knockdown protects against diet-induced obesity. Nature 508, 258–262 (2014).

15. H. Sperber et al., The metabolome regulates the epigenetic landscape during naive-to-primed human embryonic stem cell transition. Nature Cell Biology 17, 1523–1535 (2015).

16. M. A. Eckert et al., Proteomics reveals NNMT as a master metabolic regulator of cancer-associated fibroblasts. Nature 569, 723–728 (2019).

17. T. Guo et al., Quantitative proteome landscape of the NCI-60 cancer cell lines. Iscience 21, 664–680 (2019).

18. M. F. Perez, P. Sarkies, Malignancy and NF-κB signalling strengthen coordination between expression of mitochondrial and nuclear-encoded oxidative phosphorylation genes. Genome Biology 22, 1–24 (2021).

19. M. Roeßler et al., Identification of nicotinamide N-methyltransferase as a novel serum tumor marker for colorectal cancer. Clinical Cancer Research 11, 6550–6557 (2005).

20. B.-H. Lim et al., Overexpression of nicotinamide N-methyltransferase in gastric cancer tissues and its potential post-translational modification. Experimental & Molecular Medicine 38, 455–465 (2006).

21. Y. Wu, M. Siadaty, M. Berens, G. Hampton, D. Theodorescu, Overlapping gene expression profiles of cell migration and tumor invasion in human bladder cancer identify metallothionein 1E and nicotinamide N-methyltransferase as novel regulators of cell migration. Oncogene 27, 6679–6689 (2008).

22. J. Kim et al., Expression of nicotinamide N-methyltransferase in hepatocellular carcinoma is associated with poor prognosis. Journal of Experimental & Clinical Cancer Research 28, 1–9 (2009).

23. M. Tomida, I. Mikami, S. Takeuchi, H. Nishimura, H. Akiyama, Serum levels of nicotinamide N-methyltransferase in patients with lung cancer. Journal of Cancer Research and Clinical Oncology 135, 1223–1229 (2009).

24. S.-W. Tang et al., Nicotinamide N-methyltransferase induces cellular invasion through activating matrix metalloproteinase-2 expression in clear cell renal cell carcinoma cells. Carcinogenesis 32, 138–145 (2011).

25. W. Zhou et al., Nicotinamide N-methyltransferase is overexpressed in prostate cancer and correlates with prolonged progression-free and overall survival times. Oncology Letters 8, 1175–1180 (2014).

26. İ. Harmankaya et al., Nicotinamide N-methyltransferase overexpression may be associated with poor prognosis in ovarian cancer. Journal of Obstetrics and Gynaecology 41, 248–253 (2021).

27. G. Van Meer, D. R. Voelker, G. W. Feigenson, Membrane lipids: where they are and how they behave. Nature Reviews Molecular Cell Biology 9, 112–124 (2008).

28. C. J. Pynn et al., Specificity and rate of human and mouse liver and plasma phosphatidylcholine synthesis analyzed in vivo [S]. Journal of Lipid Research 52, 399–407 (2011).

29. L. M. Stead, J. T. Brosnan, M. E. Brosnan, D. E. Vance, R. L. Jacobs, Is it time to reevaluate methyl balance in humans? The American Journal of Clinical Nutrition 83, 5–10 (2006).

30. K. R. Stewart-Morgan, N. Petryk, A. Groth, Chromatin replication and epigenetic cell memory. Nature Cell Biology 22, 361–371 (2020).

31. S. Jackowski, Coordination of membrane phospholipid synthesis with the cell cycle. Journal of Biological Chemistry 269, 3858–3867 (1994).

32. I. Tseu, R. Ridsdale, J. Liu, J. Wang, M. Post, Cell cycle regulation of pulmonary phosphatidylcholine synthesis. American Journal of Respiratory Cell and Molecular Biology 26, 506–515 (2002).

33. R. C. Ramaker et al., RNA sequencing-based cell proliferation analysis across 19 cancers identifies a subset of proliferation-informative cancers with a common survival signature. Oncotarget 8, 38668 (2017).

34. V. Reinke, M. Krause, P. Okkema, Transcriptional regulation of gene expression in C. elegans. WormBook: The Online Review of C. elegans Biology (2018).

35. S. Polager, D. Ginsberg, E2F–at the crossroads of life and death. Trends in Cell Biology 18, 528–535 (2008).

36. P. D. Goetsch, J. M. Garrigues, S. Strome, Loss of the Caenorhabditis elegans pocket protein LIN-35 reveals MuvB’s innate function as the repressor of DREAM target genes. PLoS Genetics 13, e1007088 (2017).

37. I. Latorre et al., The DREAM complex promotes gene body H2A. Z for target repression. Genes & Development 29, 495–500 (2015).

38. C. Gal et al., DREAM represses distinct targets by cooperating with different THAP domain proteins. Cell Reports 37, 109835 (2021).

39. P. Badia-i-Mompel et al., decoupleR: ensemble of computational methods to infer biological activities from omics data. Bioinformatics Advances 2, vbac016 (2022).

40. N. Reverón-Gómez et al., Accurate recycling of parental histones reproduces the histone modification landscape during DNA replication. Molecular Cell 72, 239–249. e235 (2018).

41. J. Zhu et al., Transsulfuration activity can support cell growth upon extracellular cysteine limitation. Cell Metabolism 30, 865–876. e865 (2019).

42. H.-F. Zhang, R. I. K. Geltink, S. J. Parker, P. H. Sorensen, Transsulfuration, minor player or crucial for cysteine homeostasis in cancer. Trends in Cell Biology, (2022).

43. B. Henneman, C. Van Emmerik, H. van Ingen, R. T. Dame, Structure and function of archaeal histones. PLoS Genetics 14, e1007582 (2018).

44. M. I. Love, W. Huber, S. Anders, Moderated estimation of fold change and dispersion for RNA-seq data with DESeq2. Genome Biology 15, 1–21 (2014).

45. F. Ramírez, F. Dündar, S. Diehl, B. A. Grüning, T. Manke, deepTools: a flexible platform for exploring deep-sequencing data. Nucleic Acids Research 42, W187–W191 (2014).

46. L. K. Gopi, B. L. Kidder, Integrative pan cancer analysis reveals epigenomic variation in cancer type and cell specific chromatin domains. Nature Communications 12, 1–20 (2021).

47. B. Langmead, S. L. Salzberg, Fast gapped-read alignment with Bowtie 2. Nature Methods 9, 357–359 (2012).

48. R. N. Shah, A. J. Ruthenburg, Sequence deeper without sequencing more: Bayesian resolution of ambiguously mapped reads. PLoS Computational Biology 17, e1008926 (2021).

49. Y. Zhang et al., Model-based analysis of ChIP-Seq (MACS). Genome Biology 9, 1–9 (2008).

50. M. L. Bendall et al., Telescope: Characterization of the retrotranscriptome by accurate estimation of transposable element expression. PLoS Computational Biology 15, e1006453 (2019).

51. K. Narasimhan et al., Mapping and analysis of Caenorhabditis elegans transcription factor sequence specificities. Elife 4, e06967 (2015).

52. M. T. Weirauch et al., Determination and inference of eukaryotic transcription factor sequence specificity. Cell 158, 1431–1443 (2014).

53. G. Zhang, N. M. Roberto, D. Lee, S. R. Hahnel, E. C. Andersen, The impact of species-wide gene expression variation on Caenorhabditis elegans complex traits. Nature Communications 13, 3462 (2022).

54. R. Bulteau, M. Francesconi, Real age prediction from the transcriptome with RAPToR. Nature Methods 19, 969–975 (2022).

55. W. Kim, R. S. Underwood, I. Greenwald, D. D. Shaye, OrthoList 2: a new comparative genomic analysis of human and Caenorhabditis elegans genes. Genetics 210, 445–461 (2018).

56. A. Colaprico et al., TCGAbiolinks: an R/Bioconductor package for integrative analysis of TCGA data. Nucleic Acids Research 44, e71–e71 (2016).

57. L. Garcia-Alonso, C. H. Holland, M. M. Ibrahim, D. Turei, J. Saez-Rodriguez, Benchmark and integration of resources for the estimation of human transcription factor activities. Genome Research 29, 1363–1375 (2019).

58. A. D. Rouillard et al., The harmonizome: a collection of processed datasets gathered to serve and mine knowledge about genes and proteins. Database 2016, (2016).

