## Supplemental Figures and Tables for "Histone methylation has a direct metabolic role in human cells": Supp File 1. CCLE NNMT HMTs fullplots.pdf

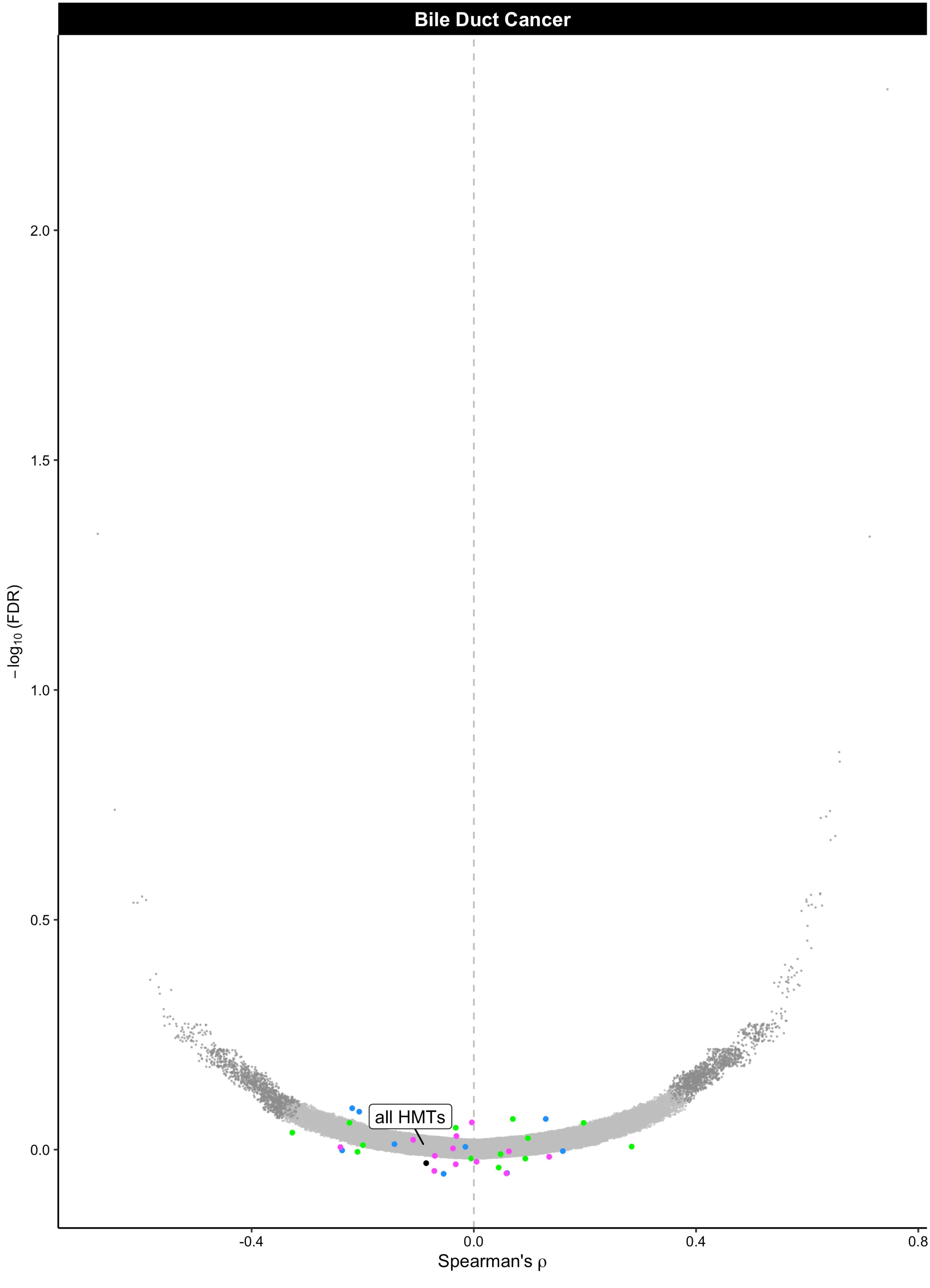

Bile Duct Cancer
all HMTs
2.0
1.5
1.0
0.5
0.0
−log10 (FDR)
-0.4
0.0
0.4
0.8
Spearman's ρ

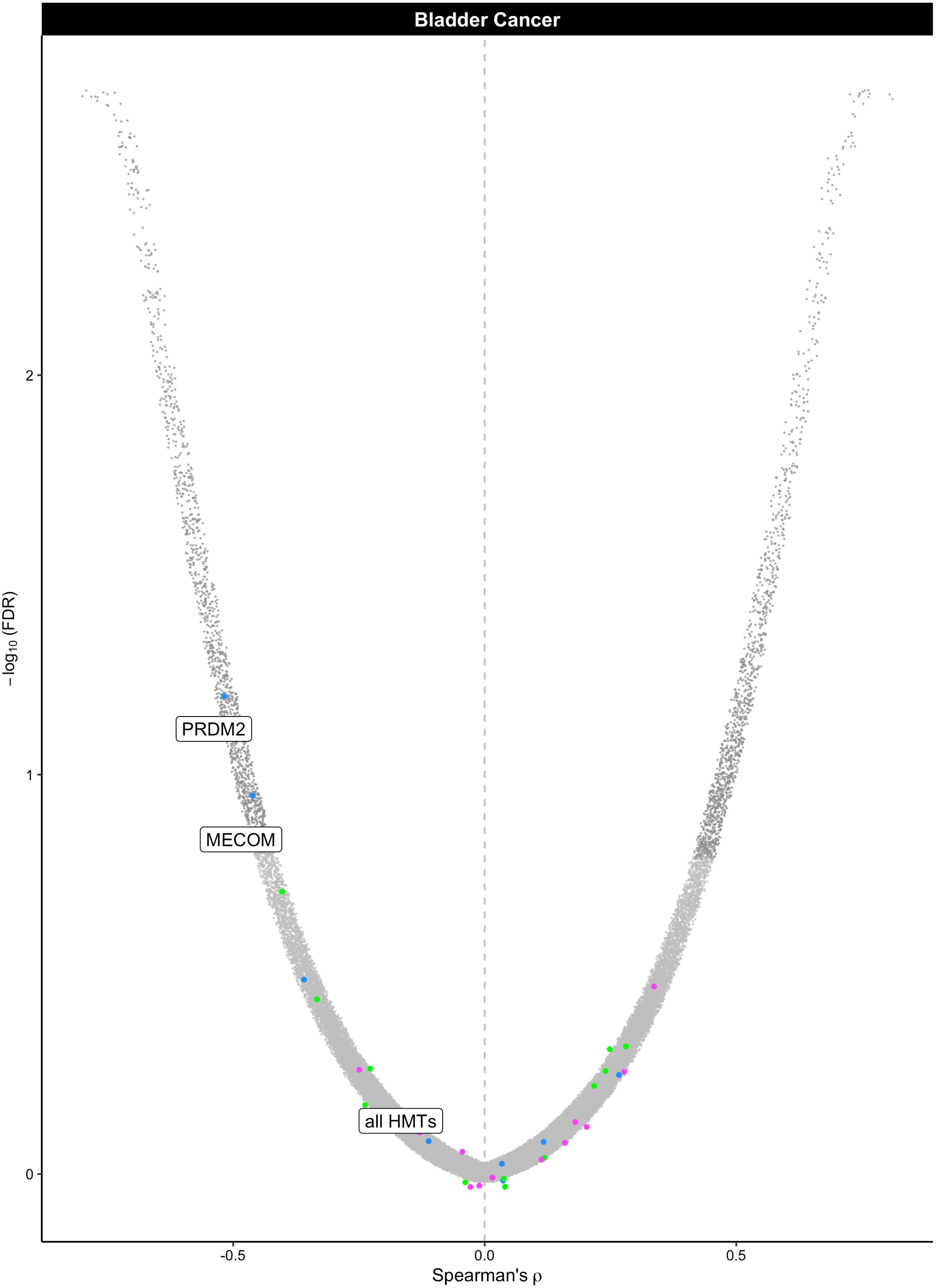

Bladder Cancer
PRDM2
MECOM
all HMTs
−log10 (FDR)
2
1
0
−0.5
0.0
0.5
Spearman's ρ

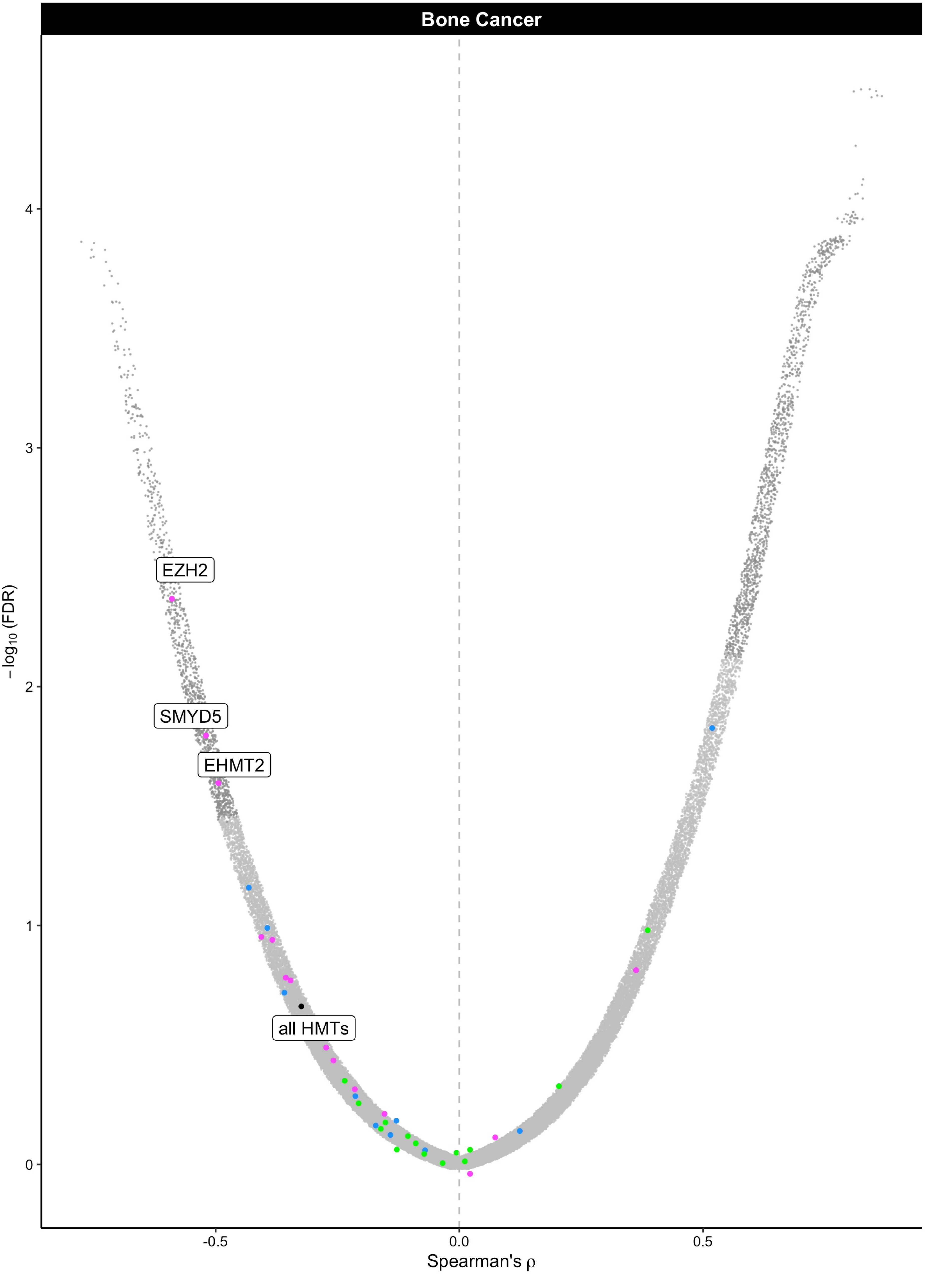

Bone Cancer
EZH2
SMYD5
EHMT2
all HMTs
−log10 (FDR)
Spearman's ρ
4
3
2
1
0
-0.5
0.0
0.5

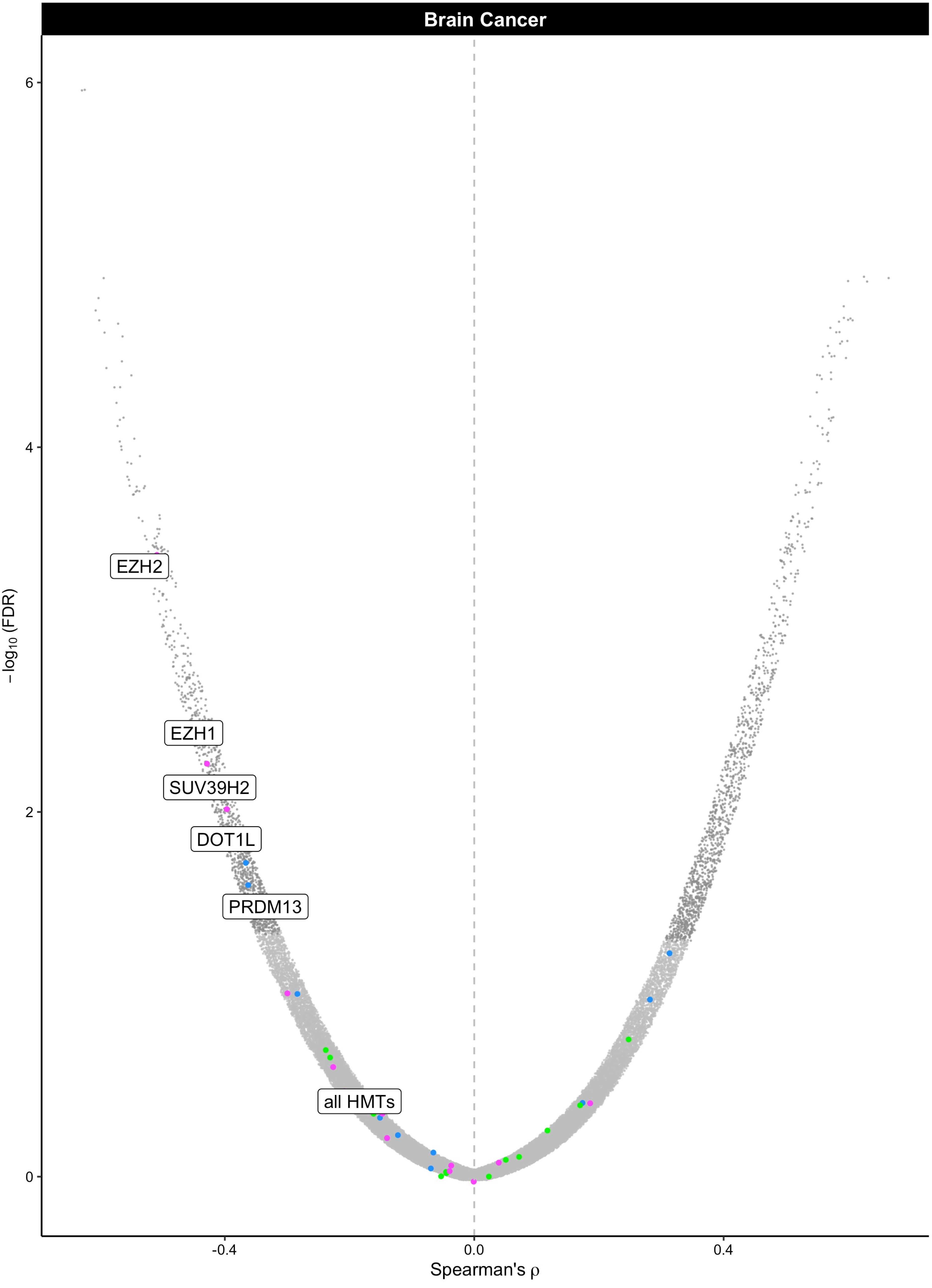

Brain Cancer
6
4
2
0
−log10 (FDR)
EZH2
EZH1
SUV39H2
DOT1L
PRDM13
all HMTs
-0.4
0.0
0.4
Spearman's ρ

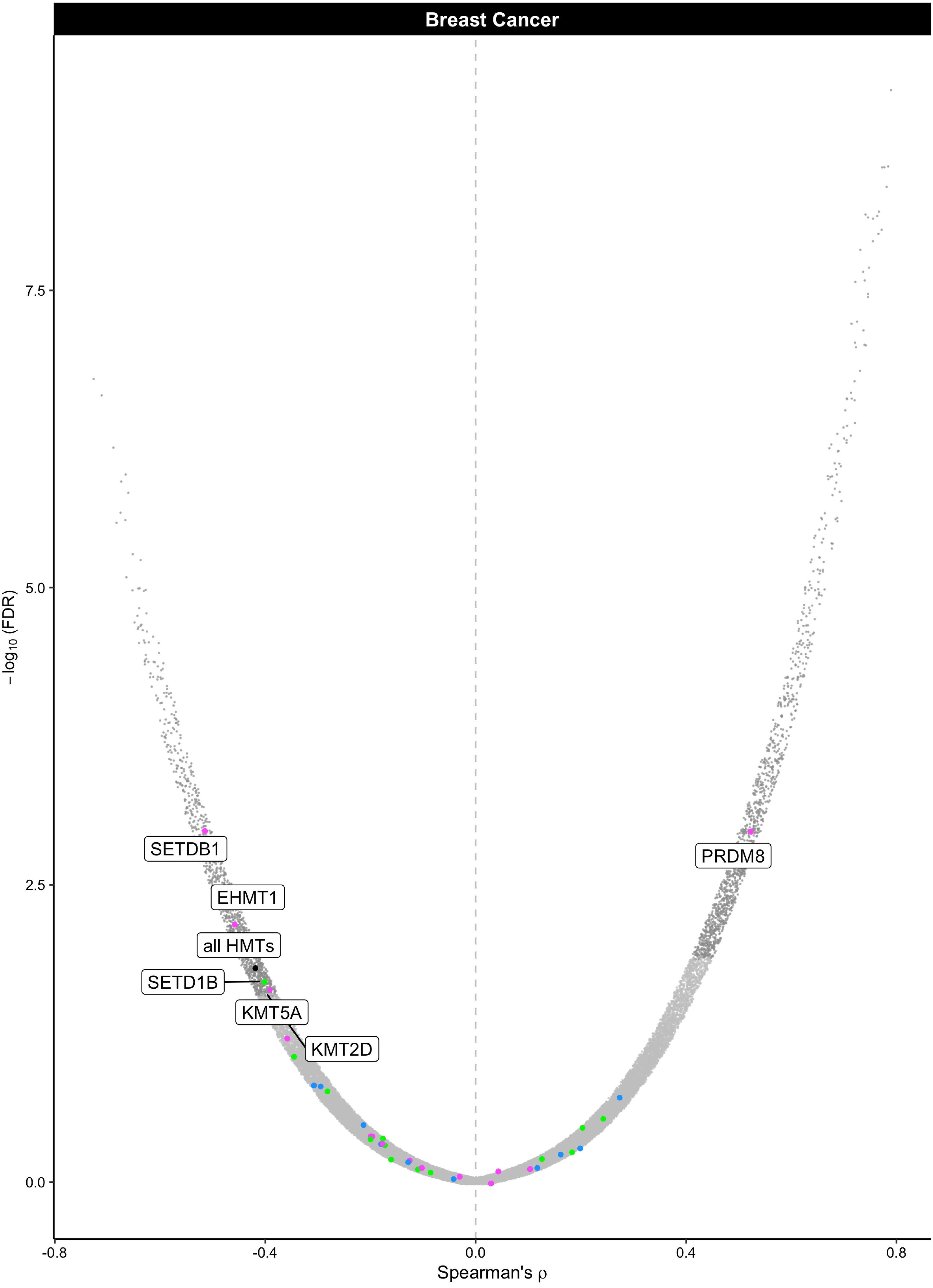
Breast Cancer
SETDB1
EHMT1
all HMTs
SETD1B
KMT5A
KMT2D
PRDM8
−log10 (FDR)
7.5
5.0
2.5
0.0
-0.8
-0.4
0.0
0.4
0.8
Spearman's ρ

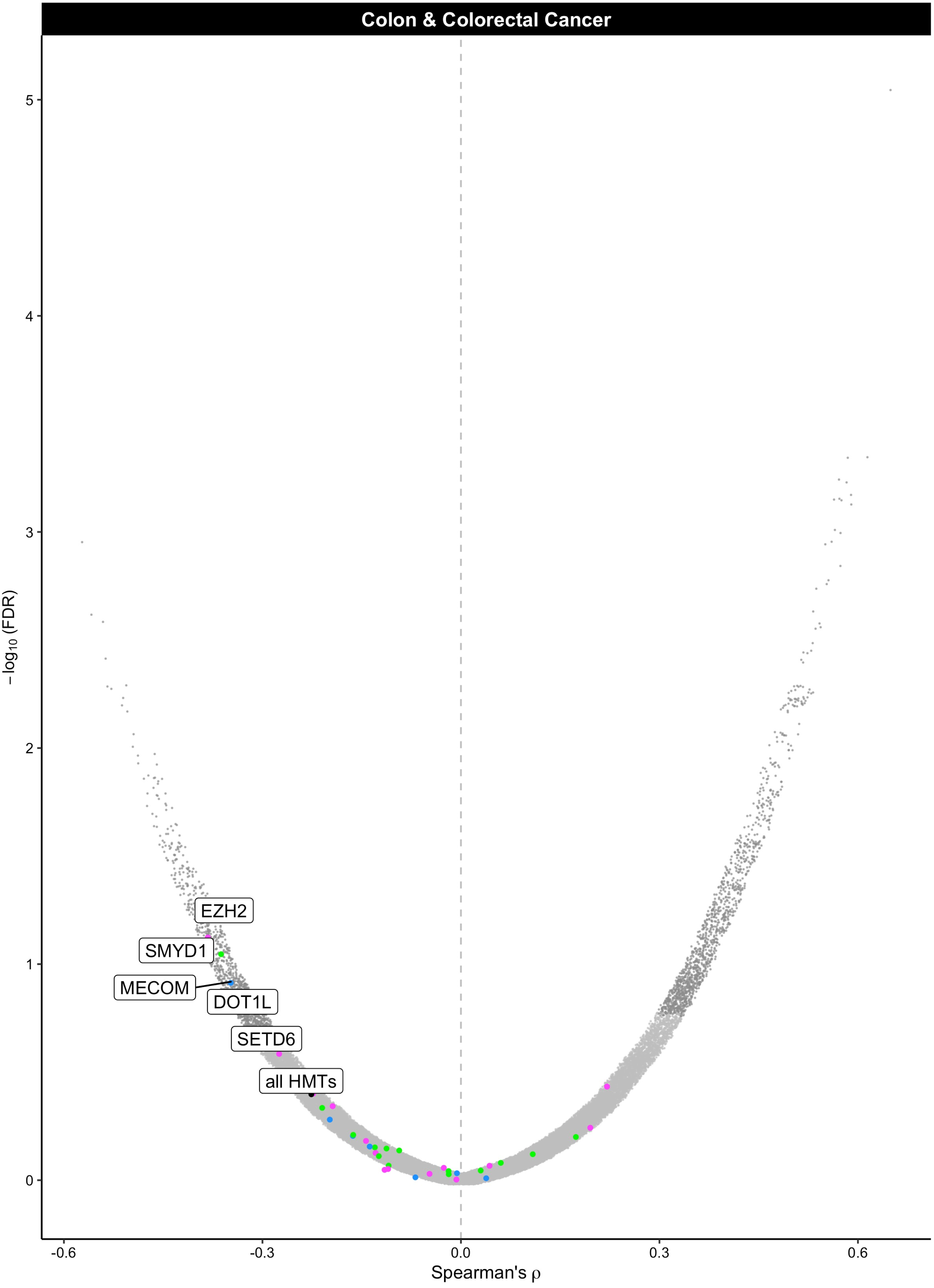
Colon & Colorectal Cancer
EZH2
SMYD1
MECOM
DOT1L
SETD6
all HMTs
−log10 (FDR)
Spearman's ρ
0
1
2
3
4
5
-0.6
-0.3
0.0
0.3
0.6

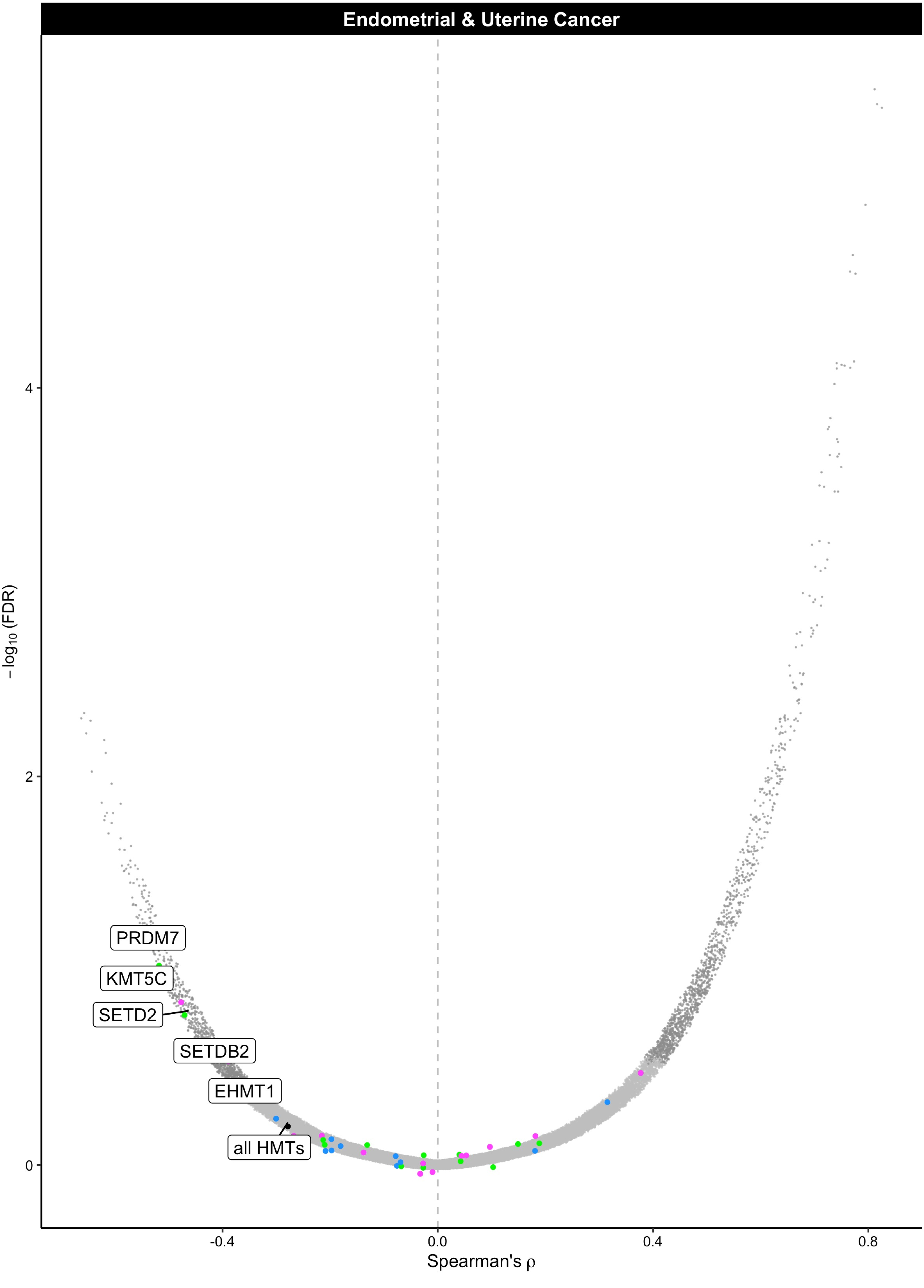
Endometrial & Uterine Cancer
PRDM7
KMT5C
SETD2
SETDB2
EHMT1
all HMTs
4
2
0
-0.4
0.0
0.4
0.8
Spearman's ρ
−log10 (FDR)

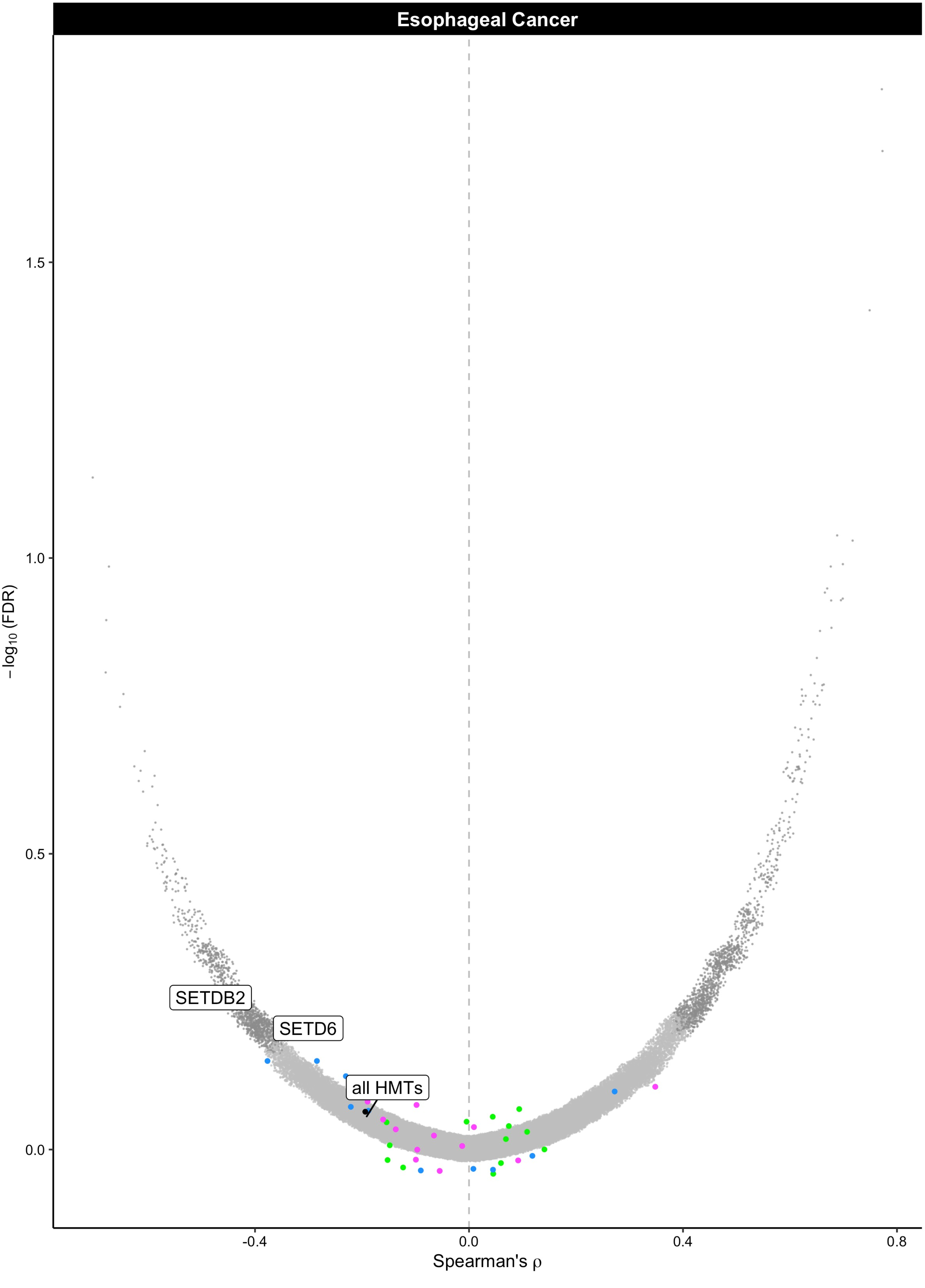

Esophageal Cancer
SETDB2
SETD6
all HMTs
1.5
1.0
0.5
0.0
−log10 (FDR)
-0.4
0.0
0.4
0.8
Spearman's ρ

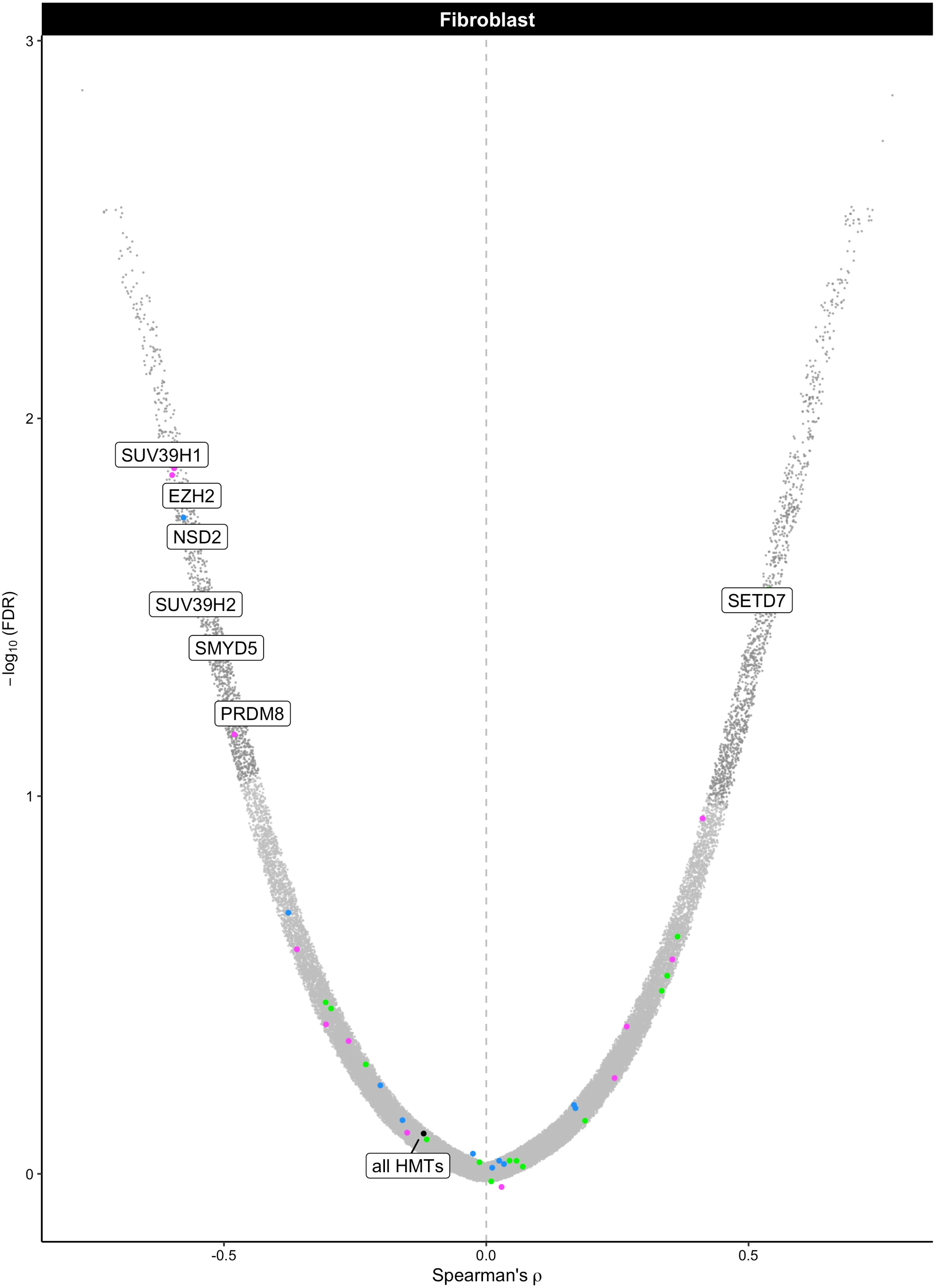

Fibroblast
3
2
1
0
−log10 (FDR)
SUV39H1
EZH2
NSD2
SUV39H2
SMYD5
PRDM8
SETD7
all HMTs
-0.5
0.0
0.5
Spearman's ρ

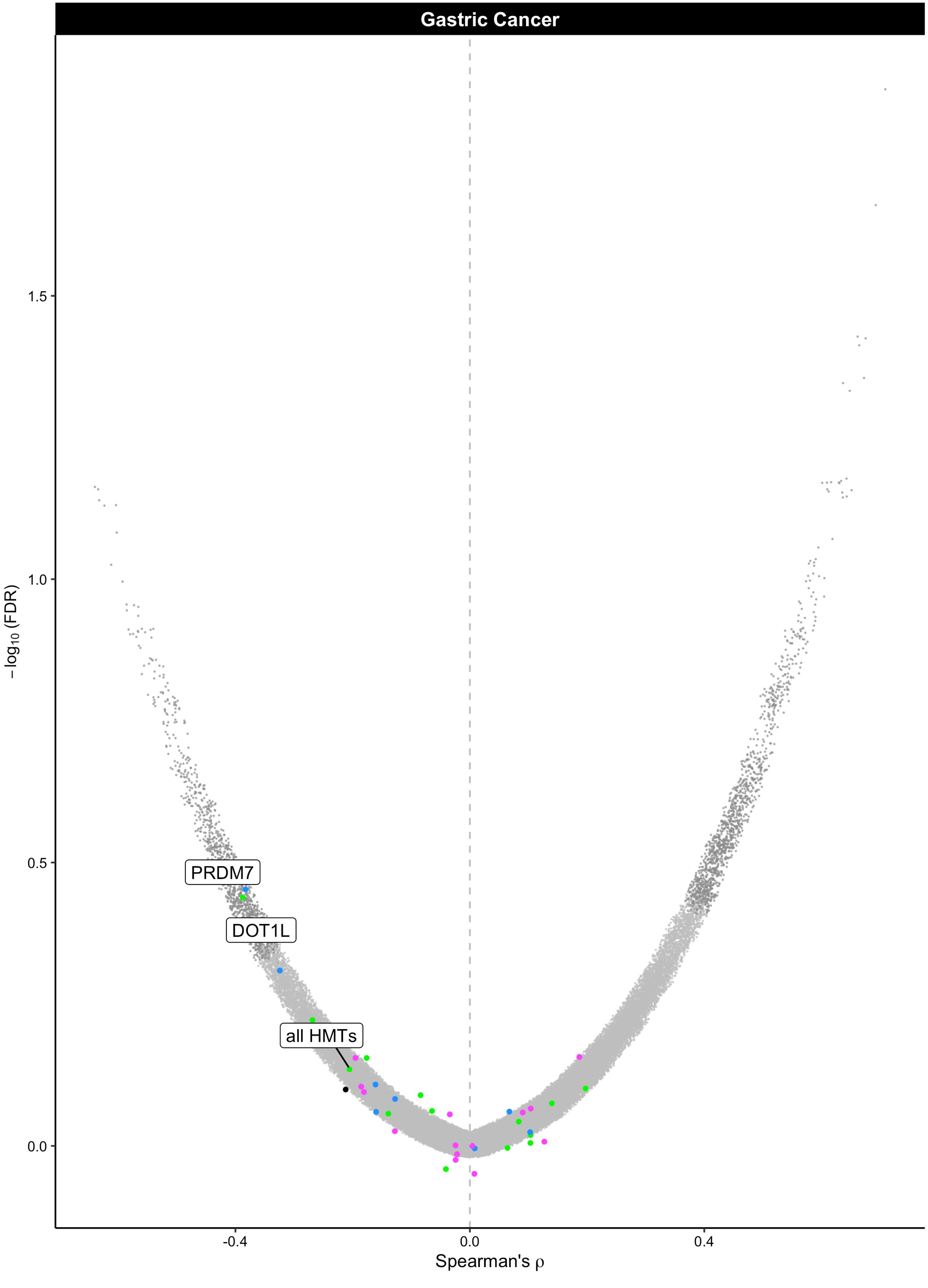
Gastric Cancer
PRDM7
DOT1L
all HMTs
1.5
1.0
0.5
0.0
-log10 (FDR)
-0.4
0.0
0.4
Spearman's ρ

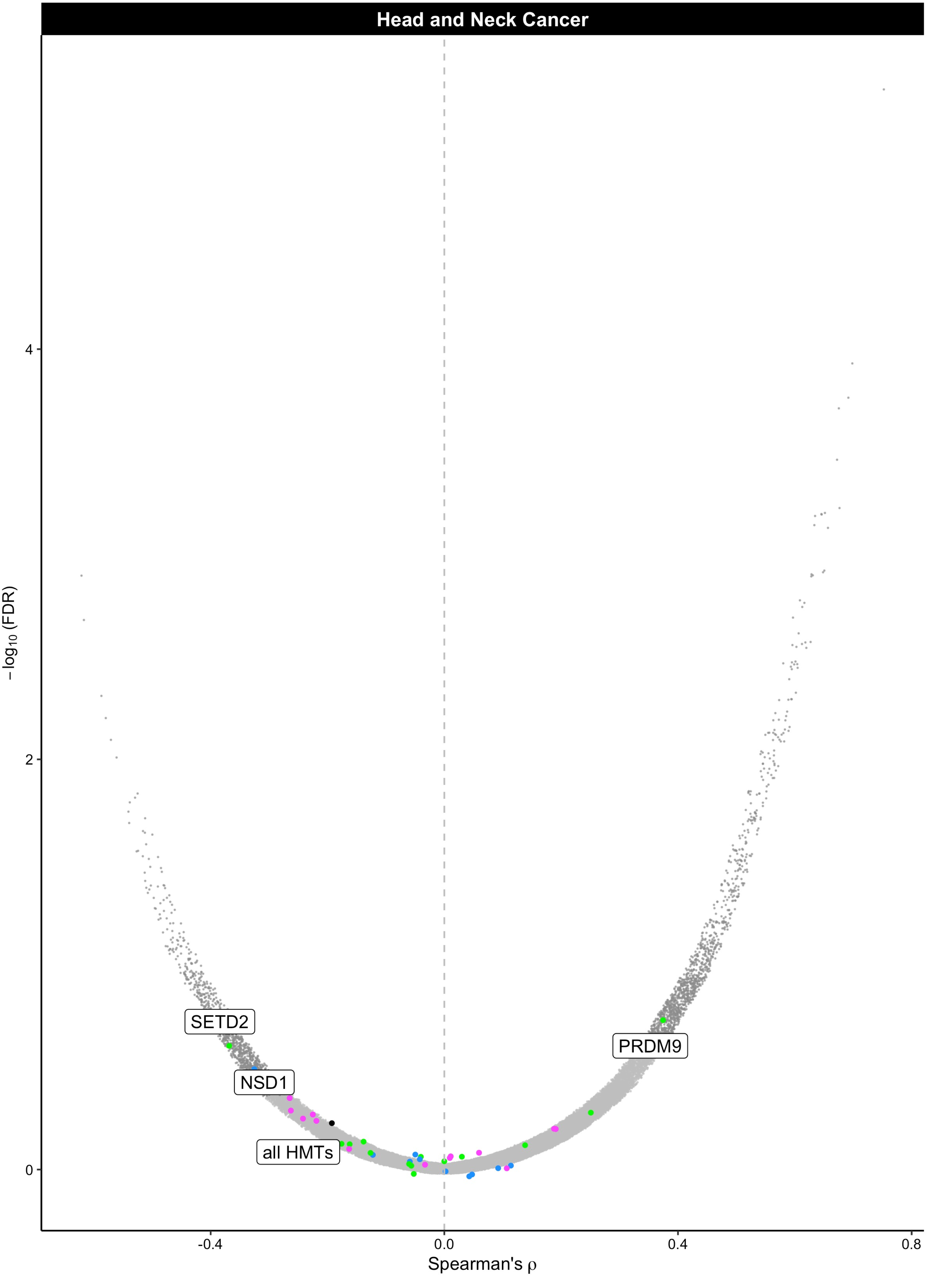

Head and Neck Cancer
−log10 (FDR)
4
2
0
SETD2
NSD1
all HMTs
PRDM9
-0.4
0.0
0.4
0.8
Spearman's ρ

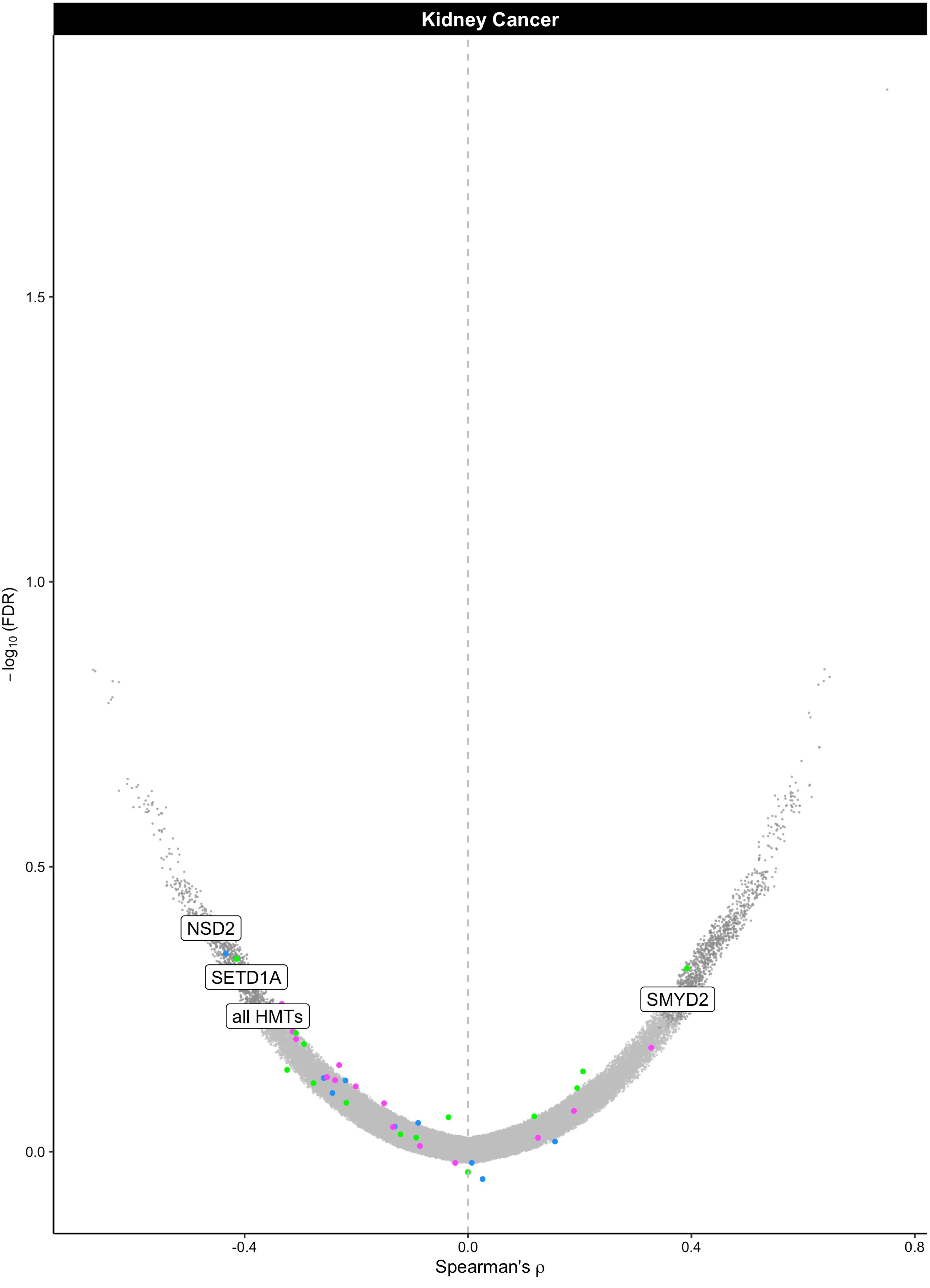

Kidney Cancer
NSD2
SETD1A
all HMTs
SMYD2
1.5
1.0
0.5
0.0
-0.4
0.0
0.4
0.8
−log10 (FDR)
Spearman's ρ

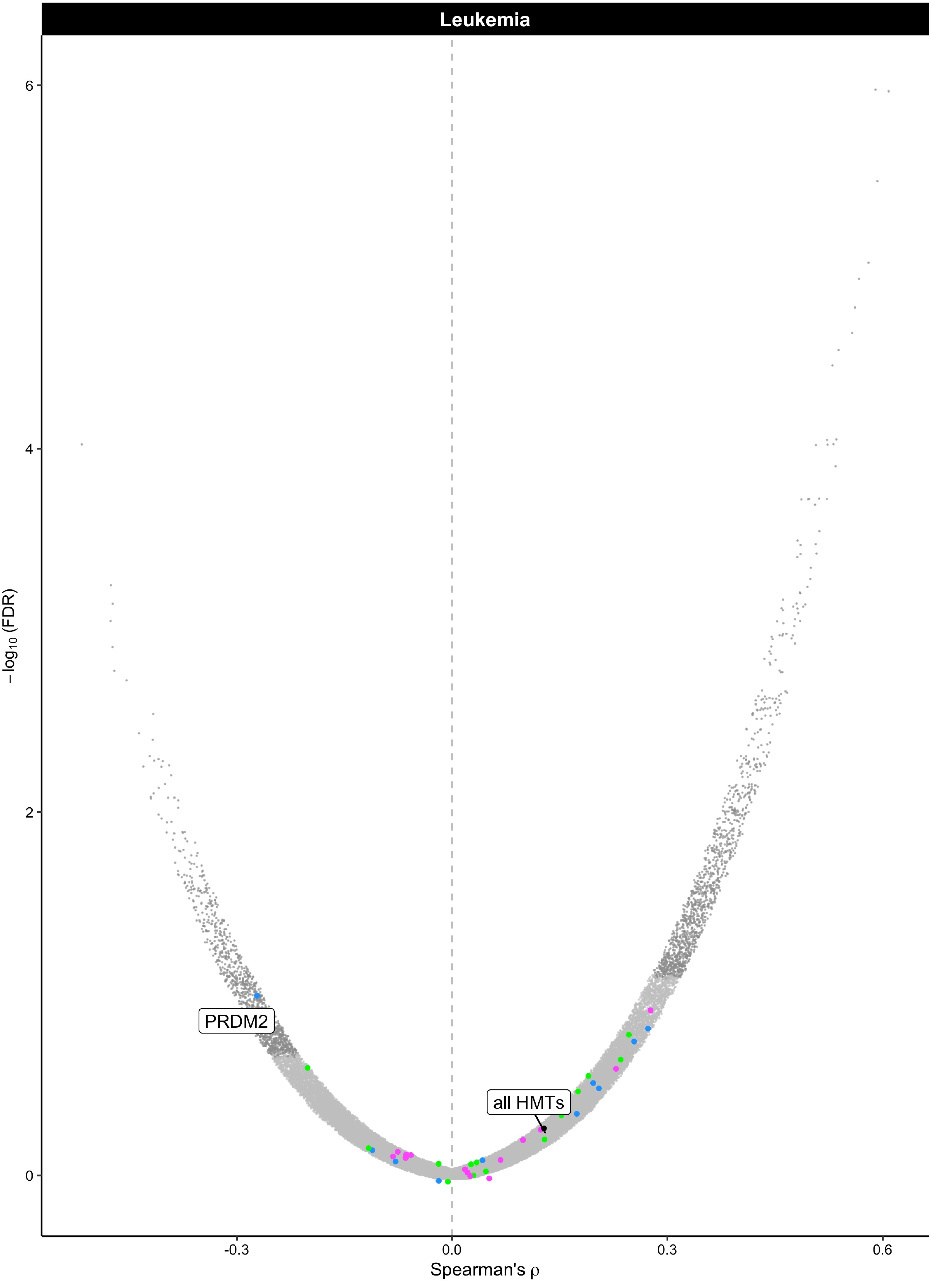

Leukemia
6
4
2
0
−log10 (FDR)
PRDM2
all HMTs
-0.3
0.0
0.3
0.6
Spearman's ρ

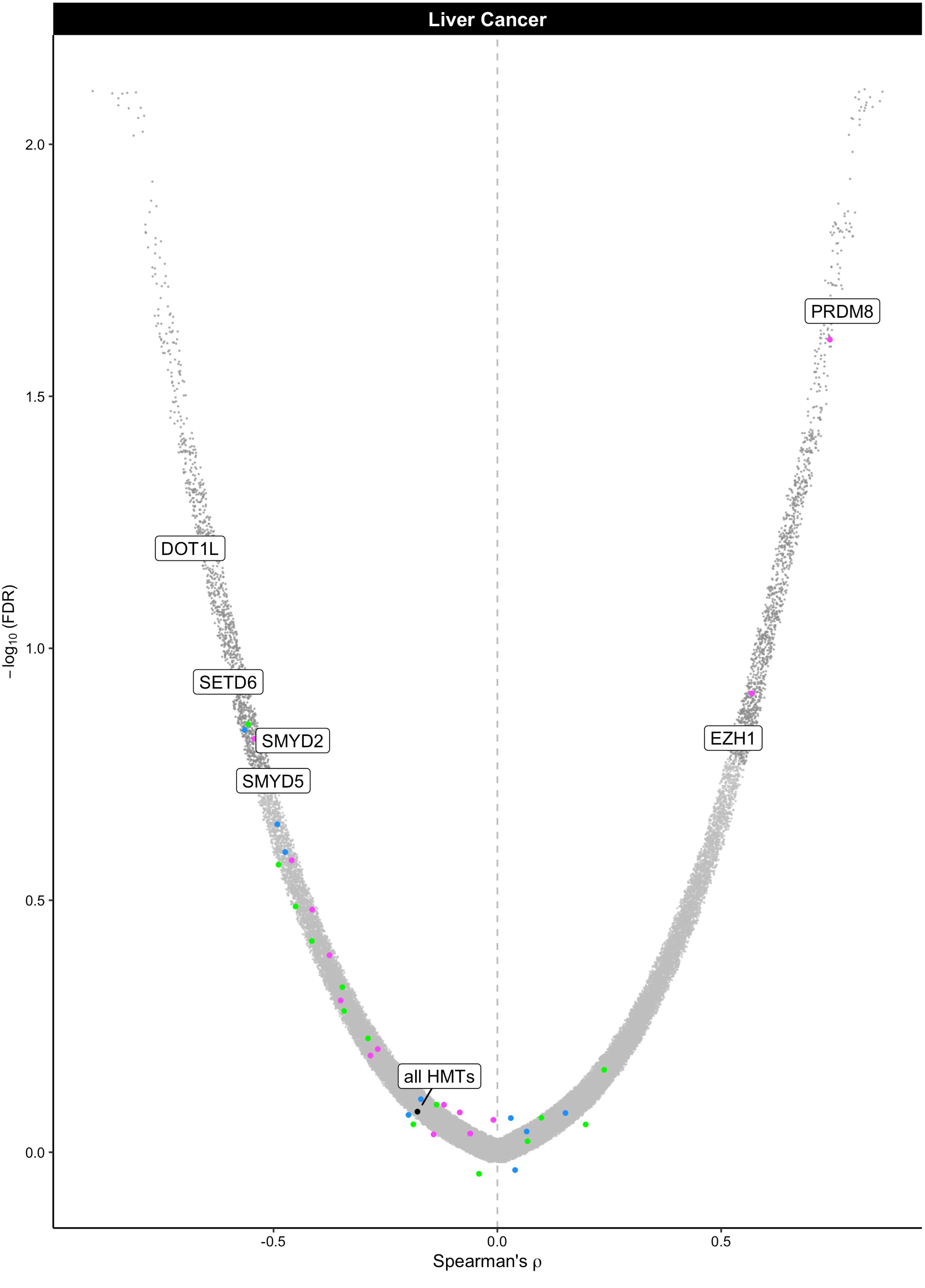

Liver Cancer
PRDM8
DOT1L
SETD6
SMYD2
SMYD5
EZH1
all HMTs
2.0
1.5
1.0
0.5
0.0
−log10 (FDR)
-0.5
0.0
0.5
Spearman's ρ

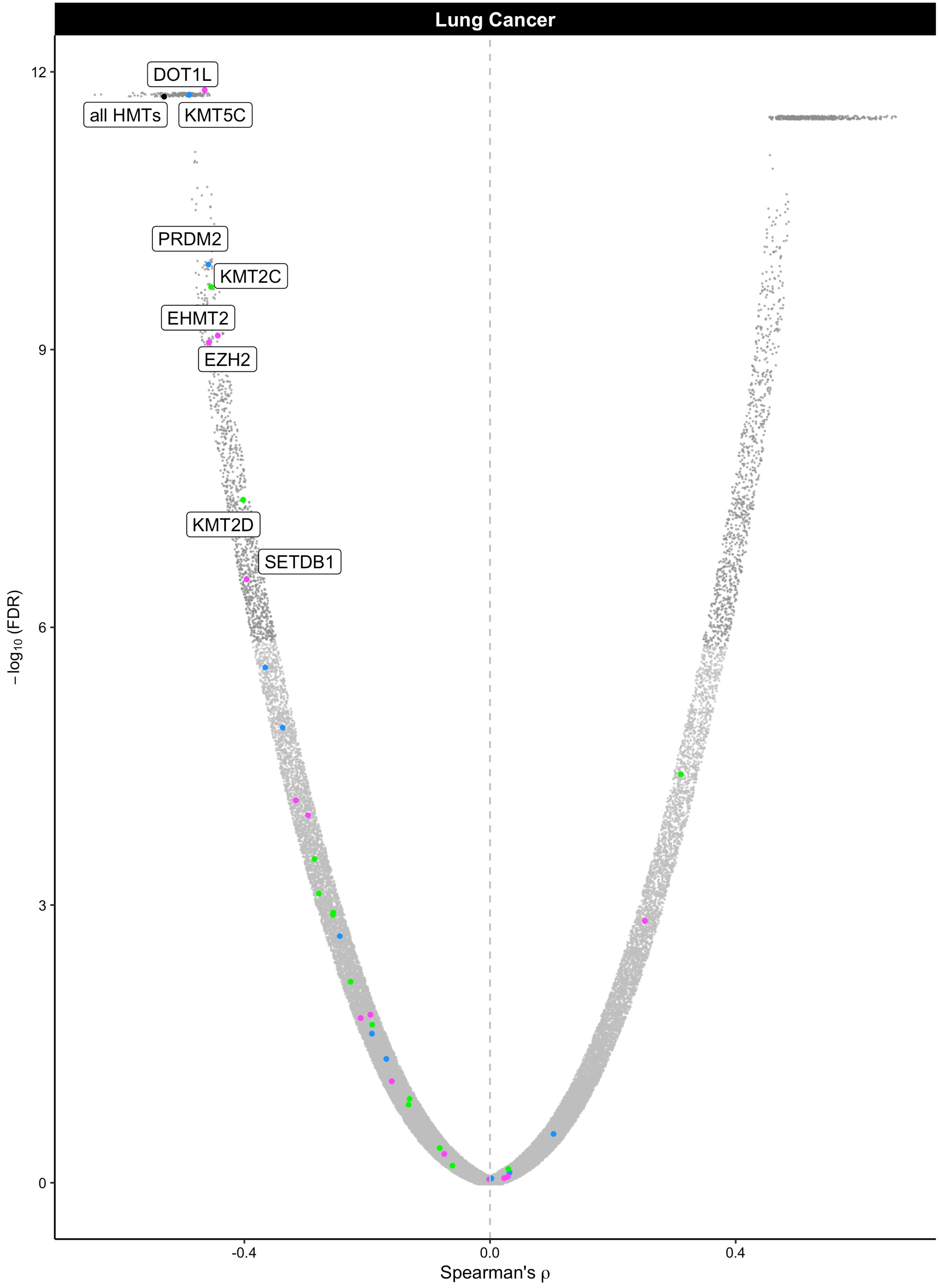
Lung Cancer
DOT1L
all HMTs
KMT5C
PRDM2
KMT2C
EHMT2
EZH2
KMT2D
SETDB1
12
9
6
3
0
−log10 (FDR)
-0.4
0.0
0.4
Spearman's ρ

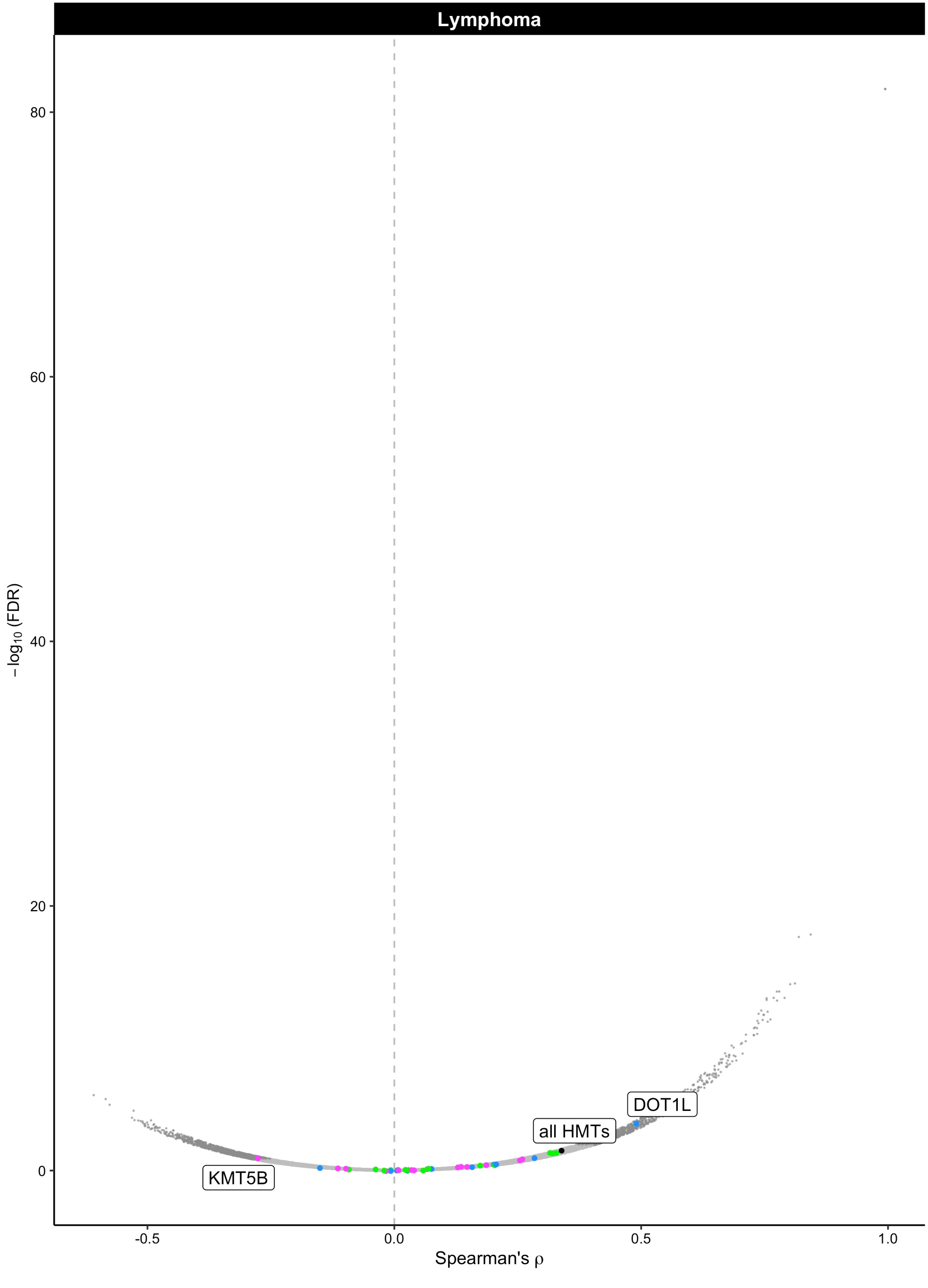

Lymphoma
80
60
40
20
0
−log10 (FDR)
DOT1L
all HMTs
KMT5B
-0.5
0.0
0.5
1.0
Spearman's ρ

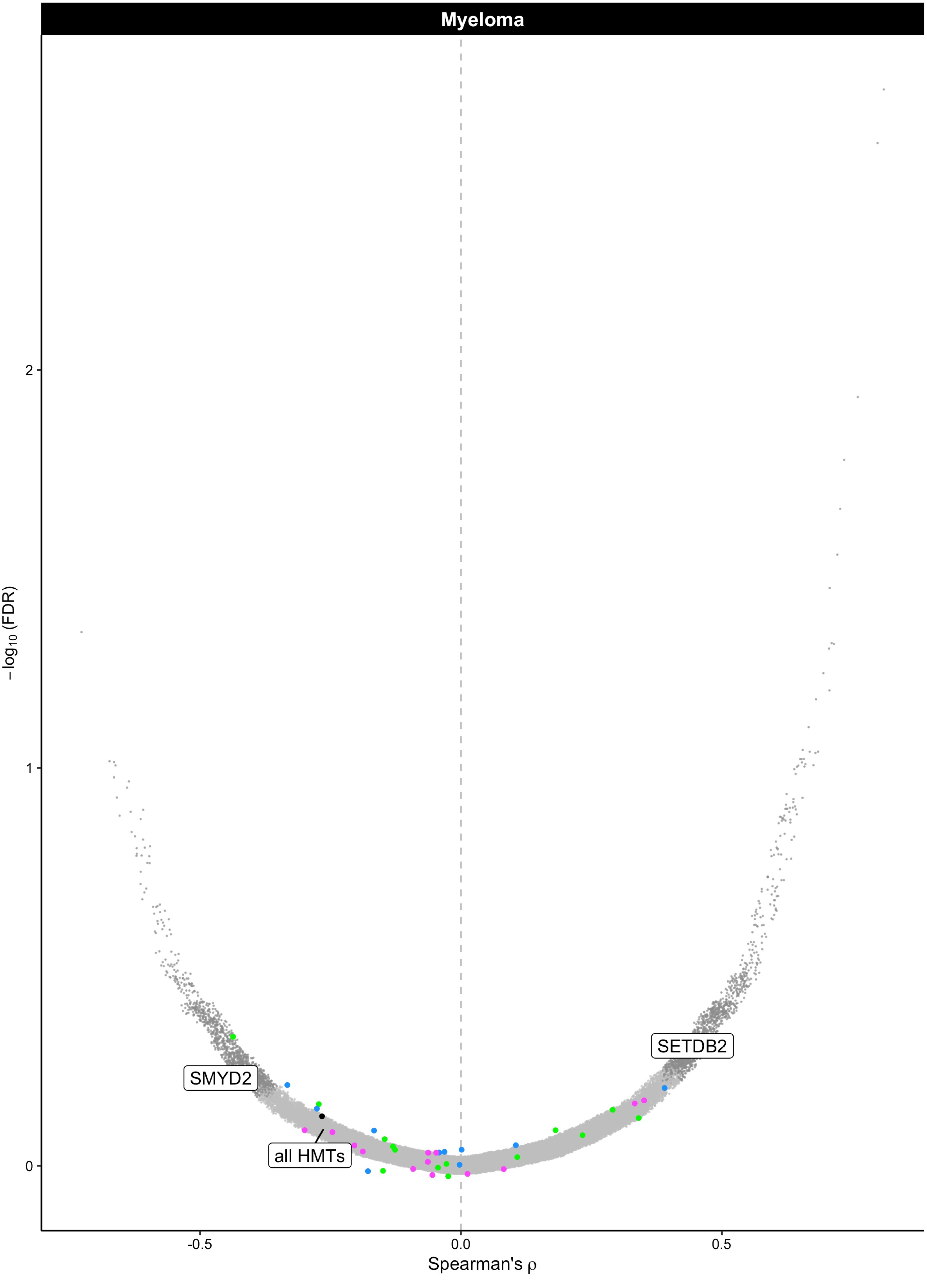

Myeloma
SETDB2
SMYD2
all HMTs
2
1
0
−log10 (FDR)
-0.5
0.0
0.5
Spearman's ρ

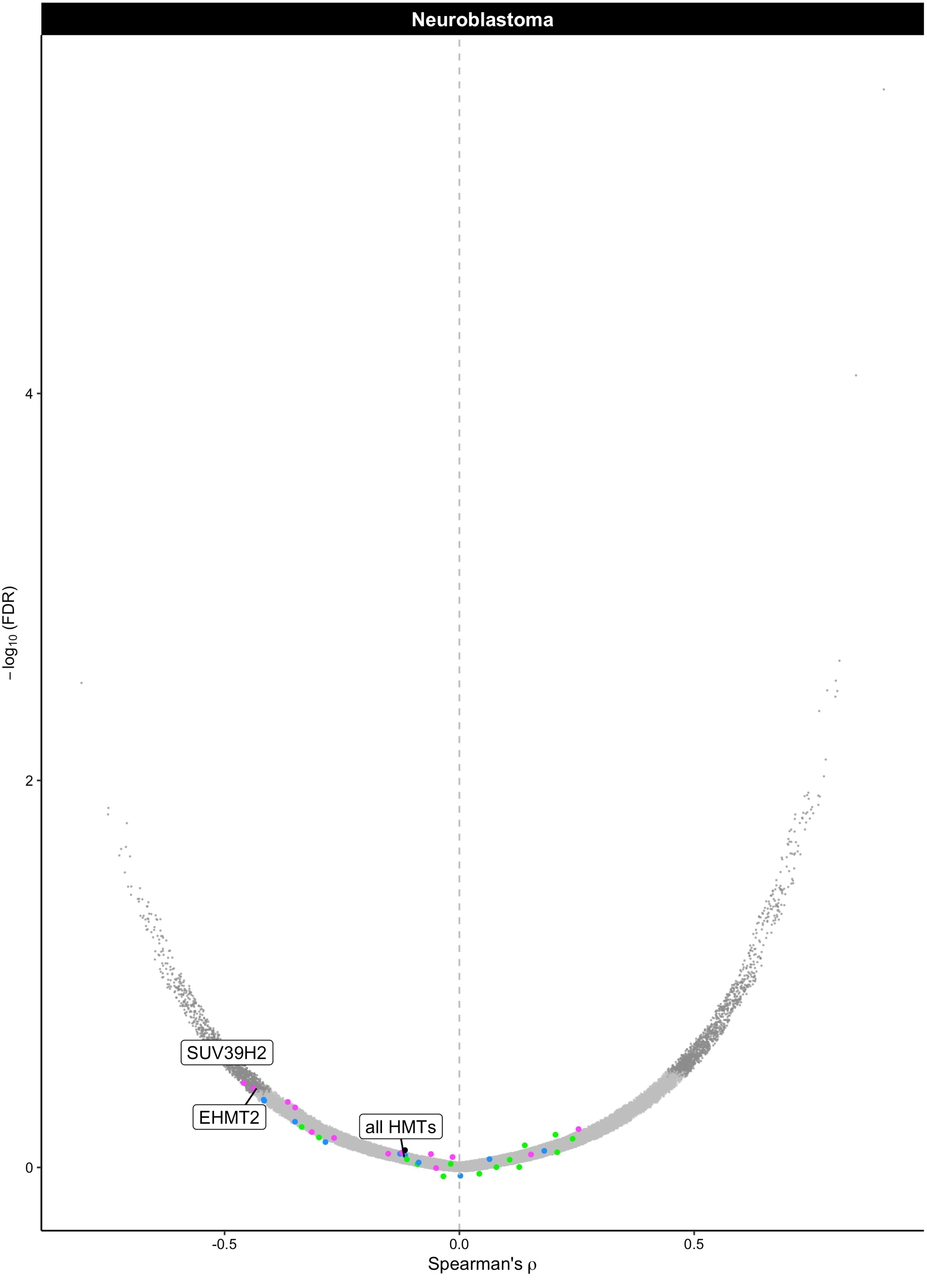

Neuroblastoma
4
2
0
−log10 (FDR)
-0.5
0.0
0.5
Spearman's ρ
SUV39H2
EHMT2
all HMTs

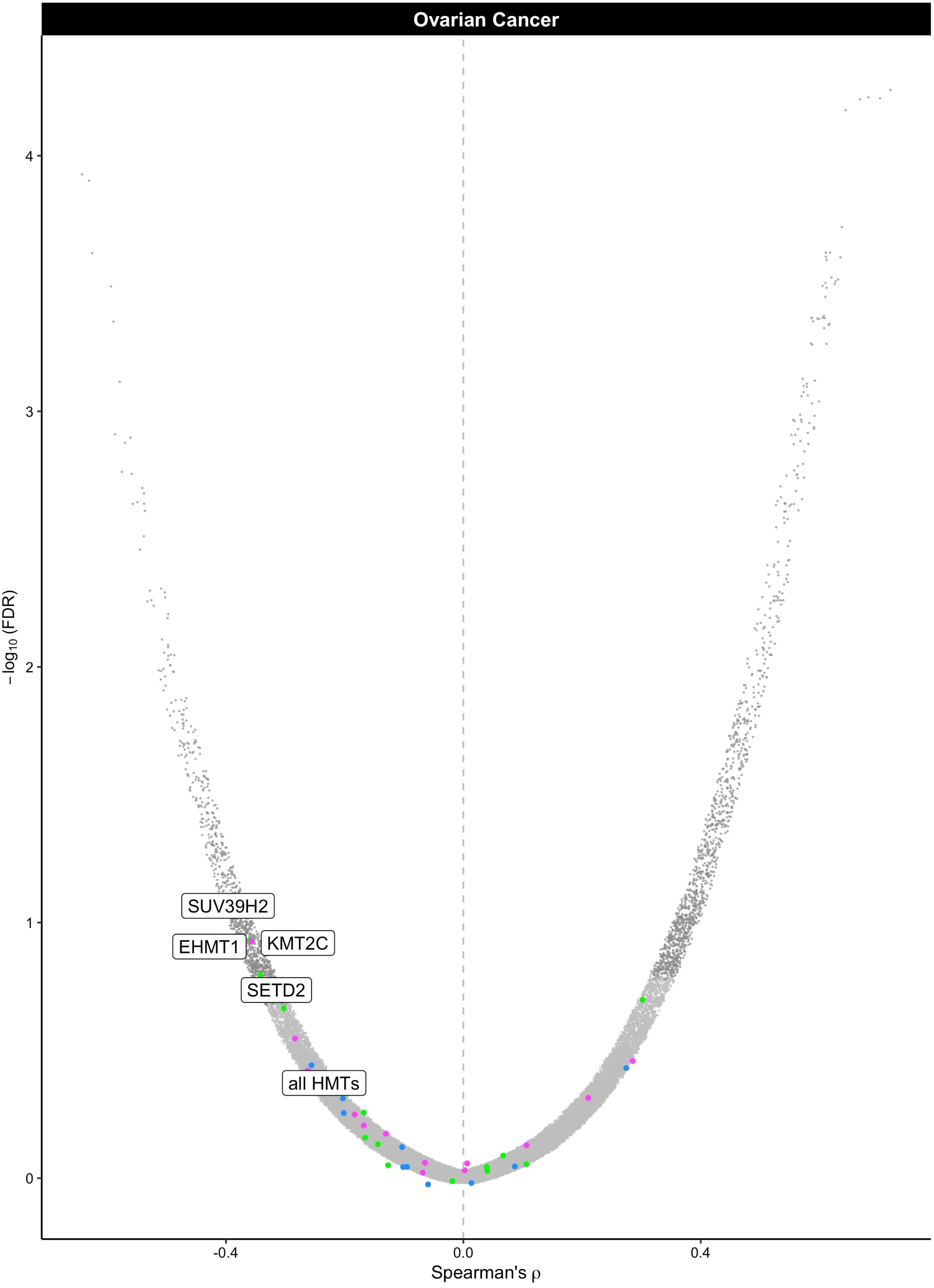

Ovarian Cancer
−log10 (FDR)
4
3
2
1
0
SUV39H2
EHMT1
KMT2C
SETD2
all HMTs
-0.4
0.0
0.4
Spearman's ρ

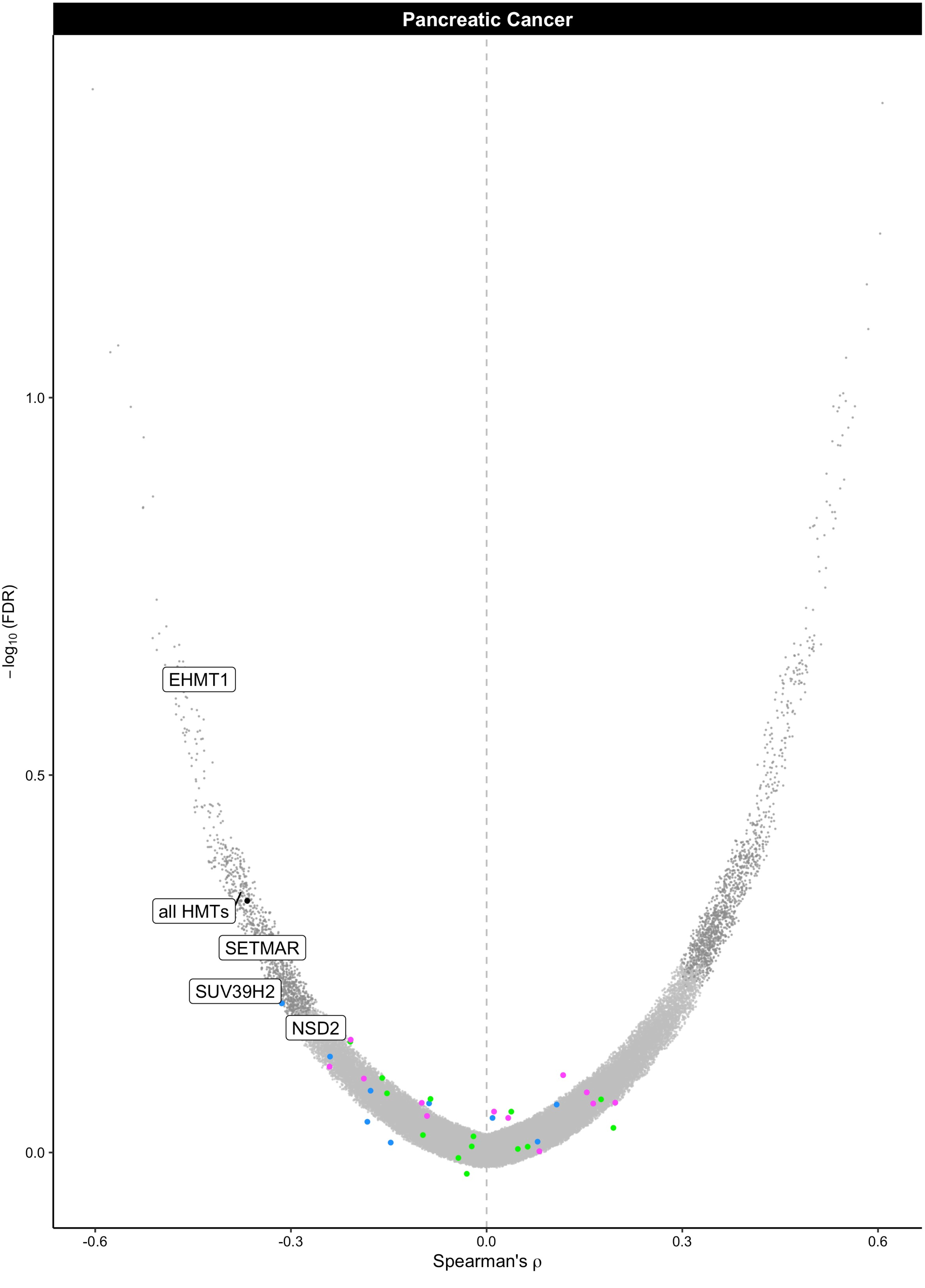
Pancreatic Cancer
−log10 (FDR)
1.0
0.5
0.0
EHMT1
all HMTs
SETMAR
SUV39H2
NSD2
-0.6
-0.3
0.0
0.3
0.6
Spearman's ρ

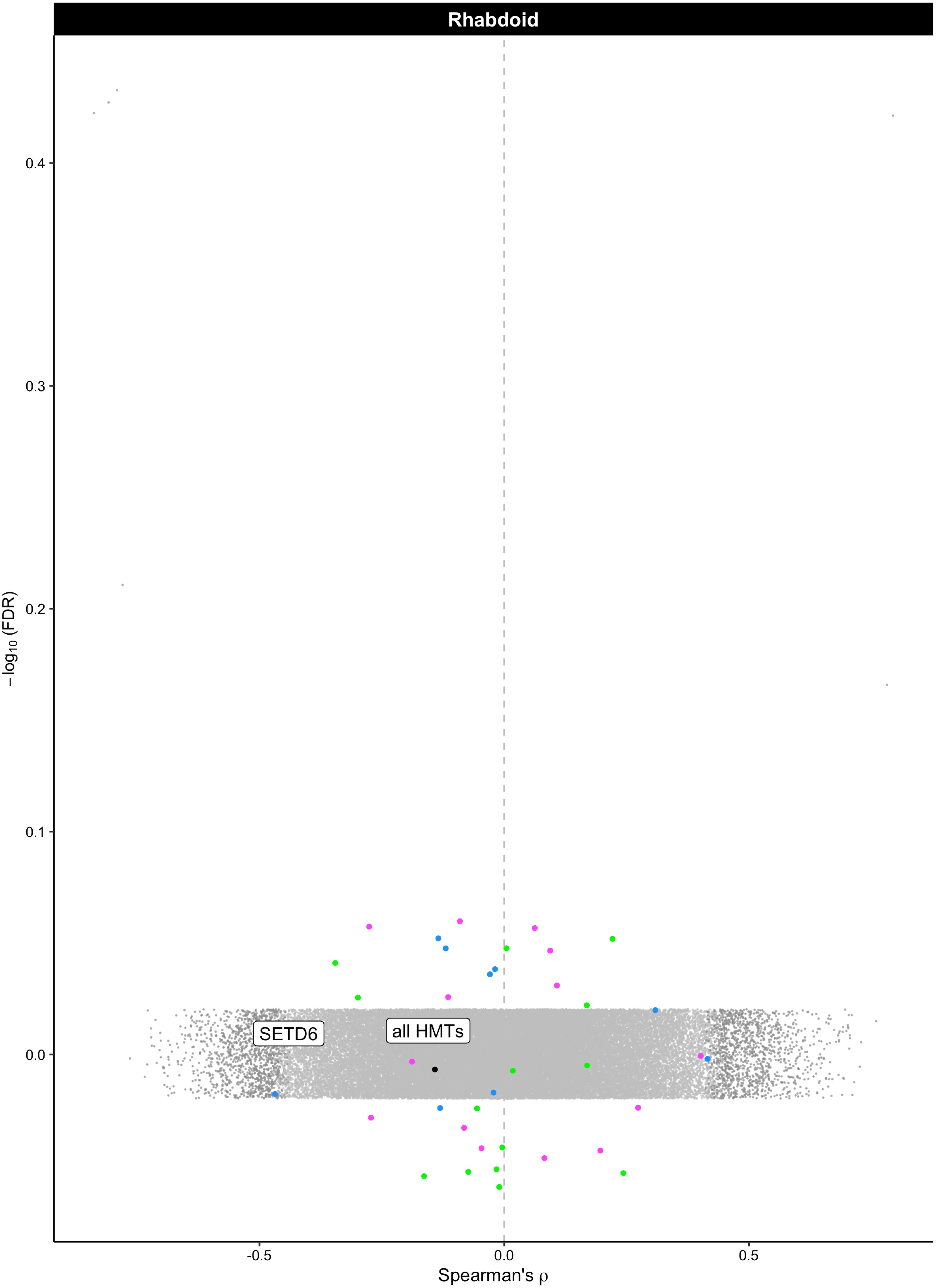
Rhabdoid
0.4
0.3
0.2
0.1
0.0
−log10 (FDR)
SETD6
all HMTs
-0.5
0.0
0.5
Spearman's ρ

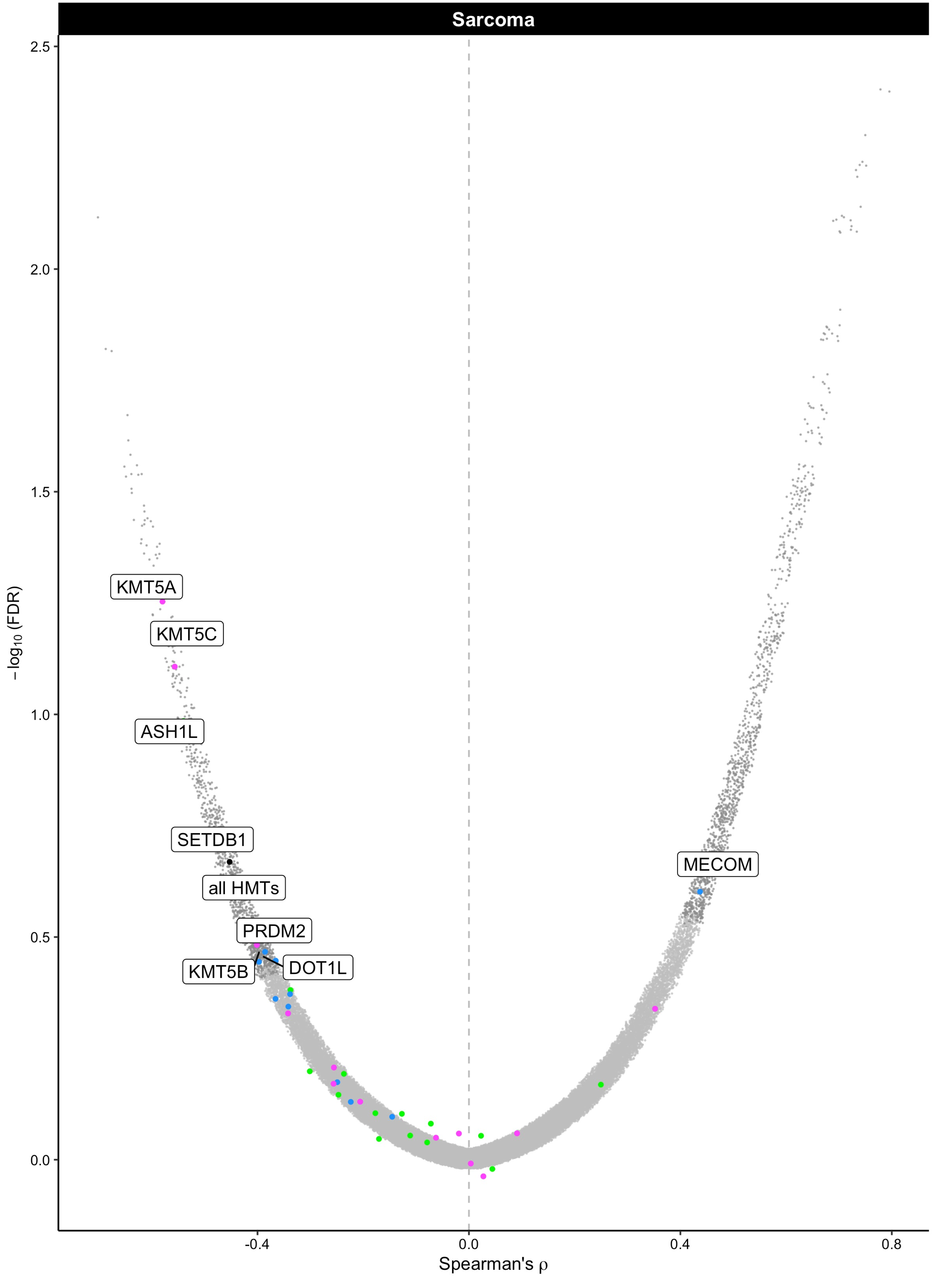

Sarcoma
2.5
2.0
1.5
1.0
0.5
0.0
−log10 (FDR)
KMT5A
KMT5C
ASH1L
SETDB1
all HMTs
PRDM2
KMT5B
DOT1L
MECOM
-0.4
0.0
0.4
0.8
Spearman's ρ

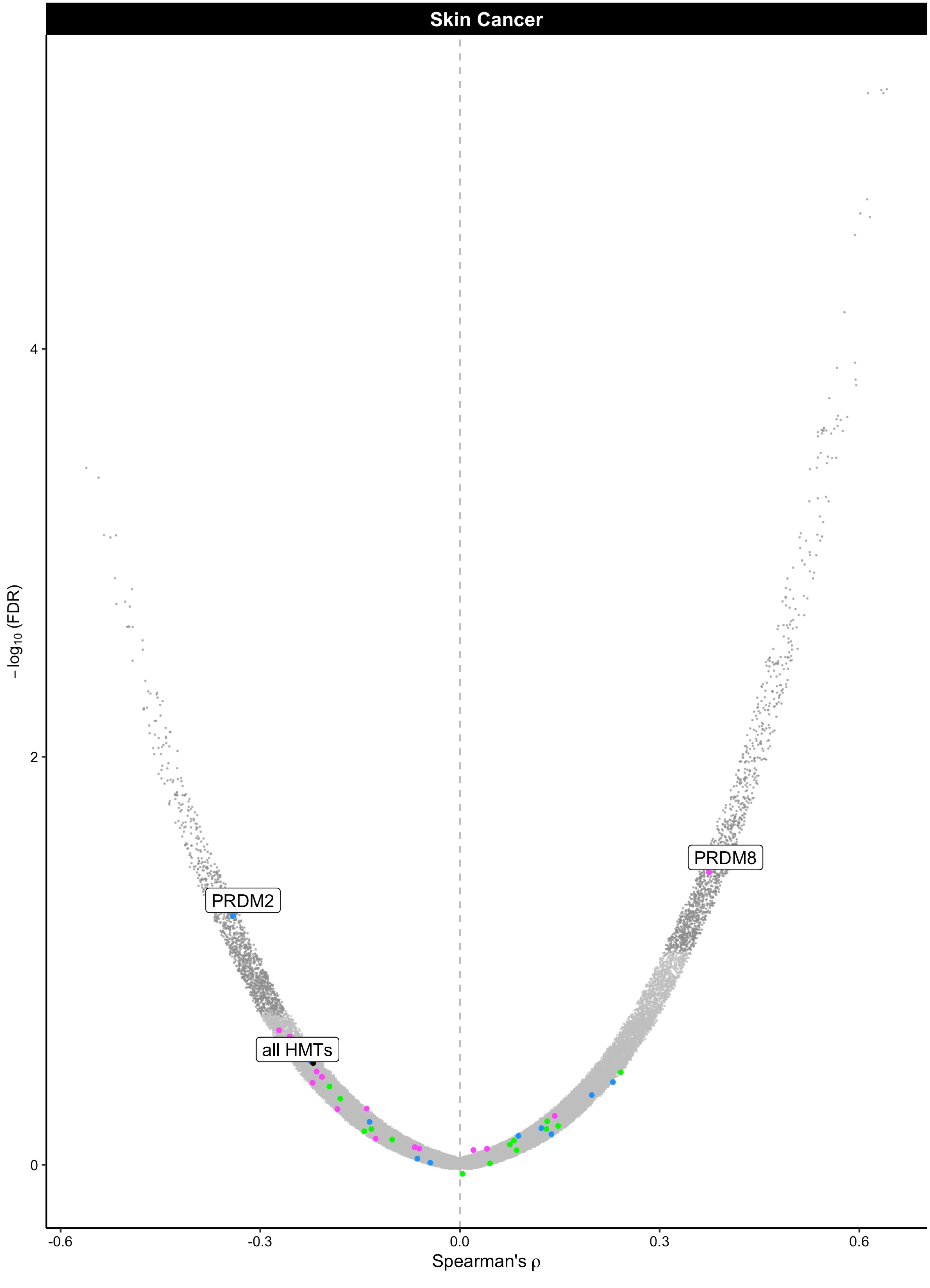

Skin Cancer
PRDM8
PRDM2
all HMTs
4
2
0
−log10 (FDR)
-0.6
-0.3
0.0
0.3
0.6
Spearman's ρ
