## Supplemental Figures and Tables for "Histone methylation has a direct metabolic role in human cells": Supp File 2. TCGA NNMT HMTs fullplots.pdf

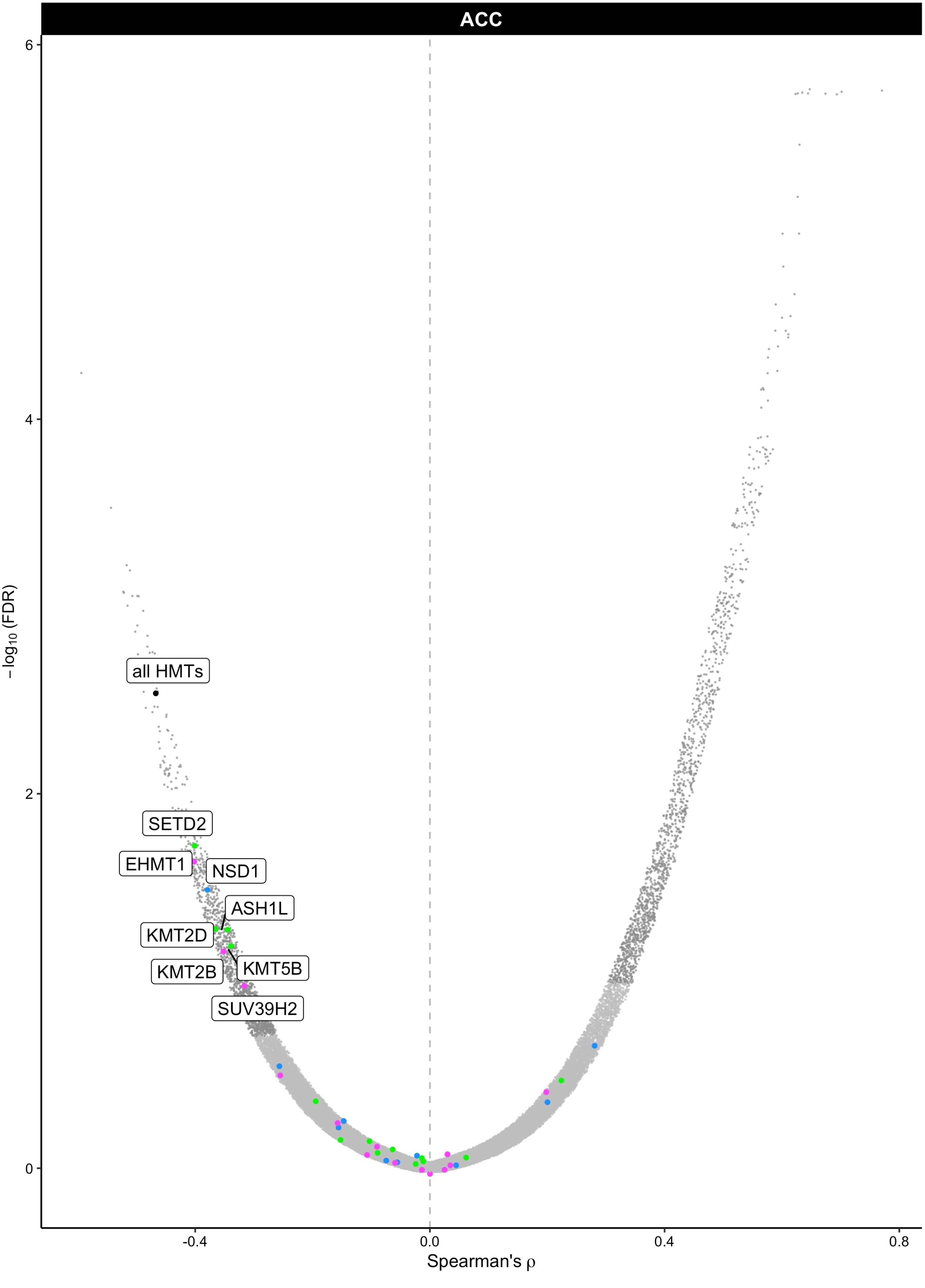

ACC
6
4
2
0
−log10 (FDR)
all HMTs
SETD2
EHMT1
NSD1
ASH1L
KMT2D
KMT2B
KMT5B
SUV39H2
-0.4
0.0
0.4
0.8
Spearman's ρ

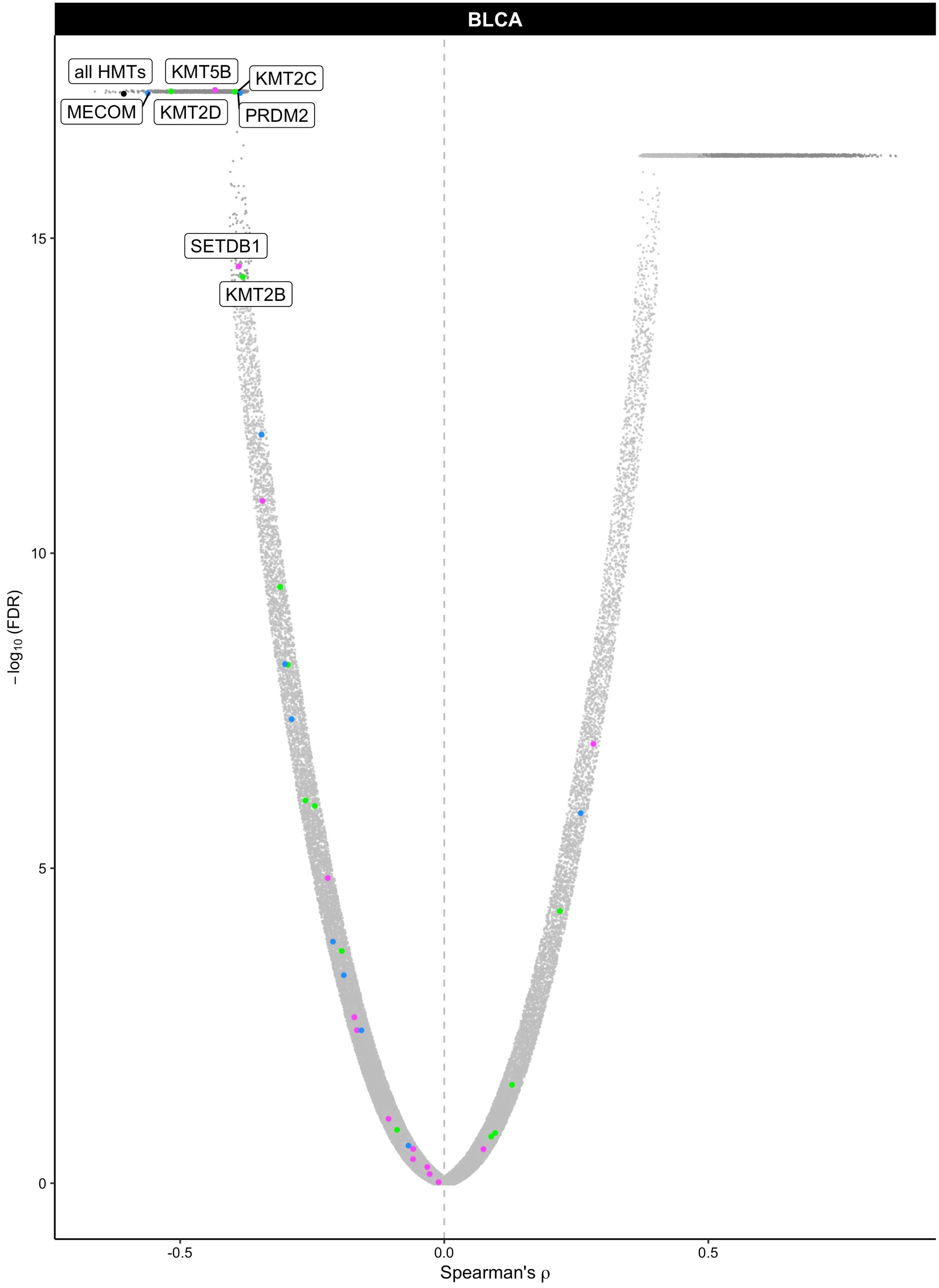

BLCA
all HMTs
KMT5B
KMT2C
MECOM
KMT2D
PRDM2
SETDB1
KMT2B
15
10
5
0
$-\log_{10}$ (FDR)
-0.5
0.0
0.5
Spearman's ρ

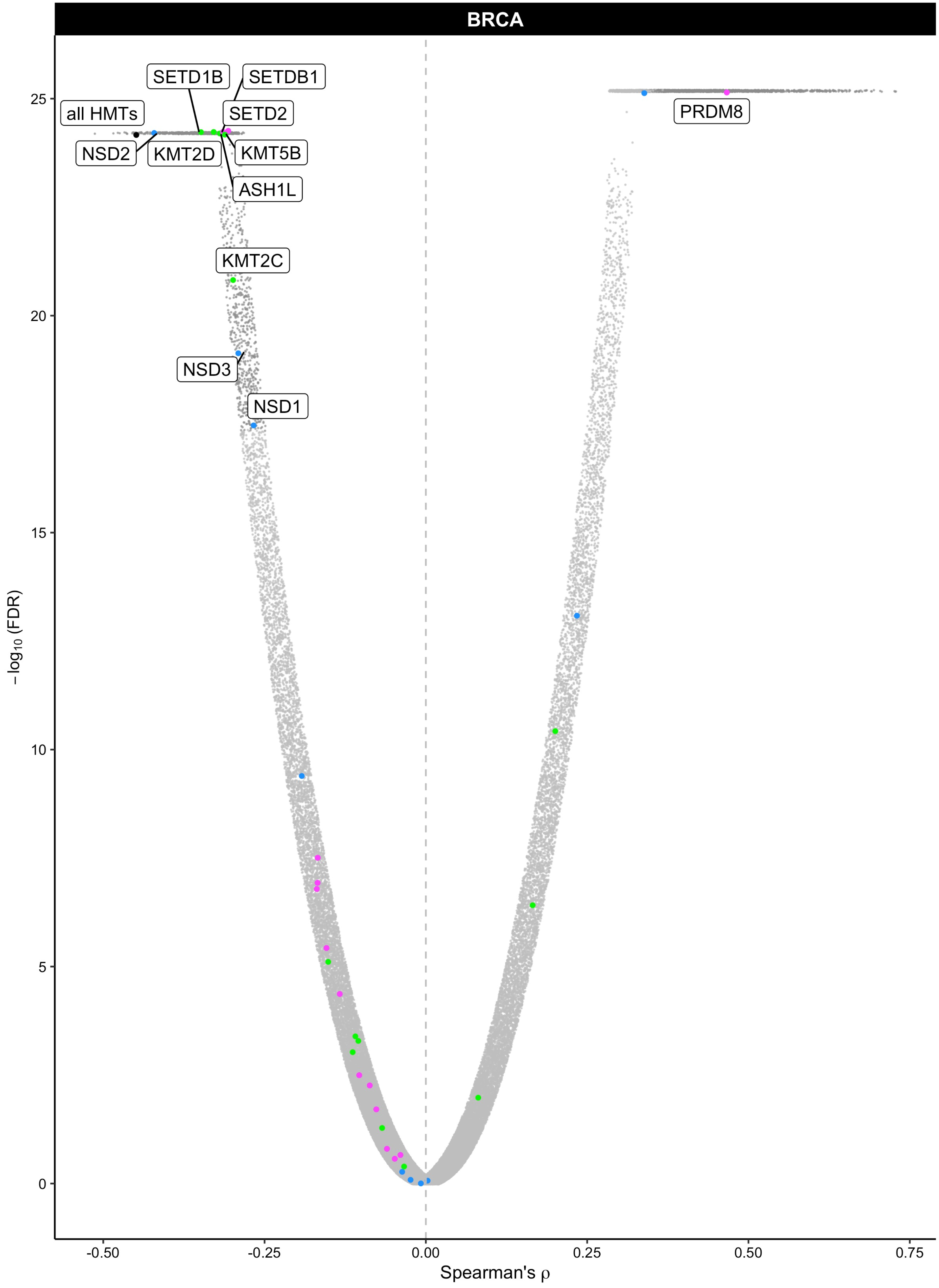

BRCA
SETD1B
SETDB1
SETD2
all HMTs
NSD2
KMT2D
KMT5B
ASH1L
KMT2C
NSD3
NSD1
PRDM8
−log10 (FDR)
Spearman's ρ
25
20
15
10
5
0
-0.50
-0.25
0.00
0.25
0.50
0.75

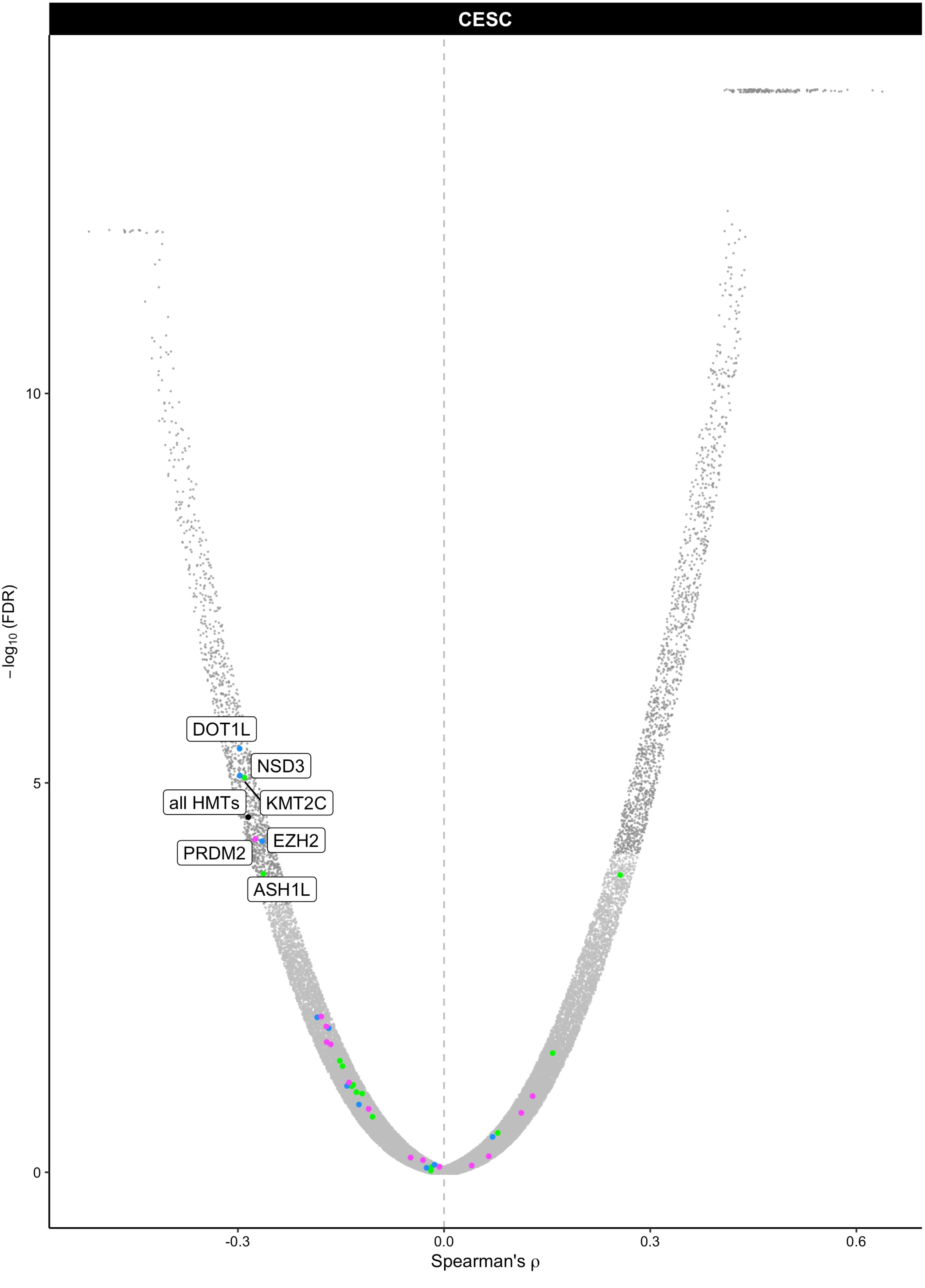

CESC
DOT1L
NSD3
all HMTs
KMT2C
EZH2
PRDM2
ASH1L
10
5
0
−log10 (FDR)
-0.3
0.0
0.3
0.6
Spearman's ρ

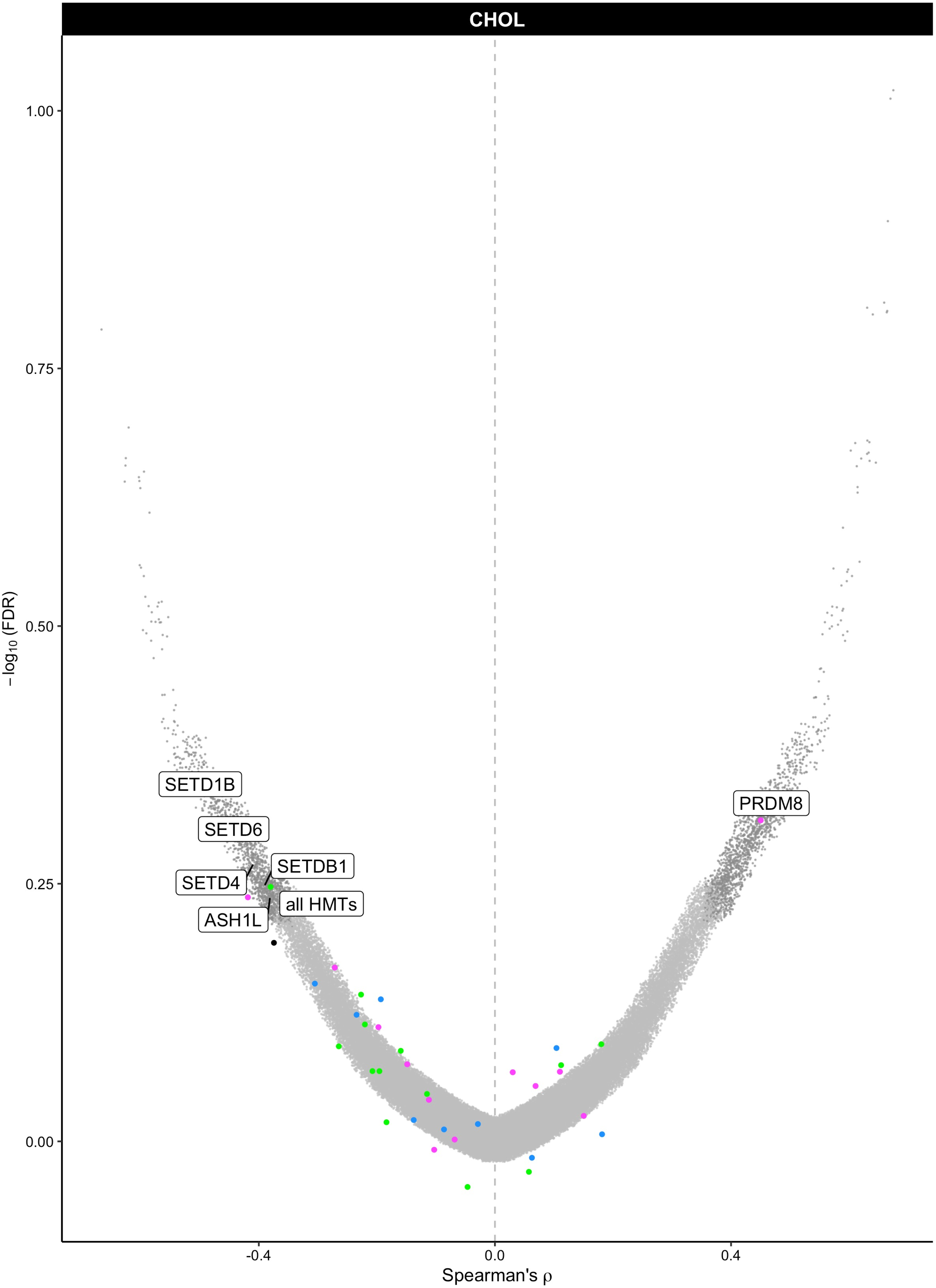
CHOL
1.00
0.75
0.50
0.25
0.00
−log10 (FDR)
SETD1B
SETD6
SETDB1
SETD4
all HMTs
ASH1L
PRDM8
-0.4
0.0
0.4
Spearman's ρ

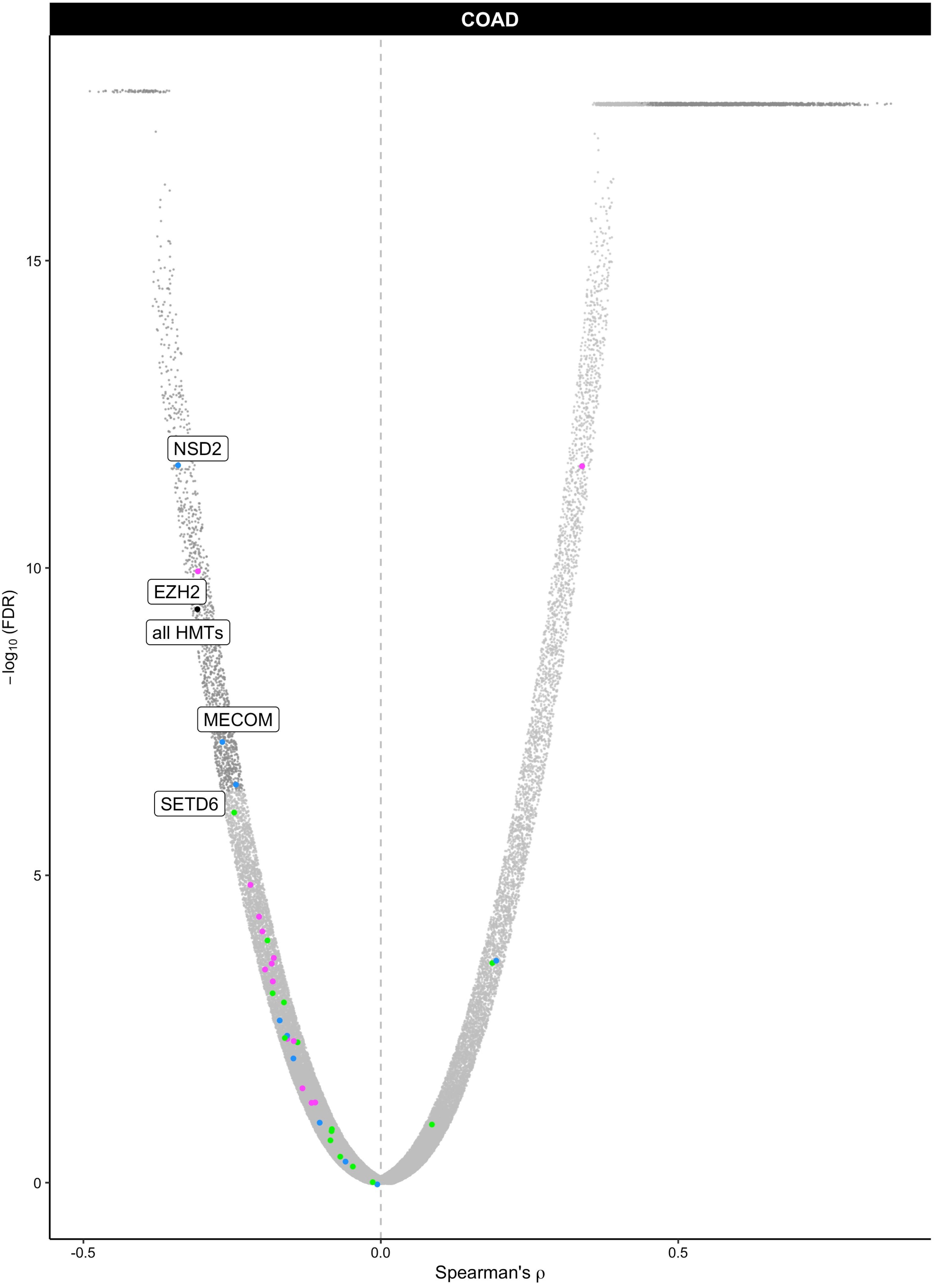
COAD
NSD2
EZH2
all HMTs
MECOM
SETD6
15
10
5
0
−log10 (FDR)
-0.5
0.0
0.5
Spearman's ρ

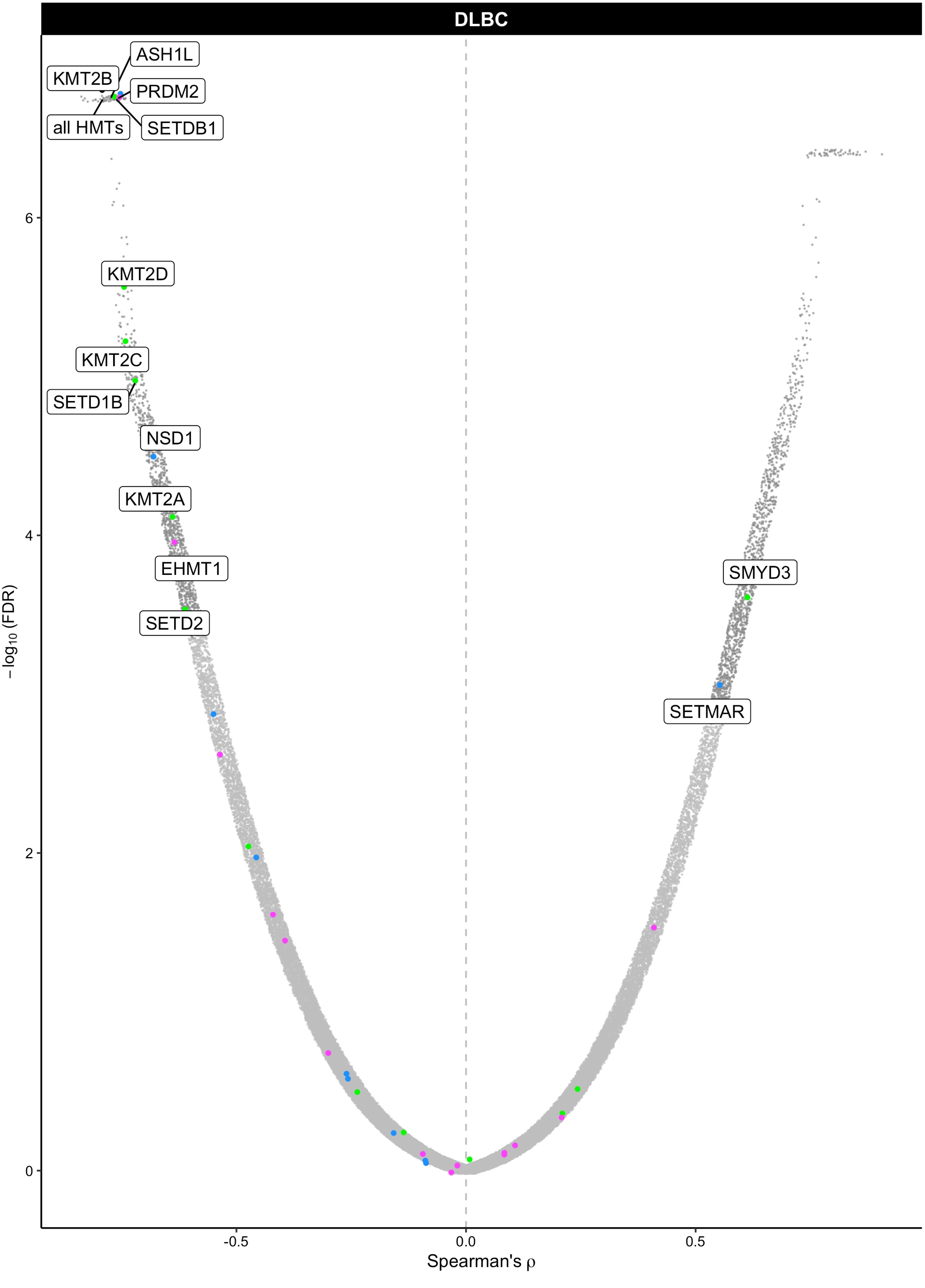

DLBC
ASH1L
KMT2B
PRDM2
all HMTs
SETDB1
KMT2D
KMT2C
SETD1B
NSD1
KMT2A
EHMT1
SETD2
SMYD3
SETMAR
6
4
2
0
−log10 (FDR)
-0.5
0.0
0.5
Spearman's ρ

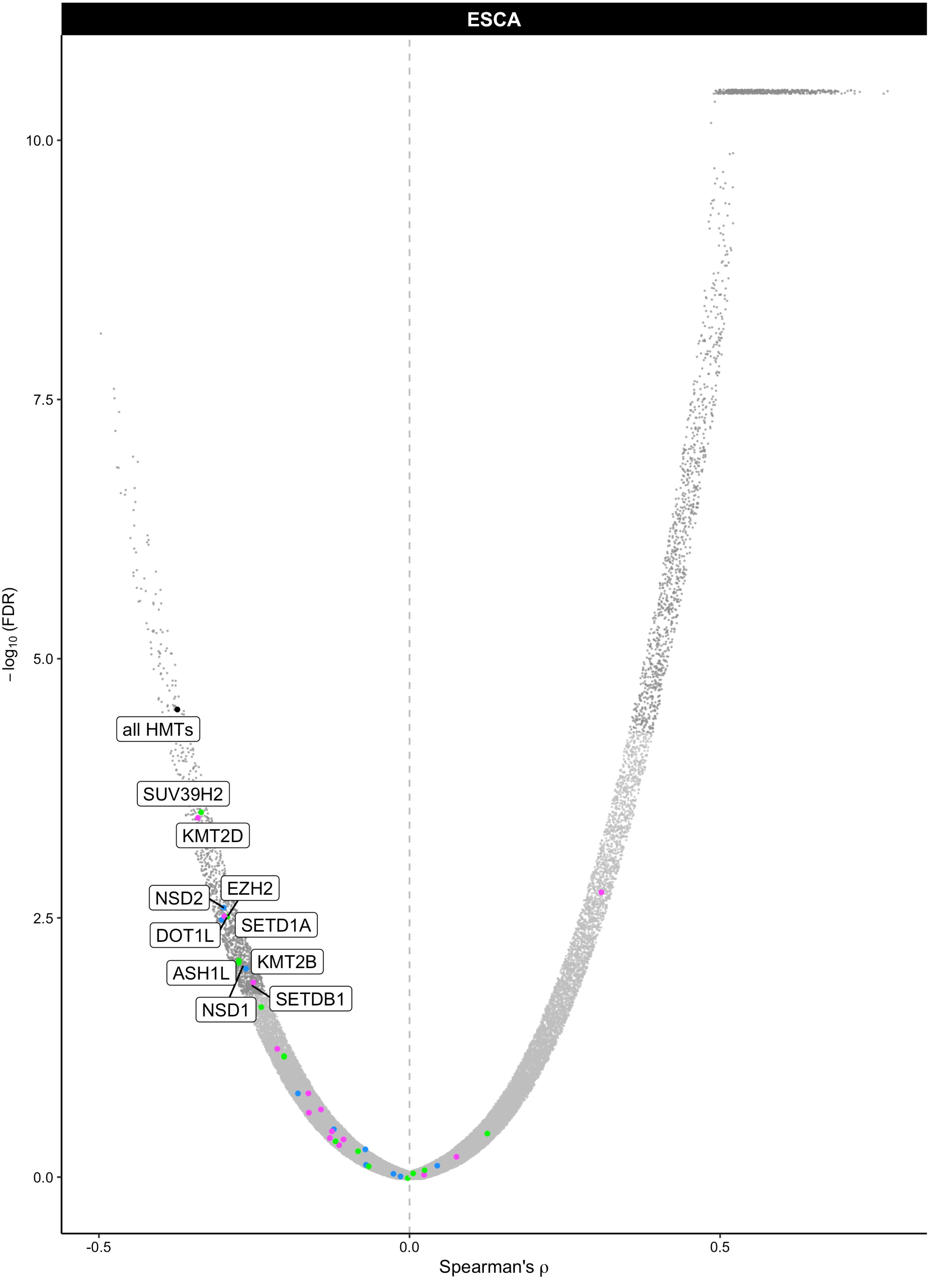

ESCA
all HMTs
SUV39H2
KMT2D
EZH2
NSD2
SETD1A
DOT1L
KMT2B
ASH1L
SETDB1
NSD1
10.0
7.5
5.0
2.5
0.0
-0.5
0.0
0.5
$-\log_{10}$(FDR)
Spearman's ρ

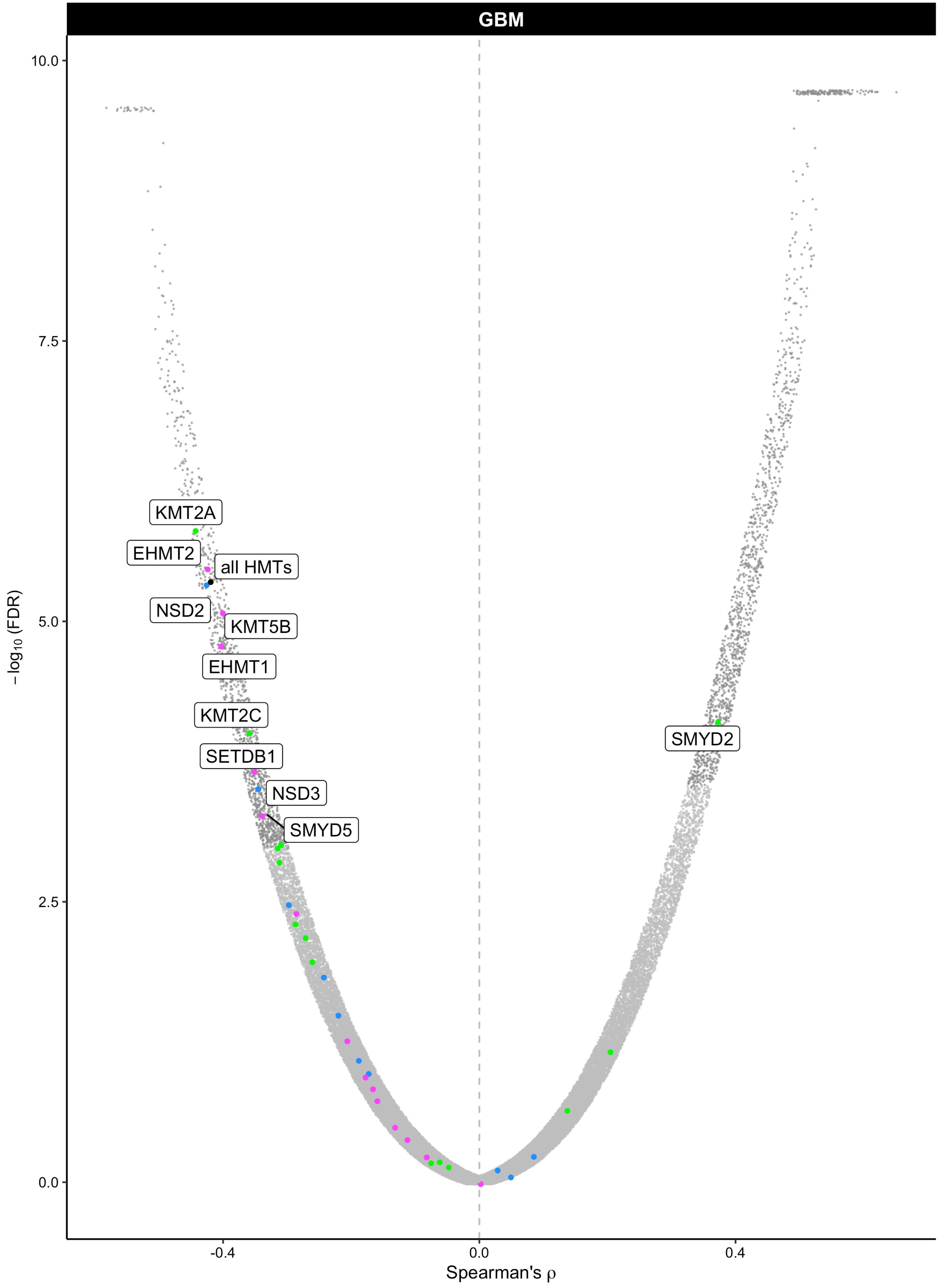

GBM
10.0
7.5
5.0
2.5
0.0
−log10 (FDR)
KMT2A
EHMT2
all HMTs
NSD2
KMT5B
EHMT1
KMT2C
SETDB1
NSD3
SMYD5
SMYD2
-0.4
0.0
0.4
Spearman's ρ

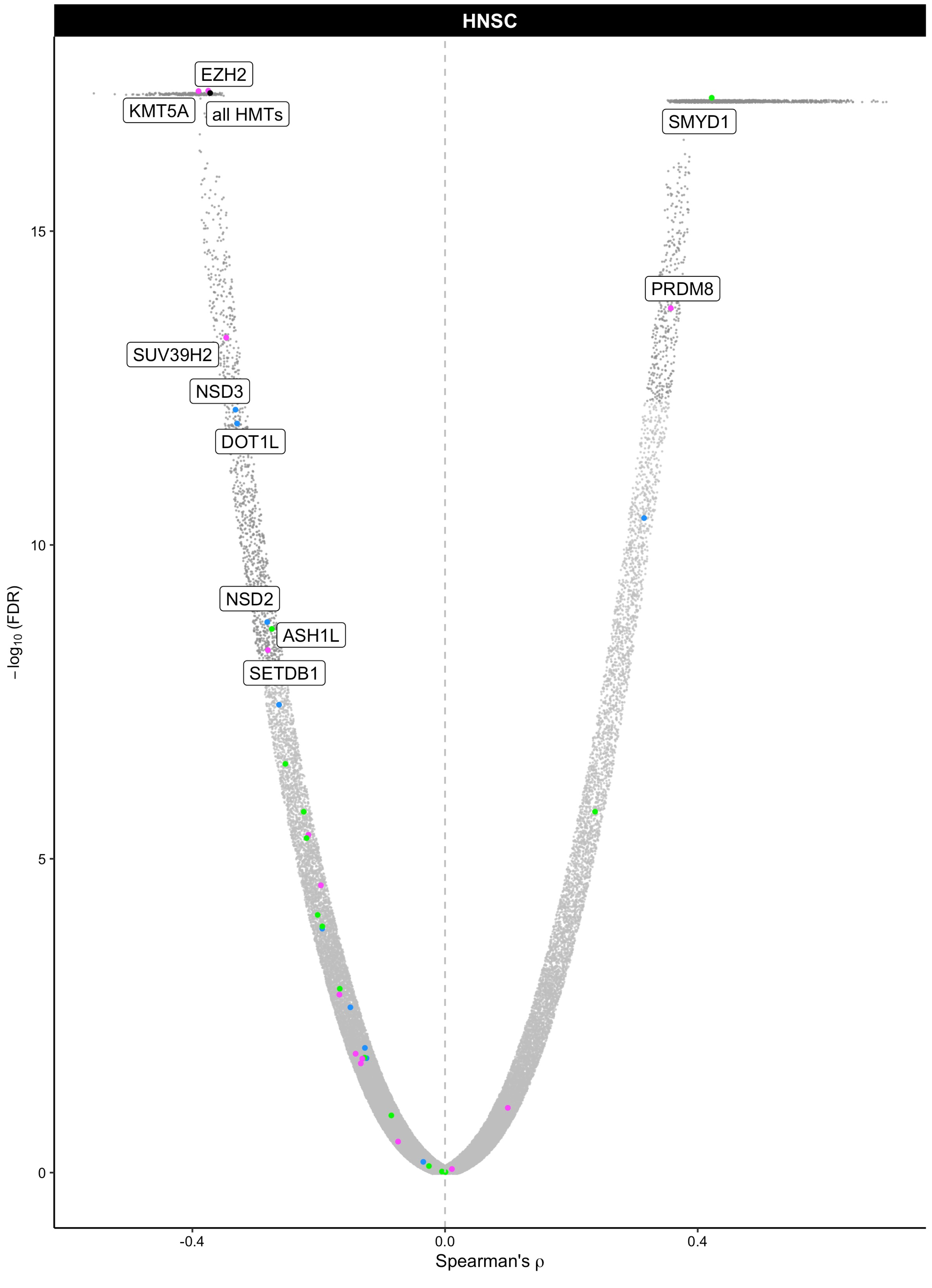

HNSC
EZH2
KMT5A
all HMTs
SMYD1
PRDM8
SUV39H2
NSD3
DOT1L
NSD2
ASH1L
SETDB1
15
10
5
0
-0.4
0.0
0.4
$-\log_{10}$ (FDR)
Spearman's ρ

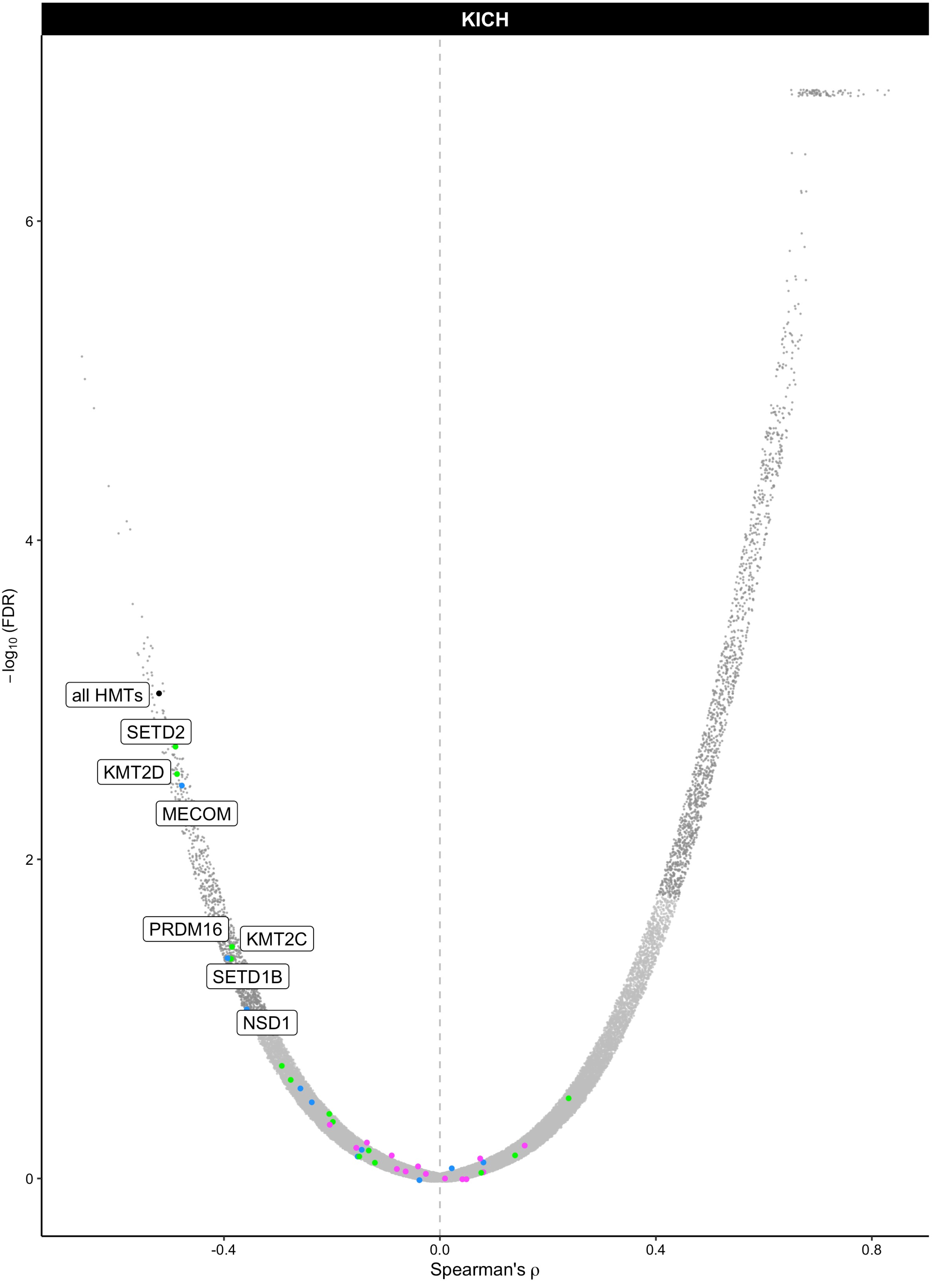

KICH
all HMTs
SETD2
KMT2D
MECOM
PRDM16
KMT2C
SETD1B
NSD1
6
4
2
0
-0.4
0.0
0.4
0.8
$-\log_{10}$(FDR)
Spearman's ρ

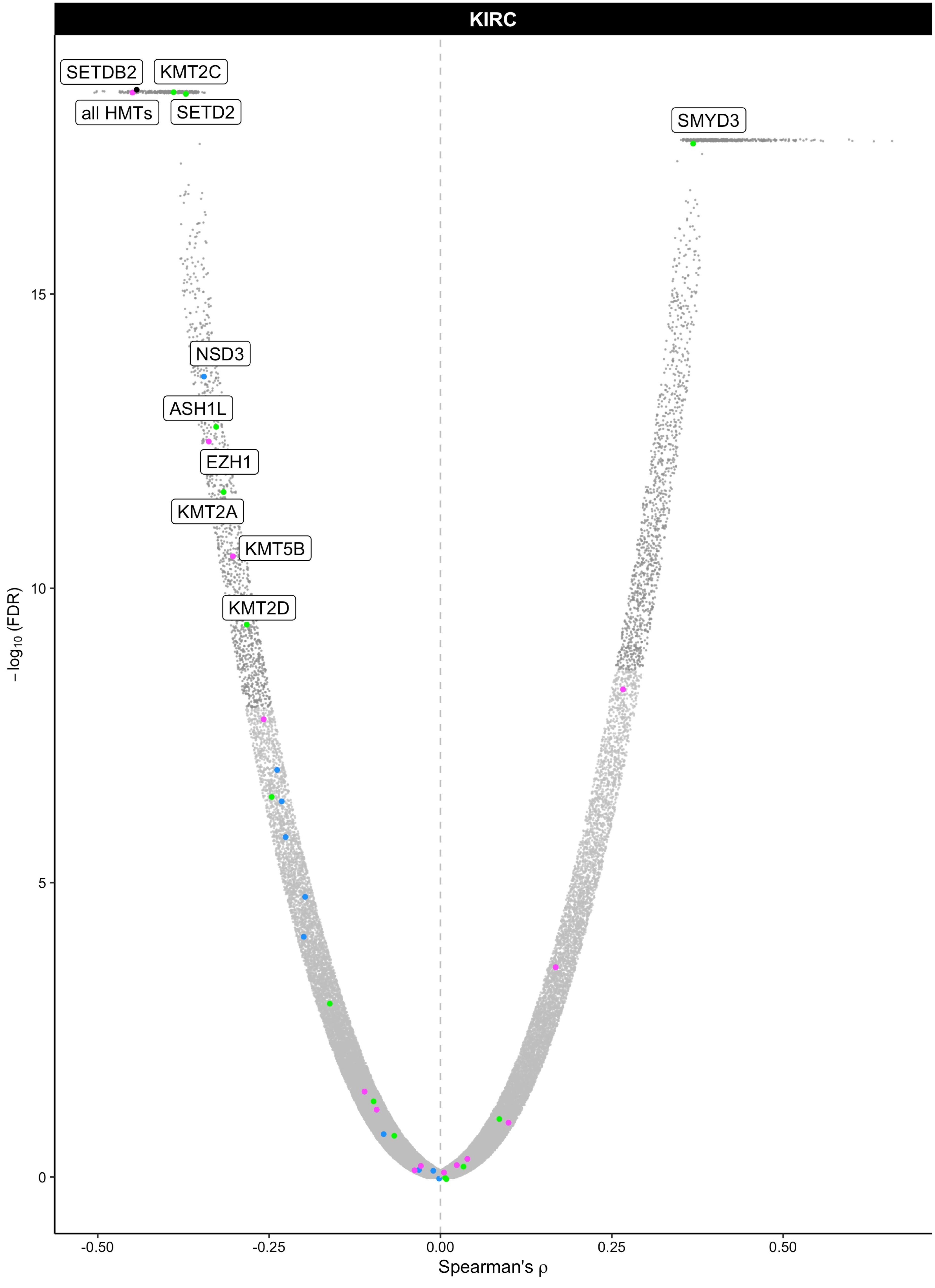

KIRC
SETDB2
KMT2C
all HMTs
SETD2
SMYD3
NSD3
ASH1L
EZH1
KMT2A
KMT5B
KMT2D
15
10
5
0
−log10 (FDR)
-0.50
-0.25
0.00
0.25
0.50
Spearman's ρ

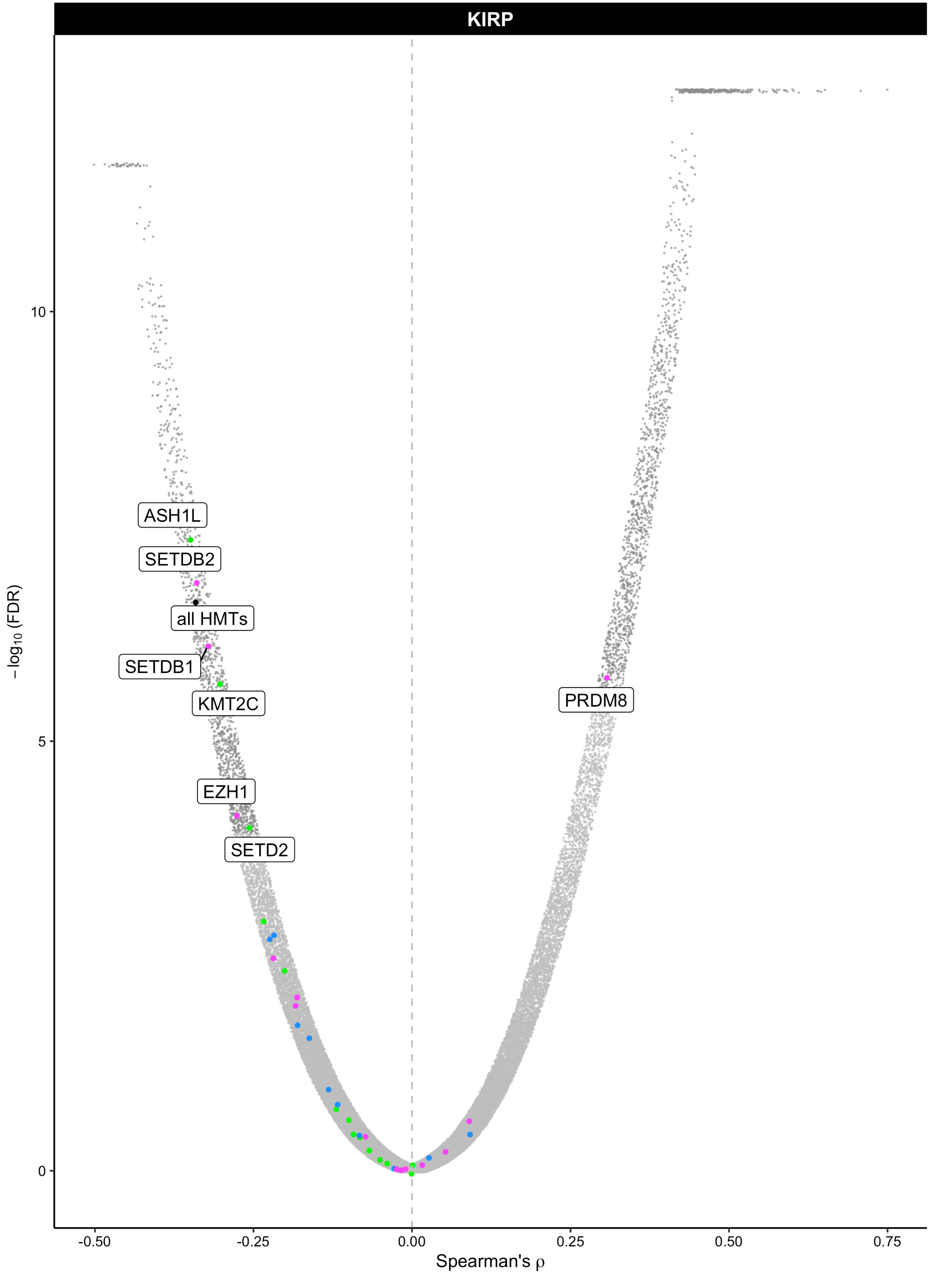

KIRP
ASH1L
SETDB2
all HMTs
SETDB1
KMT2C
EZH1
SETD2
PRDM8
10
5
0
−log10 (FDR)
-0.50
-0.25
0.00
0.25
0.50
0.75
Spearman's ρ

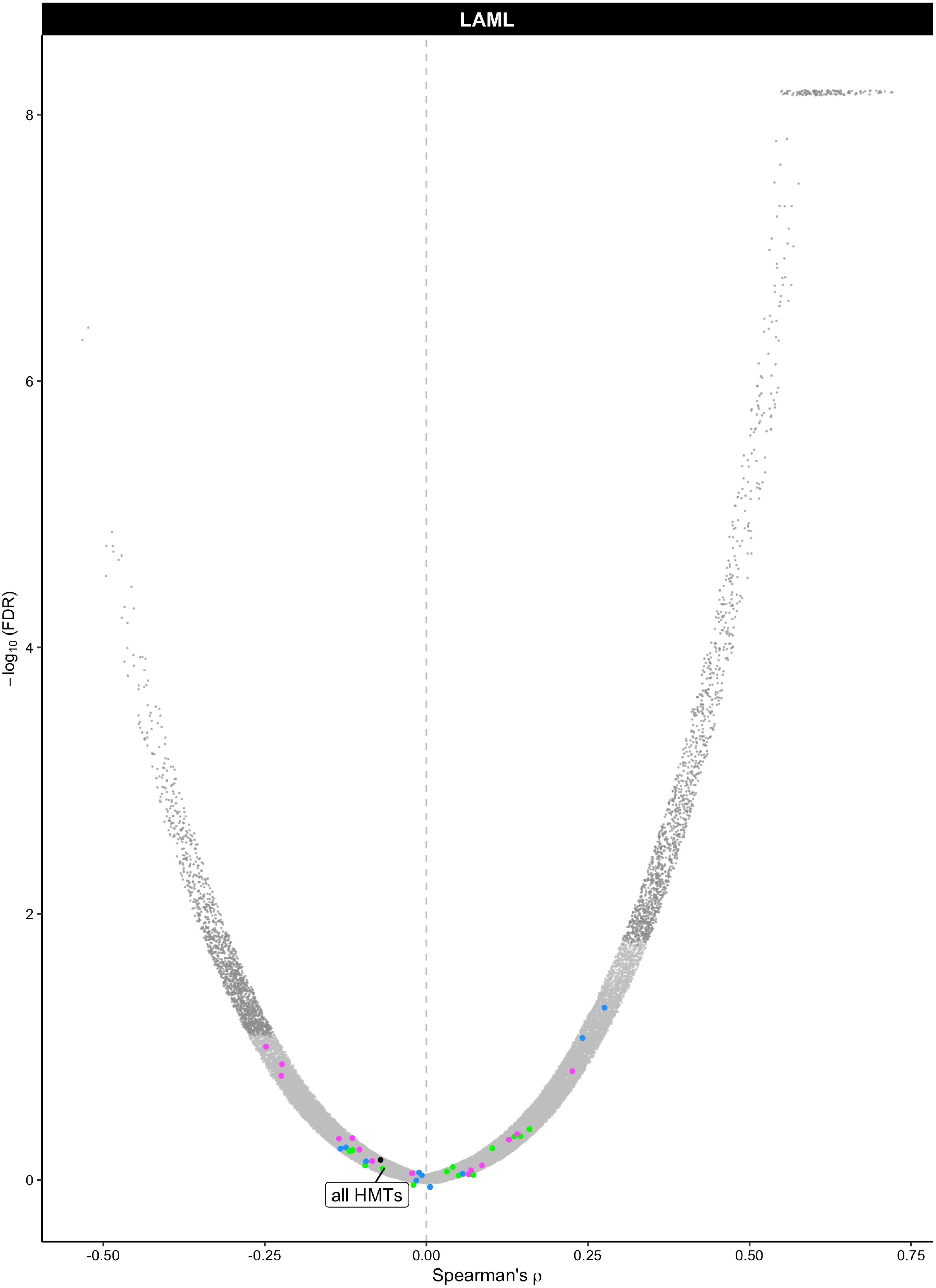
LAML
8
6
4
2
0
-0.50
-0.25
0.00
0.25
0.50
0.75
−log10 (FDR)
Spearman's ρ
all HMTs

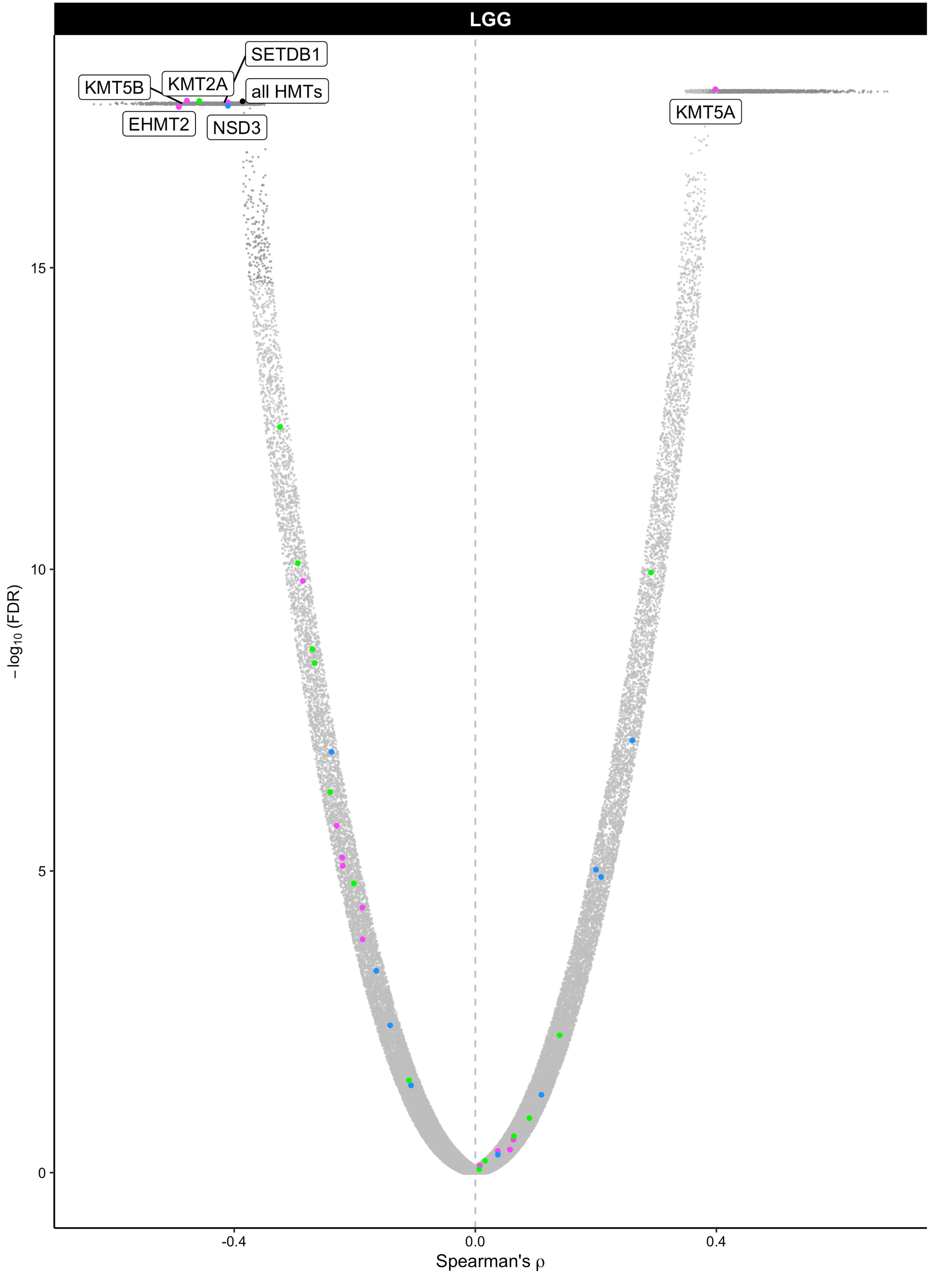

LGG
SETDB1
KMT5B
KMT2A
all HMTs
EHMT2
NSD3
KMT5A
15
10
5
0
−log10 (FDR)
-0.4
0.0
0.4
Spearman's ρ

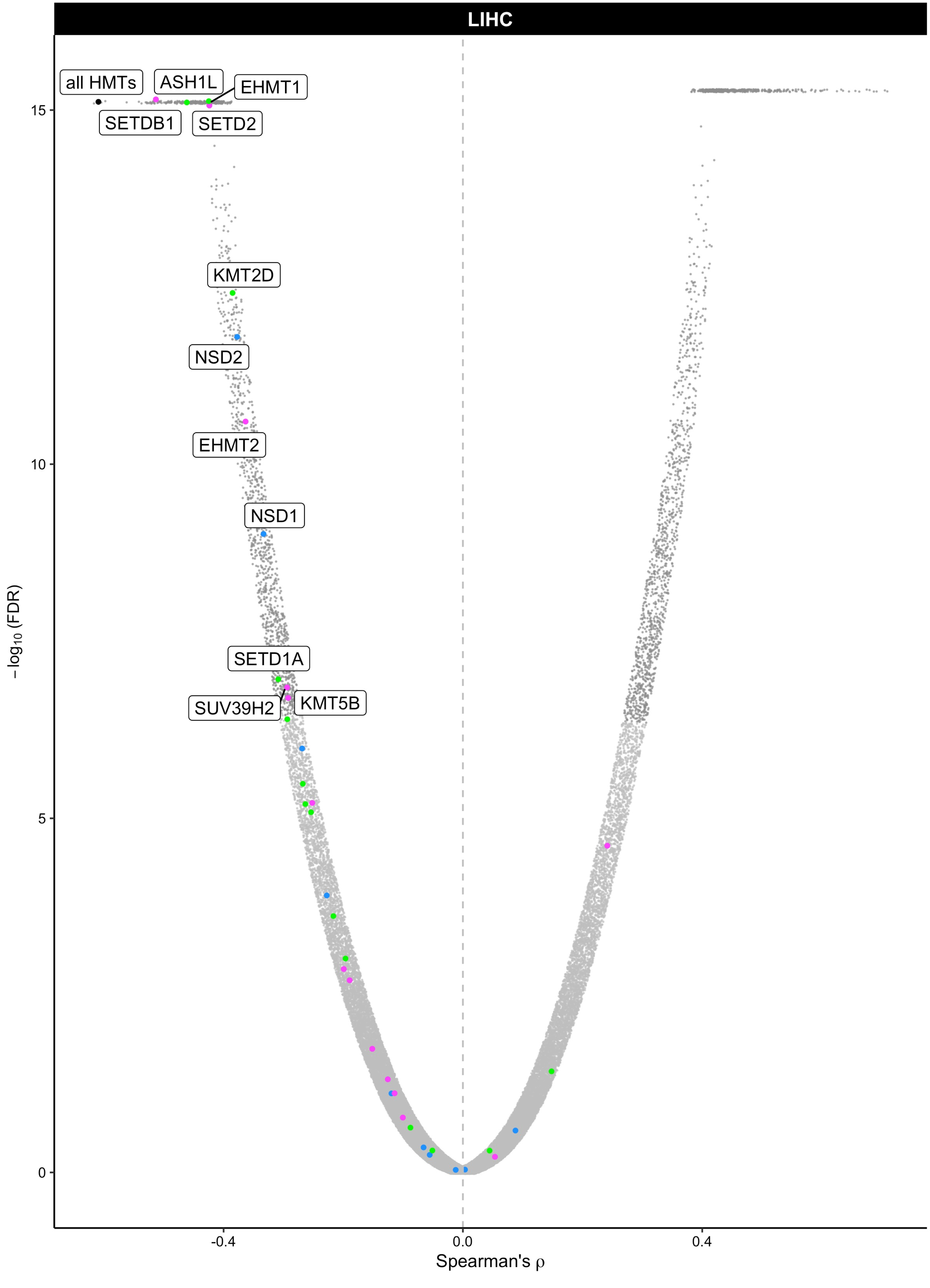

LIHC
all HMTs
ASH1L
EHMT1
SETDB1
SETD2
KMT2D
NSD2
EHMT2
NSD1
SETD1A
SUV39H2
KMT5B
15
10
5
0
-0.4
0.0
0.4
Spearman's ρ
−log10 (FDR)

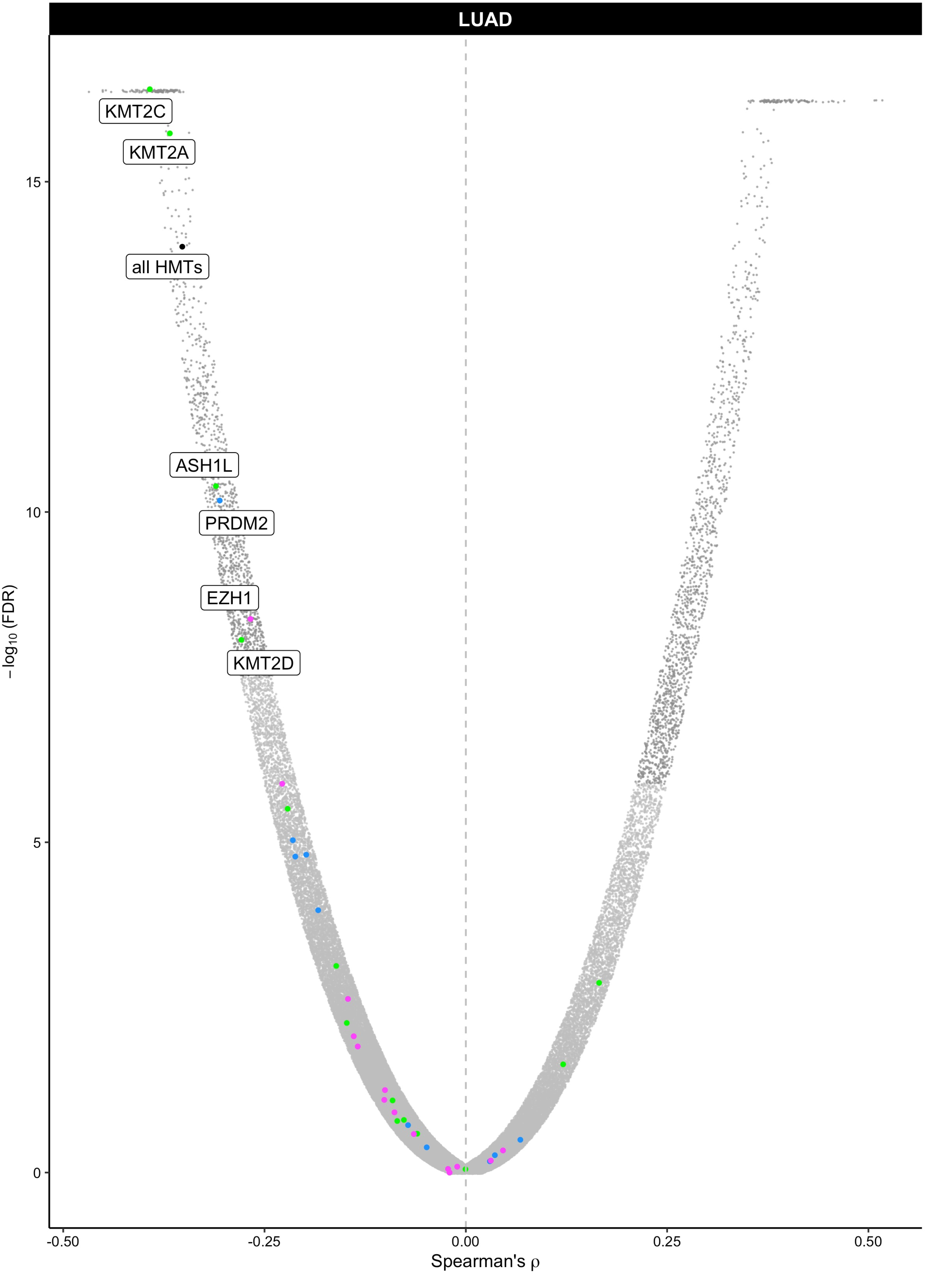
LUAD
KMT2C
KMT2A
all HMTs
ASH1L
PRDM2
EZH1
KMT2D
15
10
5
0
-log10 (FDR)
-0.50
-0.25
0.00
0.25
0.50
Spearman's ρ

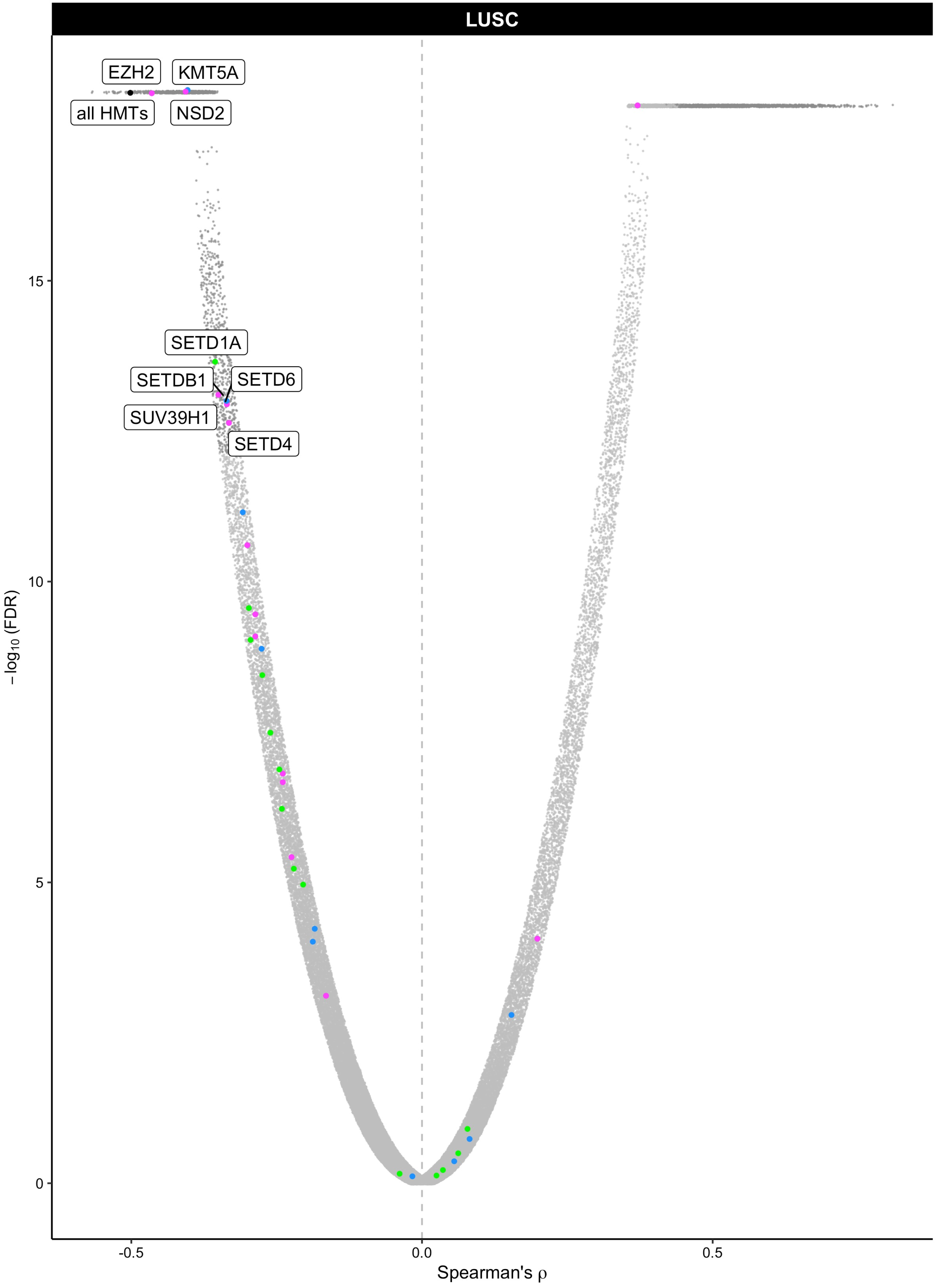

LUSC
EZH2
KMT5A
all HMTs
NSD2
SETD1A
SETDB1
SETD6
SUV39H1
SETD4
15
10
5
0
−log10 (FDR)
-0.5
0.0
0.5
Spearman's ρ

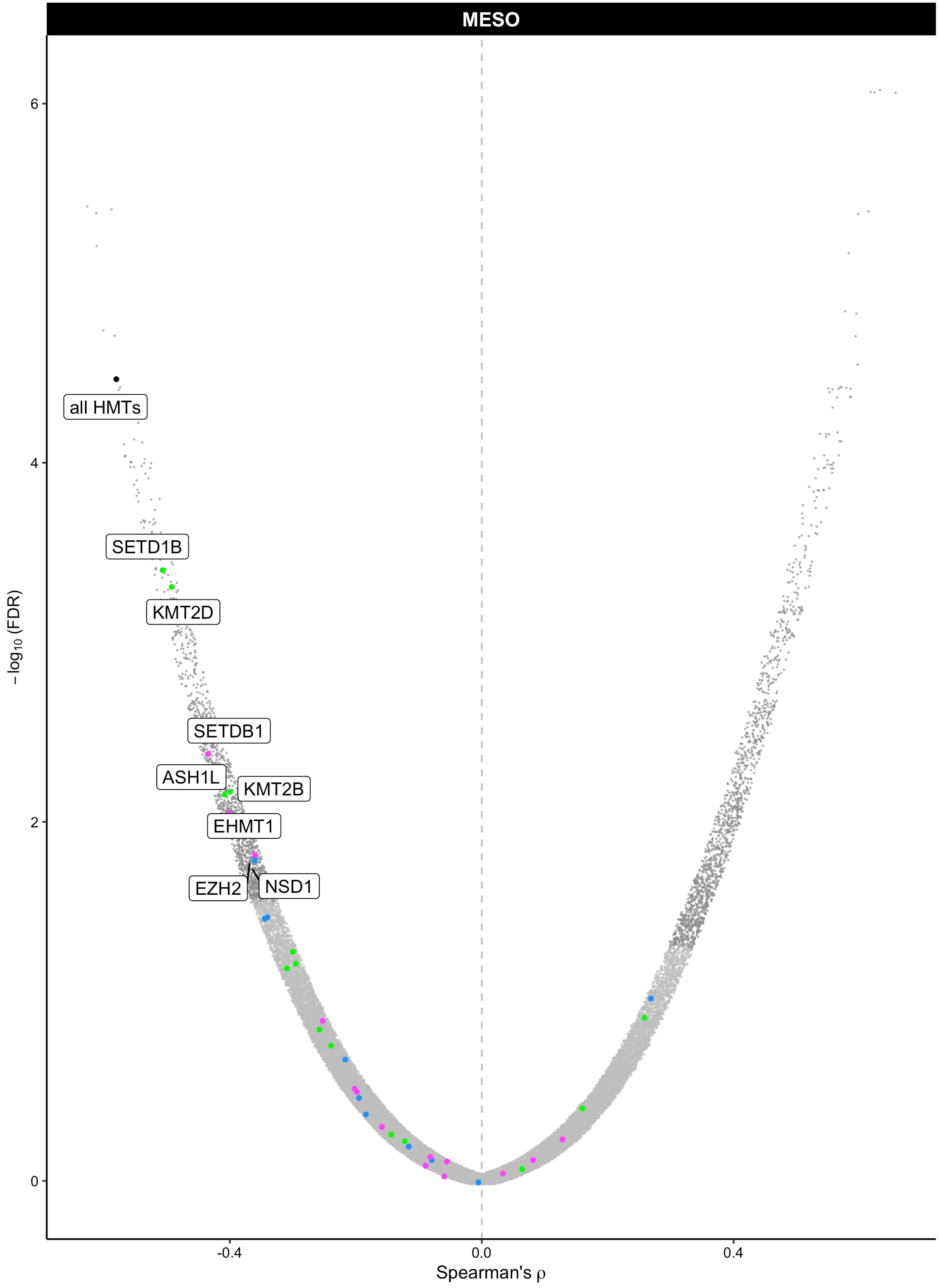
MESO
all HMTs
SETD1B
KMT2D
SETDB1
ASH1L
KMT2B
EHMT1
EZH2
NSD1
6
4
2
0
−log10 (FDR)
-0.4
0.0
0.4
Spearman's ρ

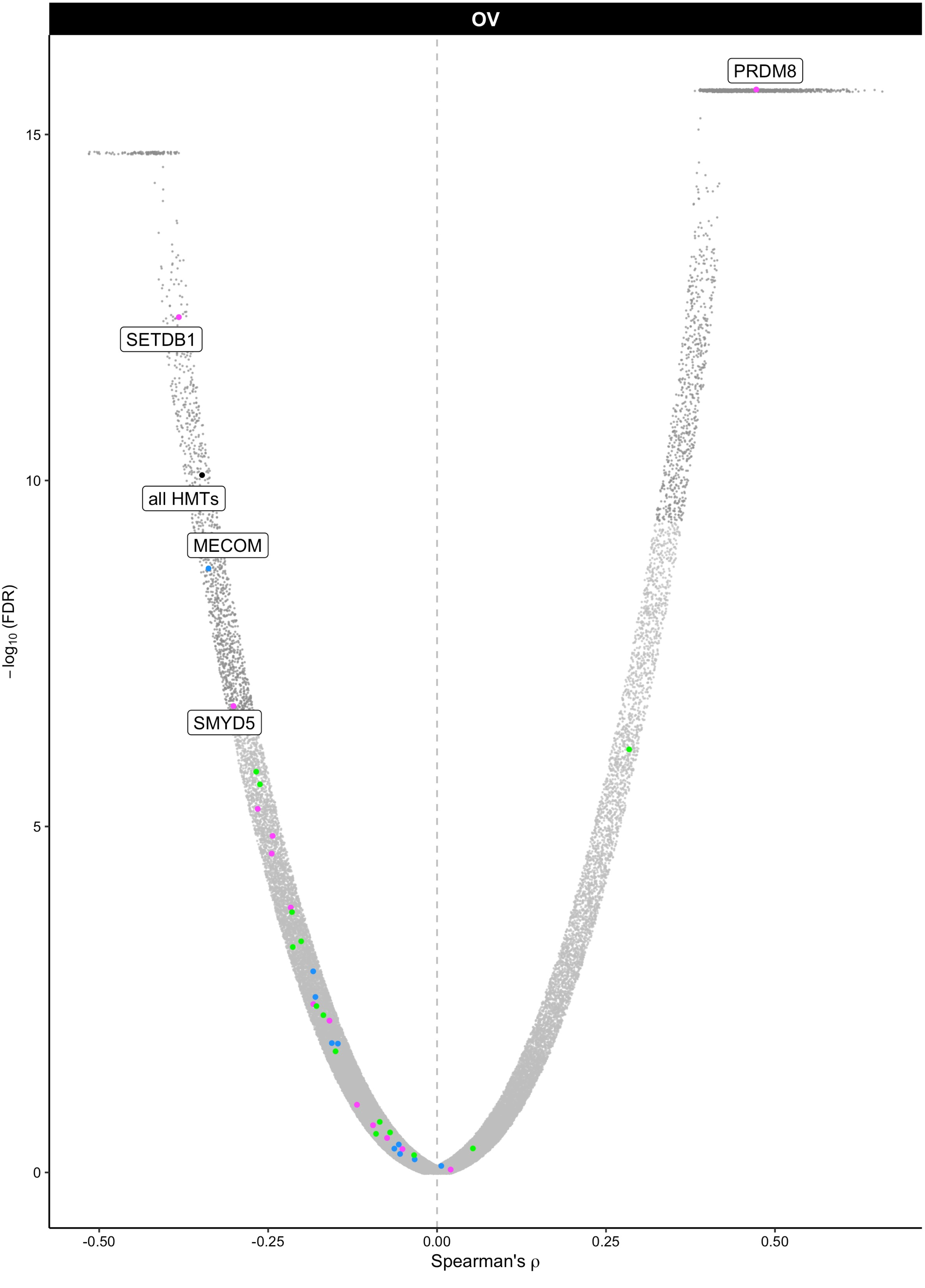

OV
PRDM8
SETDB1
all HMTs
MECOM
SMYD5
15
10
5
0
−log10 (FDR)
-0.50
-0.25
0.00
0.25
0.50
Spearman's ρ

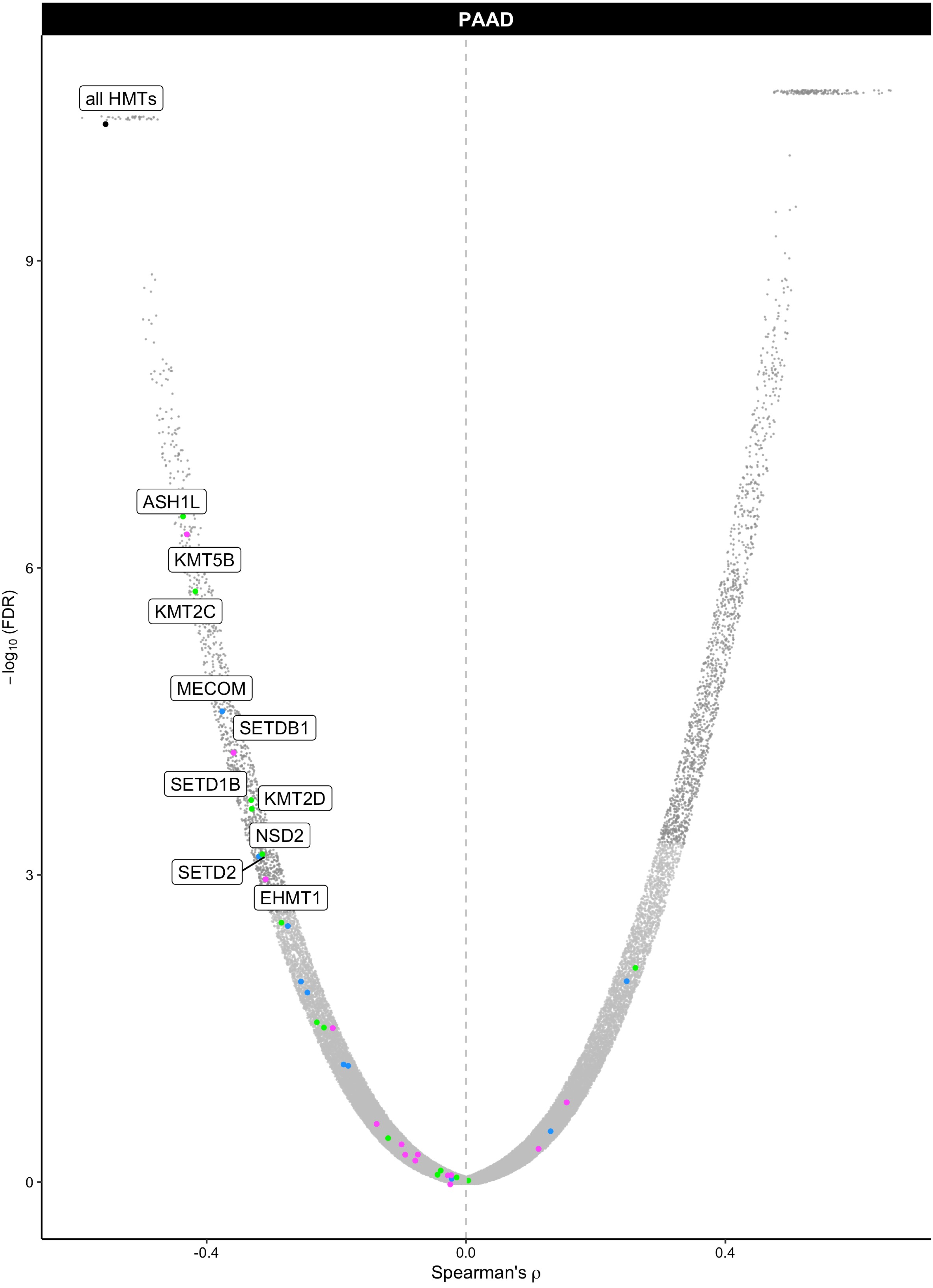

PAAD
all HMTs
ASH1L
KMT5B
KMT2C
MECOM
SETDB1
SETD1B
KMT2D
NSD2
SETD2
EHMT1
9
6
3
0
$-\log_{10}$ (FDR)
-0.4
0.0
0.4
Spearman's ρ

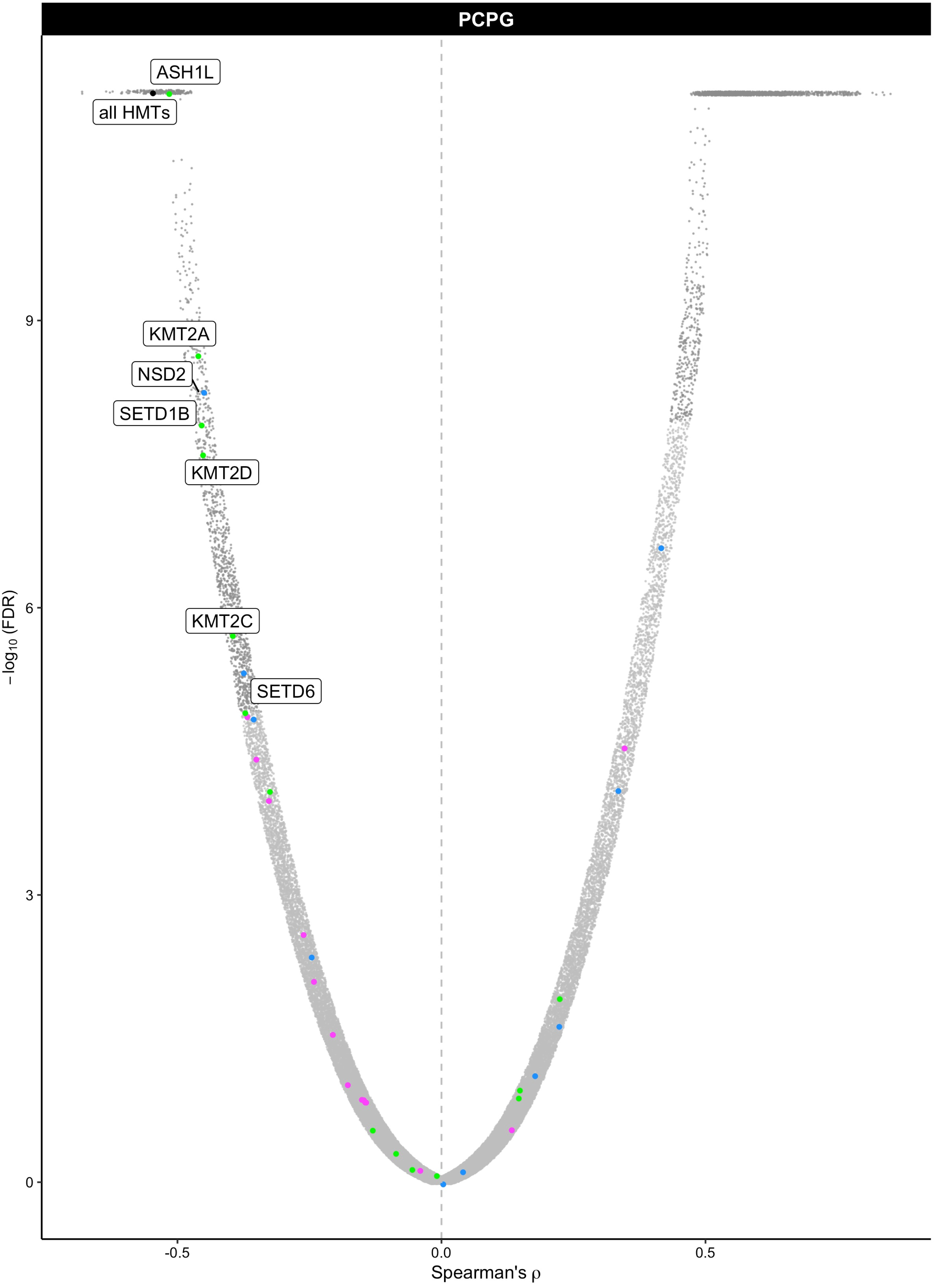

PCPG
ASH1L
all HMTs
KMT2A
NSD2
SETD1B
KMT2D
KMT2C
SETD6
9
6
3
0
$-\log_{10}$ (FDR)
-0.5
0.0
0.5
Spearman's ρ

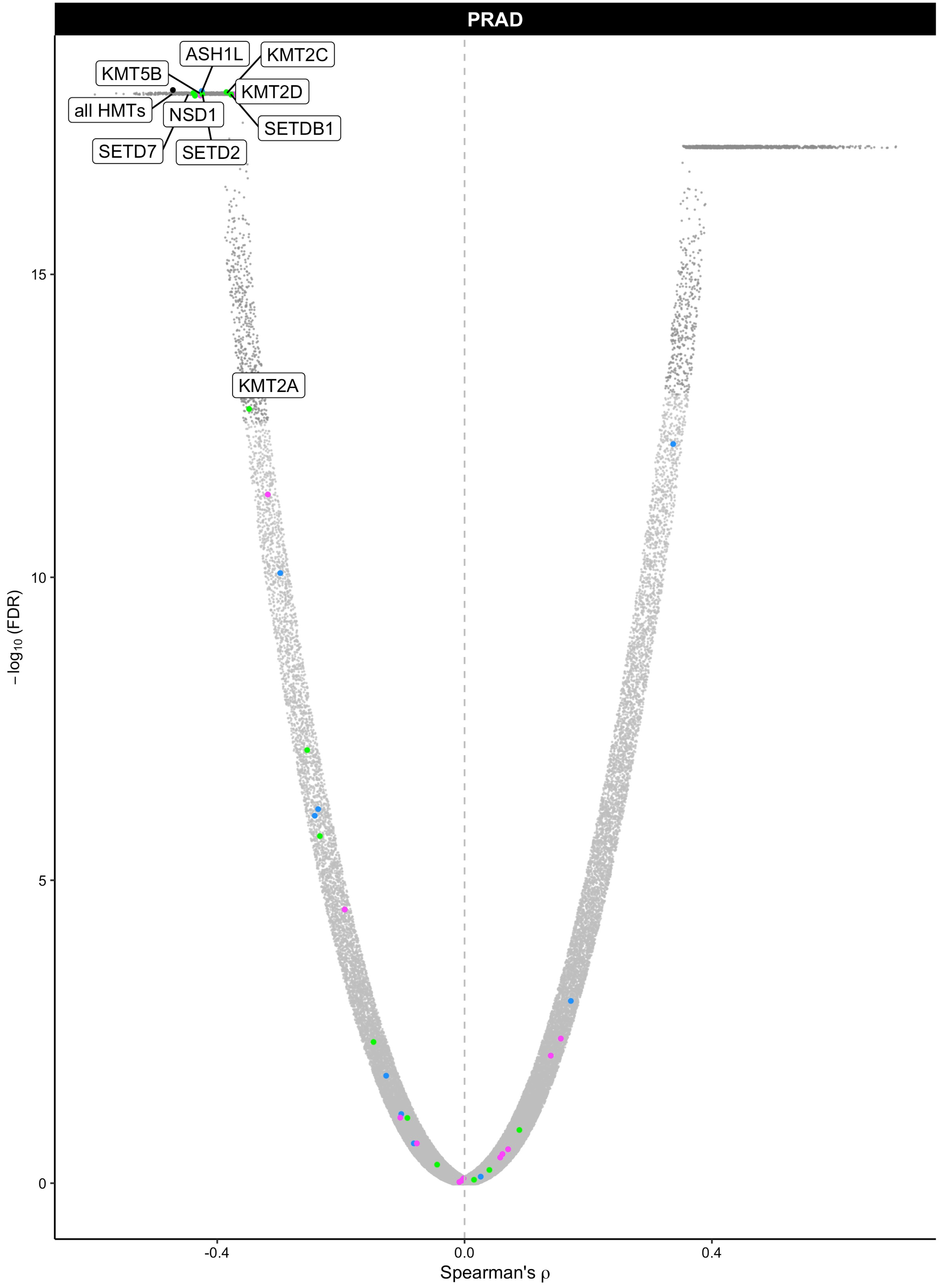

PRAD
ASH1L
KMT2C
KMT5B
KMT2D
all HMTs
NSD1
SETDB1
SETD7
SETD2
KMT2A
15
10
5
0
−log10 (FDR)
-0.4
0.0
0.4
Spearman's ρ

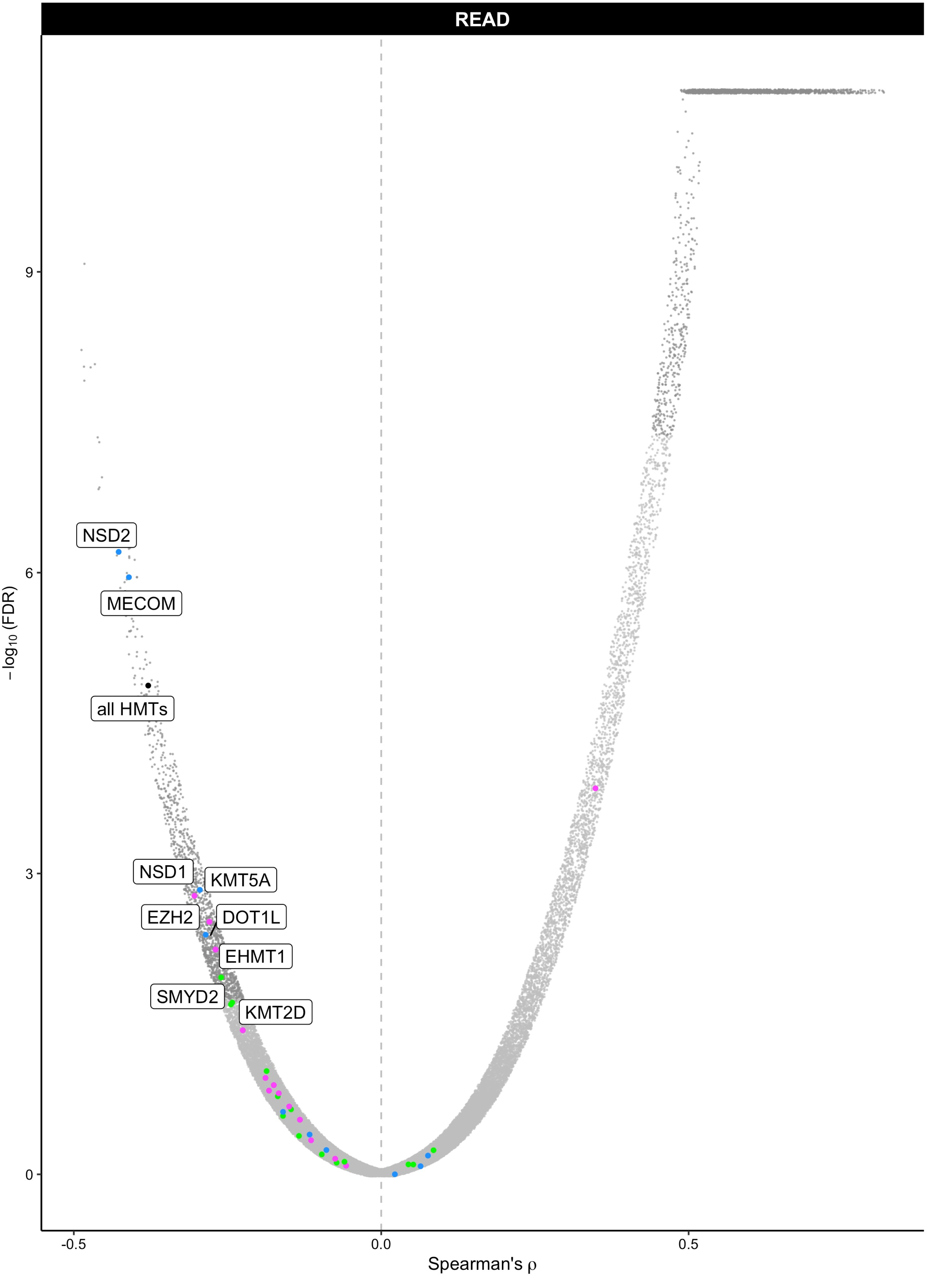
READ
NSD2
MECOM
all HMTs
NSD1
KMT5A
EZH2
DOT1L
EHMT1
SMYD2
KMT2D
9
6
3
0
-0.5
0.0
0.5
−log10 (FDR)
Spearman's ρ

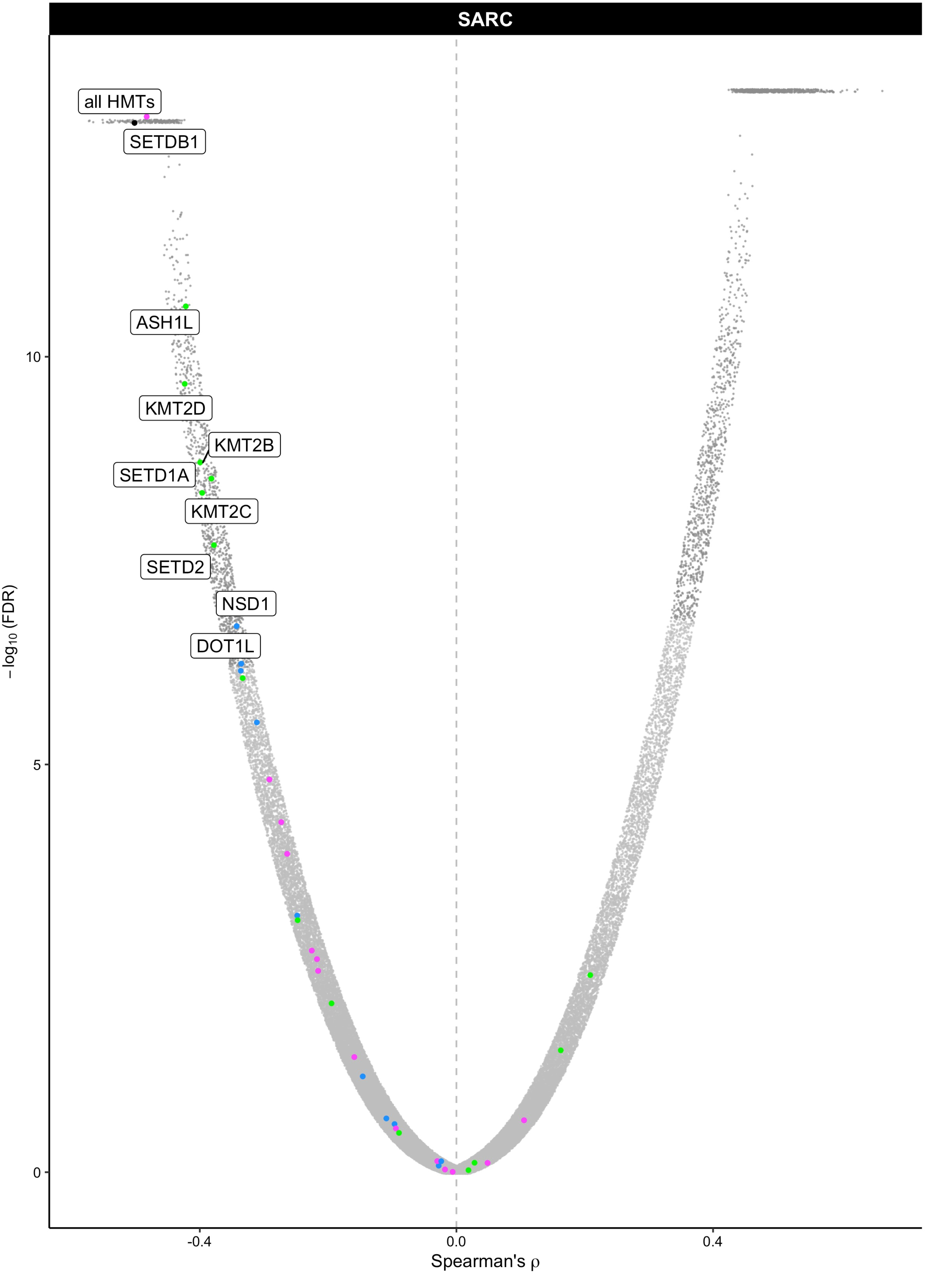
SARC
all HMTs
SETDB1
ASH1L
KMT2D
KMT2B
SETD1A
KMT2C
SETD2
NSD1
DOT1L
10
5
0
−log10 (FDR)
-0.4
0.0
0.4
Spearman's ρ

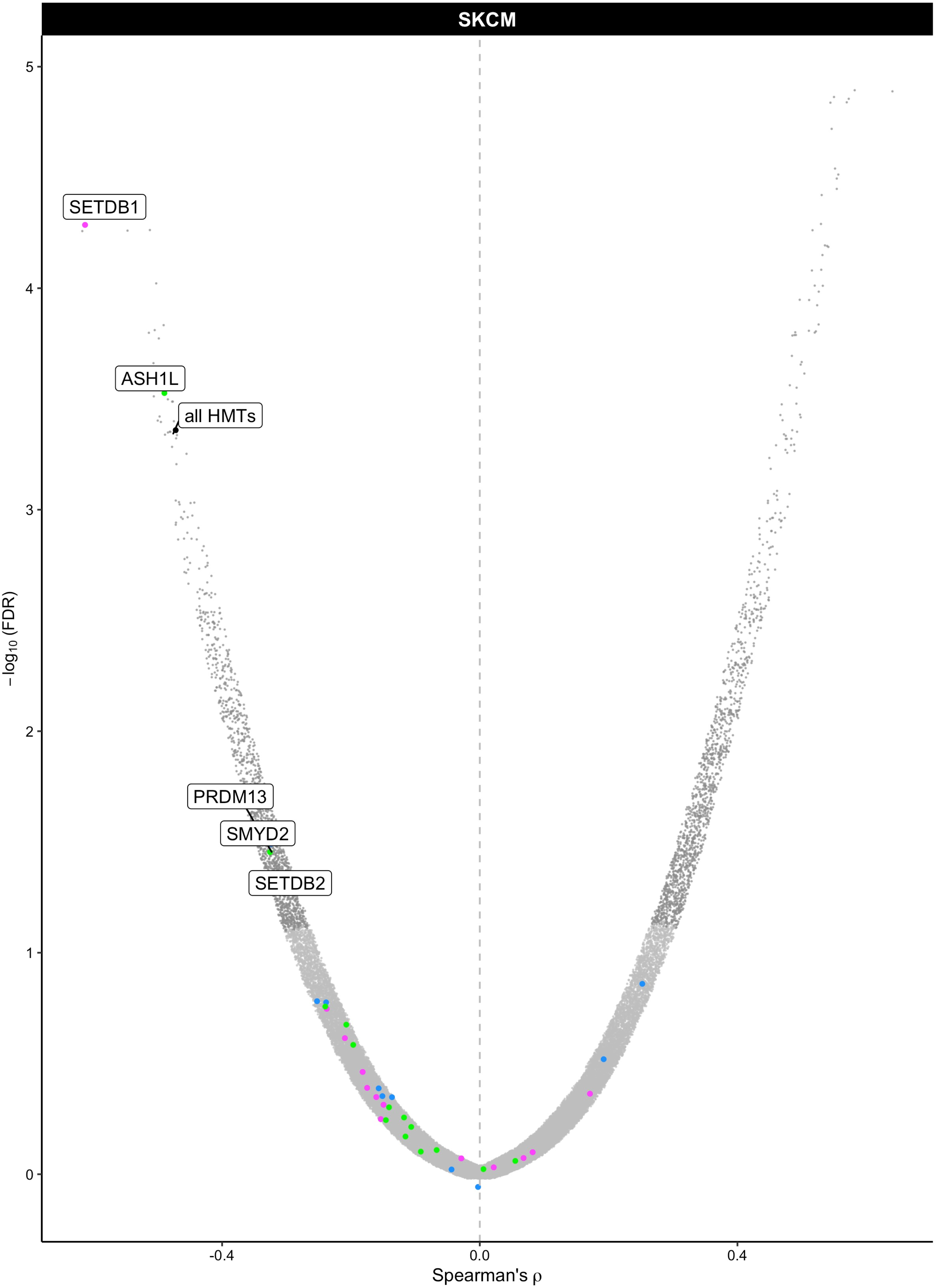

SKCM
SETDB1
ASH1L
all HMTs
PRDM13
SMYD2
SETDB2
−log10 (FDR)
Spearman's ρ
5
4
3
2
1
0
-0.4
0.0
0.4

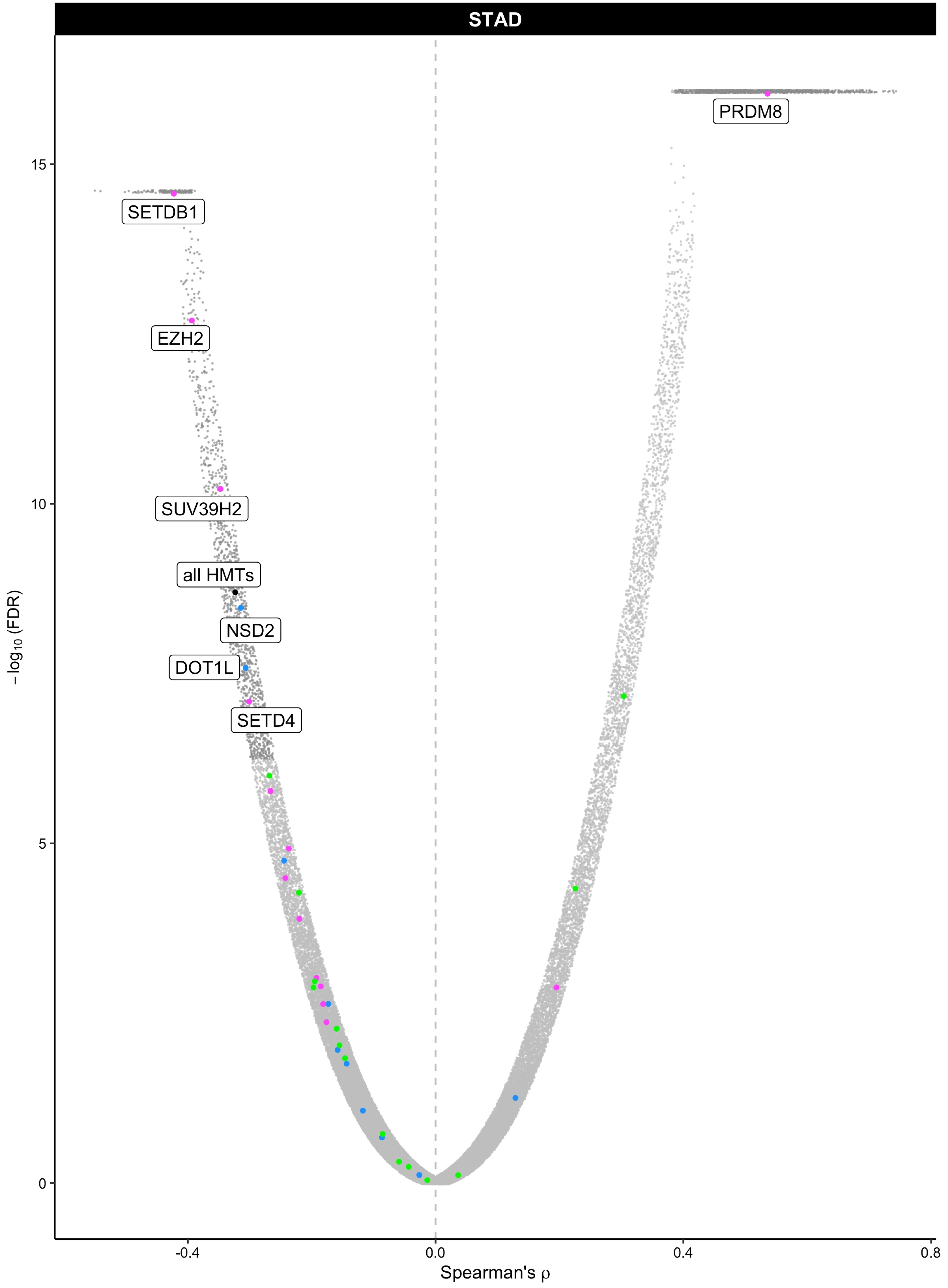
STAD
PRDM8
SETDB1
EZH2
SUV39H2
all HMTs
NSD2
DOT1L
SETD4
15
10
5
0
−log10 (FDR)
-0.4
0.0
0.4
0.8
Spearman's ρ

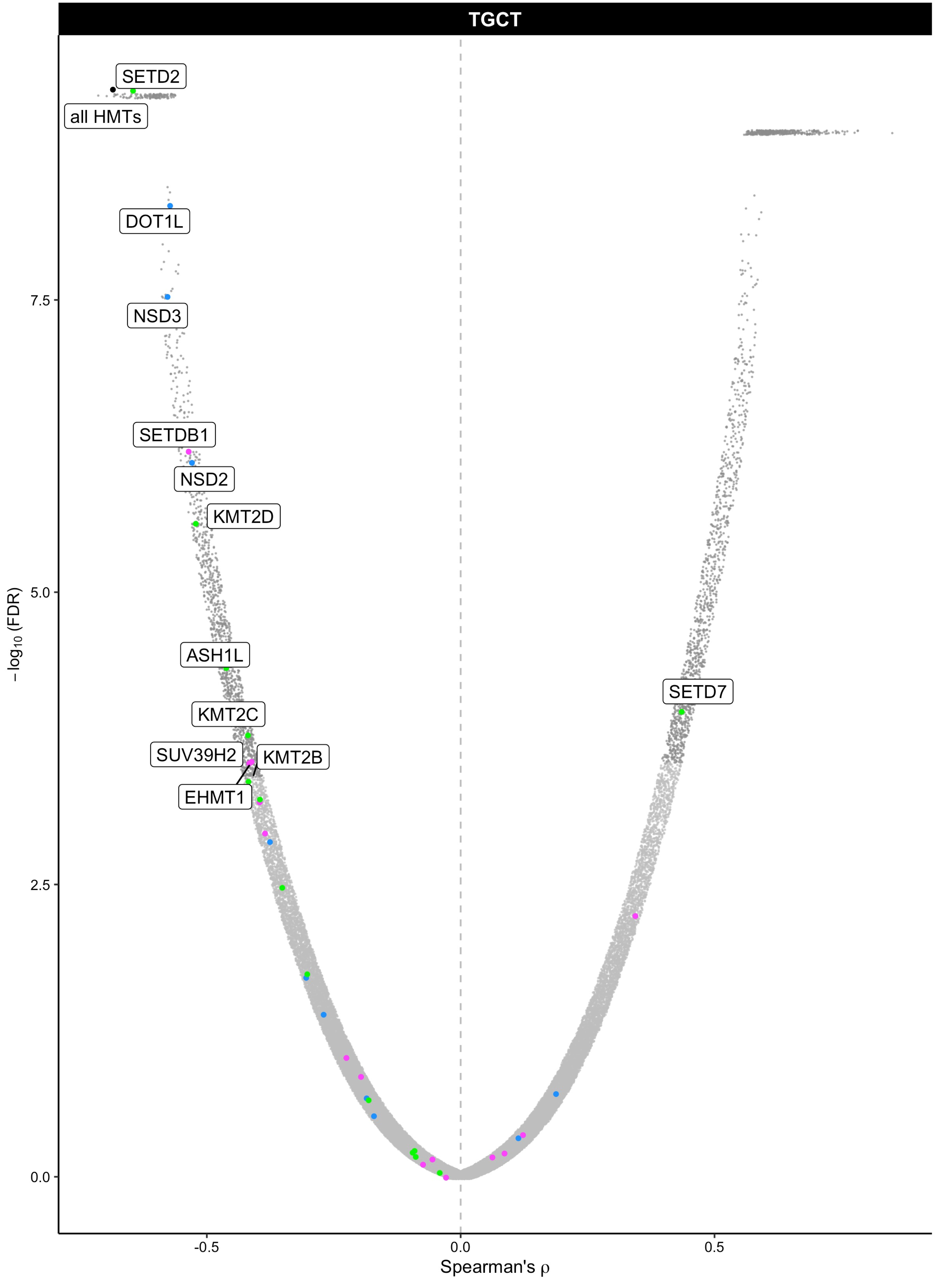

TGCT
SETD2
all HMTs
DOT1L
NSD3
SETDB1
NSD2
KMT2D
ASH1L
KMT2C
SUV39H2
KMT2B
EHMT1
SETD7
7.5
5.0
2.5
0.0
−log10 (FDR)
-0.5
0.0
0.5
Spearman's ρ

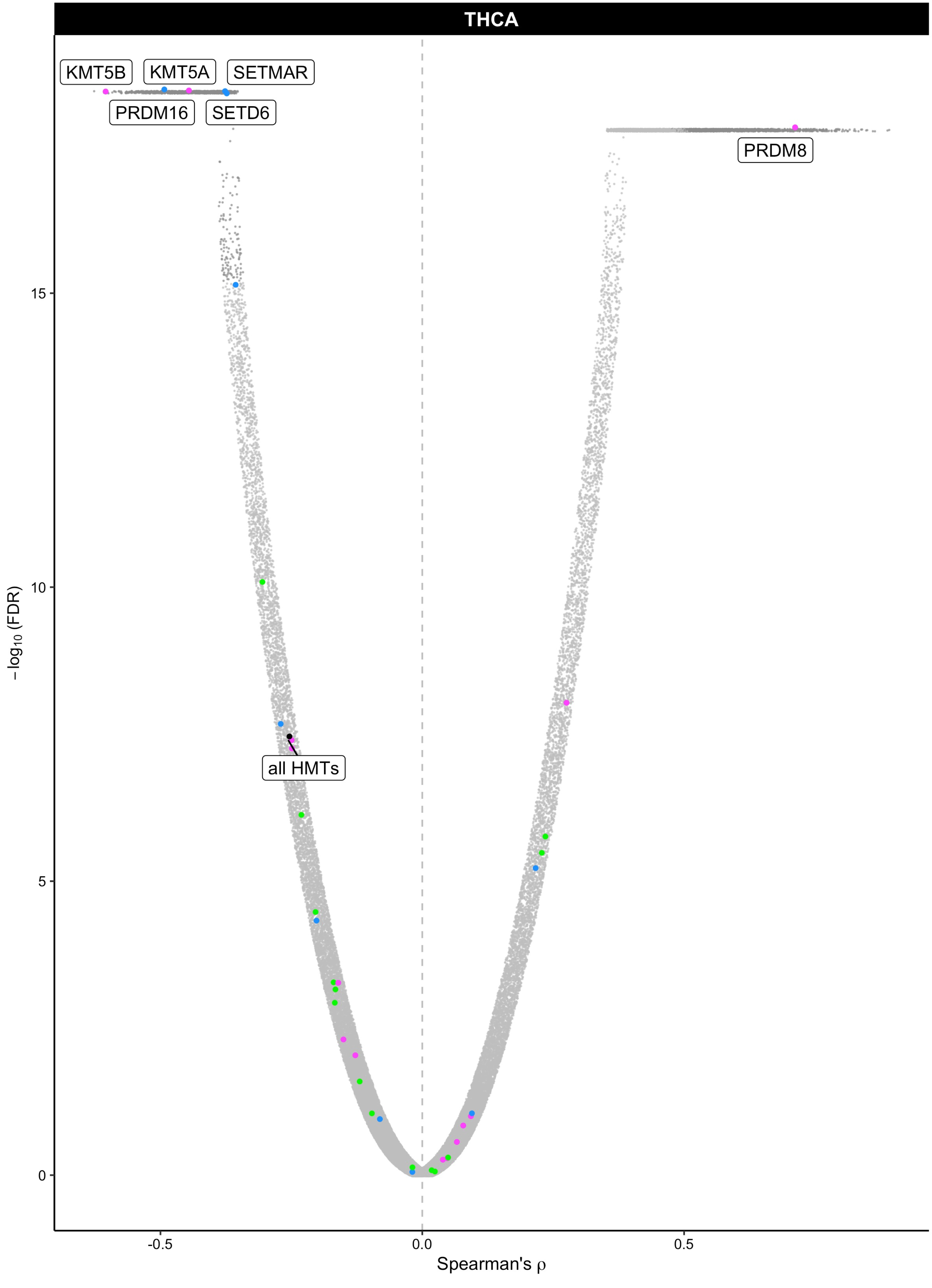
THCA
KMT5B
KMT5A
SETMAR
PRDM16
SETD6
PRDM8
all HMTs
15
10
5
0
−log10 (FDR)
-0.5
0.0
0.5
Spearman's ρ

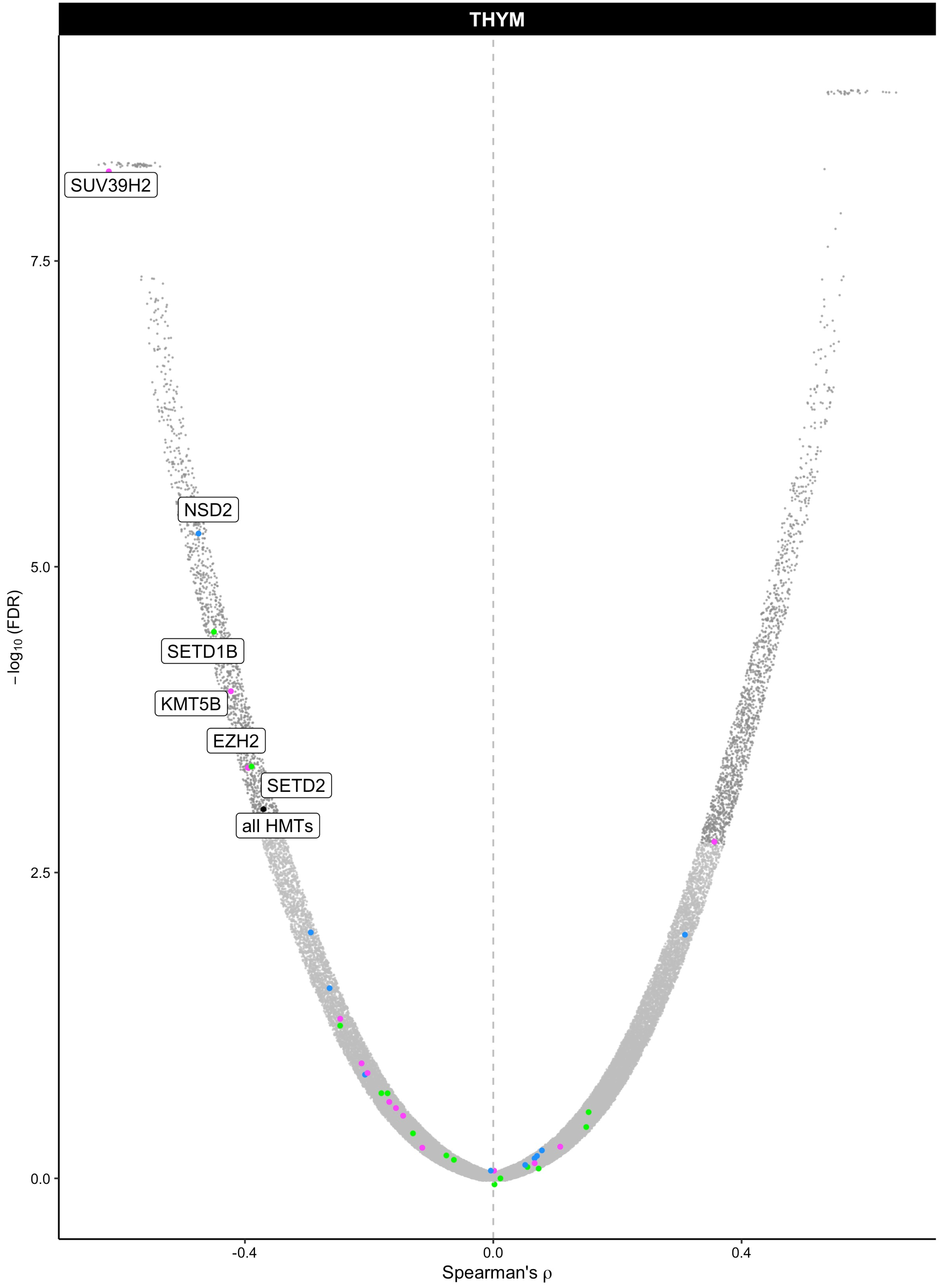
THYM
SUV39H2
NSD2
SETD1B
KMT5B
EZH2
SETD2
all HMTs
7.5
5.0
2.5
0.0
−log10 (FDR)
-0.4
0.0
0.4
Spearman's ρ

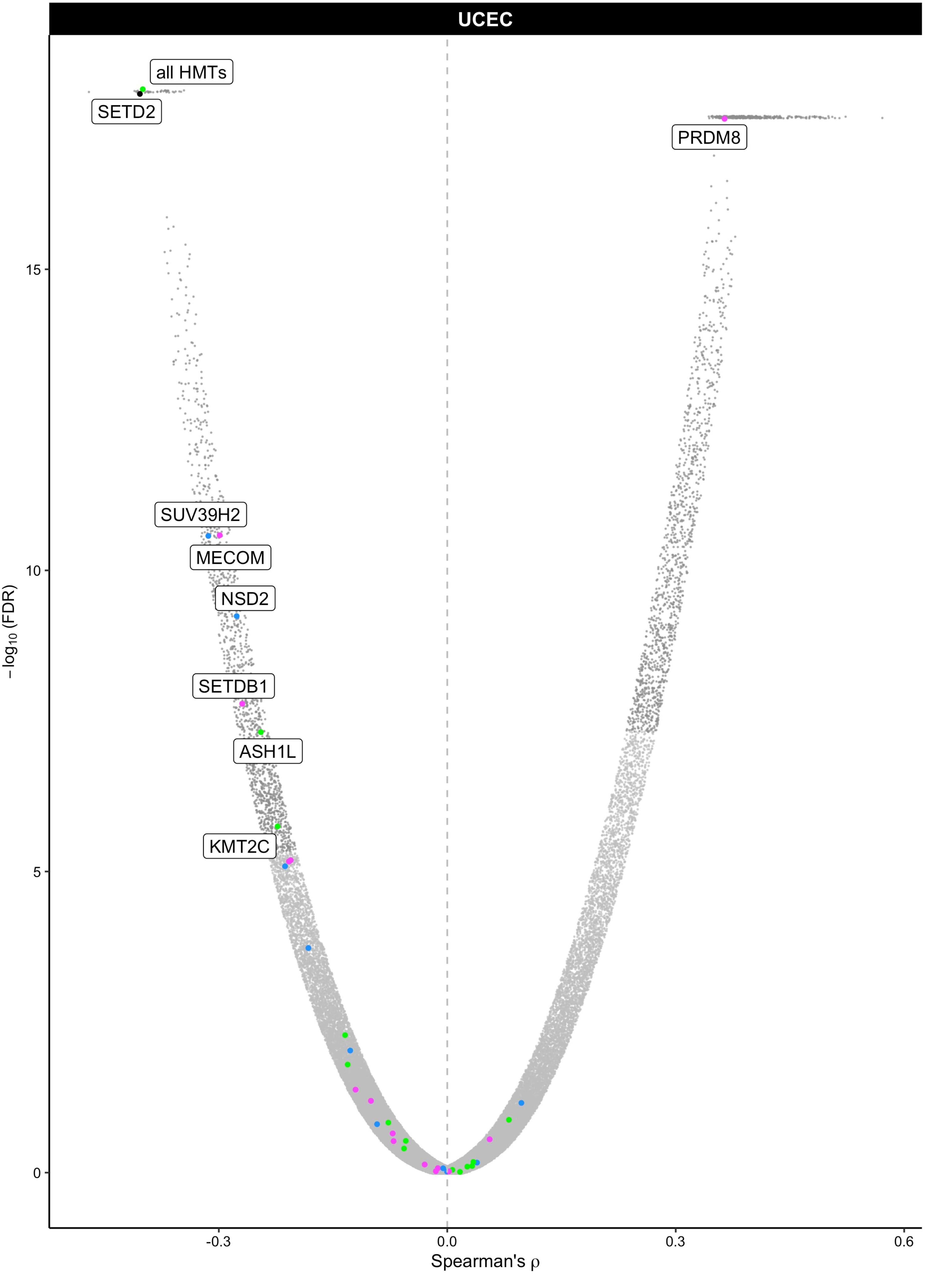
UCEC
all HMTs
SETD2
PRDM8
SUV39H2
MECOM
NSD2
SETDB1
ASH1L
KMT2C
15
10
5
0
−log10 (FDR)
-0.3
0.0
0.3
0.6
Spearman's ρ

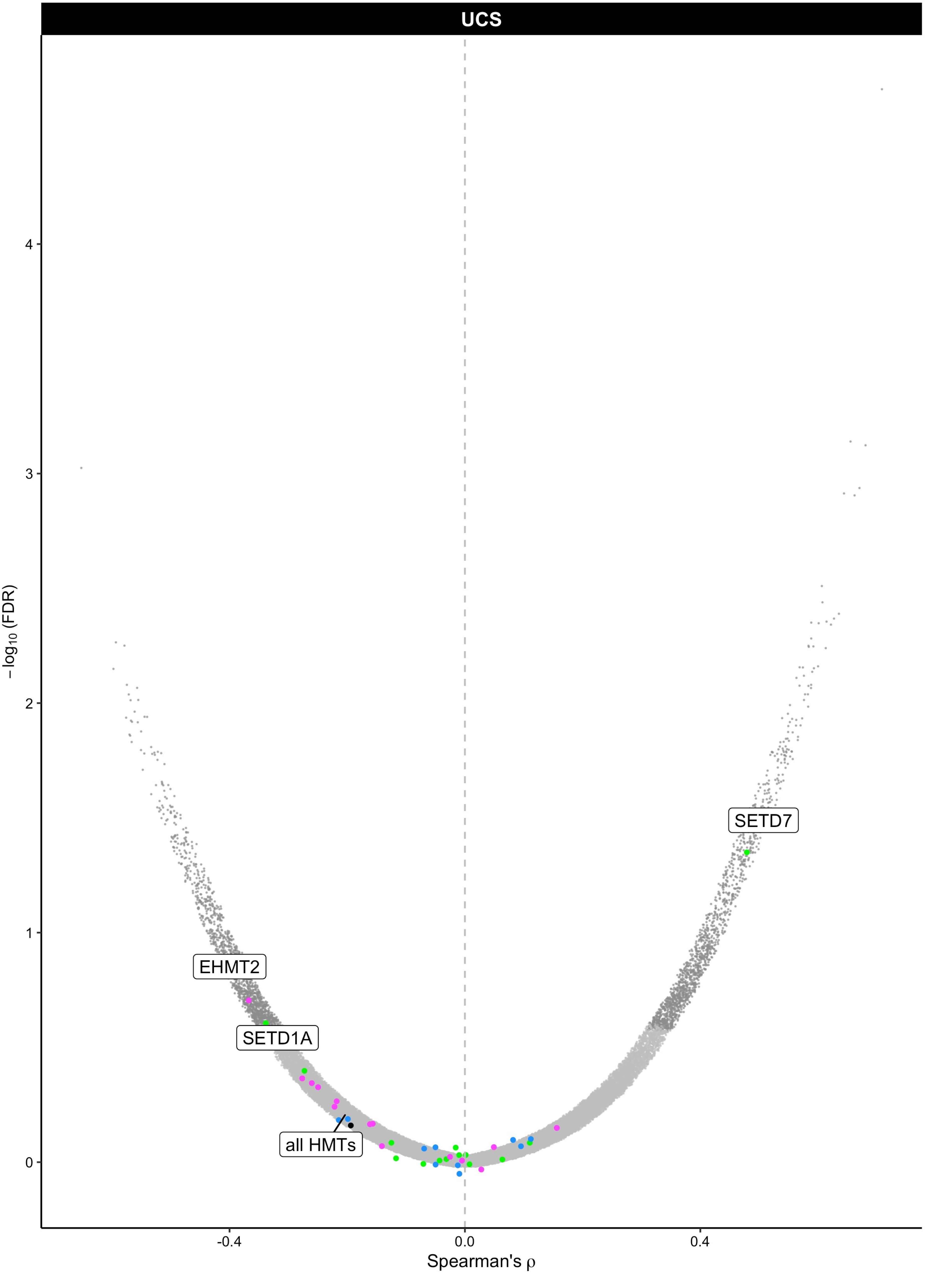

UCS
SETD7
EHMT2
SETD1A
all HMTs
4
3
2
1
0
-0.4
0.0
0.4
Spearman's ρ
−log10 (FDR)

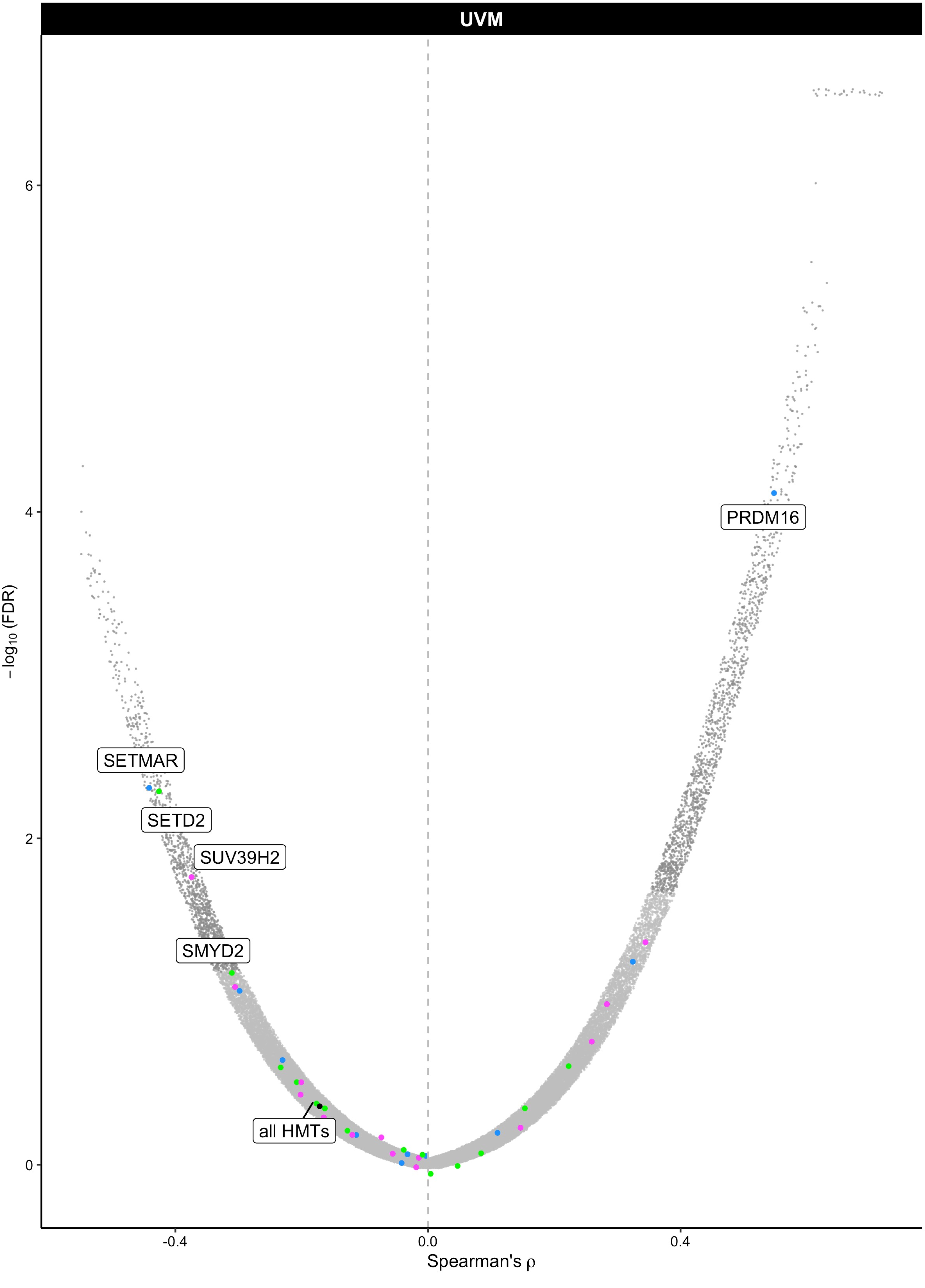

UVM
−log10 (FDR)
6
4
2
0
-0.4
0.0
0.4
Spearman's ρ
PRDM16
SETMAR
SETD2
SUV39H2
SMYD2
all HMTs
