## Supplemental Figures and Tables for "Histone methylation has a direct metabolic role in human cells": Supp File 3. GTEX NNMT HMTs fullplots.pdf

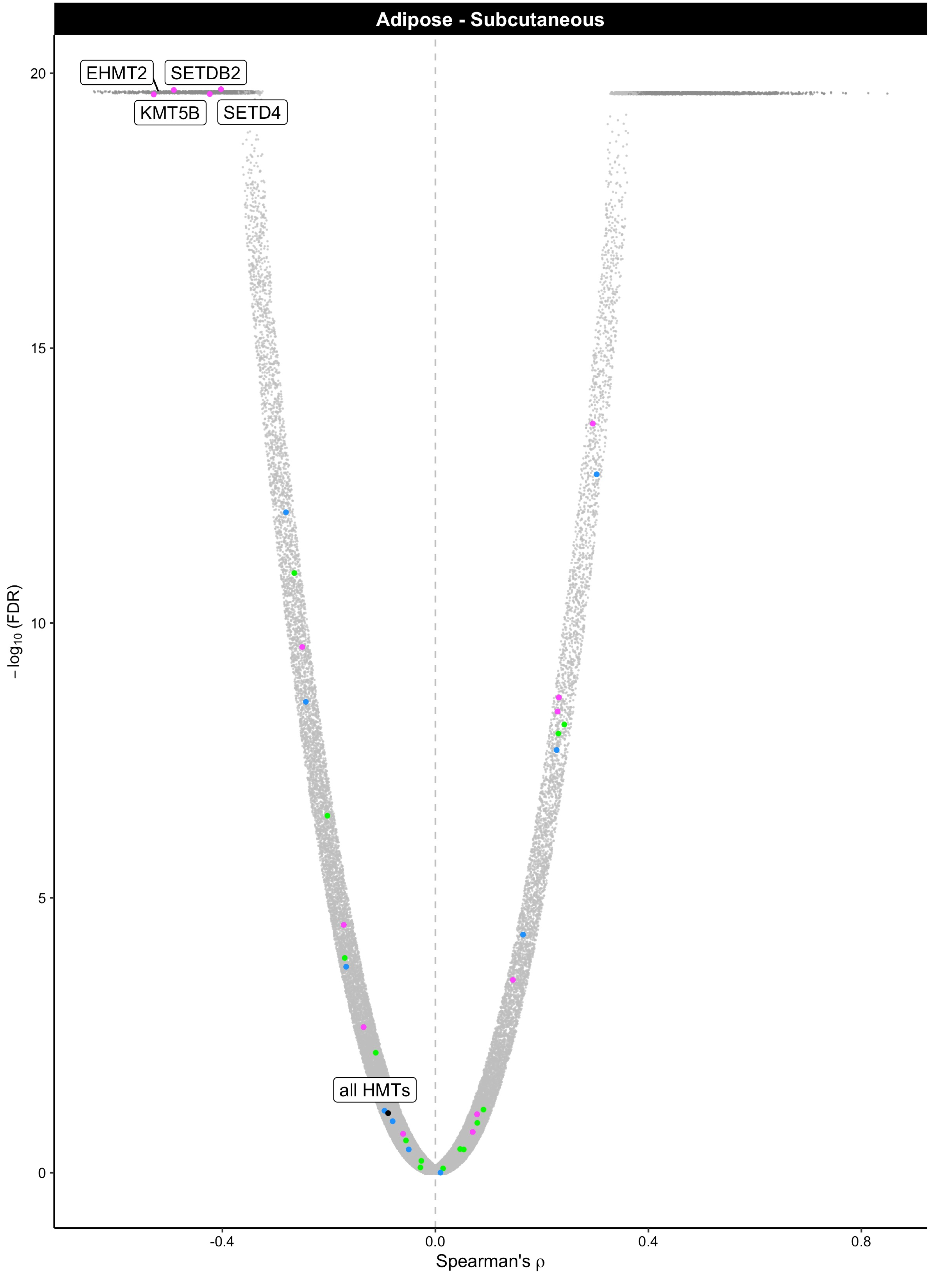

Adipose - Subcutaneous
EHMT2
SETDB2
KMT5B
SETD4
all HMTs
20
15
10
5
0
−log10 (FDR)
-0.4
0.0
0.4
0.8
Spearman's ρ

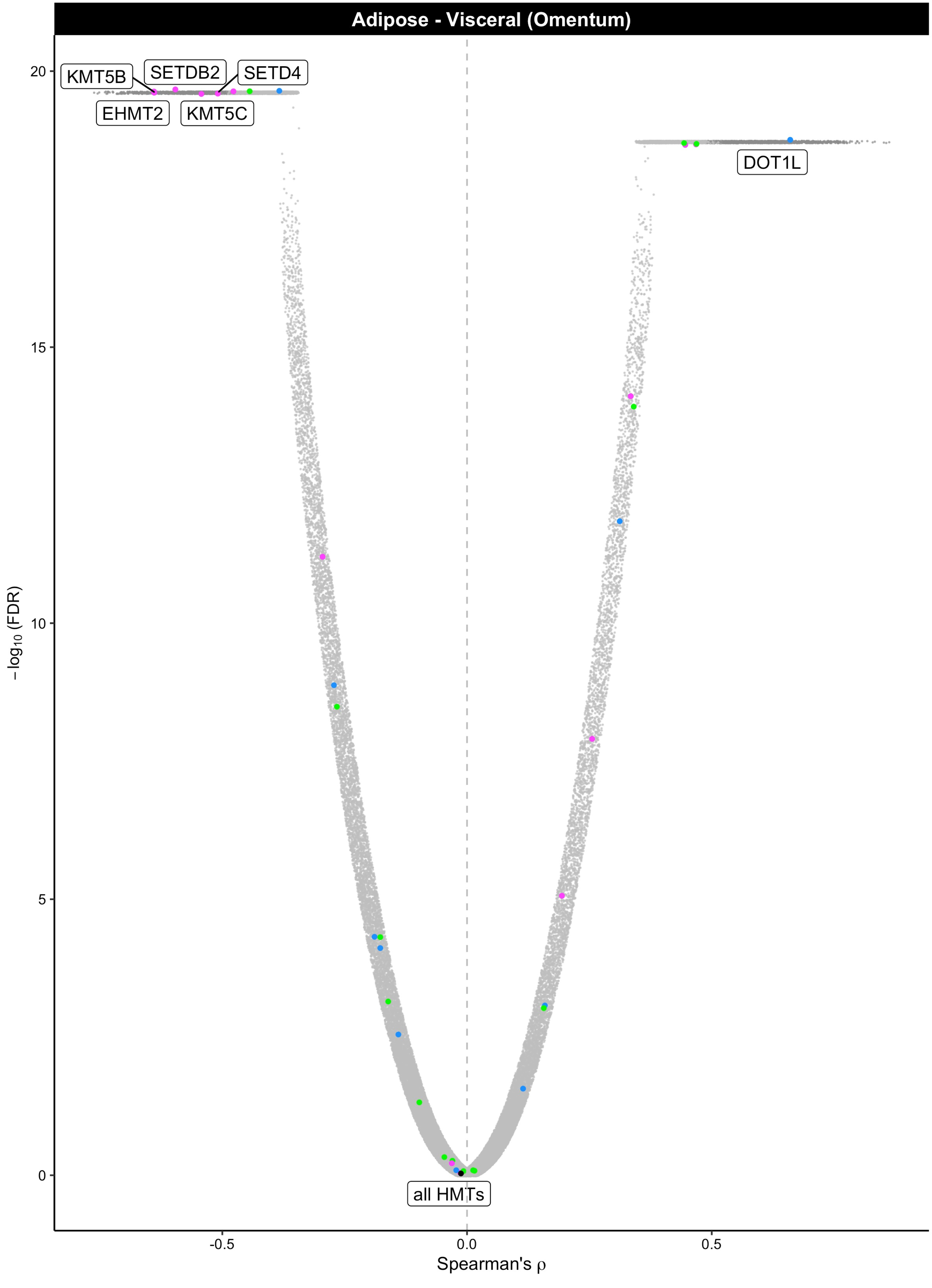

Adipose - Visceral (Omentum)
KMT5B
SETDB2
SETD4
EHMT2
KMT5C
DOT1L
all HMTs
20
15
10
5
0
−log10 (FDR)
-0.5
0.0
0.5
Spearman's ρ

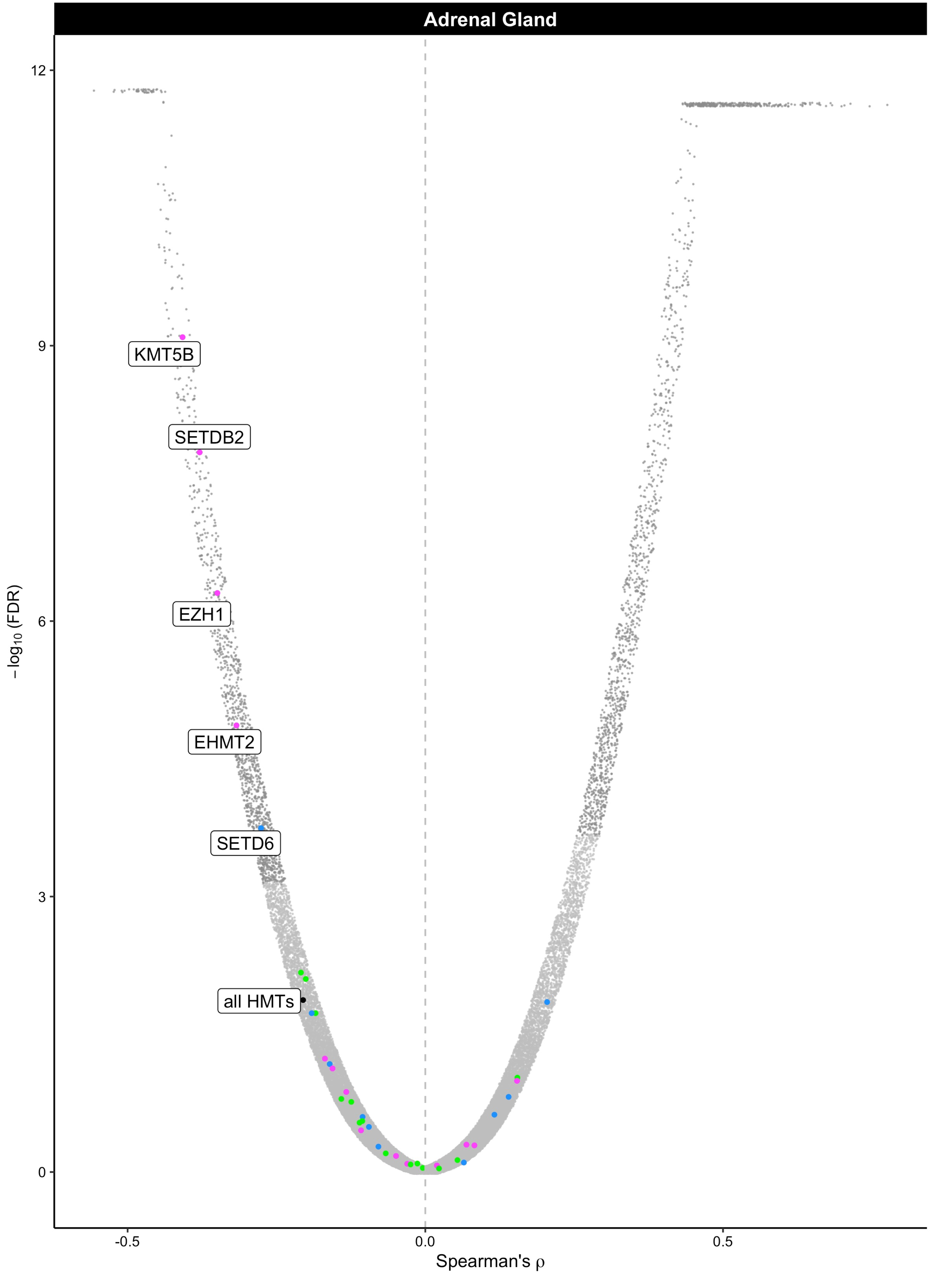
Adrenal Gland
KMT5B
SETDB2
EZH1
EHMT2
SETD6
all HMTs
−log10 (FDR)
12
9
6
3
0
-0.5
0.0
0.5
Spearman's ρ

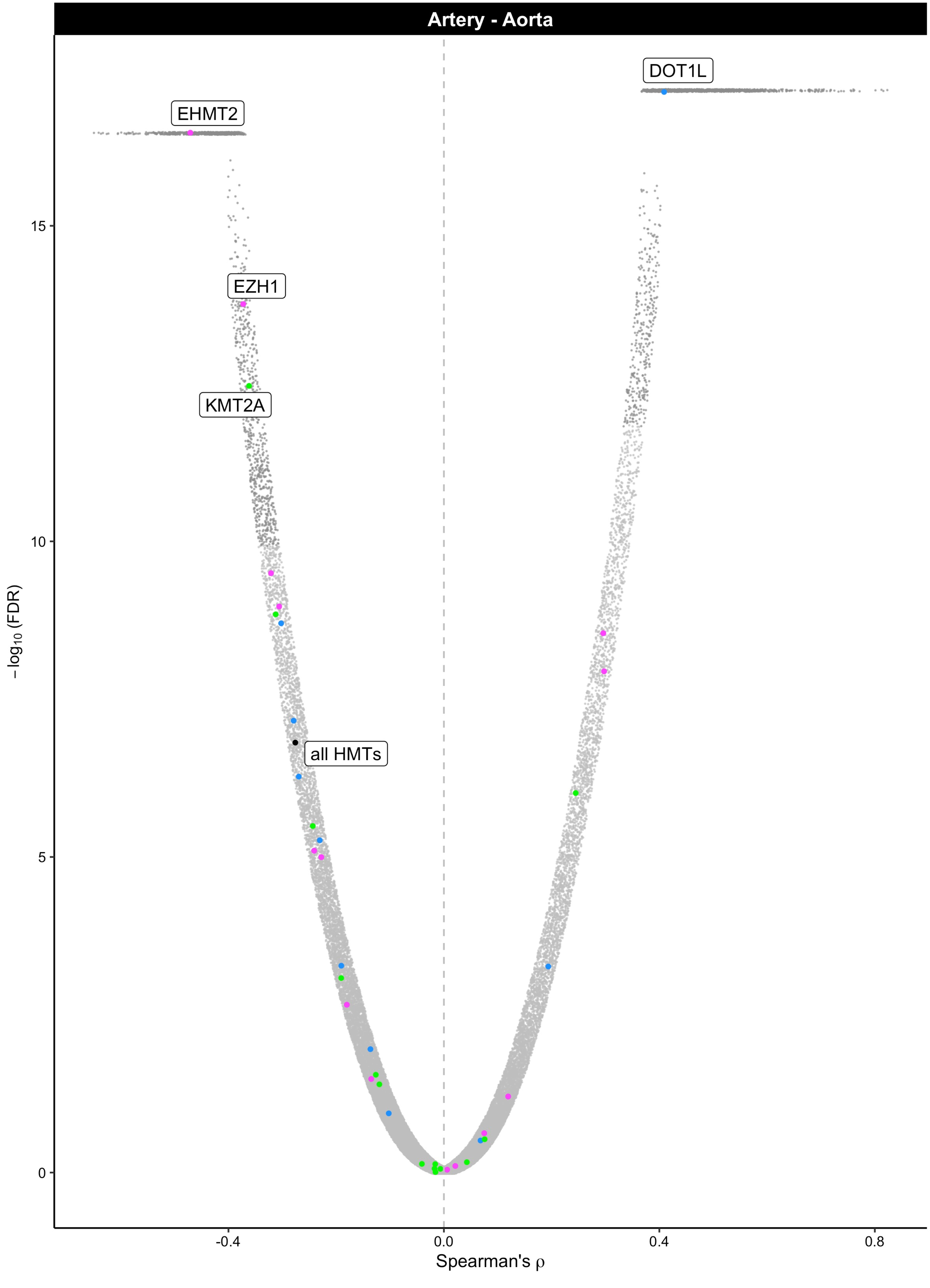

Artery - Aorta
DOT1L
EHMT2
EZH1
KMT2A
all HMTs
15
10
5
0
−log10 (FDR)
-0.4
0.0
0.4
0.8
Spearman's ρ

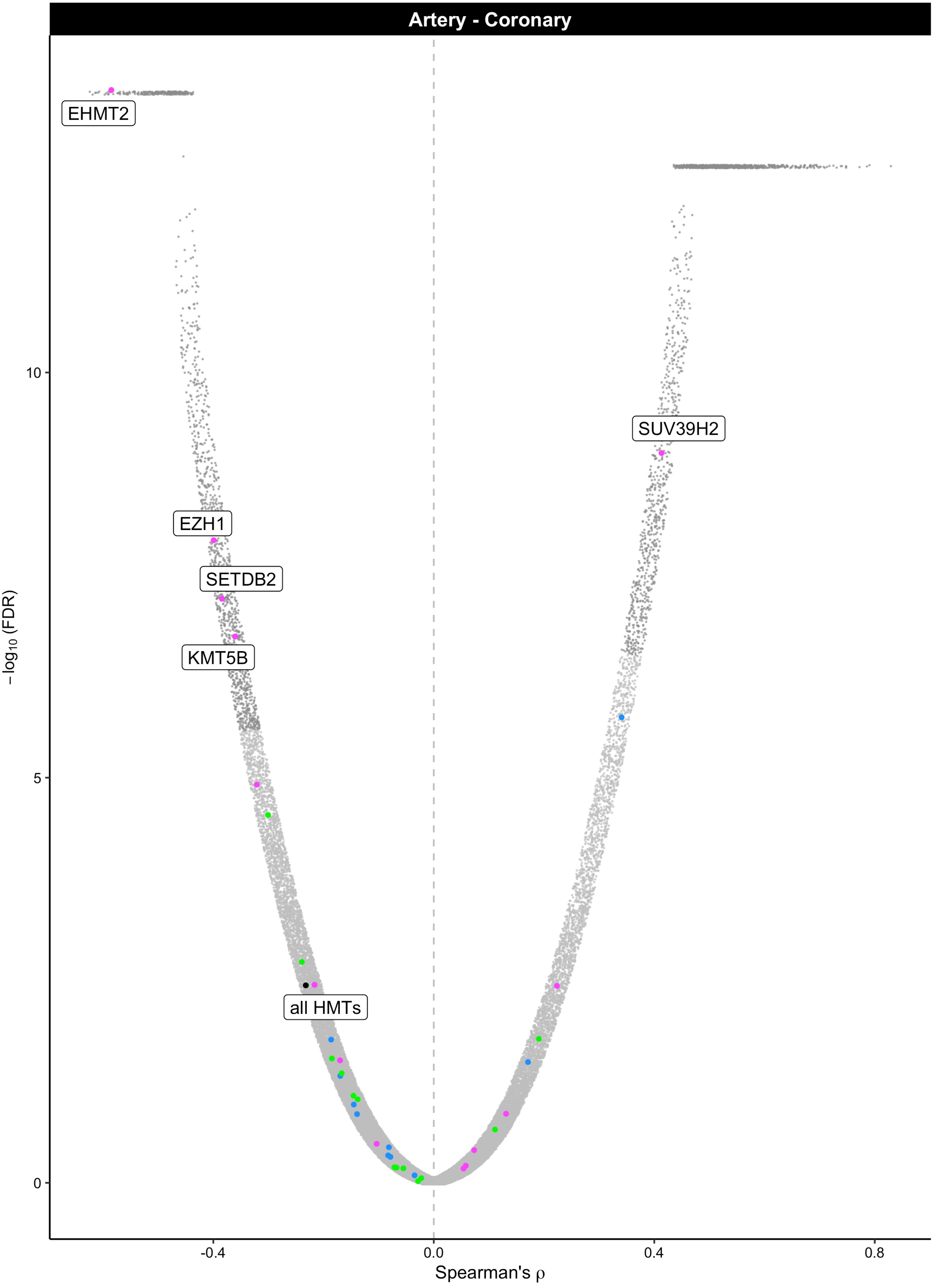

Artery - Coronary
EHMT2
SUV39H2
EZH1
SETDB2
KMT5B
all HMTs
10
5
0
-0.4
0.0
0.4
0.8
−log10 (FDR)
Spearman's ρ

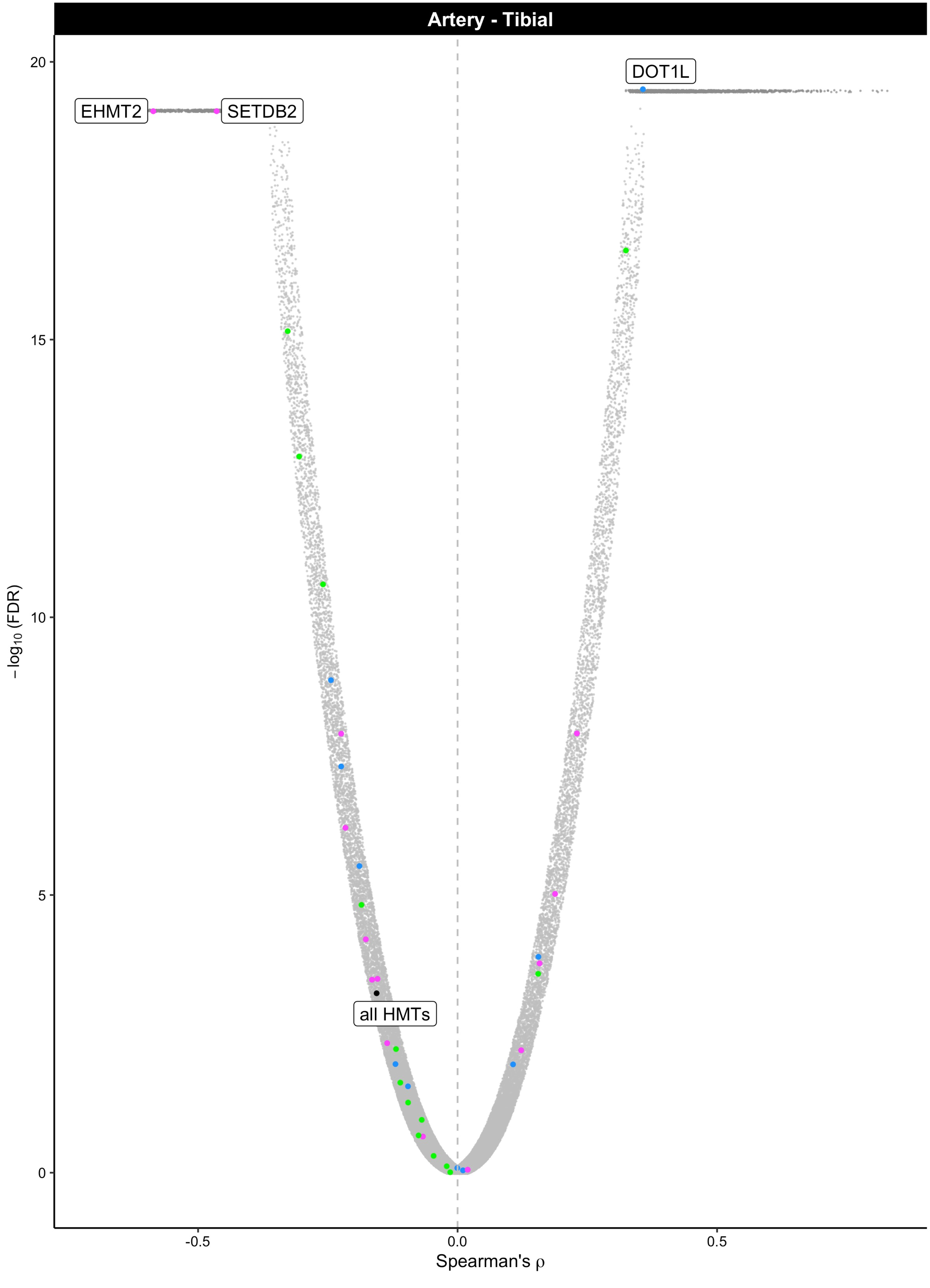

Artery - Tibial
DOT1L
EHMT2
SETDB2
all HMTs
20
15
10
5
0
−log10 (FDR)
-0.5
0.0
0.5
Spearman's ρ

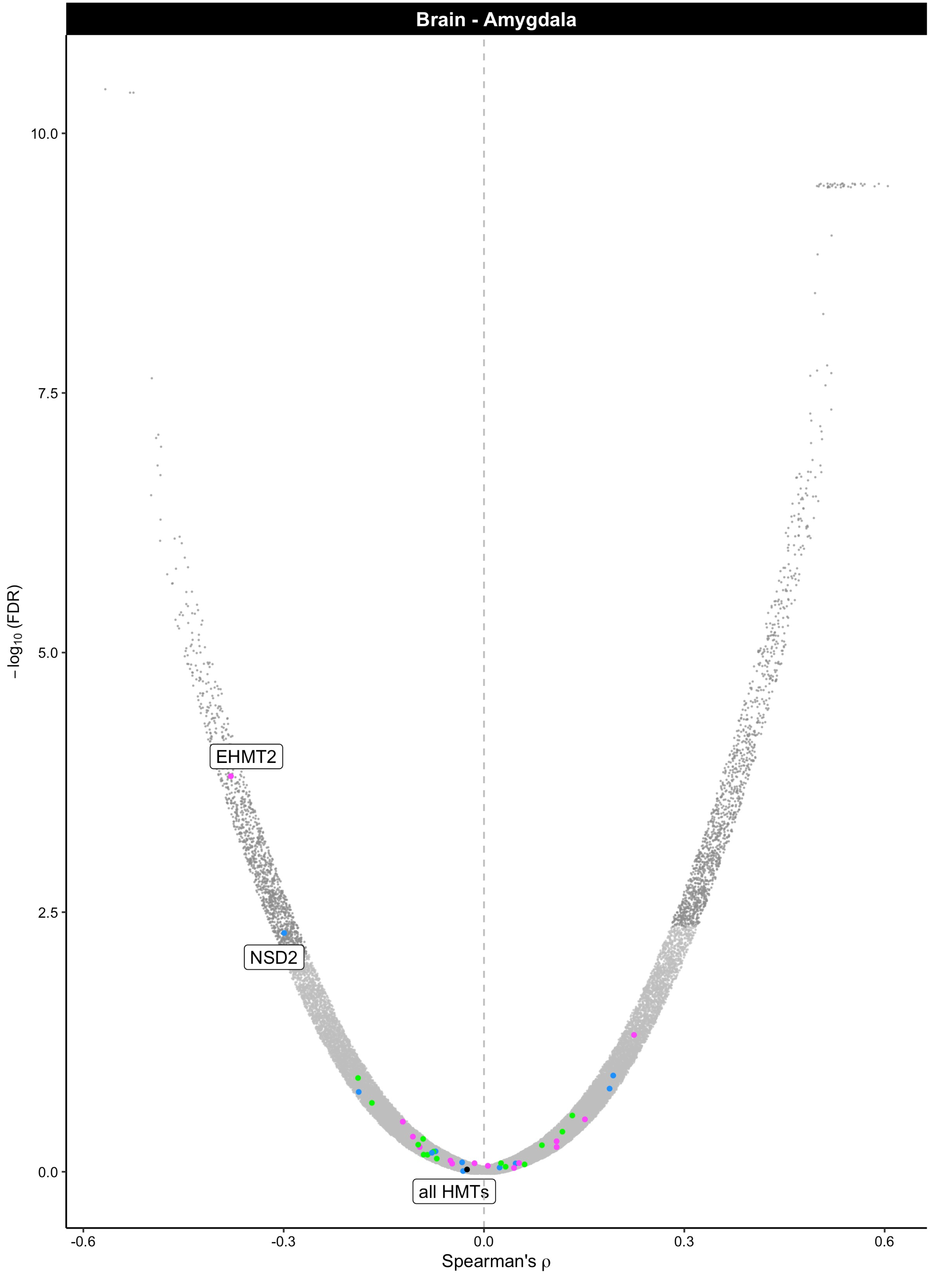
Brain - Amygdala
EHMT2
NSD2
all HMTs
−log10 (FDR)
10.0
7.5
5.0
2.5
0.0
-0.6
-0.3
0.0
0.3
0.6
Spearman's ρ

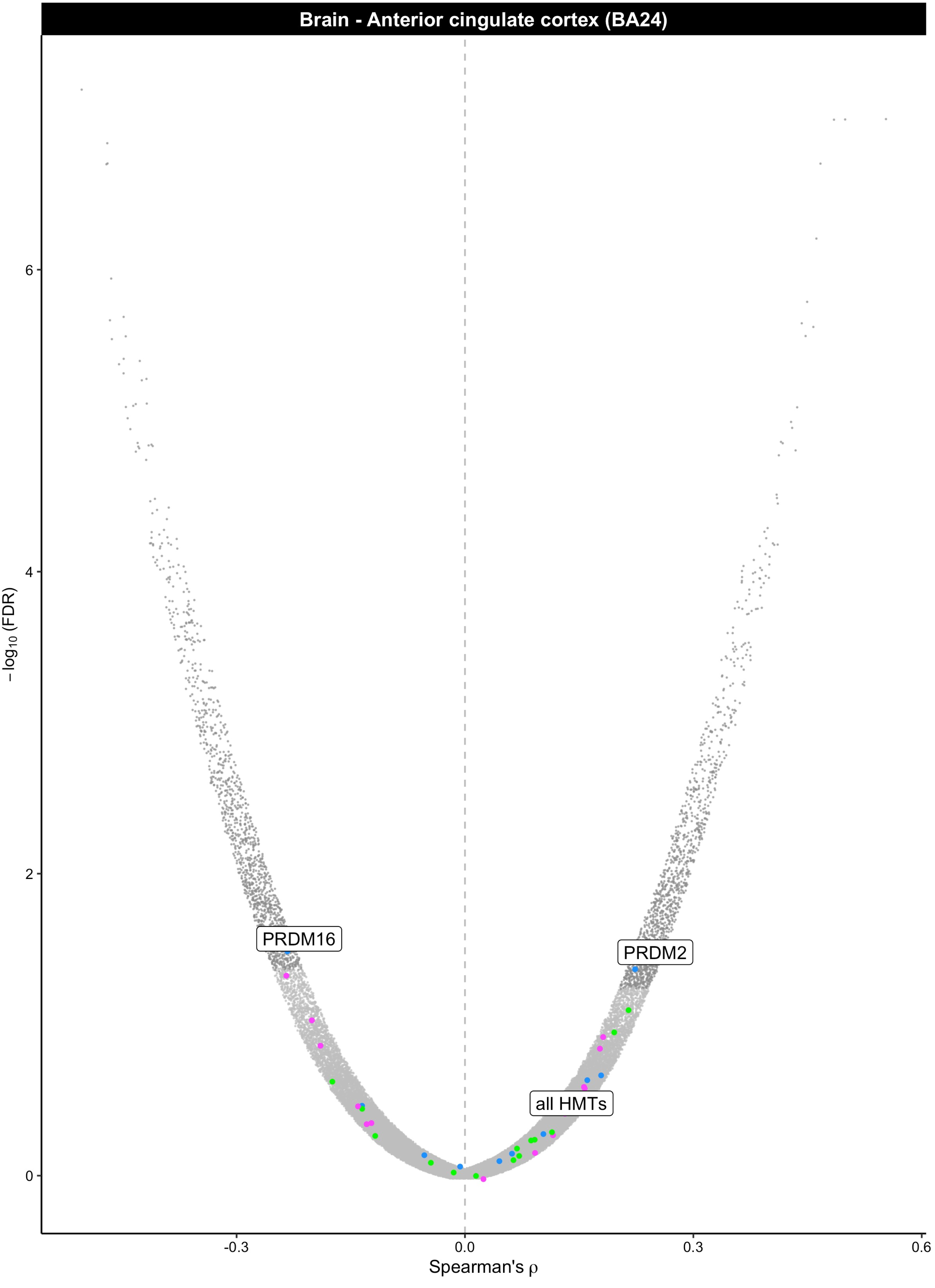
Brain - Anterior cingulate cortex (BA24)
PRDM16
PRDM2
all HMTs
−log10 (FDR)
0
2
4
6
-0.3
0.0
0.3
0.6
Spearman's ρ

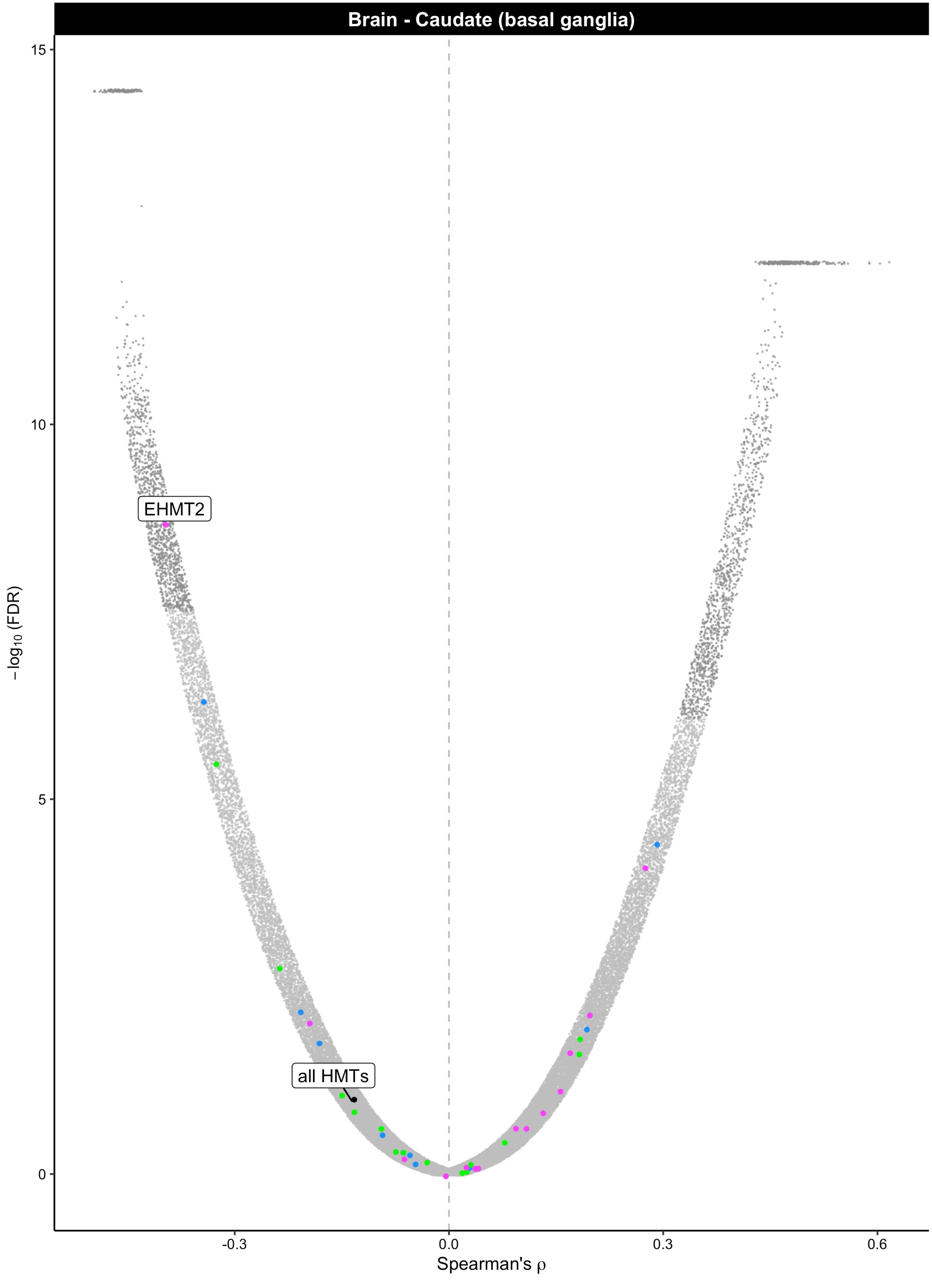
Brain - Caudate (basal ganglia)
15
10
5
0
−log10 (FDR)
EHMT2
all HMTs
-0.3
0.0
0.3
0.6
Spearman's ρ

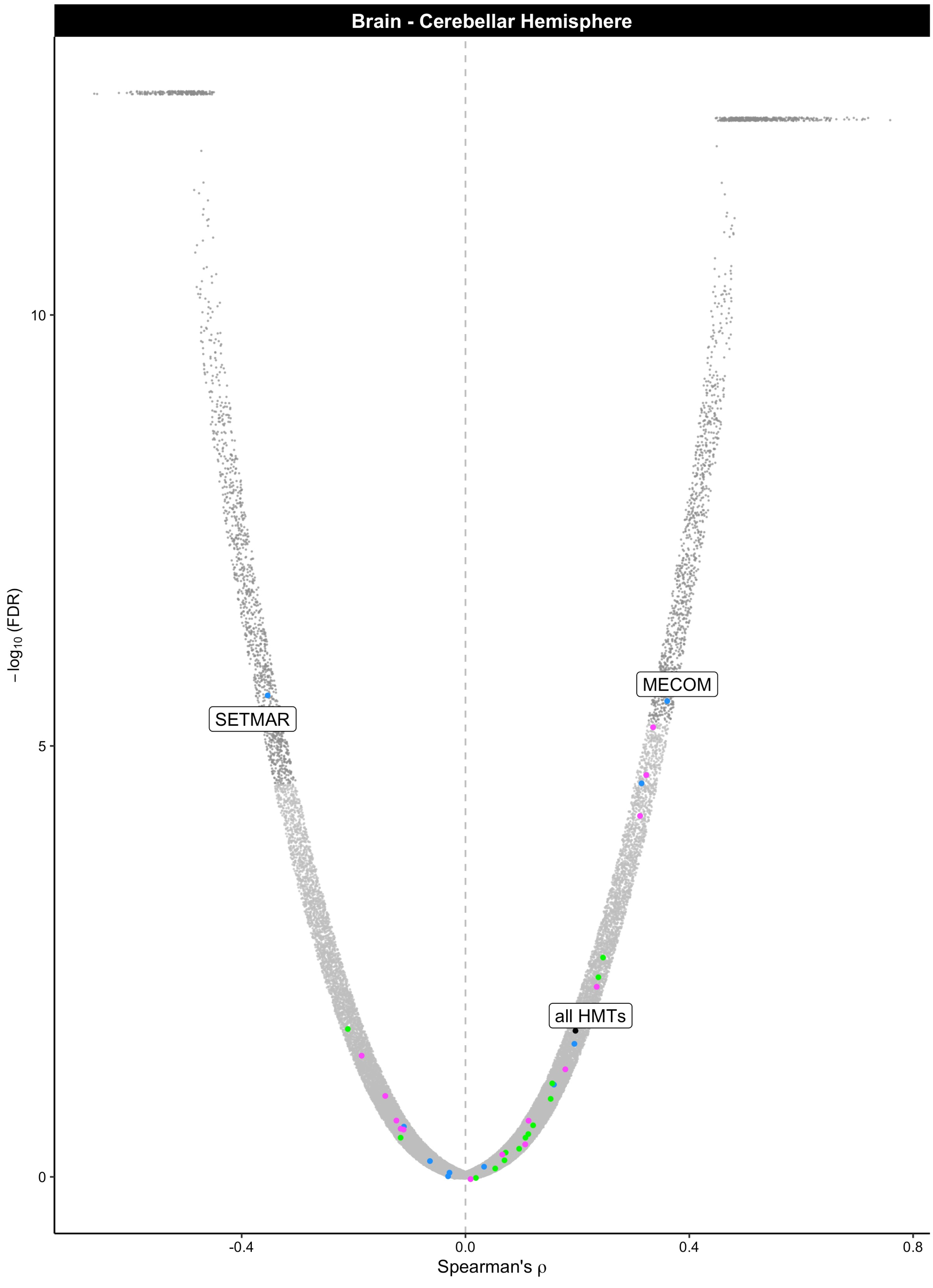

Brain - Cerebellar Hemisphere
SETMAR
MECOM
all HMTs
10
5
0
−log10 (FDR)
-0.4
0.0
0.4
0.8
Spearman's ρ

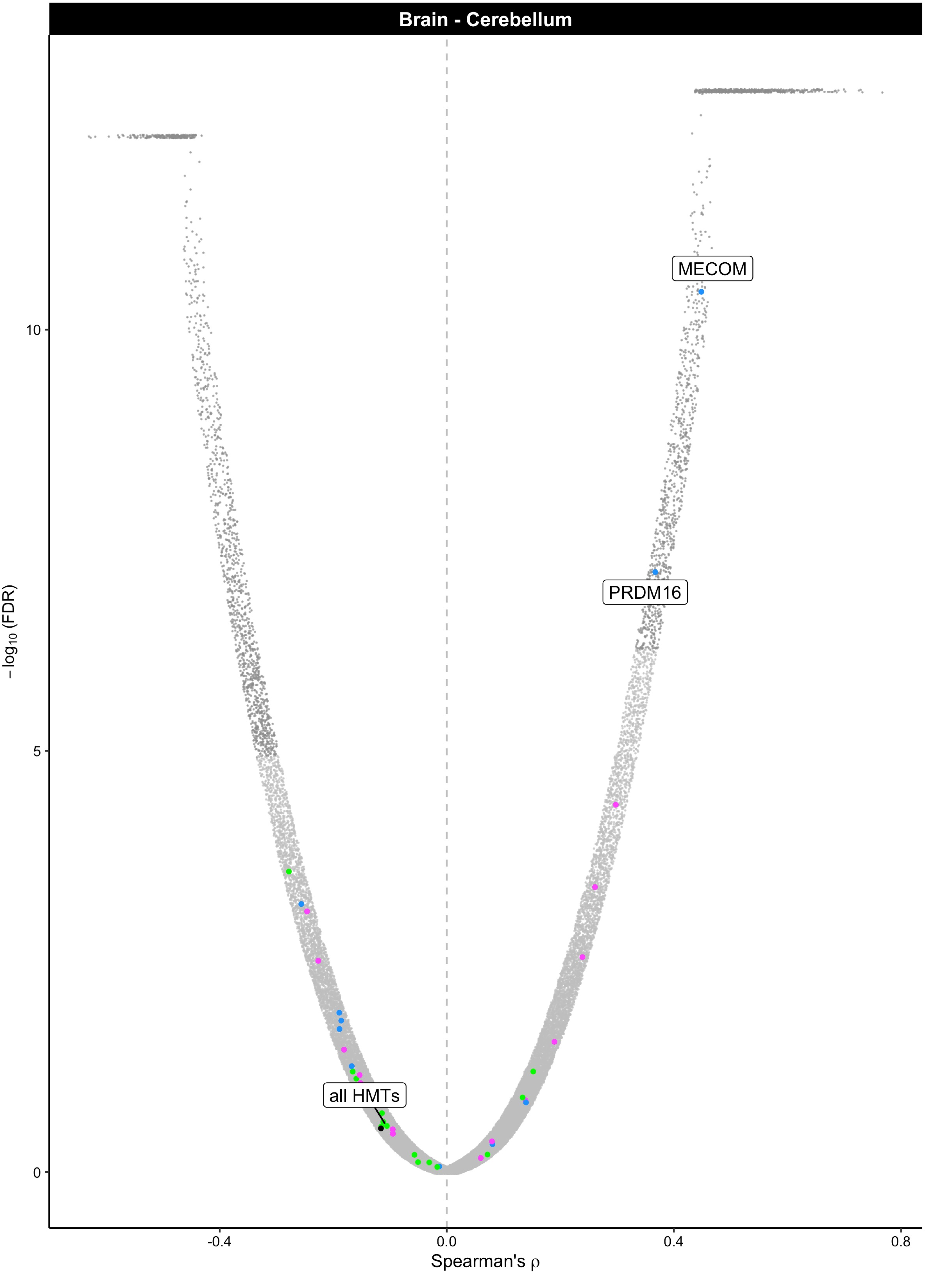

Brain - Cerebellum
MECOM
PRDM16
all HMTs
$-\log_{10}$(FDR)
10
5
0
-0.4
0.0
0.4
0.8
Spearman's ρ

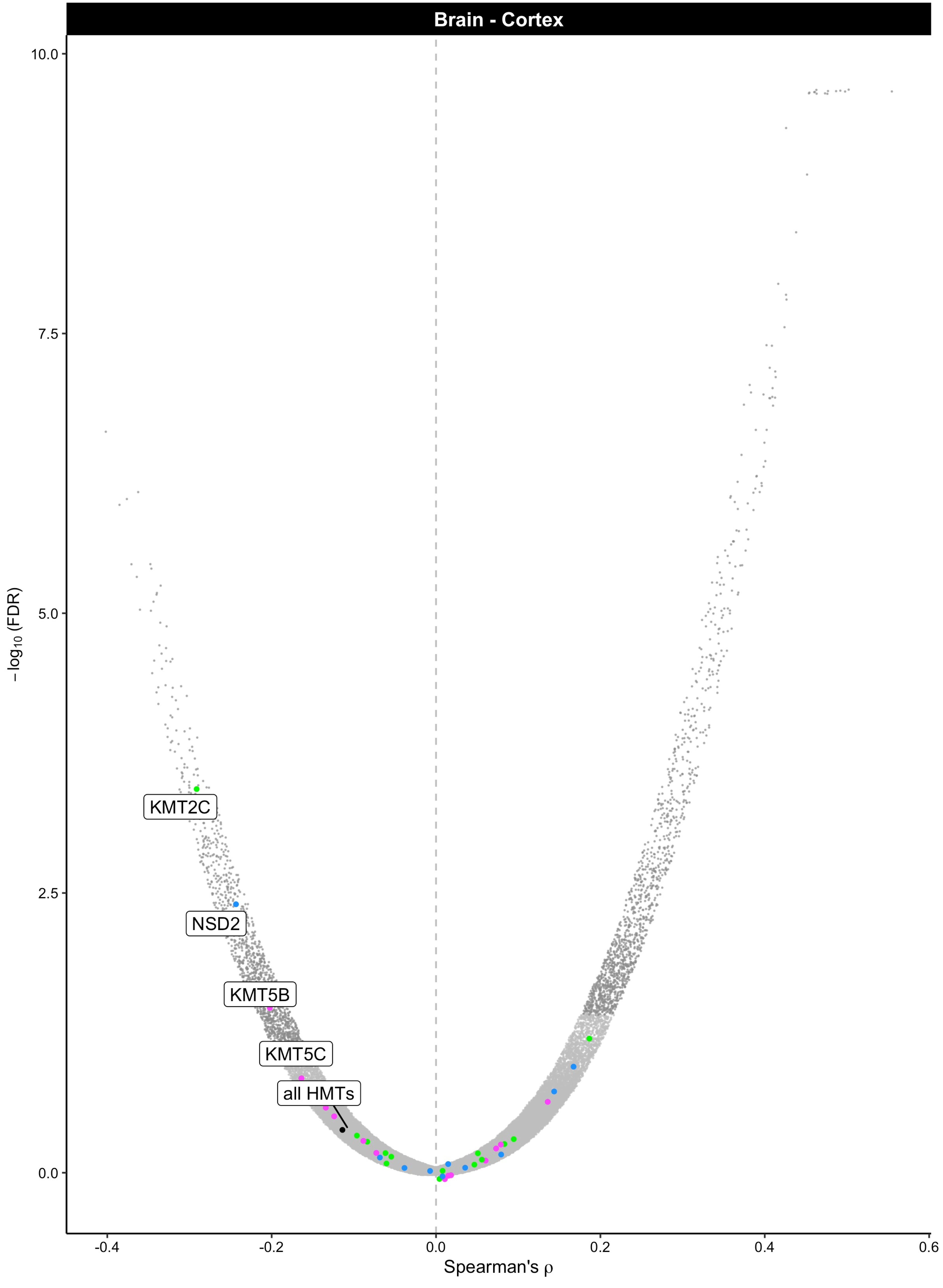

Brain - Cortex
10.0
7.5
5.0
2.5
0.0
−log10 (FDR)
-0.4
-0.2
0.0
0.2
0.4
0.6
Spearman's ρ
KMT2C
NSD2
KMT5B
KMT5C
all HMTs

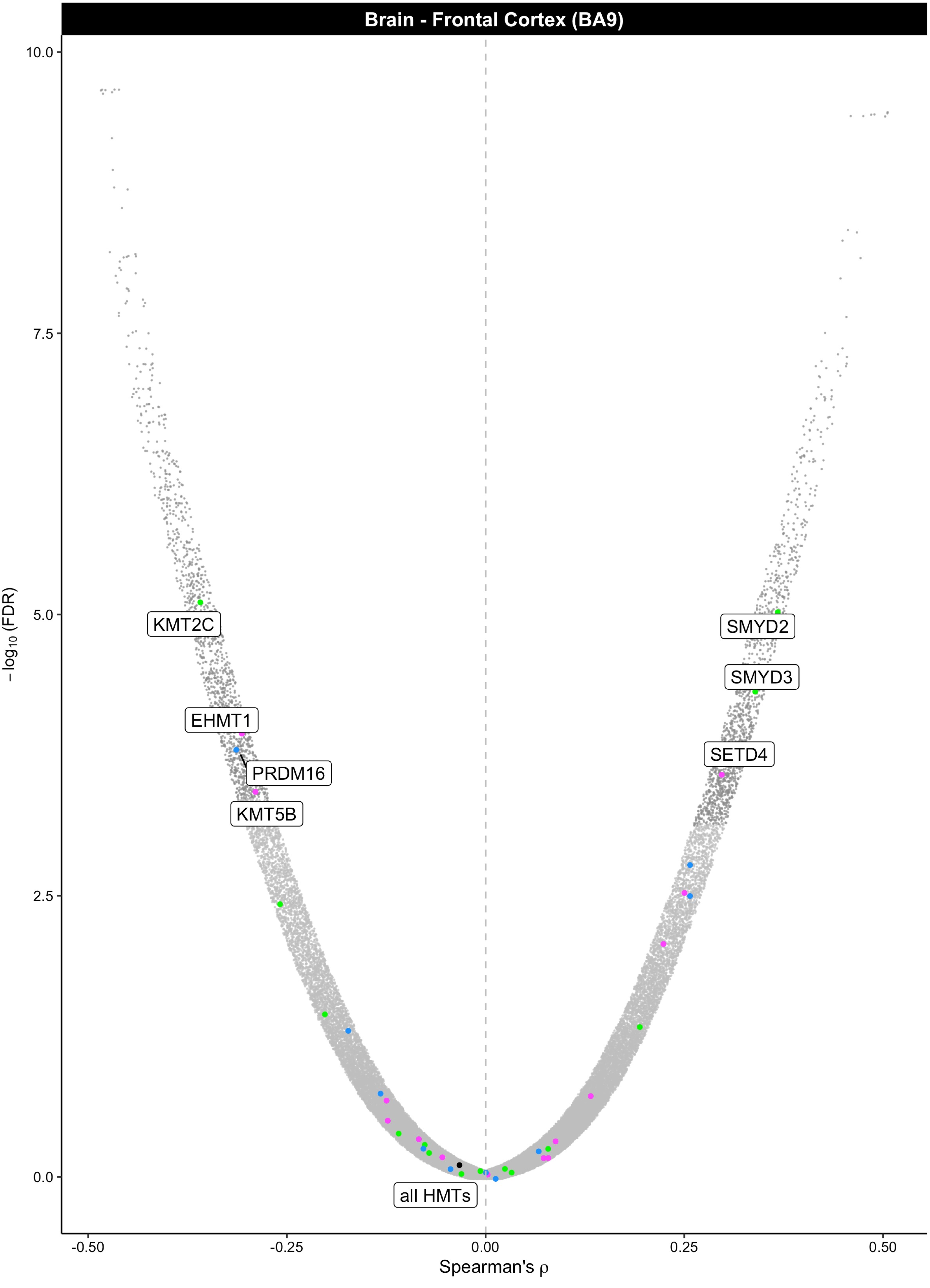

Brain - Frontal Cortex (BA9)
10.0
7.5
5.0
2.5
0.0
−log10 (FDR)
-0.50
-0.25
0.00
0.25
0.50
Spearman's ρ
KMT2C
EHMT1
PRDM16
KMT5B
SMYD2
SMYD3
SETD4
all HMTs

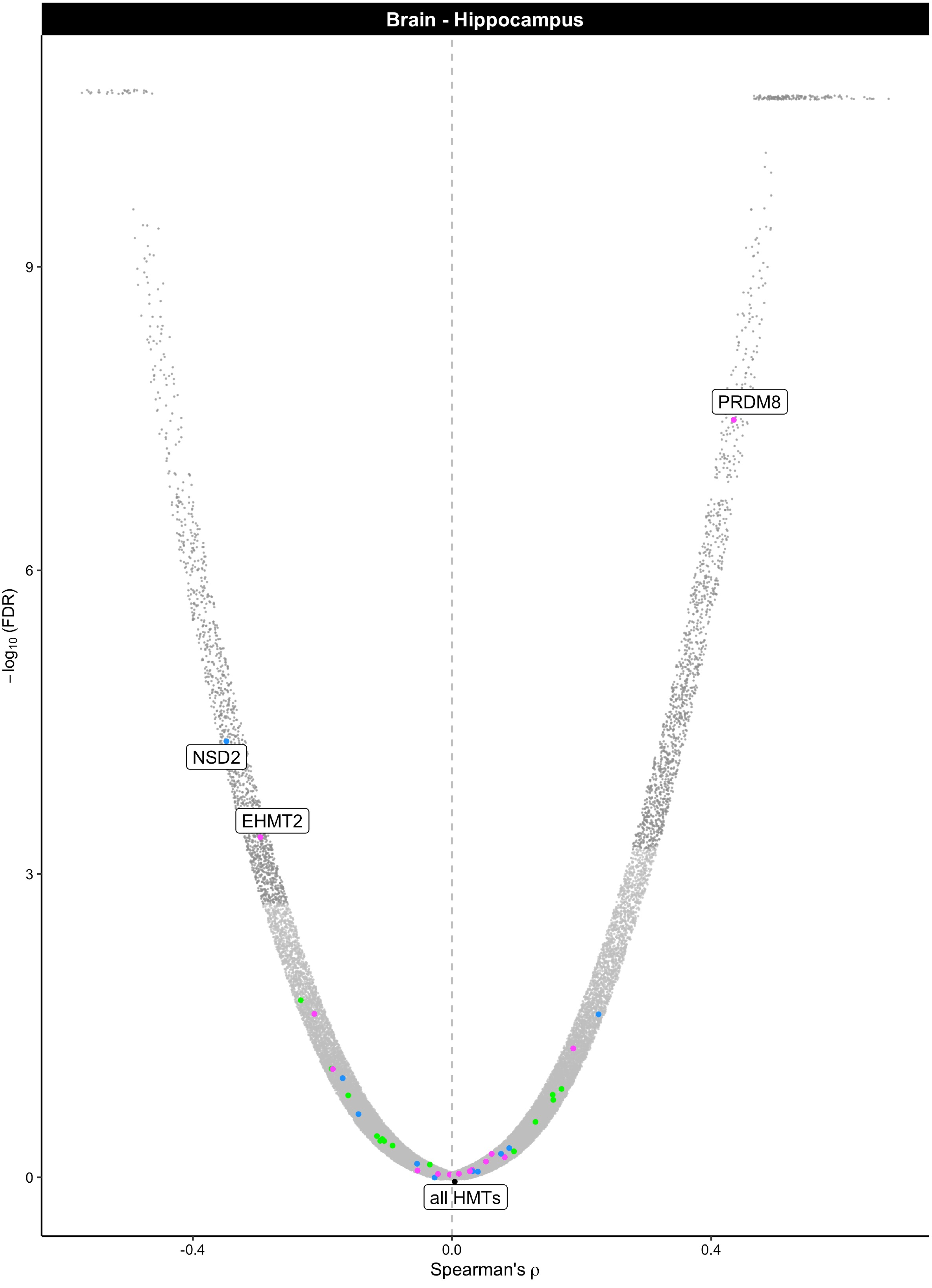

Brain - Hippocampus
PRDM8
NSD2
EHMT2
all HMTs
9
6
3
0
$-\log_{10}$(FDR)
-0.4
0.0
0.4
Spearman's ρ

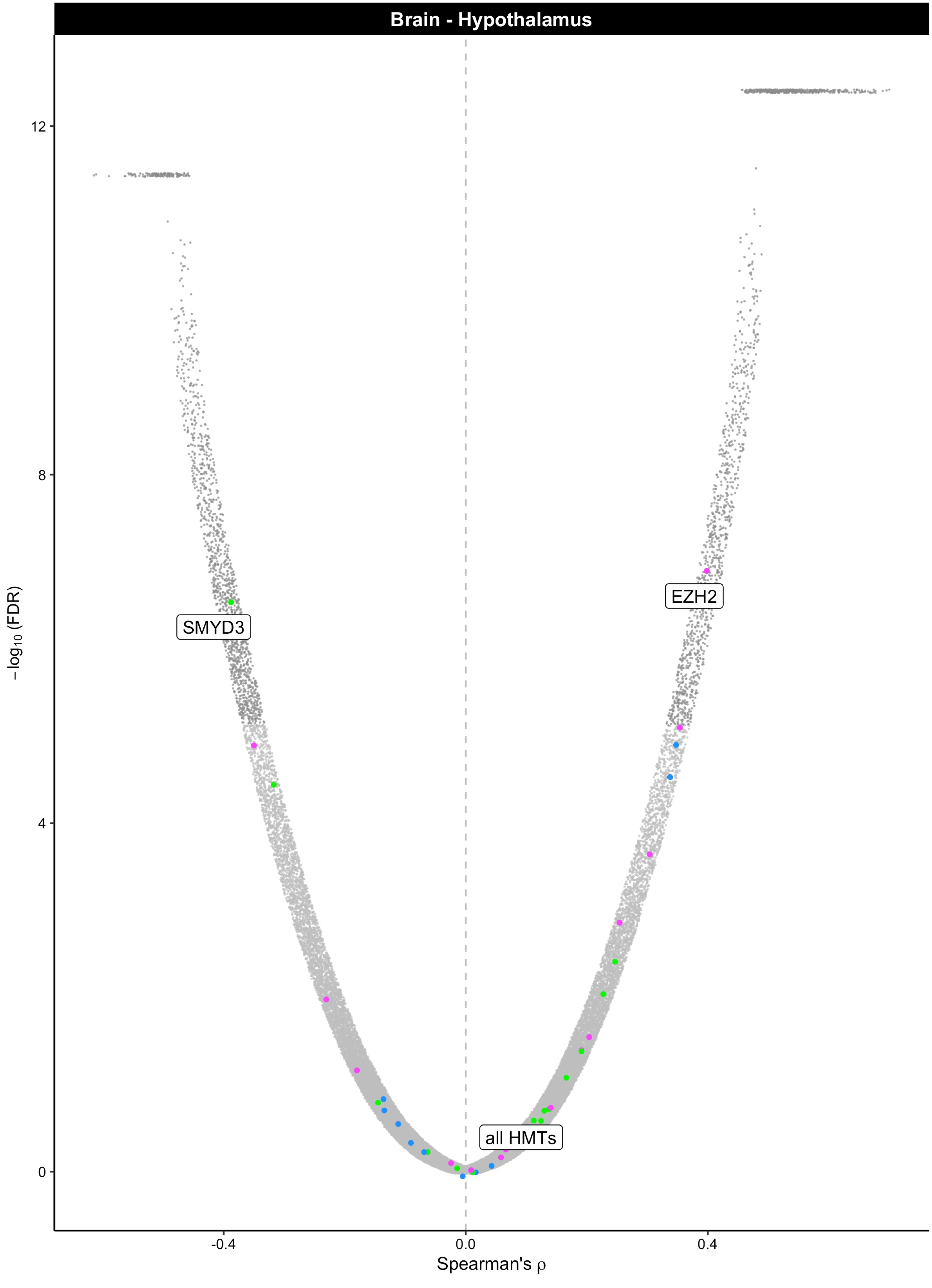

Brain - Hypothalamus
12
8
4
0
−log10 (FDR)
SMYD3
EZH2
all HMTs
-0.4
0.0
0.4
Spearman's ρ

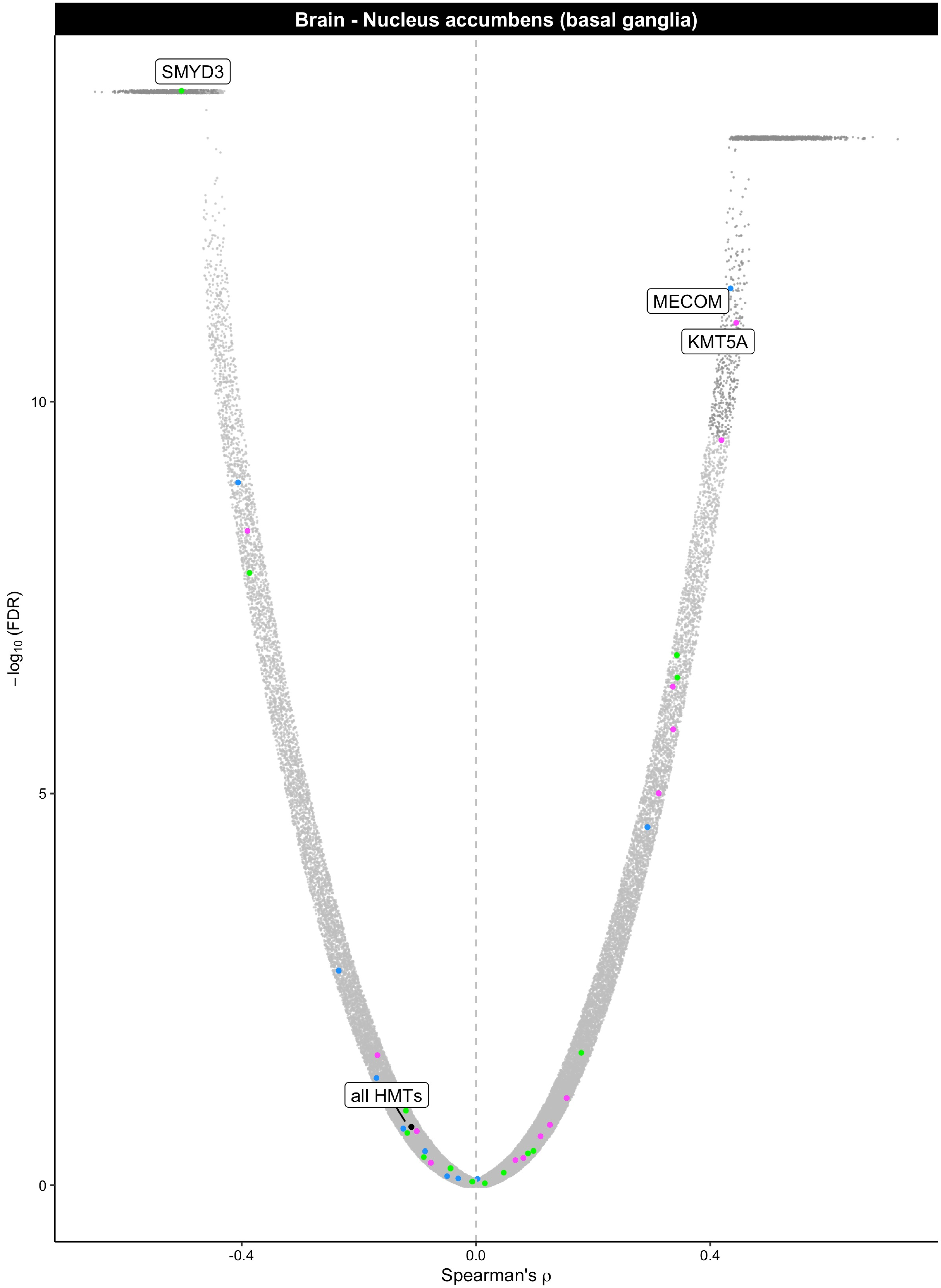
Brain - Nucleus accumbens (basal ganglia)
SMYD3
MECOM
KMT5A
all HMTs
10
5
0
−log10 (FDR)
-0.4
0.0
0.4
Spearman's ρ

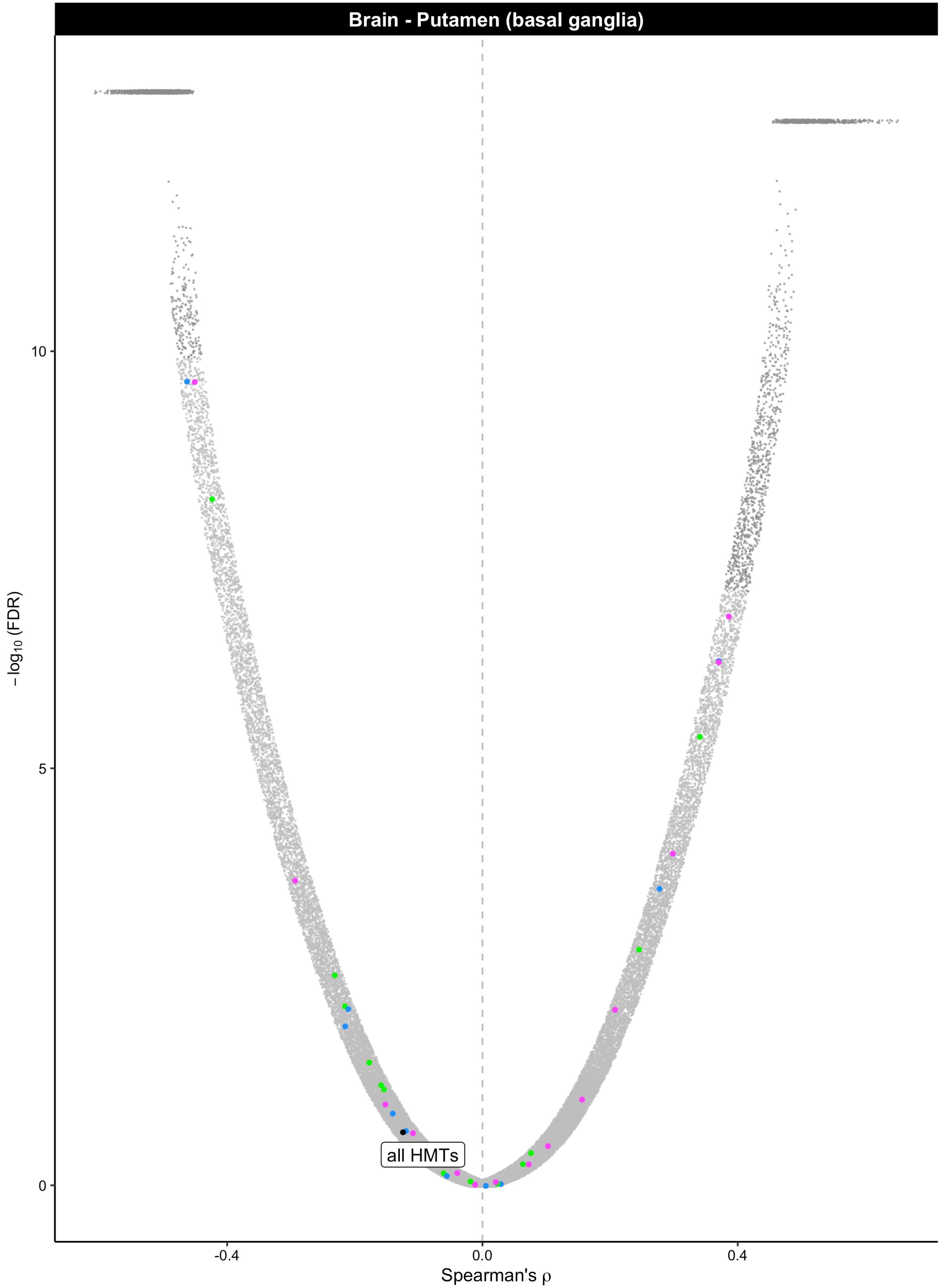

Brain - Putamen (basal ganglia)
all HMTs
$-\log_{10}$ (FDR)
10
5
0
-0.4
0.0
0.4
Spearman's ρ

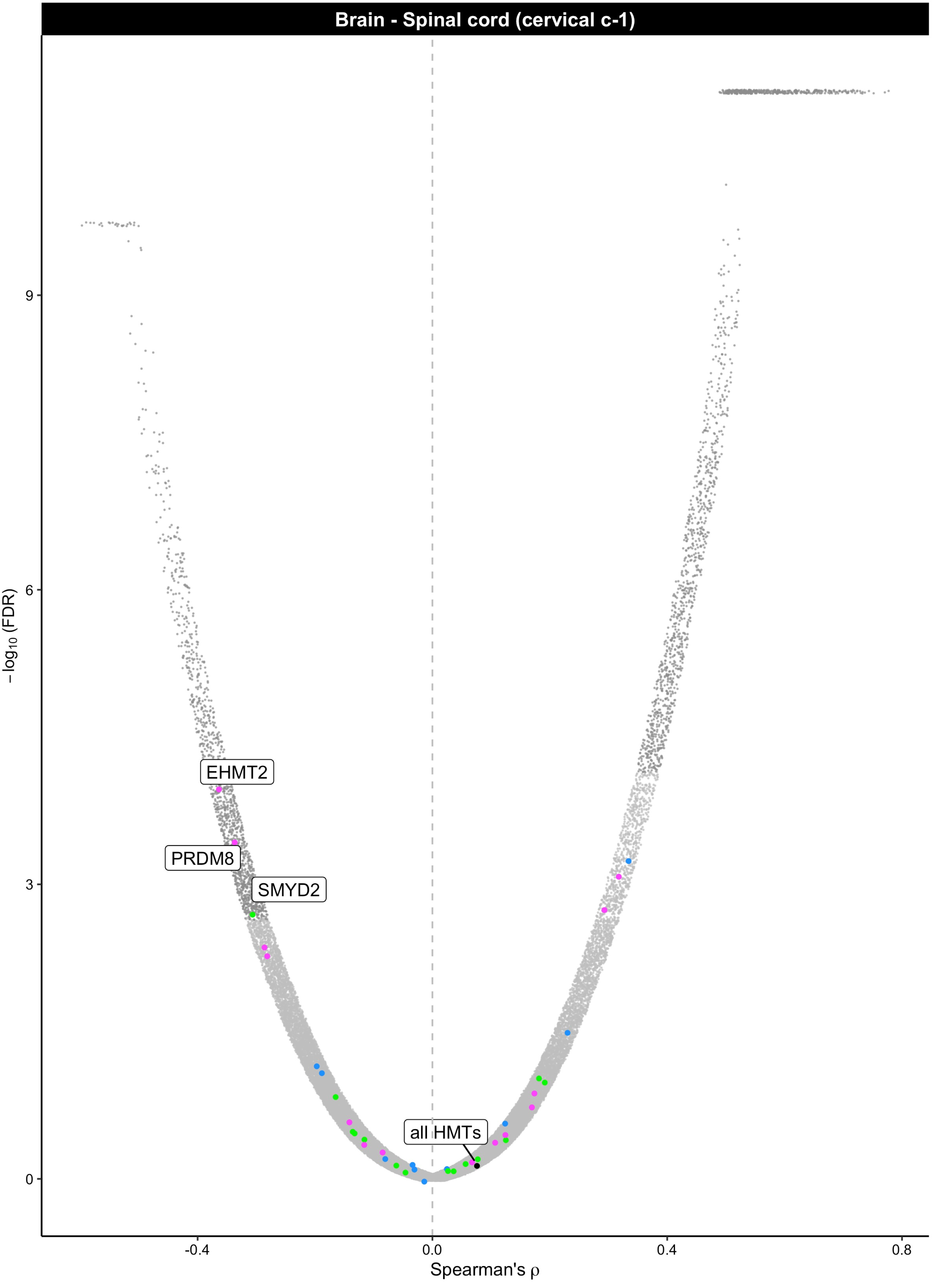

Brain - Spinal cord (cervical c-1)
EHMT2
PRDM8
SMYD2
all HMTs
9
6
3
0
-0.4
0.0
0.4
0.8
Spearman's ρ
−log10 (FDR)

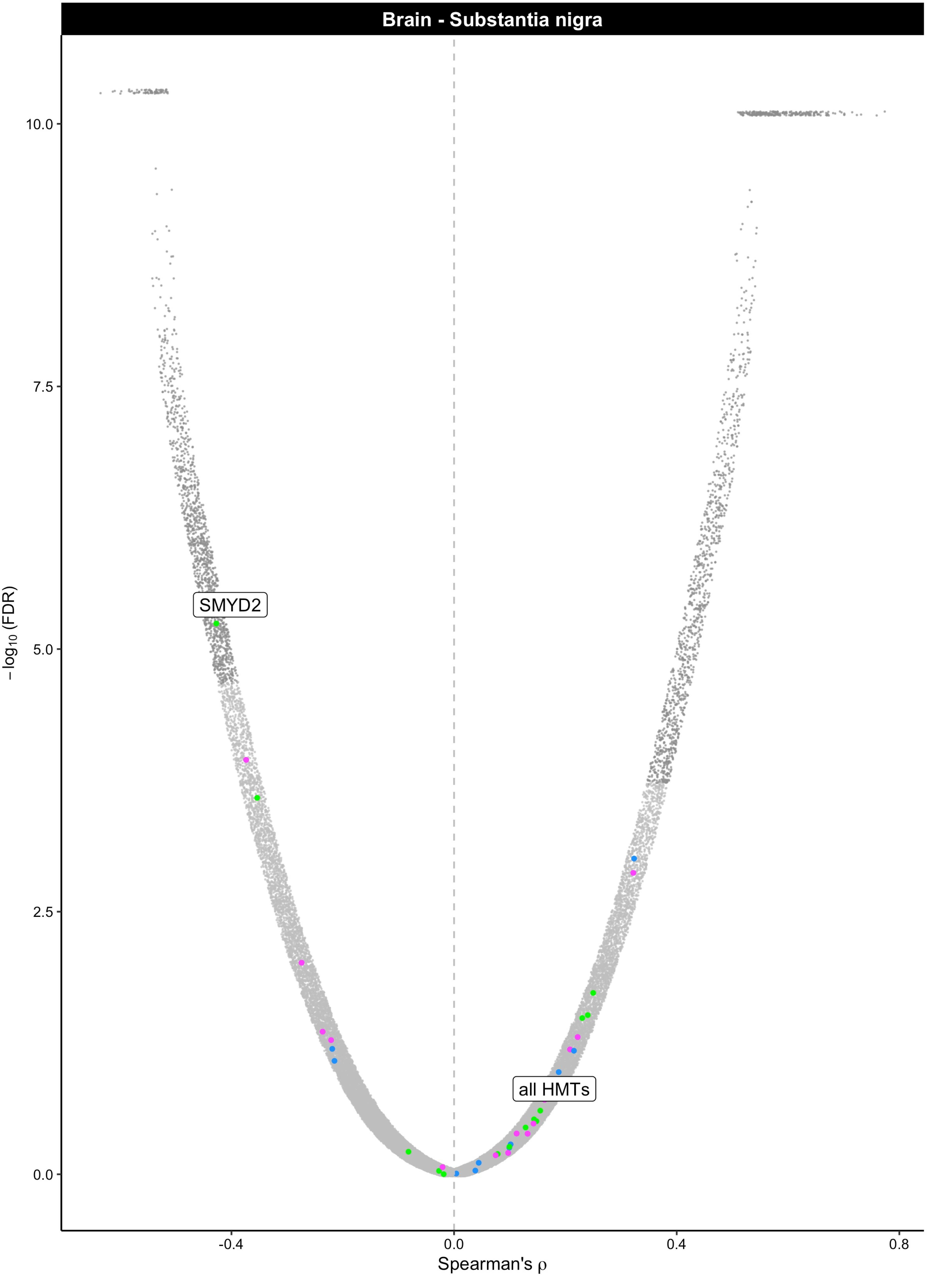

Brain - Substantia nigra
SMYD2
all HMTs
10.0
7.5
5.0
2.5
0.0
−log10 (FDR)
-0.4
0.0
0.4
0.8
Spearman's ρ

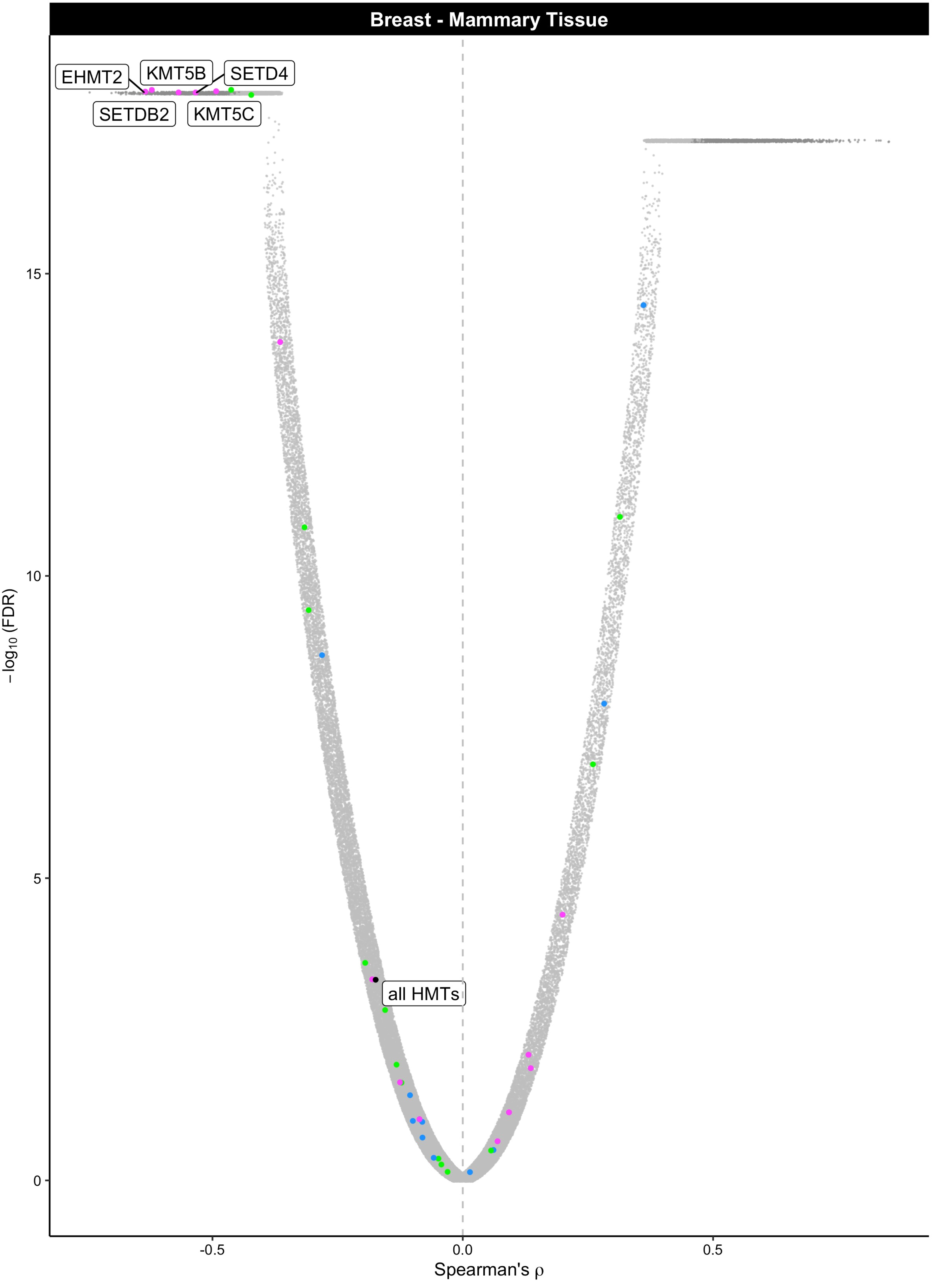

Breast - Mammary Tissue
EHMT2
KMT5B
SETD4
SETDB2
KMT5C
all HMTs
15
10
5
0
−log10 (FDR)
-0.5
0.0
0.5
Spearman's ρ

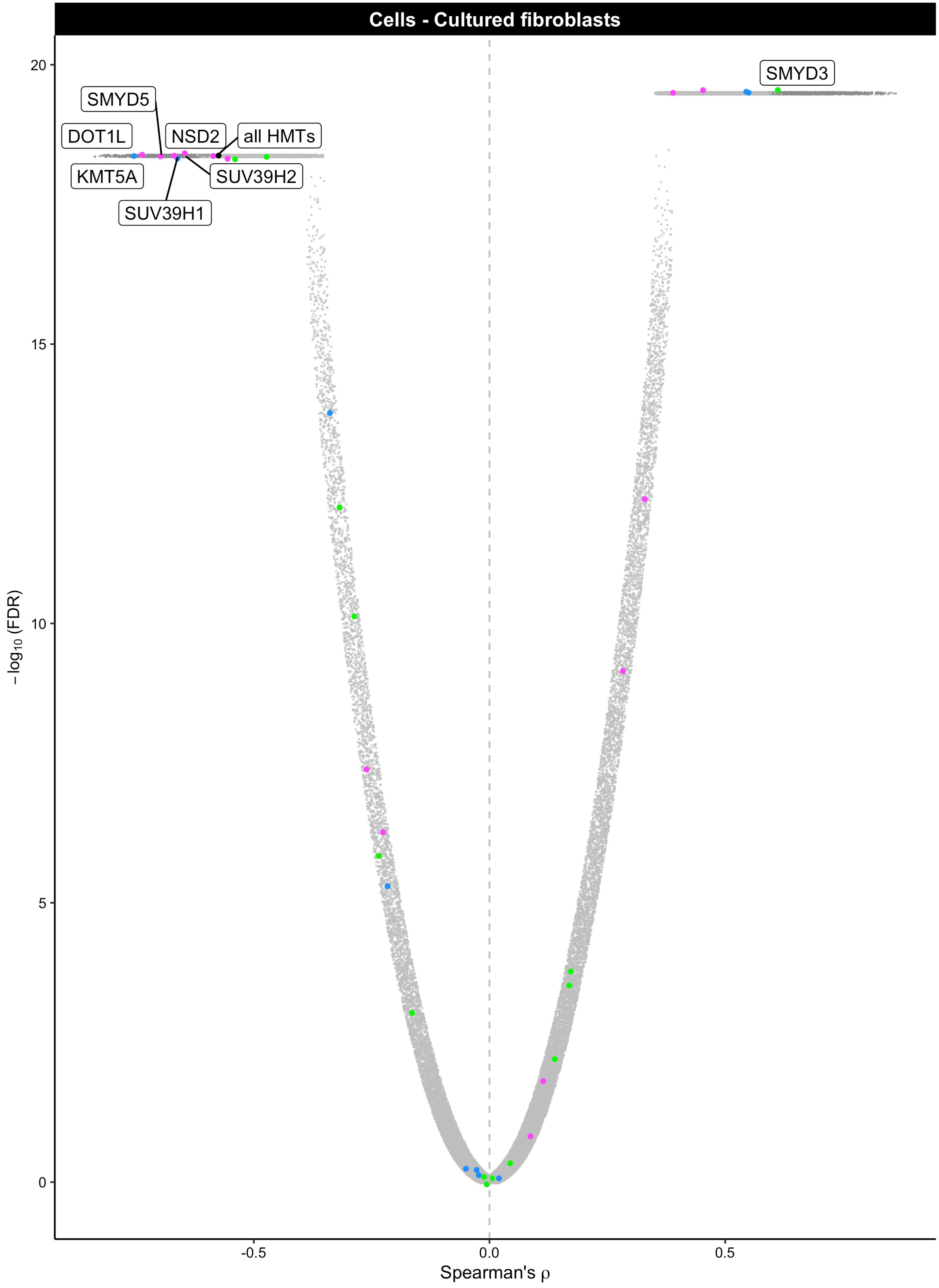

Cells - Cultured fibroblasts
SMYD3
SMYD5
DOT1L
NSD2
all HMTs
KMT5A
SUV39H2
SUV39H1
−log10 (FDR)
20
15
10
5
0
-0.5
0.0
0.5
Spearman's ρ

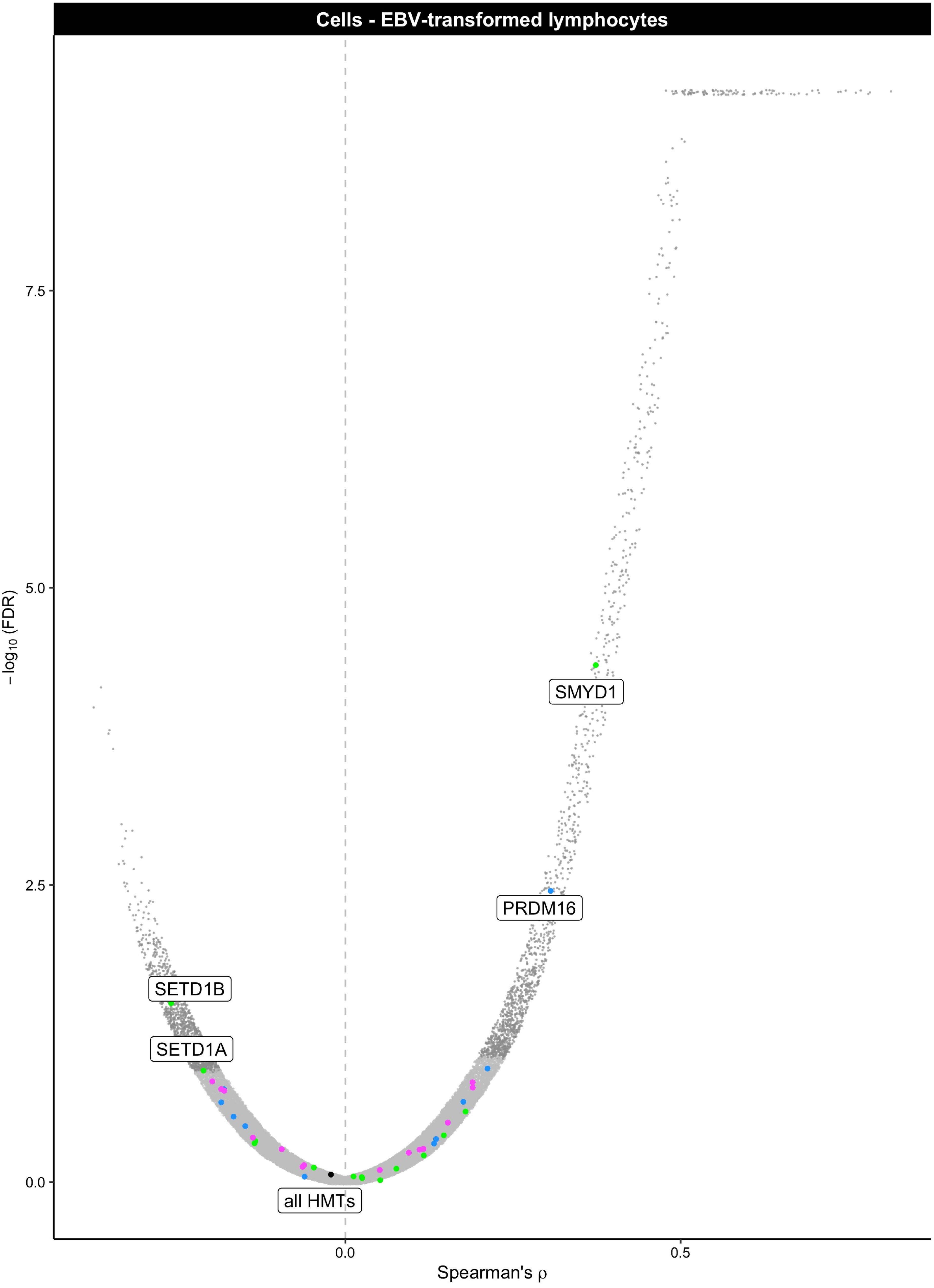
Cells - EBV-transformed lymphocytes
SMYD1
PRDM16
SETD1B
SETD1A
all HMTs
−log10 (FDR)
7.5
5.0
2.5
0.0
0.0
0.5
Spearman's ρ

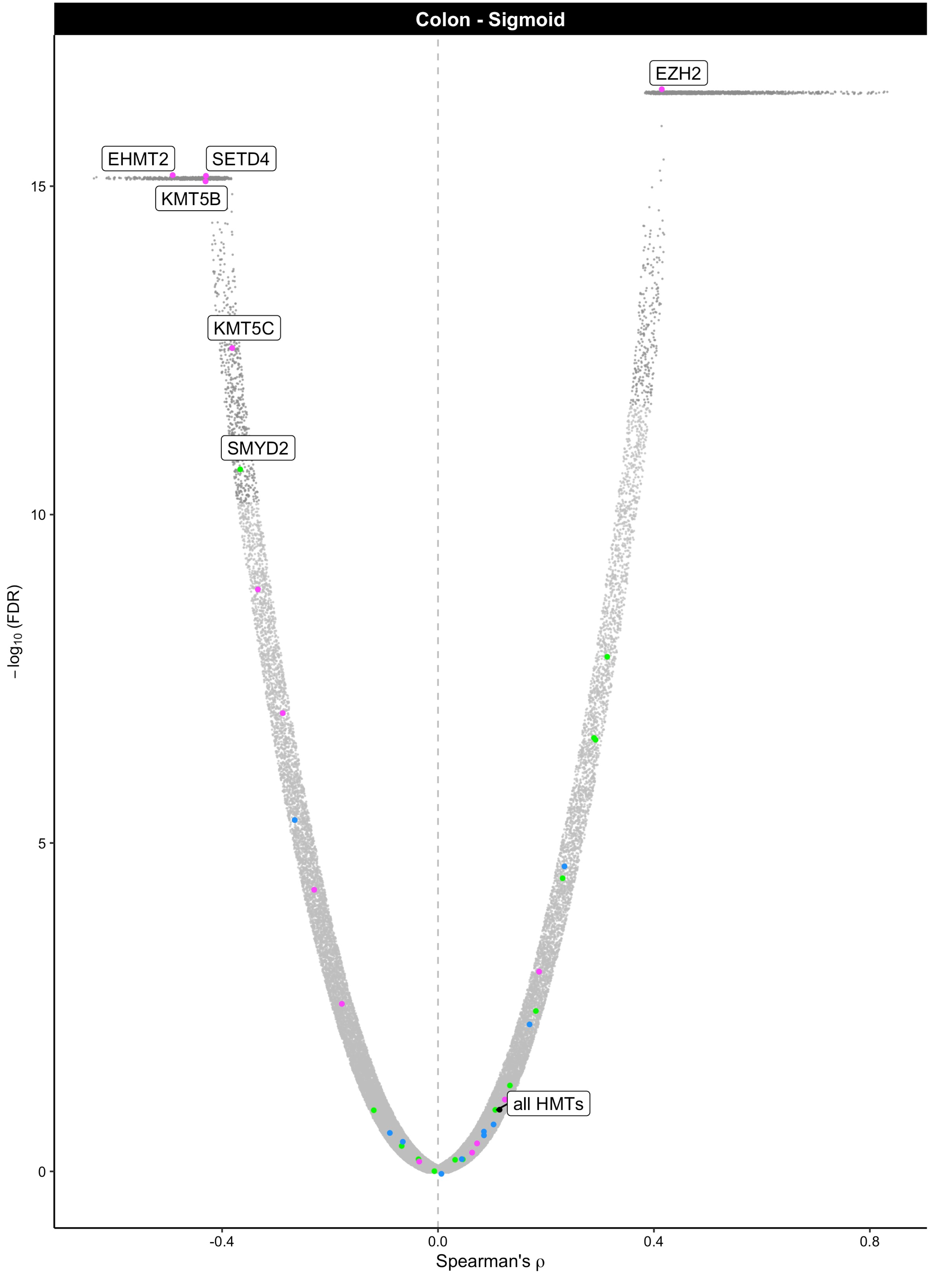

Colon - Sigmoid
EZH2
EHMT2
SETD4
KMT5B
KMT5C
SMYD2
all HMTs
−log10 (FDR)
15
10
5
0
−0.4
0.0
0.4
0.8
Spearman's ρ

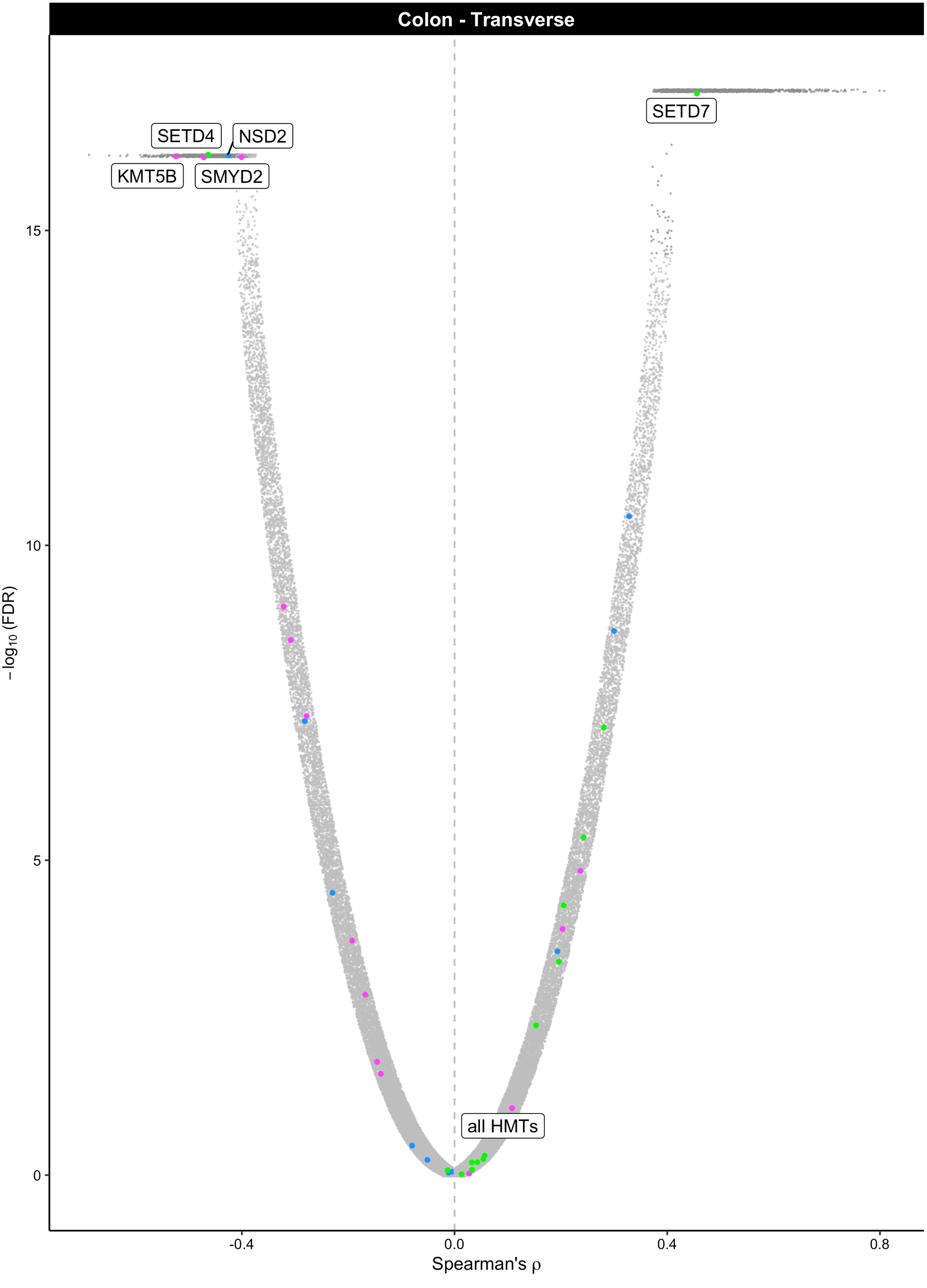

Colon - Transverse
SETD7
SETD4
NSD2
KMT5B
SMYD2
all HMTs
15
10
5
0
-log10 (FDR)
-0.4
0.0
0.4
0.8
Spearman's ρ

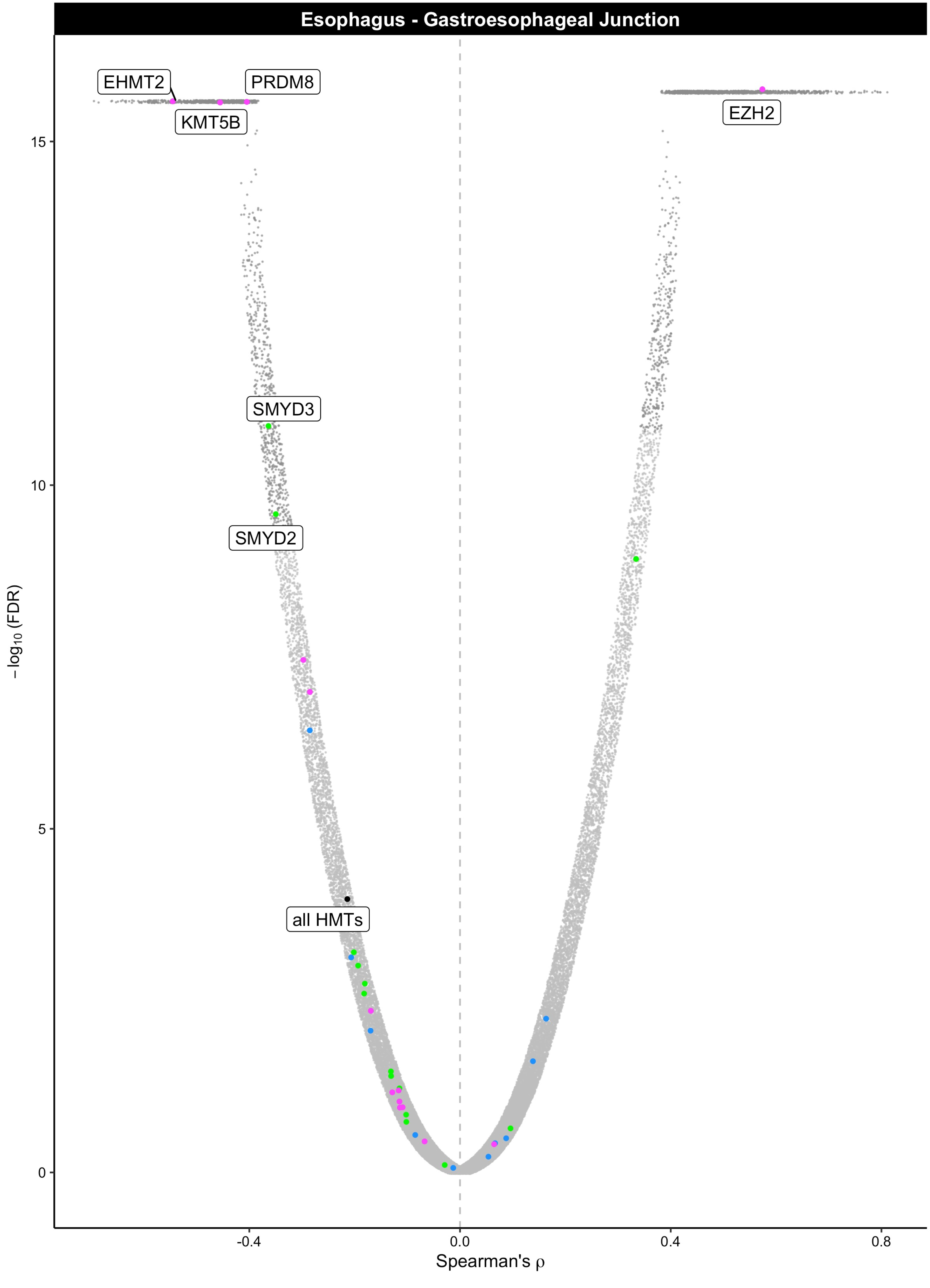

Esophagus - Gastroesophageal Junction
EHMT2
PRDM8
KMT5B
EZH2
SMYD3
SMYD2
all HMTs
15
10
5
0
−log10 (FDR)
-0.4
0.0
0.4
0.8
Spearman's ρ

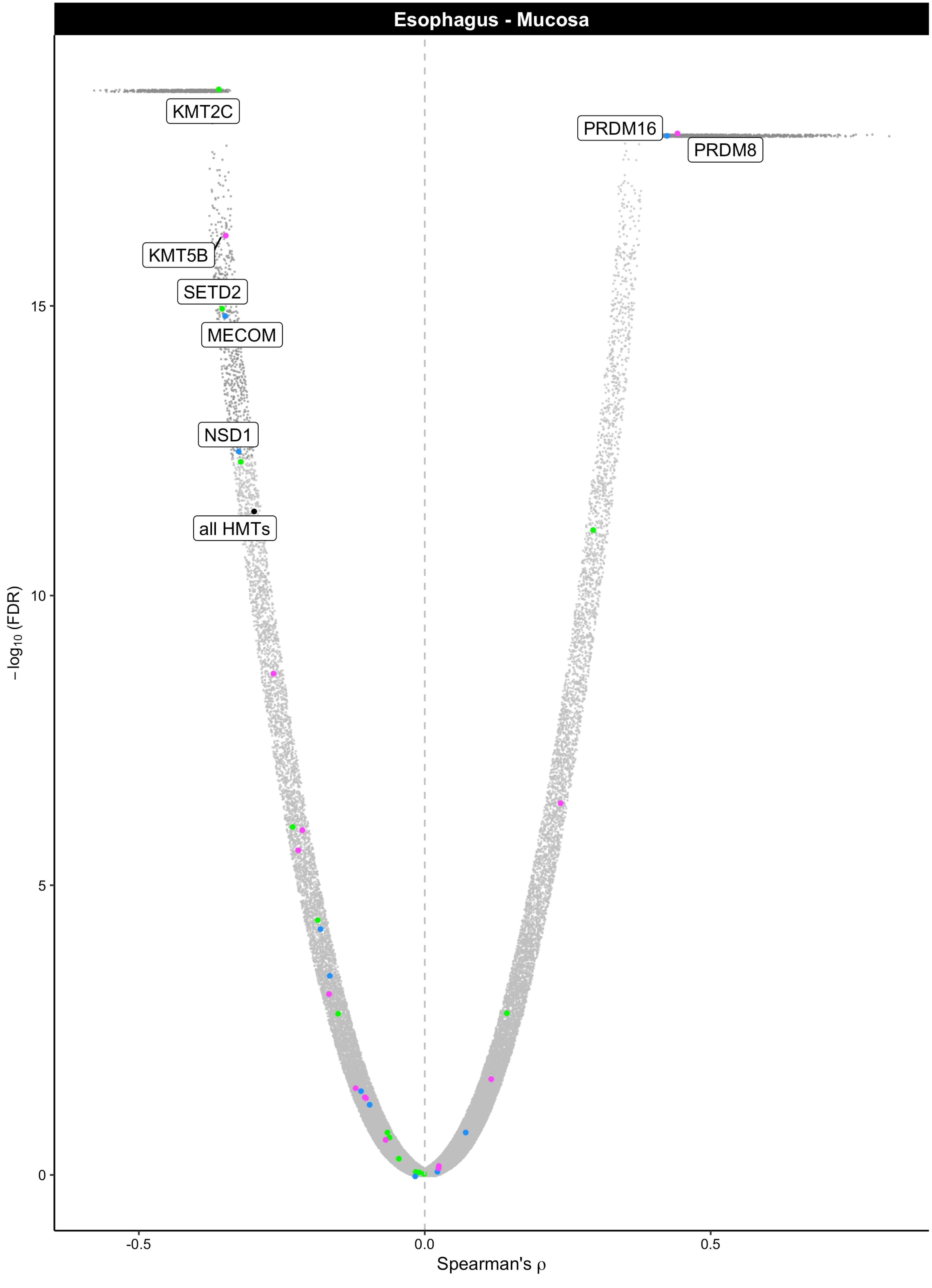

Esophagus - Mucosa
KMT2C
PRDM16
PRDM8
KMT5B
SETD2
MECOM
NSD1
all HMTs
15
10
5
0
−log10 (FDR)
-0.5
0.0
0.5
Spearman's ρ

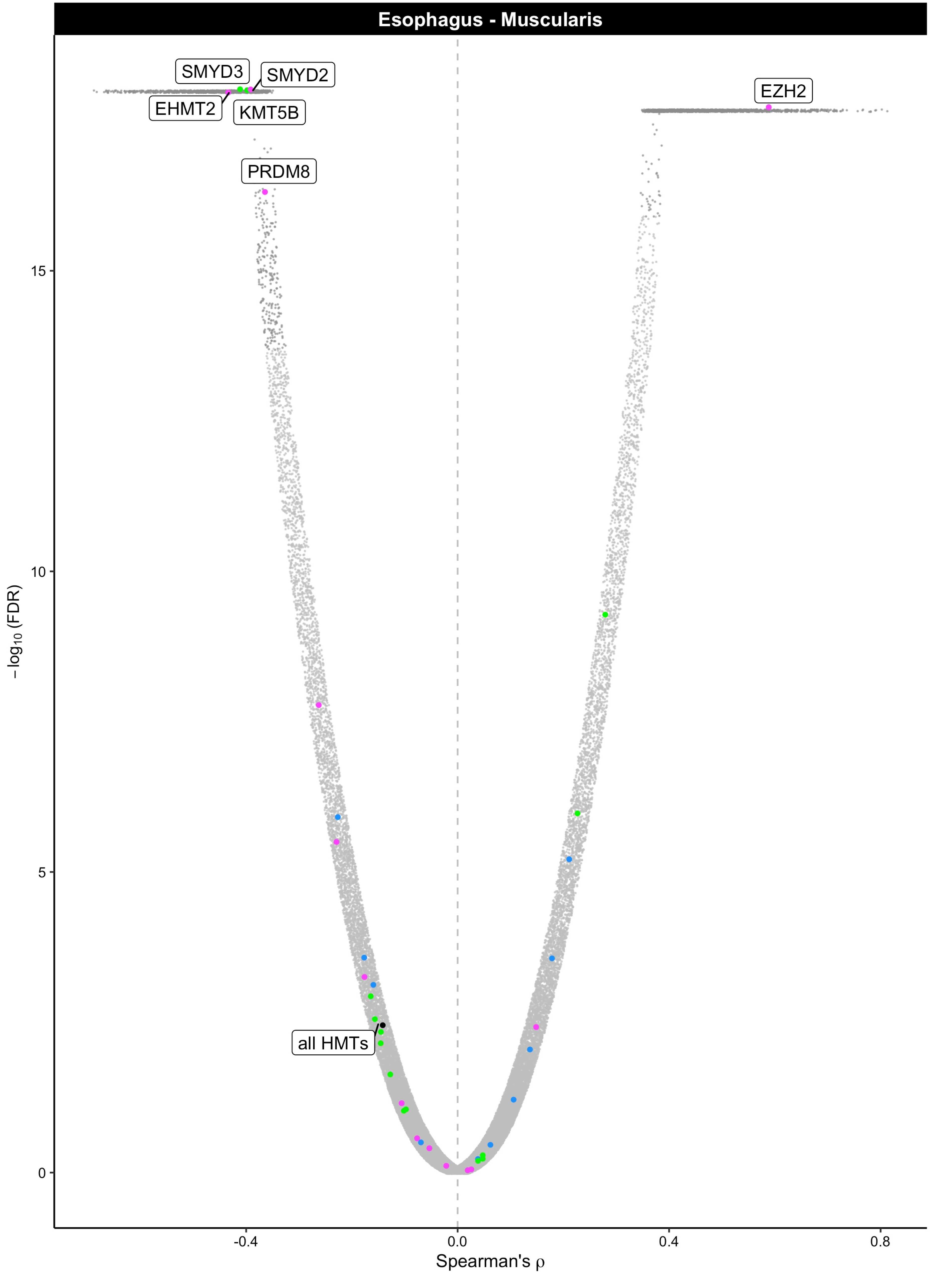

Esophagus - Muscularis
SMYD3
SMYD2
EHMT2
KMT5B
EZH2
PRDM8
all HMTs
−log10 (FDR)
15
10
5
0
-0.4
0.0
0.4
0.8
Spearman's ρ

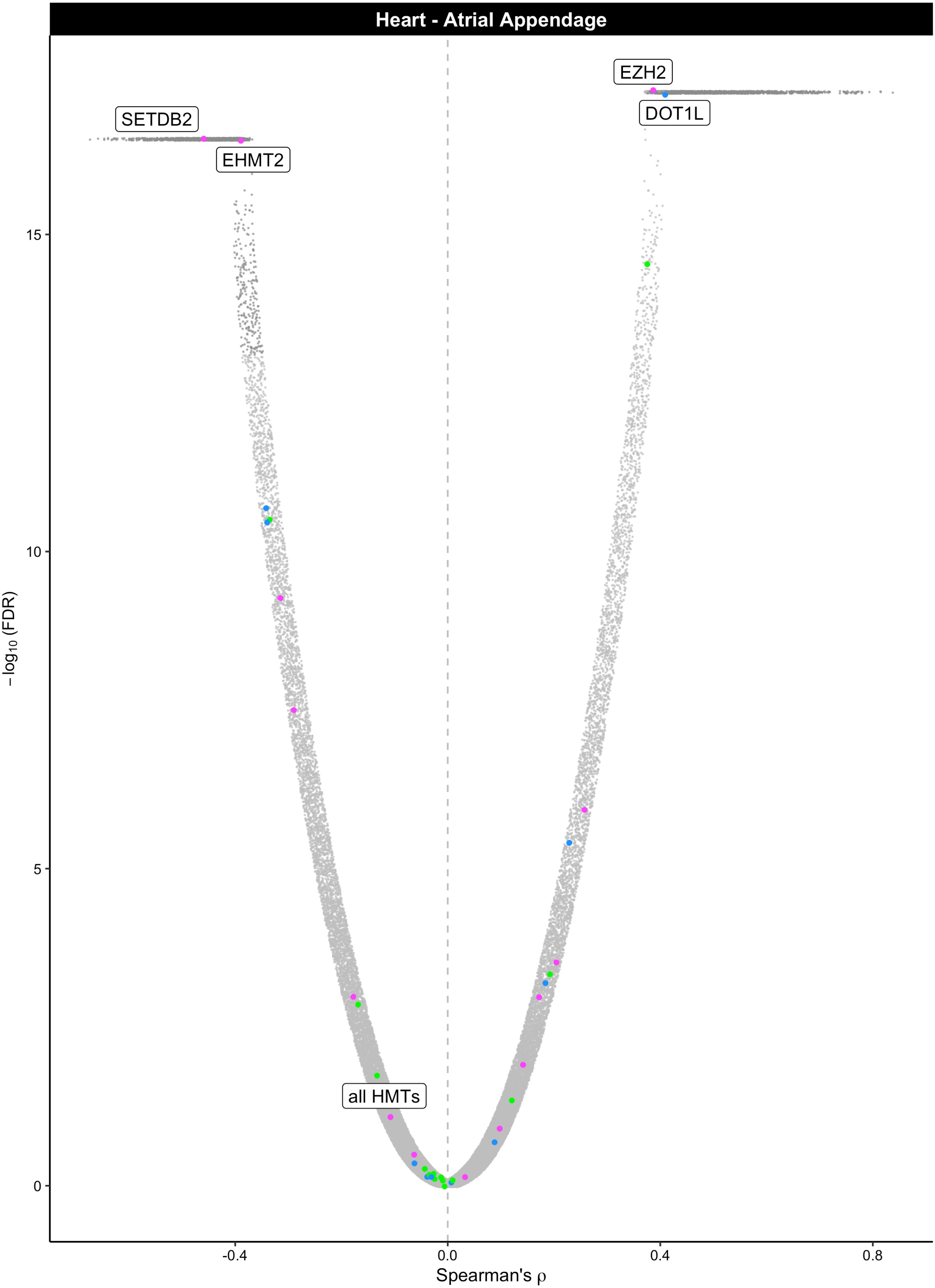

Heart - Atrial Appendage
EZH2
DOT1L
SETDB2
EHMT2
all HMTs
15
10
5
0
$-\log_{10}$(FDR)
-0.4
0.0
0.4
0.8
Spearman's ρ

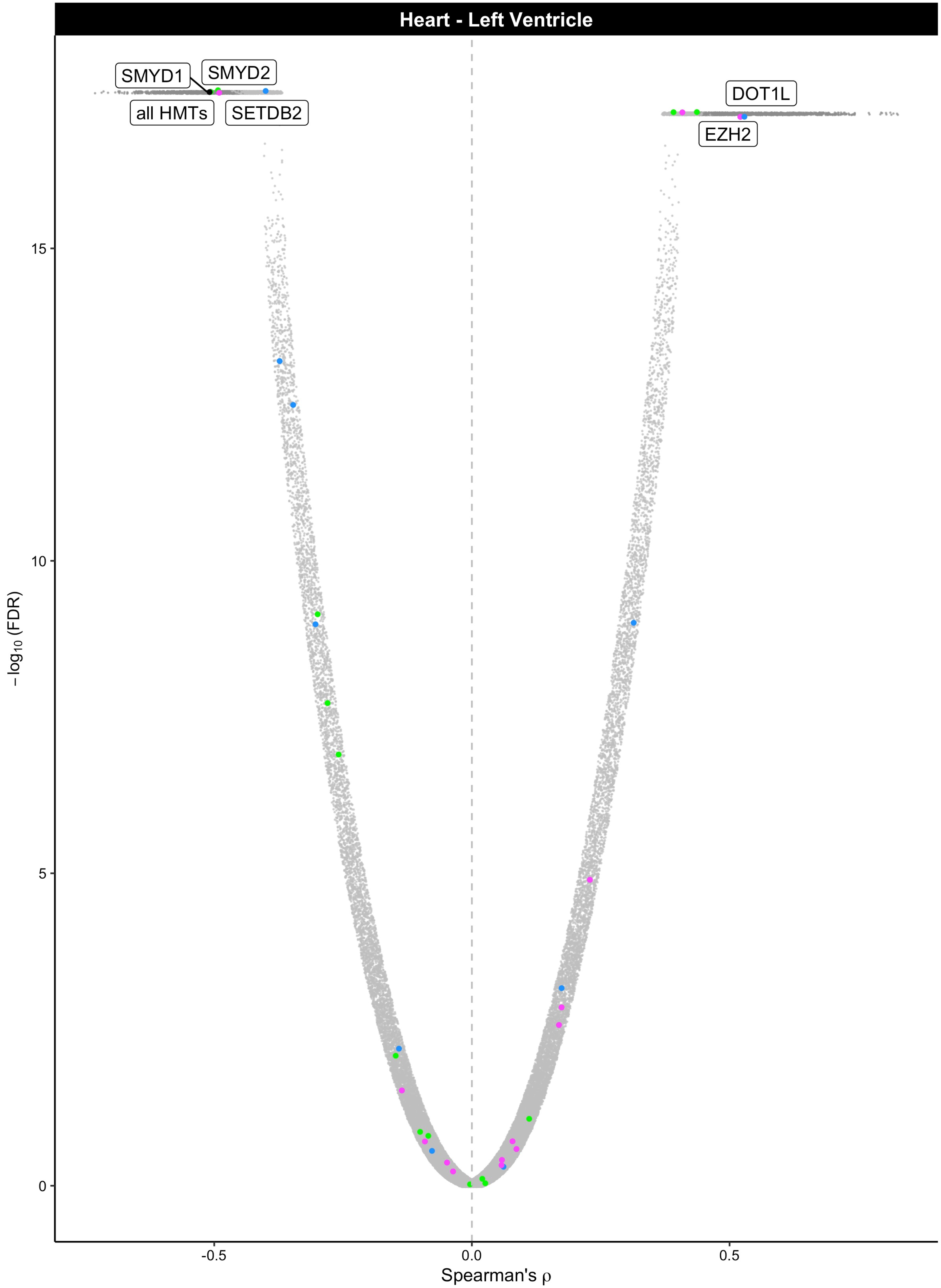
Heart - Left Ventricle
SMYD1
SMYD2
all HMTs
SETDB2
DOT1L
EZH2
15
10
5
0
−log10 (FDR)
-0.5
0.0
0.5
Spearman's ρ

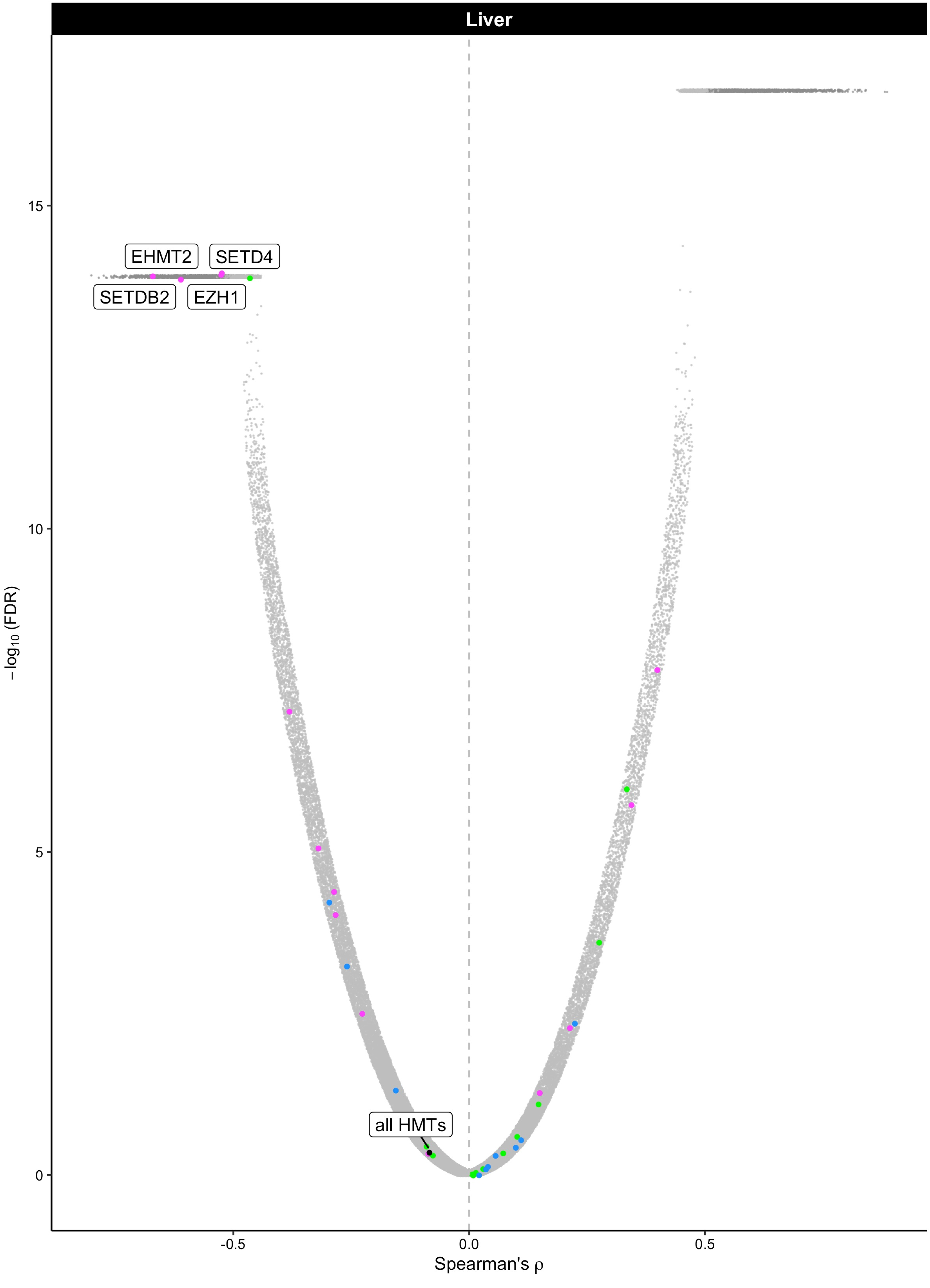

Liver
EHMT2
SETD4
SETDB2
EZH1
all HMTs
15
10
5
0
$-\log_{10}$ (FDR)
-0.5
0.0
0.5
Spearman's ρ

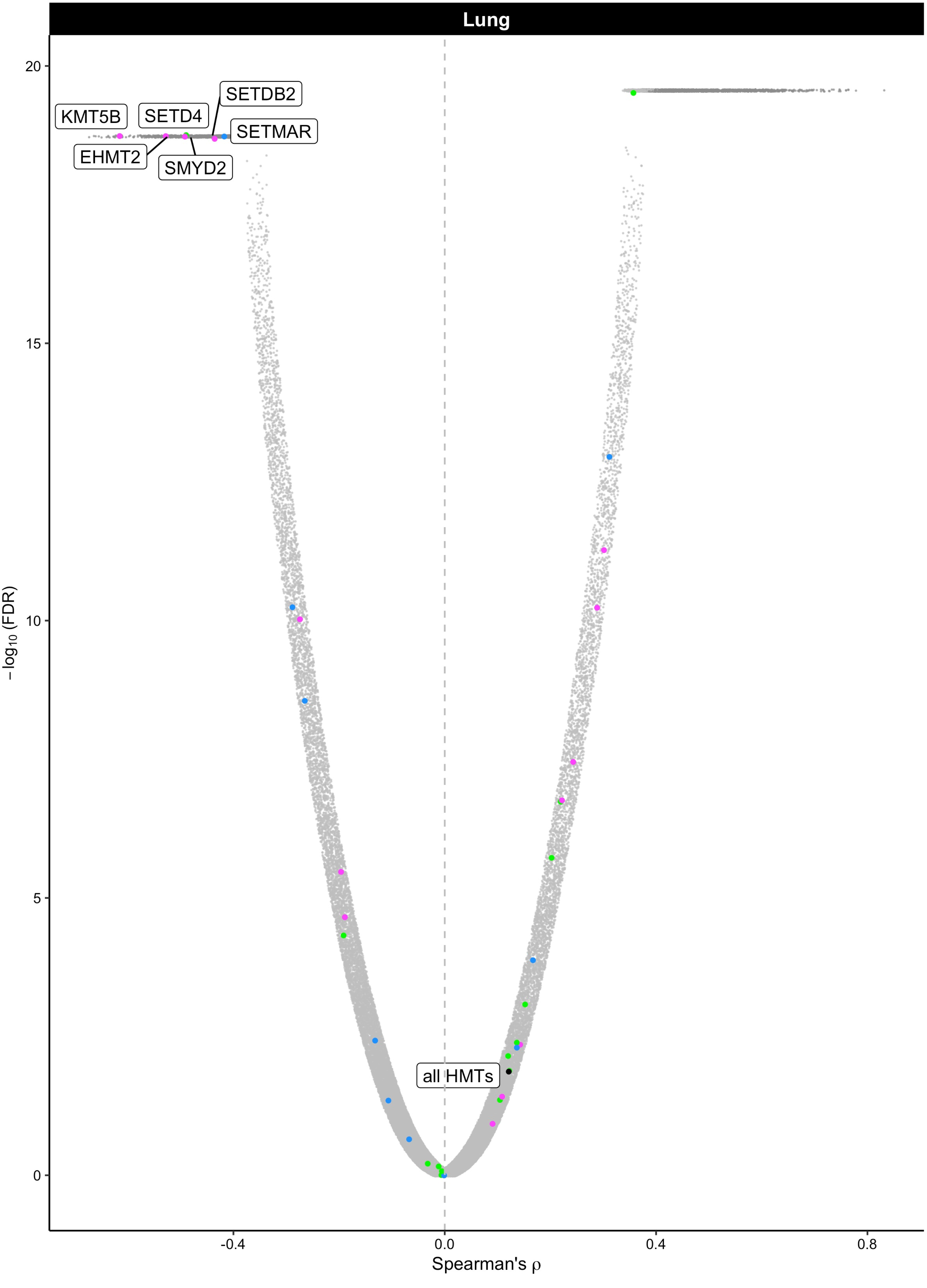

Lung
KMT5B
SETD4
SETDB2
SETMAR
EHMT2
SMYD2
all HMTs
20
15
10
5
0
−log10 (FDR)
-0.4
0.0
0.4
0.8
Spearman's ρ

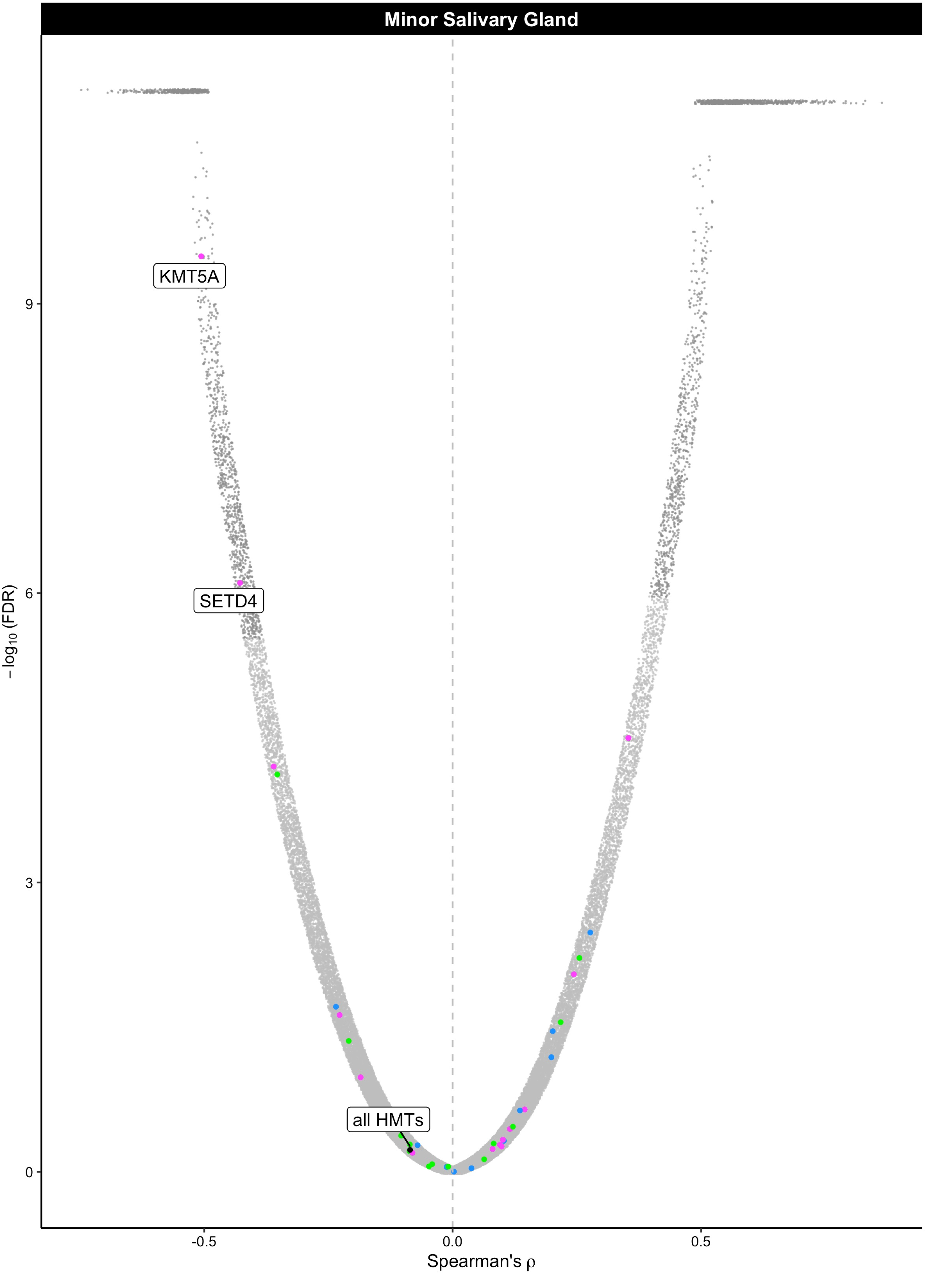
Minor Salivary Gland
KMT5A
SETD4
all HMTs
9
6
3
0
−log10 (FDR)
-0.5
0.0
0.5
Spearman's ρ

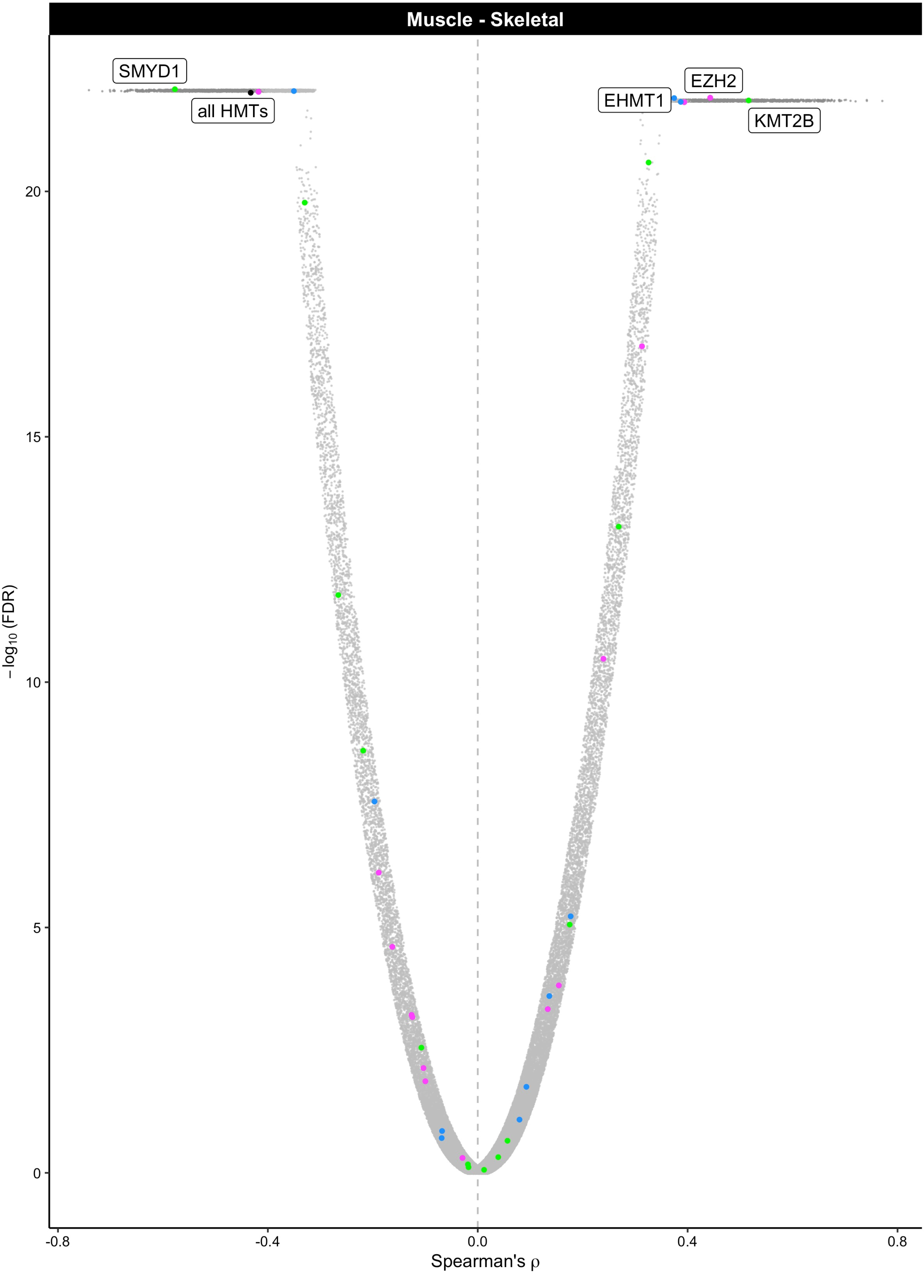

Muscle - Skeletal
SMYD1
all HMTs
EZH2
EHMT1
KMT2B
20
15
10
5
0
−log10 (FDR)
-0.8
-0.4
0.0
0.4
0.8
Spearman's ρ

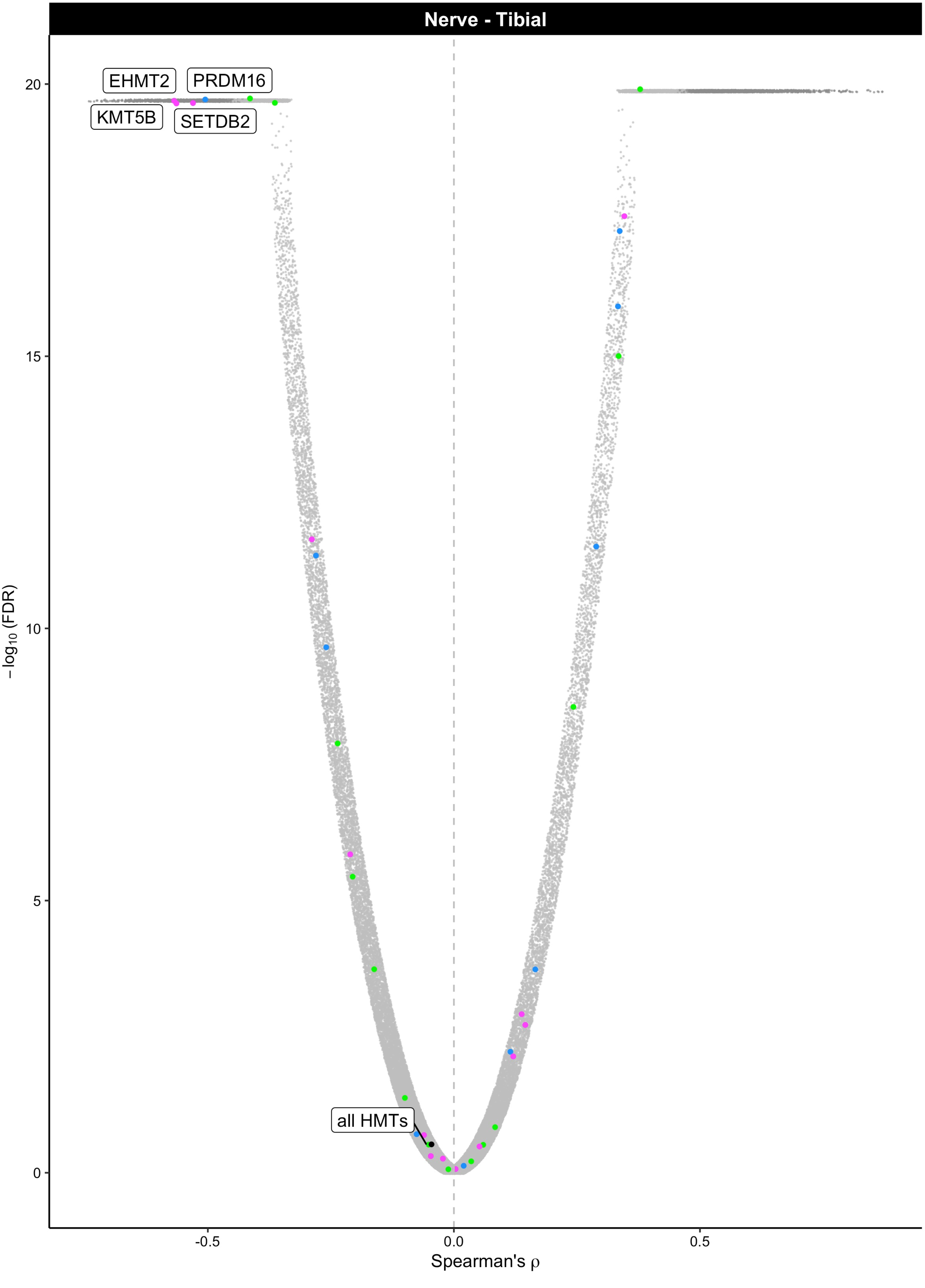

Nerve - Tibial
EHMT2
PRDM16
KMT5B
SETDB2
all HMTs
20
15
10
5
0
−log10 (FDR)
-0.5
0.0
0.5
Spearman's ρ

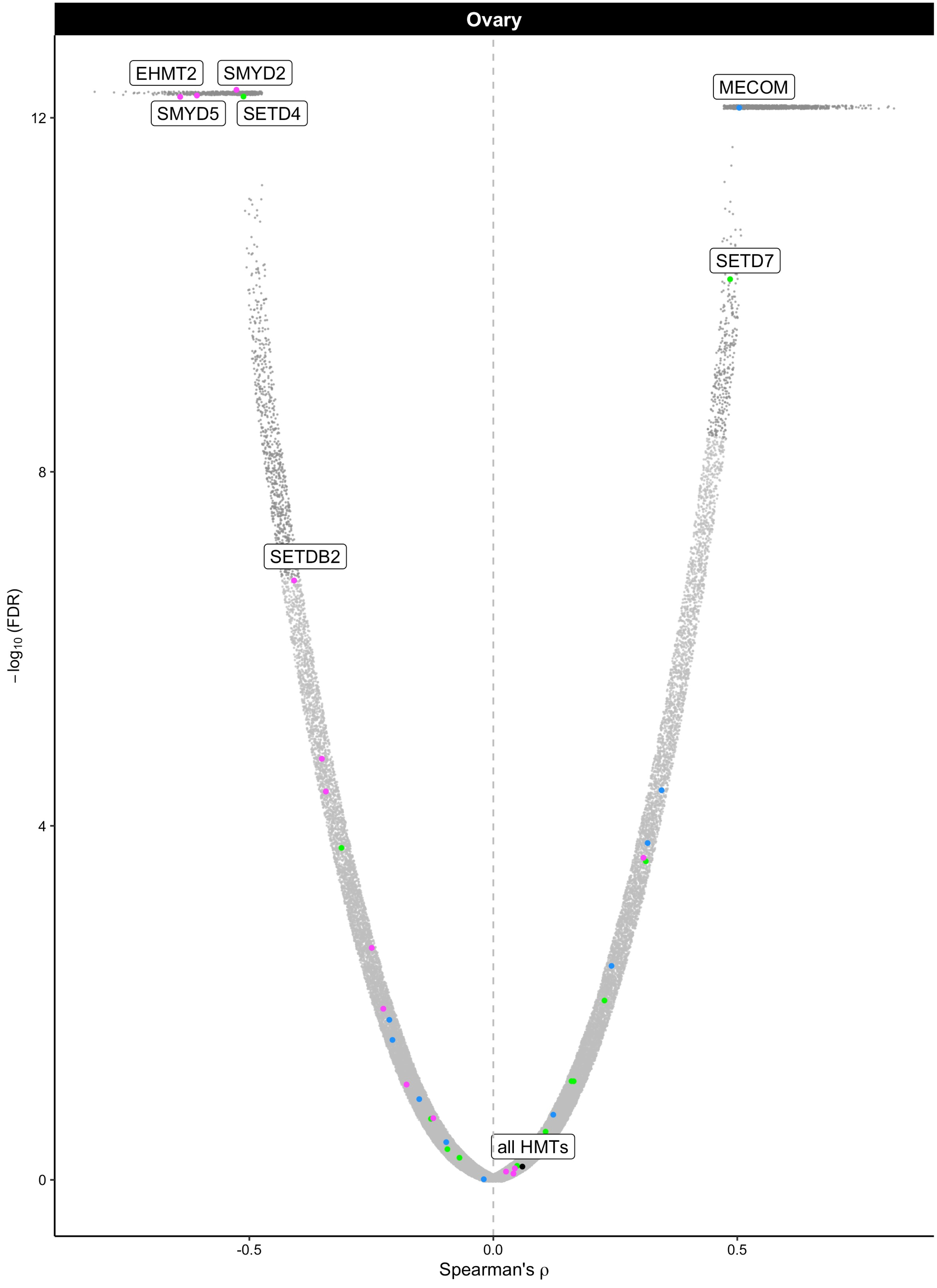

Ovary
EHMT2
SMYD2
SMYD5
SETD4
MECOM
SETD7
SETDB2
all HMTs
12
8
4
0
-0.5
0.0
0.5
$-\log_{10}$ (FDR)
Spearman's ρ

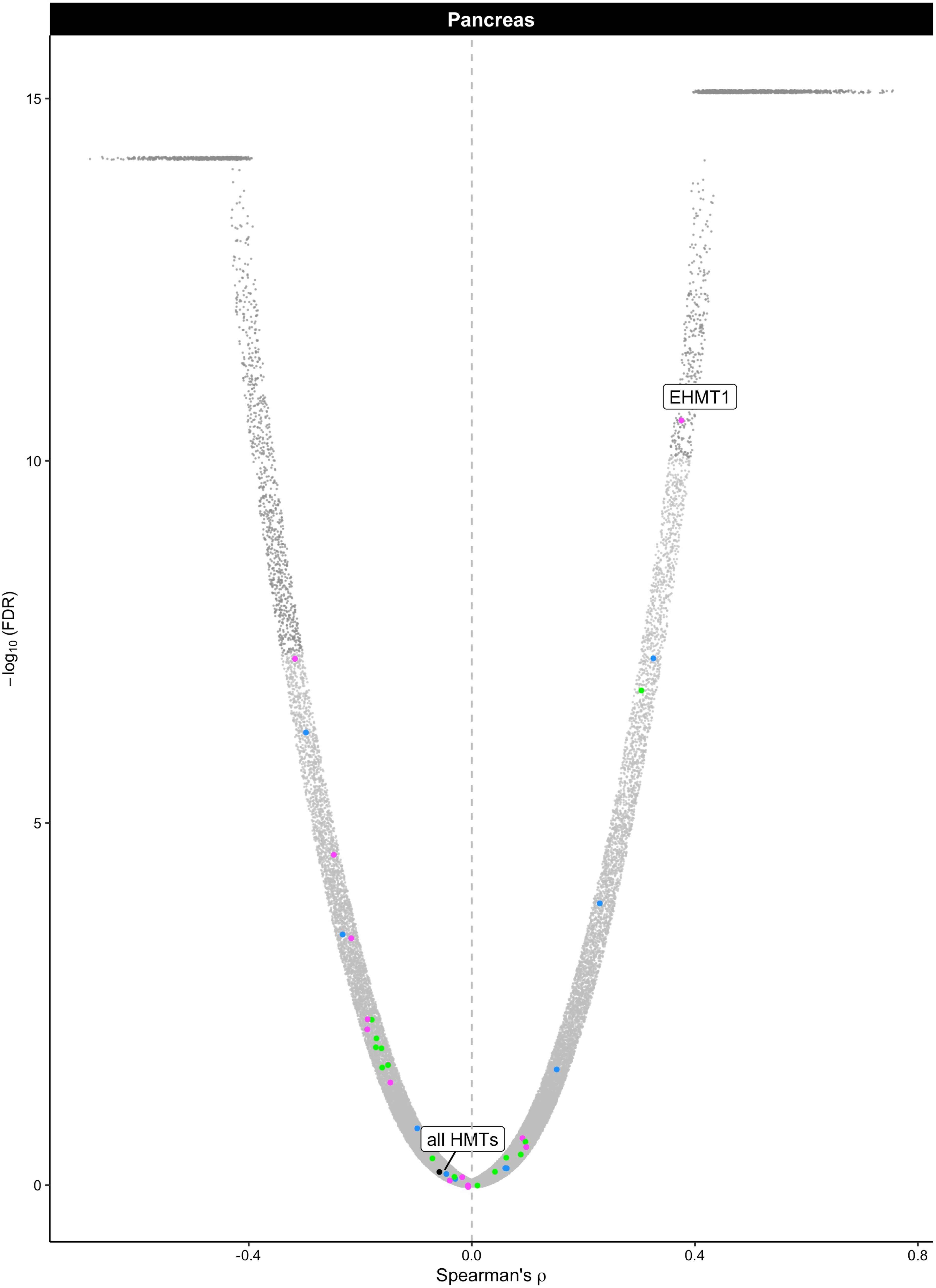

Pancreas
EHMT1
all HMTs
15
10
5
0
−log10 (FDR)
-0.4
0.0
0.4
0.8
Spearman's ρ

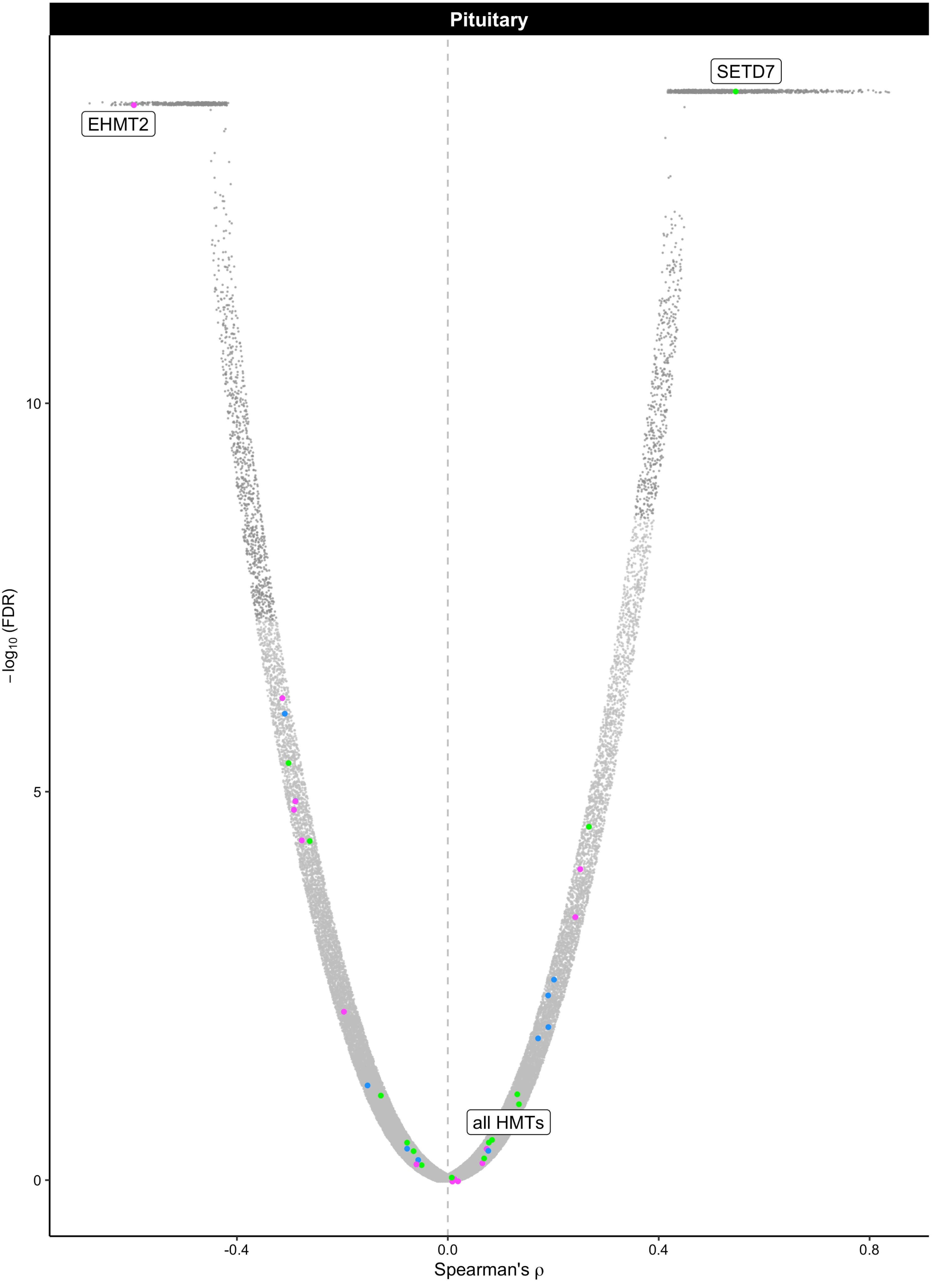

Pituitary
SETD7
EHMT2
all HMTs
10
5
0
−log10 (FDR)
-0.4
0.0
0.4
0.8
Spearman's ρ

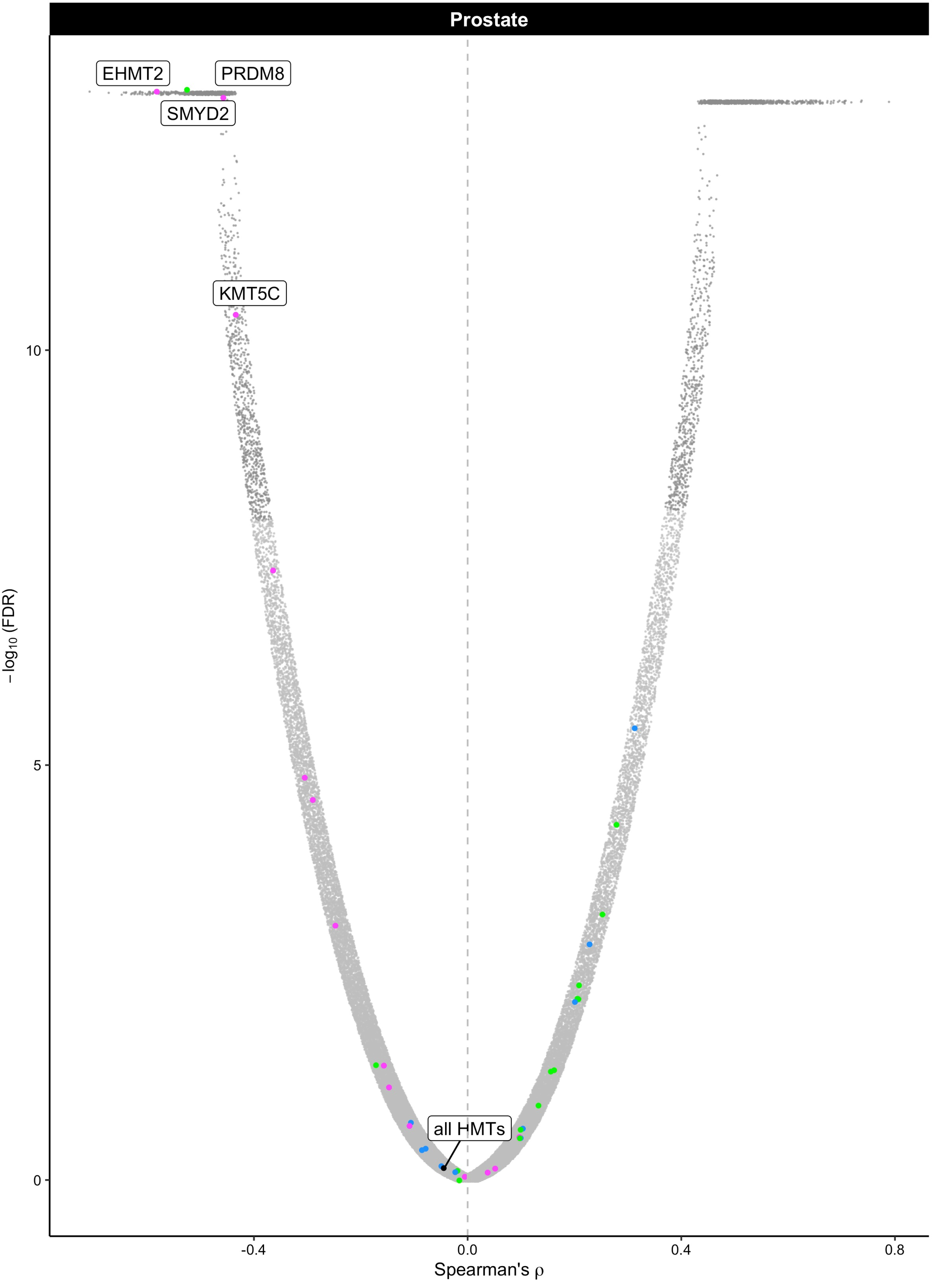

Prostate
EHMT2
PRDM8
SMYD2
KMT5C
all HMTs
10
5
0
−log10 (FDR)
-0.4
0.0
0.4
0.8
Spearman's ρ

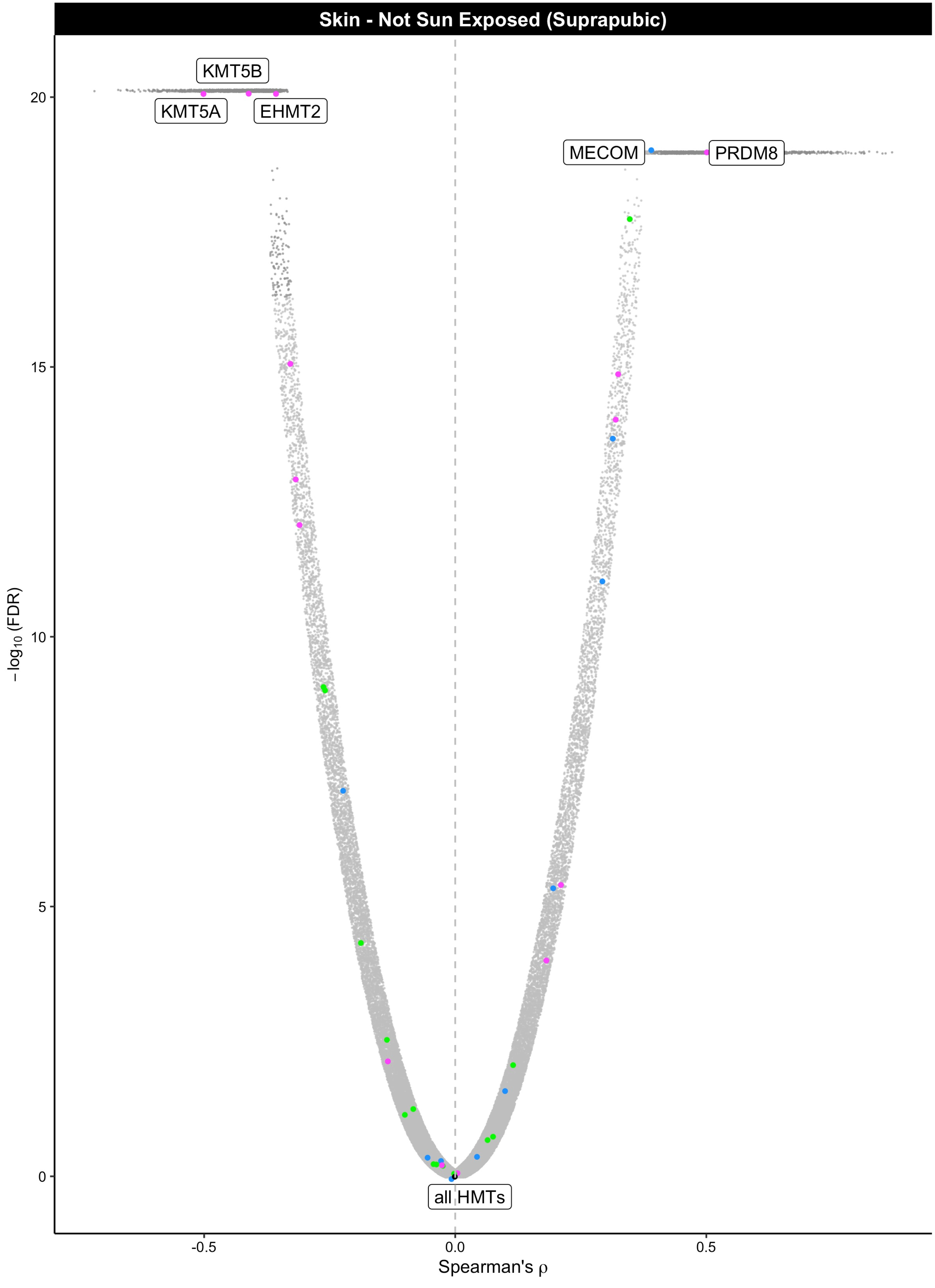

Skin - Not Sun Exposed (Suprapubic)
KMT5B
KMT5A
EHMT2
MECOM
PRDM8
all HMTs
−log10 (FDR)
20
15
10
5
0
-0.5
0.0
0.5
Spearman's ρ

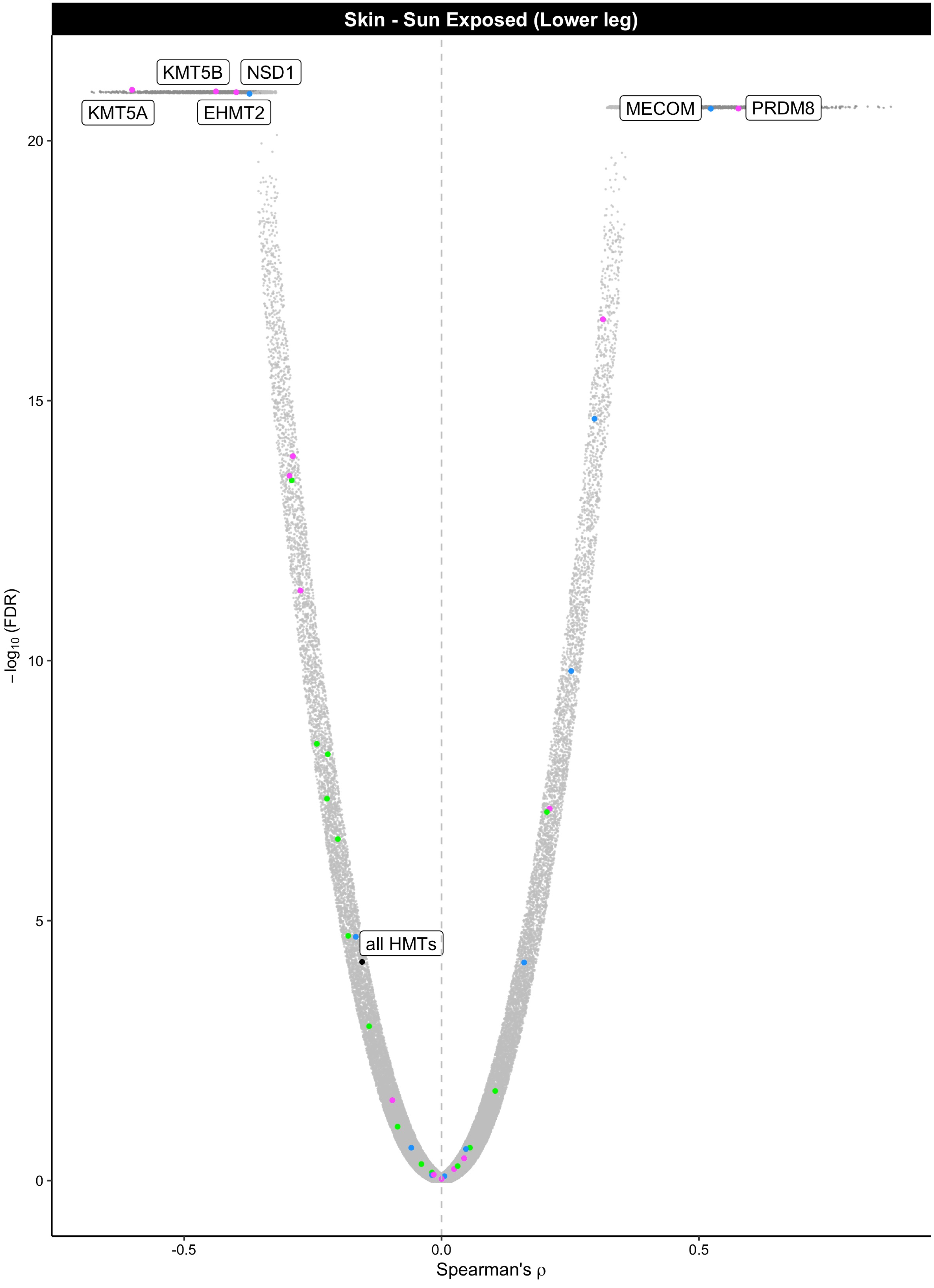

Skin - Sun Exposed (Lower leg)
KMT5B
NSD1
KMT5A
EHMT2
MECOM
PRDM8
all HMTs
20
15
10
5
0
−log10 (FDR)
-0.5
0.0
0.5
Spearman's ρ

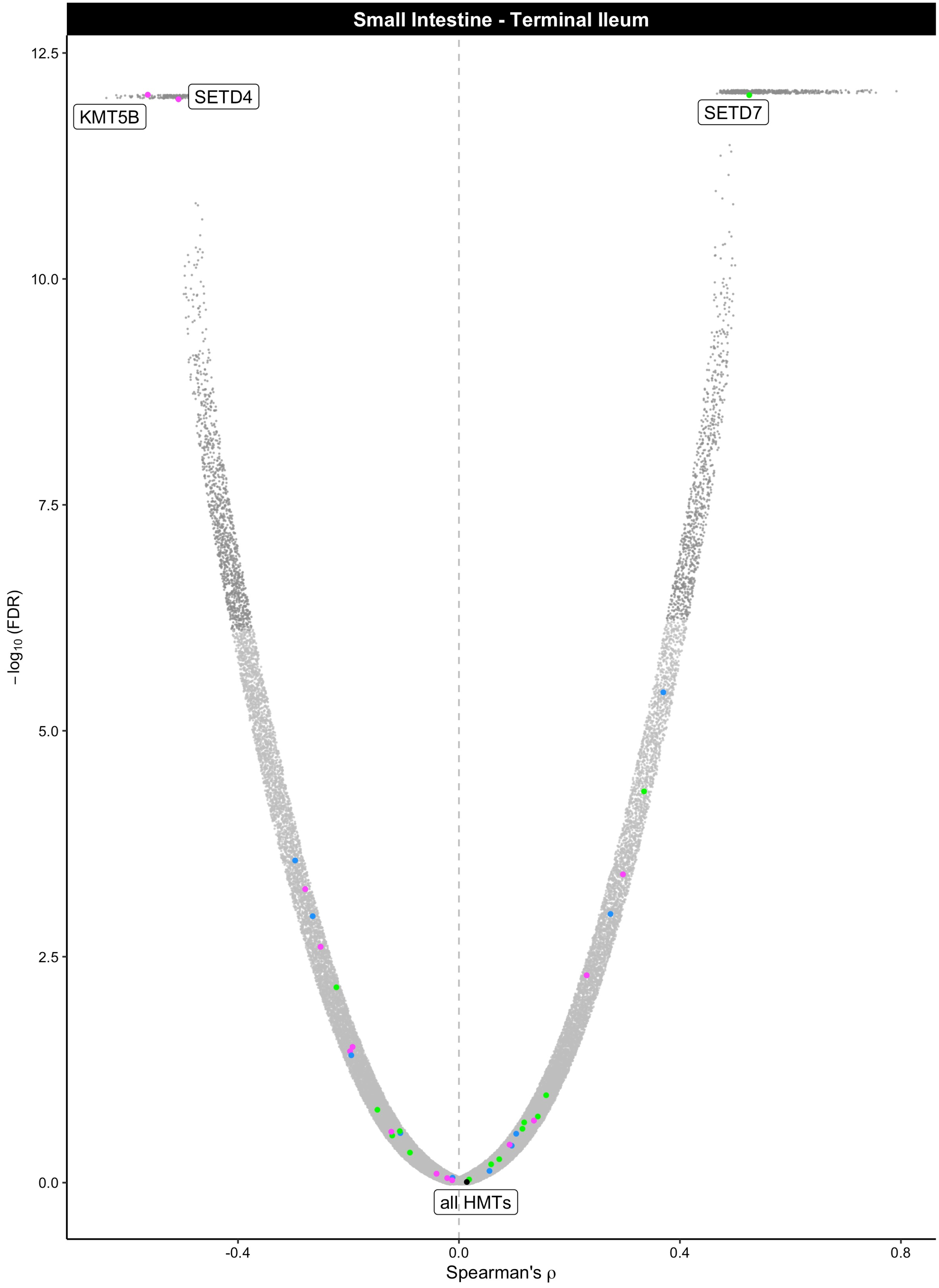

Small Intestine - Terminal Ileum
12.5
10.0
7.5
5.0
2.5
0.0
−log10 (FDR)
SETD4
KMT5B
SETD7
all HMTs
-0.4
0.0
0.4
0.8
Spearman's ρ

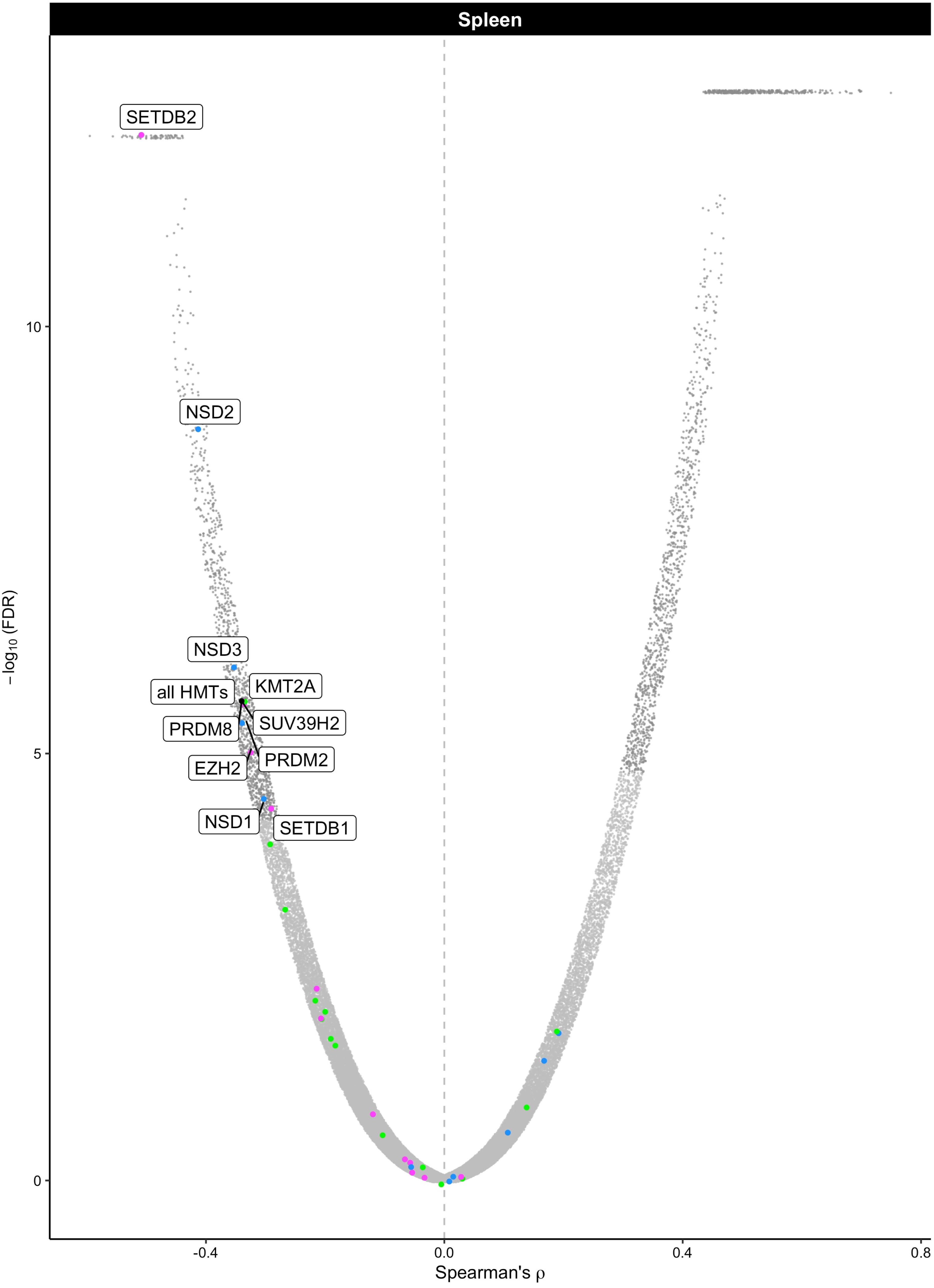
Spleen
SETDB2
NSD2
NSD3
all HMTs
KMT2A
SUV39H2
PRDM8
EZH2
PRDM2
NSD1
SETDB1
10
5
0
−log10 (FDR)
-0.4
0.0
0.4
0.8
Spearman's ρ

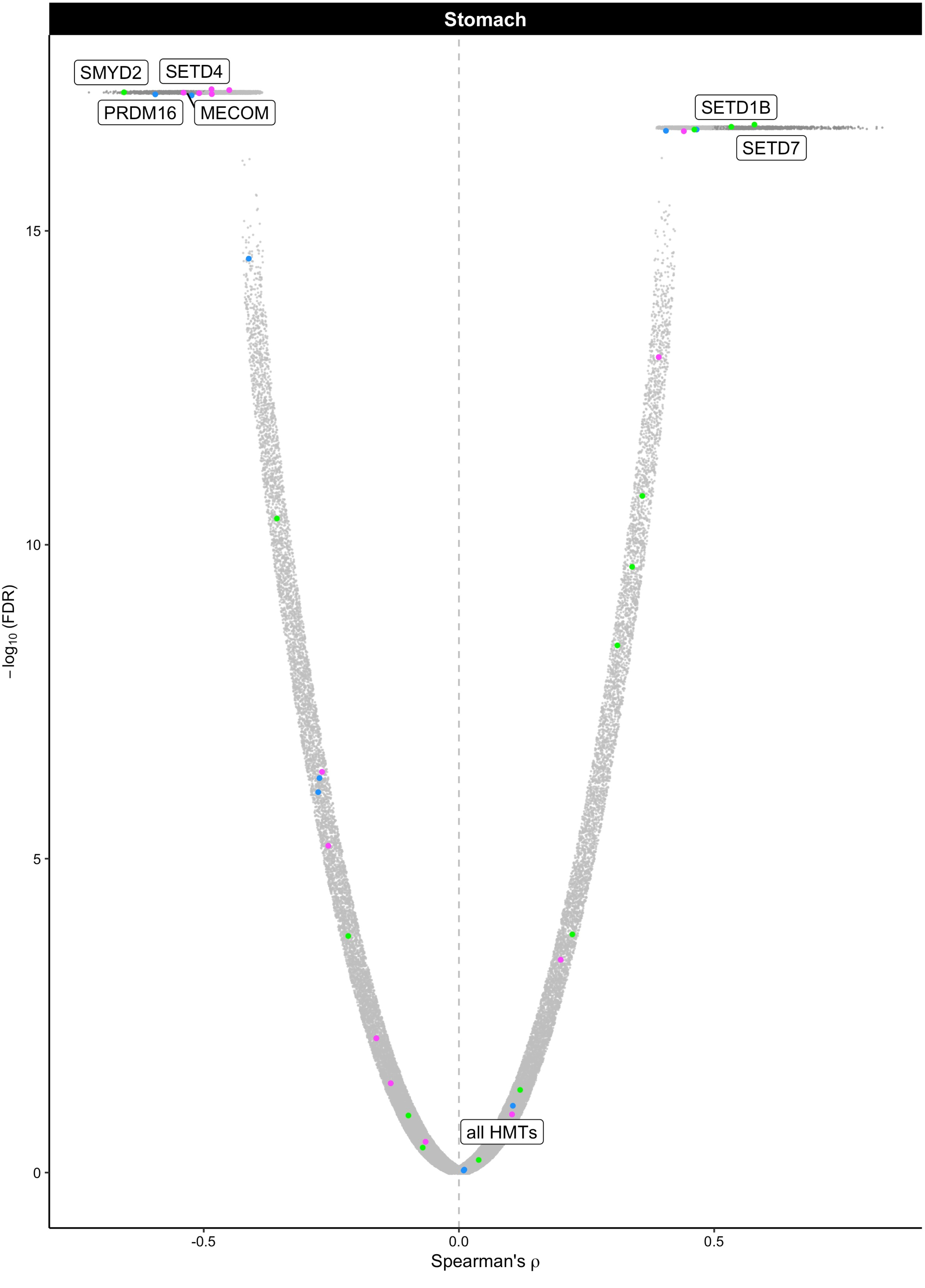
Stomach
SMYD2
SETD4
PRDM16
MECOM
SETD1B
SETD7
all HMTs
15
10
5
0
-0.5
0.0
0.5
−log10 (FDR)
Spearman's ρ

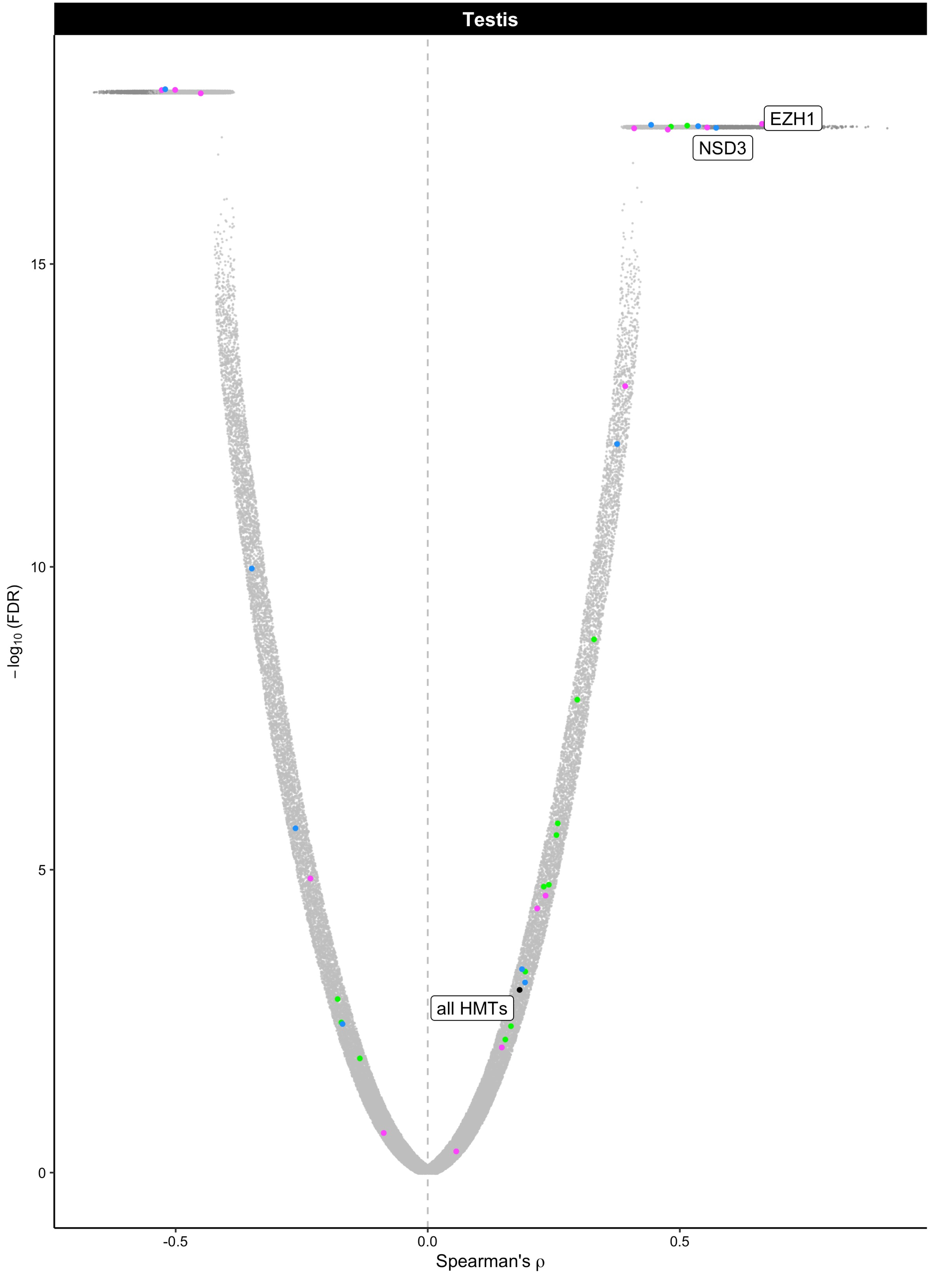

Testis
EZH1
NSD3
all HMTs
−log10 (FDR)
Spearman's ρ
15
10
5
0
-0.5
0.0
0.5

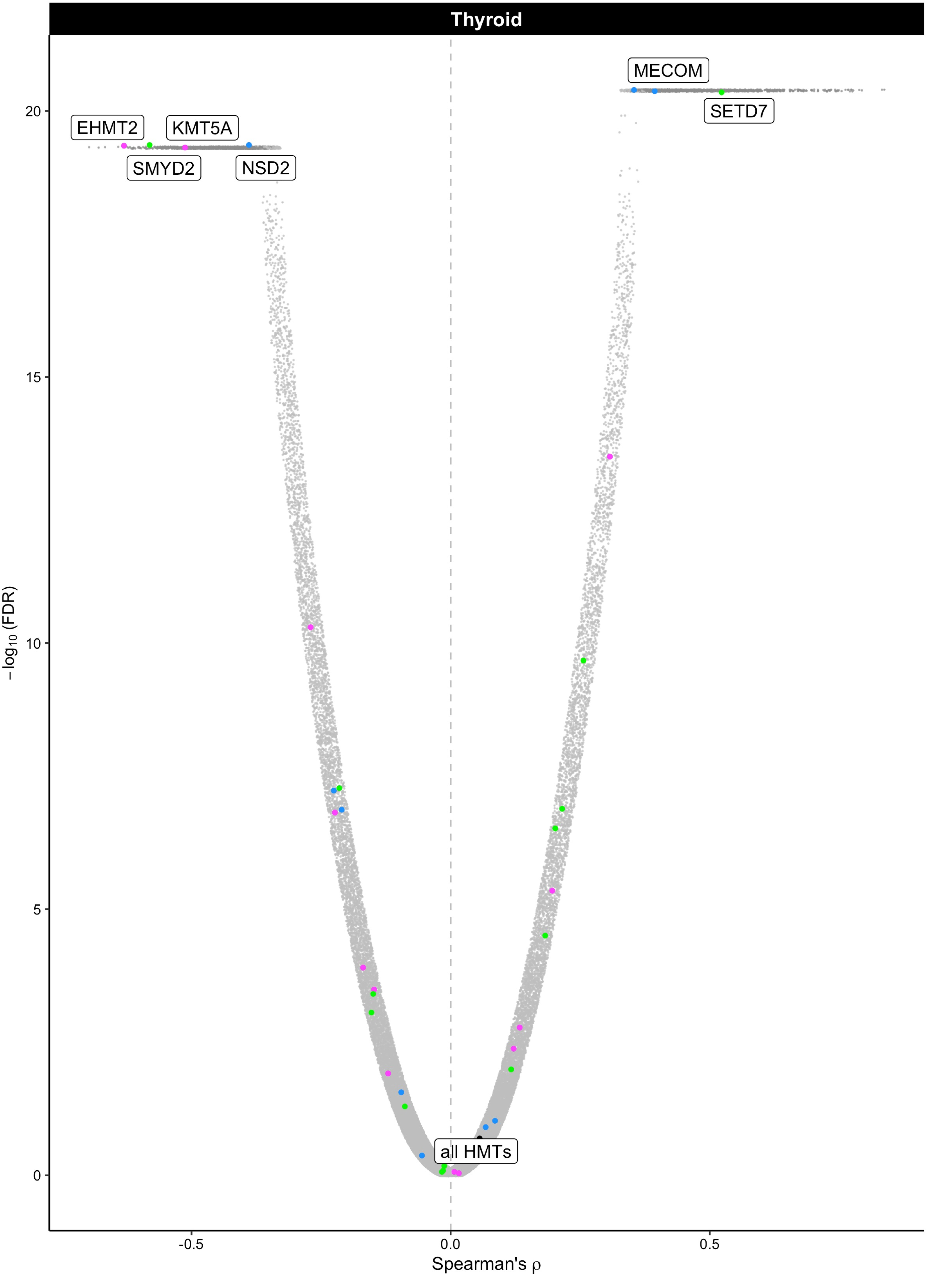

Thyroid
MECOM
SETD7
EHMT2
KMT5A
SMYD2
NSD2
all HMTs
20
15
10
5
0
−log10 (FDR)
-0.5
0.0
0.5
Spearman's ρ

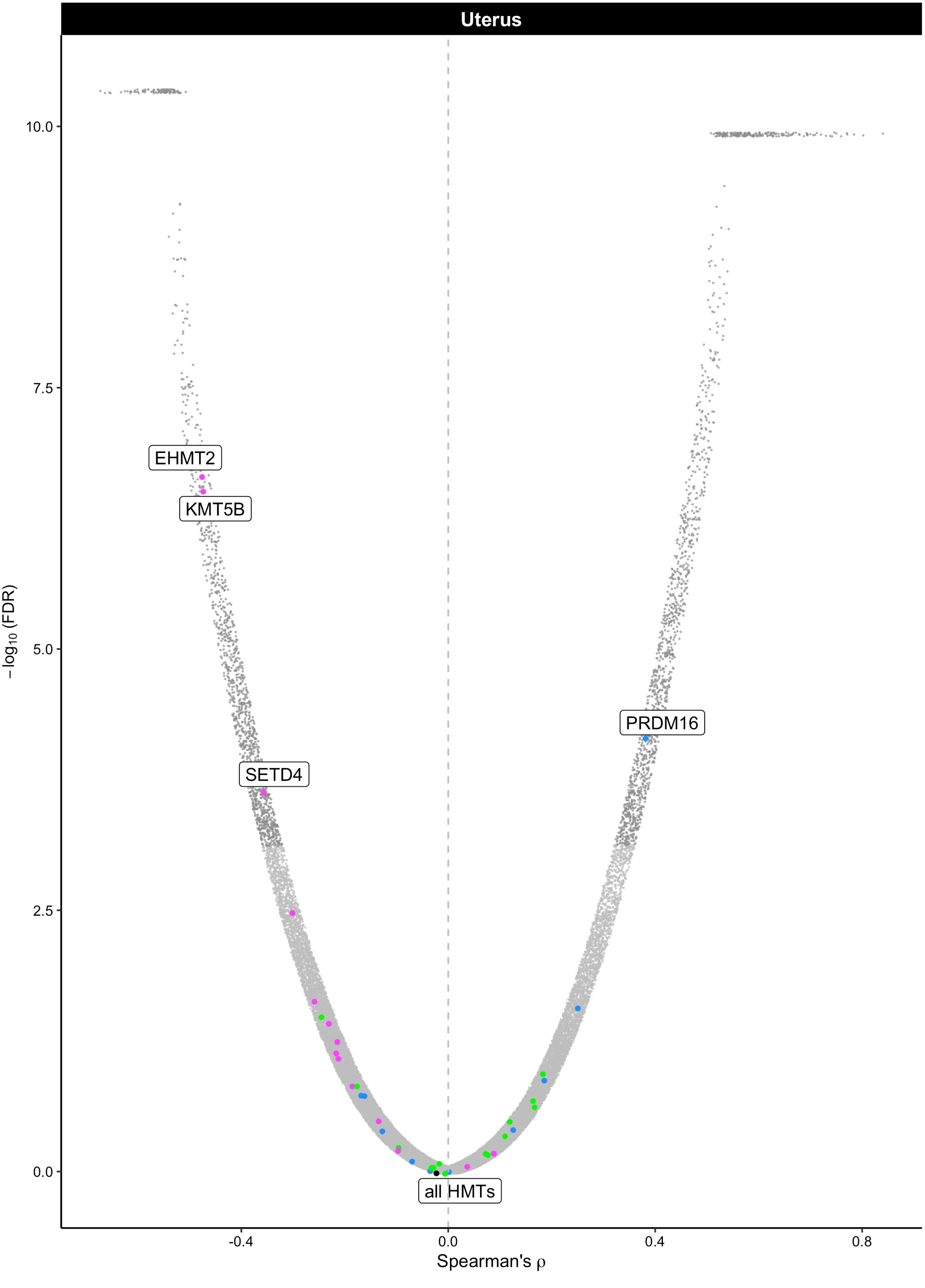

Uterus
EHMT2
KMT5B
SETD4
PRDM16
all HMTs
10.0
7.5
5.0
2.5
0.0
-0.4
0.0
0.4
0.8
$-\log_{10}$ (FDR)
Spearman's ρ

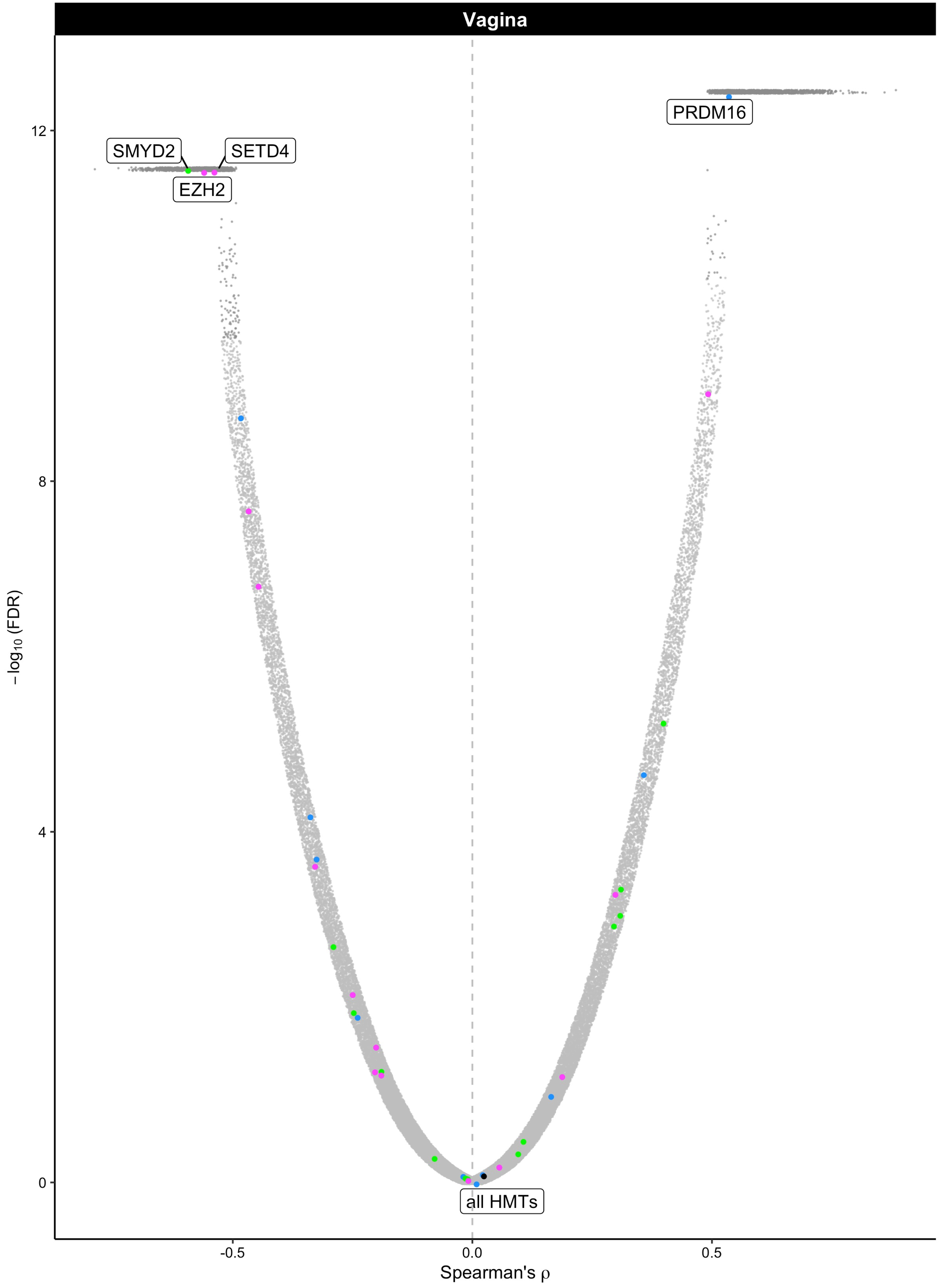

Vagina
SMYD2
SETD4
EZH2
PRDM16
all HMTs
12
8
4
0
-0.5
0.0
0.5
−log10 (FDR)
Spearman's ρ

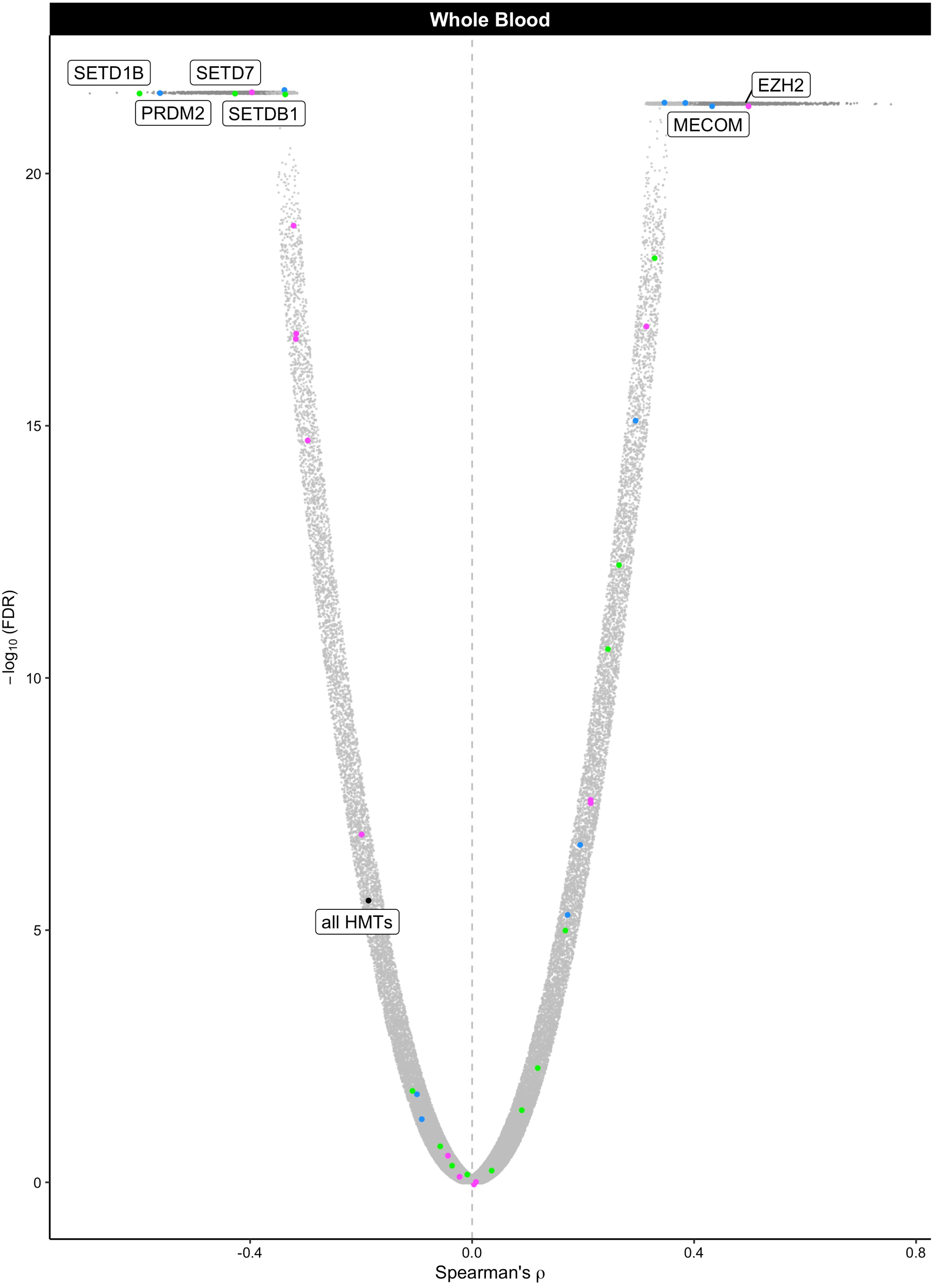

Whole Blood
SETD1B
SETD7
PRDM2
SETDB1
EZH2
MECOM
all HMTs
−log10 (FDR)
20
15
10
5
0
-0.4
0.0
0.4
0.8
Spearman's ρ
