## Supplemental Figures and Tables for "Histone methylation has a direct metabolic role in human cells": Supp File 4. GTEX_PEMT_HMTs_fullplots.pdf

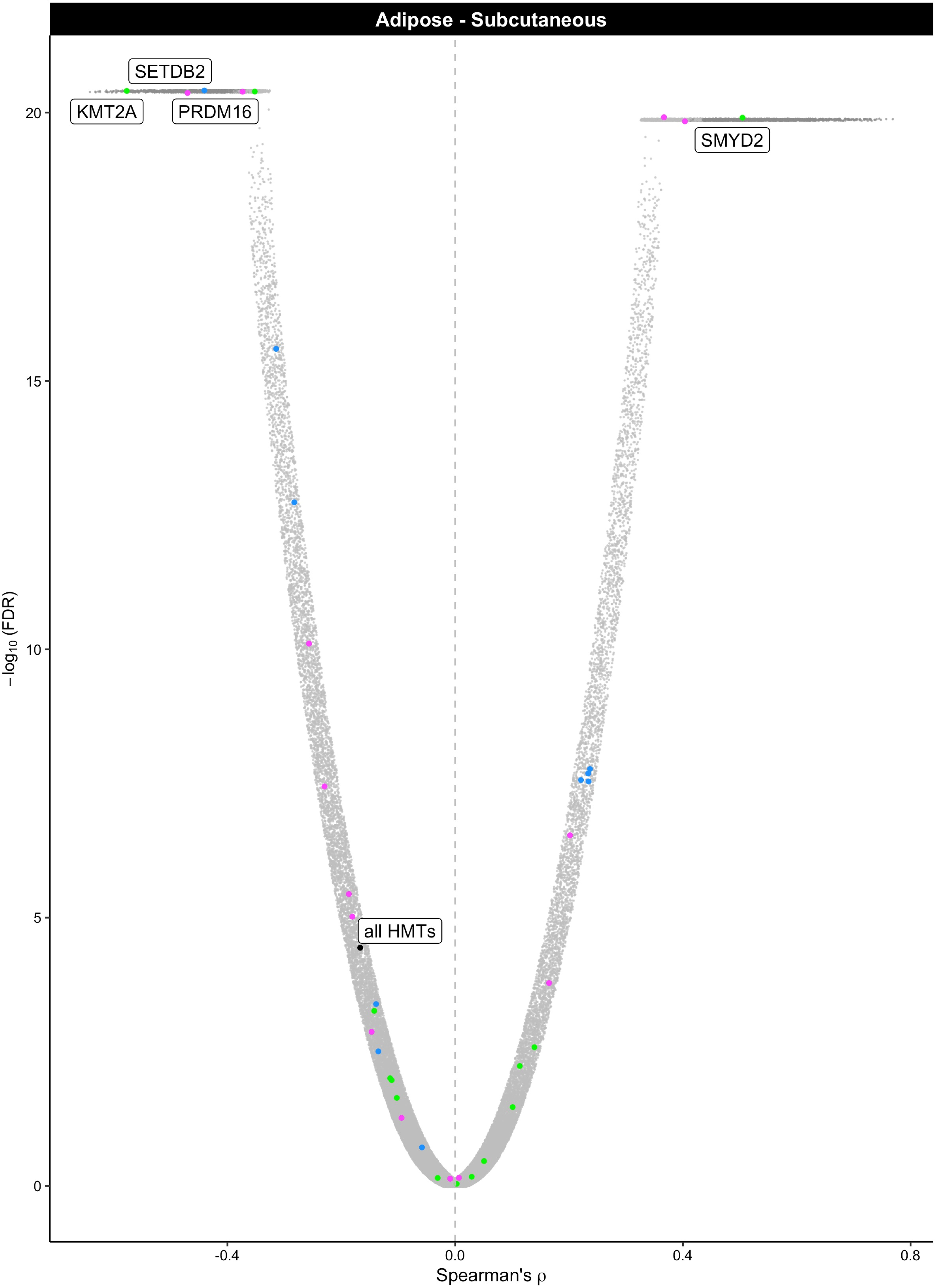

Adipose - Subcutaneous
SETDB2
KMT2A
PRDM16
SMYD2
all HMTs
20
15
10
5
0
−log10 (FDR)
-0.4
0.0
0.4
0.8
Spearman's ρ

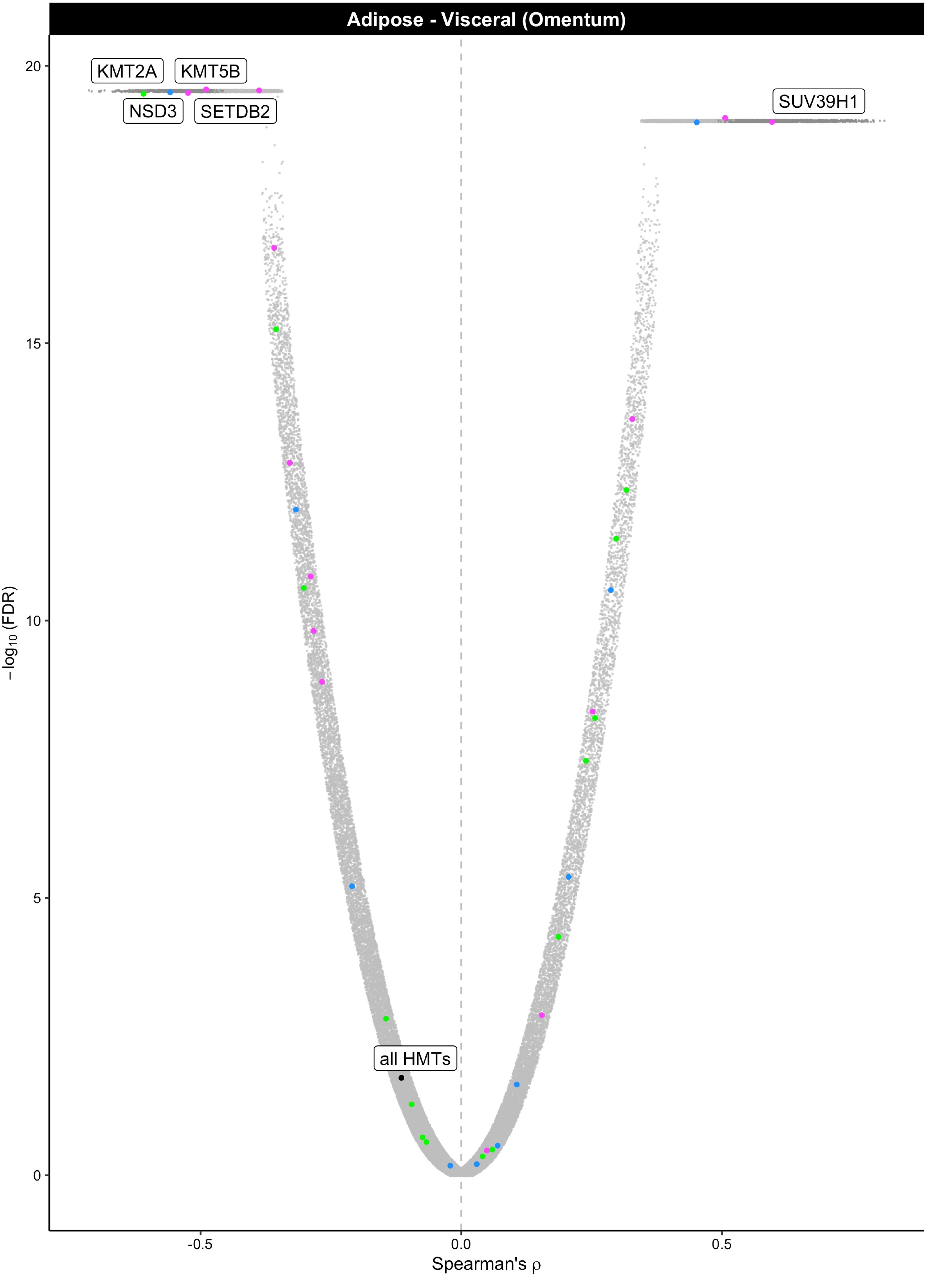

Adipose - Visceral (Omentum)
KMT2A
KMT5B
NSD3
SETDB2
SUV39H1
all HMTs
20
15
10
5
0
-log10 (FDR)
-0.5
0.0
0.5
Spearman's ρ

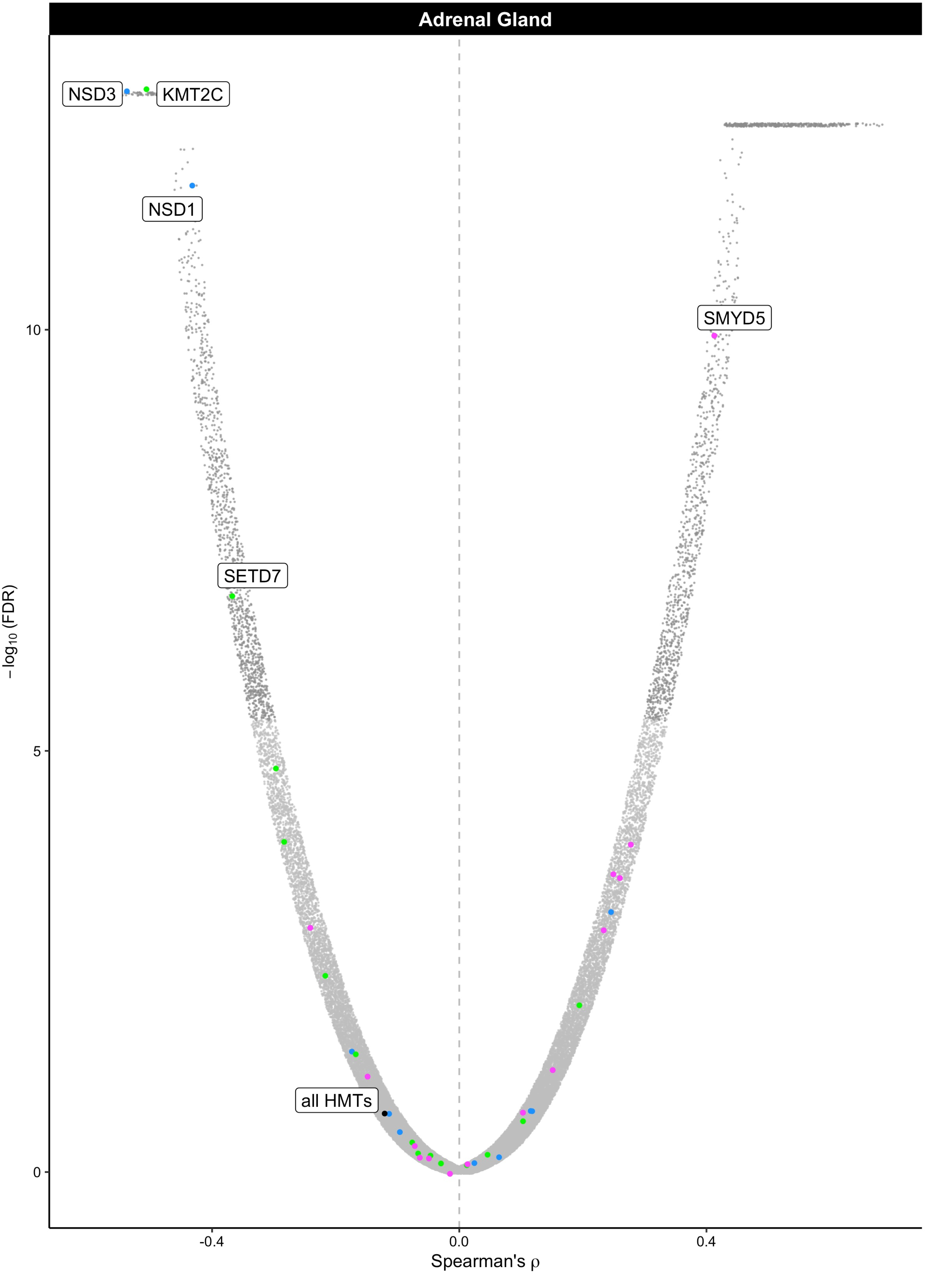

Adrenal Gland
NSD3
KMT2C
NSD1
SMYD5
SETD7
all HMTs
10
5
0
-0.4
0.0
0.4
−log10 (FDR)
Spearman's ρ

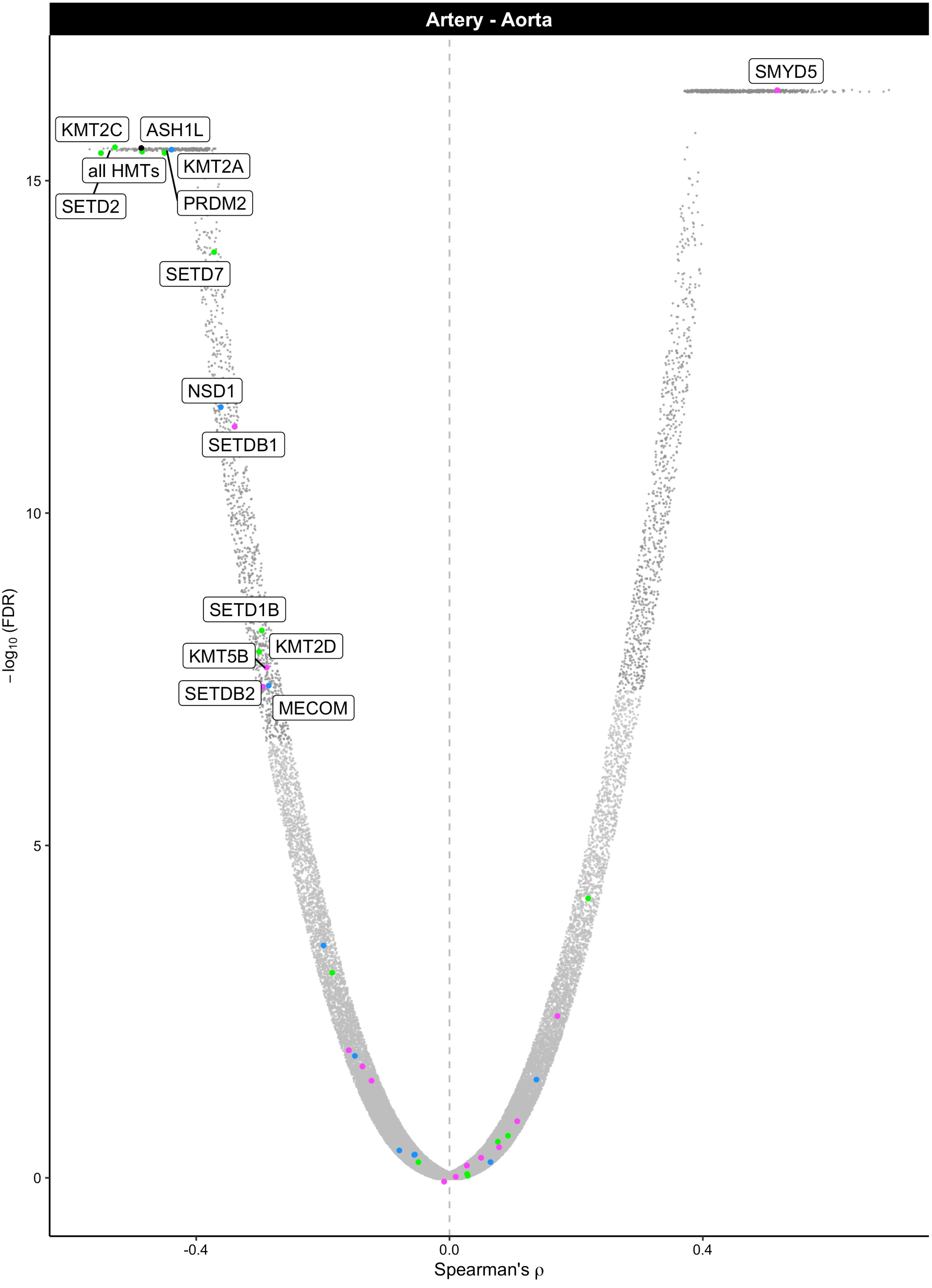

Artery - Aorta
SMYD5
KMT2C
ASH1L
all HMTs
KMT2A
SETD2
PRDM2
SETD7
NSD1
SETDB1
SETD1B
KMT2D
KMT5B
SETDB2
MECOM
15
10
5
0
−log10 (FDR)
-0.4
0.0
0.4
Spearman's ρ

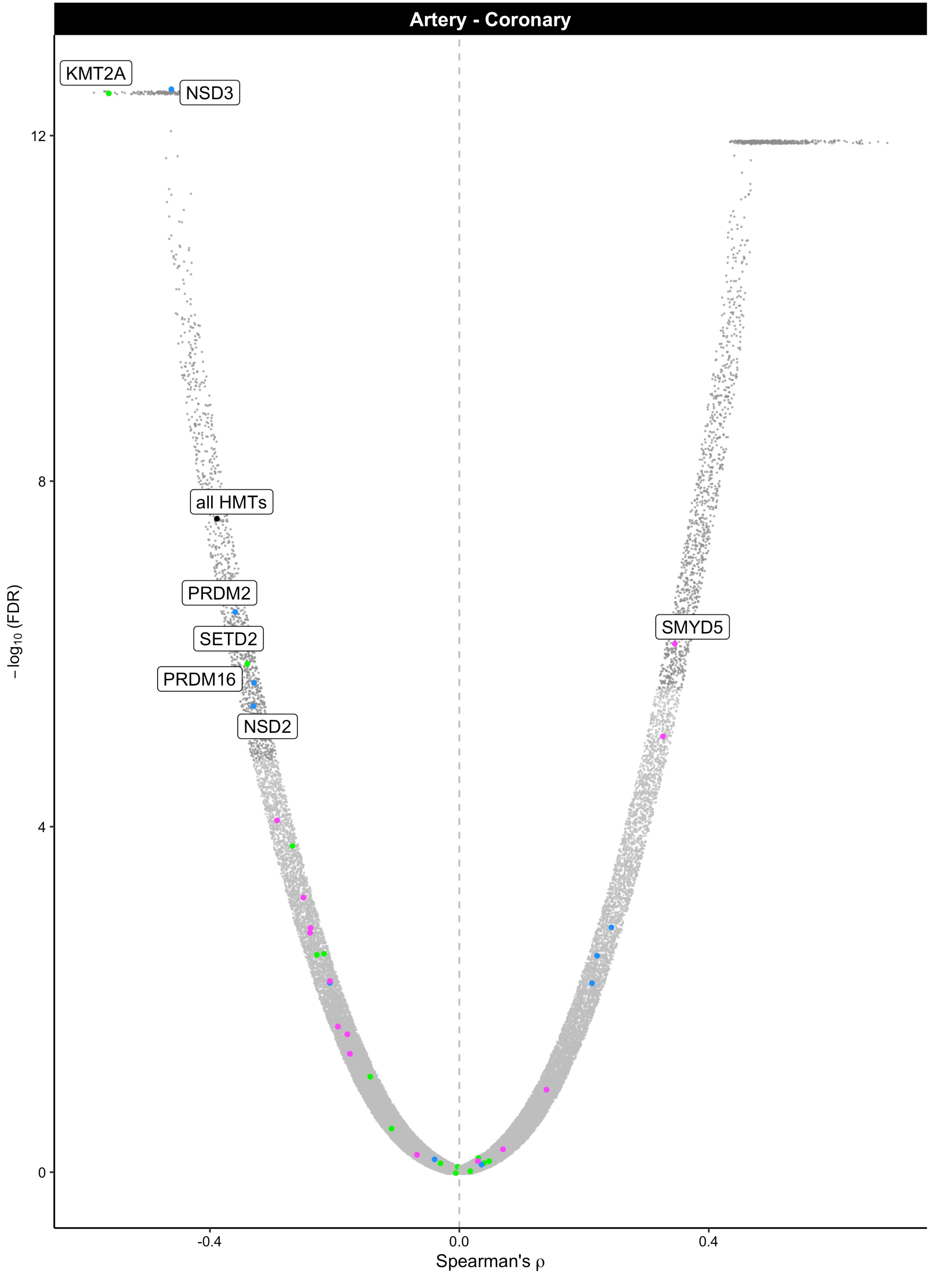

Artery - Coronary
KMT2A
NSD3
all HMTs
PRDM2
SETD2
PRDM16
NSD2
SMYD5
12
8
4
0
-0.4
0.0
0.4
$-\log_{10}$ (FDR)
Spearman's ρ

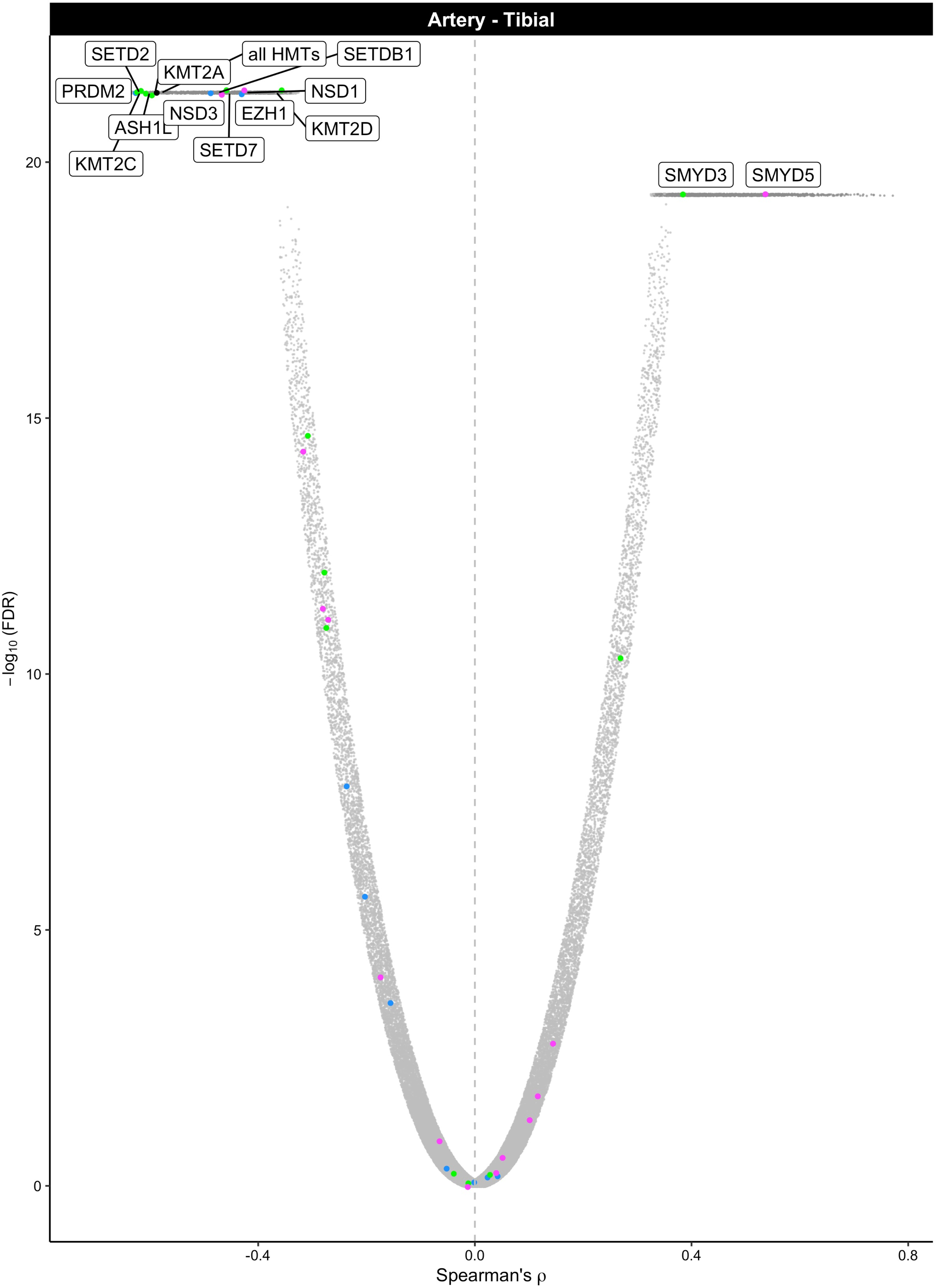

Artery - Tibial
SETD2
all HMTs
SETDB1
KMT2A
PRDM2
NSD1
NSD3
EZH1
ASH1L
KMT2D
SETD7
KMT2C
SMYD3
SMYD5
20
15
10
5
0
−log10 (FDR)
-0.4
0.0
0.4
0.8
Spearman's ρ

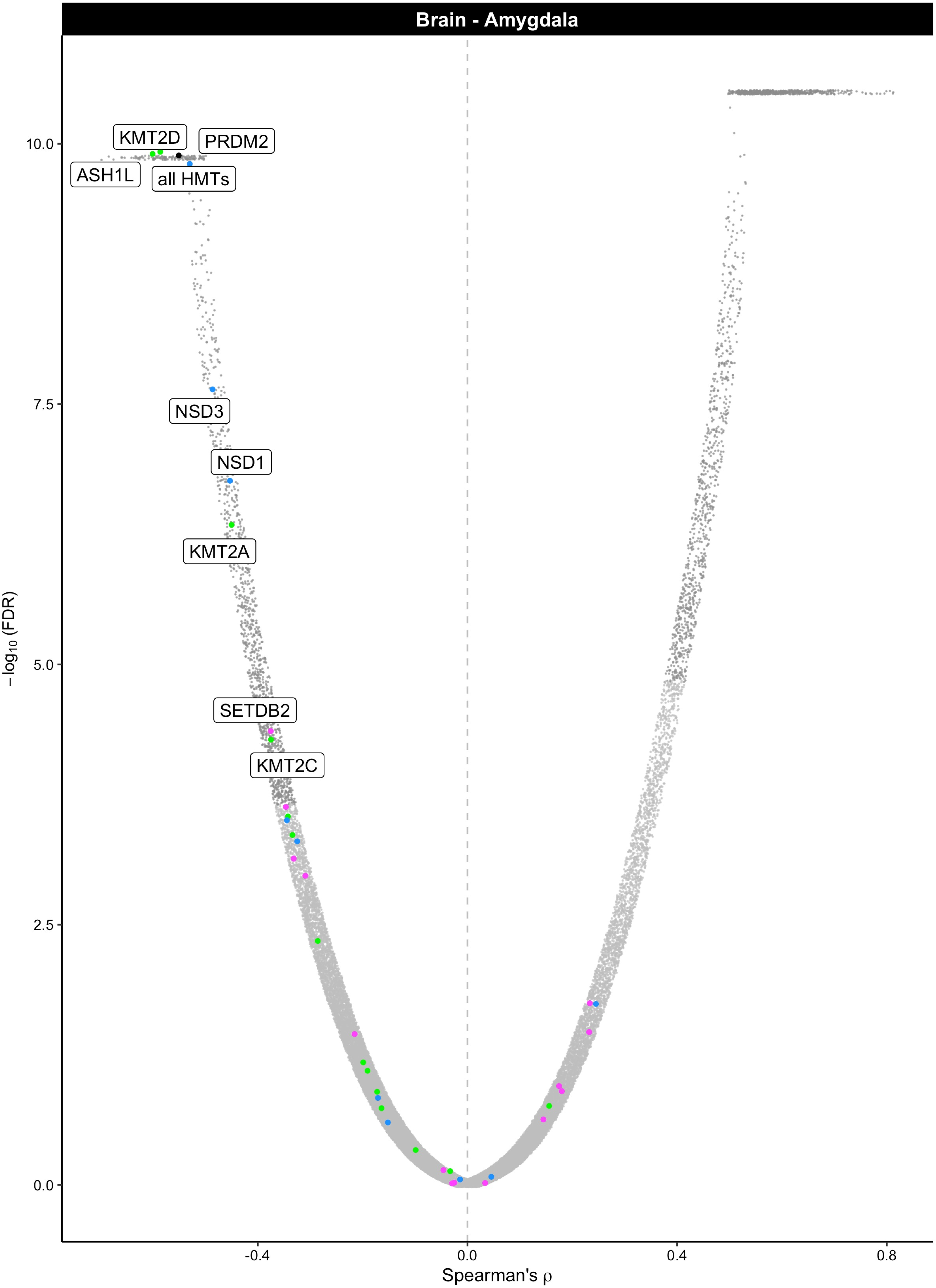

Brain - Amygdala
KMT2D
PRDM2
ASH1L
all HMTs
NSD3
NSD1
KMT2A
SETDB2
KMT2C
10.0
7.5
5.0
2.5
0.0
−log10 (FDR)
-0.4
0.0
0.4
0.8
Spearman's ρ

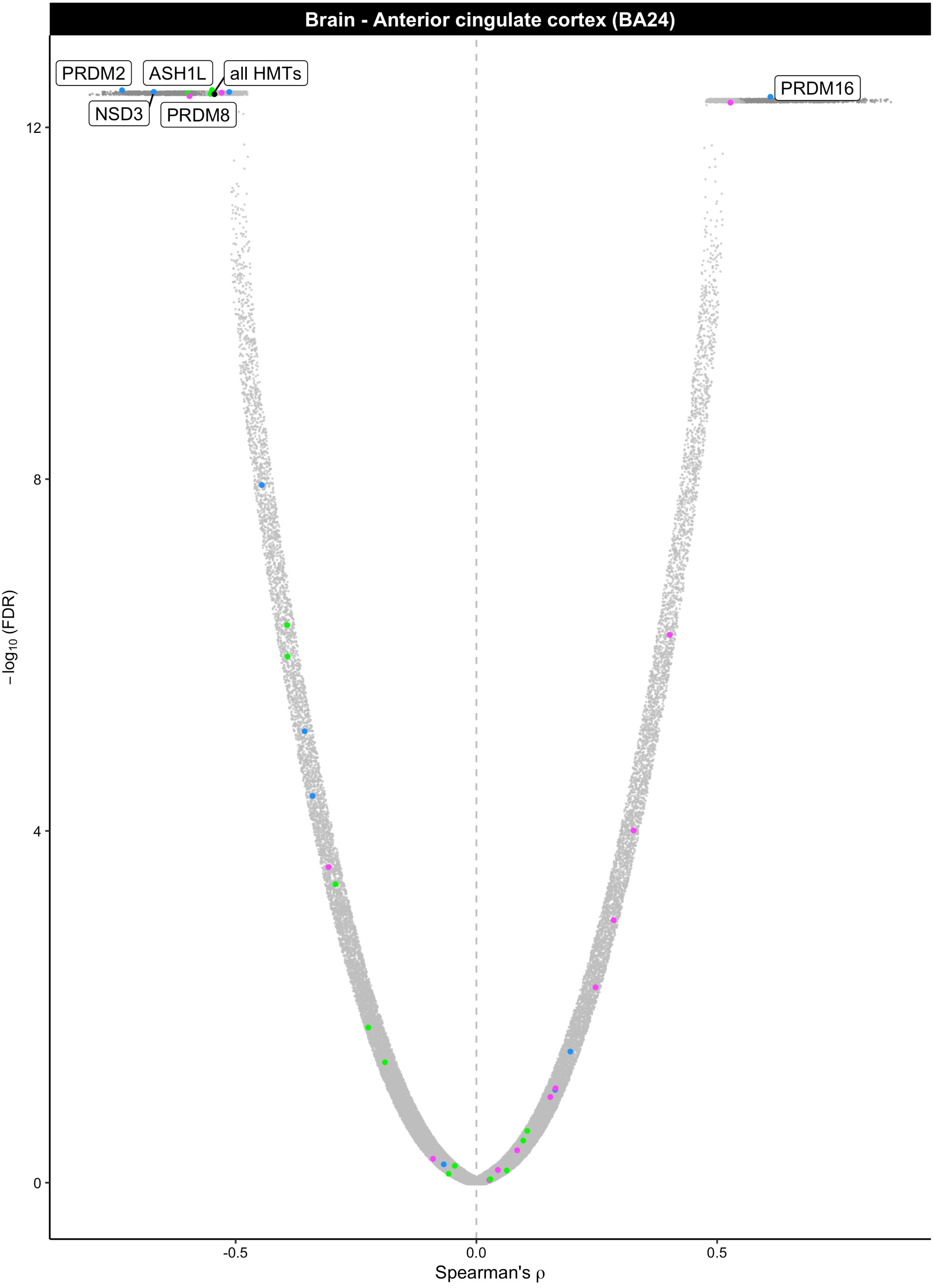
Brain - Anterior cingulate cortex (BA24)
PRDM2
ASH1L
all HMTs
NSD3
PRDM8
PRDM16
12
8
4
0
−log10 (FDR)
-0.5
0.0
0.5
Spearman's ρ

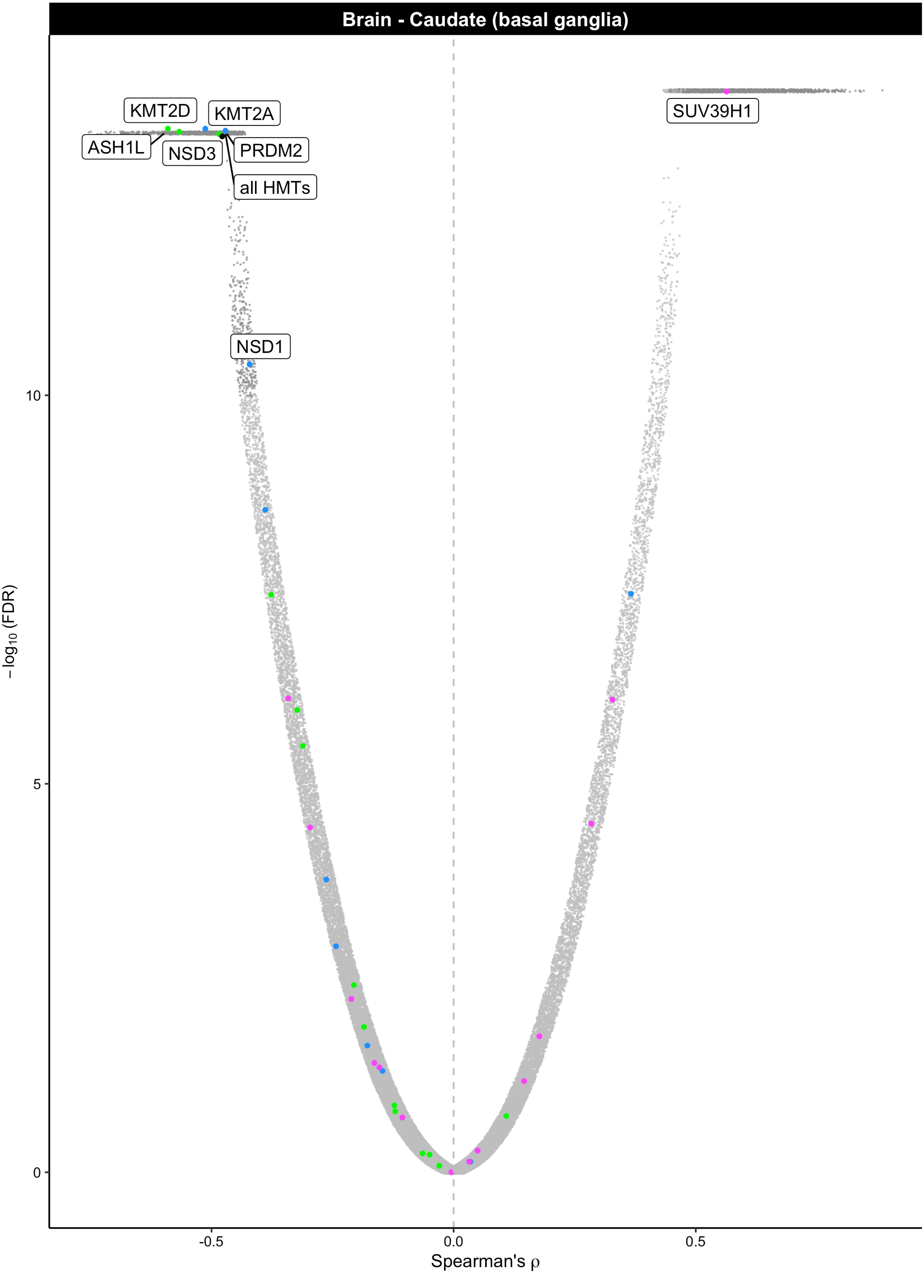

Brain - Caudate (basal ganglia)
KMT2D
KMT2A
ASH1L
NSD3
PRDM2
all HMTs
SUV39H1
NSD1
10
5
0
-0.5
0.0
0.5
−log10 (FDR)
Spearman's ρ

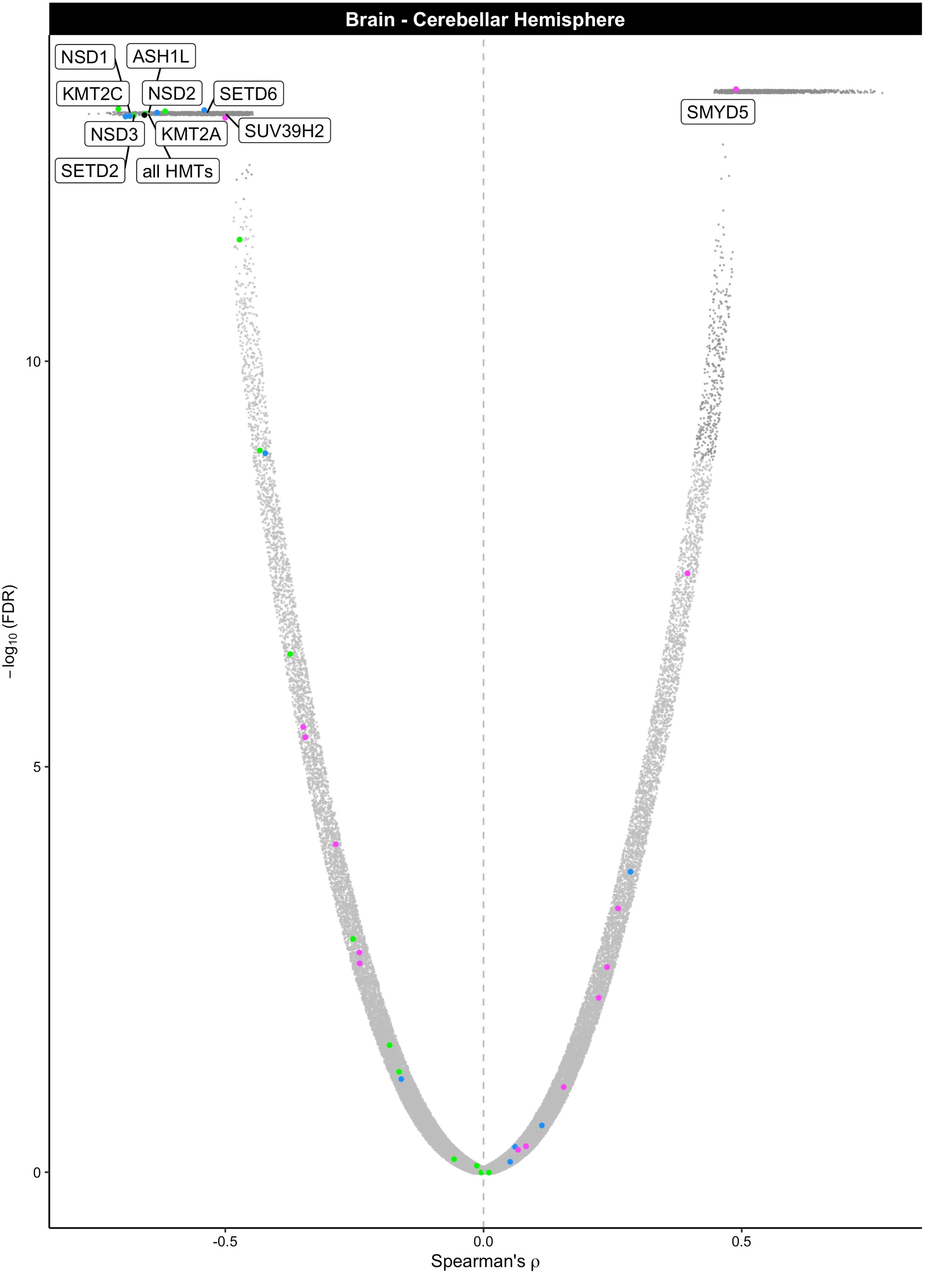

Brain - Cerebellar Hemisphere
NSD1
ASH1L
KMT2C
NSD2
SETD6
SMYD5
NSD3
KMT2A
SUV39H2
SETD2
all HMTs
10
5
0
-0.5
0.0
0.5
$-\log_{10}$ (FDR)
Spearman's ρ

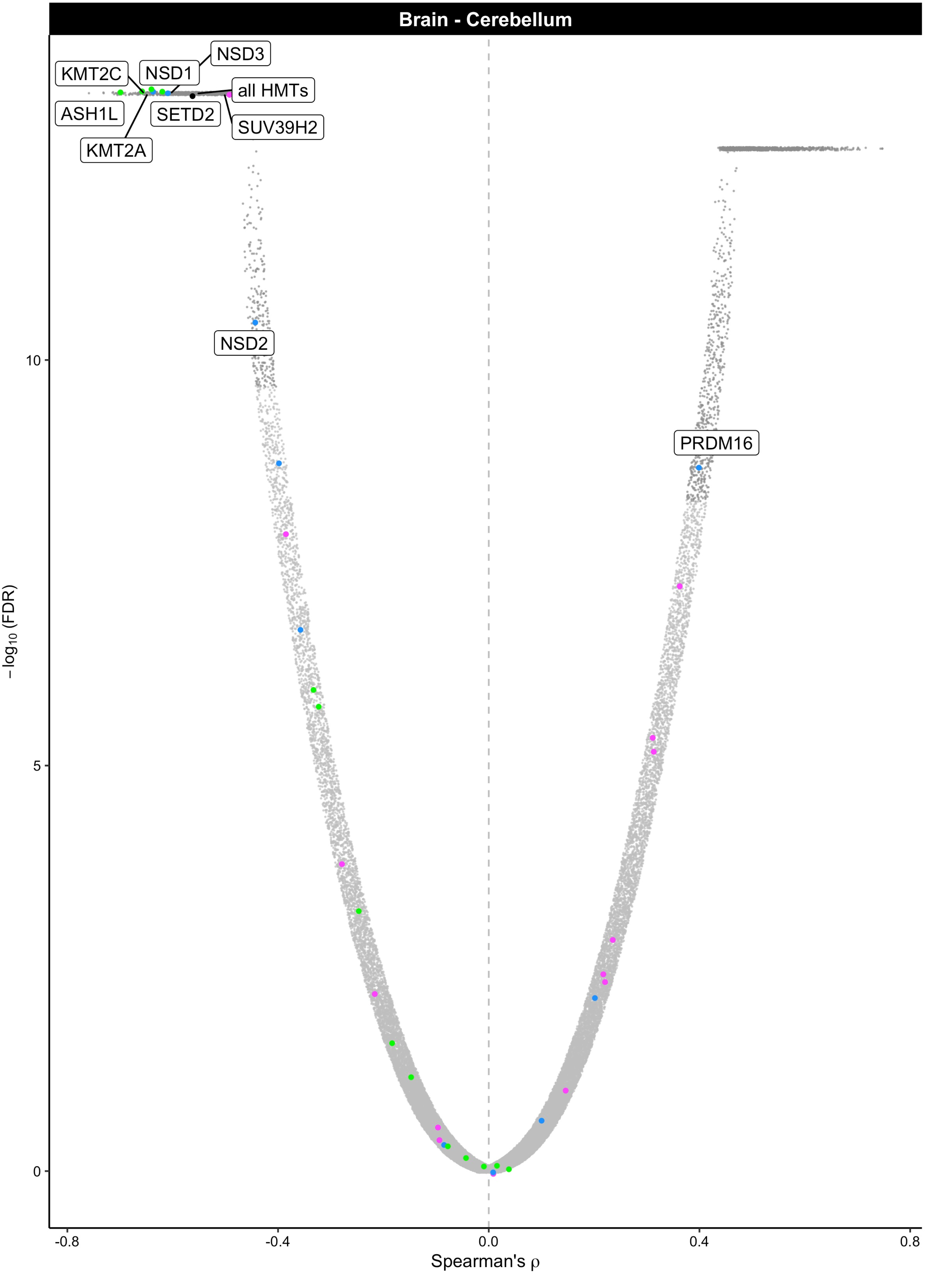

Brain - Cerebellum
NSD3
KMT2C
NSD1
all HMTs
ASH1L
SETD2
SUV39H2
KMT2A
NSD2
PRDM16
10
5
0
−log10 (FDR)
-0.8
-0.4
0.0
0.4
0.8
Spearman's ρ

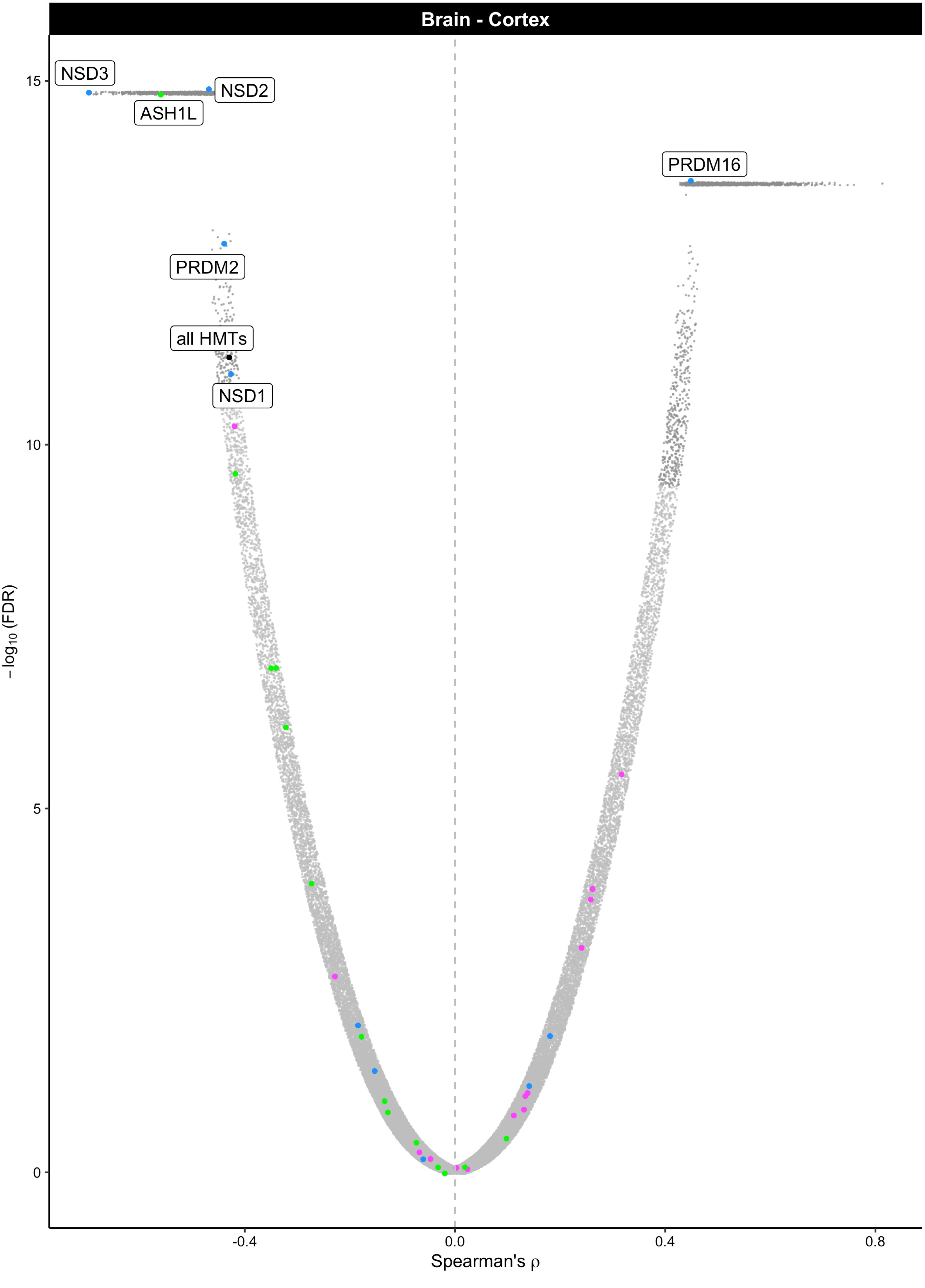
Brain - Cortex
NSD3
NSD2
ASH1L
PRDM16
PRDM2
all HMTs
NSD1
15
10
5
0
$-\log_{10}$(FDR)
-0.4
0.0
0.4
0.8
Spearman's ρ

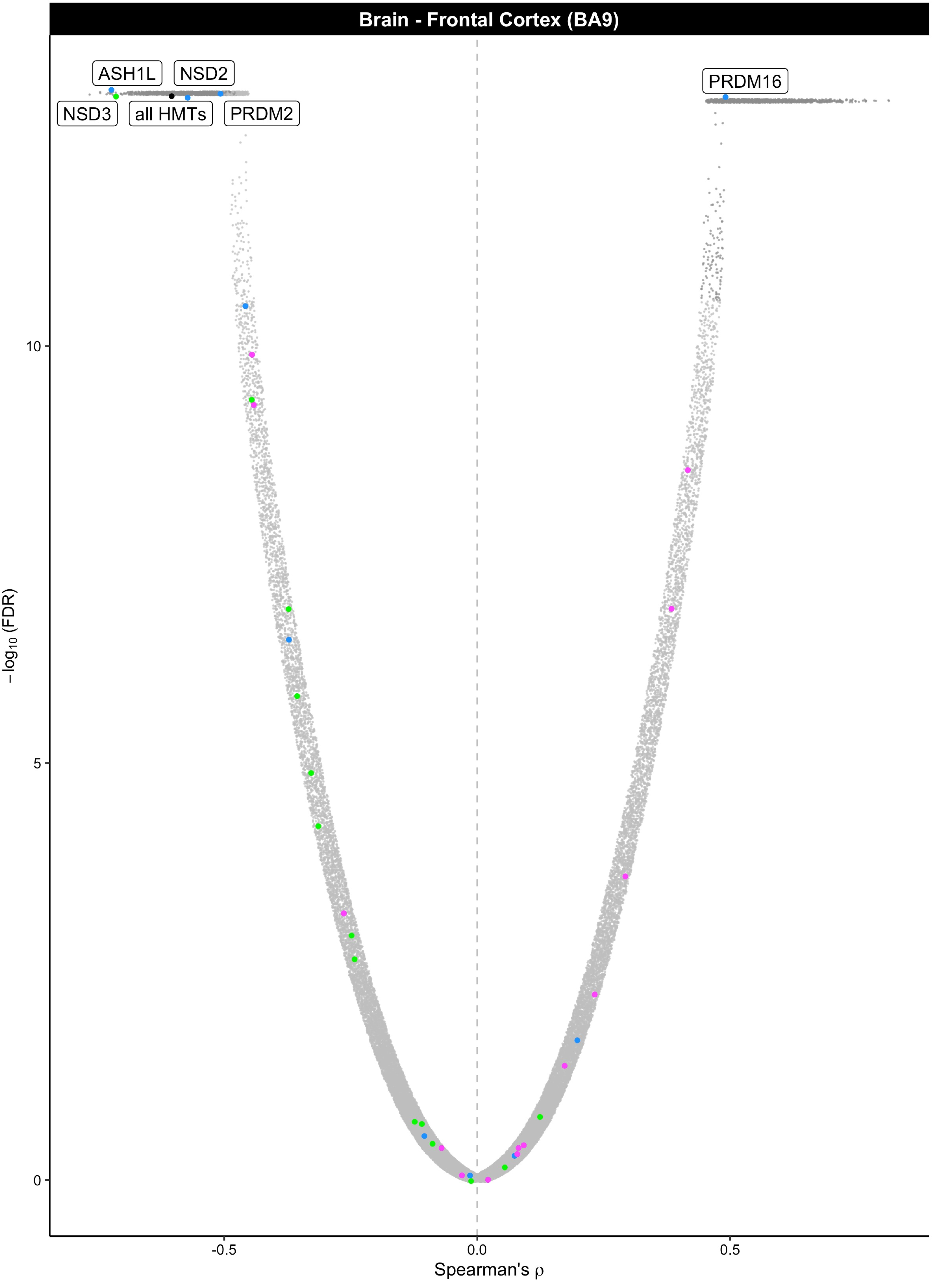

Brain - Frontal Cortex (BA9)
ASH1L
NSD2
PRDM16
NSD3
all HMTs
PRDM2
10
5
0
-0.5
0.0
0.5
−log10 (FDR)
Spearman's ρ

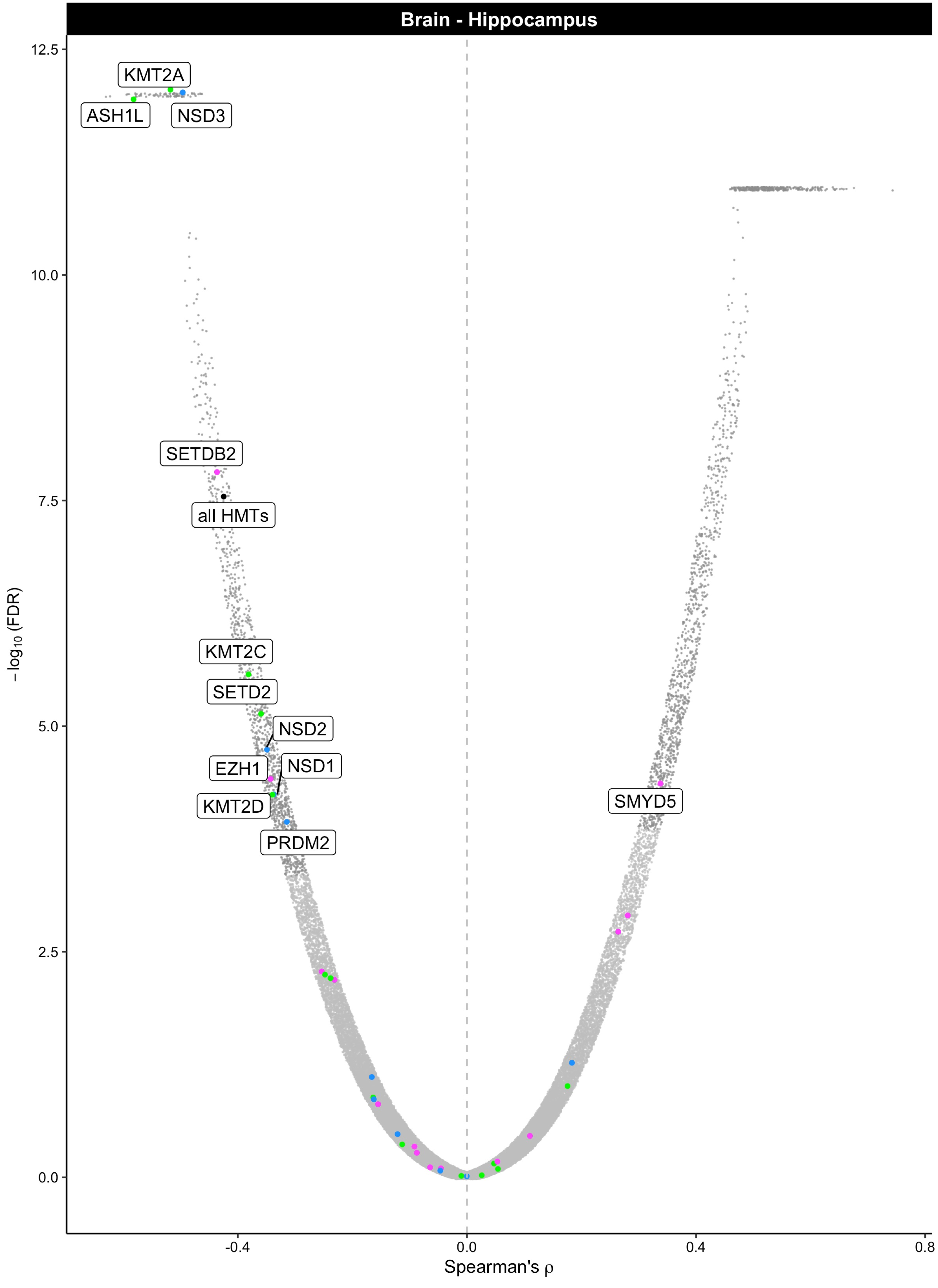

Brain - Hippocampus
12.5
10.0
7.5
5.0
2.5
0.0
−log10 (FDR)
KMT2A
ASH1L
NSD3
SETDB2
all HMTs
KMT2C
SETD2
NSD2
EZH1
NSD1
KMT2D
PRDM2
SMYD5
-0.4
0.0
0.4
0.8
Spearman's ρ

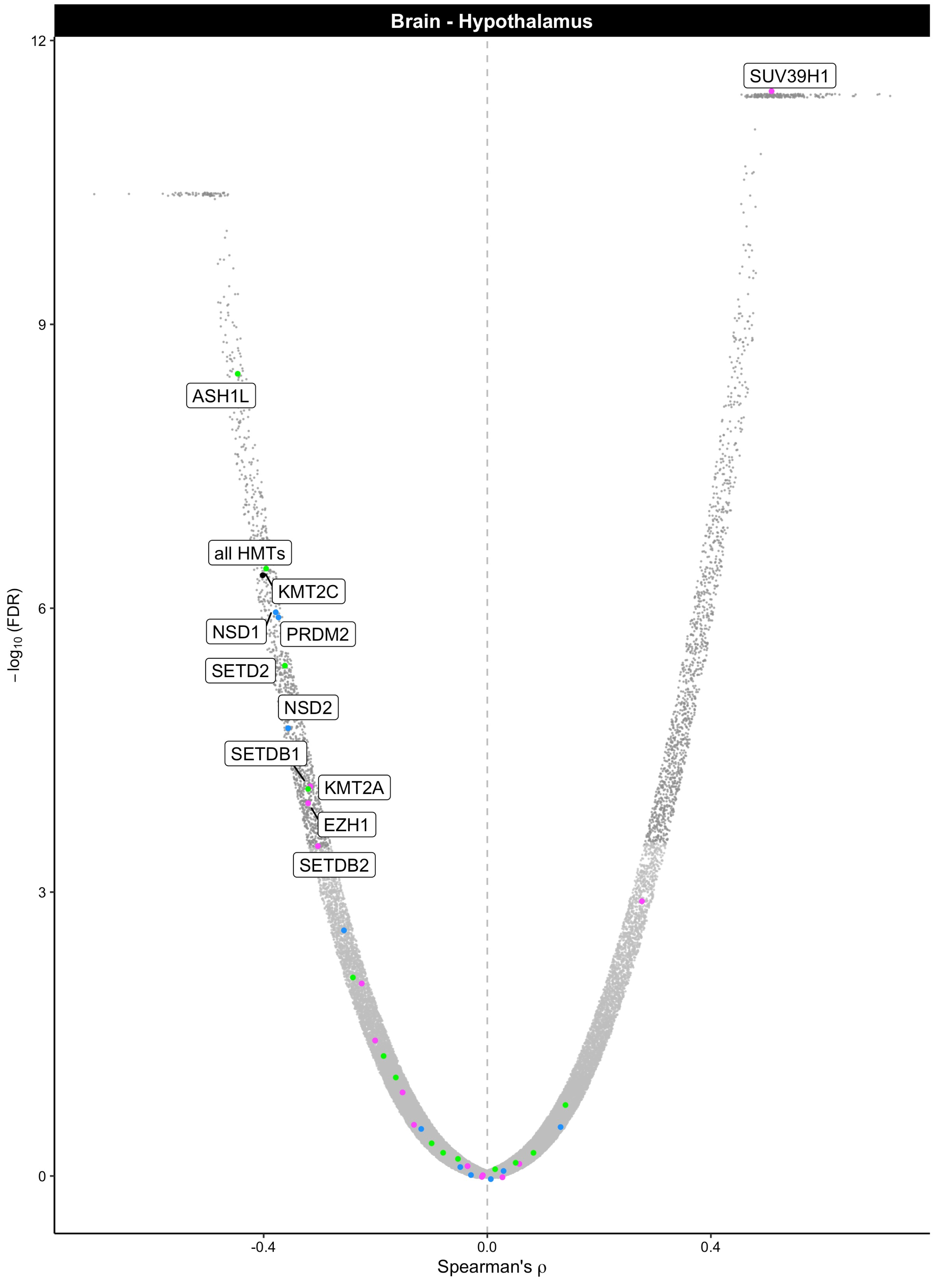
Brain - Hypothalamus
SUV39H1
ASH1L
all HMTs
KMT2C
NSD1
PRDM2
SETD2
NSD2
SETDB1
KMT2A
EZH1
SETDB2
12
9
6
3
0
-0.4
0.0
0.4
−log10 (FDR)
Spearman's ρ

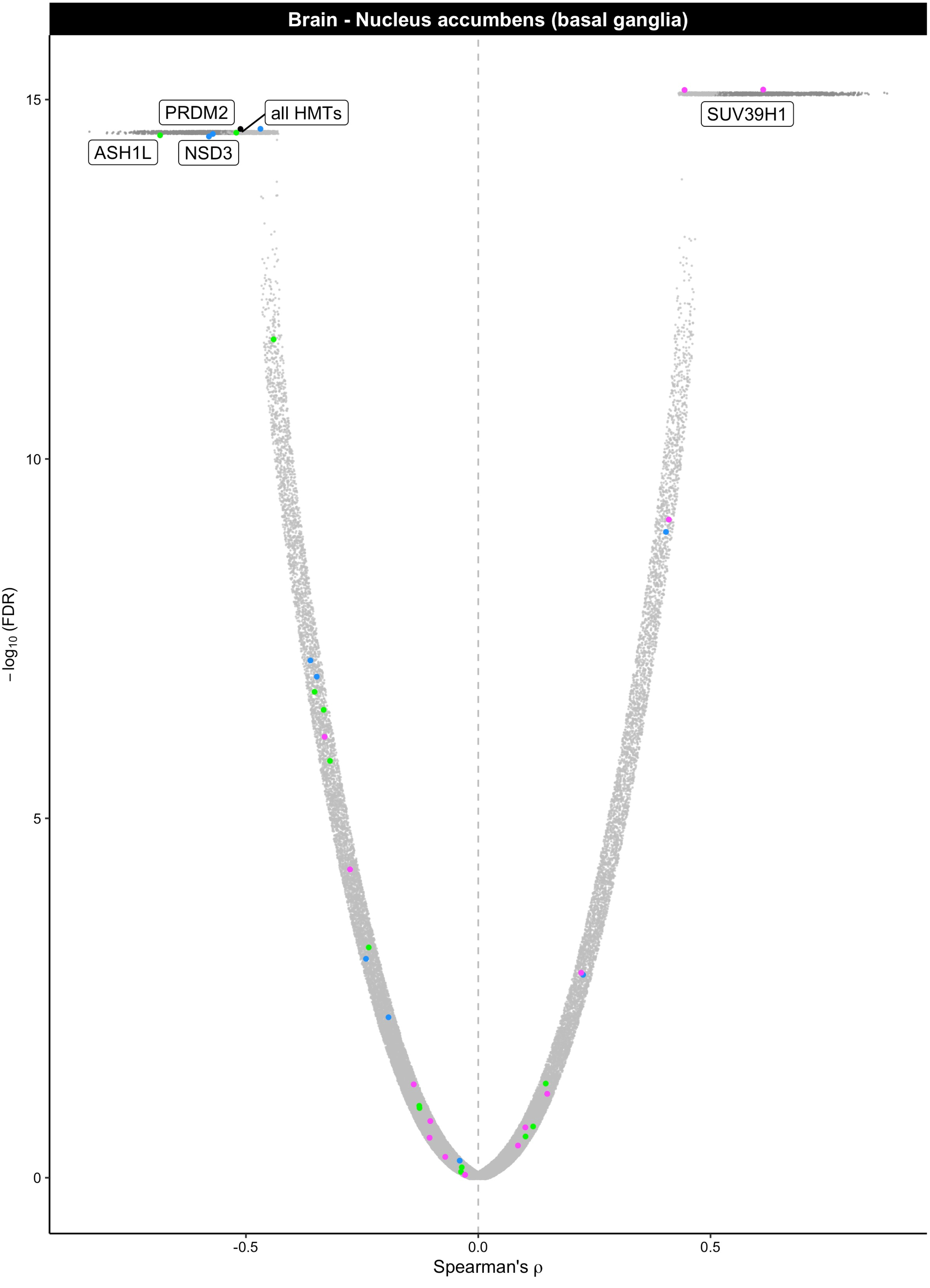

Brain - Nucleus accumbens (basal ganglia)
PRDM2
all HMTs
ASH1L
NSD3
SUV39H1
15
10
5
0
-log10 (FDR)
-0.5
0.0
0.5
Spearman's ρ

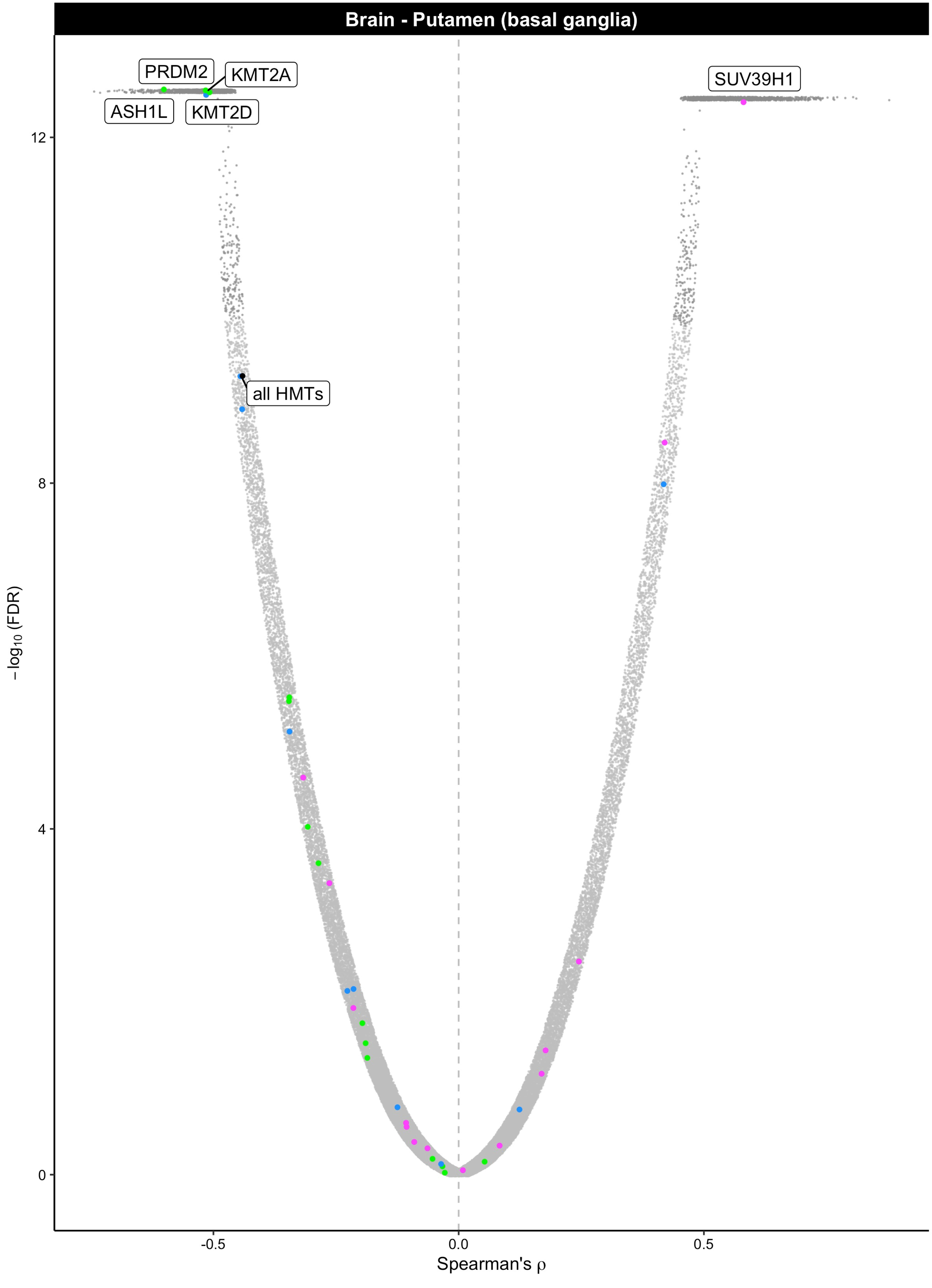

Brain - Putamen (basal ganglia)
PRDM2
KMT2A
ASH1L
KMT2D
SUV39H1
all HMTs
12
8
4
0
−log10 (FDR)
-0.5
0.0
0.5
Spearman's ρ

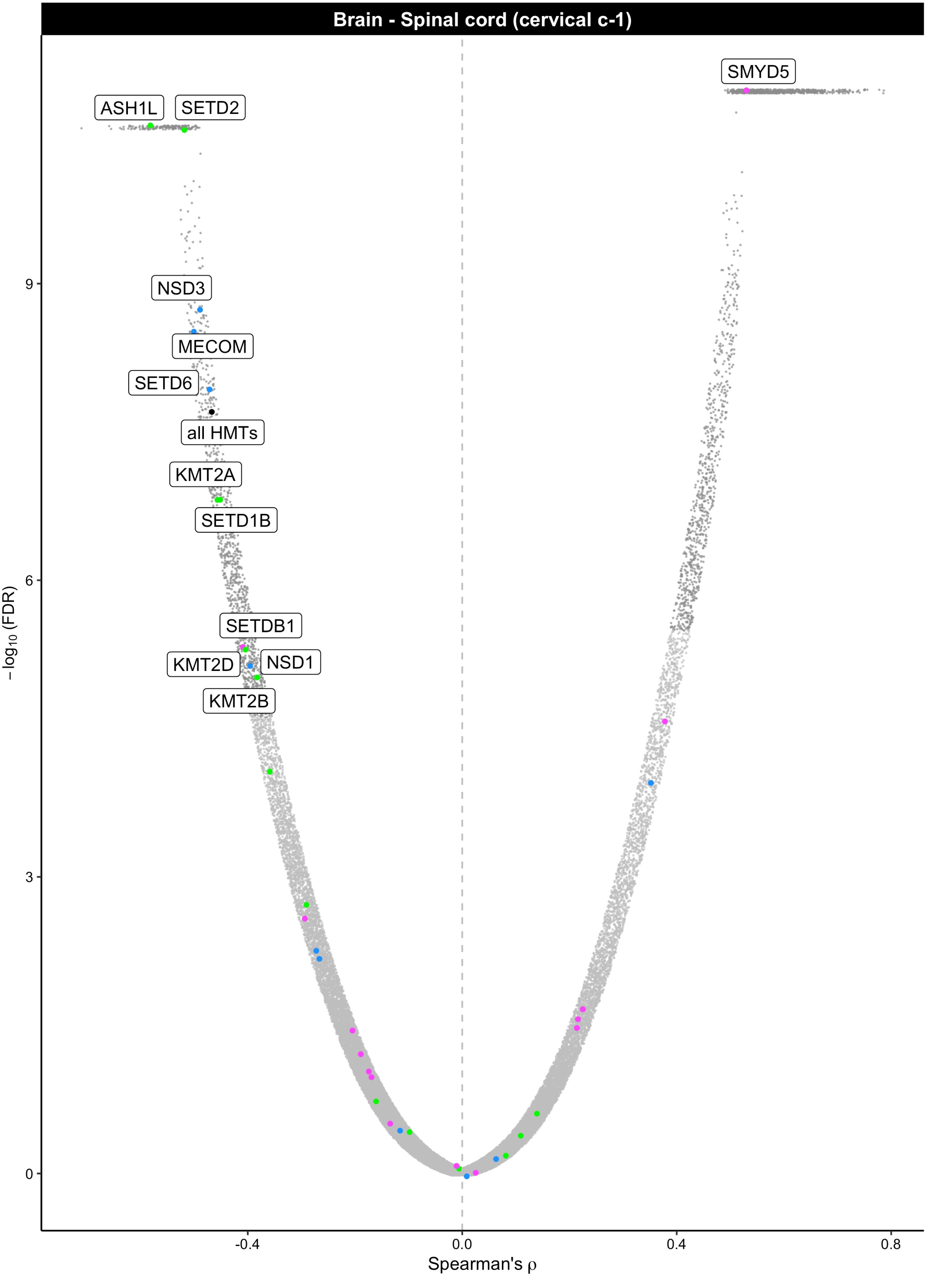
Brain - Spinal cord (cervical c-1)
SMYD5
ASH1L
SETD2
NSD3
MECOM
SETD6
all HMTs
KMT2A
SETD1B
SETDB1
KMT2D
NSD1
KMT2B
9
6
3
0
−log10 (FDR)
-0.4
0.0
0.4
0.8
Spearman's ρ

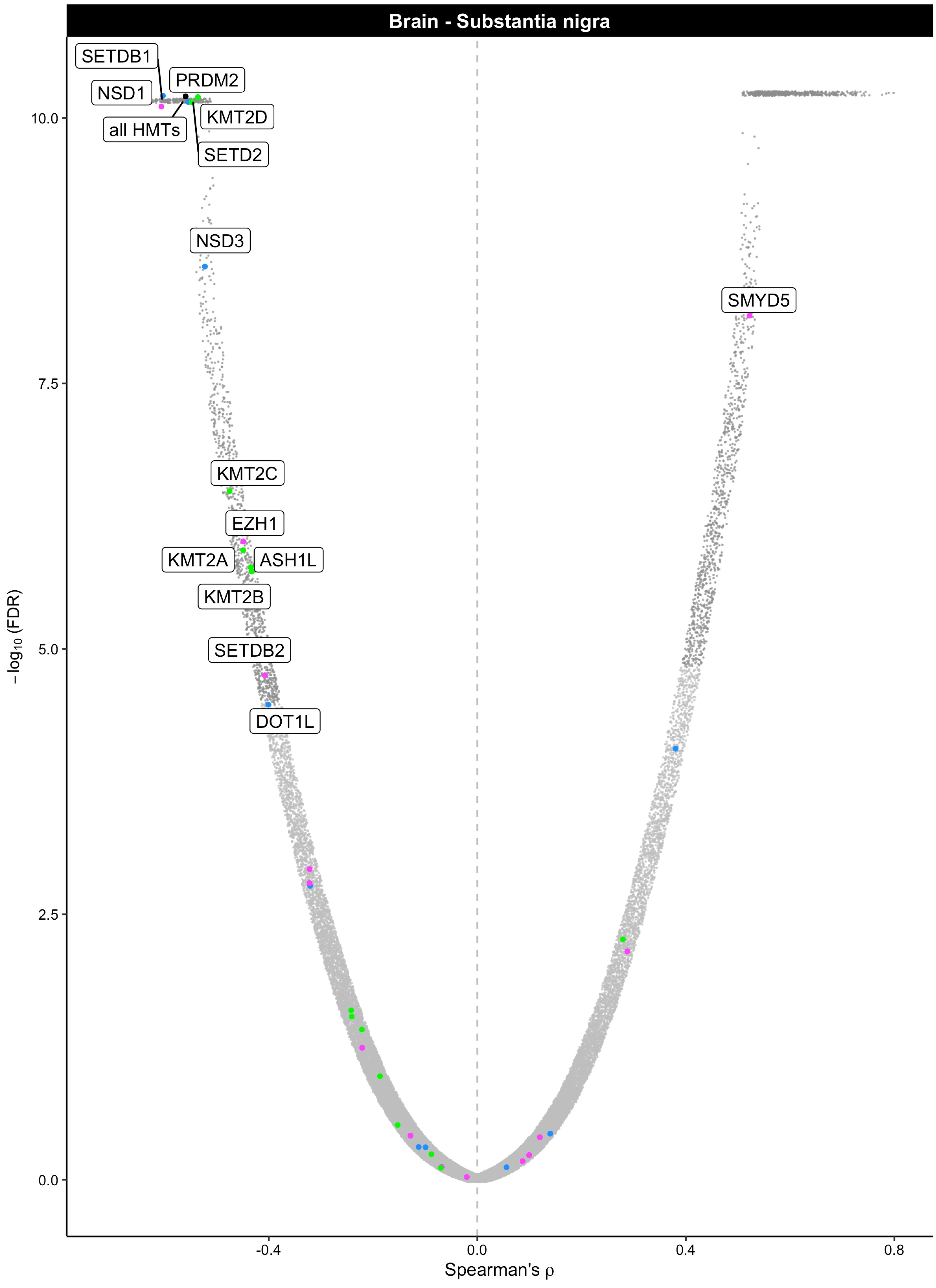

Brain - Substantia nigra
SETDB1
NSD1
PRDM2
all HMTs
KMT2D
SETD2
NSD3
SMYD5
KMT2C
EZH1
KMT2A
ASH1L
KMT2B
SETDB2
DOT1L
10.0
7.5
5.0
2.5
0.0
-0.4
0.0
0.4
0.8
−log10 (FDR)
Spearman's ρ

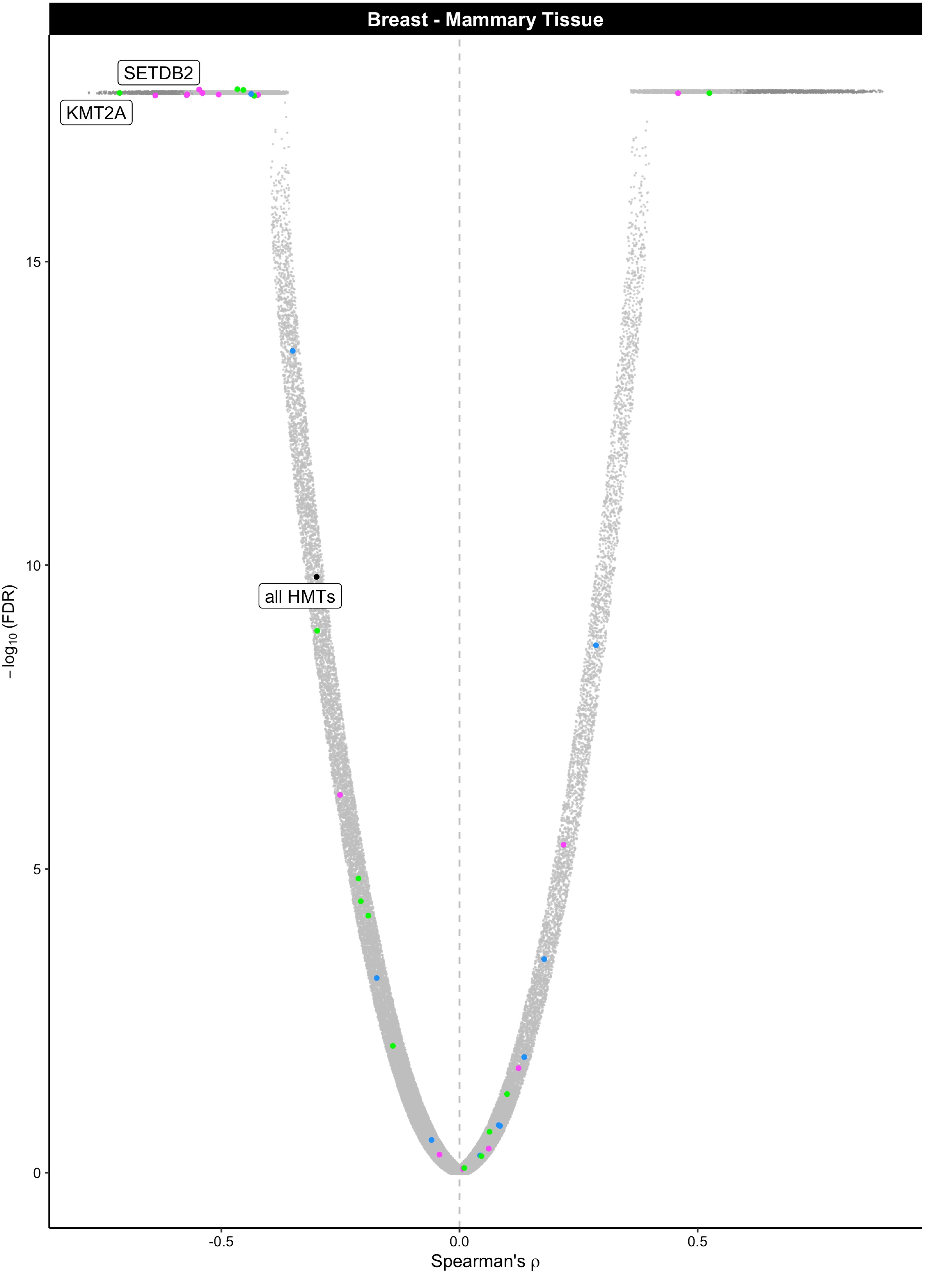

Breast - Mammary Tissue
SETDB2
KMT2A
all HMTs
15
10
5
0
−log10 (FDR)
-0.5
0.0
0.5
Spearman's ρ

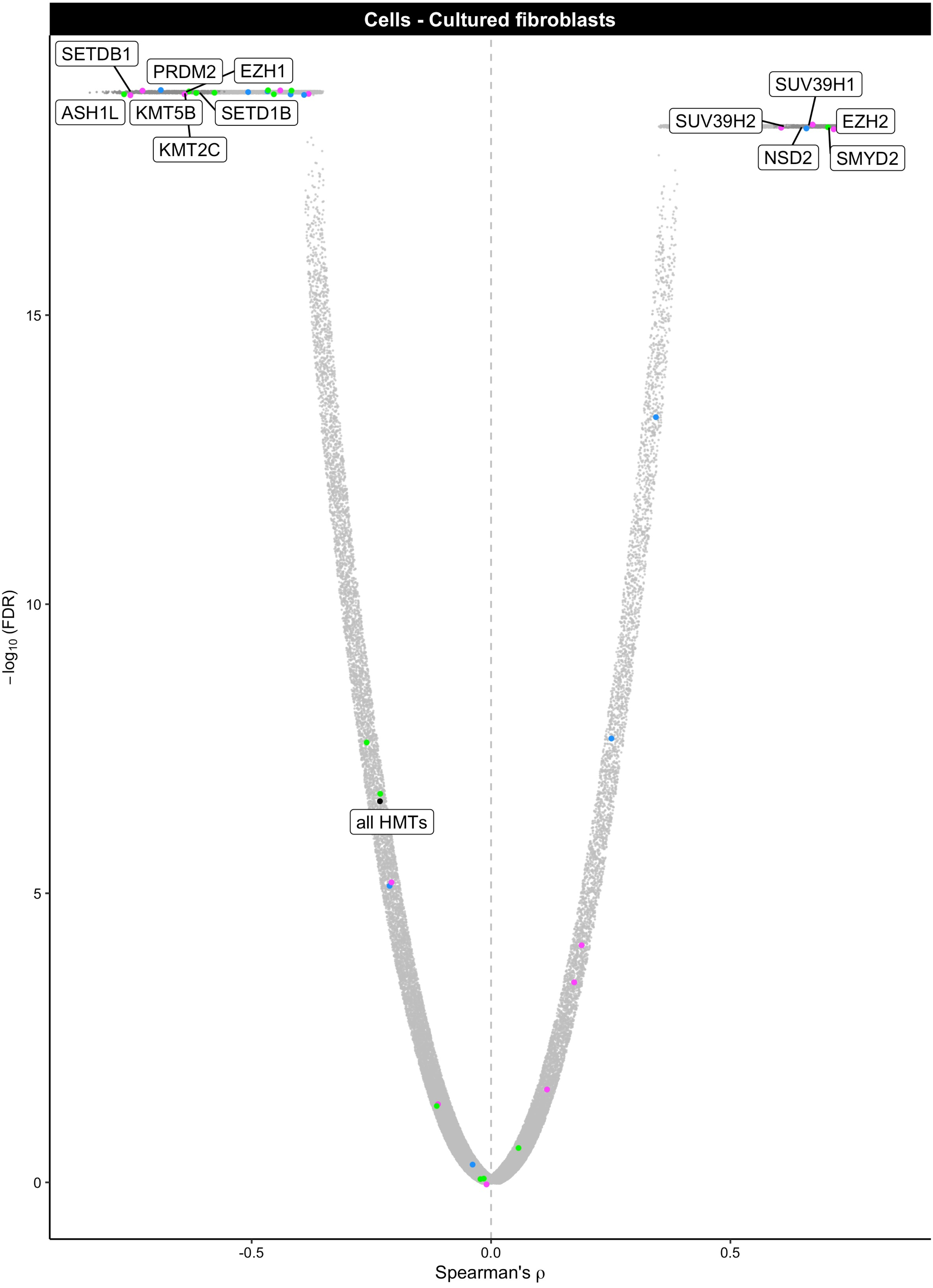

Cells - Cultured fibroblasts
SETDB1
PRDM2
EZH1
ASH1L
KMT5B
SETD1B
KMT2C
SUV39H1
SUV39H2
EZH2
NSD2
SMYD2
all HMTs
15
10
5
0
$-\log_{10}$ (FDR)
-0.5
0.0
0.5
Spearman's ρ

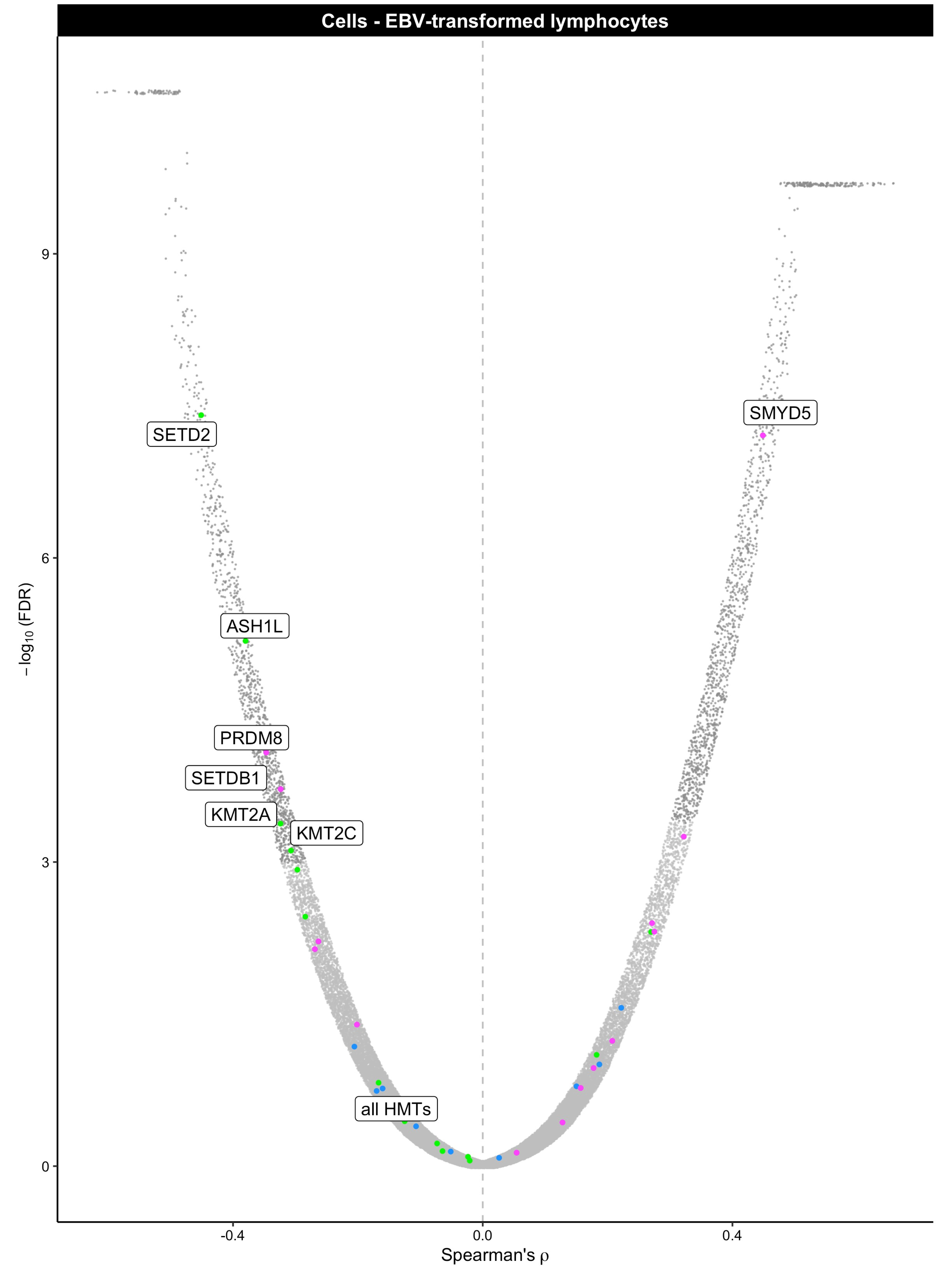
Cells - EBV-transformed lymphocytes
SETD2
SMYD5
ASH1L
PRDM8
SETDB1
KMT2A
KMT2C
all HMTs
9
6
3
0
−log10 (FDR)
-0.4
0.0
0.4
Spearman's ρ

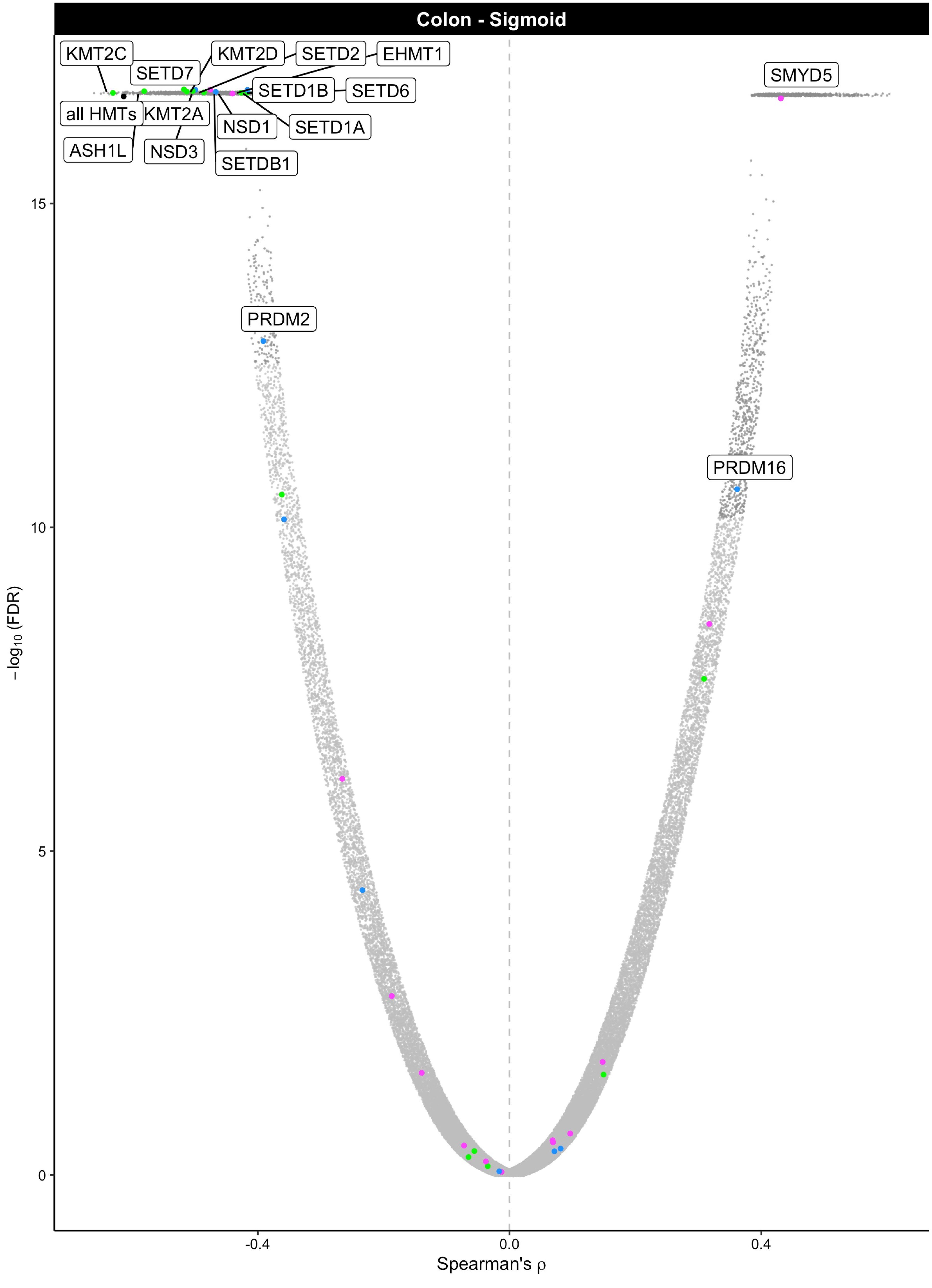

Colon - Sigmoid
KMT2C
SETD7
KMT2D
SETD2
EHMT1
SMYD5
SETD1B
SETD6
all HMTs
KMT2A
NSD1
SETD1A
ASH1L
NSD3
SETDB1
PRDM2
PRDM16
15
10
5
0
−log10 (FDR)
-0.4
0.0
0.4
Spearman's ρ

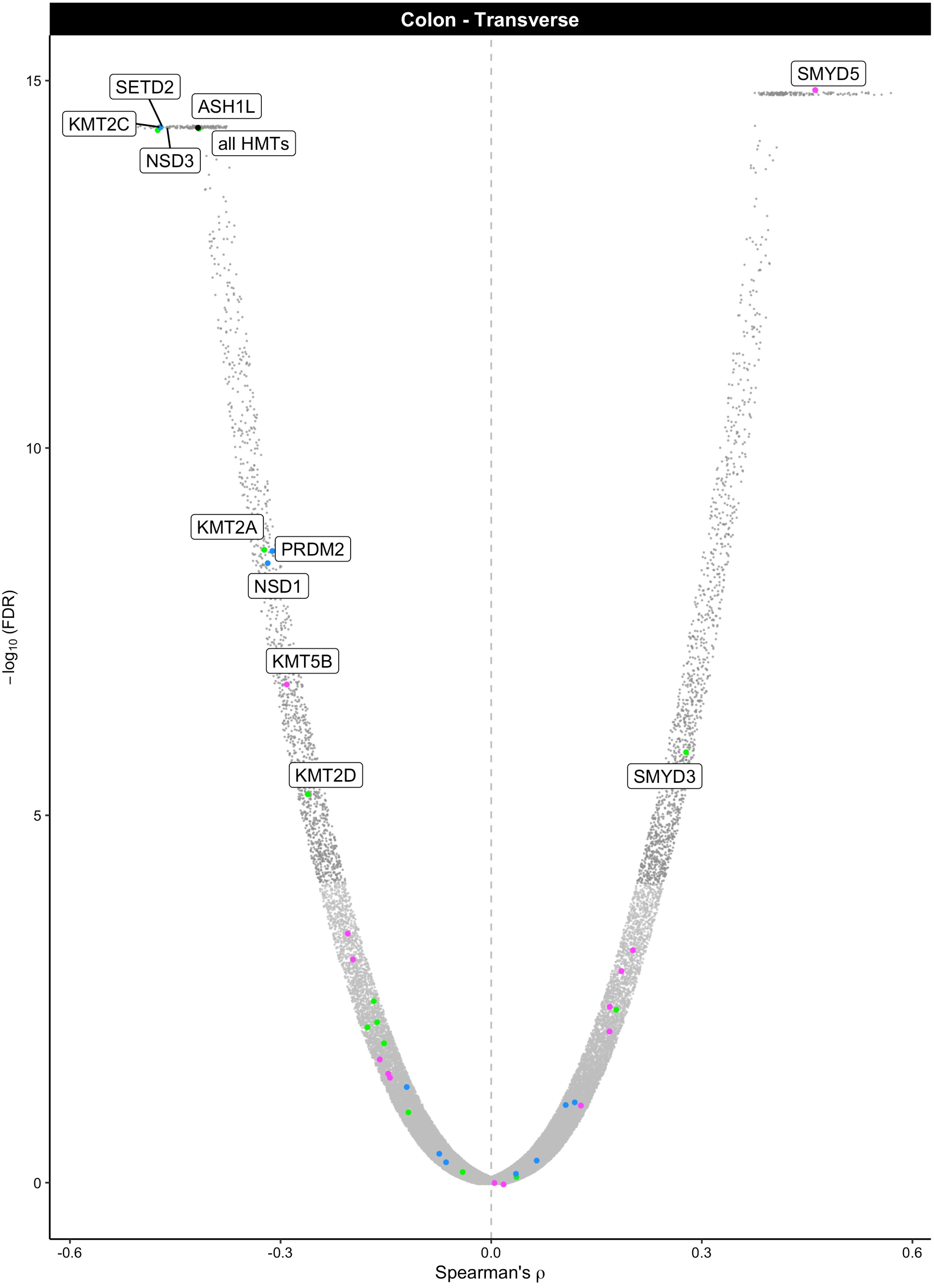

Colon - Transverse
SMYD5
SETD2
ASH1L
KMT2C
all HMTs
NSD3
KMT2A
PRDM2
NSD1
KMT5B
KMT2D
SMYD3
15
10
5
0
−log10 (FDR)
-0.6
-0.3
0.0
0.3
0.6
Spearman's ρ

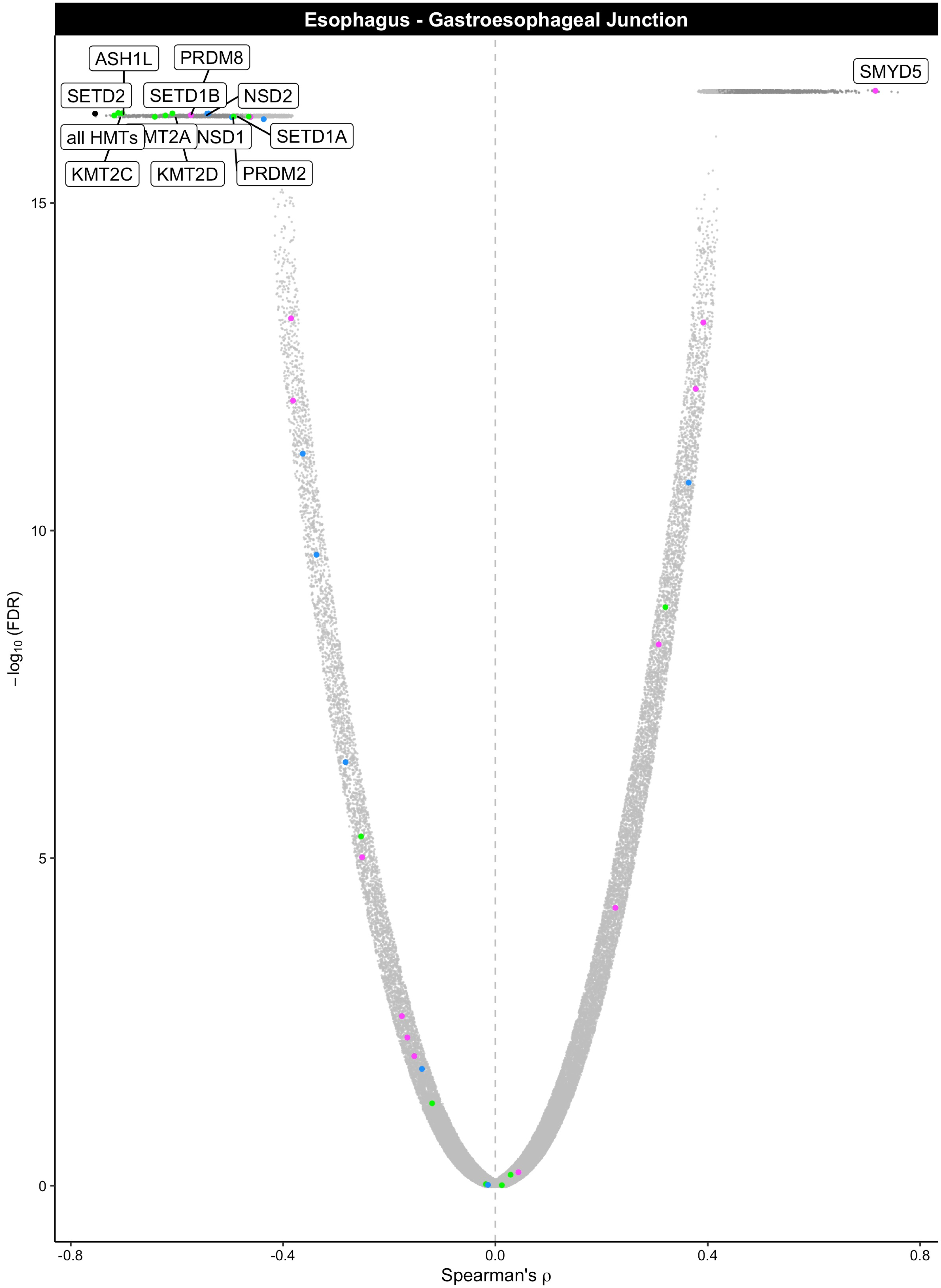
Esophagus - Gastroesophageal Junction
ASH1L
PRDM8
SETD2
SETD1B
NSD2
SMYD5
all HMTs
KMT2A
NSD1
SETD1A
KMT2C
KMT2D
PRDM2
15
10
5
0
−log10 (FDR)
-0.8
-0.4
0.0
0.4
0.8
Spearman's ρ

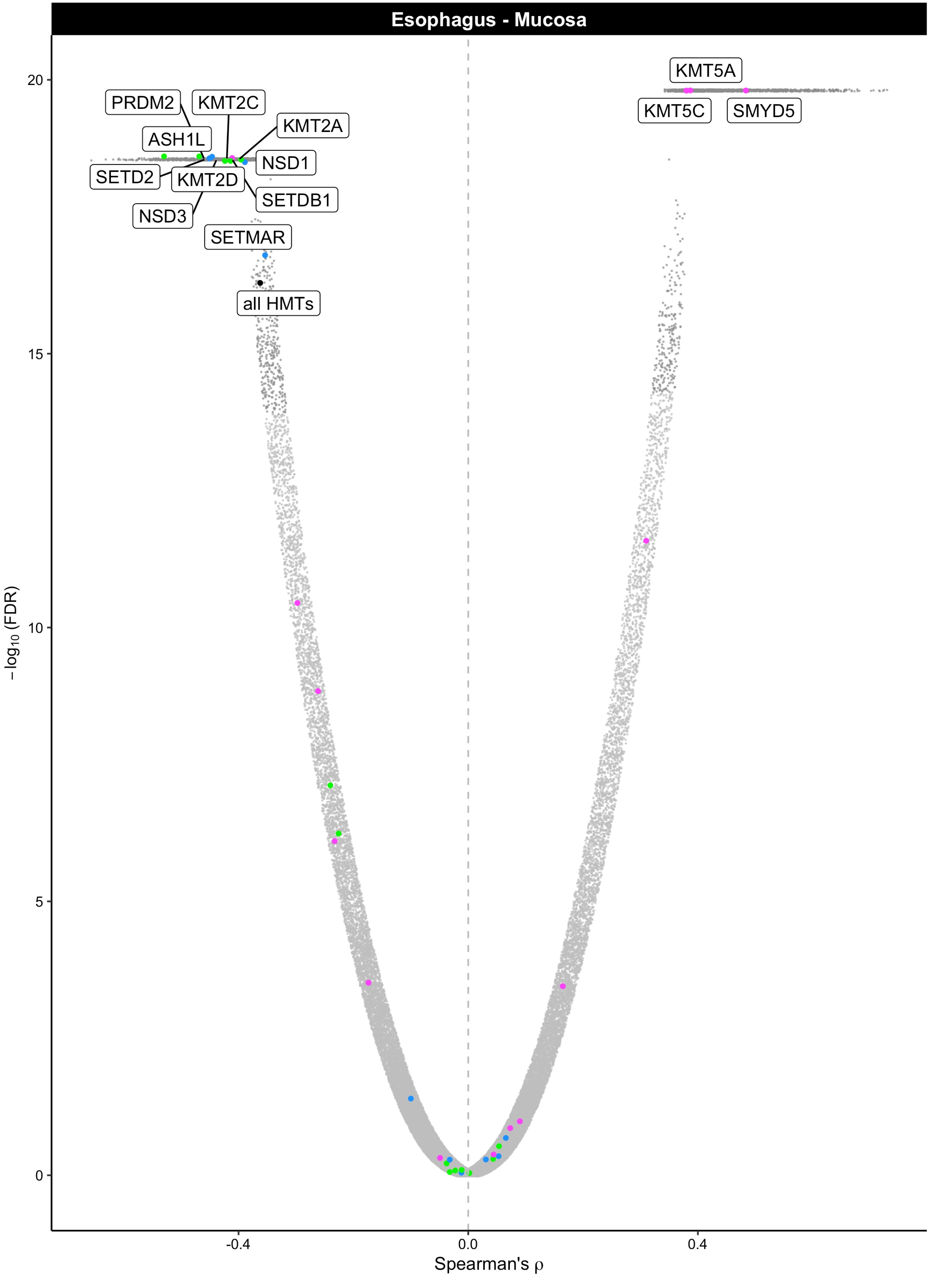

Esophagus - Mucosa
KMT5A
KMT5C
SMYD5
PRDM2
KMT2C
KMT2A
ASH1L
NSD1
SETD2
KMT2D
SETDB1
NSD3
SETMAR
all HMTs
20
15
10
5
0
−log10 (FDR)
−0.4
0.0
0.4
Spearman's ρ

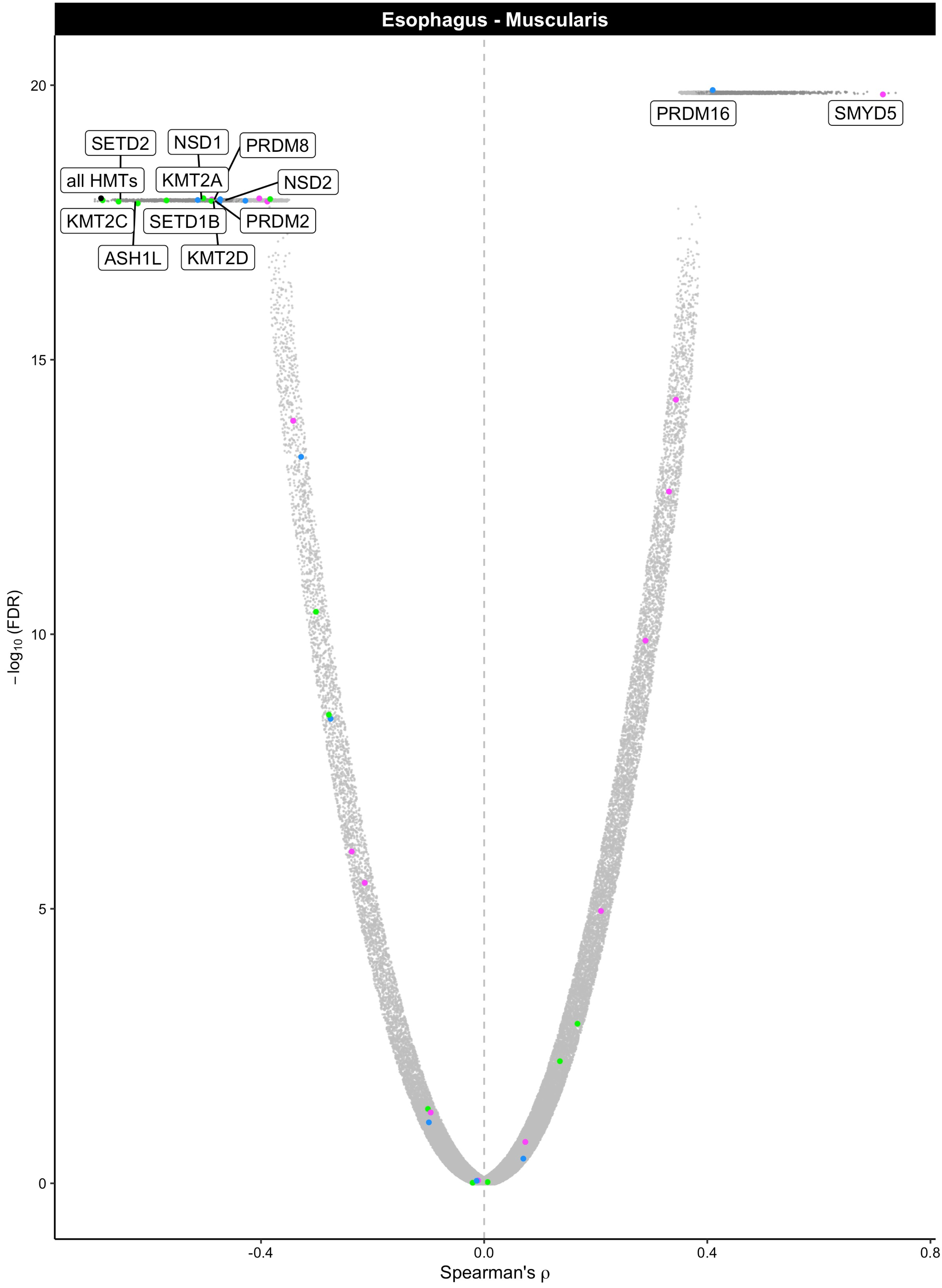

Esophagus - Muscularis
PRDM16
SMYD5
SETD2
NSD1
PRDM8
all HMTs
KMT2A
NSD2
KMT2C
SETD1B
PRDM2
ASH1L
KMT2D
20
15
10
5
0
−log10 (FDR)
-0.4
0.0
0.4
0.8
Spearman's ρ

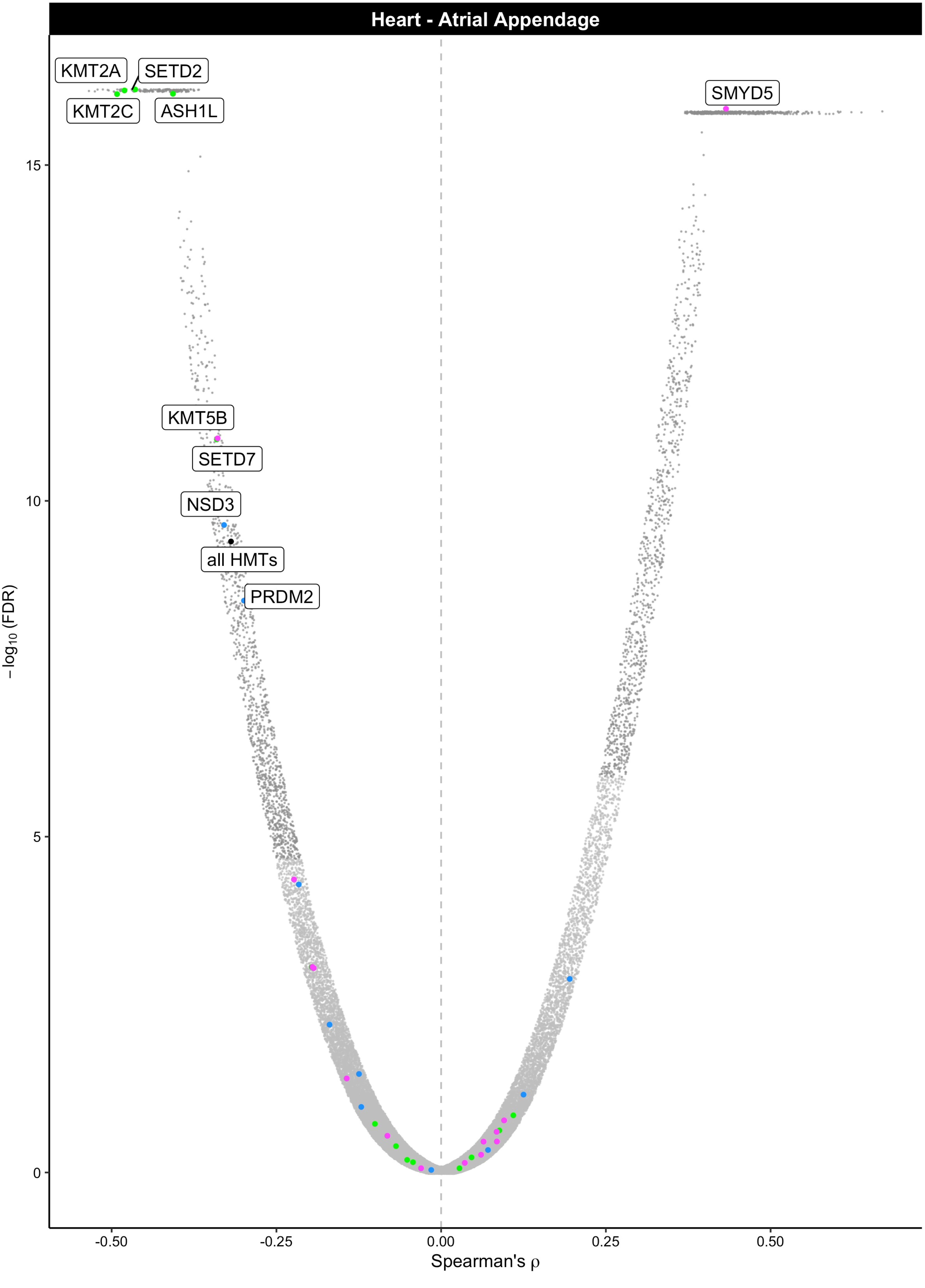

Heart - Atrial Appendage
KMT2A
SETD2
KMT2C
ASH1L
SMYD5
KMT5B
SETD7
NSD3
all HMTs
PRDM2
−log10 (FDR)
15
10
5
0
-0.50
-0.25
0.00
0.25
0.50
Spearman's ρ

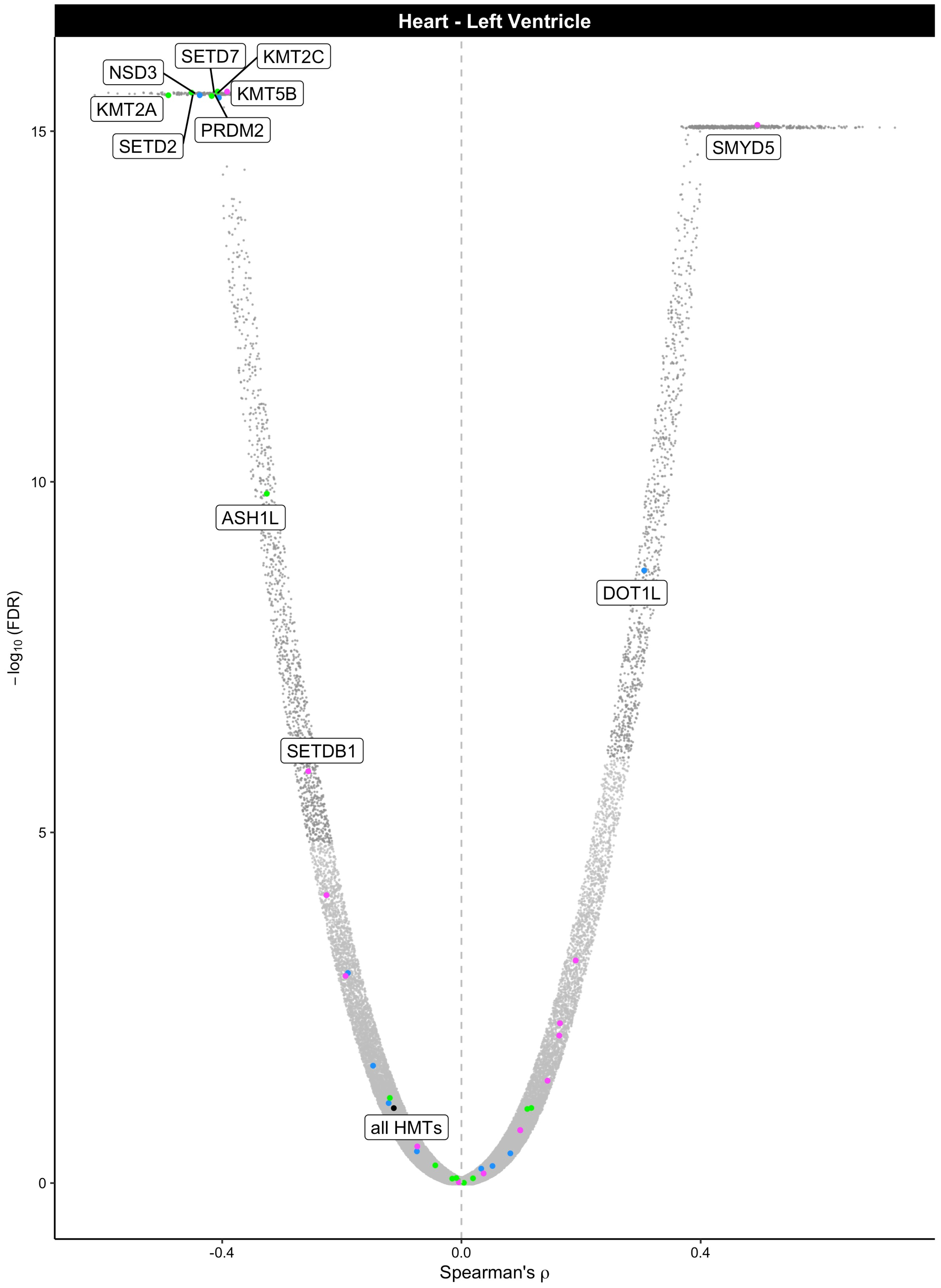

Heart - Left Ventricle
NSD3
SETD7
KMT2C
KMT5B
KMT2A
PRDM2
SETD2
SMYD5
ASH1L
DOT1L
SETDB1
all HMTs
15
10
5
0
−log10 (FDR)
-0.4
0.0
0.4
Spearman's ρ

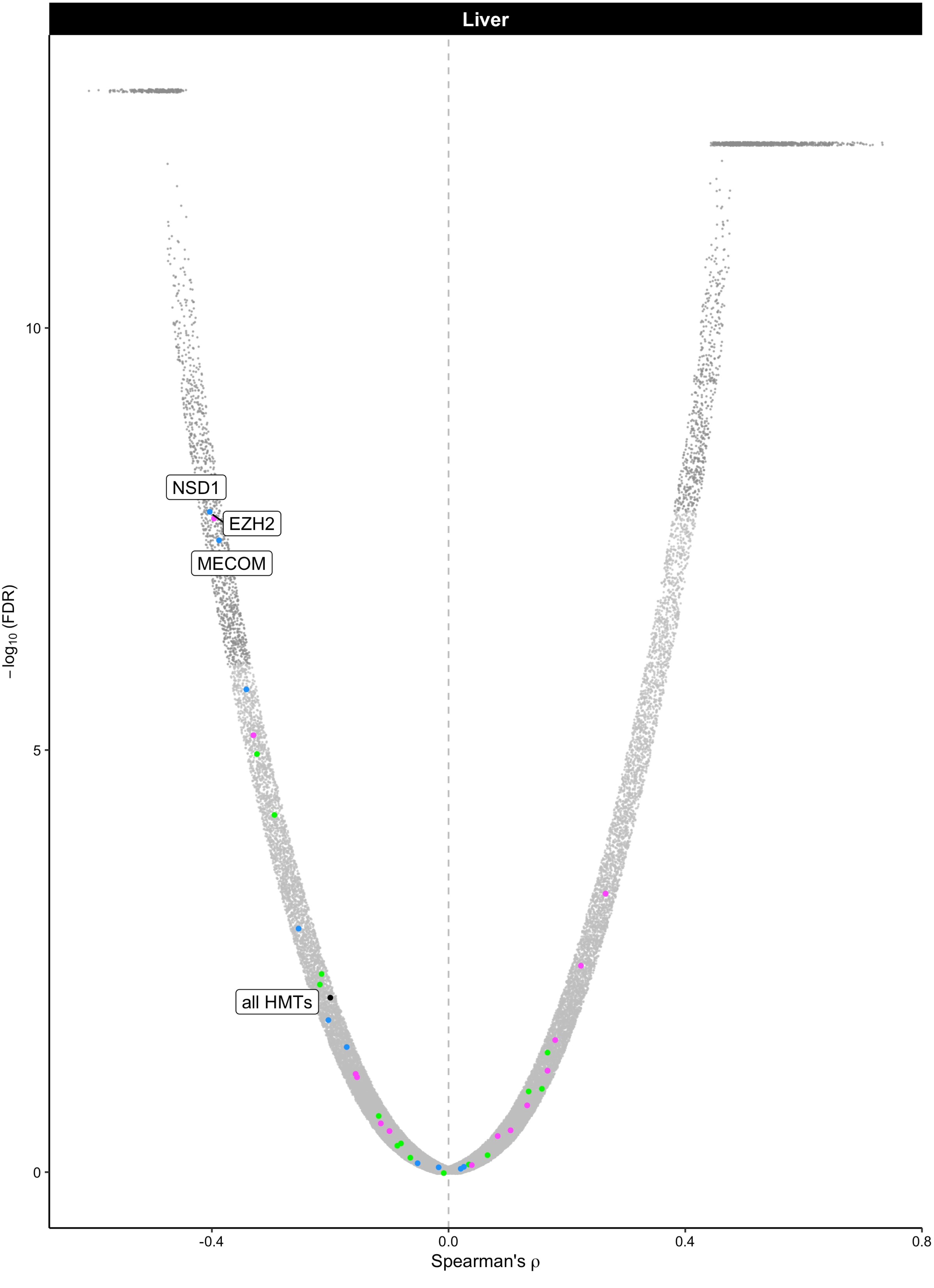
Liver
NSD1
EZH2
MECOM
all HMTs
−log₁₀ (FDR)
10
5
0
-0.4
0.0
0.4
0.8
Spearman's ρ

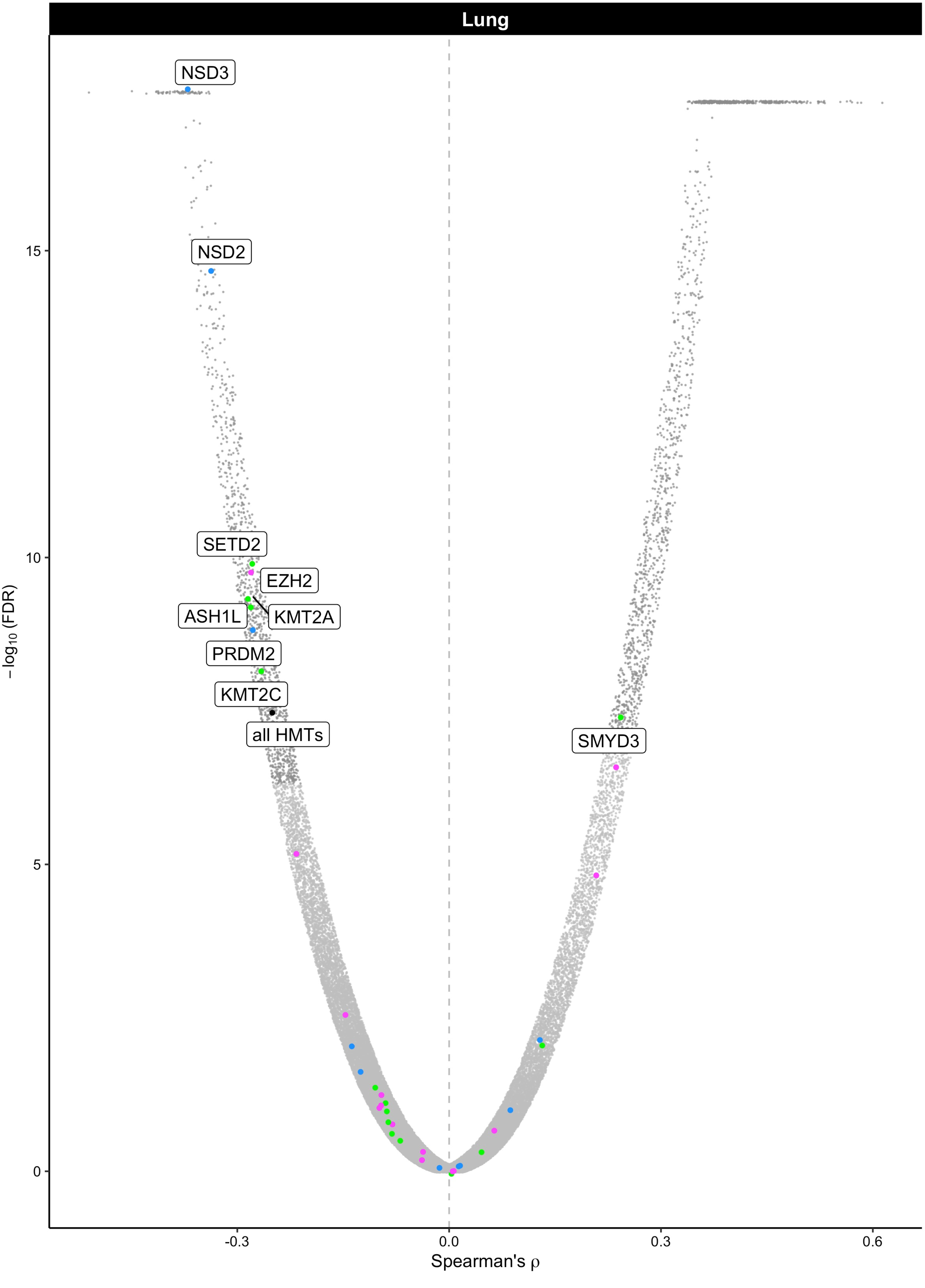

Lung
NSD3
NSD2
SETD2
EZH2
ASH1L
KMT2A
PRDM2
KMT2C
all HMTs
SMYD3
15
10
5
0
−log$_{10}$ (FDR)
-0.3
0.0
0.3
0.6
Spearman's ρ

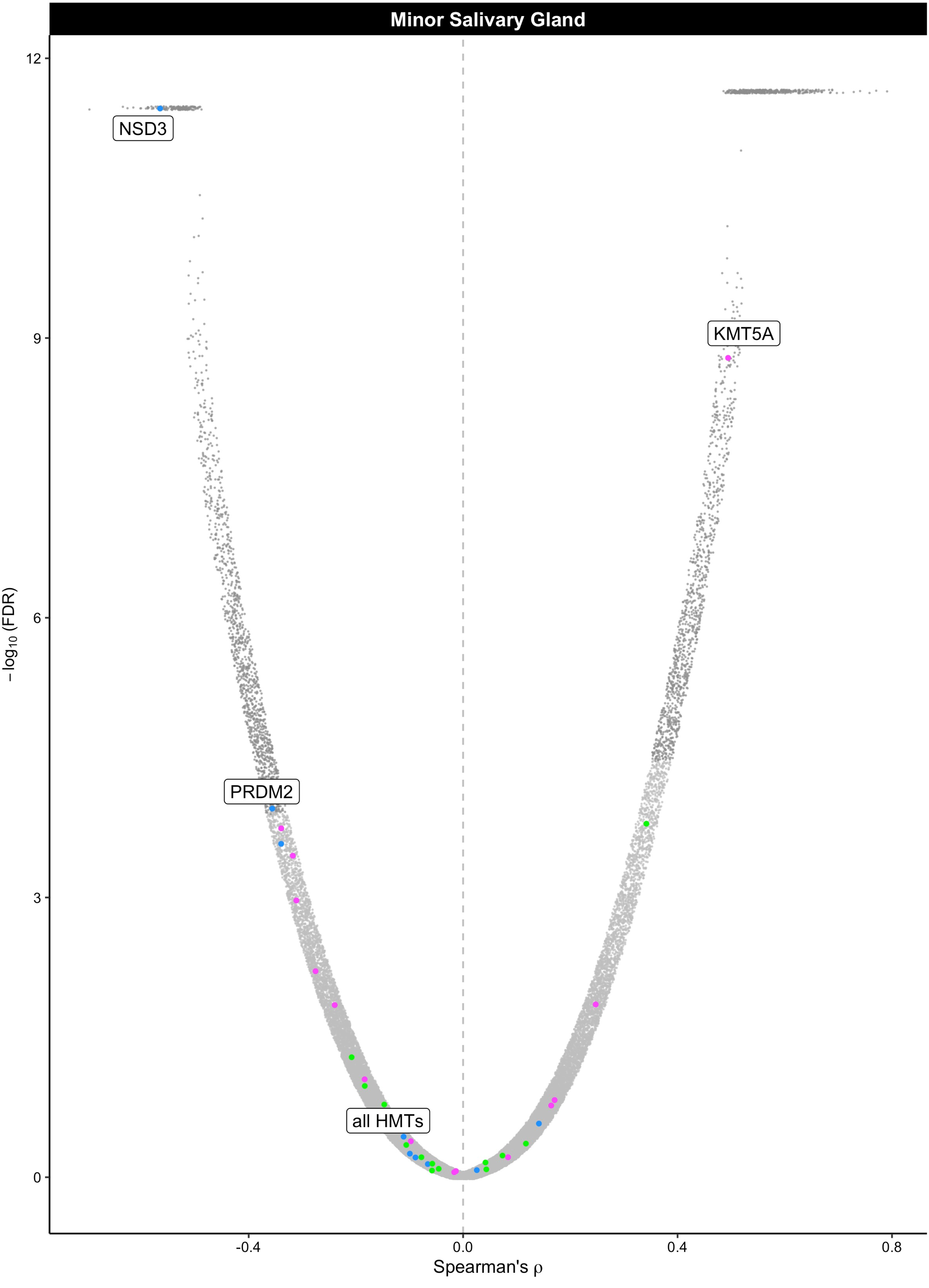

Minor Salivary Gland
NSD3
KMT5A
PRDM2
all HMTs
12
9
6
3
0
-0.4
0.0
0.4
0.8
Spearman's ρ
−log10 (FDR)

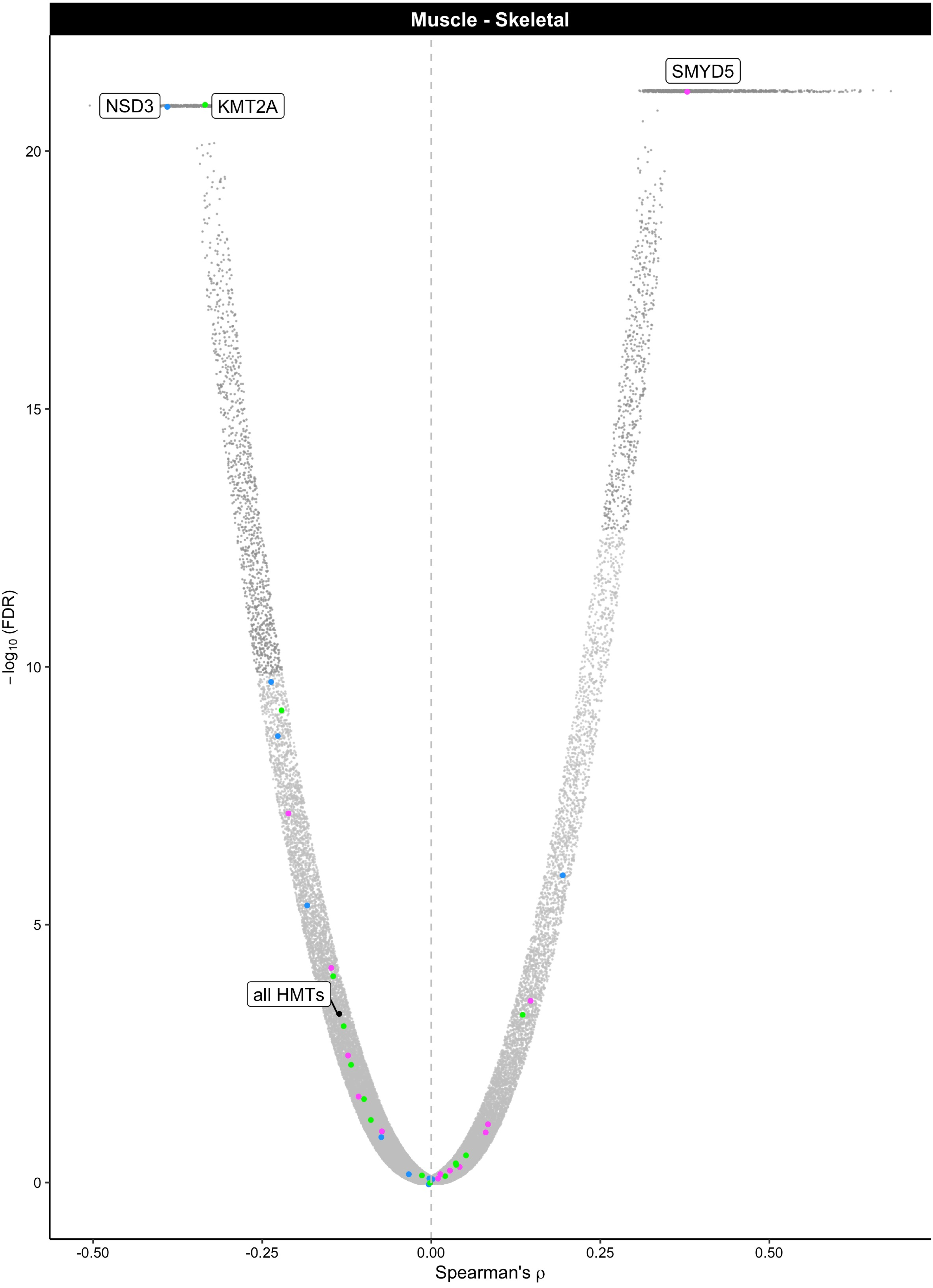
Muscle - Skeletal
SMYD5
NSD3
KMT2A
all HMTs
20
15
10
5
0
−log10 (FDR)
-0.50
-0.25
0.00
0.25
0.50
Spearman's ρ

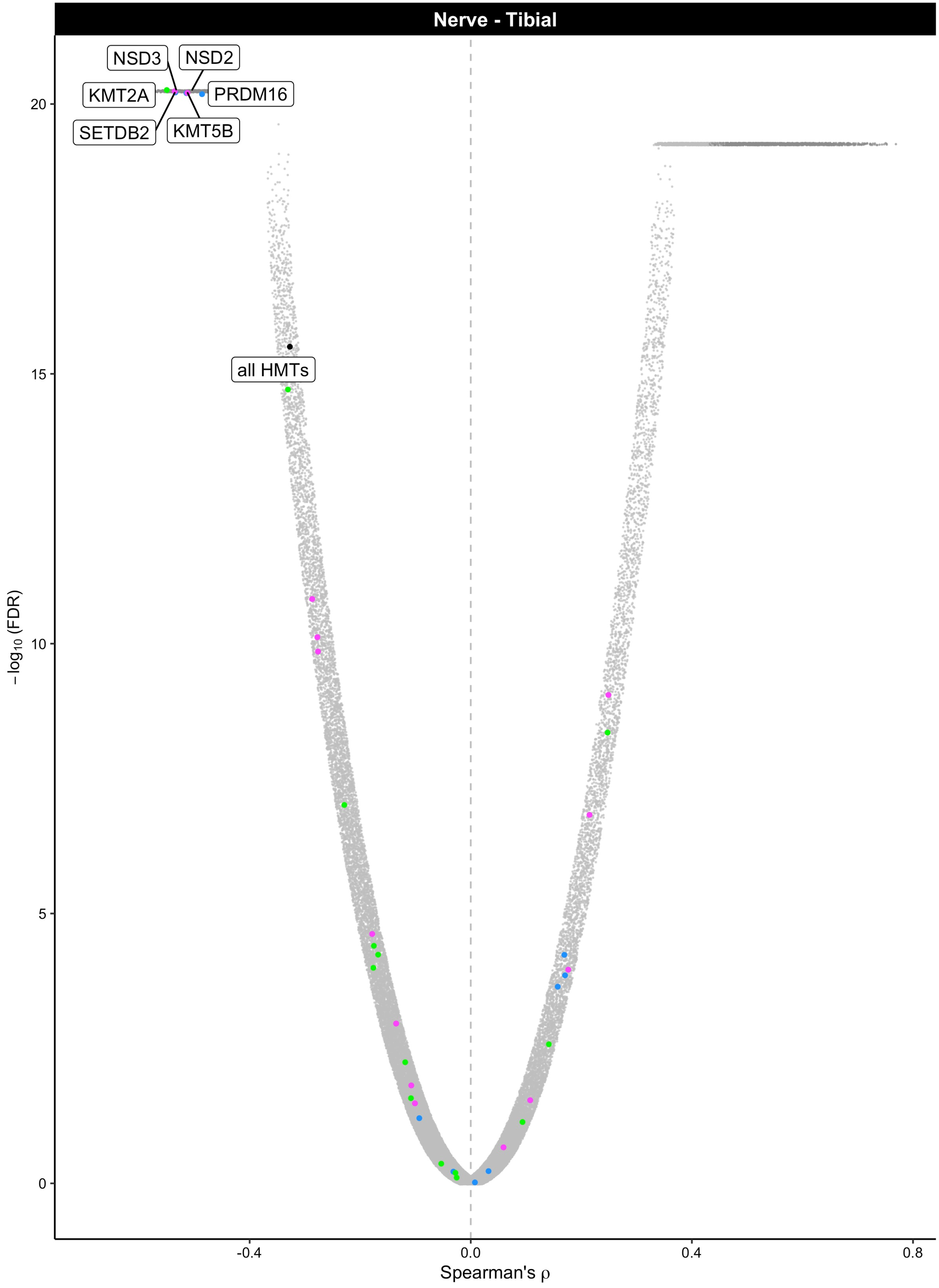

Nerve - Tibial
NSD3
NSD2
KMT2A
PRDM16
SETDB2
KMT5B
all HMTs
20
15
10
5
0
−log10 (FDR)
-0.4
0.0
0.4
0.8
Spearman's ρ

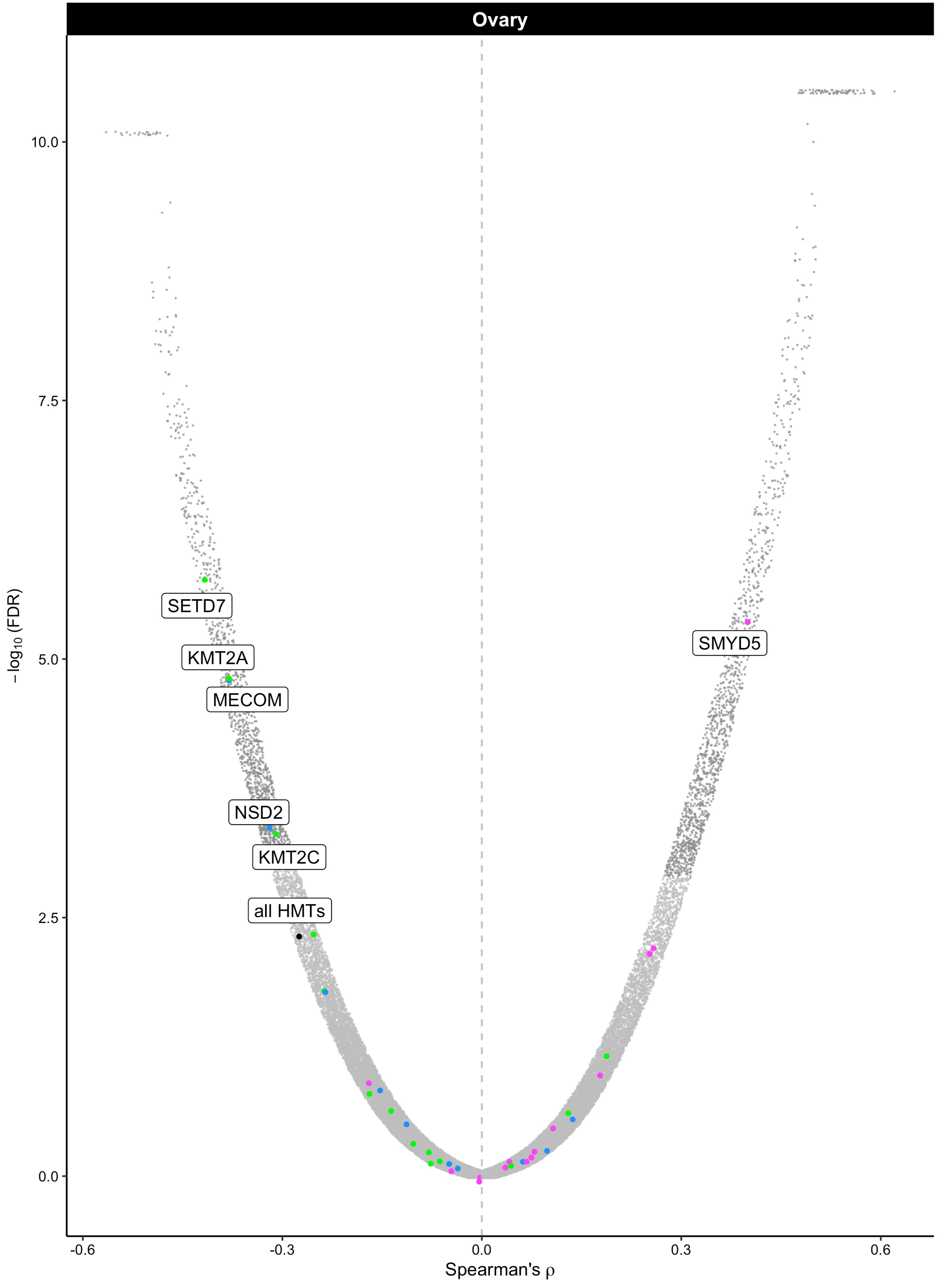

Ovary
SETD7
KMT2A
MECOM
NSD2
KMT2C
all HMTs
SMYD5
10.0
7.5
5.0
2.5
0.0
−log10 (FDR)
-0.6
-0.3
0.0
0.3
0.6
Spearman's ρ

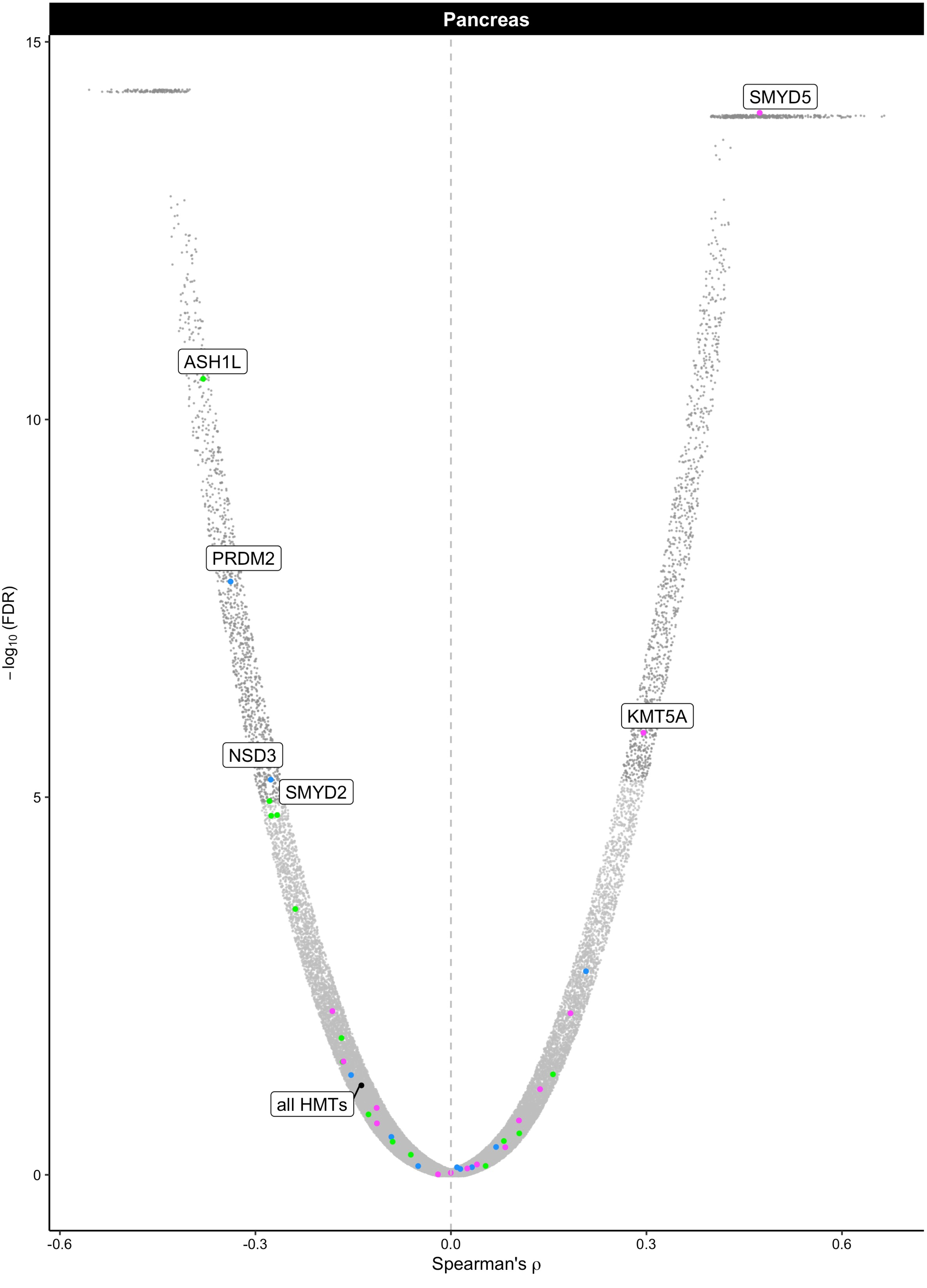
Pancreas
15
10
5
0
−log10 (FDR)
SMYD5
ASH1L
PRDM2
KMT5A
NSD3
SMYD2
all HMTs
-0.6
-0.3
0.0
0.3
0.6
Spearman's ρ

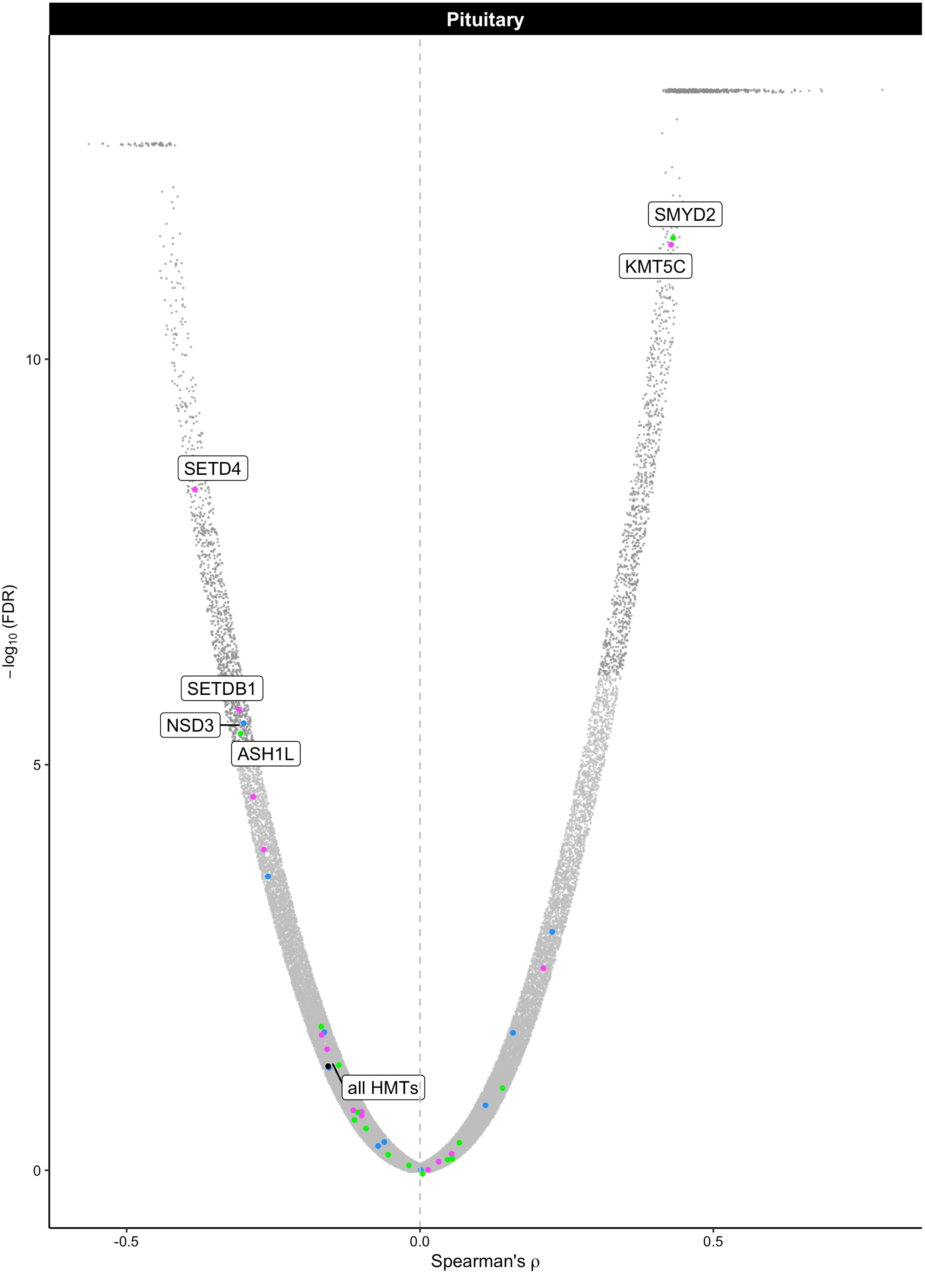

Pituitary
SMYD2
KMT5C
SETD4
SETDB1
NSD3
ASH1L
all HMTs
10
5
0
−log10 (FDR)
-0.5
0.0
0.5
Spearman's ρ

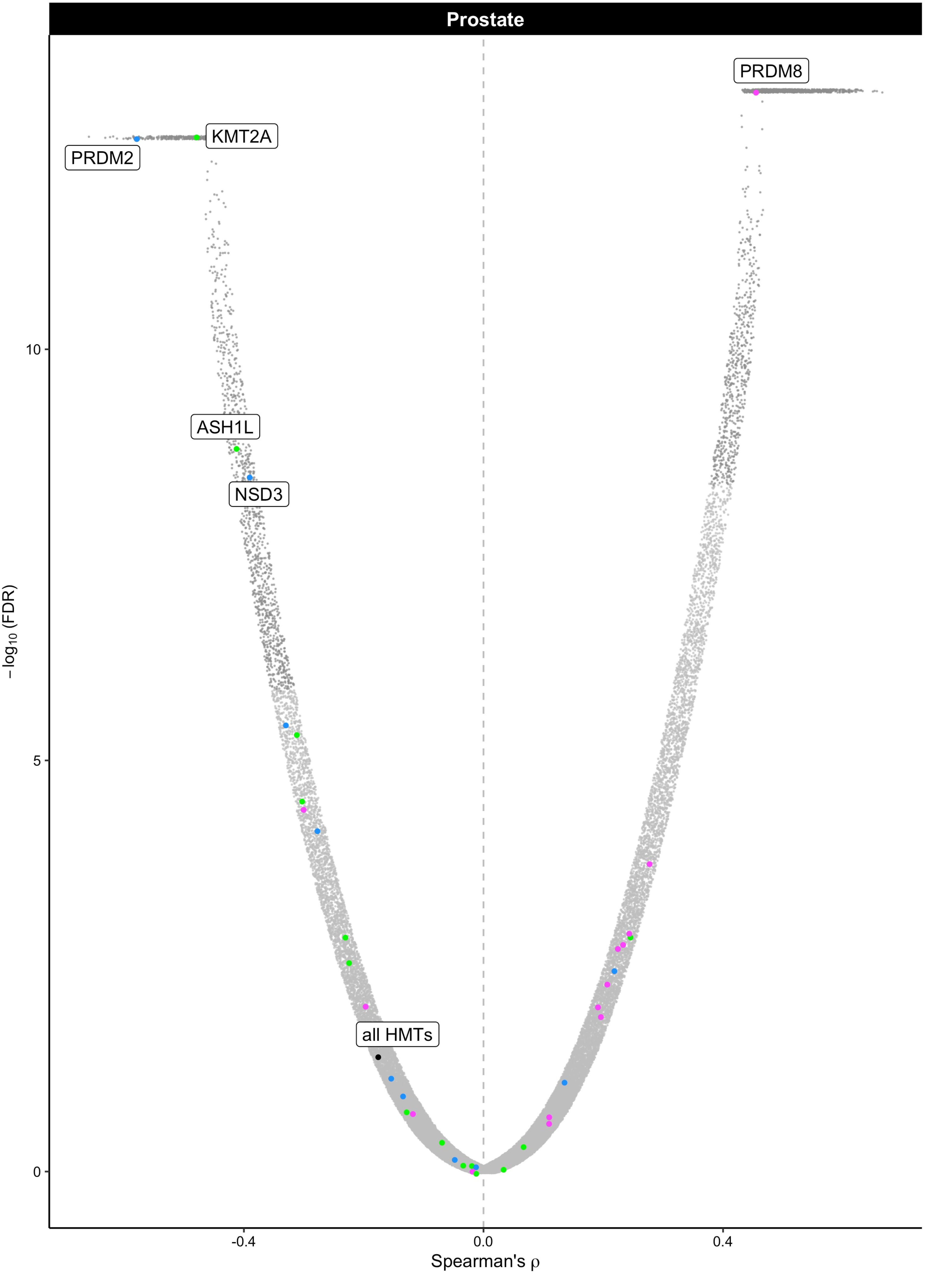

Prostate
PRDM8
KMT2A
PRDM2
ASH1L
NSD3
all HMTs
10
5
0
−log10 (FDR)
-0.4
0.0
0.4
Spearman's ρ

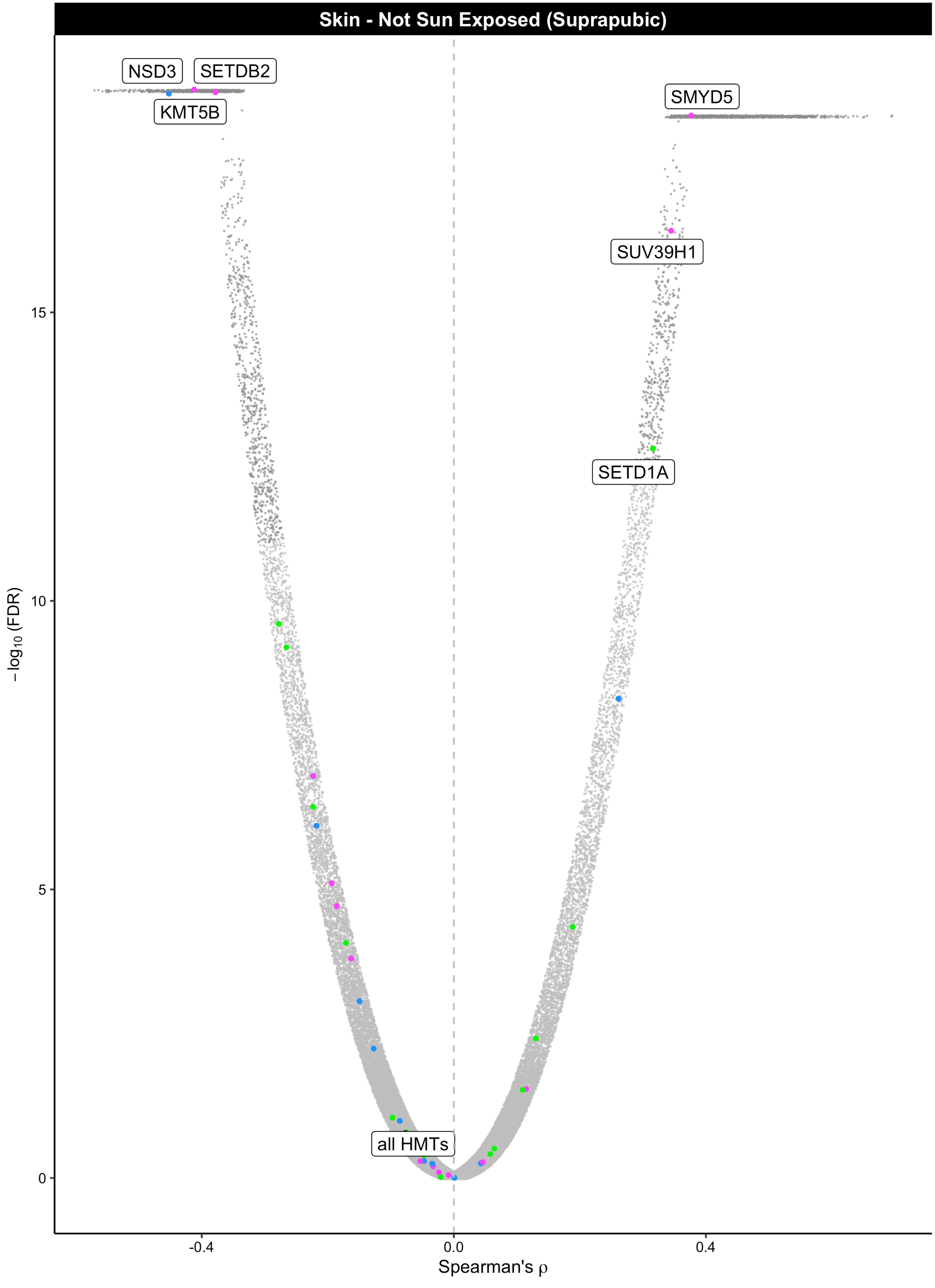

Skin - Not Sun Exposed (Suprapubic)
NSD3
SETDB2
KMT5B
SMYD5
SUV39H1
SETD1A
all HMTs
15
10
5
0
$-\log_{10}$ (FDR)
-0.4
0.0
0.4
Spearman's ρ

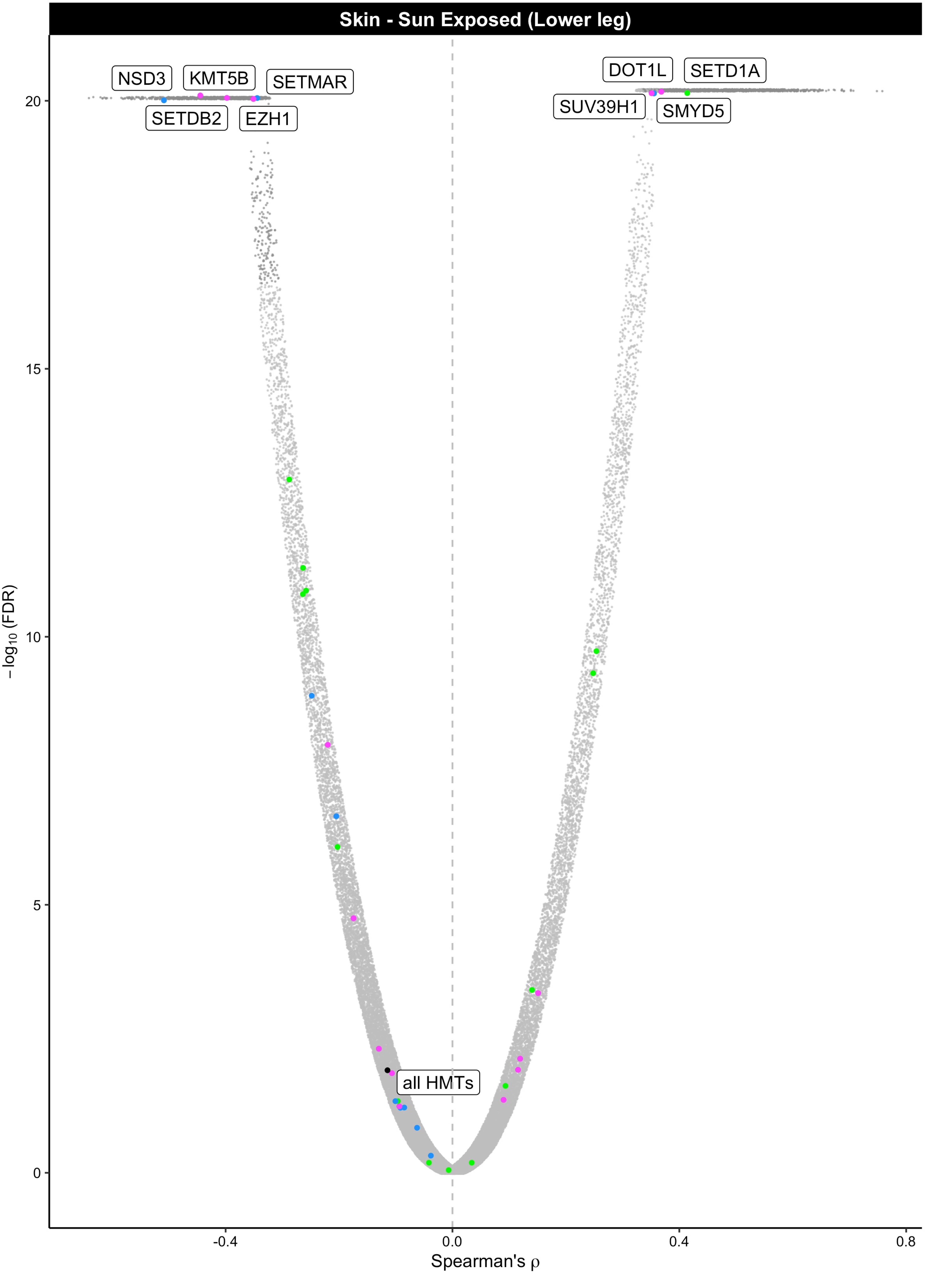

Skin - Sun Exposed (Lower leg)
NSD3
KMT5B
SETMAR
SETDB2
EZH1
DOT1L
SETD1A
SUV39H1
SMYD5
all HMTs
20
15
10
5
0
−log10 (FDR)
-0.4
0.0
0.4
0.8
Spearman's ρ

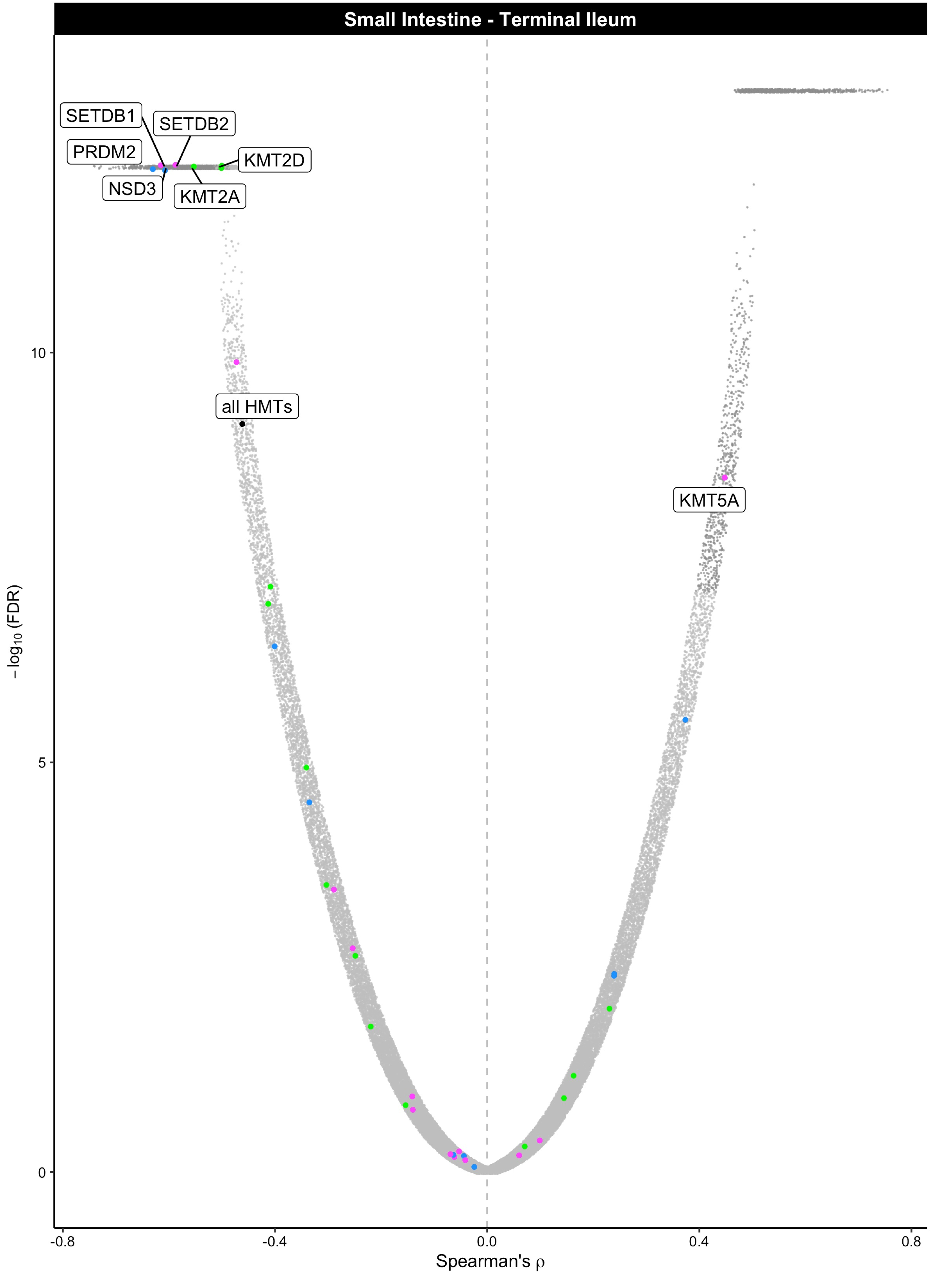
Small Intestine - Terminal Ileum
SETDB1
SETDB2
PRDM2
KMT2D
NSD3
KMT2A
all HMTs
KMT5A
10
5
0
−log10 (FDR)
-0.8
-0.4
0.0
0.4
0.8
Spearman's ρ

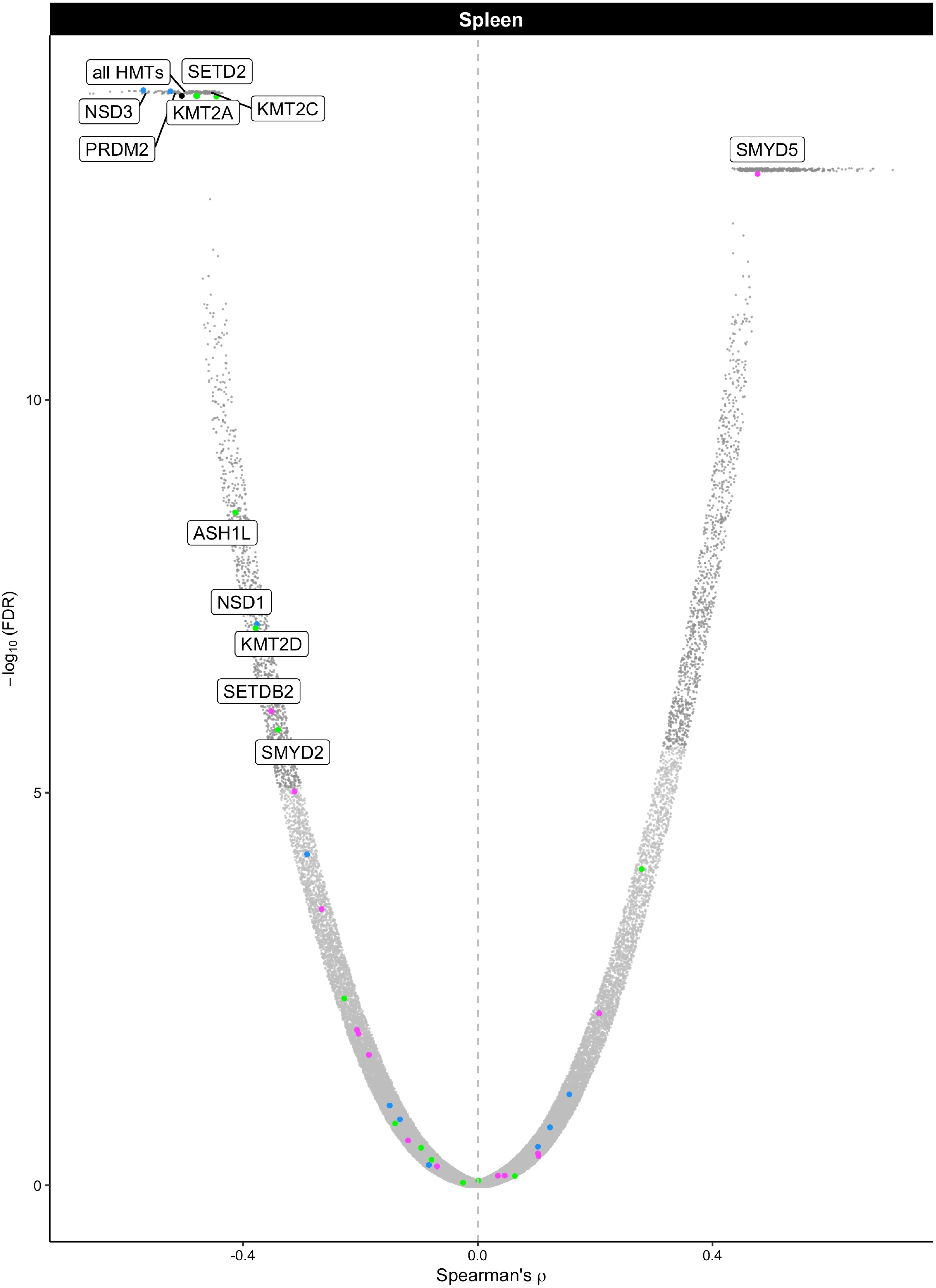

Spleen
all HMTs
SETD2
NSD3
KMT2A
KMT2C
PRDM2
SMYD5
ASH1L
NSD1
KMT2D
SETDB2
SMYD2
10
5
0
−log10 (FDR)
-0.4
0.0
0.4
Spearman's ρ

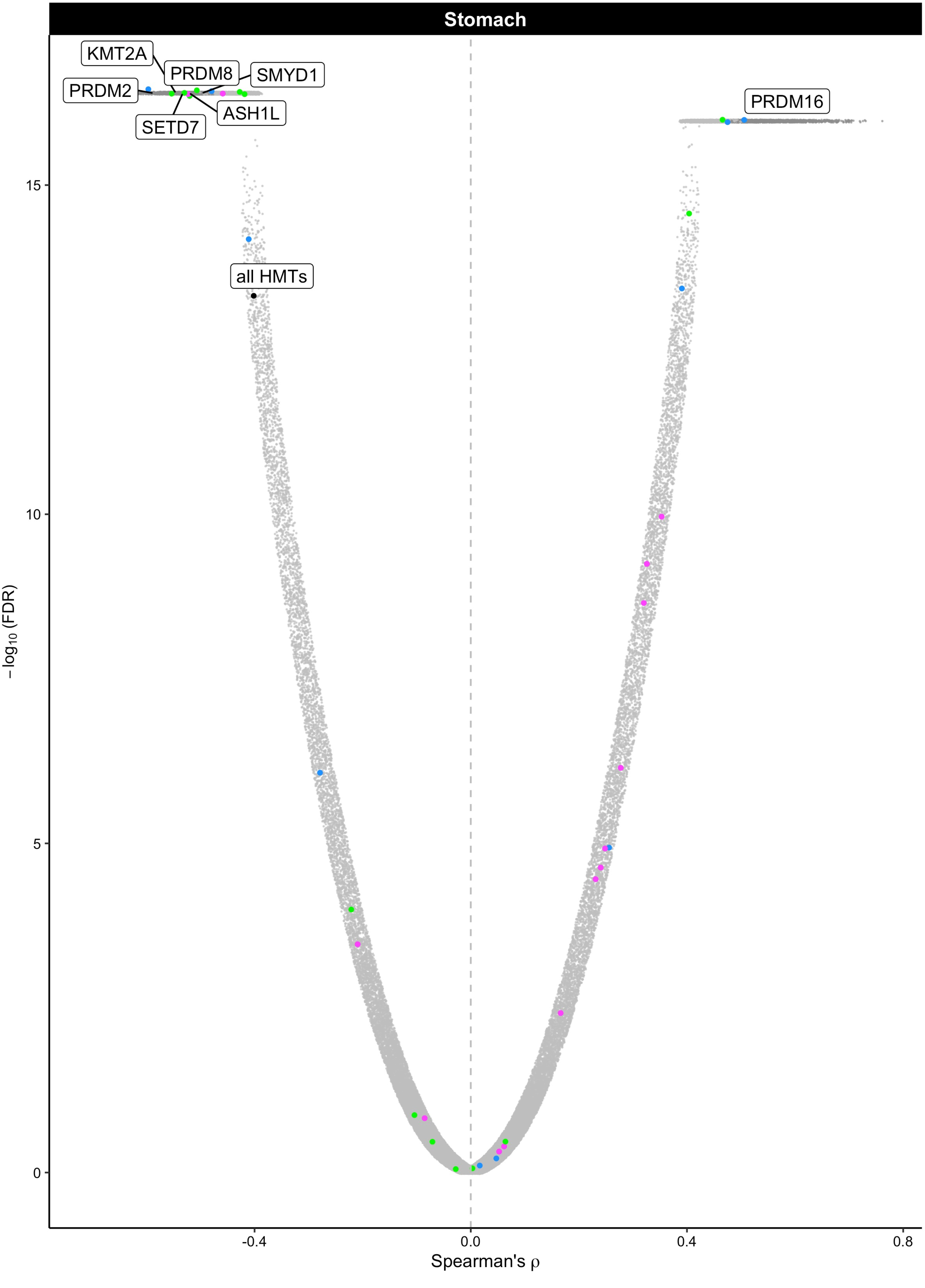

Stomach
KMT2A
PRDM8
SMYD1
PRDM2
ASH1L
SETD7
PRDM16
all HMTs
15
10
5
0
−log10 (FDR)
-0.4
0.0
0.4
0.8
Spearman's ρ

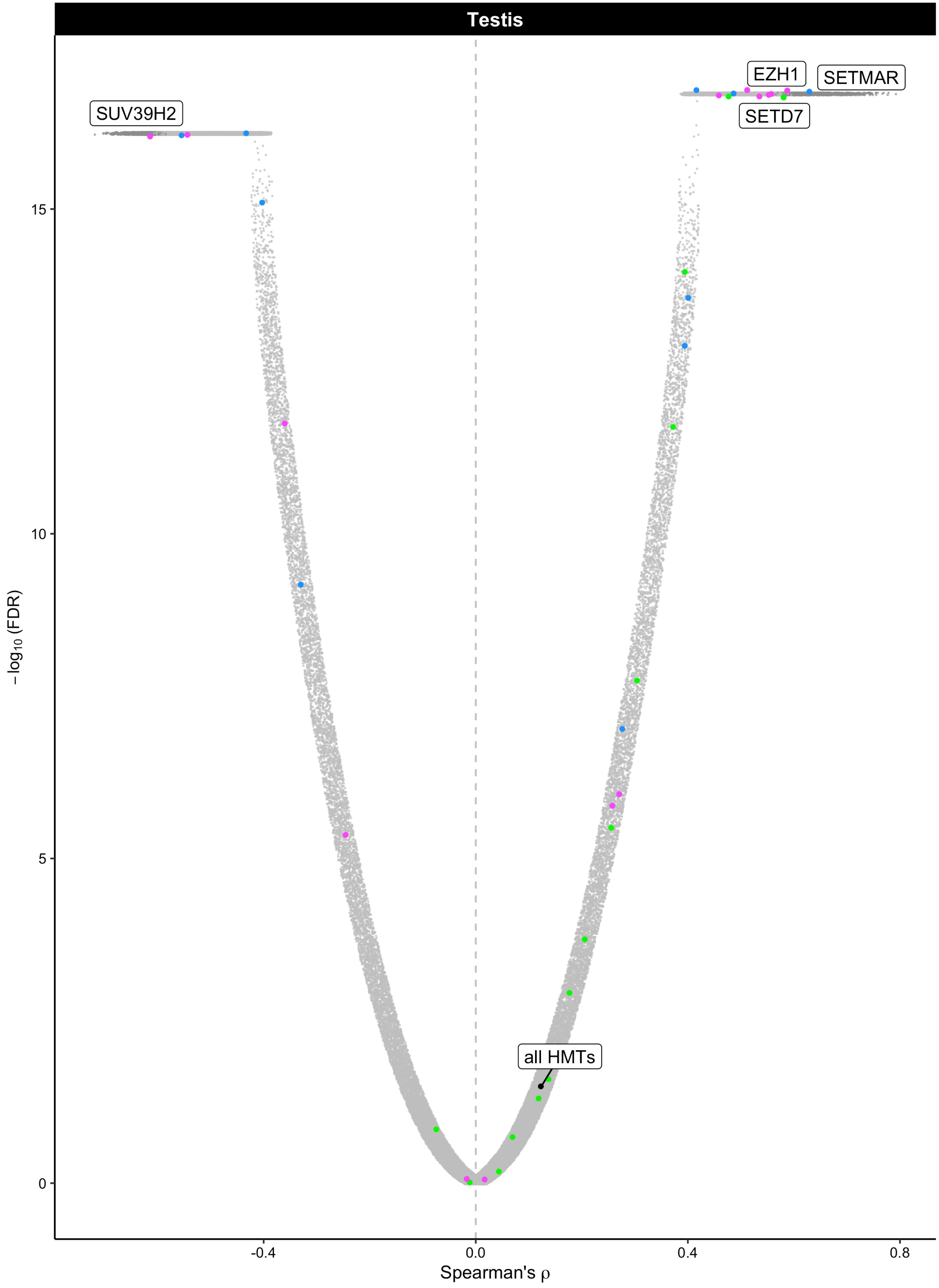

Testis
EZH1
SETMAR
SUV39H2
SETD7
all HMTs
15
10
5
0
−log10 (FDR)
-0.4
0.0
0.4
0.8
Spearman's ρ

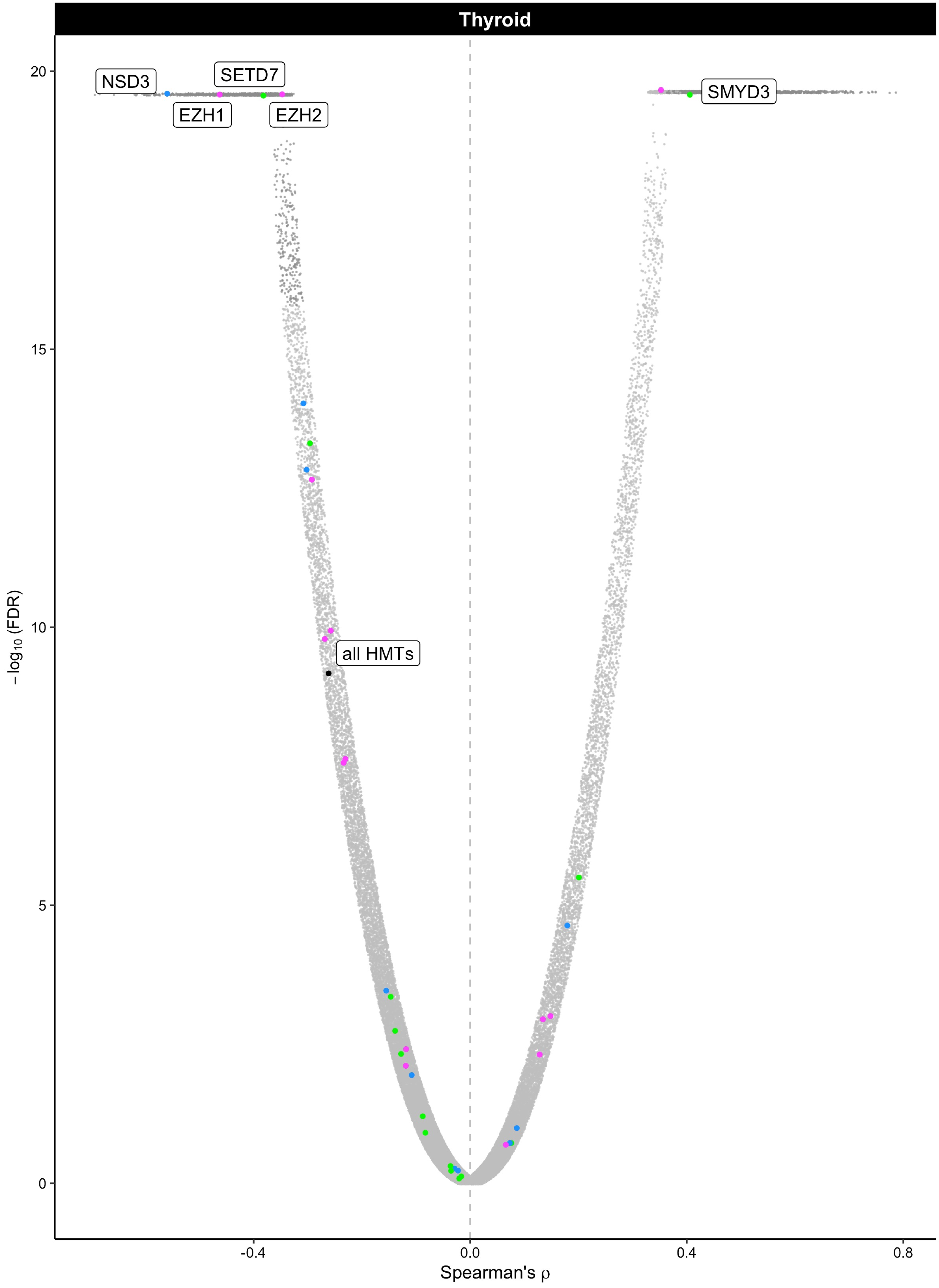

Thyroid
NSD3
SETD7
EZH1
EZH2
SMYD3
all HMTs
20
15
10
5
0
−log10 (FDR)
-0.4
0.0
0.4
0.8
Spearman's ρ

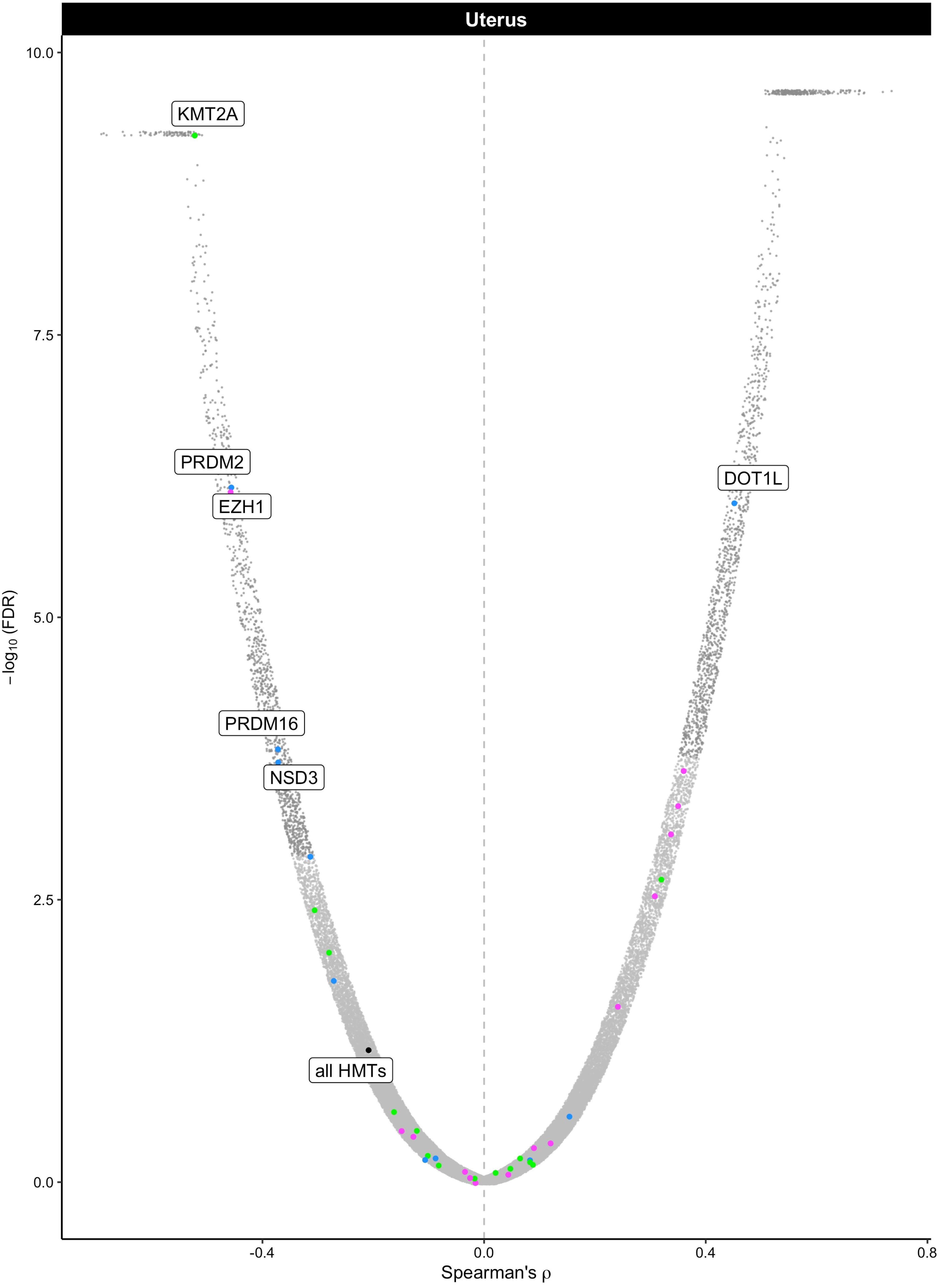

Uterus
KMT2A
PRDM2
EZH1
PRDM16
NSD3
DOT1L
all HMTs
10.0
7.5
5.0
2.5
0.0
−log10 (FDR)
-0.4
0.0
0.4
0.8
Spearman's ρ

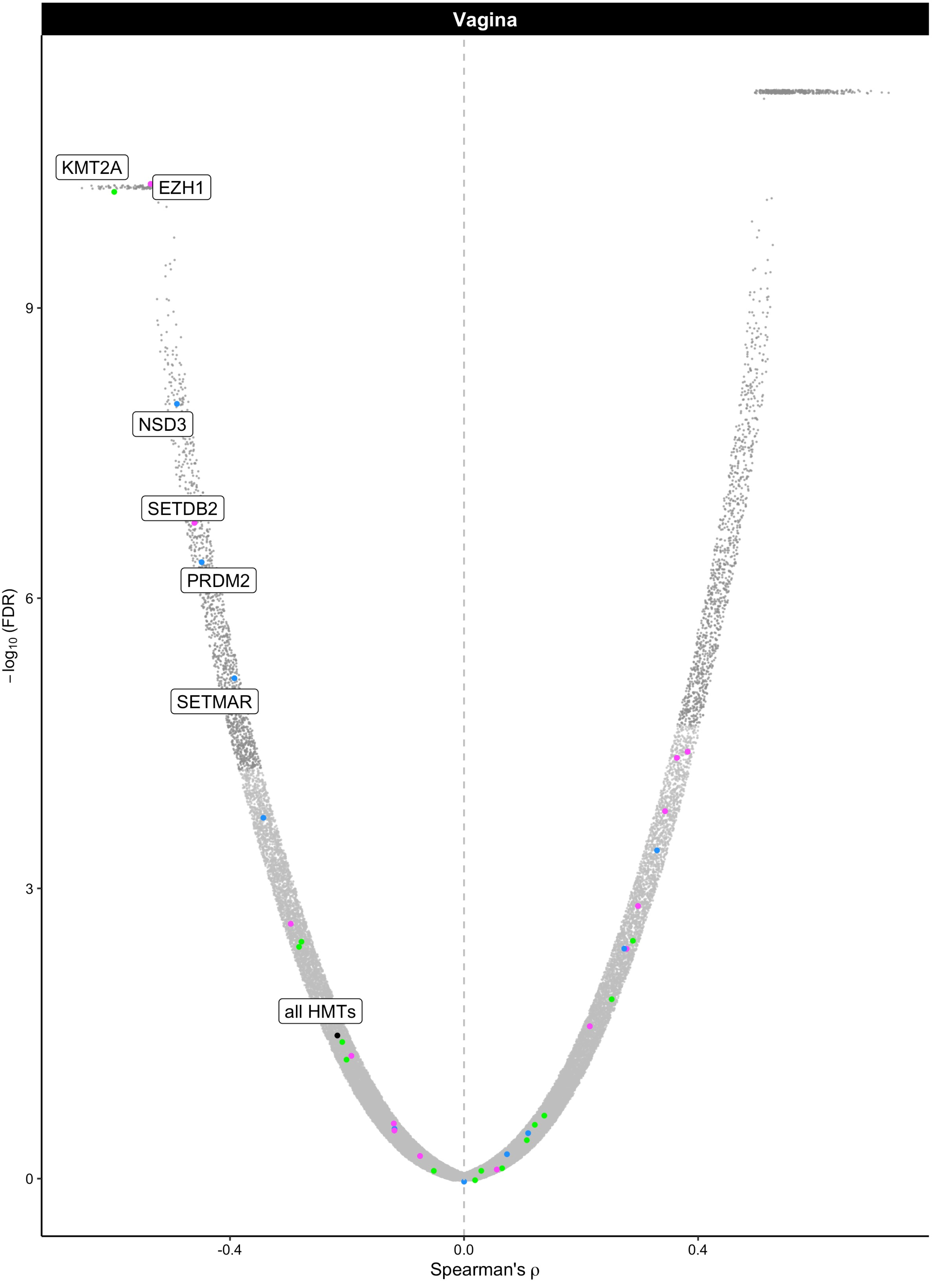

Vagina
KMT2A
EZH1
NSD3
SETDB2
PRDM2
SETMAR
all HMTs
9
6
3
0
−log10 (FDR)
-0.4
0.0
0.4
Spearman's ρ

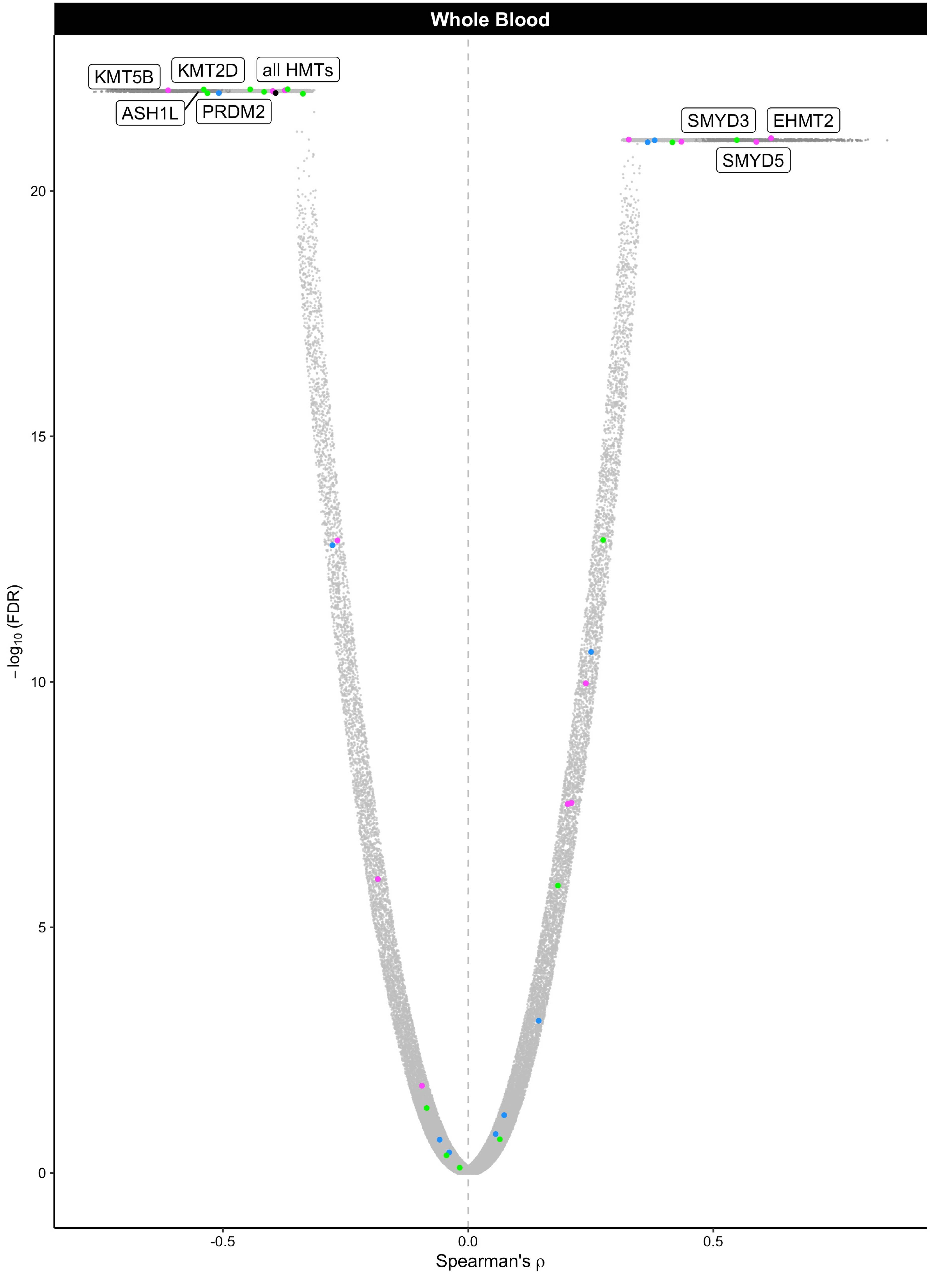

Whole Blood
KMT5B
KMT2D
all HMTs
ASH1L
PRDM2
SMYD3
EHMT2
SMYD5
20
15
10
5
0
−log10 (FDR)
-0.5
0.0
0.5
Spearman's ρ
