## Supplemental Figures and Tables for "Histone methylation has a direct metabolic role in human cells": Supp File 5. GTEX tissue HMT correlations.pdf

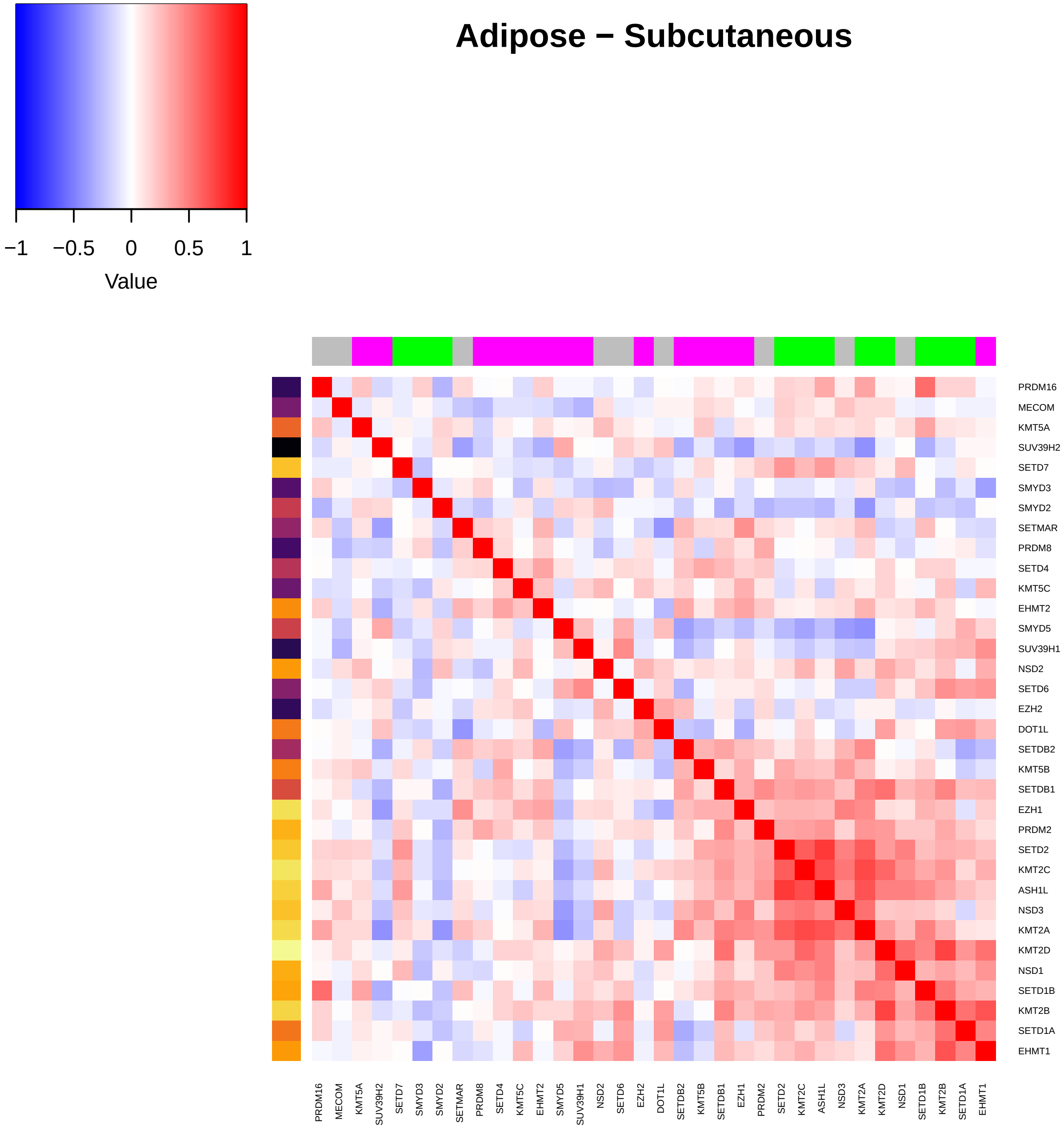
Adipose − Subcutaneous
−1
−0.5
0
0.5
1
Value
PRDM16
MECOM
KMT5A
SUV39H2
SETD7
SMYD3
SMYD2
SETMAR
PRDM8
SETD4
KMT5C
EHMT2
SMYD5
SUV39H1
NSD2
SETD6
EZH2
DOT1L
SETDB2
KMT5B
SETDB1
EZH1
PRDM2
SETD2
KMT2C
ASH1L
NSD3
KMT2A
KMT2D
NSD1
SETD1B
KMT2B
SETD1A
EHMT1

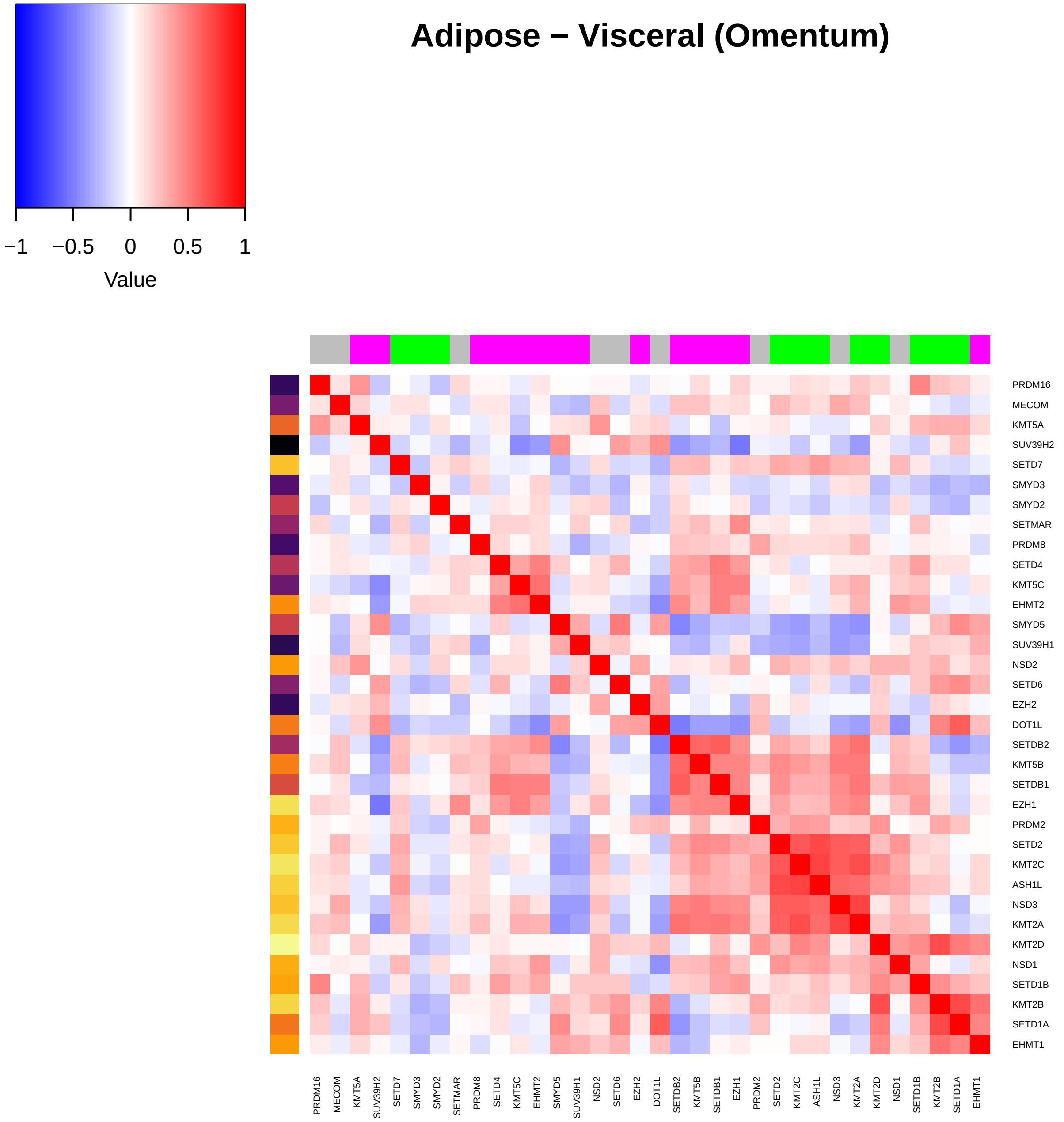
Adipose − Visceral (Omentum)
−1
−0.5
0
0.5
1
Value
PRDM16
MECOM
KMT5A
SUV39H2
SETD7
SMYD3
SMYD2
SETMAR
PRDM8
SETD4
KMT5C
EHMT2
SMYD5
SUV39H1
NSD2
SETD6
EZH2
DOT1L
SETDB2
KMT5B
SETDB1
EZH1
PRDM2
SETD2
KMT2C
ASH1L
NSD3
KMT2A
KMT2D
NSD1
SETD1B
KMT2B
SETD1A
EHMT1

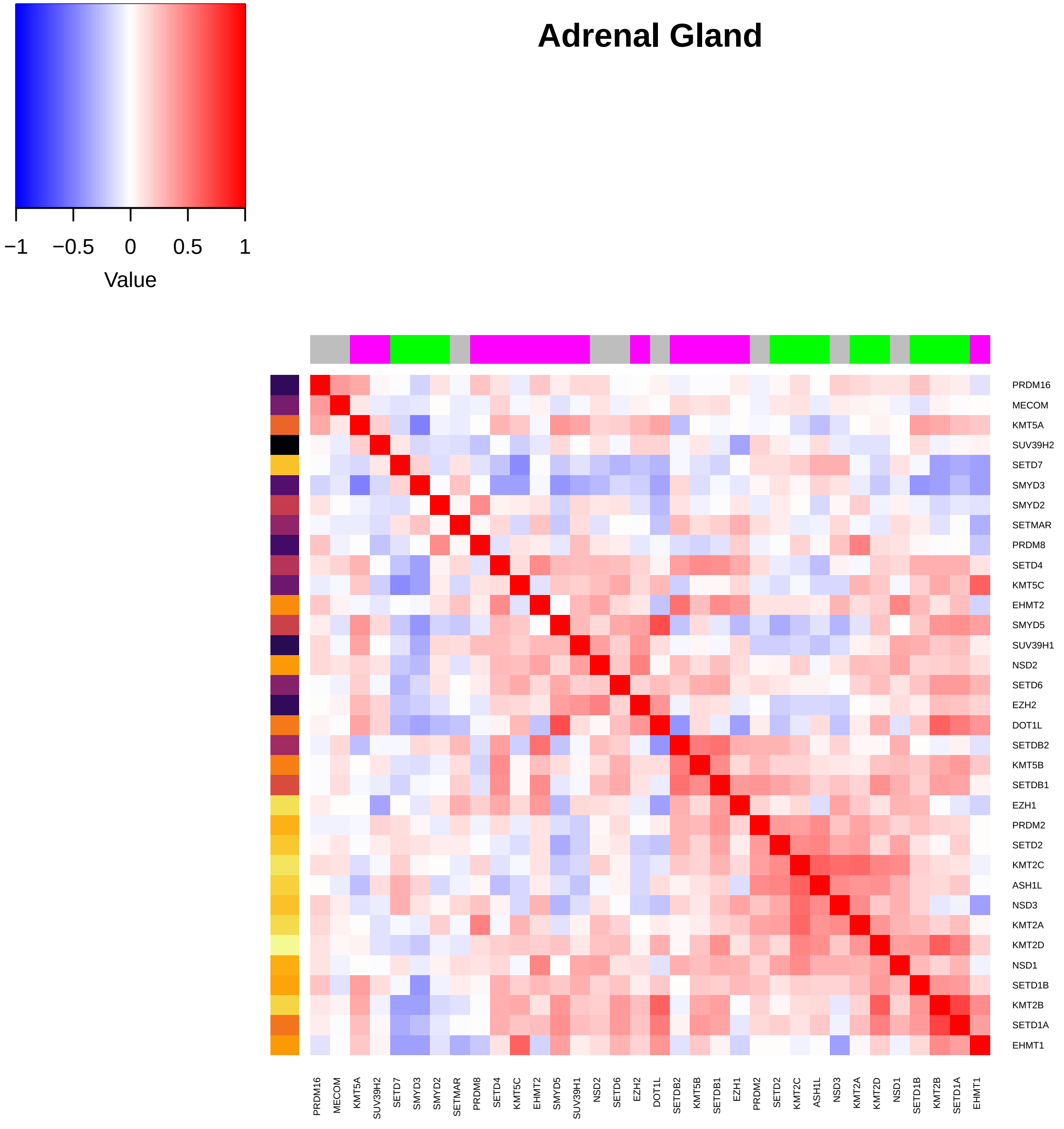

Adrenal Gland
−1
−0.5
0
0.5
1
Value
PRDM16
MECOM
KMT5A
SUV39H2
SETD7
SMYD3
SMYD2
SETMAR
PRDM8
SETD4
KMT5C
EHMT2
SMYD5
SUV39H1
NSD2
SETD6
EZH2
DOT1L
SETDB2
KMT5B
SETDB1
EZH1
PRDM2
SETD2
KMT2C
ASH1L
NSD3
KMT2A
KMT2D
NSD1
SETD1B
KMT2B
SETD1A
EHMT1

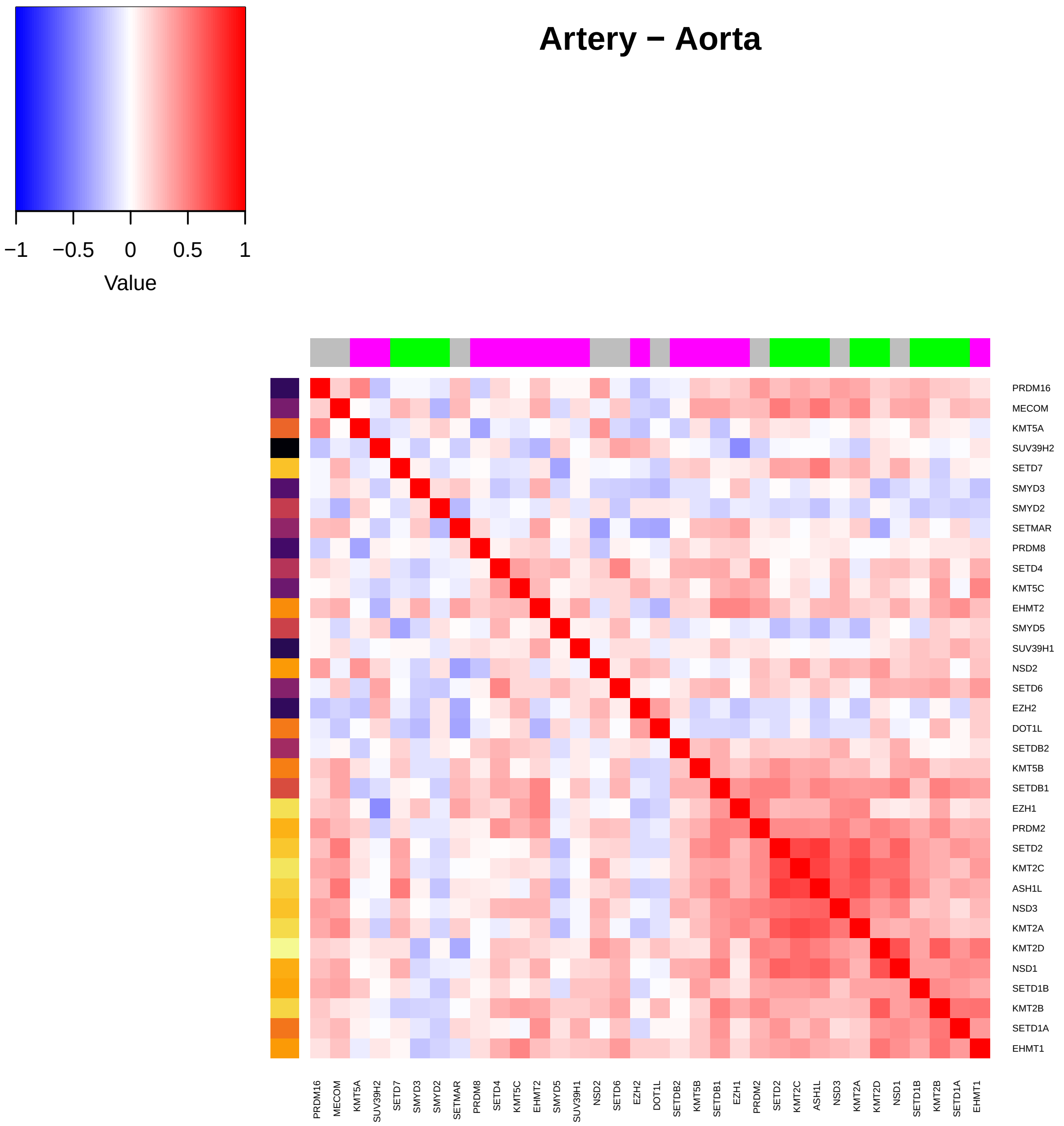
Artery − Aorta
−1
−0.5
0
0.5
1
Value
PRDM16
MECOM
KMT5A
SUV39H2
SETD7
SMYD3
SMYD2
SETMAR
PRDM8
SETD4
KMT5C
EHMT2
SMYD5
SUV39H1
NSD2
SETD6
EZH2
DOT1L
SETDB2
KMT5B
SETDB1
EZH1
PRDM2
SETD2
KMT2C
ASH1L
NSD3
KMT2A
KMT2D
NSD1
SETD1B
KMT2B
SETD1A
EHMT1

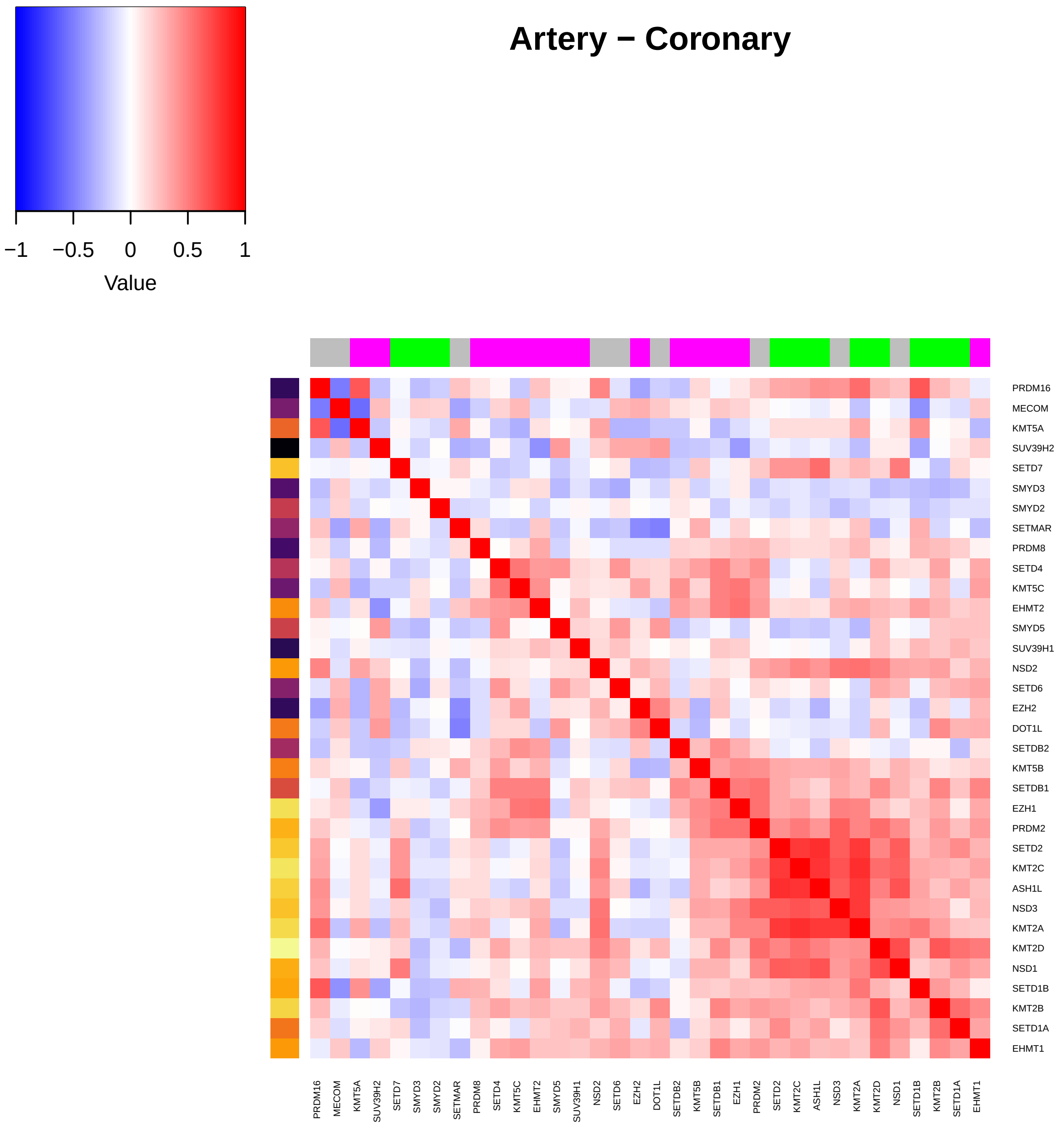

Artery – Coronary
−1
−0.5
0
0.5
1
Value
PRDM16
MECOM
KMT5A
SUV39H2
SETD7
SMYD3
SMYD2
SETMAR
PRDM8
SETD4
KMT5C
EHMT2
SMYD5
SUV39H1
NSD2
SETD6
EZH2
DOT1L
SETDB2
KMT5B
SETDB1
EZH1
PRDM2
SETD2
KMT2C
ASH1L
NSD3
KMT2A
KMT2D
NSD1
SETD1B
KMT2B
SETD1A
EHMT1

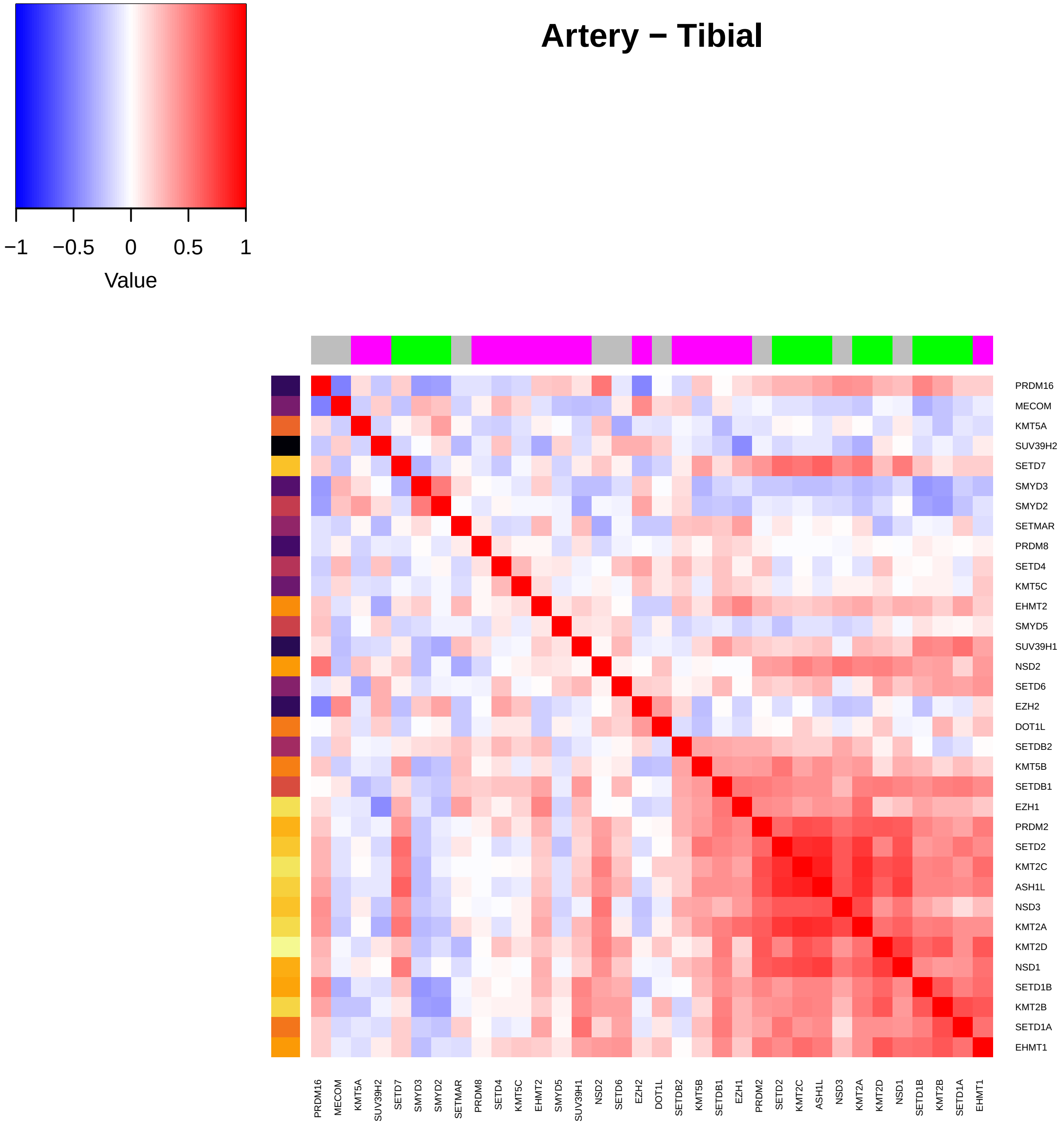

Artery – Tibial
−1
−0.5
0
0.5
1
Value
PRDM16
MECOM
KMT5A
SUV39H2
SETD7
SMYD3
SMYD2
SETMAR
PRDM8
SETD4
KMT5C
EHMT2
SMYD5
SUV39H1
NSD2
SETD6
EZH2
DOT1L
SETDB2
KMT5B
SETDB1
EZH1
PRDM2
SETD2
KMT2C
ASH1L
NSD3
KMT2A
KMT2D
NSD1
SETD1B
KMT2B
SETD1A
EHMT1

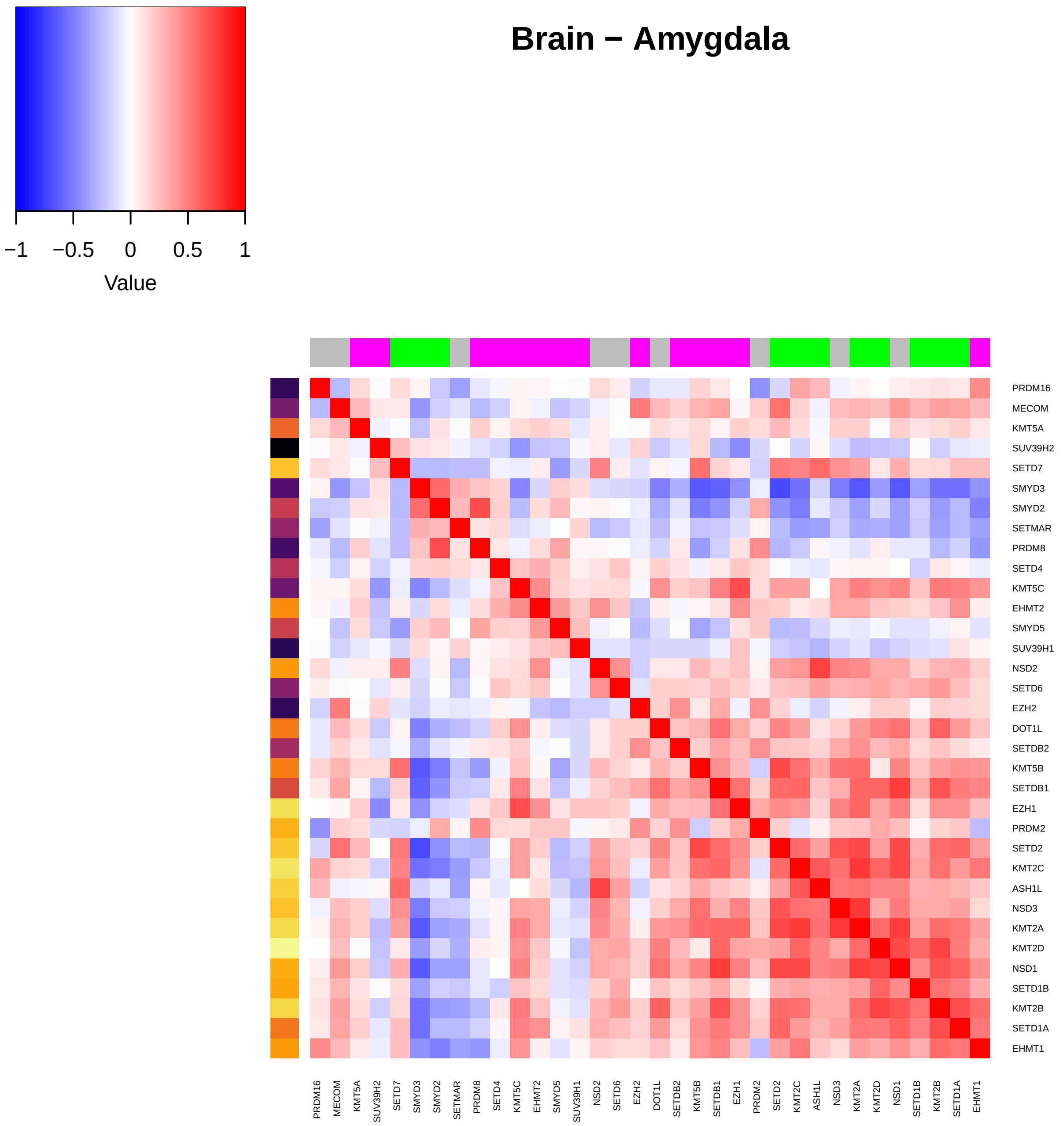

Brain − Amygdala
−1
−0.5
0
0.5
1
Value
PRDM16
MECOM
KMT5A
SUV39H2
SETD7
SMYD3
SMYD2
SETMAR
PRDM8
SETD4
KMT5C
EHMT2
SMYD5
SUV39H1
NSD2
SETD6
EZH2
DOT1L
SETDB2
KMT5B
SETDB1
EZH1
PRDM2
SETD2
KMT2C
ASH1L
NSD3
KMT2A
KMT2D
NSD1
SETD1B
KMT2B
SETD1A
EHMT1

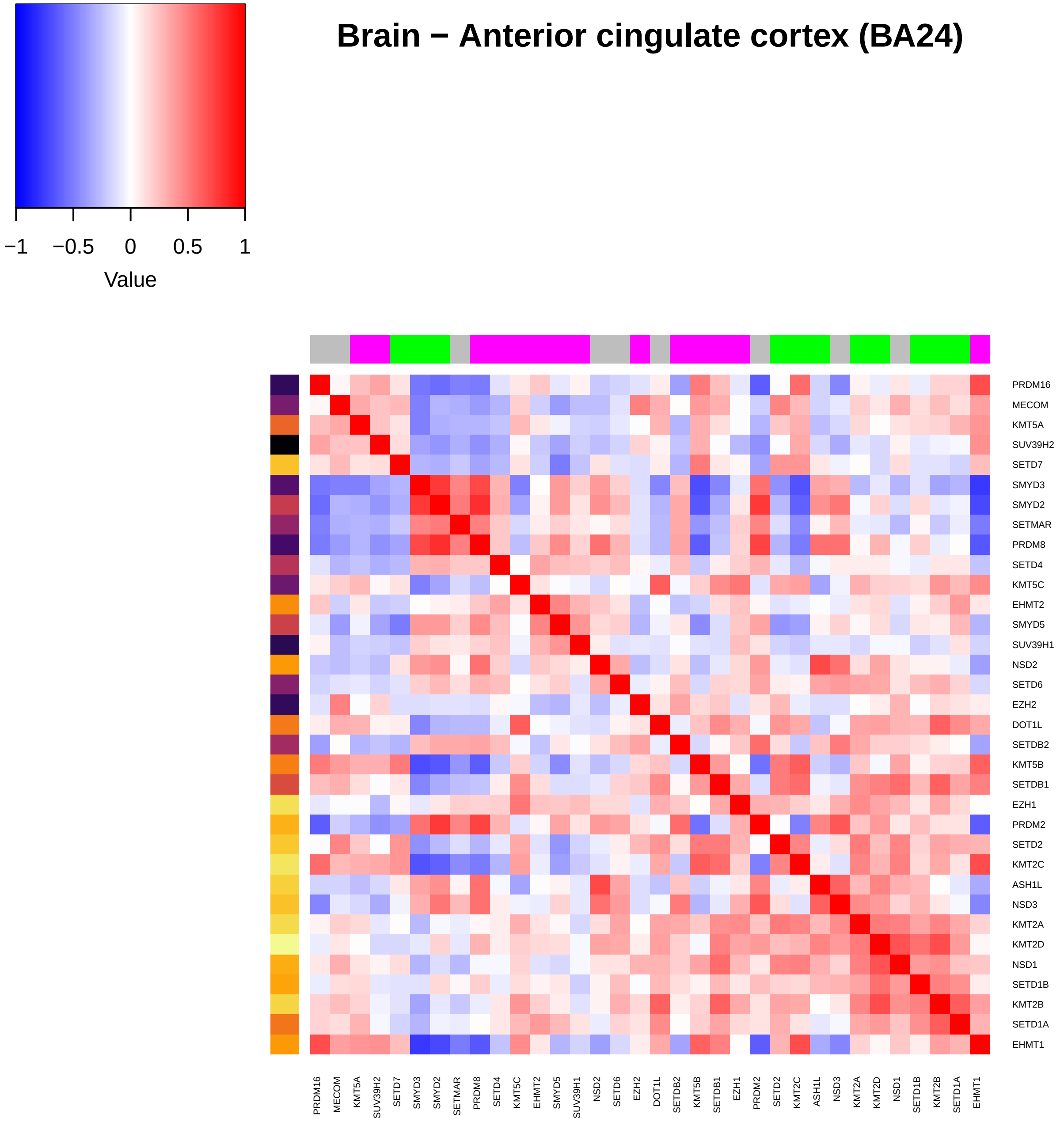

Brain − Anterior cingulate cortex (BA24)
−1
−0.5
0
0.5
1
Value
PRDM16
MECOM
KMT5A
SUV39H2
SETD7
SMYD3
SMYD2
SETMAR
PRDM8
SETD4
KMT5C
EHMT2
SMYD5
SUV39H1
NSD2
SETD6
EZH2
DOT1L
SETDB2
KMT5B
SETDB1
EZH1
PRDM2
SETD2
KMT2C
ASH1L
NSD3
KMT2A
KMT2D
NSD1
SETD1B
KMT2B
SETD1A
EHMT1

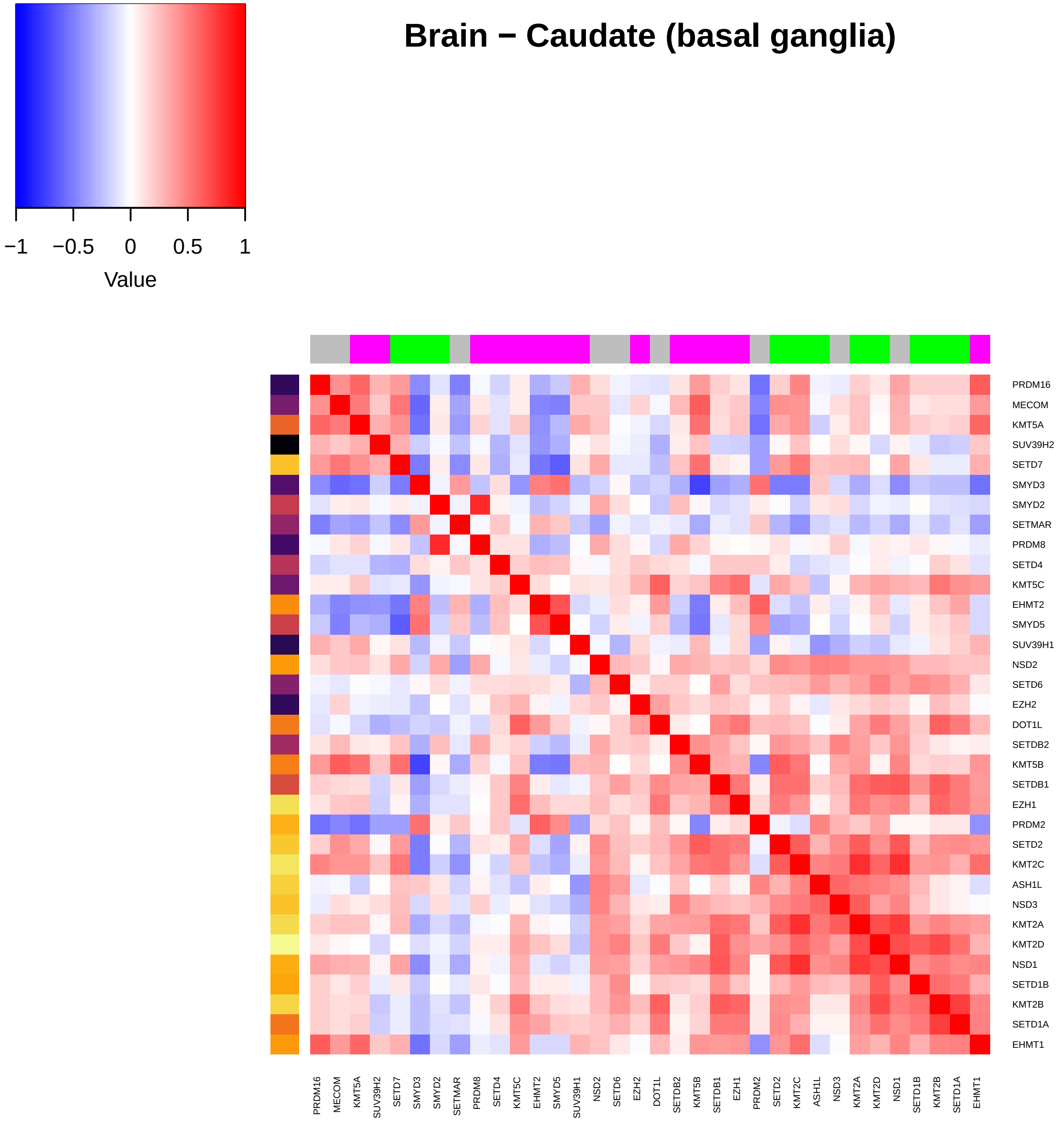

Brain − Caudate (basal ganglia)
−1
−0.5
0
0.5
1
Value
PRDM16
MECOM
KMT5A
SUV39H2
SETD7
SMYD3
SMYD2
SETMAR
PRDM8
SETD4
KMT5C
EHMT2
SMYD5
SUV39H1
NSD2
SETD6
EZH2
DOT1L
SETDB2
KMT5B
SETDB1
EZH1
PRDM2
SETD2
KMT2C
ASH1L
NSD3
KMT2A
KMT2D
NSD1
SETD1B
KMT2B
SETD1A
EHMT1

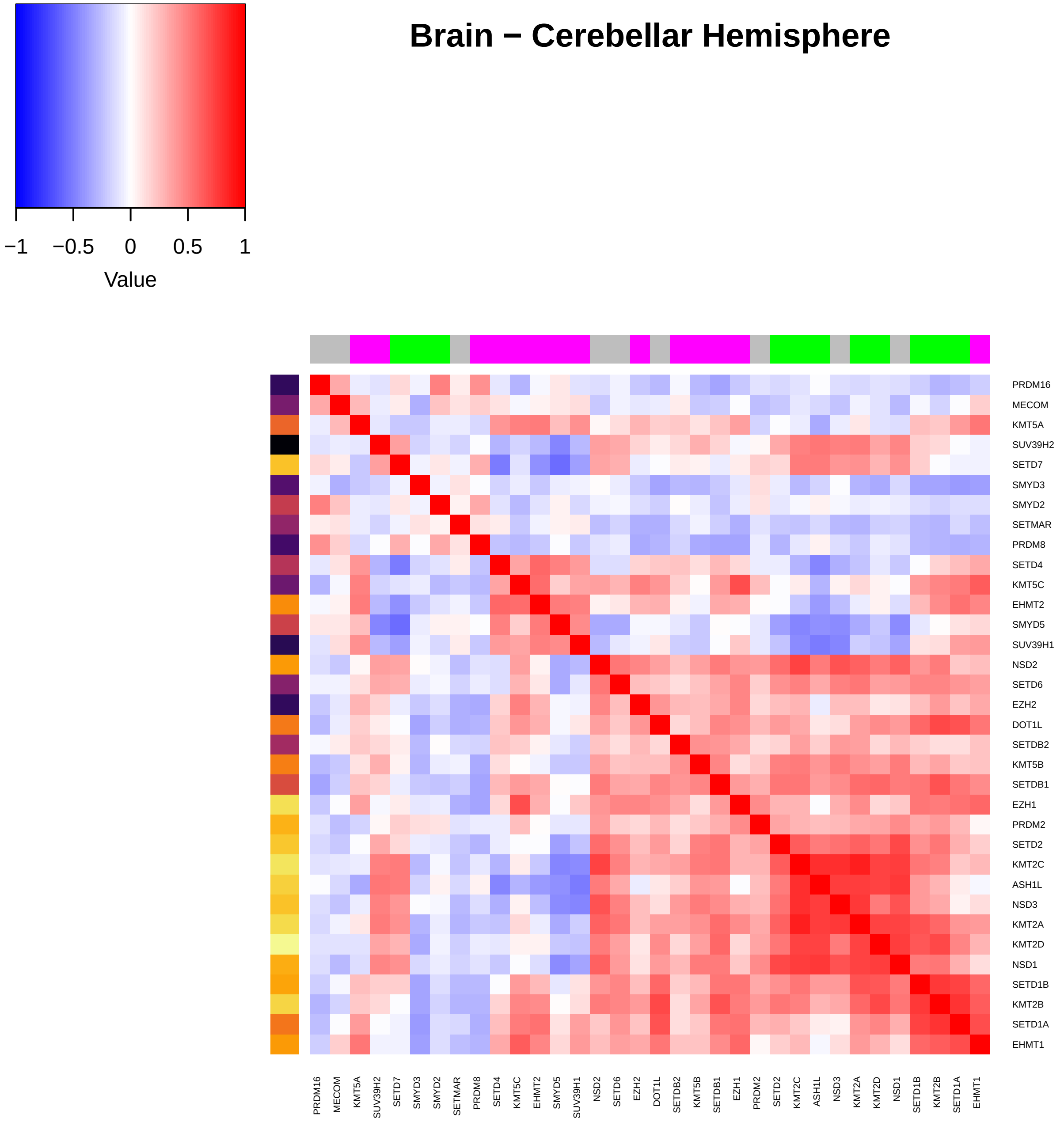
Brain − Cerebellar Hemisphere
−1 −0.5 0 0.5 1
Value
PRDM16
MECOM
KMT5A
SUV39H2
SETD7
SMYD3
SMYD2
SETMAR
PRDM8
SETD4
KMT5C
EHMT2
SMYD5
SUV39H1
NSD2
SETD6
EZH2
DOT1L
SETDB2
KMT5B
SETDB1
EZH1
PRDM2
SETD2
KMT2C
ASH1L
NSD3
KMT2A
KMT2D
NSD1
SETD1B
KMT2B
SETD1A
EHMT1
PRDM16 MECOM KMT5A SUV39H2 SETD7 SMYD3 SMYD2 SETMAR PRDM8 SETD4 KMT5C EHMT2 SMYD5 SUV39H1 NSD2 SETD6 EZH2 DOT1L SETDB2 KMT5B SETDB1 EZH1 PRDM2 SETD2 KMT2C ASH1L NSD3 KMT2A KMT2D NSD1 SETD1B KMT2B SETD1A EHMT1

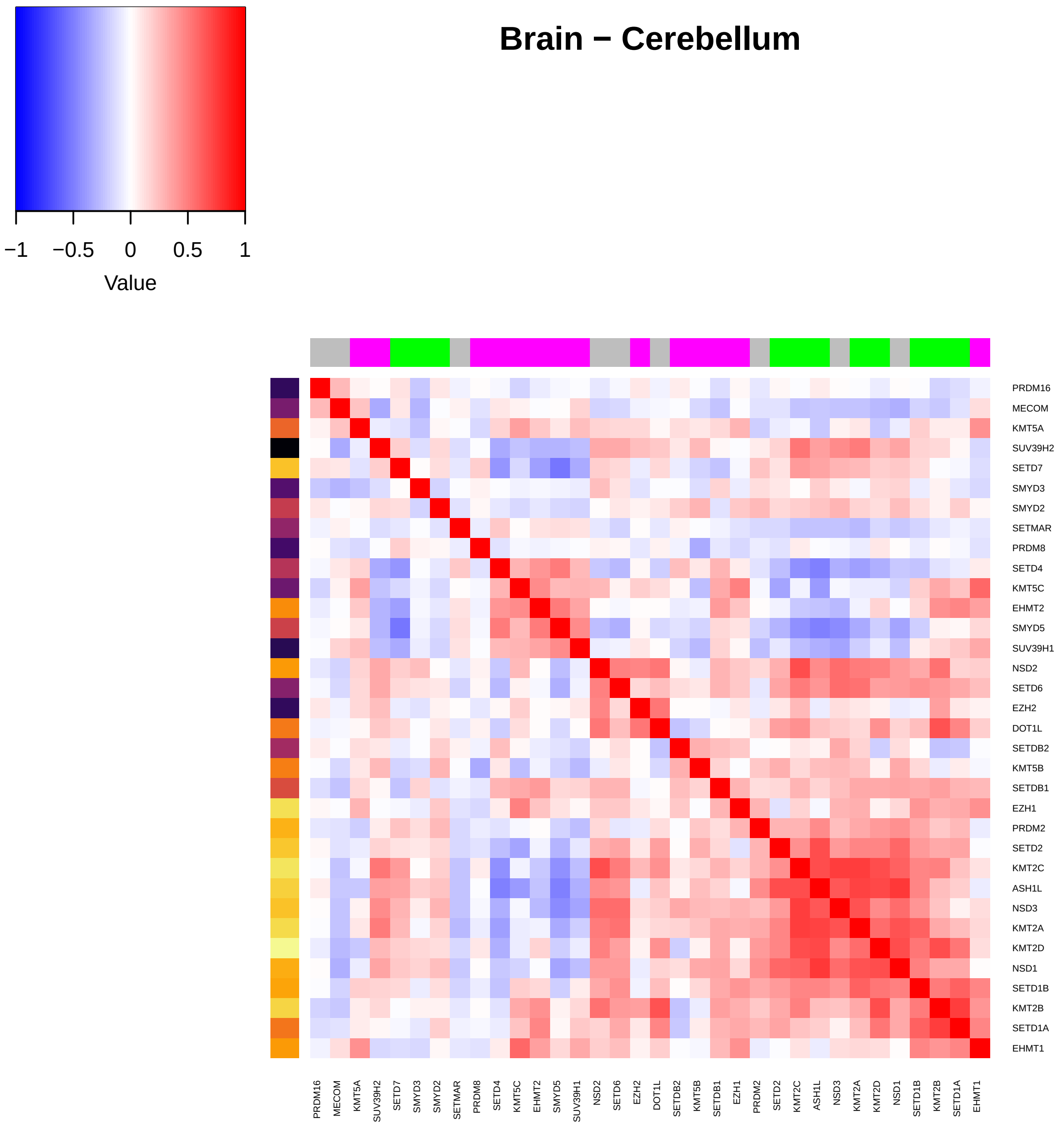

Brain − Cerebellum
−1
−0.5
0
0.5
1
Value
PRDM16
MECOM
KMT5A
SUV39H2
SETD7
SMYD3
SMYD2
SETMAR
PRDM8
SETD4
KMT5C
EHMT2
SMYD5
SUV39H1
NSD2
SETD6
EZH2
DOT1L
SETDB2
KMT5B
SETDB1
EZH1
PRDM2
SETD2
KMT2C
ASH1L
NSD3
KMT2A
KMT2D
NSD1
SETD1B
KMT2B
SETD1A
EHMT1

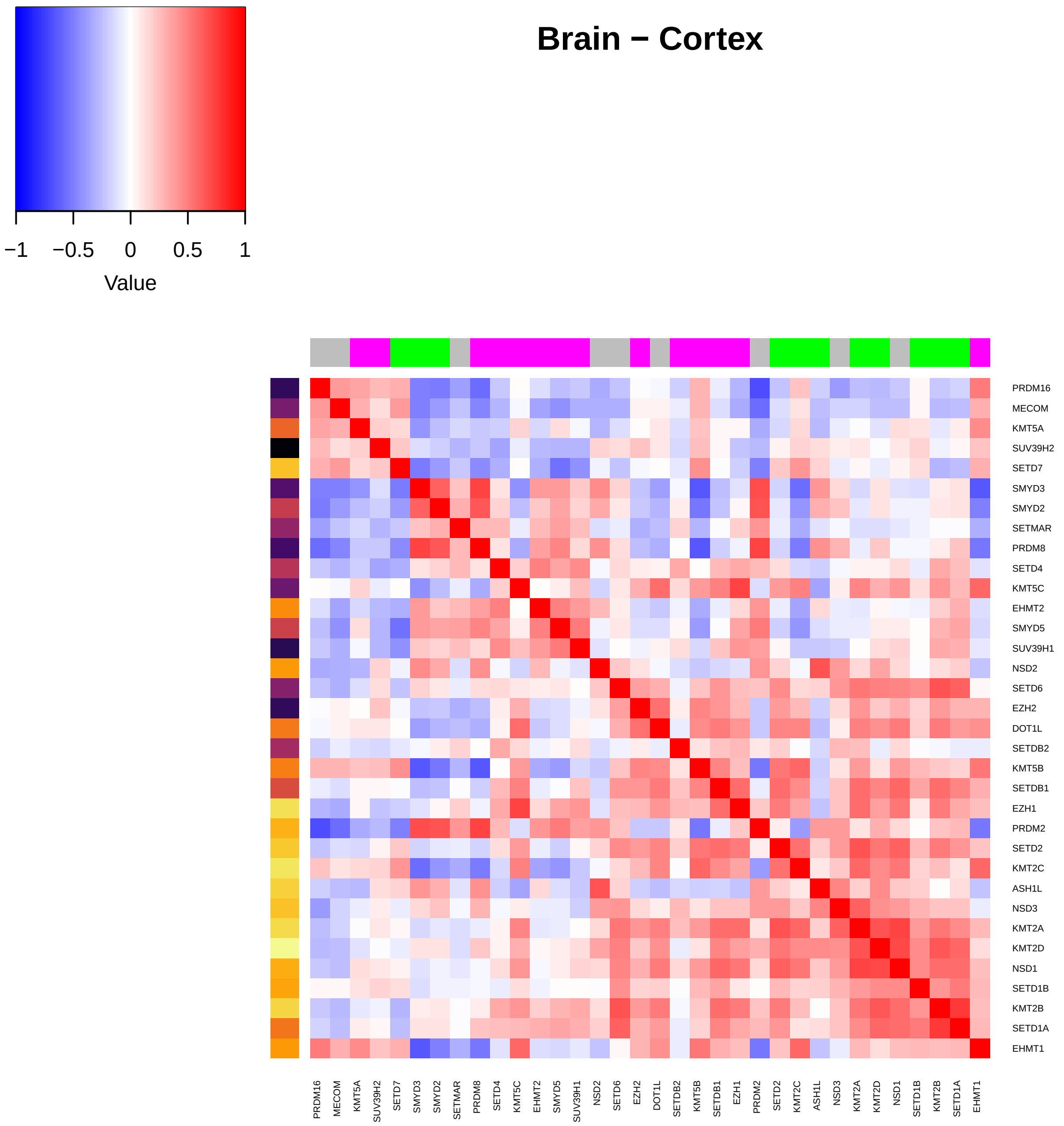
Brain − Cortex
Value
−1
−0.5
0
0.5
1
PRDM16
MECOM
KMT5A
SUV39H2
SETD7
SMYD3
SMYD2
SETMAR
PRDM8
SETD4
KMT5C
EHMT2
SMYD5
SUV39H1
NSD2
SETD6
EZH2
DOT1L
SETDB2
KMT5B
SETDB1
EZH1
PRDM2
SETD2
KMT2C
ASH1L
NSD3
KMT2A
KMT2D
NSD1
SETD1B
KMT2B
SETD1A
EHMT1

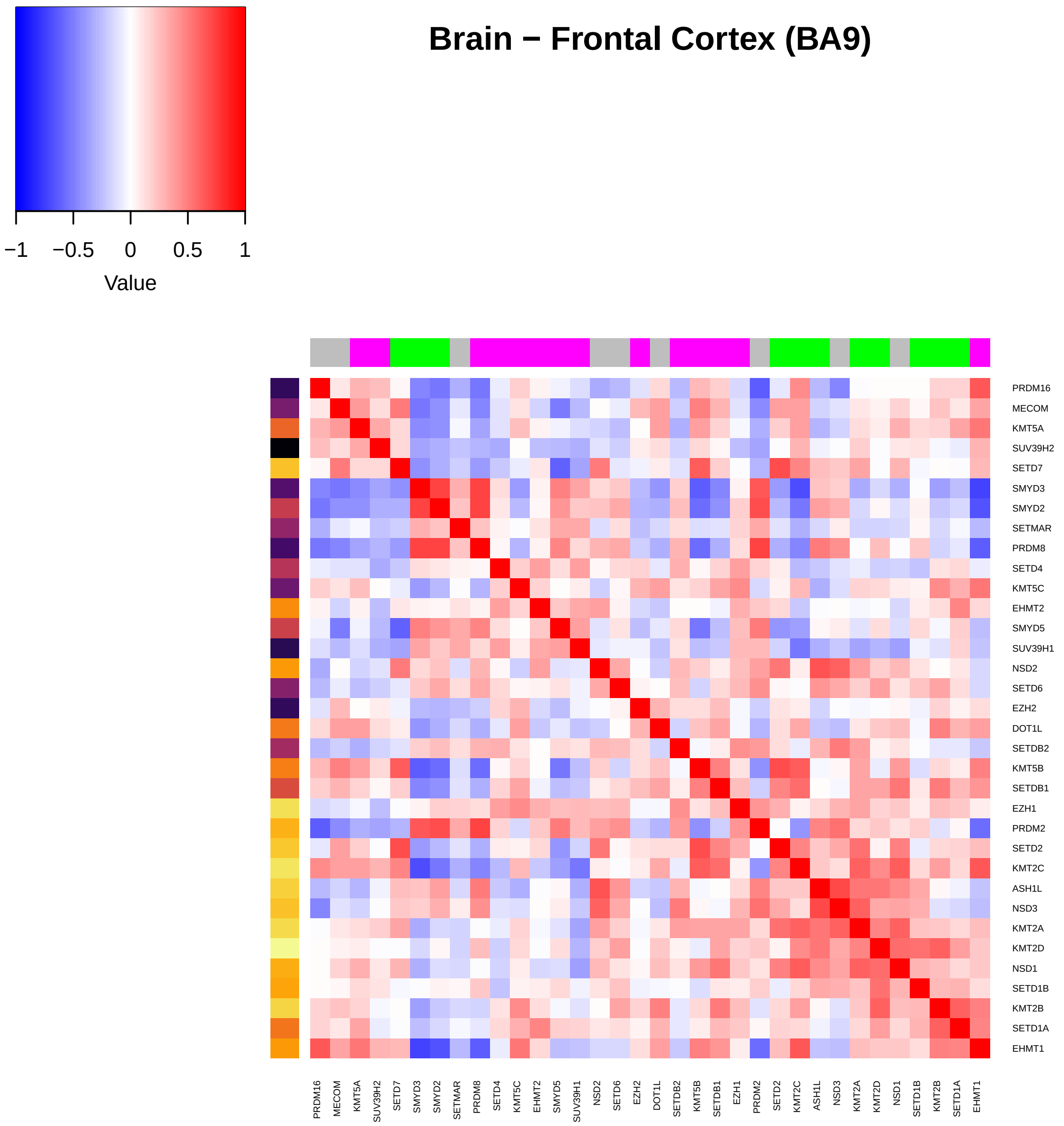

Brain − Frontal Cortex (BA9)
−1
−0.5
0
0.5
1
Value
PRDM16
MECOM
KMT5A
SUV39H2
SETD7
SMYD3
SMYD2
SETMAR
PRDM8
SETD4
KMT5C
EHMT2
SMYD5
SUV39H1
NSD2
SETD6
EZH2
DOT1L
SETDB2
KMT5B
SETDB1
EZH1
PRDM2
SETD2
KMT2C
ASH1L
NSD3
KMT2A
KMT2D
NSD1
SETD1B
KMT2B
SETD1A
EHMT1

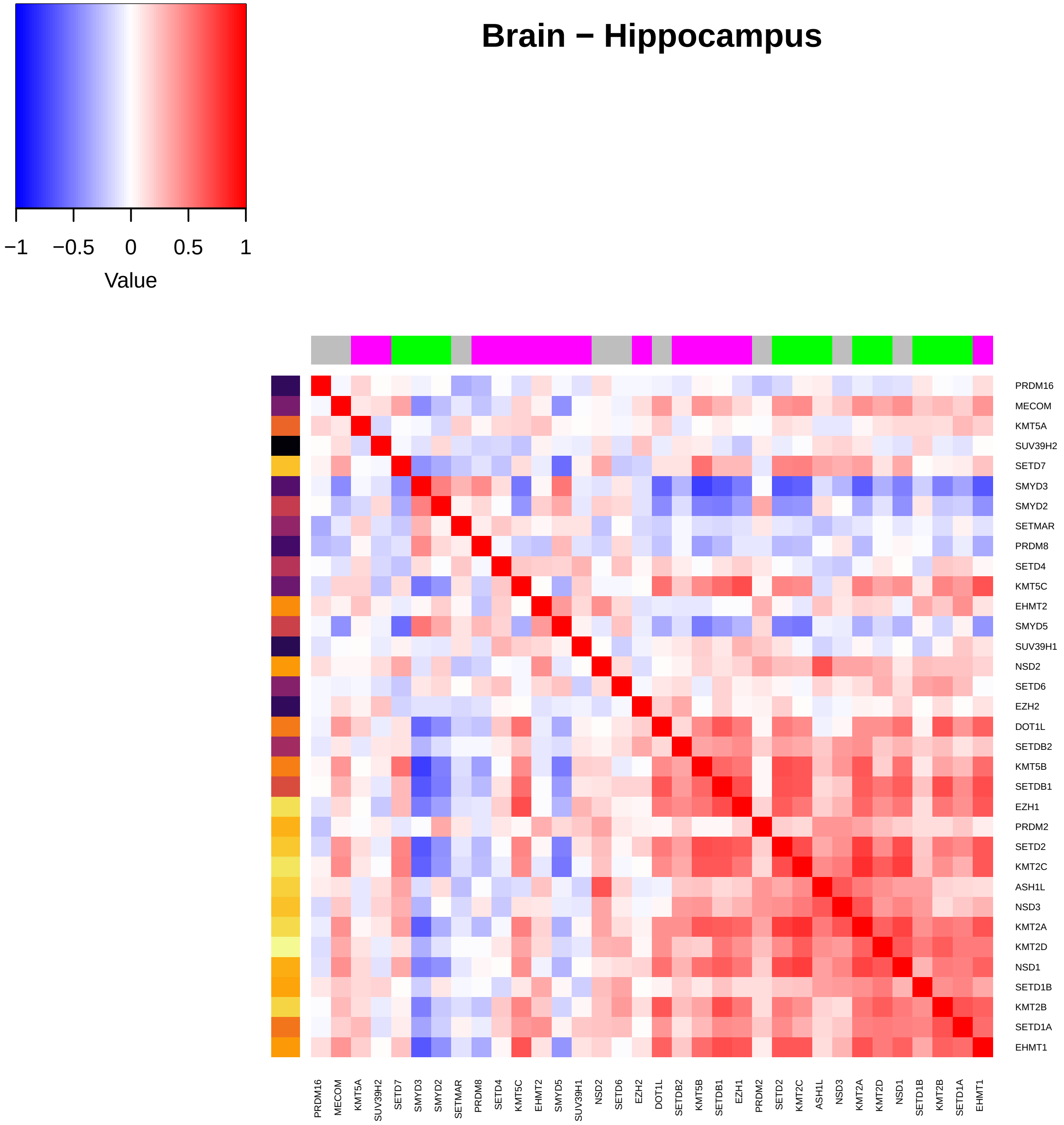
Brain − Hippocampus
−1
−0.5
0
0.5
1
Value
PRDM16
MECOM
KMT5A
SUV39H2
SETD7
SMYD3
SMYD2
SETMAR
PRDM8
SETD4
KMT5C
EHMT2
SMYD5
SUV39H1
NSD2
SETD6
EZH2
DOT1L
SETDB2
KMT5B
SETDB1
EZH1
PRDM2
SETD2
KMT2C
ASH1L
NSD3
KMT2A
KMT2D
NSD1
SETD1B
KMT2B
SETD1A
EHMT1

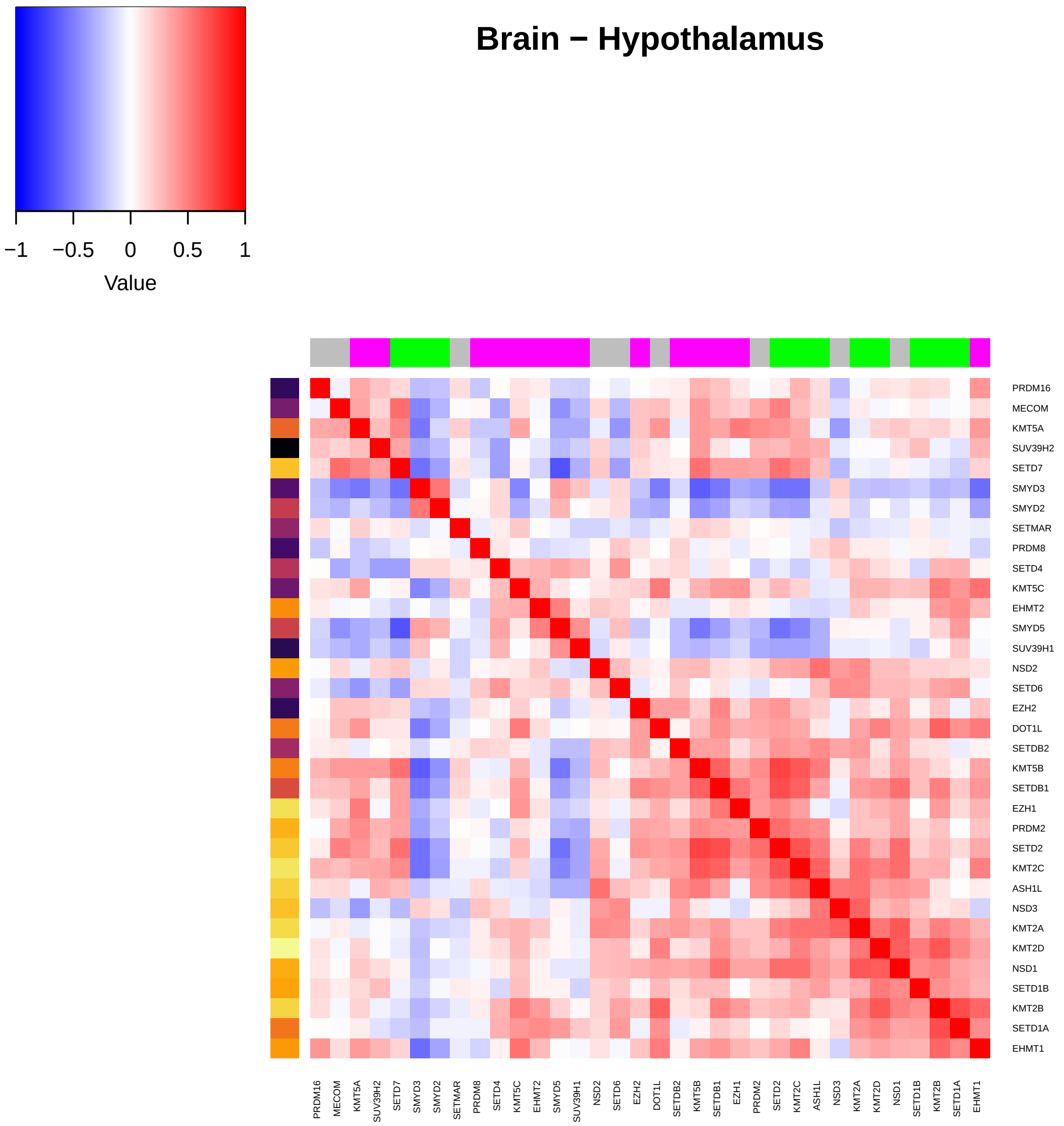
Brain – Hypothalamus
Value
−1
−0.5
0
0.5
1
PRDM16
MECOM
KMT5A
SUV39H2
SETD7
SMYD3
SMYD2
SETMAR
PRDM8
SETD4
KMT5C
EHMT2
SMYD5
SUV39H1
NSD2
SETD6
EZH2
DOT1L
SETDB2
KMT5B
SETDB1
EZH1
PRDM2
SETD2
KMT2C
ASH1L
NSD3
KMT2A
KMT2D
NSD1
SETD1B
KMT2B
SETD1A
EHMT1

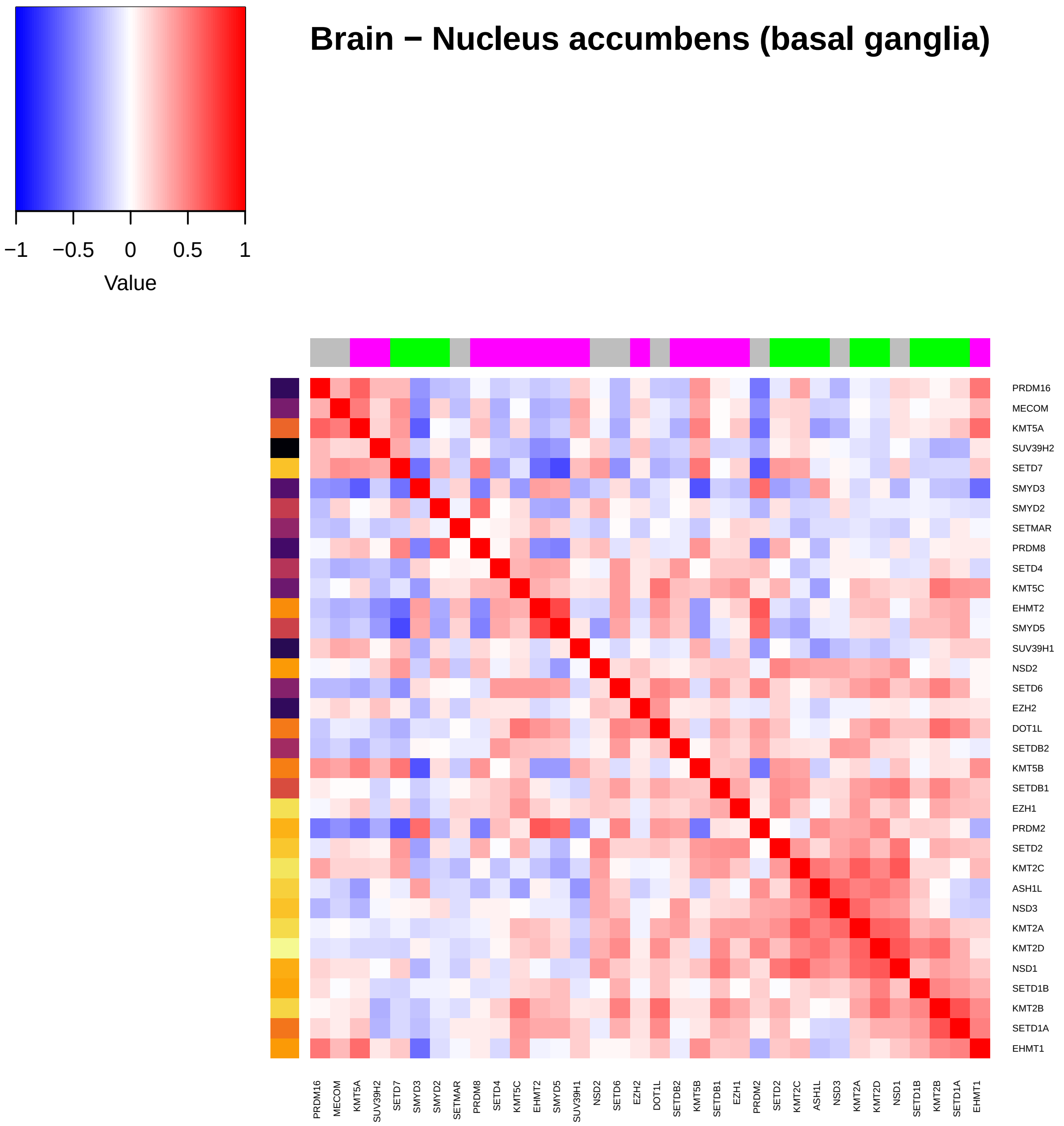

Brain − Nucleus accumbens (basal ganglia)
−1
−0.5
0
0.5
1
Value
PRDM16
MECOM
KMT5A
SUV39H2
SETD7
SMYD3
SMYD2
SETMAR
PRDM8
SETD4
KMT5C
EHMT2
SMYD5
SUV39H1
NSD2
SETD6
EZH2
DOT1L
SETDB2
KMT5B
SETDB1
EZH1
PRDM2
SETD2
KMT2C
ASH1L
NSD3
KMT2A
KMT2D
NSD1
SETD1B
KMT2B
SETD1A
EHMT1

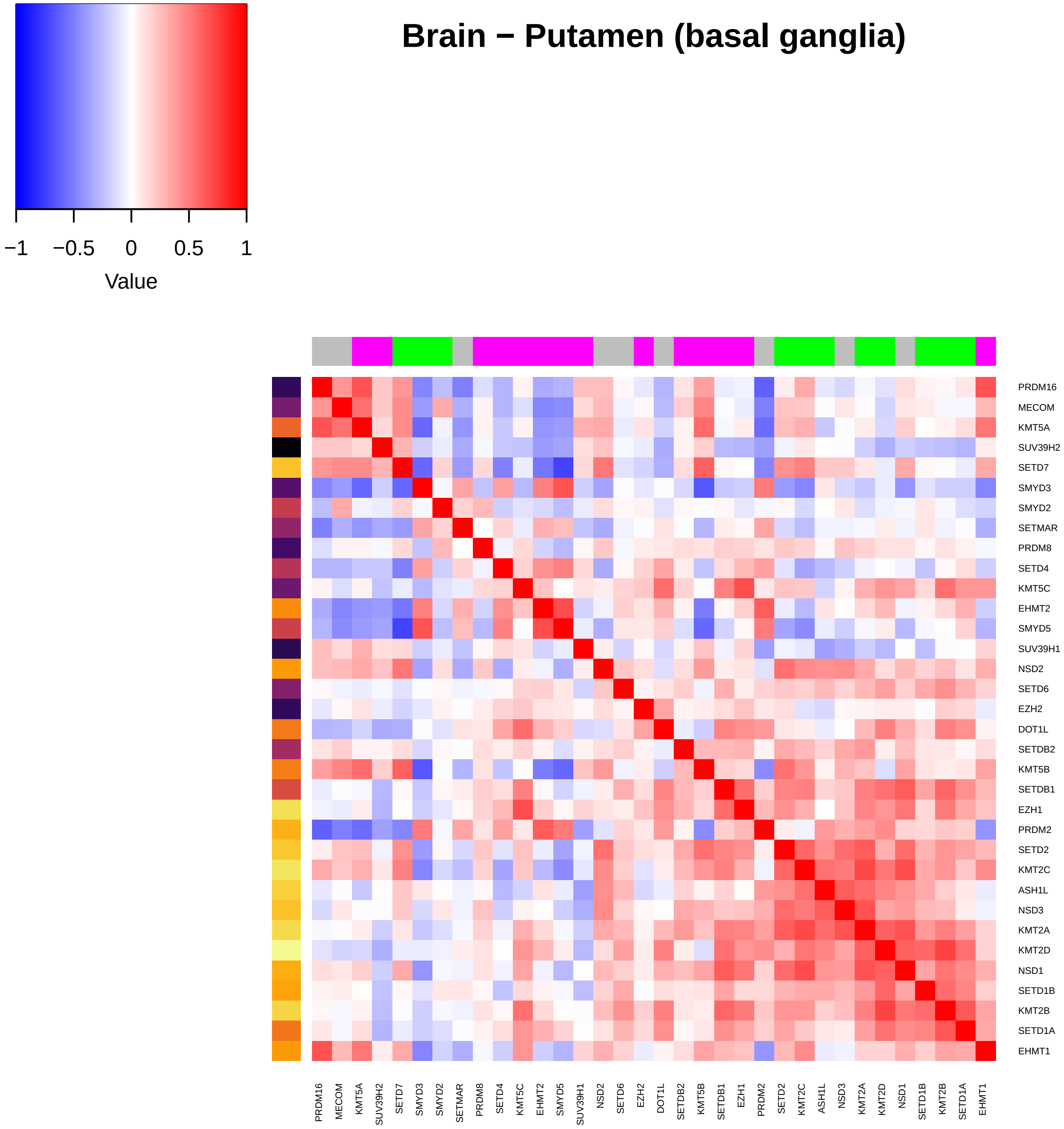
Brain − Putamen (basal ganglia)
−1
−0.5
0
0.5
1
Value
PRDM16
MECOM
KMT5A
SUV39H2
SETD7
SMYD3
SMYD2
SETMAR
PRDM8
SETD4
KMT5C
EHMT2
SMYD5
SUV39H1
NSD2
SETD6
EZH2
DOT1L
SETDB2
KMT5B
SETDB1
EZH1
PRDM2
SETD2
KMT2C
ASH1L
NSD3
KMT2A
KMT2D
NSD1
SETD1B
KMT2B
SETD1A
EHMT1

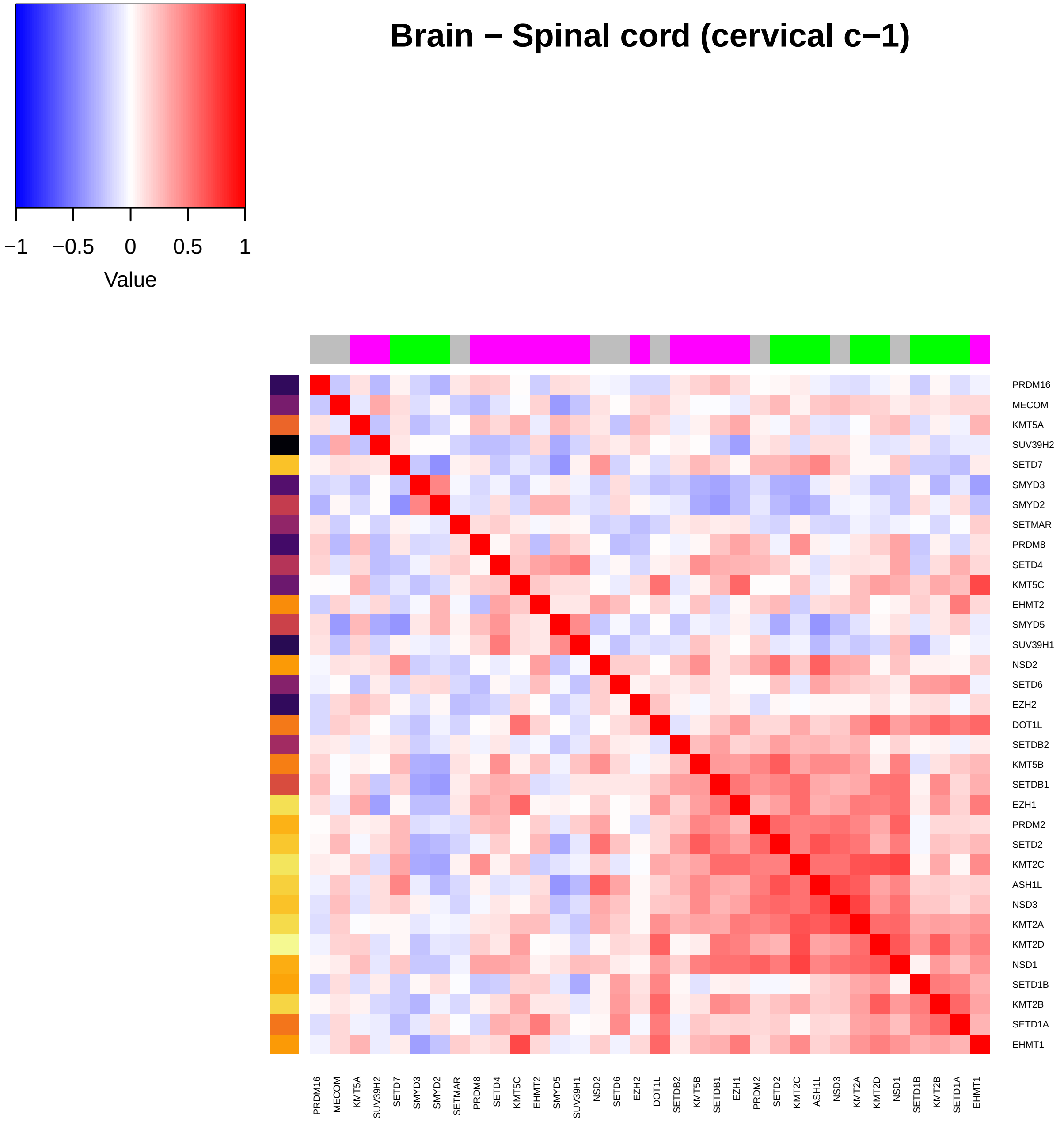

Brain − Spinal cord (cervical c−1)
−1
−0.5
0
0.5
1
Value
PRDM16
MECOM
KMT5A
SUV39H2
SETD7
SMYD3
SMYD2
SETMAR
PRDM8
SETD4
KMT5C
EHMT2
SMYD5
SUV39H1
NSD2
SETD6
EZH2
DOT1L
SETDB2
KMT5B
SETDB1
EZH1
PRDM2
SETD2
KMT2C
ASH1L
NSD3
KMT2A
KMT2D
NSD1
SETD1B
KMT2B
SETD1A
EHMT1

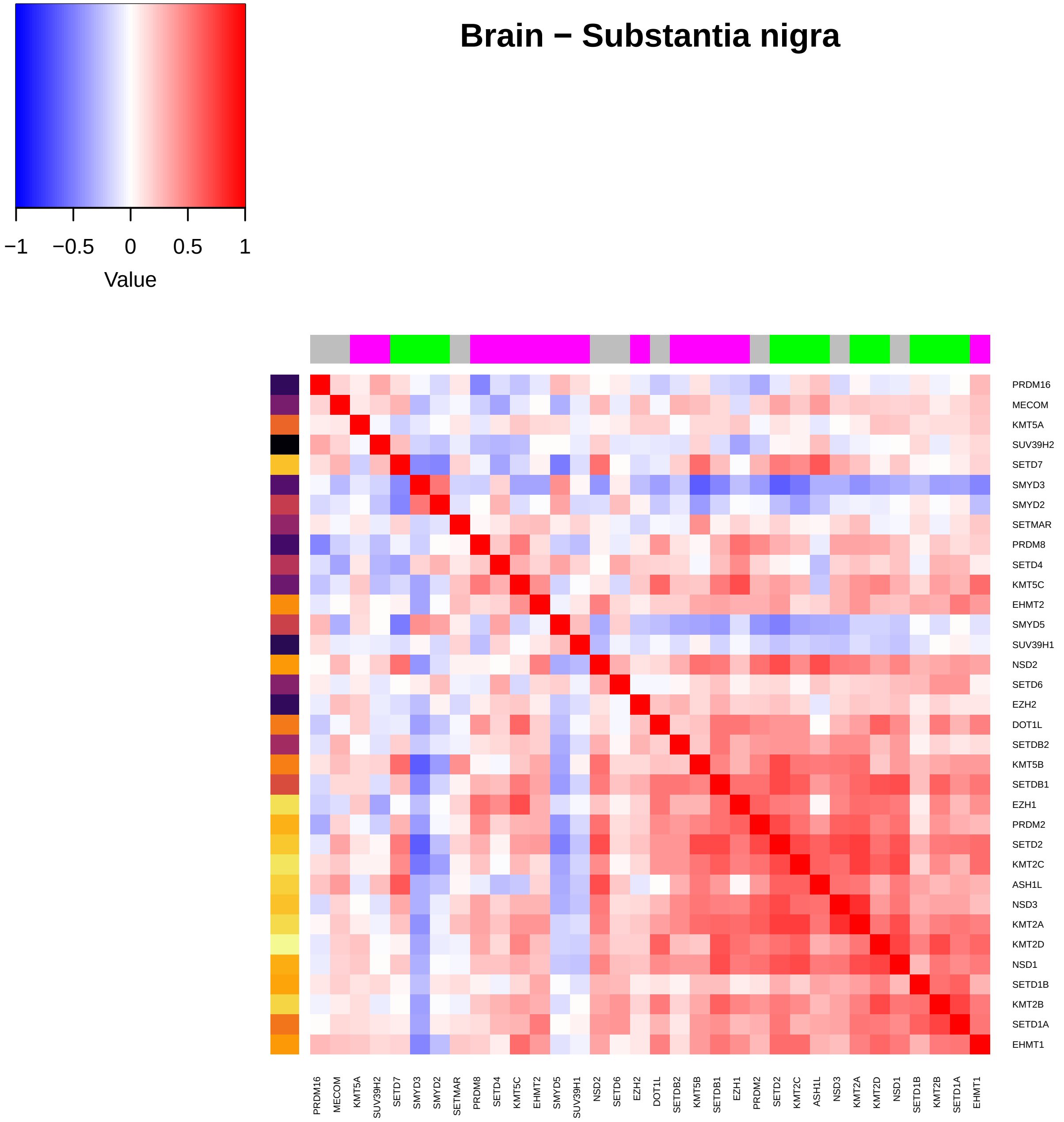

Brain − Substantia nigra
−1
−0.5
0
0.5
1
Value
PRDM16
MECOM
KMT5A
SUV39H2
SETD7
SMYD3
SMYD2
SETMAR
PRDM8
SETD4
KMT5C
EHMT2
SMYD5
SUV39H1
NSD2
SETD6
EZH2
DOT1L
SETDB2
KMT5B
SETDB1
EZH1
PRDM2
SETD2
KMT2C
ASH1L
NSD3
KMT2A
KMT2D
NSD1
SETD1B
KMT2B
SETD1A
EHMT1

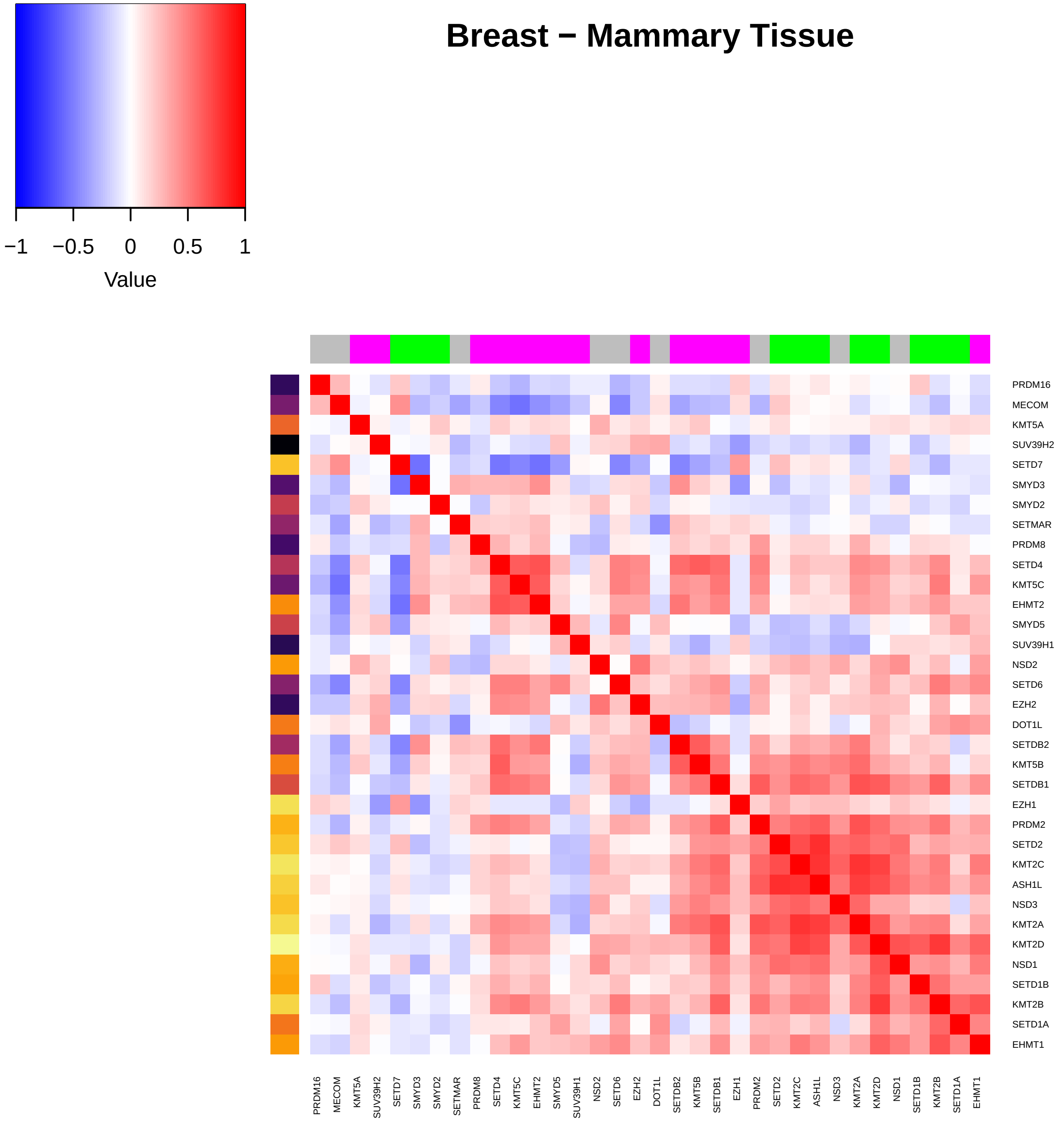
Breast − Mammary Tissue
−1
−0.5
0
0.5
1
Value
PRDM16
MECOM
KMT5A
SUV39H2
SETD7
SMYD3
SMYD2
SETMAR
PRDM8
SETD4
KMT5C
EHMT2
SMYD5
SUV39H1
NSD2
SETD6
EZH2
DOT1L
SETDB2
KMT5B
SETDB1
EZH1
PRDM2
SETD2
KMT2C
ASH1L
NSD3
KMT2A
KMT2D
NSD1
SETD1B
KMT2B
SETD1A
EHMT1

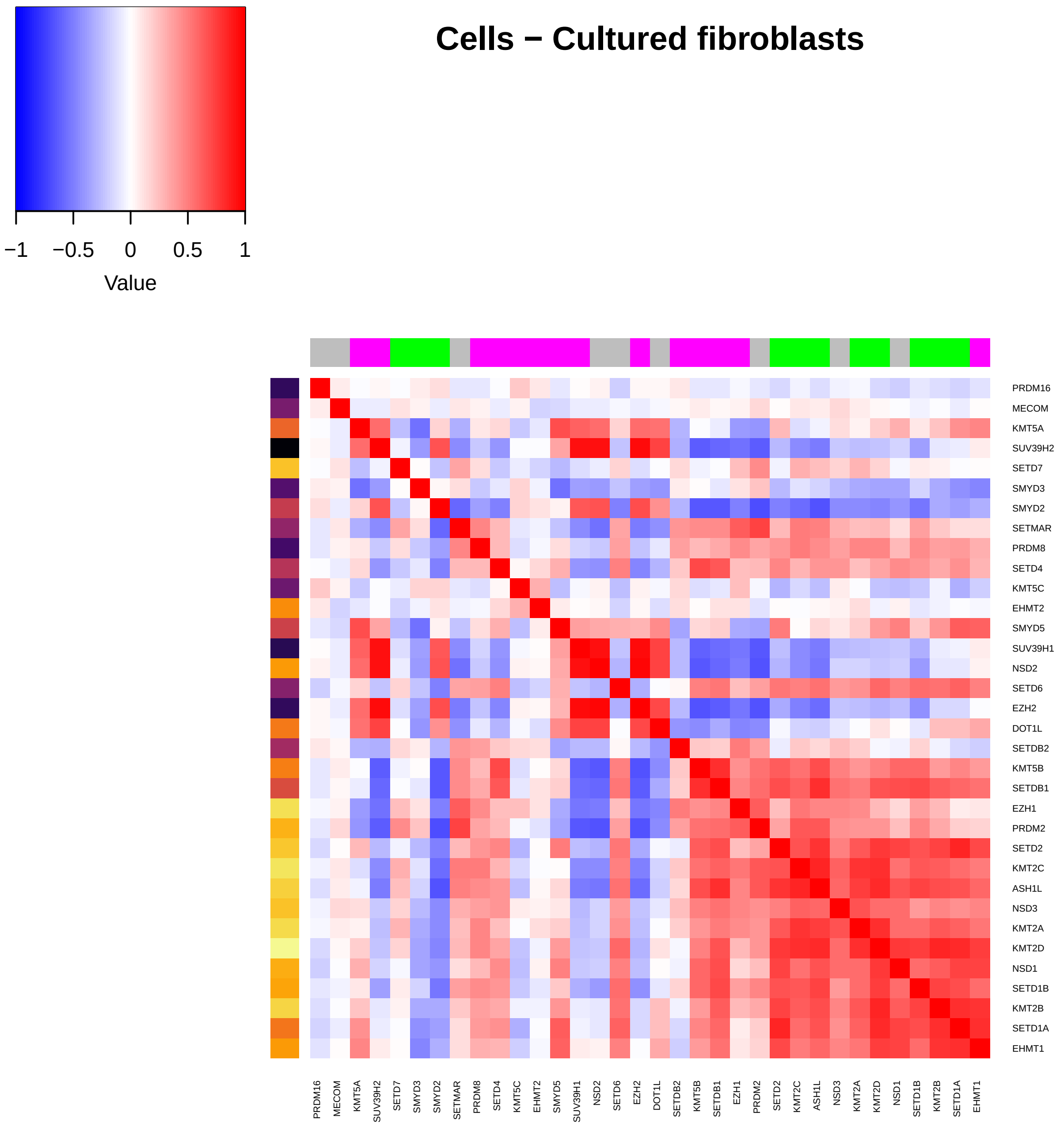

Cells − Cultured fibroblasts
−1
−0.5
0
0.5
1
Value
PRDM16
MECOM
KMT5A
SUV39H2
SETD7
SMYD3
SMYD2
SETMAR
PRDM8
SETD4
KMT5C
EHMT2
SMYD5
SUV39H1
NSD2
SETD6
EZH2
DOT1L
SETDB2
KMT5B
SETDB1
EZH1
PRDM2
SETD2
KMT2C
ASH1L
NSD3
KMT2A
KMT2D
NSD1
SETD1B
KMT2B
SETD1A
EHMT1

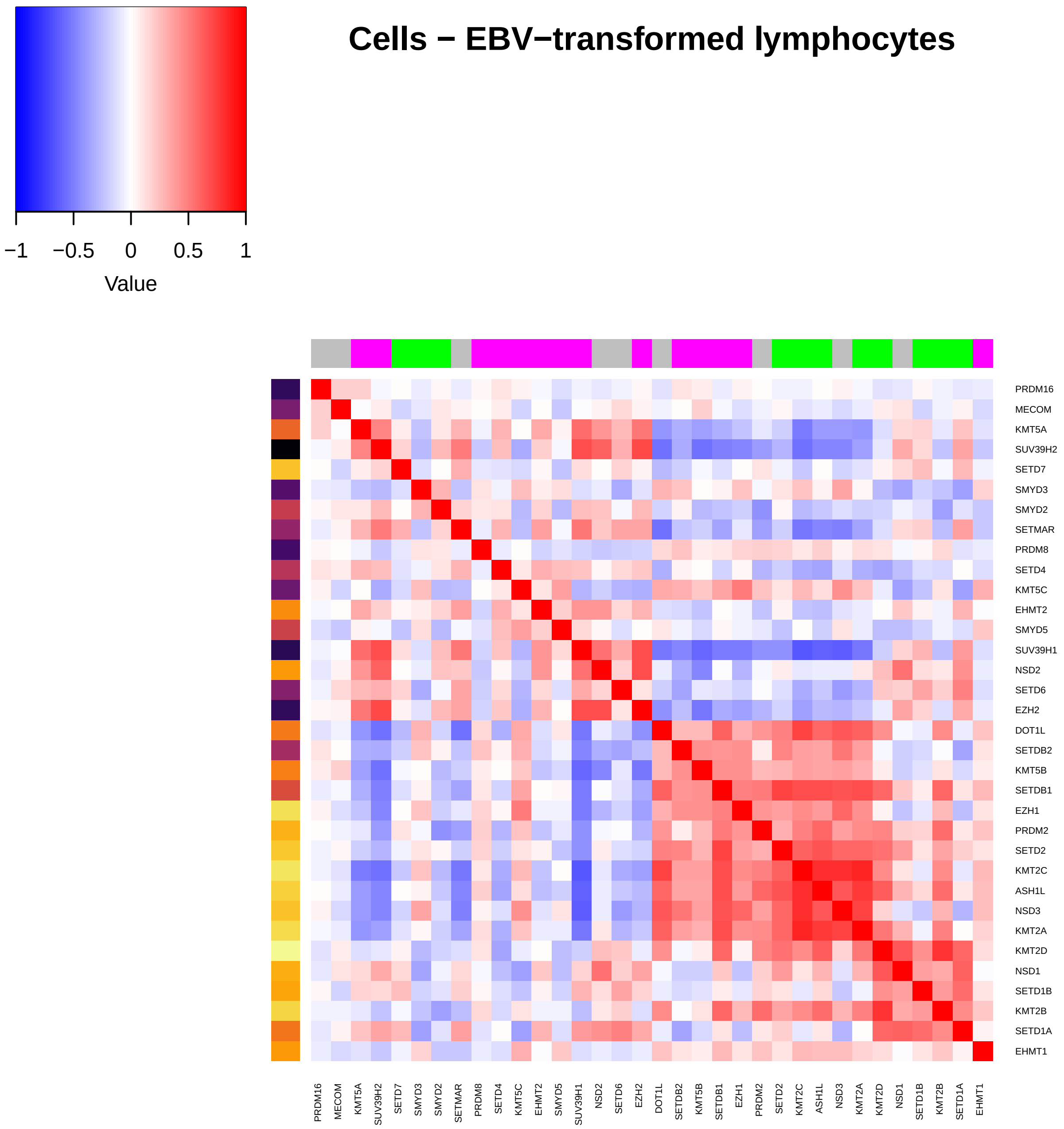
Cells − EBV−transformed lymphocytes
−1
−0.5
0
0.5
1
Value
PRDM16
MECOM
KMT5A
SUV39H2
SETD7
SMYD3
SMYD2
SETMAR
PRDM8
SETD4
KMT5C
EHMT2
SMYD5
SUV39H1
NSD2
SETD6
EZH2
DOT1L
SETDB2
KMT5B
SETDB1
EZH1
PRDM2
SETD2
KMT2C
ASH1L
NSD3
KMT2A
KMT2D
NSD1
SETD1B
KMT2B
SETD1A
EHMT1

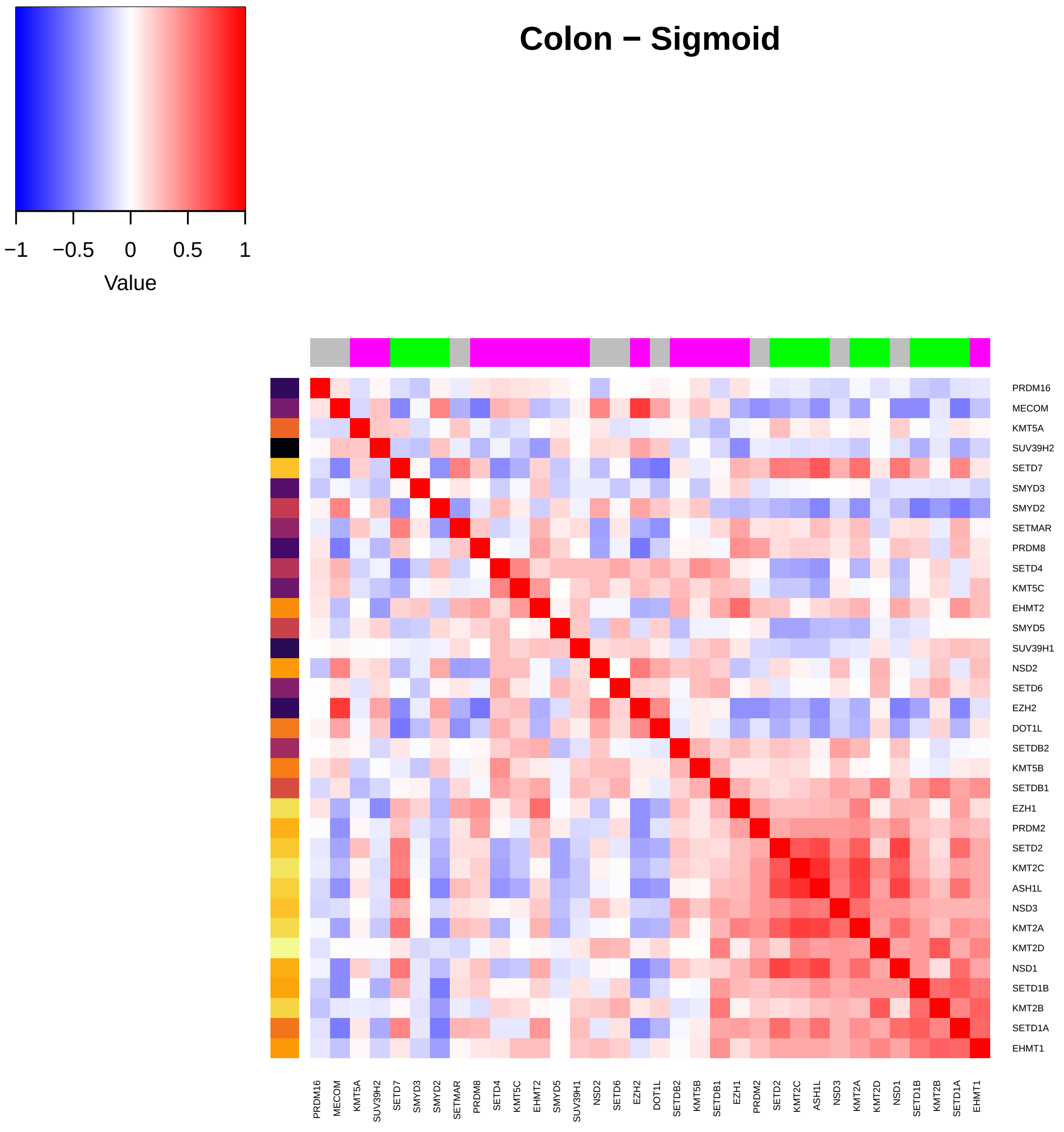

Colon − Sigmoid
−1
−0.5
0
0.5
1
Value
PRDM16
MECOM
KMT5A
SUV39H2
SETD7
SMYD3
SMYD2
SETMAR
PRDM8
SETD4
KMT5C
EHMT2
SMYD5
SUV39H1
NSD2
SETD6
EZH2
DOT1L
SETDB2
KMT5B
SETDB1
EZH1
PRDM2
SETD2
KMT2C
ASH1L
NSD3
KMT2A
KMT2D
NSD1
SETD1B
KMT2B
SETD1A
EHMT1

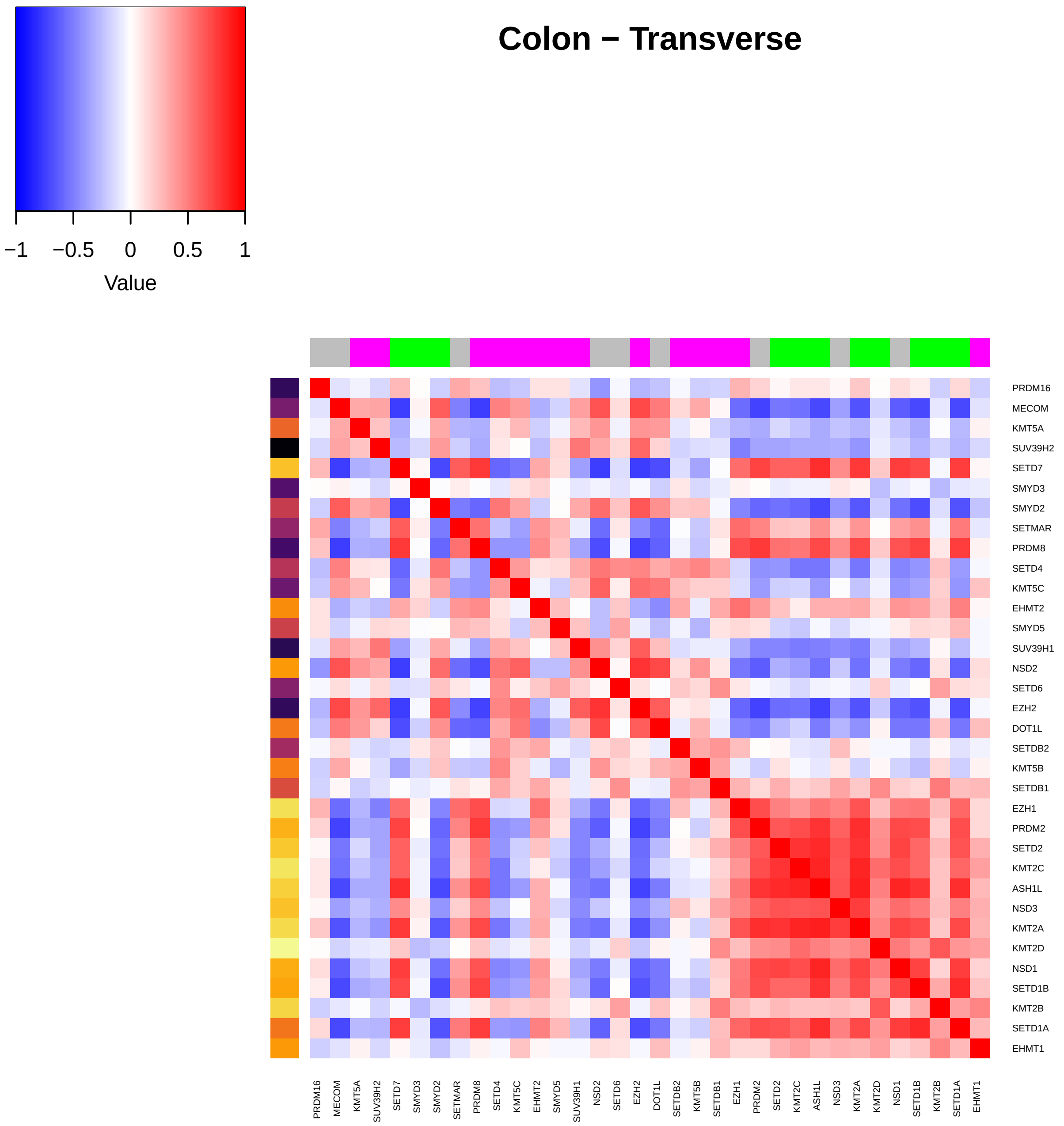
Colon – Transverse
−1
−0.5
0
0.5
1
Value
PRDM16
MECOM
KMT5A
SUV39H2
SETD7
SMYD3
SMYD2
SETMAR
PRDM8
SETD4
KMT5C
EHMT2
SMYD5
SUV39H1
NSD2
SETD6
EZH2
DOT1L
SETDB2
KMT5B
SETDB1
EZH1
PRDM2
SETD2
KMT2C
ASH1L
NSD3
KMT2A
KMT2D
NSD1
SETD1B
KMT2B
SETD1A
EHMT1

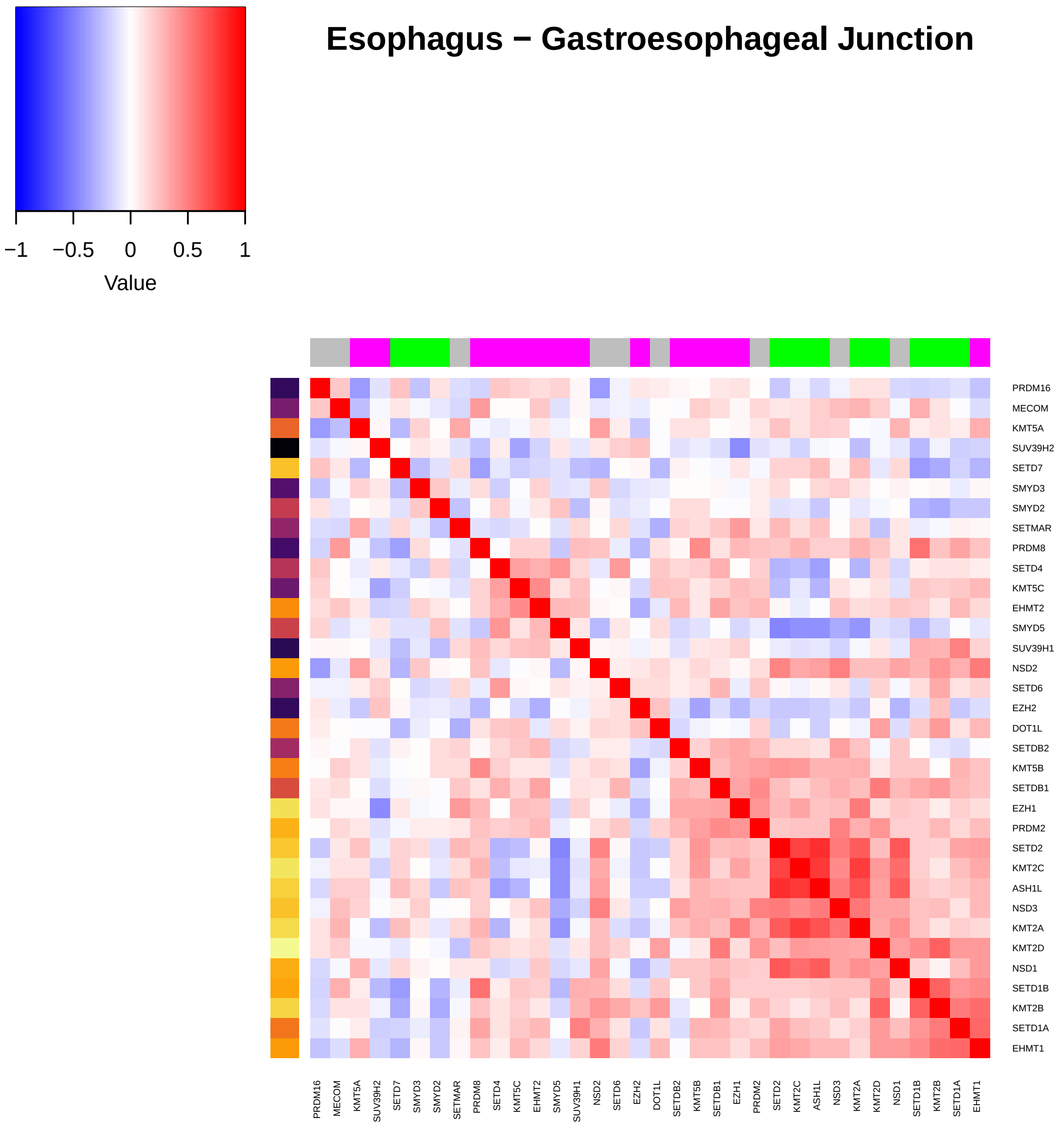
Esophagus − Gastroesophageal Junction
−1
−0.5
0
0.5
1
Value
PRDM16
MECOM
KMT5A
SUV39H2
SETD7
SMYD3
SMYD2
SETMAR
PRDM8
SETD4
KMT5C
EHMT2
SMYD5
SUV39H1
NSD2
SETD6
EZH2
DOT1L
SETDB2
KMT5B
SETDB1
EZH1
PRDM2
SETD2
KMT2C
ASH1L
NSD3
KMT2A
KMT2D
NSD1
SETD1B
KMT2B
SETD1A
EHMT1

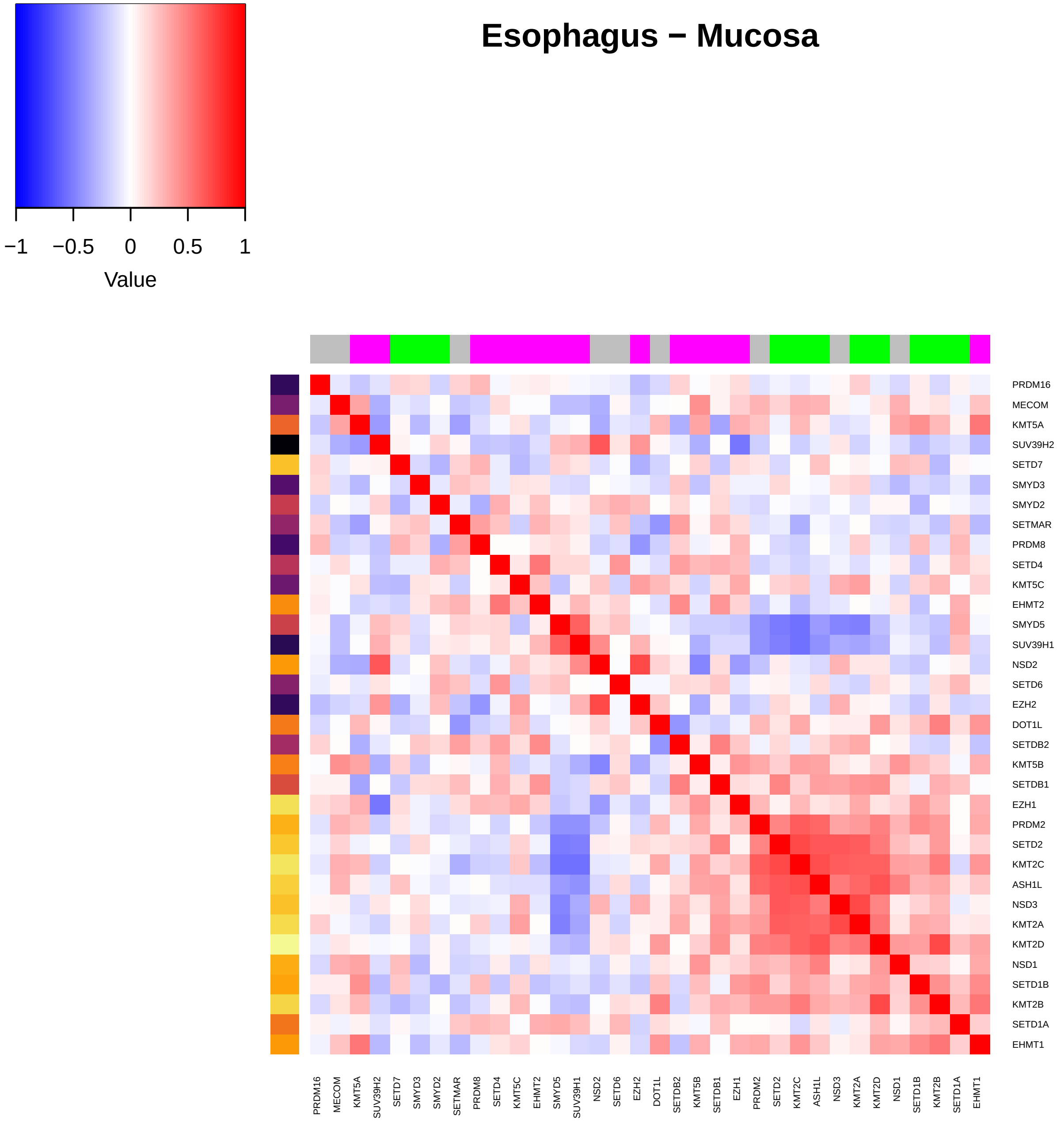

Esophagus − Mucosa
−1
−0.5
0
0.5
1
Value
PRDM16
MECOM
KMT5A
SUV39H2
SETD7
SMYD3
SMYD2
SETMAR
PRDM8
SETD4
KMT5C
EHMT2
SMYD5
SUV39H1
NSD2
SETD6
EZH2
DOT1L
SETDB2
KMT5B
SETDB1
EZH1
PRDM2
SETD2
KMT2C
ASH1L
NSD3
KMT2A
KMT2D
NSD1
SETD1B
KMT2B
SETD1A
EHMT1

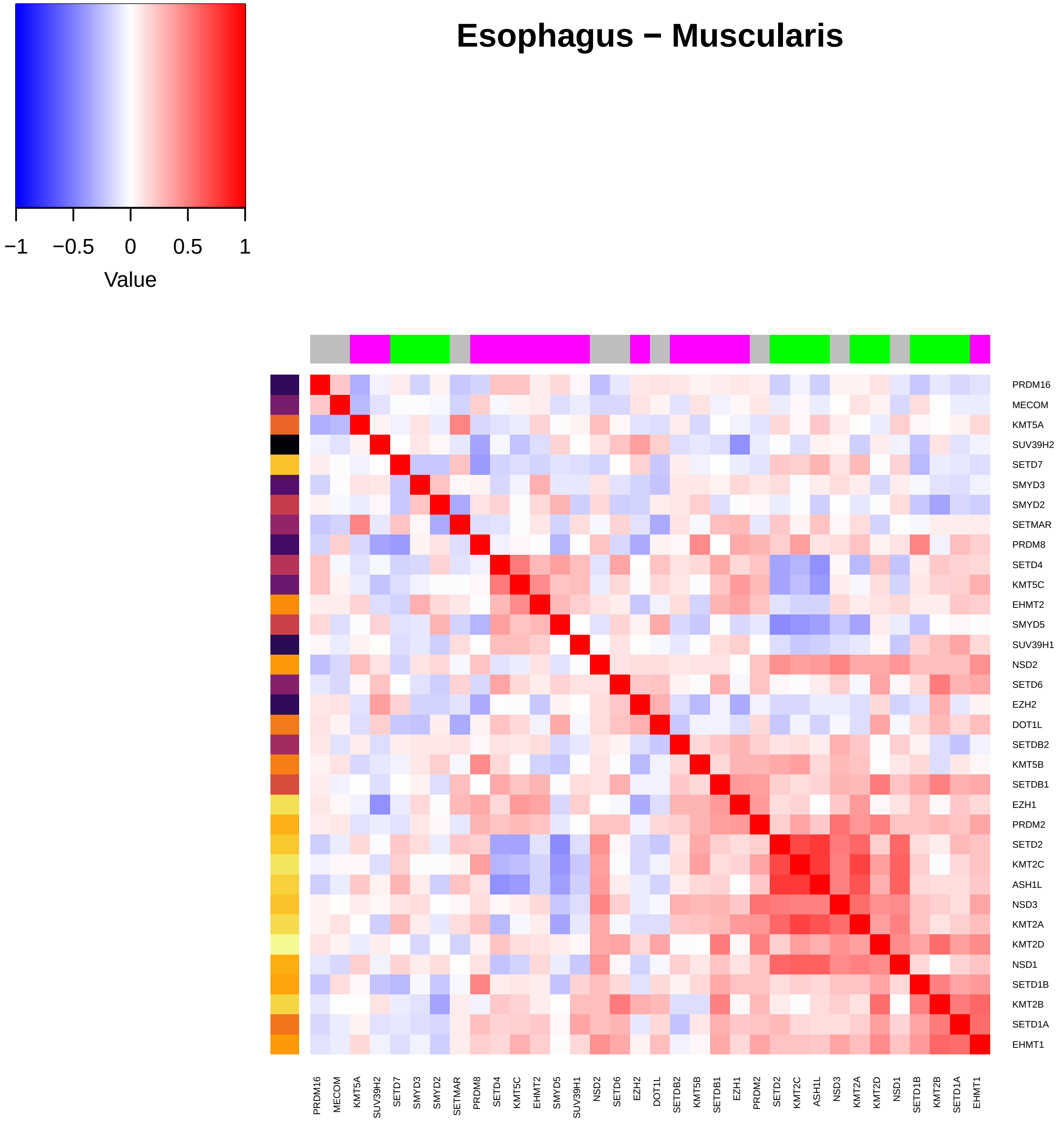

Value
−1
−0.5
0
0.5
1
Esophagus − Muscularis
PRDM16
MECOM
KMT5A
SUV39H2
SETD7
SMYD3
SMYD2
SETMAR
PRDM8
SETD4
KMT5C
EHMT2
SMYD5
SUV39H1
NSD2
SETD6
EZH2
DOT1L
SETDB2
KMT5B
SETDB1
EZH1
PRDM2
SETD2
KMT2C
ASH1L
NSD3
KMT2A
KMT2D
NSD1
SETD1B
KMT2B
SETD1A
EHMT1

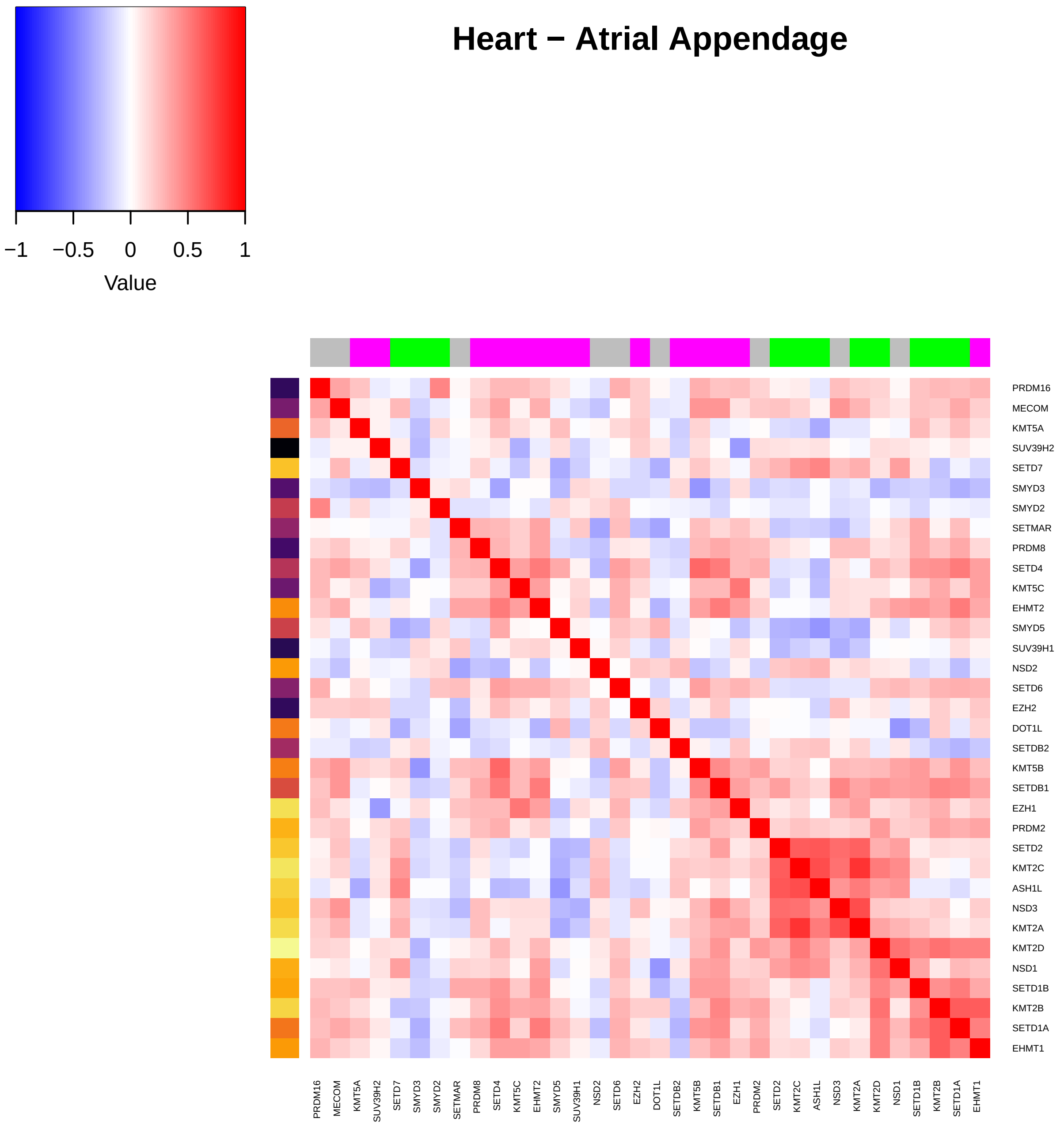
Heart − Atrial Appendage
−1
−0.5
0
0.5
1
Value
PRDM16
MECOM
KMT5A
SUV39H2
SETD7
SMYD3
SMYD2
SETMAR
PRDM8
SETD4
KMT5C
EHMT2
SMYD5
SUV39H1
NSD2
SETD6
EZH2
DOT1L
SETDB2
KMT5B
SETDB1
EZH1
PRDM2
SETD2
KMT2C
ASH1L
NSD3
KMT2A
KMT2D
NSD1
SETD1B
KMT2B
SETD1A
EHMT1

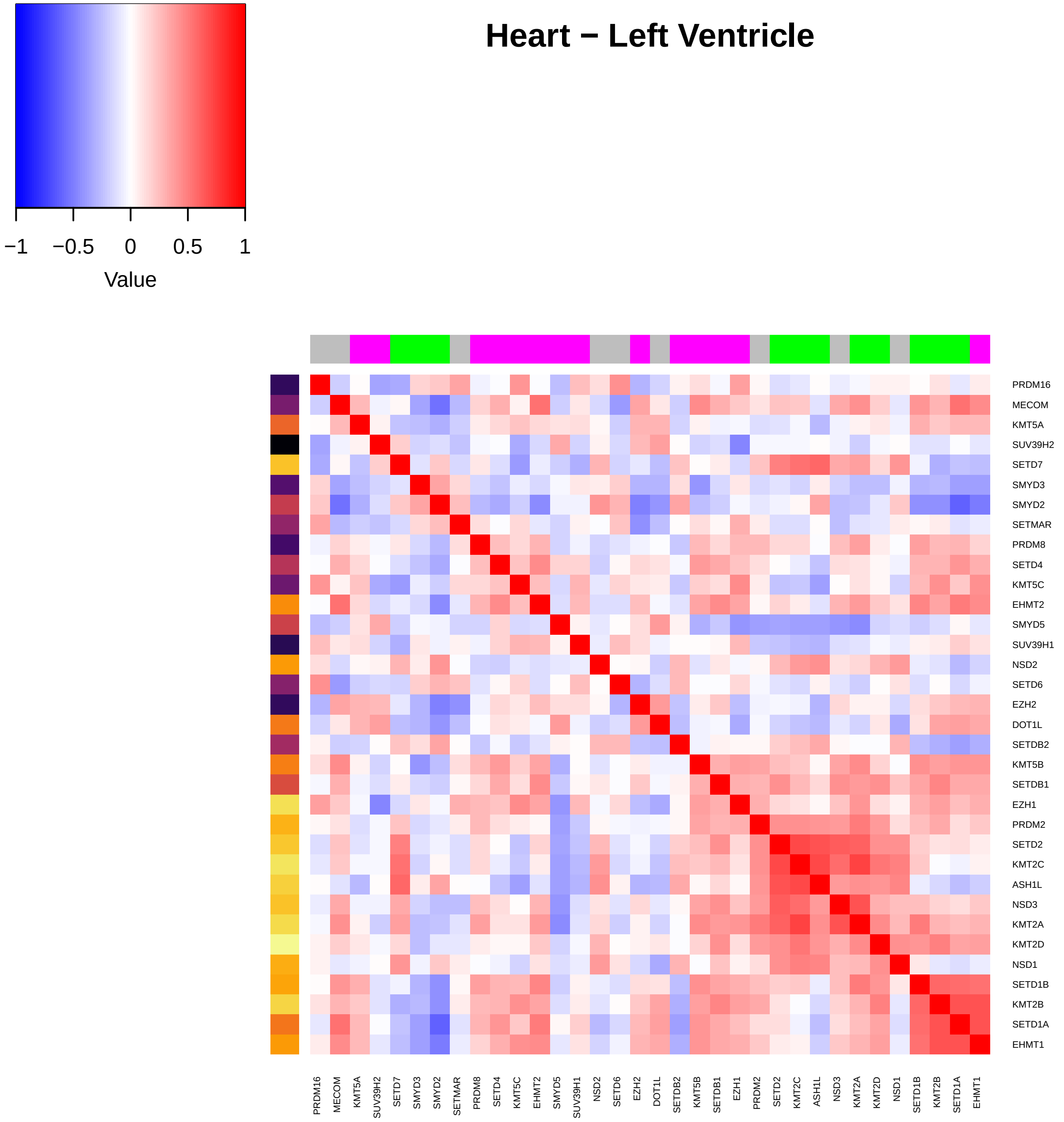

Heart − Left Ventricle
−1
−0.5
0
0.5
1
Value
PRDM16
MECOM
KMT5A
SUV39H2
SETD7
SMYD3
SMYD2
SETMAR
PRDM8
SETD4
KMT5C
EHMT2
SMYD5
SUV39H1
NSD2
SETD6
EZH2
DOT1L
SETDB2
KMT5B
SETDB1
EZH1
PRDM2
SETD2
KMT2C
ASH1L
NSD3
KMT2A
KMT2D
NSD1
SETD1B
KMT2B
SETD1A
EHMT1

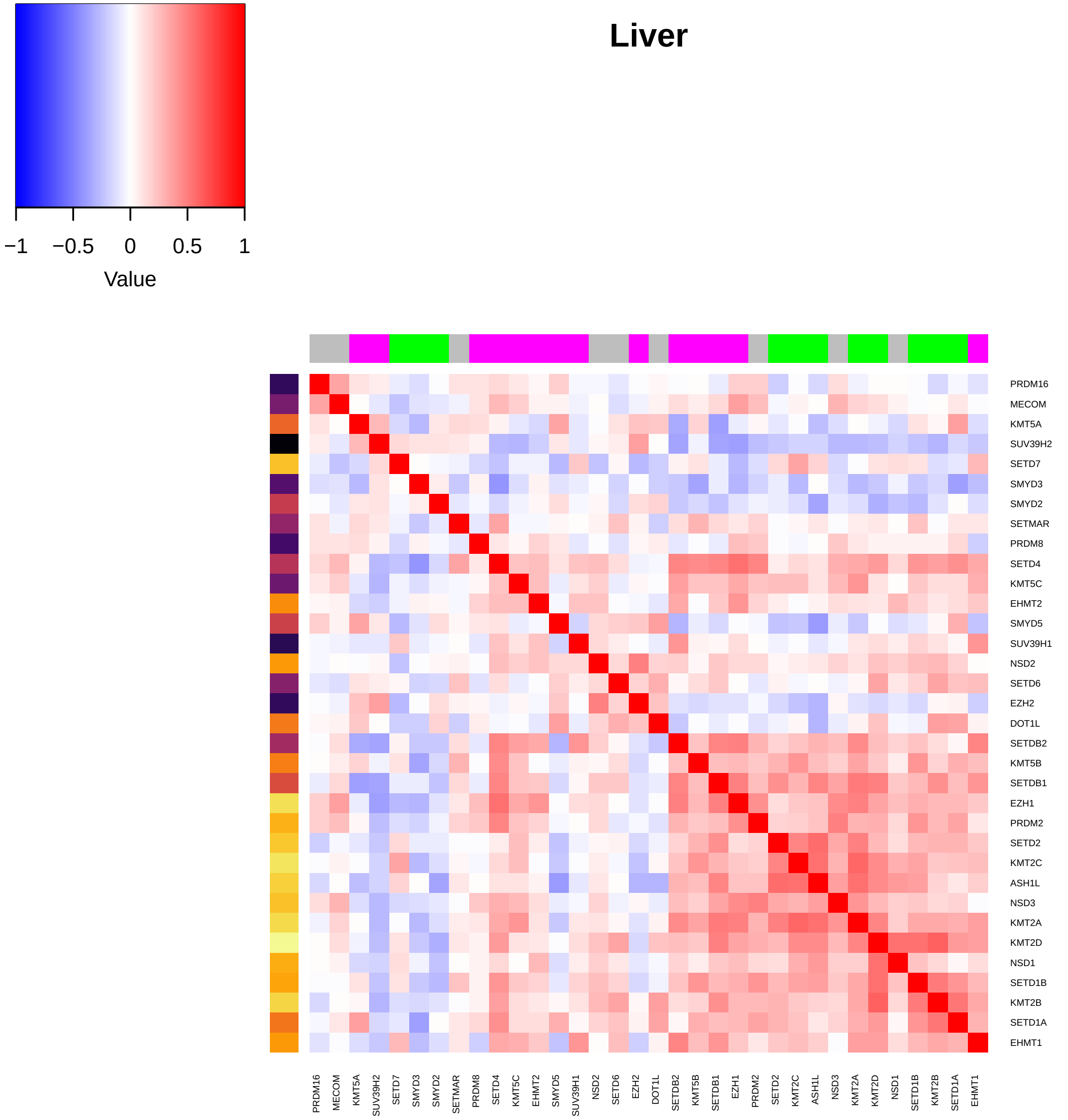

Liver
−1
−0.5
0
0.5
1
Value
PRDM16
MECOM
KMT5A
SUV39H2
SETD7
SMYD3
SMYD2
SETMAR
PRDM8
SETD4
KMT5C
EHMT2
SMYD5
SUV39H1
NSD2
SETD6
EZH2
DOT1L
SETDB2
KMT5B
SETDB1
EZH1
PRDM2
SETD2
KMT2C
ASH1L
NSD3
KMT2A
KMT2D
NSD1
SETD1B
KMT2B
SETD1A
EHMT1

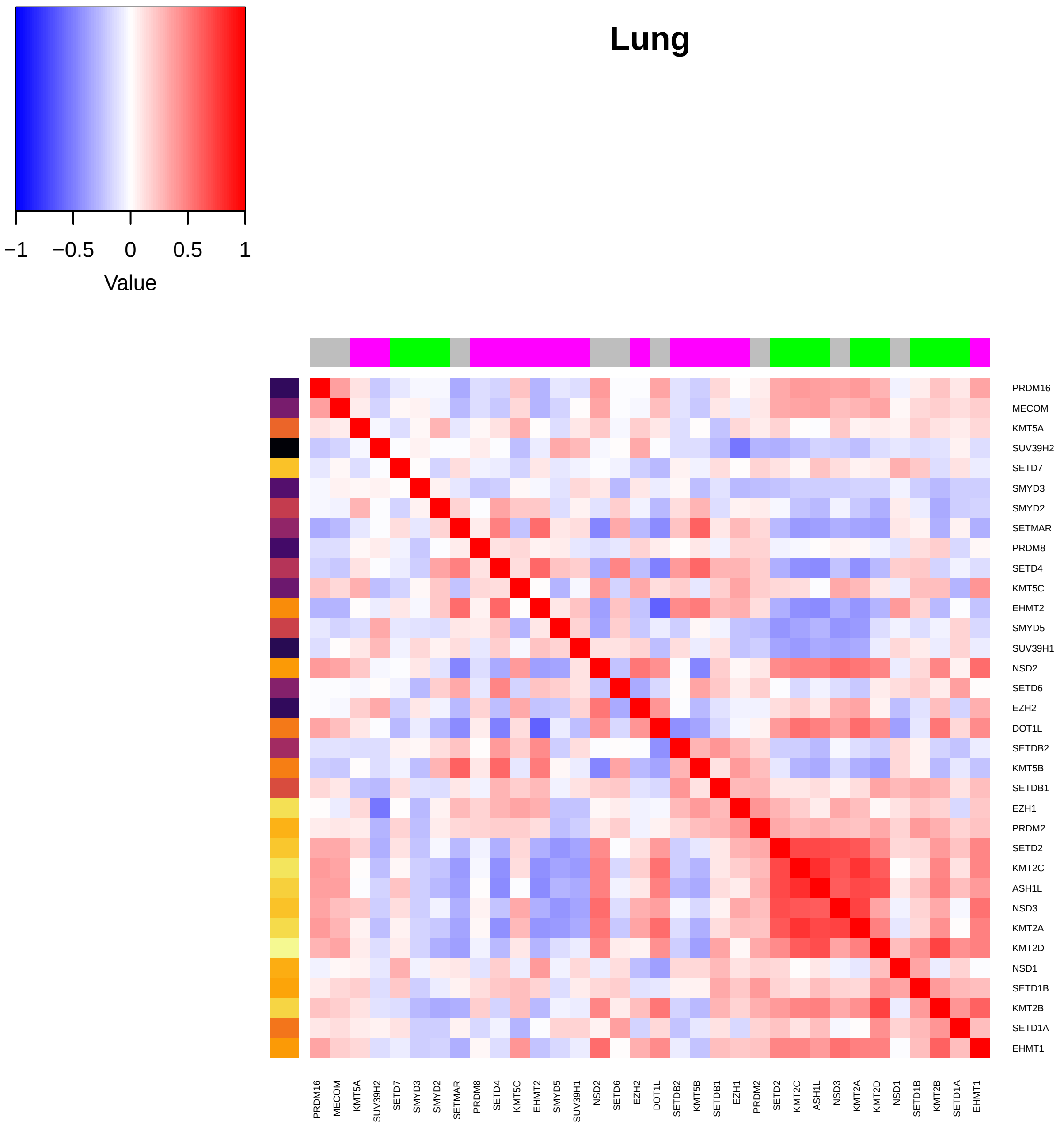
Lung
Value
−1
−0.5
0
0.5
1
PRDM16
MECOM
KMT5A
SUV39H2
SETD7
SMYD3
SMYD2
SETMAR
PRDM8
SETD4
KMT5C
EHMT2
SMYD5
SUV39H1
NSD2
SETD6
EZH2
DOT1L
SETDB2
KMT5B
SETDB1
EZH1
PRDM2
SETD2
KMT2C
ASH1L
NSD3
KMT2A
KMT2D
NSD1
SETD1B
KMT2B
SETD1A
EHMT1

### Minor Salivary Gland

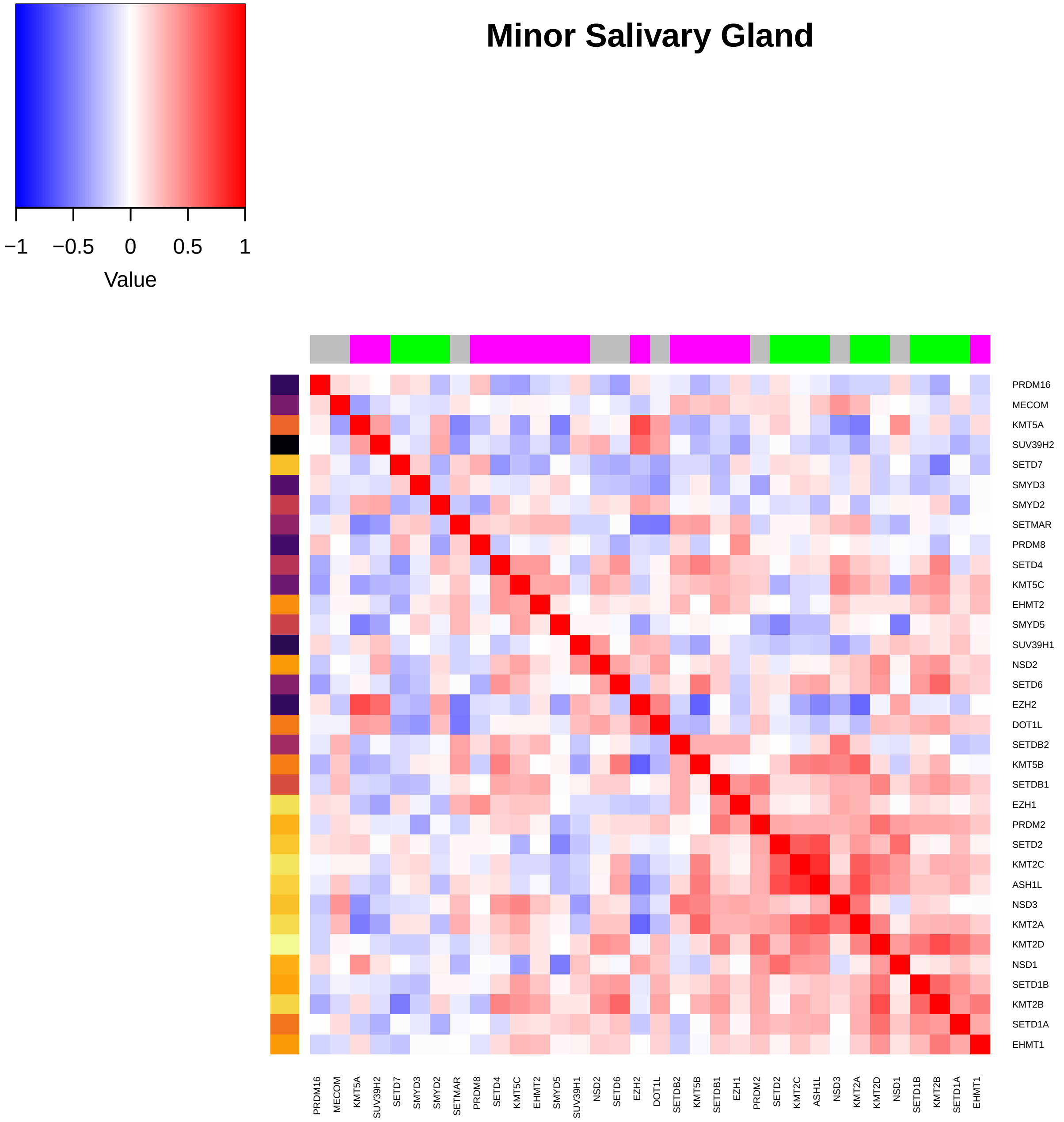

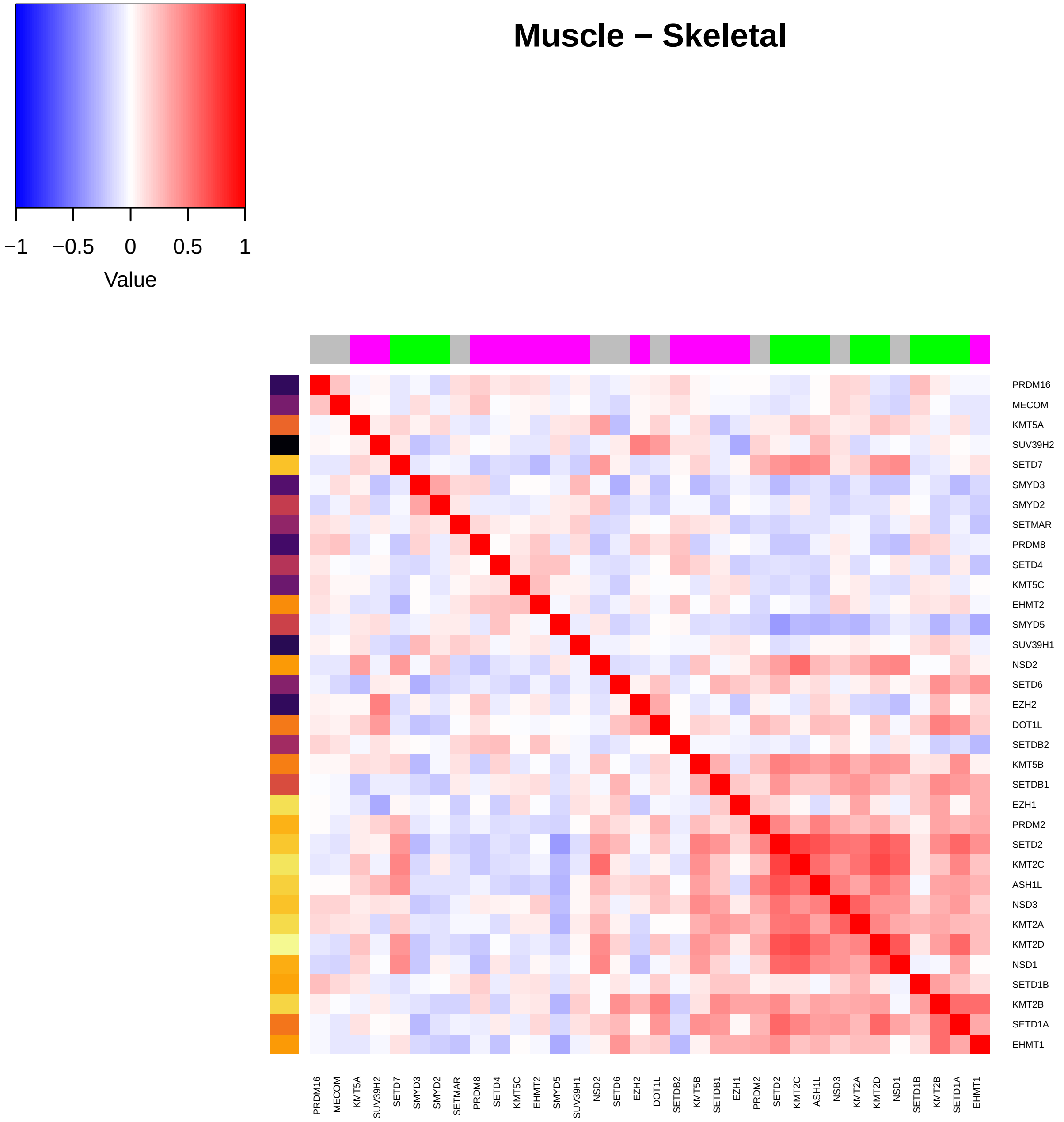

Muscle − Skeletal
Value
−1
−0.5
0
0.5
1
PRDM16
MECOM
KMT5A
SUV39H2
SETD7
SMYD3
SMYD2
SETMAR
PRDM8
SETD4
KMT5C
EHMT2
SMYD5
SUV39H1
NSD2
SETD6
EZH2
DOT1L
SETDB2
KMT5B
SETDB1
EZH1
PRDM2
SETD2
KMT2C
ASH1L
NSD3
KMT2A
KMT2D
NSD1
SETD1B
KMT2B
SETD1A
EHMT1

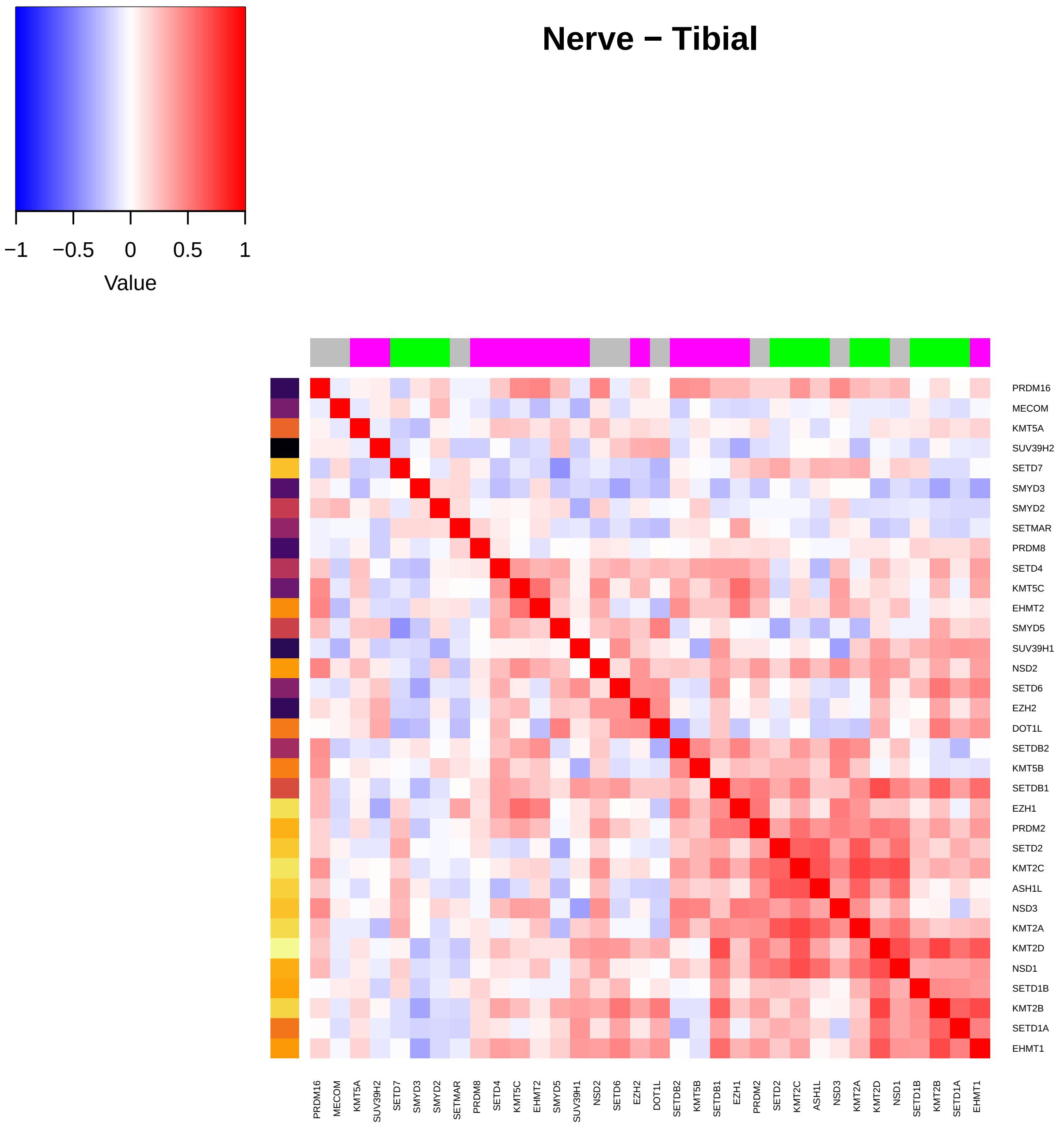

Nerve − Tibial
−1
−0.5
0
0.5
1
Value
PRDM16
MECOM
KMT5A
SUV39H2
SETD7
SMYD3
SMYD2
SETMAR
PRDM8
SETD4
KMT5C
EHMT2
SMYD5
SUV39H1
NSD2
SETD6
EZH2
DOT1L
SETDB2
KMT5B
SETDB1
EZH1
PRDM2
SETD2
KMT2C
ASH1L
NSD3
KMT2A
KMT2D
NSD1
SETD1B
KMT2B
SETD1A
EHMT1

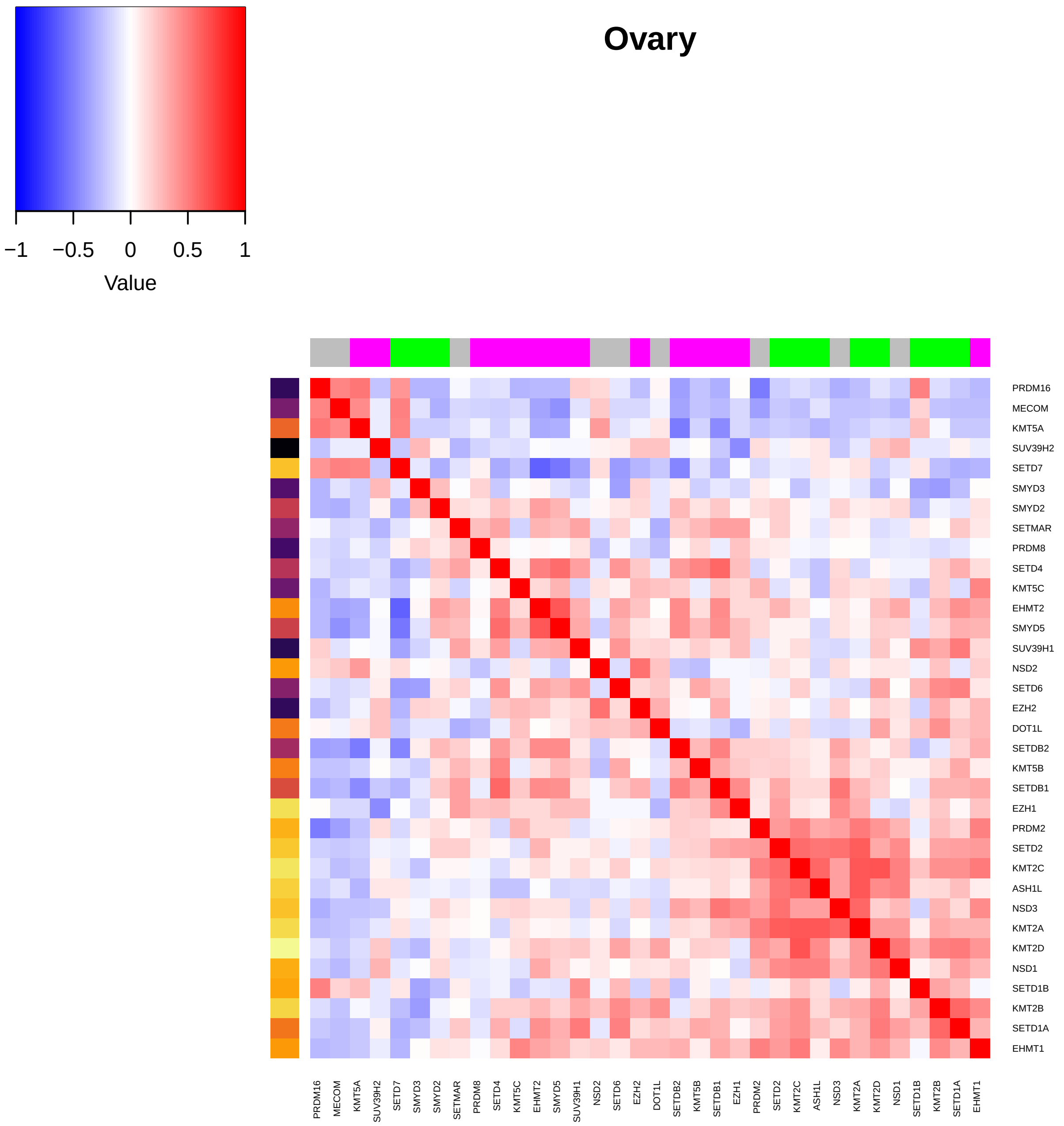

Ovary
−1
−0.5
0
0.5
1
Value
PRDM16
MECOM
KMT5A
SUV39H2
SETD7
SMYD3
SMYD2
SETMAR
PRDM8
SETD4
KMT5C
EHMT2
SMYD5
SUV39H1
NSD2
SETD6
EZH2
DOT1L
SETDB2
KMT5B
SETDB1
EZH1
PRDM2
SETD2
KMT2C
ASH1L
NSD3
KMT2A
KMT2D
NSD1
SETD1B
KMT2B
SETD1A
EHMT1

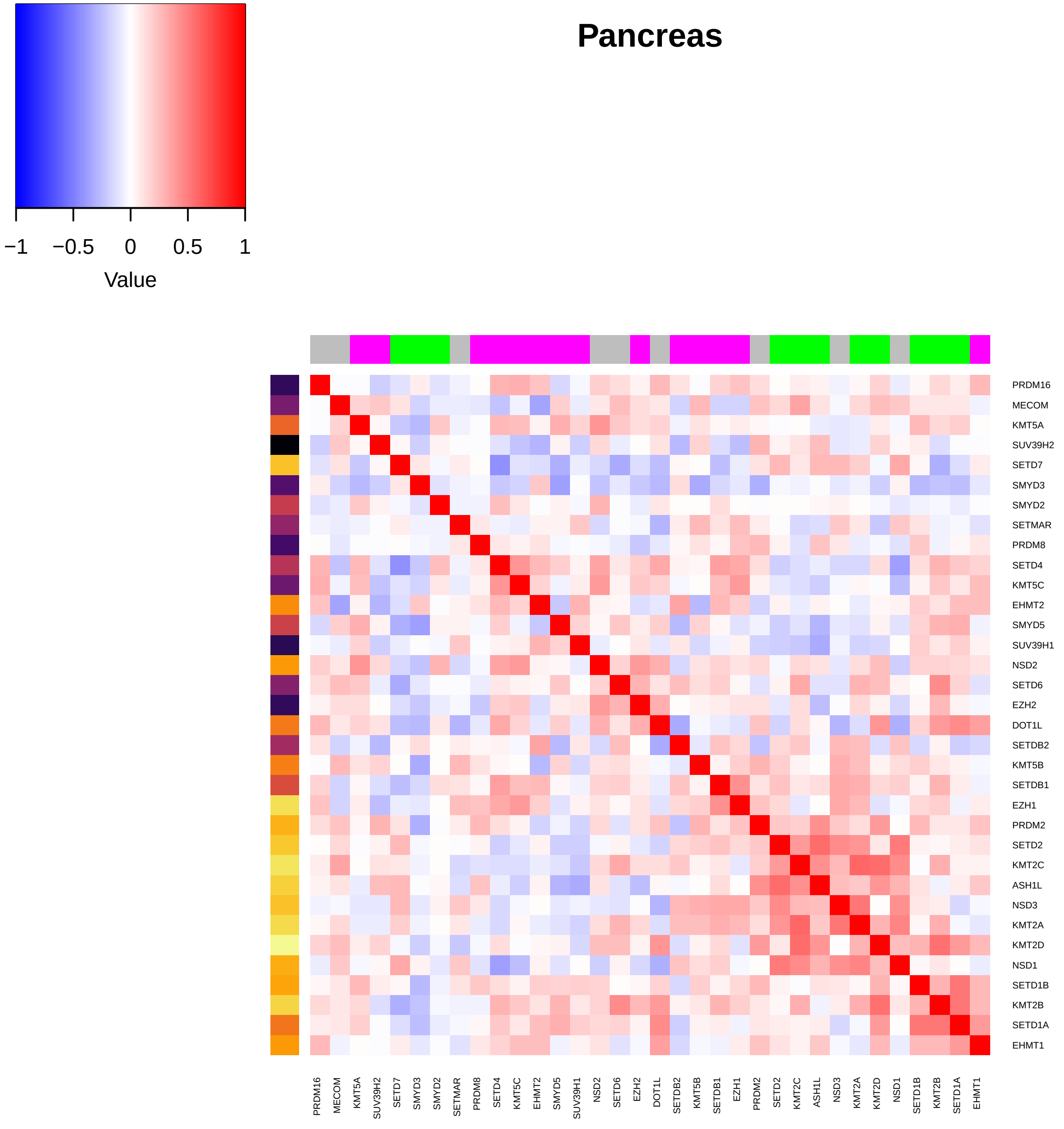

Pancreas
−1
−0.5
0
0.5
1
Value
PRDM16
MECOM
KMT5A
SUV39H2
SETD7
SMYD3
SMYD2
SETMAR
PRDM8
SETD4
KMT5C
EHMT2
SMYD5
SUV39H1
NSD2
SETD6
EZH2
DOT1L
SETDB2
KMT5B
SETDB1
EZH1
PRDM2
SETD2
KMT2C
ASH1L
NSD3
KMT2A
KMT2D
NSD1
SETD1B
KMT2B
SETD1A
EHMT1

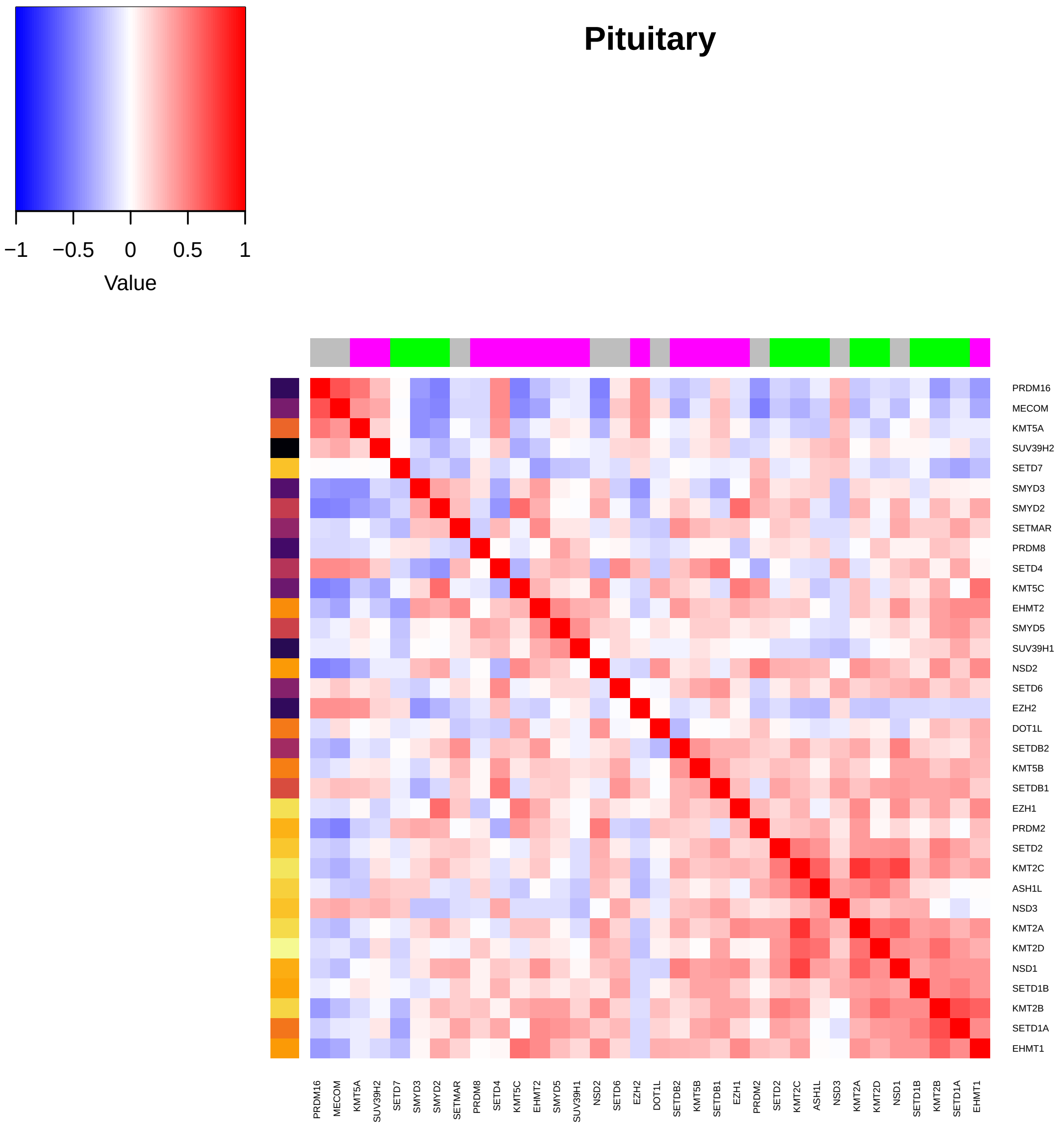

Pituitary
−1
−0.5
0
0.5
1
Value
PRDM16
MECOM
KMT5A
SUV39H2
SETD7
SMYD3
SMYD2
SETMAR
PRDM8
SETD4
KMT5C
EHMT2
SMYD5
SUV39H1
NSD2
SETD6
EZH2
DOT1L
SETDB2
KMT5B
SETDB1
EZH1
PRDM2
SETD2
KMT2C
ASH1L
NSD3
KMT2A
KMT2D
NSD1
SETD1B
KMT2B
SETD1A
EHMT1

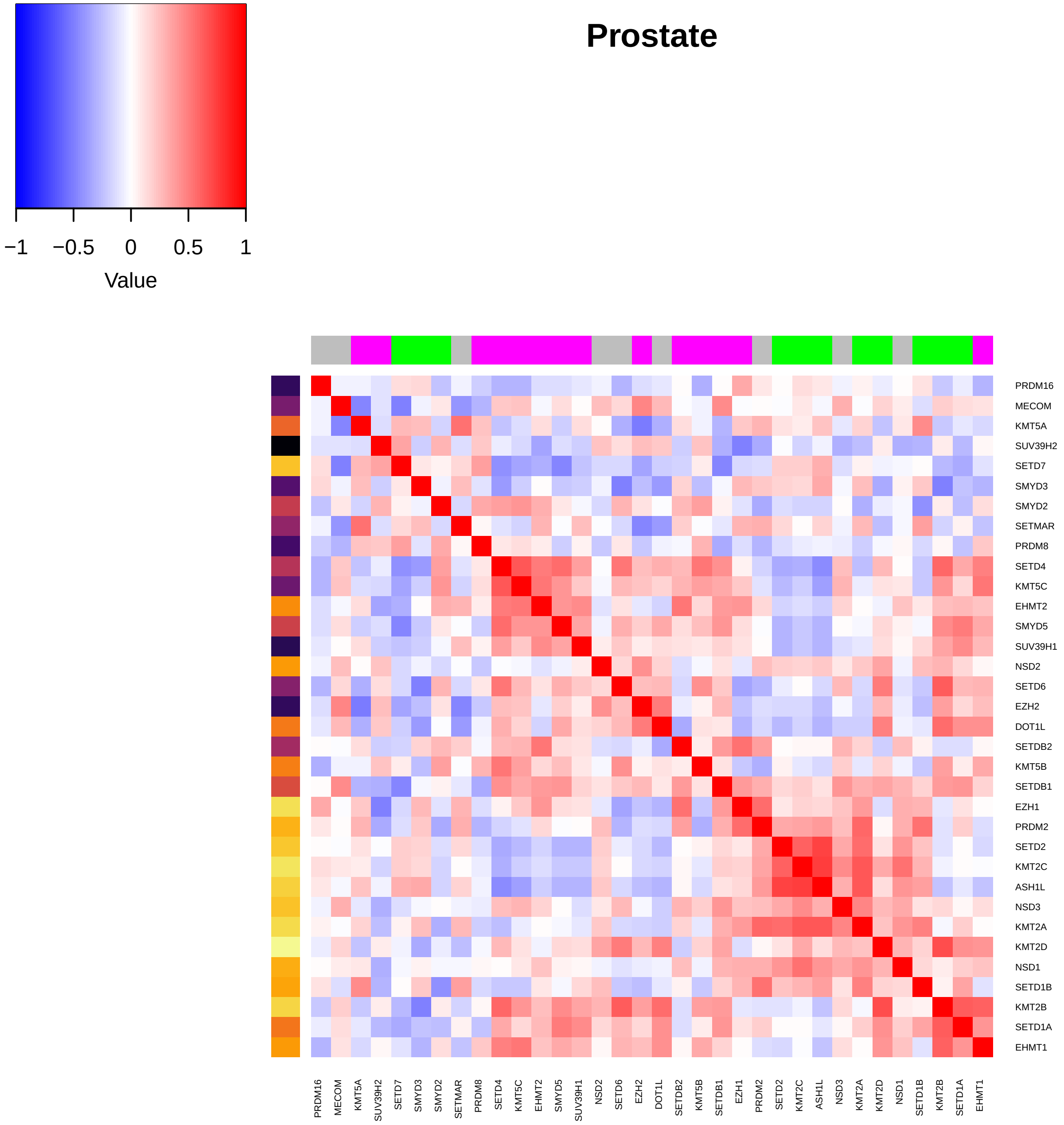
Prostate
−1
−0.5
0
0.5
1
Value
PRDM16
MECOM
KMT5A
SUV39H2
SETD7
SMYD3
SMYD2
SETMAR
PRDM8
SETD4
KMT5C
EHMT2
SMYD5
SUV39H1
NSD2
SETD6
EZH2
DOT1L
SETDB2
KMT5B
SETDB1
EZH1
PRDM2
SETD2
KMT2C
ASH1L
NSD3
KMT2A
KMT2D
NSD1
SETD1B
KMT2B
SETD1A
EHMT1

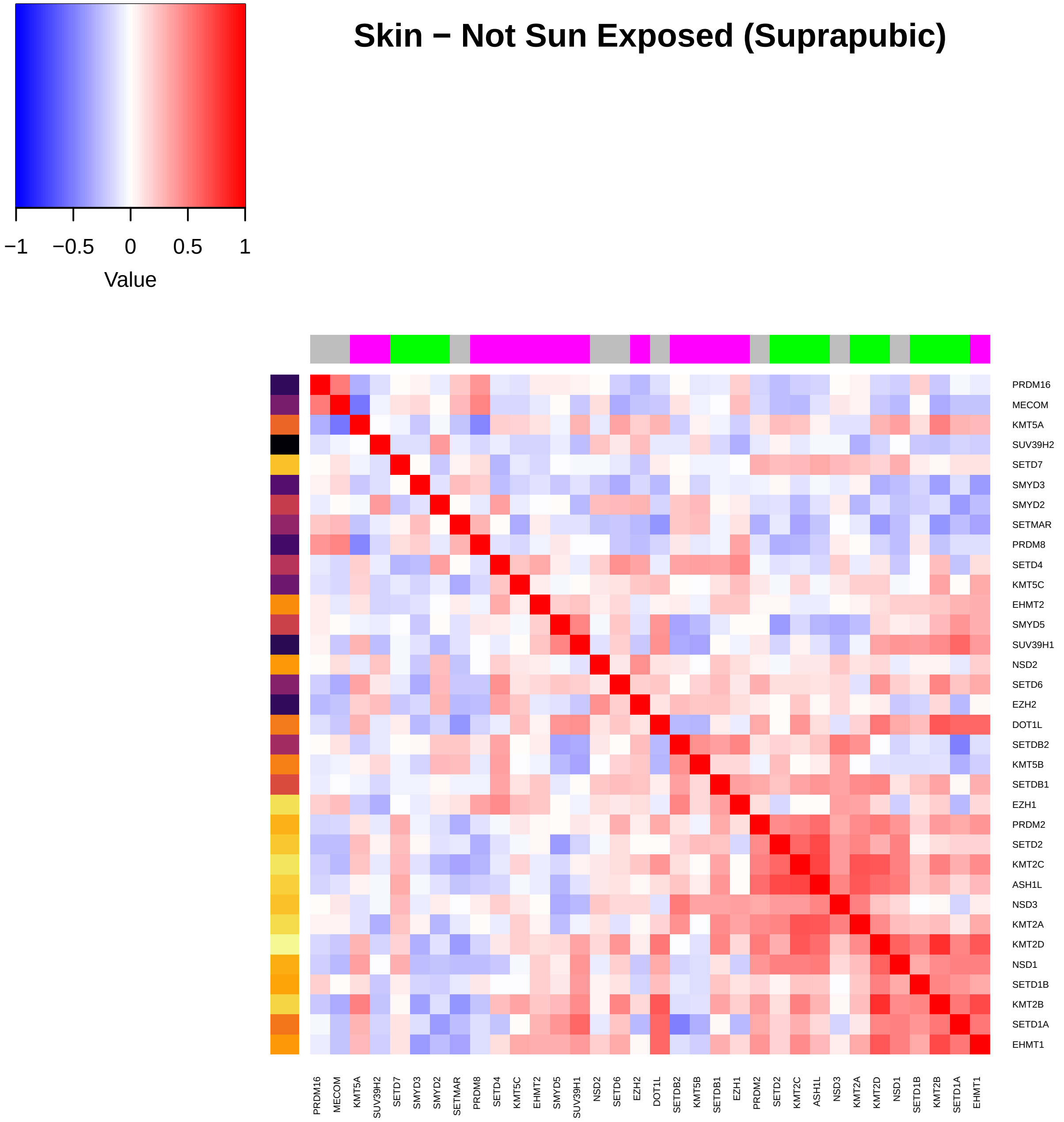

Skin − Not Sun Exposed (Suprapubic)
−1
−0.5
0
0.5
1
Value
PRDM16
MECOM
KMT5A
SUV39H2
SETD7
SMYD3
SMYD2
SETMAR
PRDM8
SETD4
KMT5C
EHMT2
SMYD5
SUV39H1
NSD2
SETD6
EZH2
DOT1L
SETDB2
KMT5B
SETDB1
EZH1
PRDM2
SETD2
KMT2C
ASH1L
NSD3
KMT2A
KMT2D
NSD1
SETD1B
KMT2B
SETD1A
EHMT1

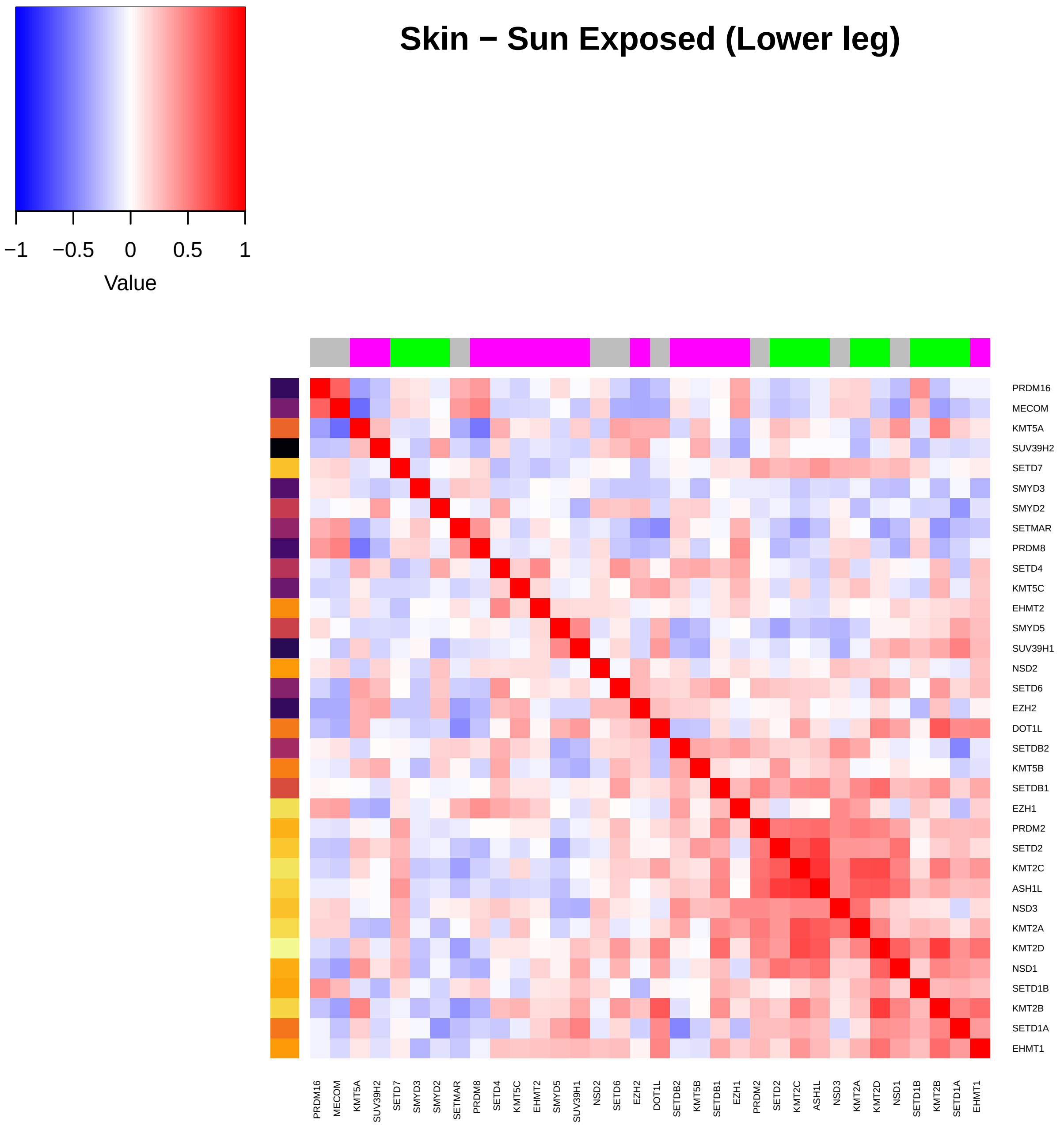

Skin − Sun Exposed (Lower leg)
−1
−0.5
0
0.5
1
Value
PRDM16
MECOM
KMT5A
SUV39H2
SETD7
SMYD3
SMYD2
SETMAR
PRDM8
SETD4
KMT5C
EHMT2
SMYD5
SUV39H1
NSD2
SETD6
EZH2
DOT1L
SETDB2
KMT5B
SETDB1
EZH1
PRDM2
SETD2
KMT2C
ASH1L
NSD3
KMT2A
KMT2D
NSD1
SETD1B
KMT2B
SETD1A
EHMT1

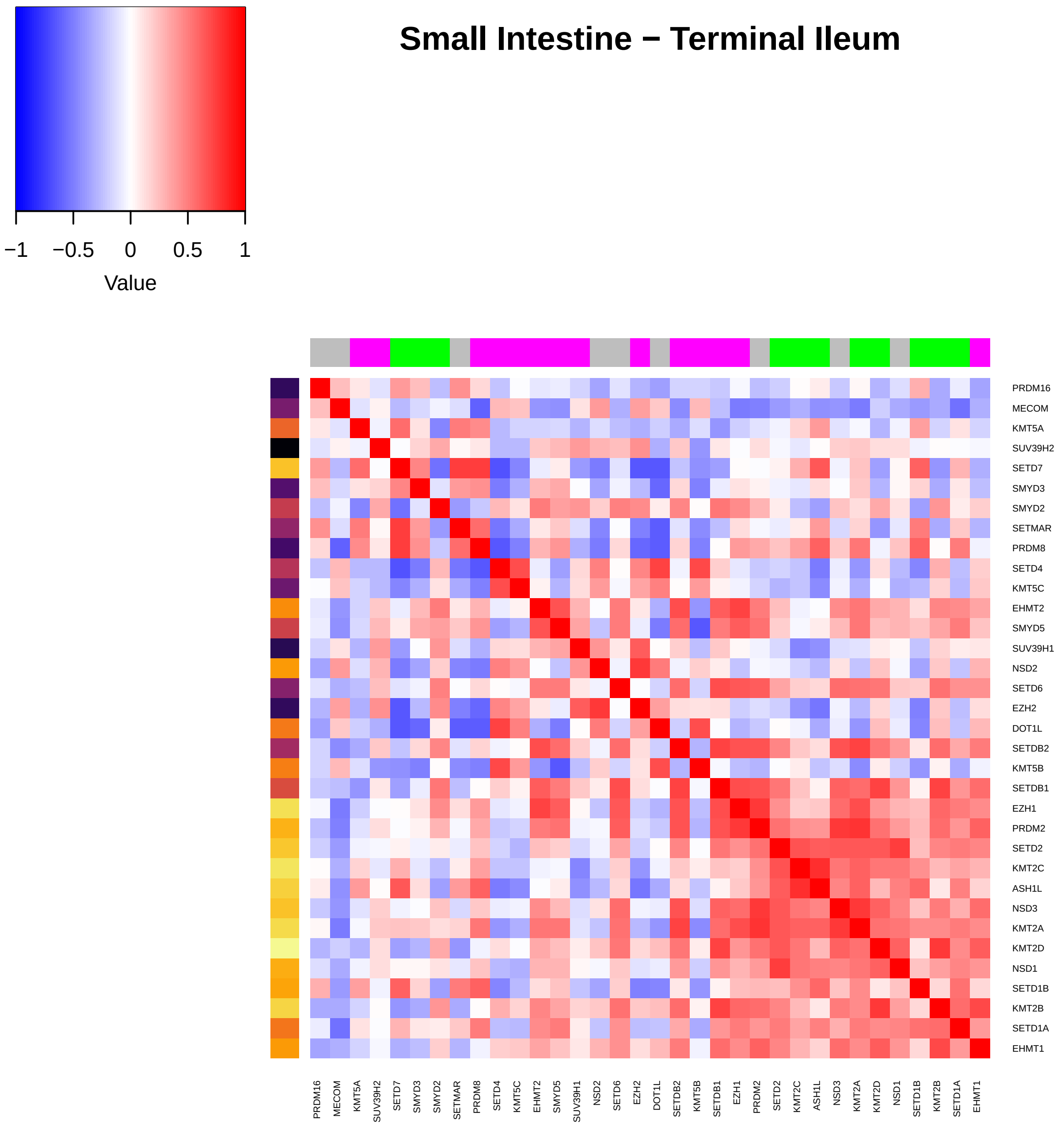
Small Intestine − Terminal Ileum
Value
−1
−0.5
0
0.5
1
PRDM16
MECOM
KMT5A
SUV39H2
SETD7
SMYD3
SMYD2
SETMAR
PRDM8
SETD4
KMT5C
EHMT2
SMYD5
SUV39H1
NSD2
SETD6
EZH2
DOT1L
SETDB2
KMT5B
SETDB1
EZH1
PRDM2
SETD2
KMT2C
ASH1L
NSD3
KMT2A
KMT2D
NSD1
SETD1B
KMT2B
SETD1A
EHMT1

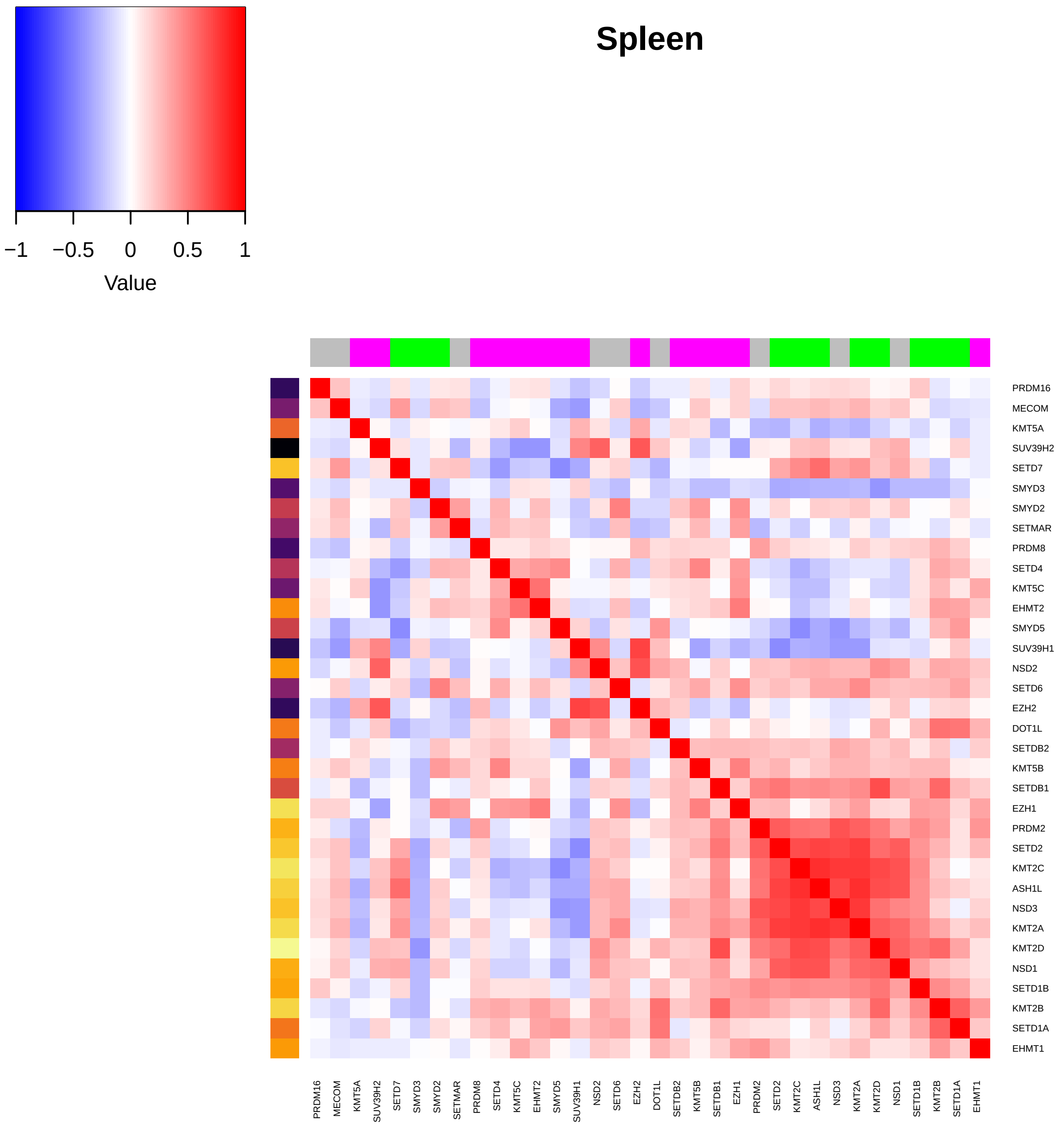
Spleen
−1
−0.5
0
0.5
1
Value
PRDM16
MECOM
KMT5A
SUV39H2
SETD7
SMYD3
SMYD2
SETMAR
PRDM8
SETD4
KMT5C
EHMT2
SMYD5
SUV39H1
NSD2
SETD6
EZH2
DOT1L
SETDB2
KMT5B
SETDB1
EZH1
PRDM2
SETD2
KMT2C
ASH1L
NSD3
KMT2A
KMT2D
NSD1
SETD1B
KMT2B
SETD1A
EHMT1

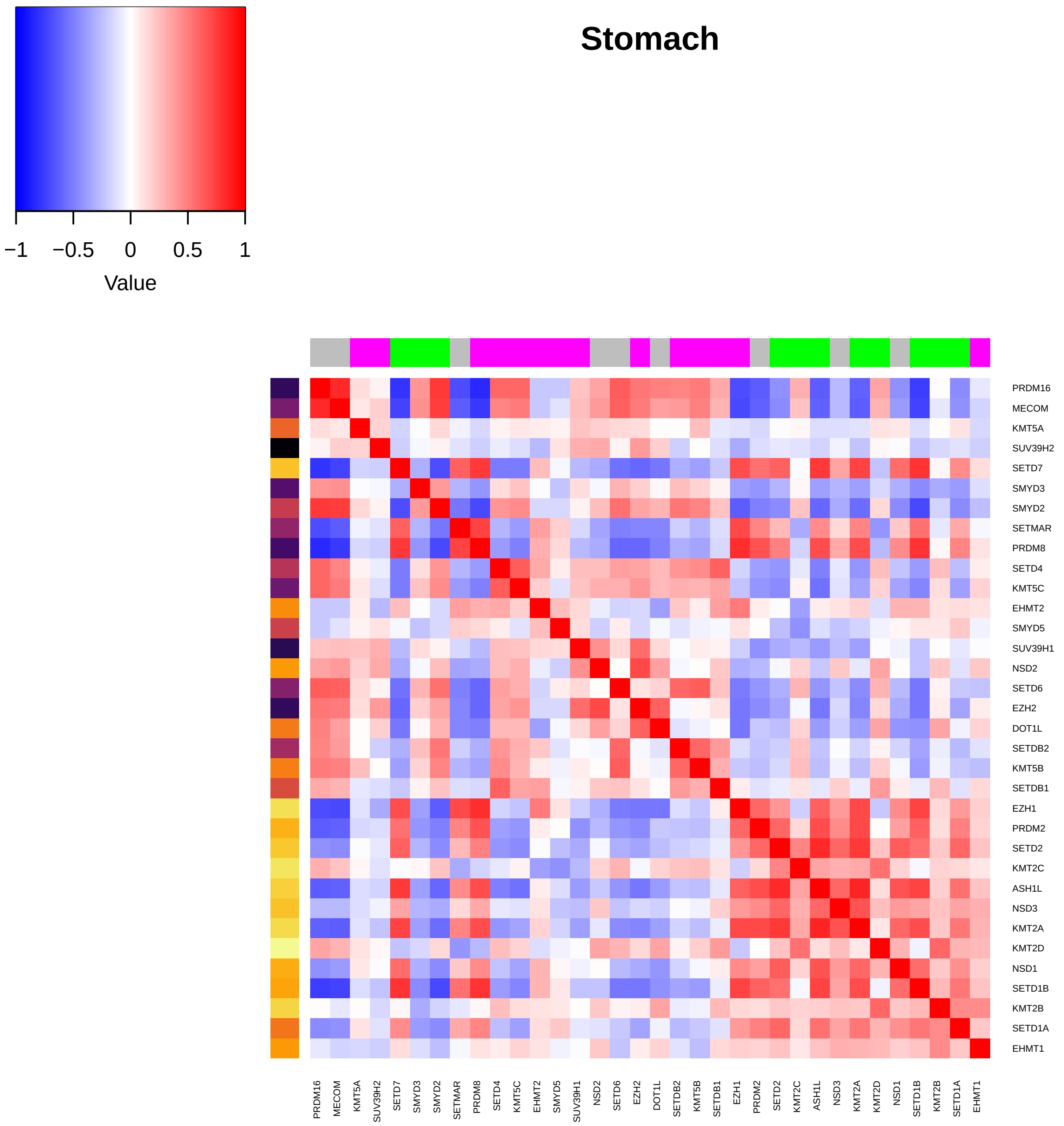

Stomach
−1
−0.5
0
0.5
1
Value
PRDM16
MECOM
KMT5A
SUV39H2
SETD7
SMYD3
SMYD2
SETMAR
PRDM8
SETD4
KMT5C
EHMT2
SMYD5
SUV39H1
NSD2
SETD6
EZH2
DOT1L
SETDB2
KMT5B
SETDB1
EZH1
PRDM2
SETD2
KMT2C
ASH1L
NSD3
KMT2A
KMT2D
NSD1
SETD1B
KMT2B
SETD1A
EHMT1

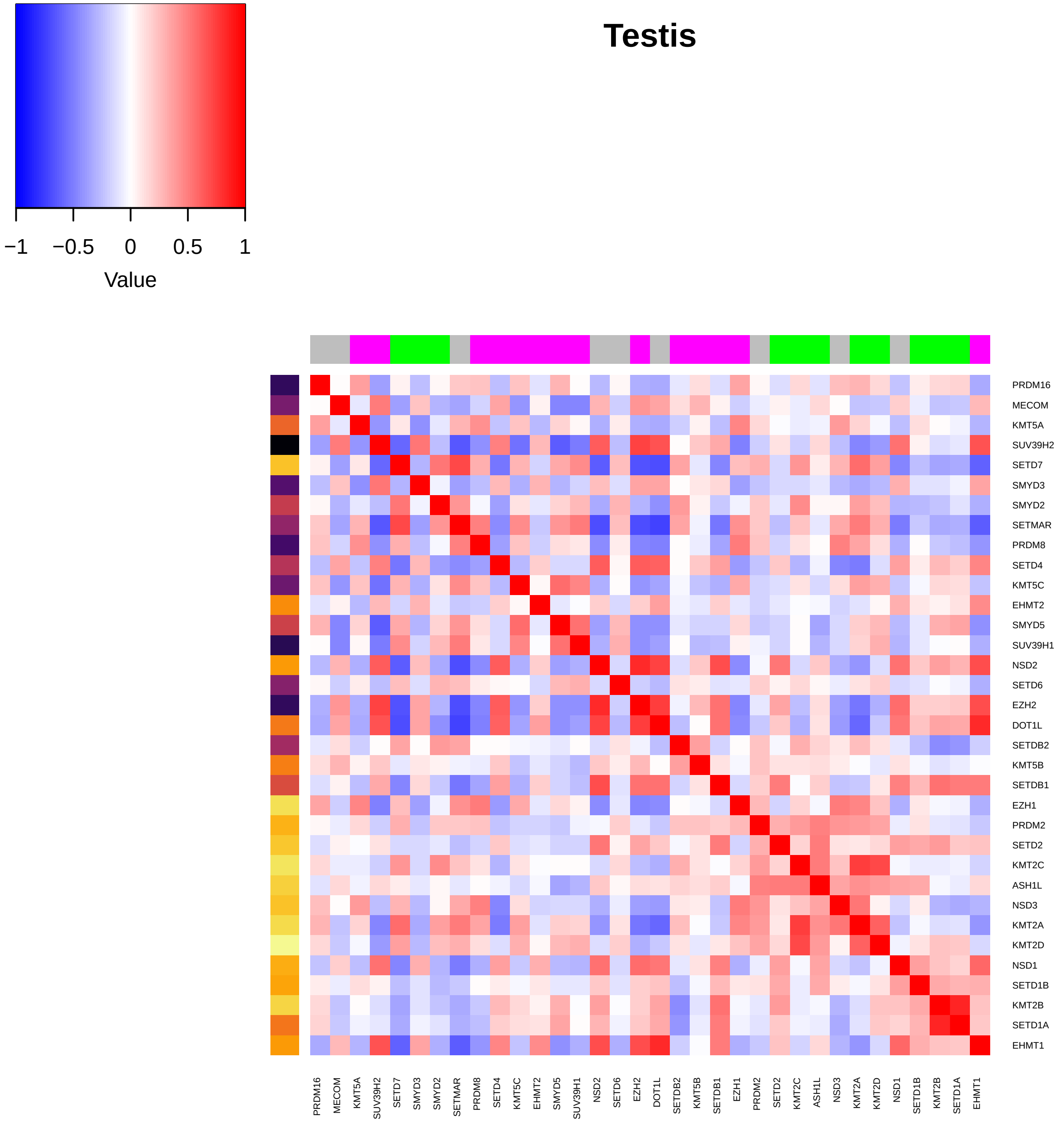
Testis
−1
−0.5
0
0.5
1
Value
PRDM16
MECOM
KMT5A
SUV39H2
SETD7
SMYD3
SMYD2
SETMAR
PRDM8
SETD4
KMT5C
EHMT2
SMYD5
SUV39H1
NSD2
SETD6
EZH2
DOT1L
SETDB2
KMT5B
SETDB1
EZH1
PRDM2
SETD2
KMT2C
ASH1L
NSD3
KMT2A
KMT2D
NSD1
SETD1B
KMT2B
SETD1A
EHMT1

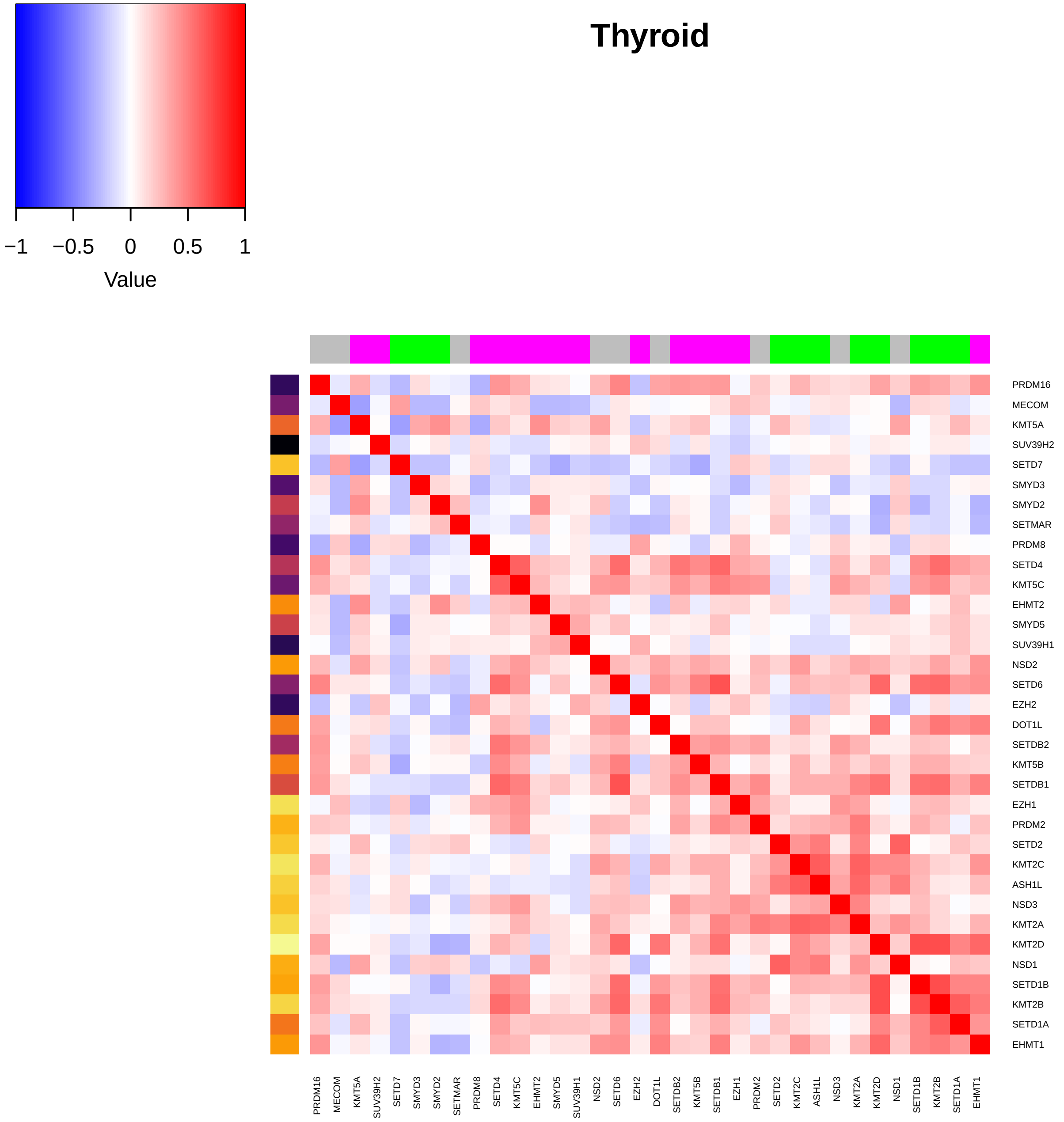
Thyroid
−1
−0.5
0
0.5
1
Value
PRDM16
MECOM
KMT5A
SUV39H2
SETD7
SMYD3
SMYD2
SETMAR
PRDM8
SETD4
KMT5C
EHMT2
SMYD5
SUV39H1
NSD2
SETD6
EZH2
DOT1L
SETDB2
KMT5B
SETDB1
EZH1
PRDM2
SETD2
KMT2C
ASH1L
NSD3
KMT2A
KMT2D
NSD1
SETD1B
KMT2B
SETD1A
EHMT1

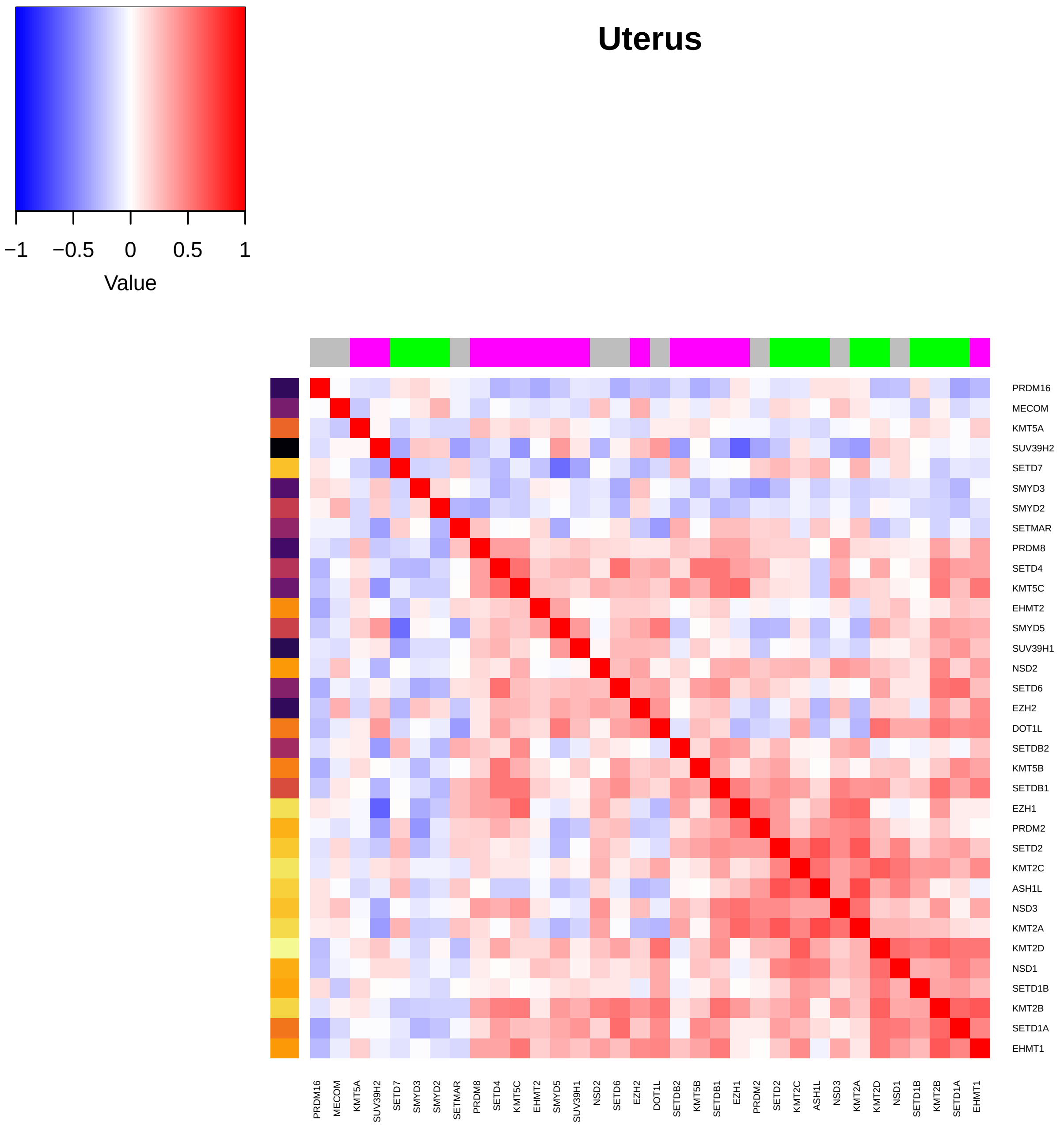
Uterus
−1 −0.5 0 0.5 1
Value
PRDM16
MECOM
KMT5A
SUV39H2
SETD7
SMYD3
SMYD2
SETMAR
PRDM8
SETD4
KMT5C
EHMT2
SMYD5
SUV39H1
NSD2
SETD6
EZH2
DOT1L
SETDB2
KMT5B
SETDB1
EZH1
PRDM2
SETD2
KMT2C
ASH1L
NSD3
KMT2A
KMT2D
NSD1
SETD1B
KMT2B
SETD1A
EHMT1

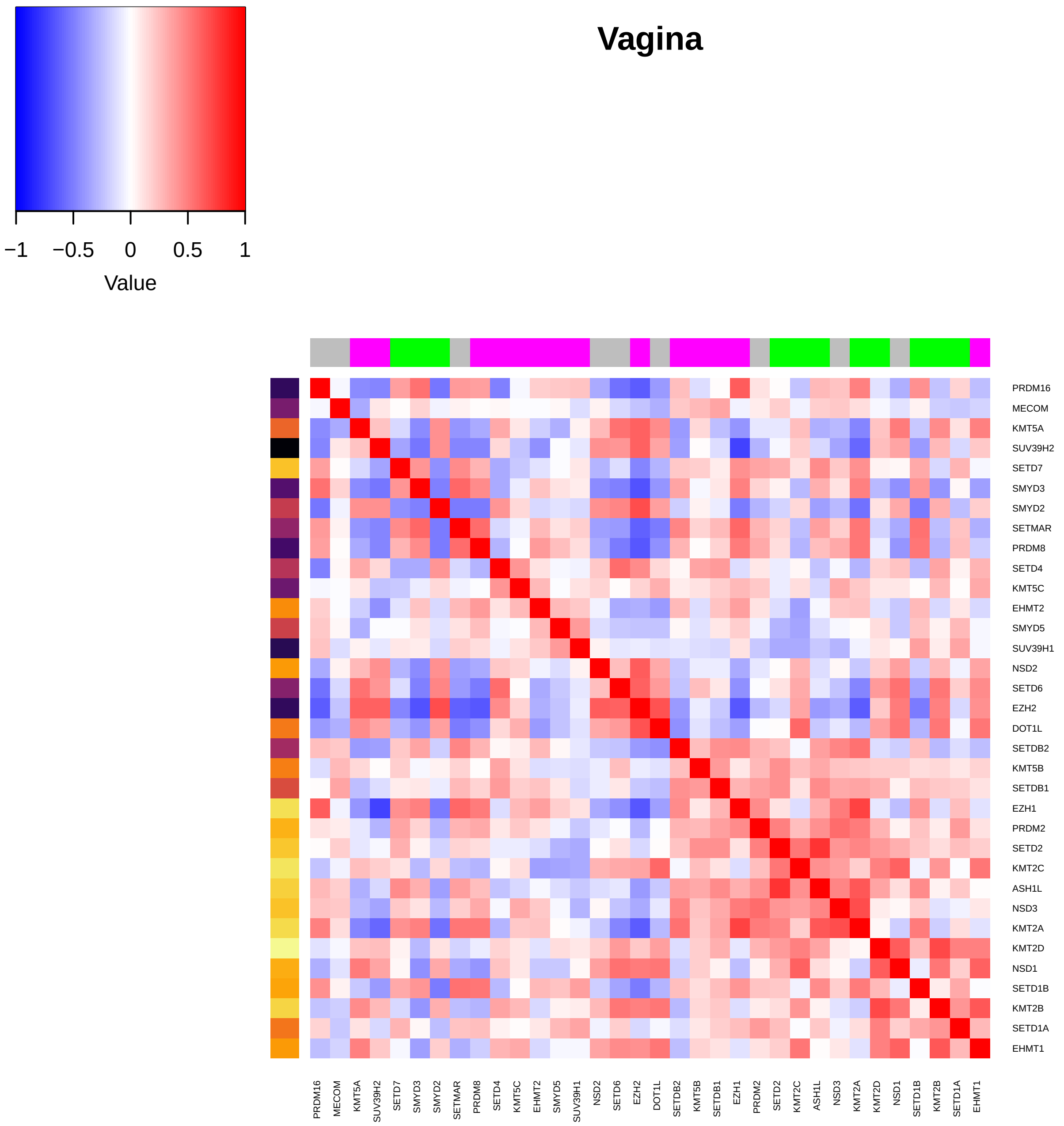
Vagina
Value
−1
−0.5
0
0.5
1
PRDM16
MECOM
KMT5A
SUV39H2
SETD7
SMYD3
SMYD2
SETMAR
PRDM8
SETD4
KMT5C
EHMT2
SMYD5
SUV39H1
NSD2
SETD6
EZH2
DOT1L
SETDB2
KMT5B
SETDB1
EZH1
PRDM2
SETD2
KMT2C
ASH1L
NSD3
KMT2A
KMT2D
NSD1
SETD1B
KMT2B
SETD1A
EHMT1

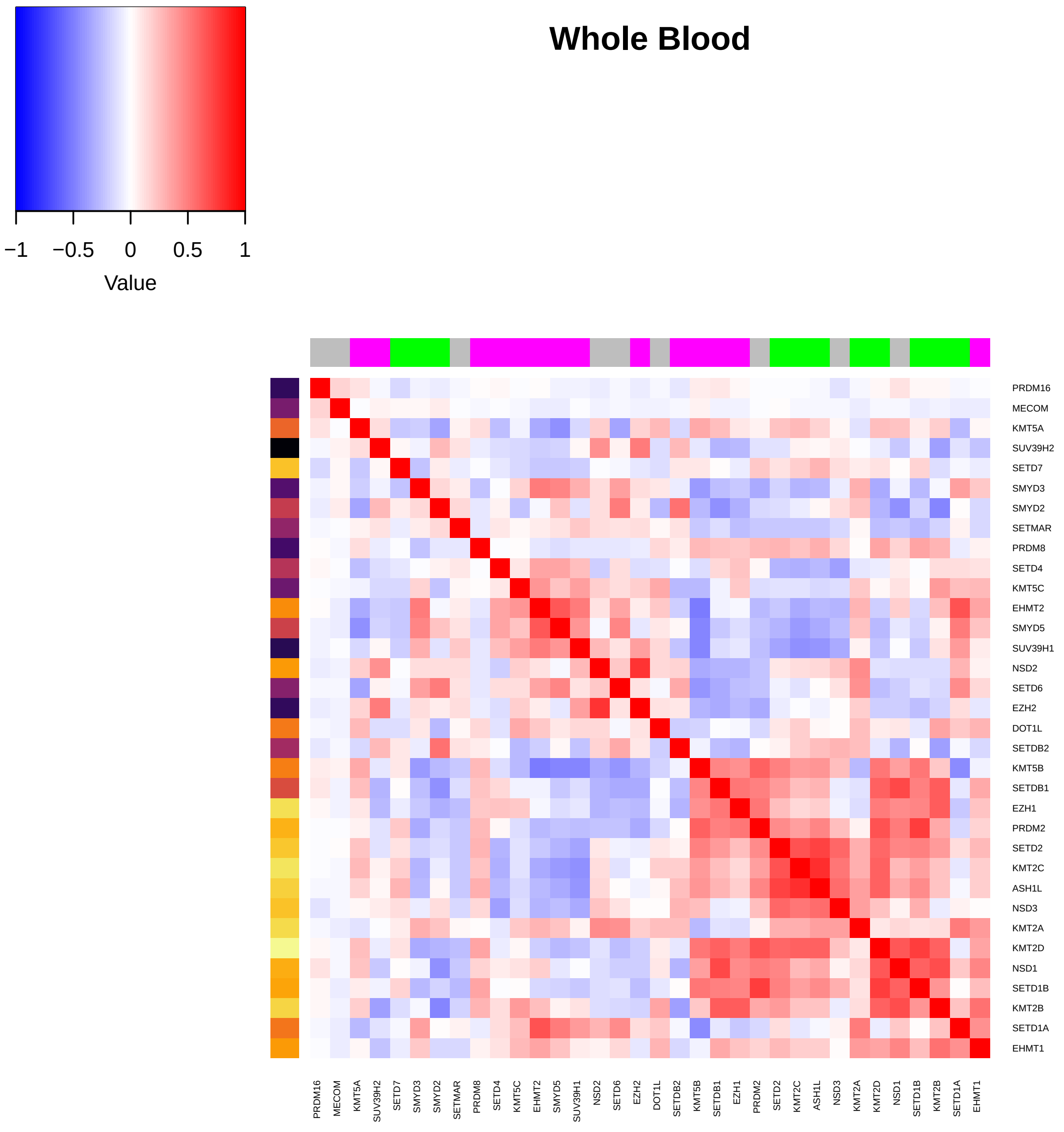
Whole Blood
Value
−1
−0.5
0
0.5
1
PRDM16
MECOM
KMT5A
SUV39H2
SETD7
SMYD3
SMYD2
SETMAR
PRDM8
SETD4
KMT5C
EHMT2
SMYD5
SUV39H1
NSD2
SETD6
EZH2
DOT1L
SETDB2
KMT5B
SETDB1
EZH1
PRDM2
SETD2
KMT2C
ASH1L
NSD3
KMT2A
KMT2D
NSD1
SETD1B
KMT2B
SETD1A
EHMT1
