## Supplemental Figures and Tables for "Histone methylation has a direct metabolic role in human cells": Supp File 6. TCGA HMTcorrelations tissues fullplots.pdf

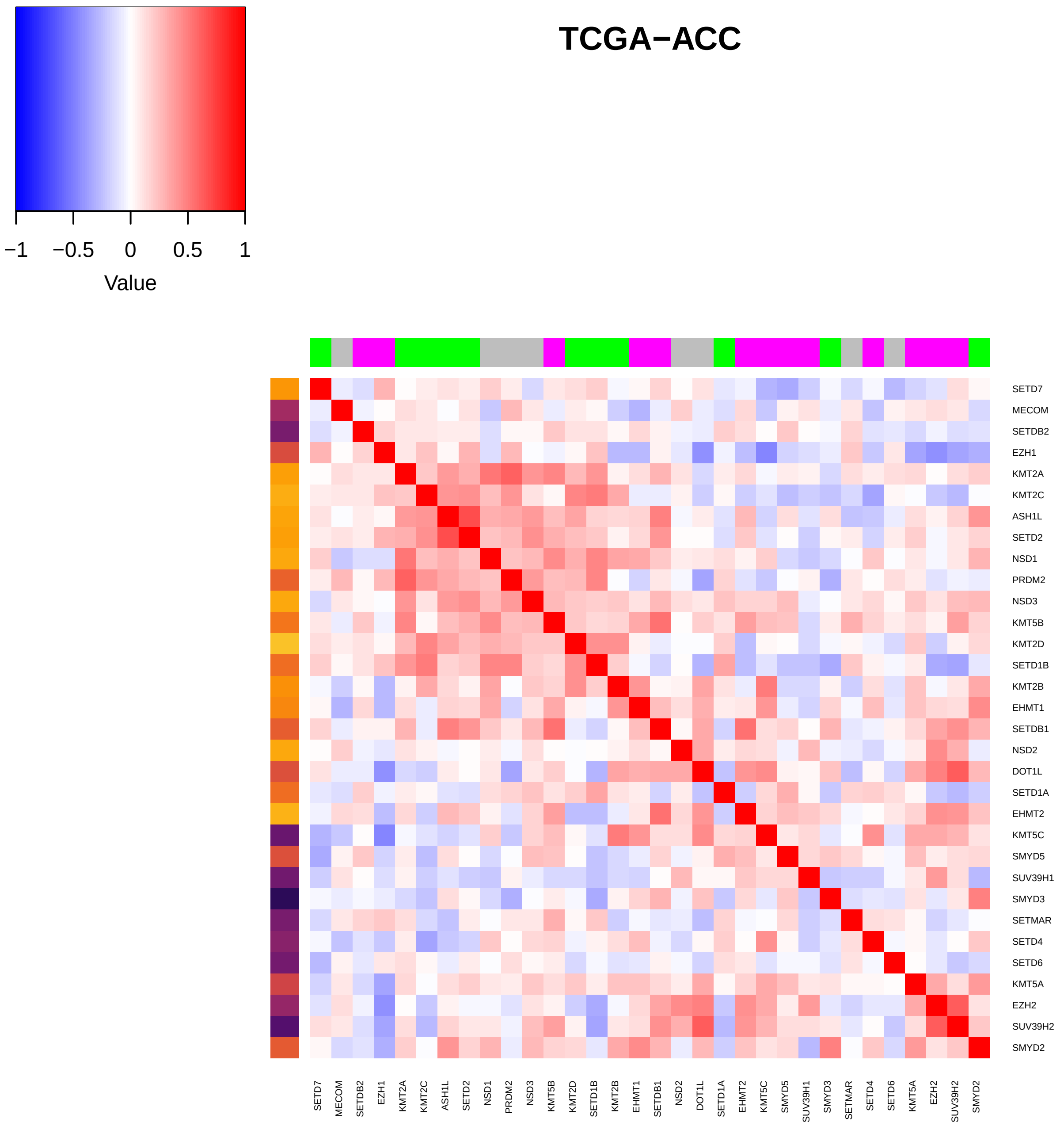
TCGA−ACC
Value
−1
−0.5
0
0.5
1
SETD7
MECOM
SETDB2
EZH1
KMT2A
KMT2C
ASH1L
SETD2
NSD1
PRDM2
NSD3
KMT5B
KMT2D
SETD1B
KMT2B
EHMT1
SETDB1
NSD2
DOT1L
SETD1A
EHMT2
KMT5C
SMYD5
SUV39H1
SMYD3
SETMAR
SETD4
SETD6
KMT5A
EZH2
SUV39H2
SMYD2

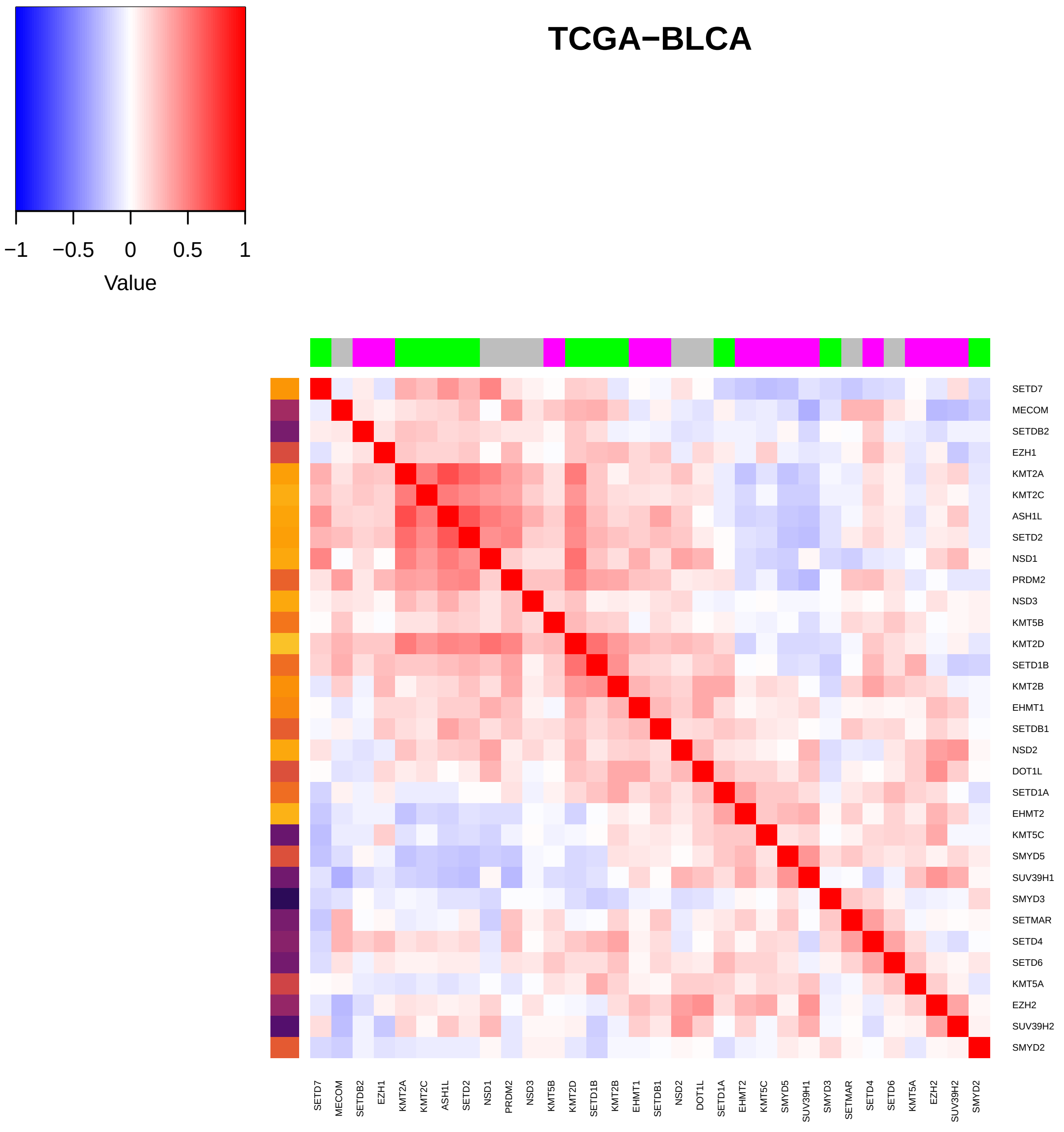
TCGA−BLCA
−1
−0.5
0
0.5
1
Value
SETD7
MECOM
SETDB2
EZH1
KMT2A
KMT2C
ASH1L
SETD2
NSD1
PRDM2
NSD3
KMT5B
KMT2D
SETD1B
KMT2B
EHMT1
SETDB1
NSD2
DOT1L
SETD1A
EHMT2
KMT5C
SMYD5
SUV39H1
SMYD3
SETMAR
SETD4
SETD6
KMT5A
EZH2
SUV39H2
SMYD2

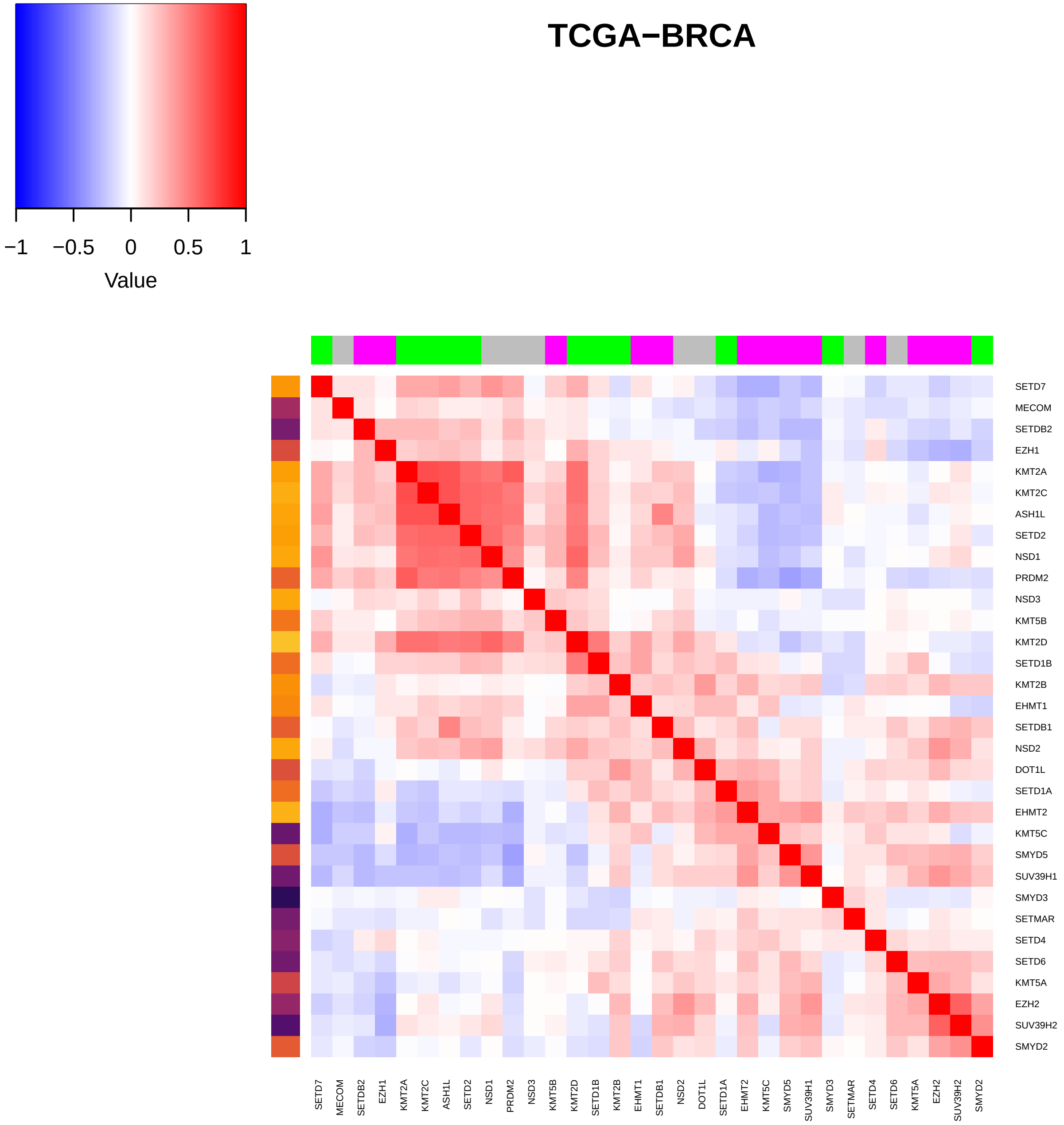
TCGA−BRCA
−1
−0.5
0
0.5
1
Value
SETD7
MECOM
SETDB2
EZH1
KMT2A
KMT2C
ASH1L
SETD2
NSD1
PRDM2
NSD3
KMT5B
KMT2D
SETD1B
KMT2B
EHMT1
SETDB1
NSD2
DOT1L
SETD1A
EHMT2
KMT5C
SMYD5
SUV39H1
SMYD3
SETMAR
SETD4
SETD6
KMT5A
EZH2
SUV39H2
SMYD2

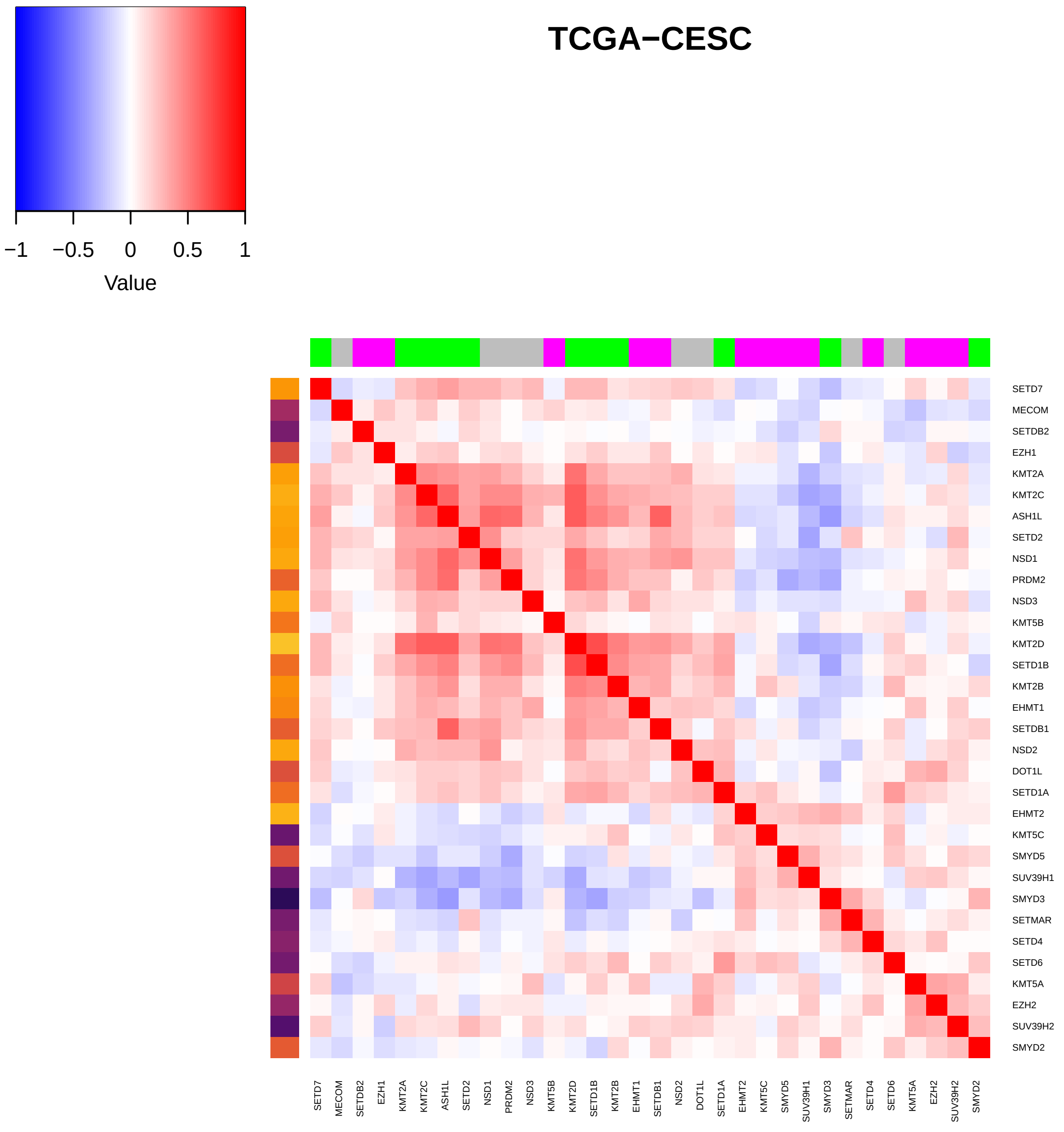
TCGA−CESC
−1
−0.5
0
0.5
1
Value
SETD7
MECOM
SETDB2
EZH1
KMT2A
KMT2C
ASH1L
SETD2
NSD1
PRDM2
NSD3
KMT5B
KMT2D
SETD1B
KMT2B
EHMT1
SETDB1
NSD2
DOT1L
SETD1A
EHMT2
KMT5C
SMYD5
SUV39H1
SMYD3
SETMAR
SETD4
SETD6
KMT5A
EZH2
SUV39H2
SMYD2

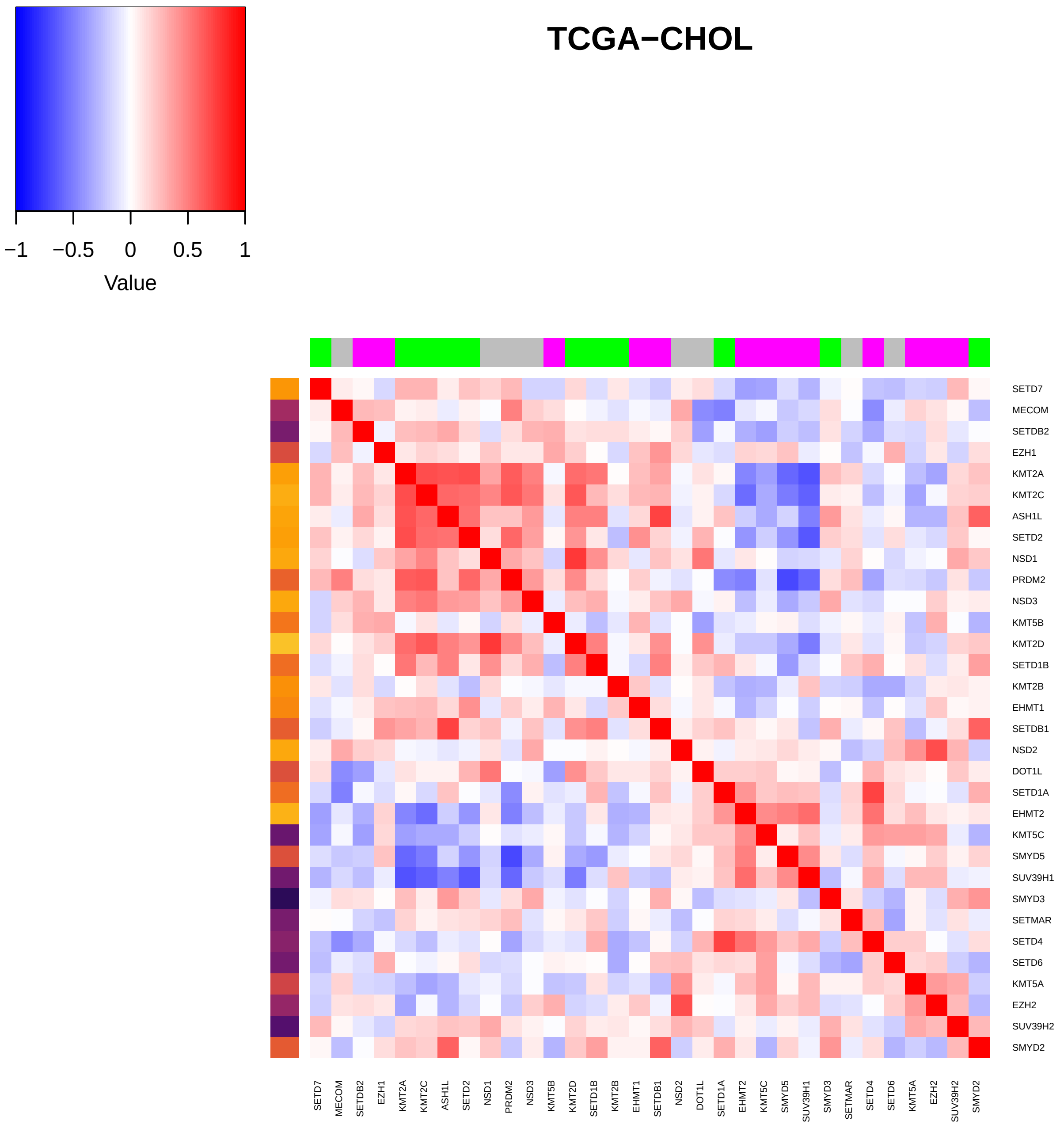

TCGA−CHOL
−1
−0.5
0
0.5
1
Value
SETD7
MECOM
SETDB2
EZH1
KMT2A
KMT2C
ASH1L
SETD2
NSD1
PRDM2
NSD3
KMT5B
KMT2D
SETD1B
KMT2B
EHMT1
SETDB1
NSD2
DOT1L
SETD1A
EHMT2
KMT5C
SMYD5
SUV39H1
SMYD3
SETMAR
SETD4
SETD6
KMT5A
EZH2
SUV39H2
SMYD2

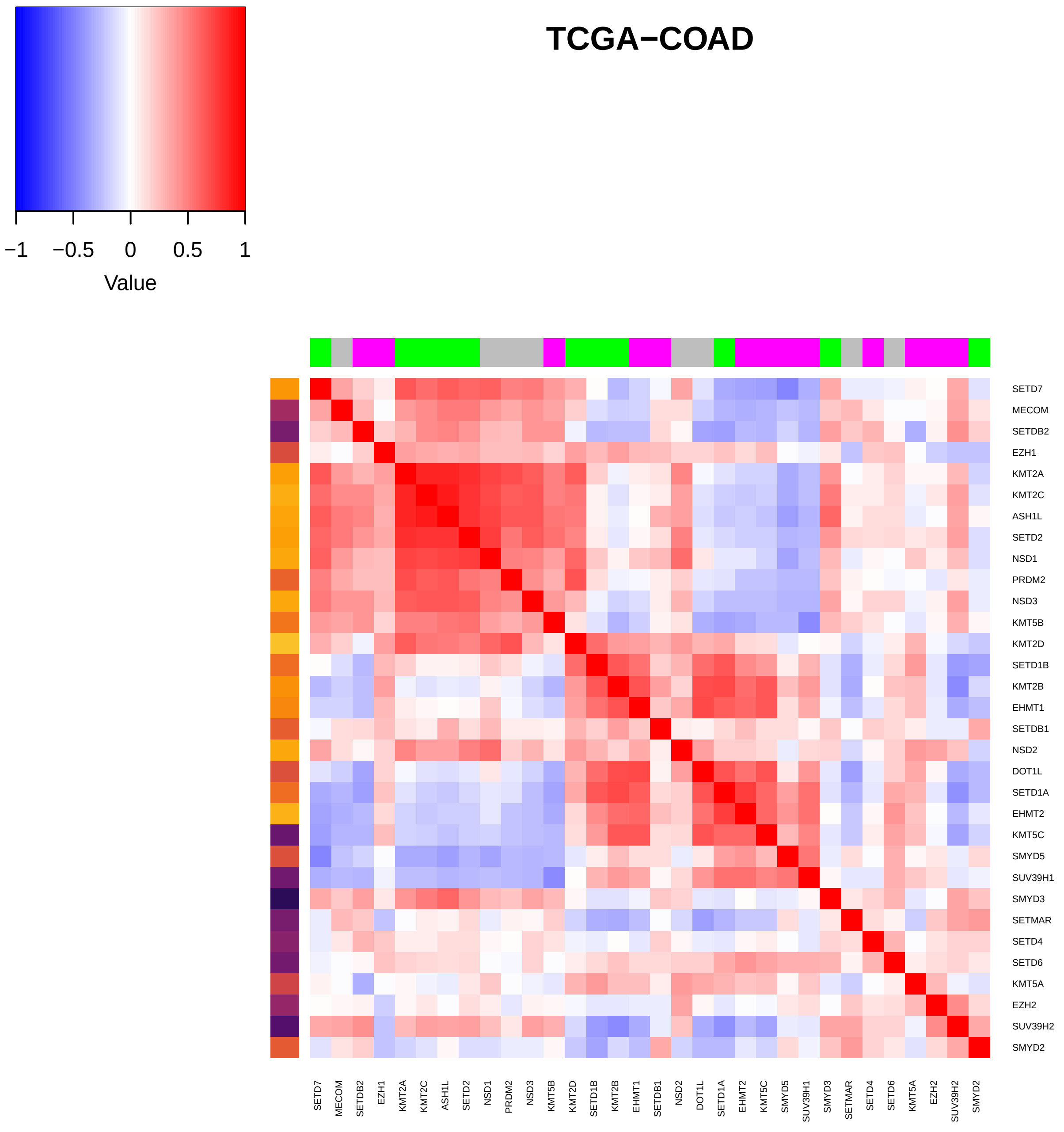
TCGA−COAD
Value
−1
−0.5
0
0.5
1
SETD7
MECOM
SETDB2
EZH1
KMT2A
KMT2C
ASH1L
SETD2
NSD1
PRDM2
NSD3
KMT5B
KMT2D
SETD1B
KMT2B
EHMT1
SETDB1
NSD2
DOT1L
SETD1A
EHMT2
KMT5C
SMYD5
SUV39H1
SMYD3
SETMAR
SETD4
SETD6
KMT5A
EZH2
SUV39H2
SMYD2

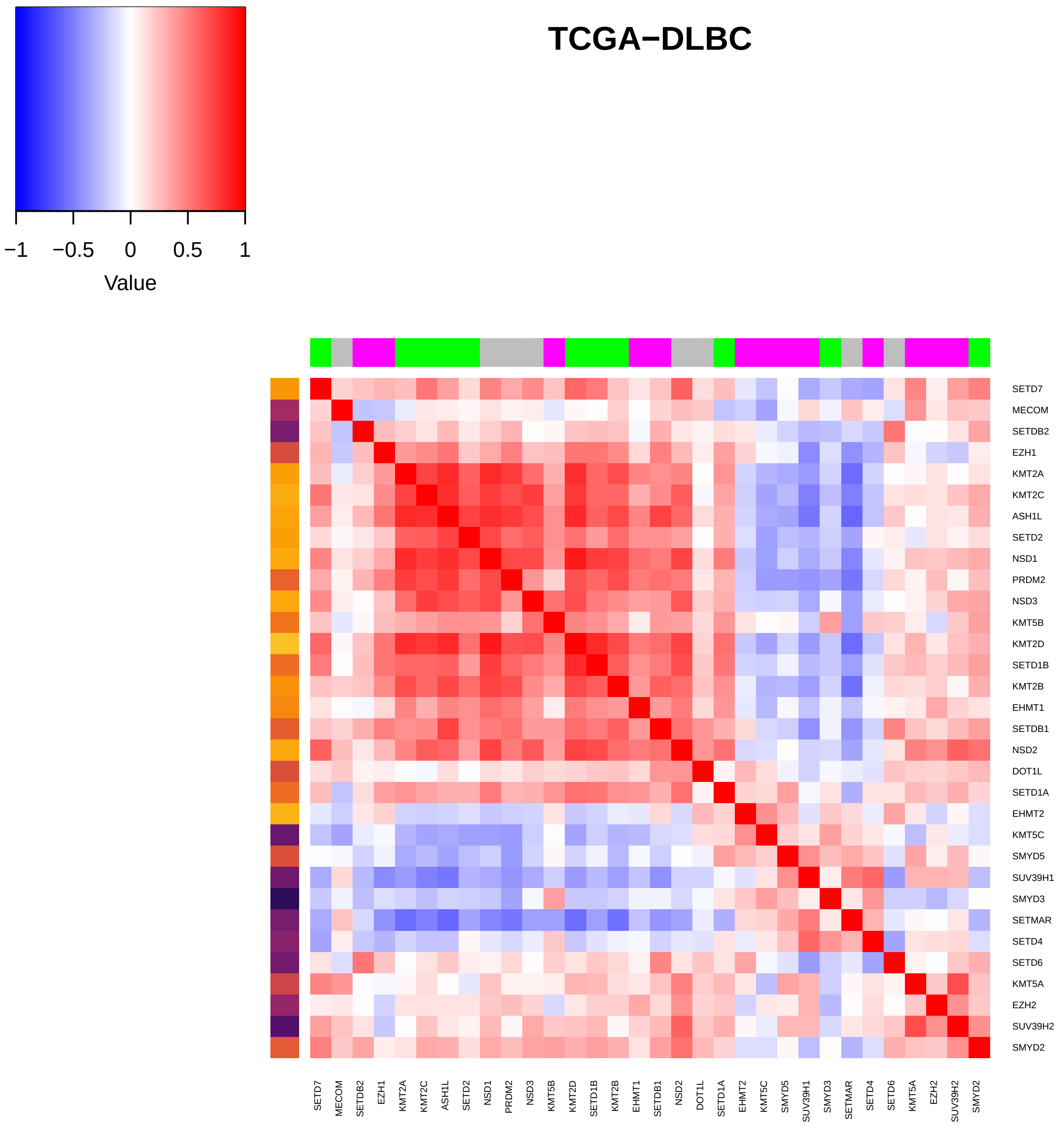
TCGA−DLBC
−1
−0.5
0
0.5
1
Value
SETD7
MECOM
SETDB2
EZH1
KMT2A
KMT2C
ASH1L
SETD2
NSD1
PRDM2
NSD3
KMT5B
KMT2D
SETD1B
KMT2B
EHMT1
SETDB1
NSD2
DOT1L
SETD1A
EHMT2
KMT5C
SMYD5
SUV39H1
SMYD3
SETMAR
SETD4
SETD6
KMT5A
EZH2
SUV39H2
SMYD2

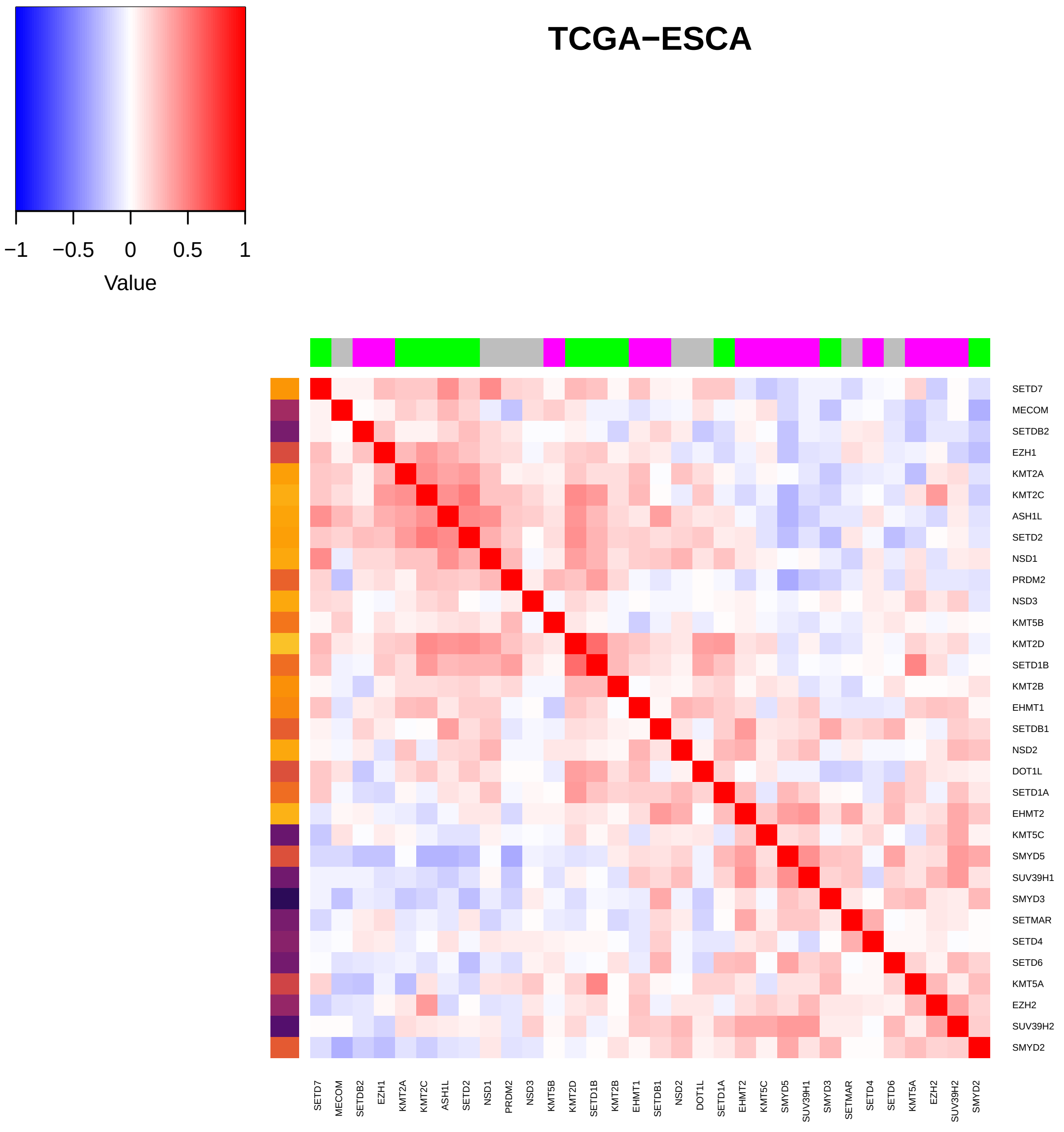

TCGA−ESCA
−1
−0.5
0
0.5
1
Value
SETD7
MECOM
SETDB2
EZH1
KMT2A
KMT2C
ASH1L
SETD2
NSD1
PRDM2
NSD3
KMT5B
KMT2D
SETD1B
KMT2B
EHMT1
SETDB1
NSD2
DOT1L
SETD1A
EHMT2
KMT5C
SMYD5
SUV39H1
SMYD3
SETMAR
SETD4
SETD6
KMT5A
EZH2
SUV39H2
SMYD2

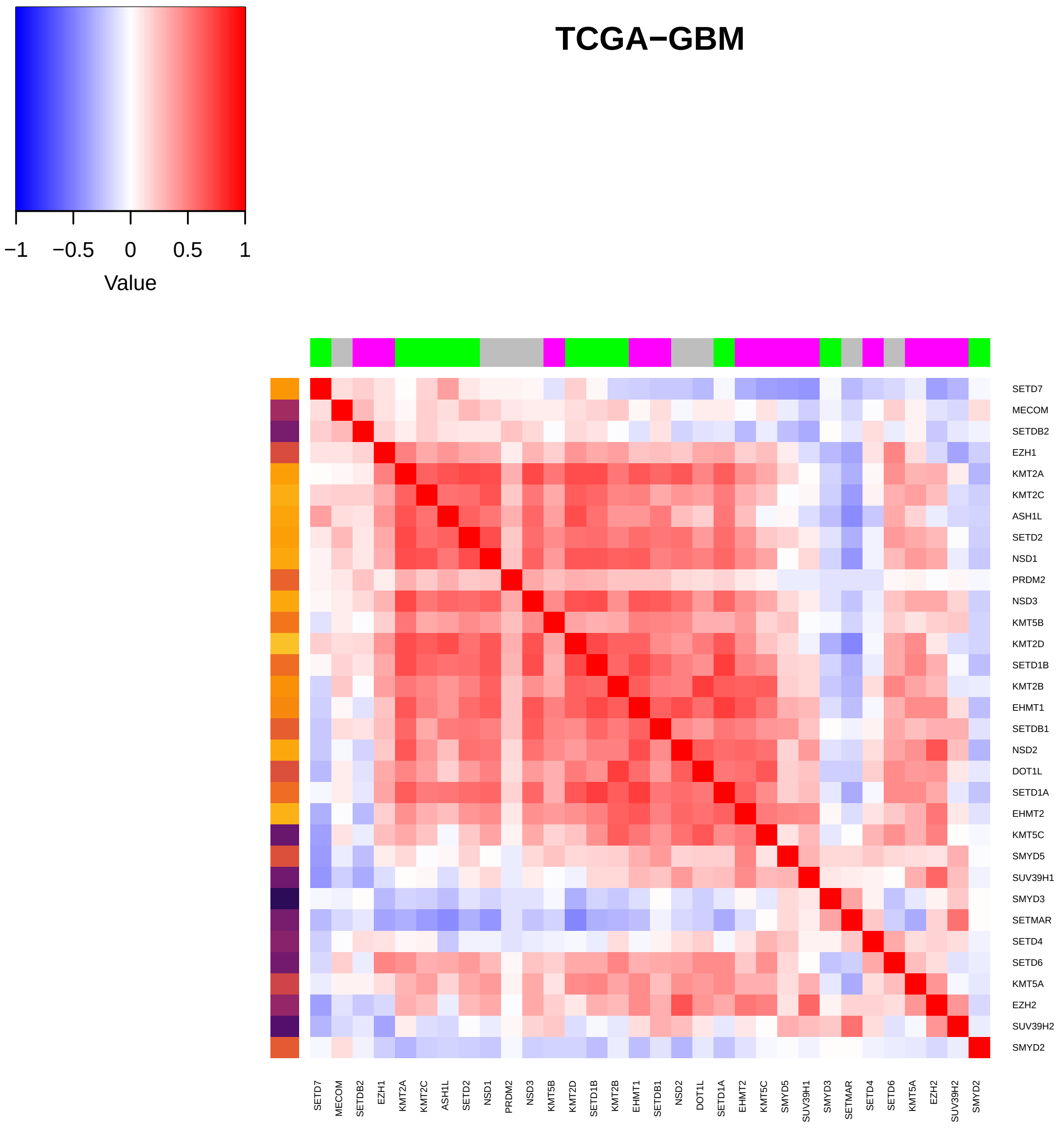

TCGA−GBM
Value
−1
−0.5
0
0.5
1
SETD7
MECOM
SETDB2
EZH1
KMT2A
KMT2C
ASH1L
SETD2
NSD1
PRDM2
NSD3
KMT5B
KMT2D
SETD1B
KMT2B
EHMT1
SETDB1
NSD2
DOT1L
SETD1A
EHMT2
KMT5C
SMYD5
SUV39H1
SMYD3
SETMAR
SETD4
SETD6
KMT5A
EZH2
SUV39H2
SMYD2

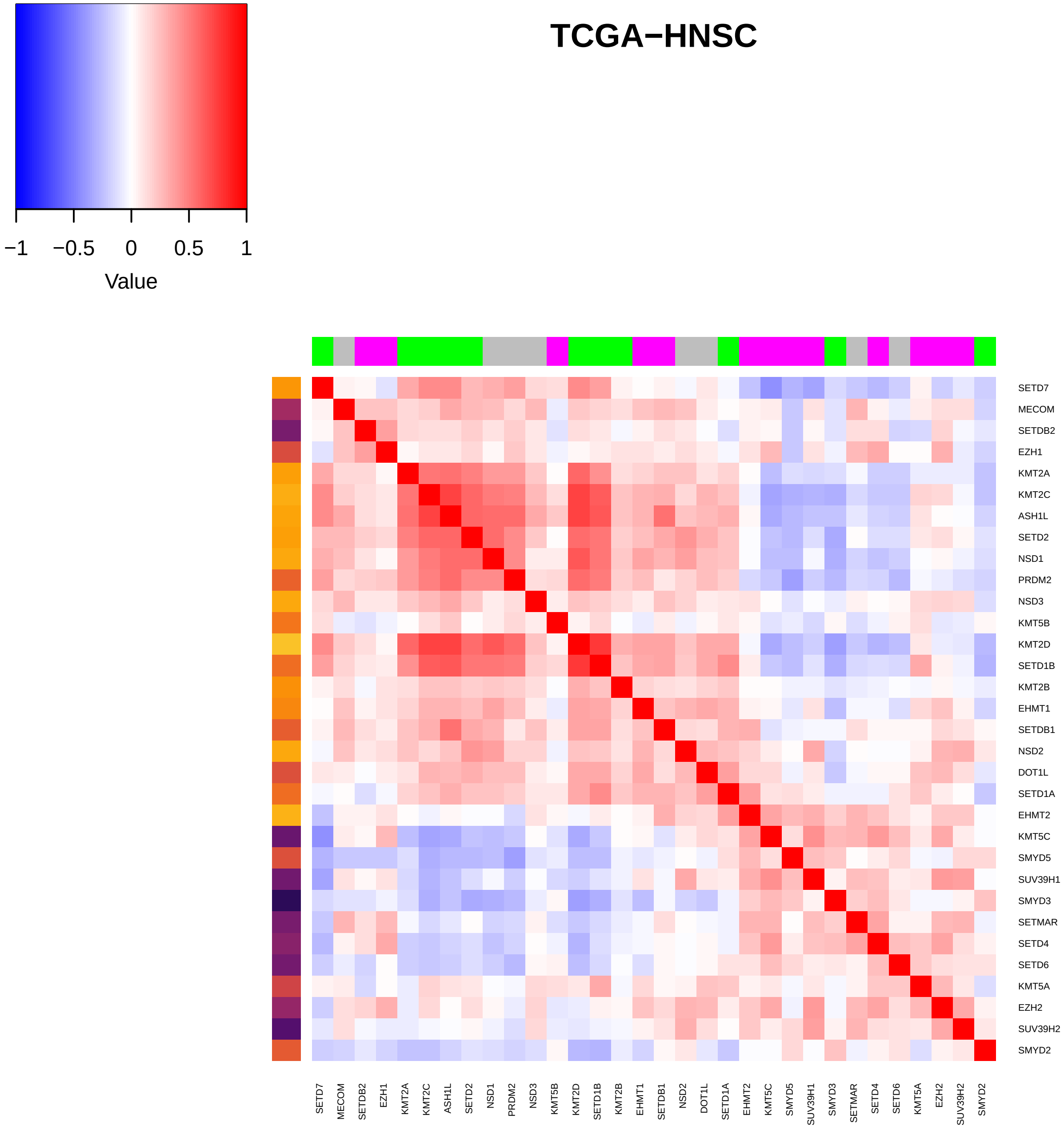

TCGA−HNSC
−1
−0.5
0
0.5
1
Value
SETD7
MECOM
SETDB2
EZH1
KMT2A
KMT2C
ASH1L
SETD2
NSD1
PRDM2
NSD3
KMT5B
KMT2D
SETD1B
KMT2B
EHMT1
SETDB1
NSD2
DOT1L
SETD1A
EHMT2
KMT5C
SMYD5
SUV39H1
SMYD3
SETMAR
SETD4
SETD6
KMT5A
EZH2
SUV39H2
SMYD2

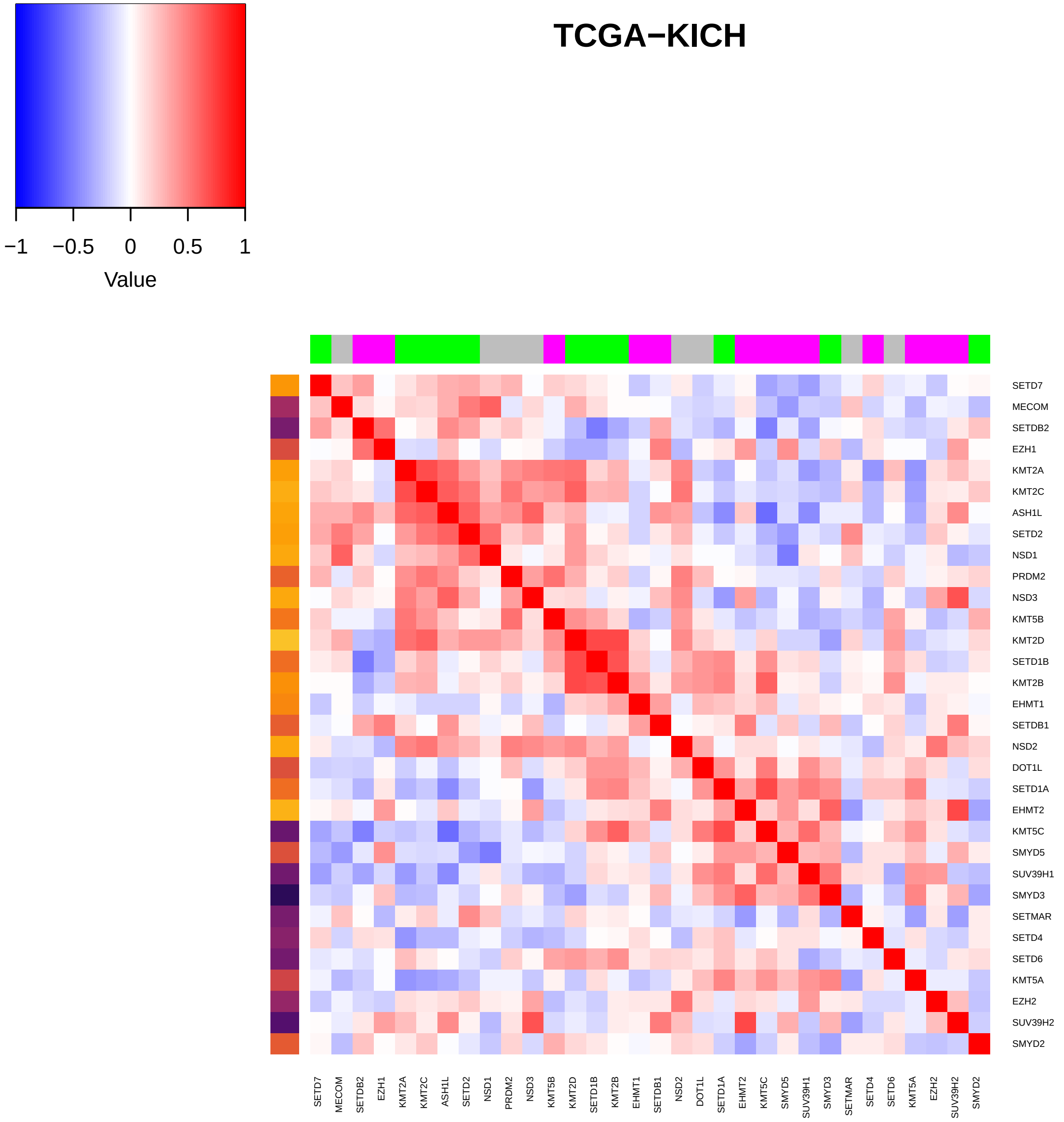
TCGA−KICH
−1
−0.5
0
0.5
1
Value
SETD7
MECOM
SETDB2
EZH1
KMT2A
KMT2C
ASH1L
SETD2
NSD1
PRDM2
NSD3
KMT5B
KMT2D
SETD1B
KMT2B
EHMT1
SETDB1
NSD2
DOT1L
SETD1A
EHMT2
KMT5C
SMYD5
SUV39H1
SMYD3
SETMAR
SETD4
SETD6
KMT5A
EZH2
SUV39H2
SMYD2

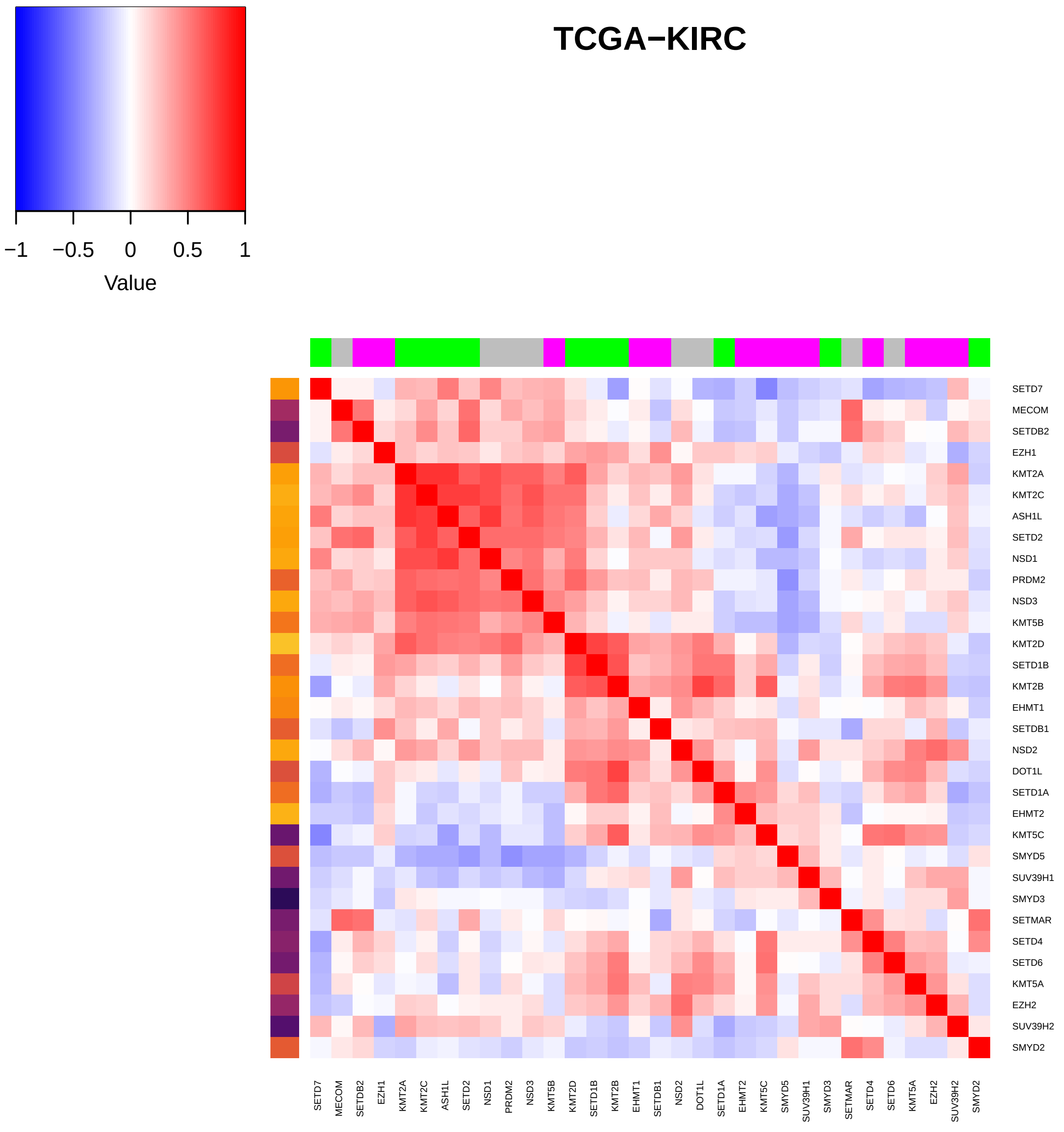
TCGA−KIRC
−1
−0.5
0
0.5
1
Value
SETD7
MECOM
SETDB2
EZH1
KMT2A
KMT2C
ASH1L
SETD2
NSD1
PRDM2
NSD3
KMT5B
KMT2D
SETD1B
KMT2B
EHMT1
SETDB1
NSD2
DOT1L
SETD1A
EHMT2
KMT5C
SMYD5
SUV39H1
SMYD3
SETMAR
SETD4
SETD6
KMT5A
EZH2
SUV39H2
SMYD2

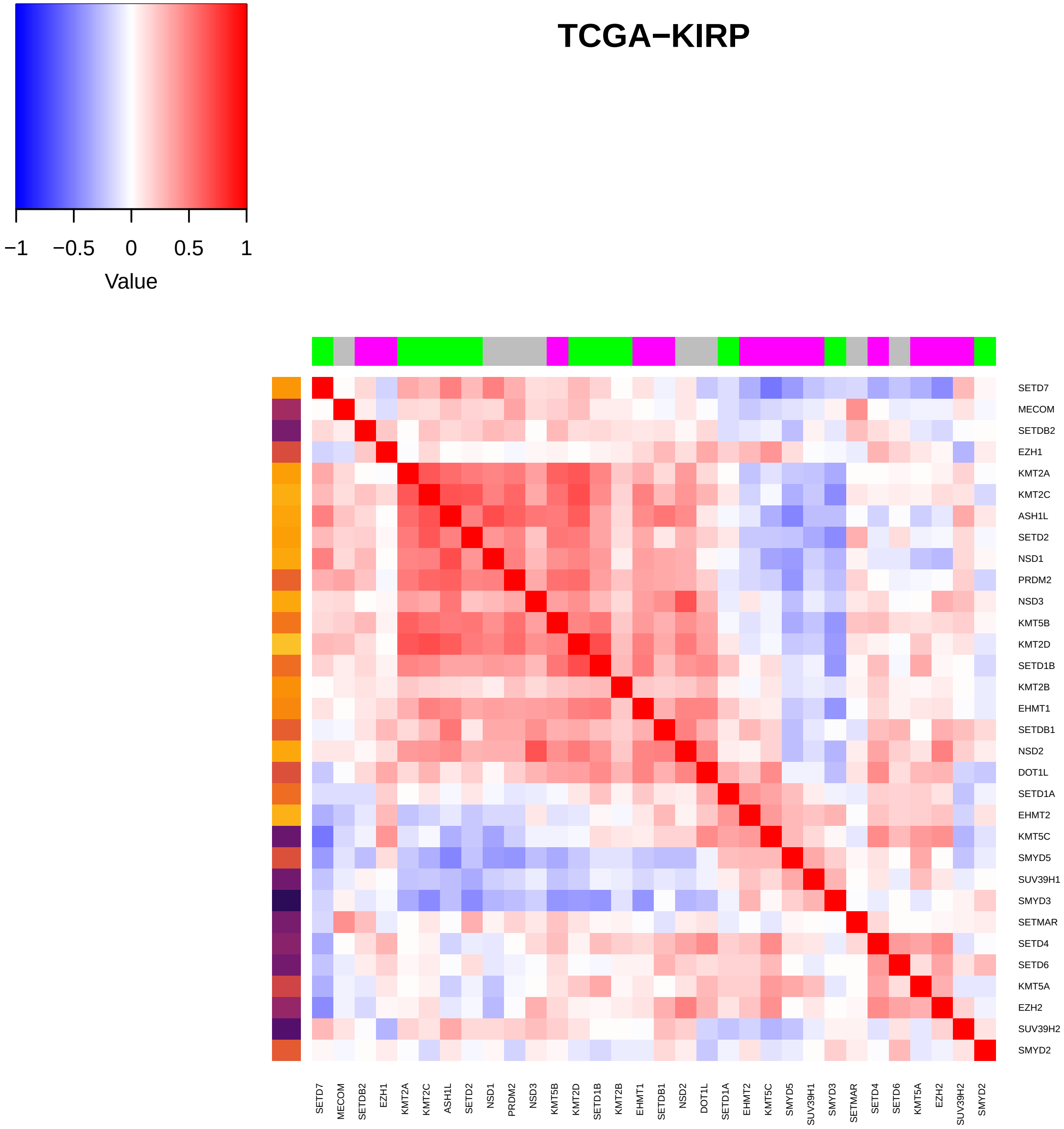

TCGA−KIRP
−1
−0.5
0
0.5
1
Value
SETD7
MECOM
SETDB2
EZH1
KMT2A
KMT2C
ASH1L
SETD2
NSD1
PRDM2
NSD3
KMT5B
KMT2D
SETD1B
KMT2B
EHMT1
SETDB1
NSD2
DOT1L
SETD1A
EHMT2
KMT5C
SMYD5
SUV39H1
SMYD3
SETMAR
SETD4
SETD6
KMT5A
EZH2
SUV39H2
SMYD2

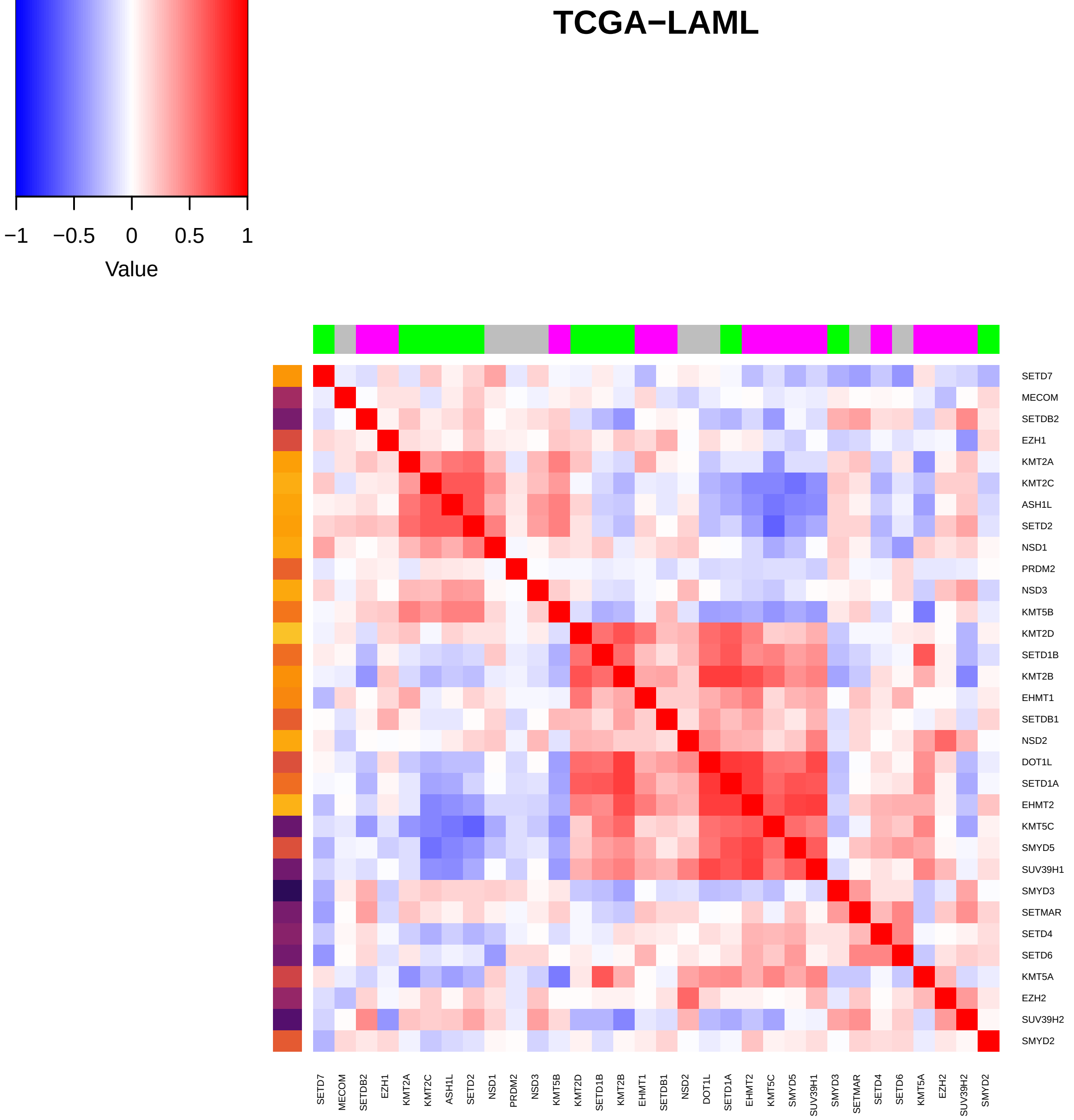
TCGA−LAML
−1
−0.5
0
0.5
1
Value
SETD7
MECOM
SETDB2
EZH1
KMT2A
KMT2C
ASH1L
SETD2
NSD1
PRDM2
NSD3
KMT5B
KMT2D
SETD1B
KMT2B
EHMT1
SETDB1
NSD2
DOT1L
SETD1A
EHMT2
KMT5C
SMYD5
SUV39H1
SMYD3
SETMAR
SETD4
SETD6
KMT5A
EZH2
SUV39H2
SMYD2

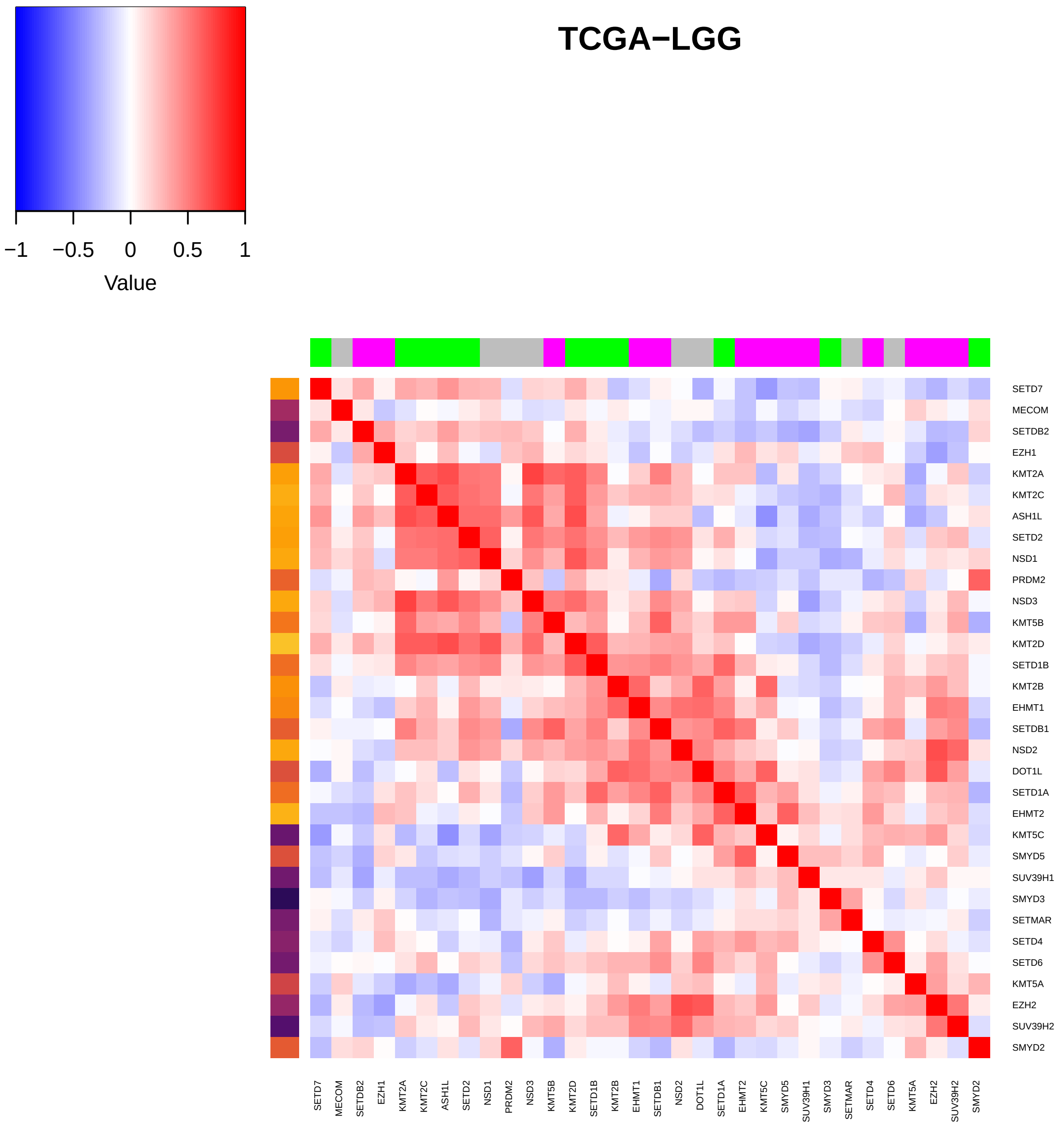

TCGA−LGG
−1
−0.5
0
0.5
1
Value
SETD7
MECOM
SETDB2
EZH1
KMT2A
KMT2C
ASH1L
SETD2
NSD1
PRDM2
NSD3
KMT5B
KMT2D
SETD1B
KMT2B
EHMT1
SETDB1
NSD2
DOT1L
SETD1A
EHMT2
KMT5C
SMYD5
SUV39H1
SMYD3
SETMAR
SETD4
SETD6
KMT5A
EZH2
SUV39H2
SMYD2

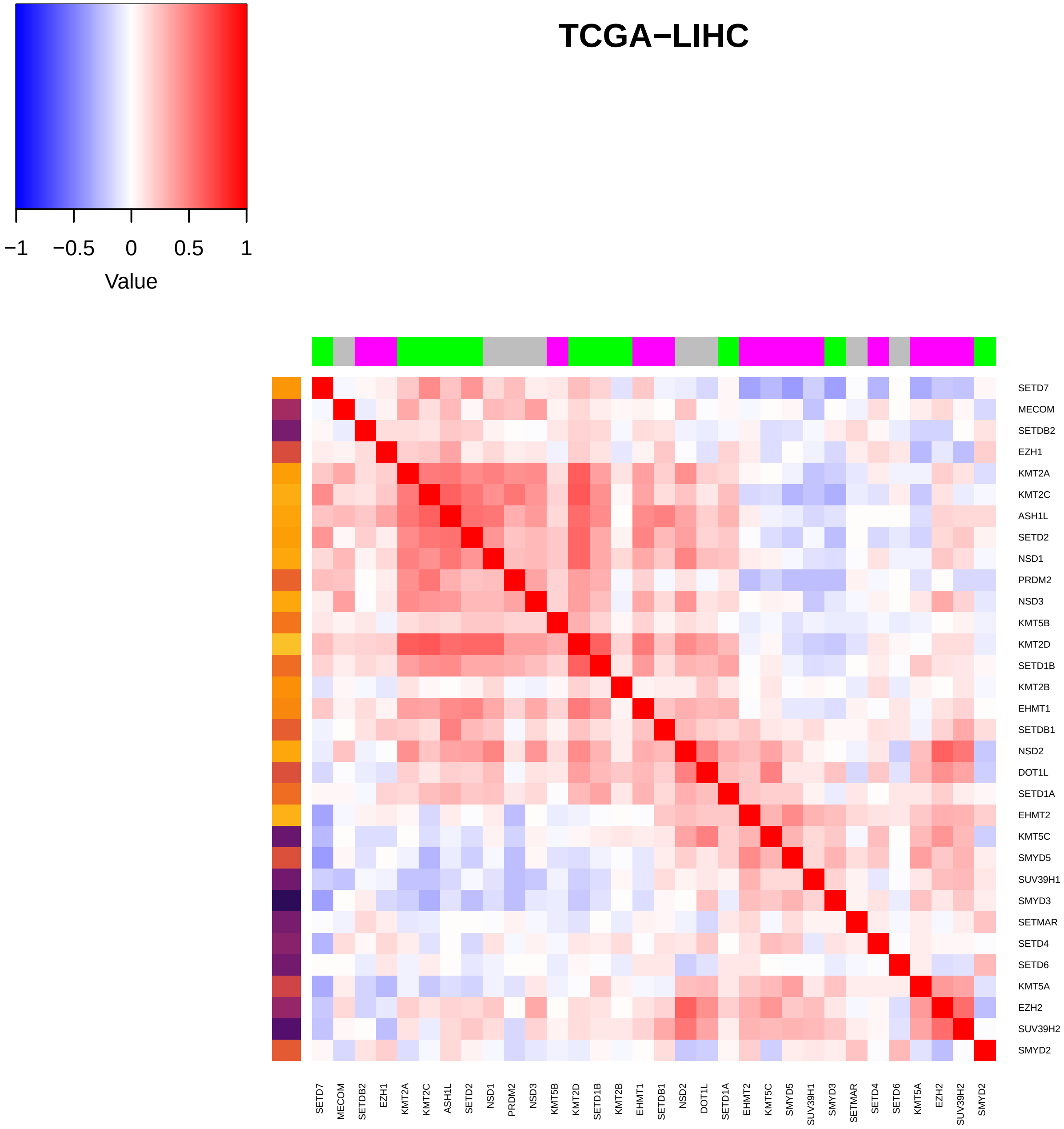

TCGA−LIHC
−1
−0.5
0
0.5
1
Value
SETD7
MECOM
SETDB2
EZH1
KMT2A
KMT2C
ASH1L
SETD2
NSD1
PRDM2
NSD3
KMT5B
KMT2D
SETD1B
KMT2B
EHMT1
SETDB1
NSD2
DOT1L
SETD1A
EHMT2
KMT5C
SMYD5
SUV39H1
SMYD3
SETMAR
SETD4
SETD6
KMT5A
EZH2
SUV39H2
SMYD2

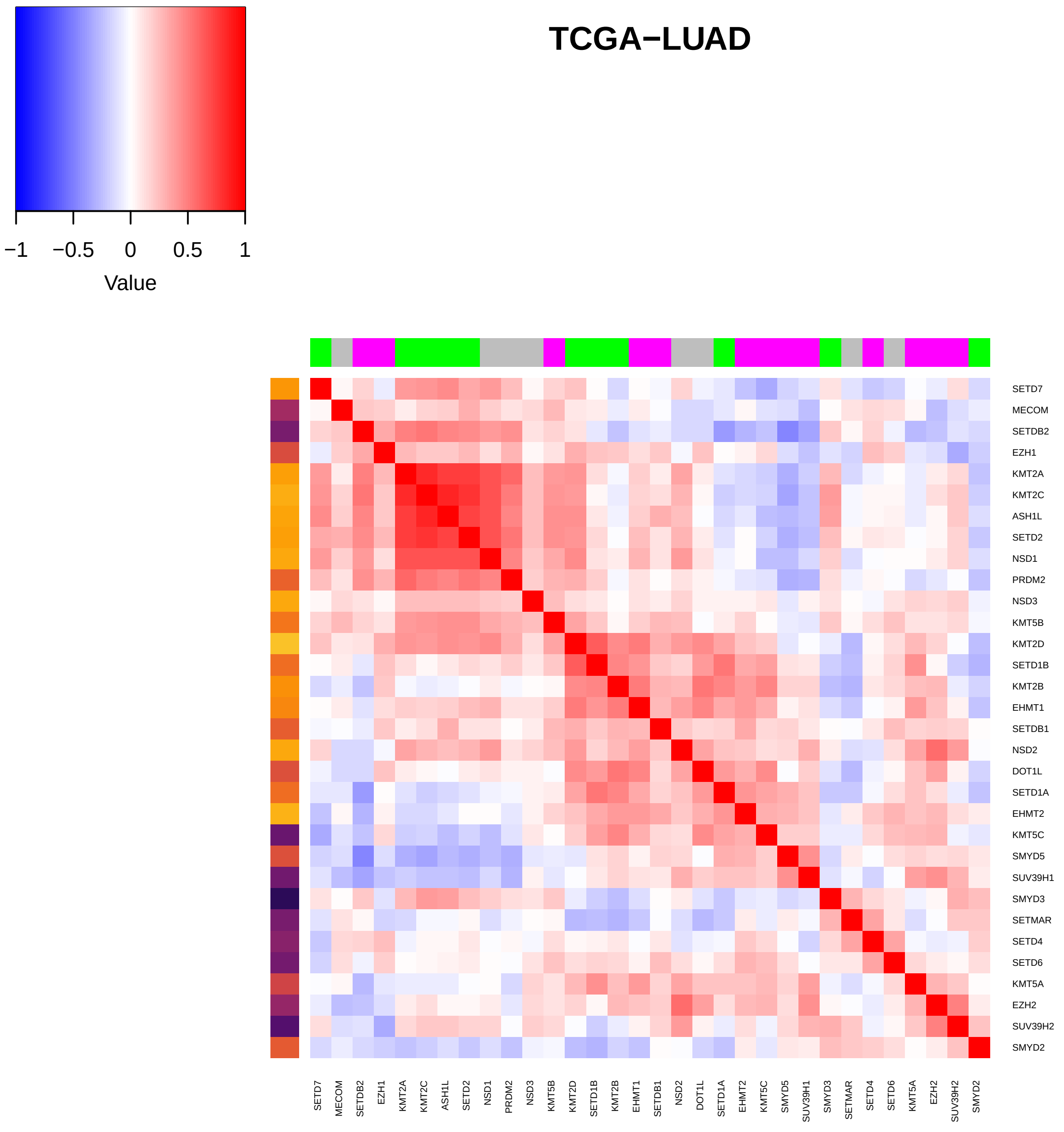

TCGA−LUAD
−1
−0.5
0
0.5
1
Value
SETD7
MECOM
SETDB2
EZH1
KMT2A
KMT2C
ASH1L
SETD2
NSD1
PRDM2
NSD3
KMT5B
KMT2D
SETD1B
KMT2B
EHMT1
SETDB1
NSD2
DOT1L
SETD1A
EHMT2
KMT5C
SMYD5
SUV39H1
SMYD3
SETMAR
SETD4
SETD6
KMT5A
EZH2
SUV39H2
SMYD2

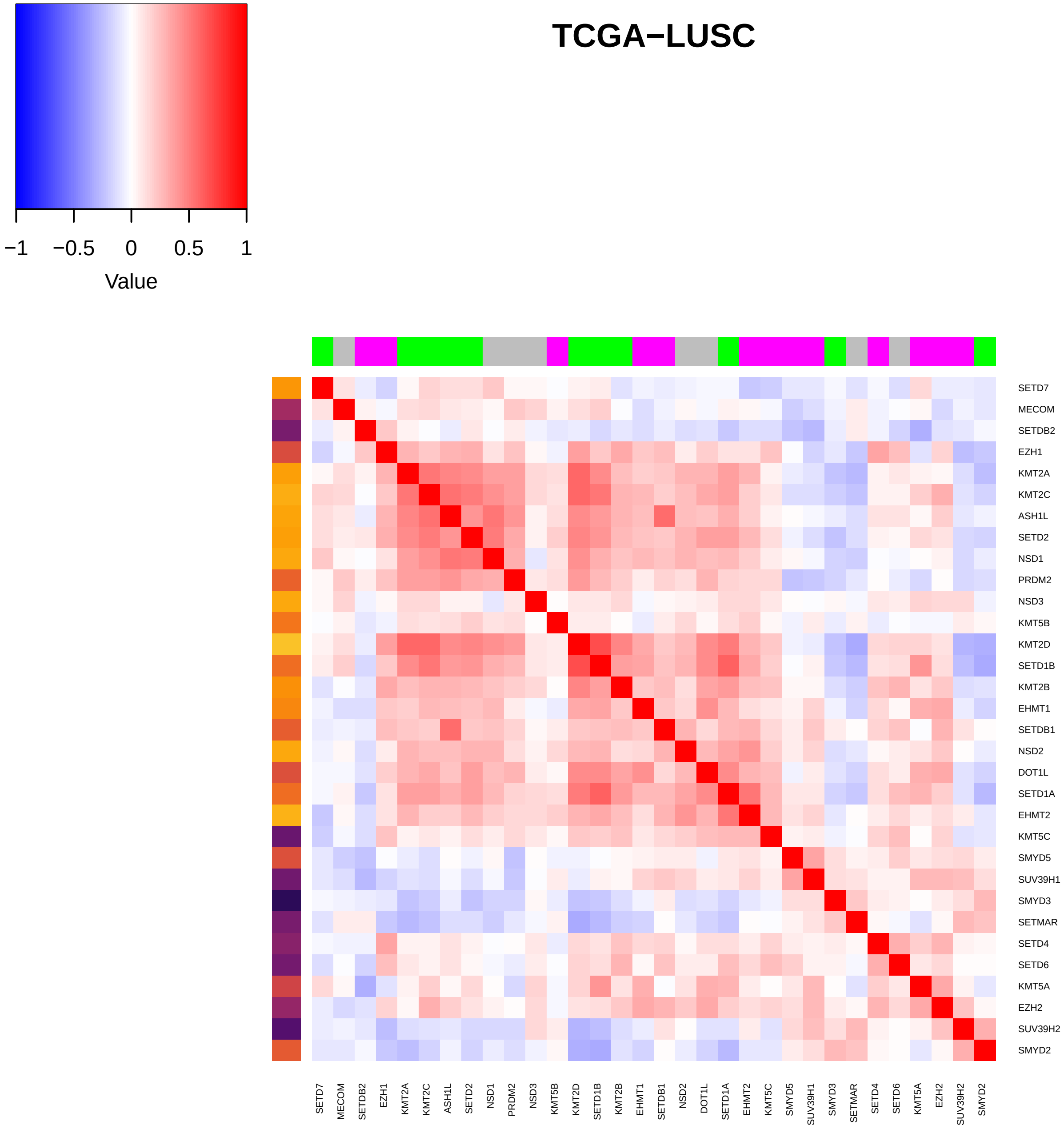

TCGA−LUSC
−1
−0.5
0
0.5
1
Value
SETD7
MECOM
SETDB2
EZH1
KMT2A
KMT2C
ASH1L
SETD2
NSD1
PRDM2
NSD3
KMT5B
KMT2D
SETD1B
KMT2B
EHMT1
SETDB1
NSD2
DOT1L
SETD1A
EHMT2
KMT5C
SMYD5
SUV39H1
SMYD3
SETMAR
SETD4
SETD6
KMT5A
EZH2
SUV39H2
SMYD2

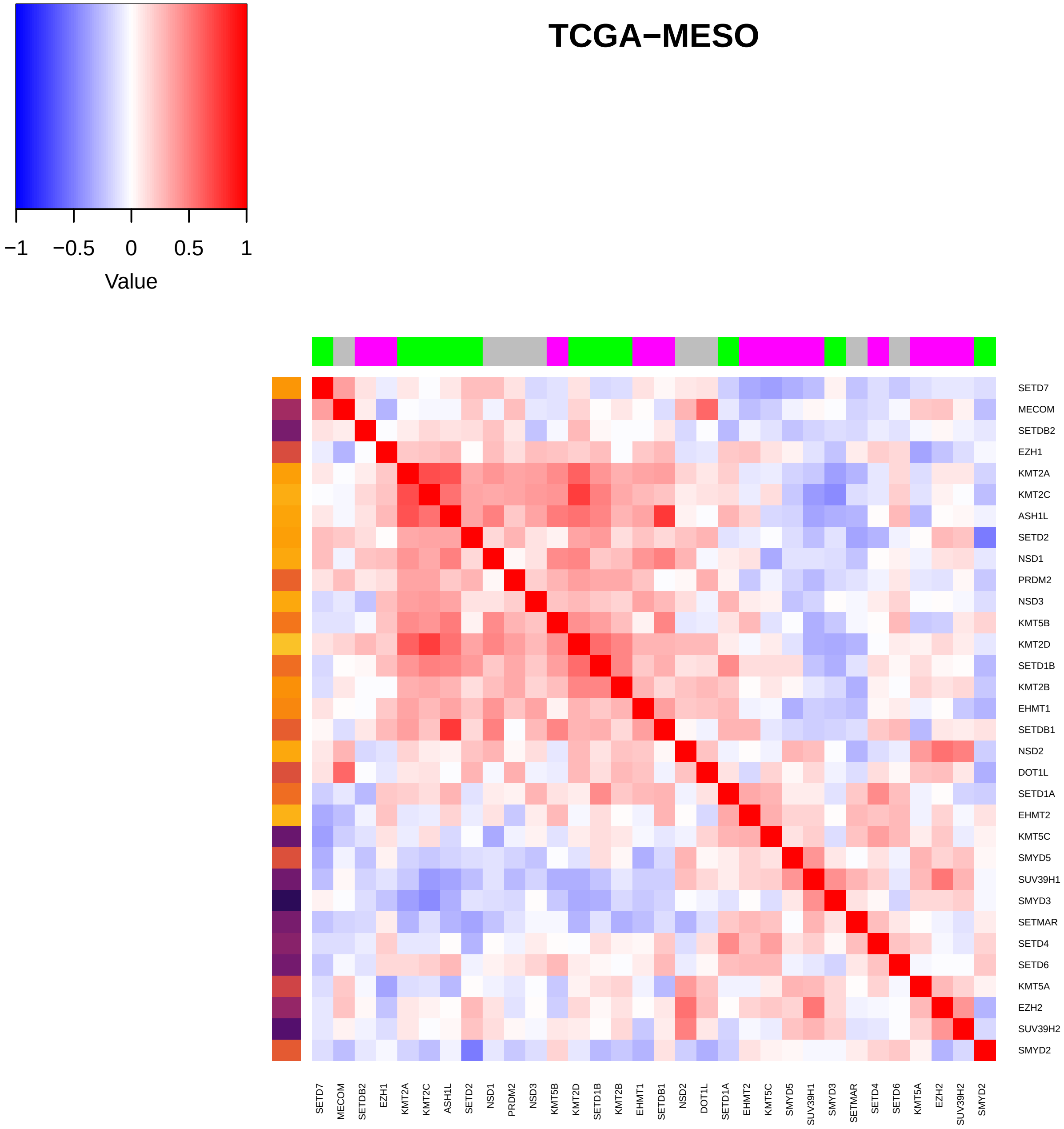

TCGA−MESO
−1
−0.5
0
0.5
1
Value
SETD7
MECOM
SETDB2
EZH1
KMT2A
KMT2C
ASH1L
SETD2
NSD1
PRDM2
NSD3
KMT5B
KMT2D
SETD1B
KMT2B
EHMT1
SETDB1
NSD2
DOT1L
SETD1A
EHMT2
KMT5C
SMYD5
SUV39H1
SMYD3
SETMAR
SETD4
SETD6
KMT5A
EZH2
SUV39H2
SMYD2

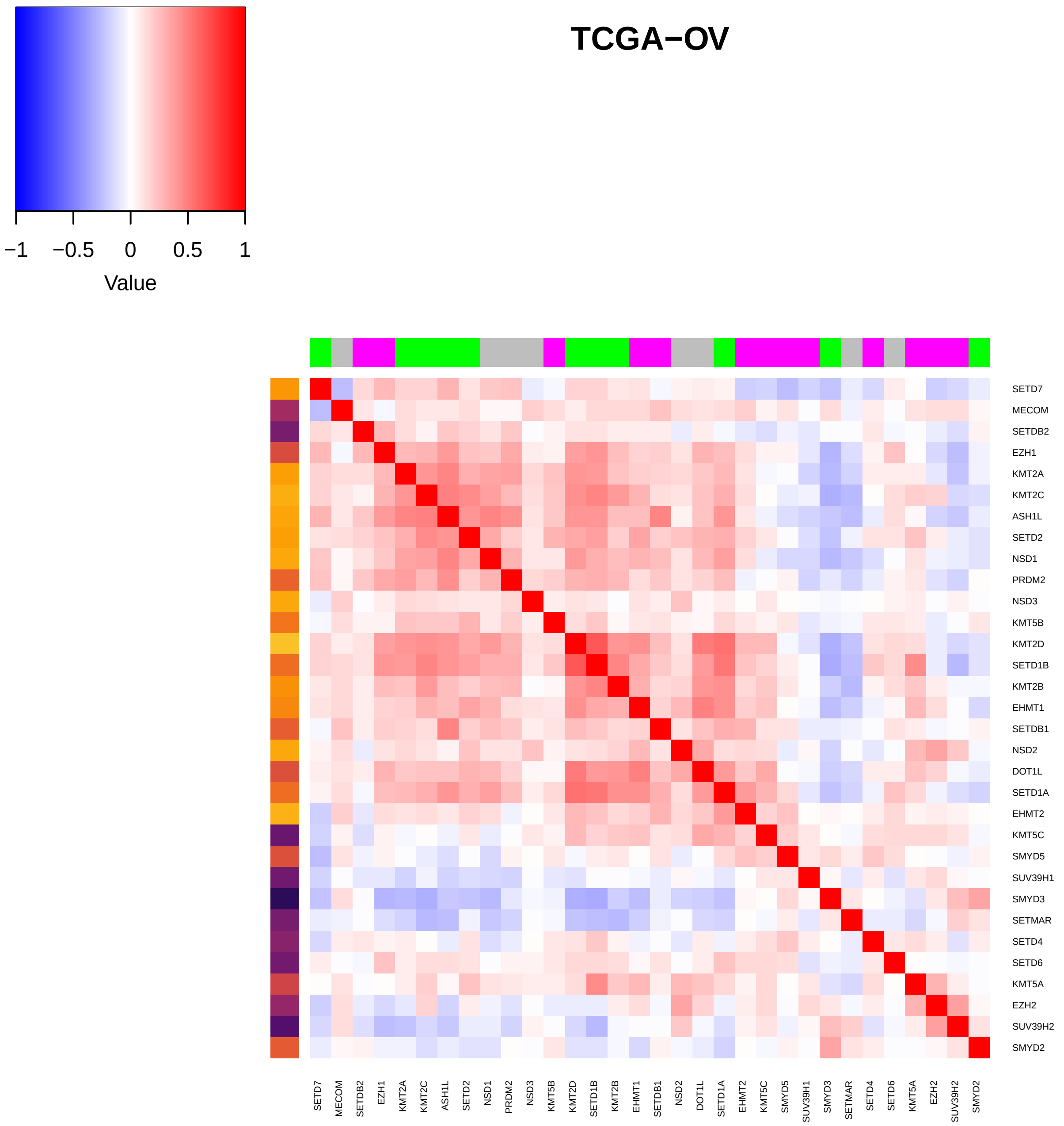

TCGA−OV
−1
−0.5
0
0.5
1
Value
SETD7
MECOM
SETDB2
EZH1
KMT2A
KMT2C
ASH1L
SETD2
NSD1
PRDM2
NSD3
KMT5B
KMT2D
SETD1B
KMT2B
EHMT1
SETDB1
NSD2
DOT1L
SETD1A
EHMT2
KMT5C
SMYD5
SUV39H1
SMYD3
SETMAR
SETD4
SETD6
KMT5A
EZH2
SUV39H2
SMYD2

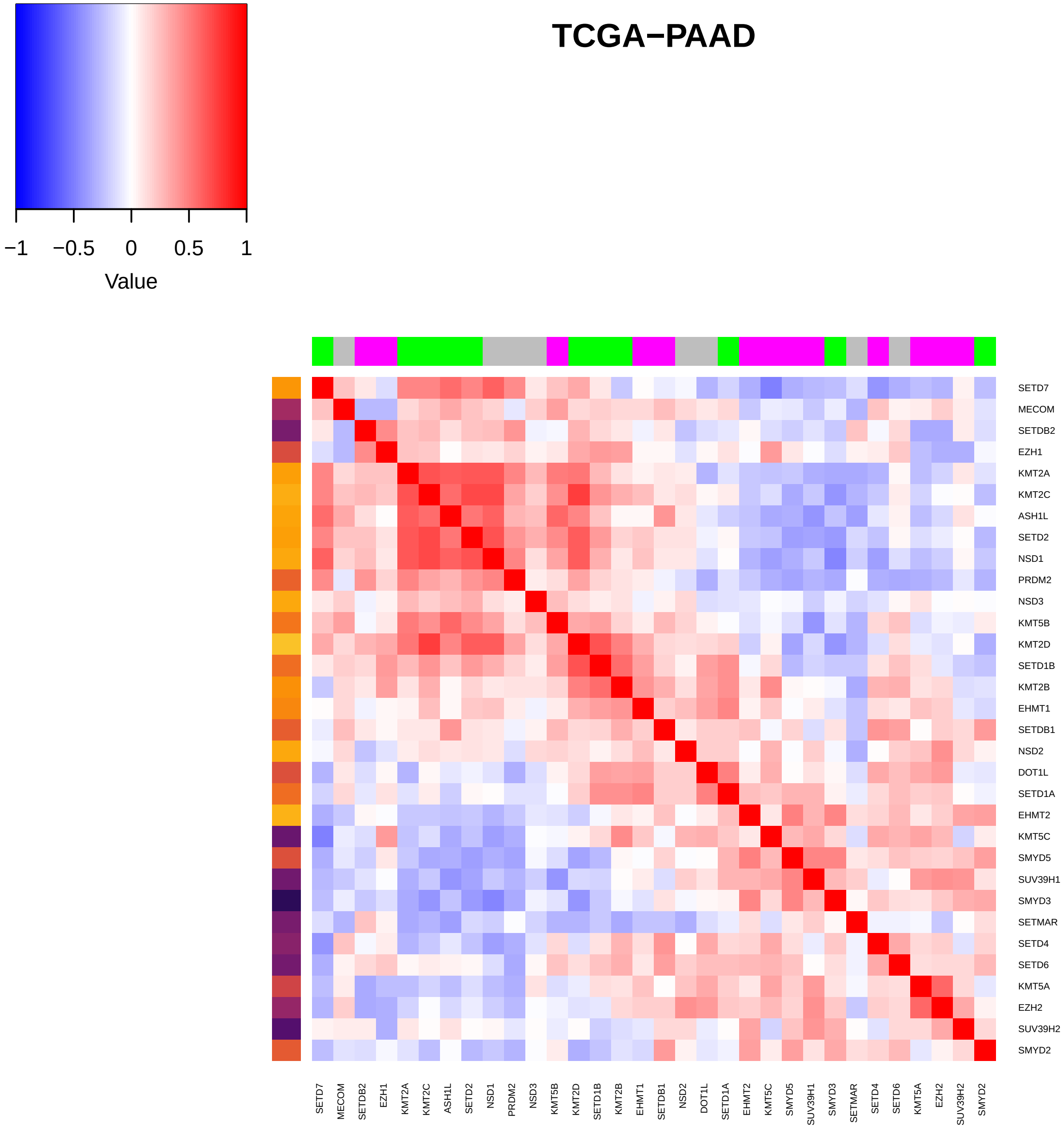

TCGA−PAAD
−1
−0.5
0
0.5
1
Value
SETD7
MECOM
SETDB2
EZH1
KMT2A
KMT2C
ASH1L
SETD2
NSD1
PRDM2
NSD3
KMT5B
KMT2D
SETD1B
KMT2B
EHMT1
SETDB1
NSD2
DOT1L
SETD1A
EHMT2
KMT5C
SMYD5
SUV39H1
SMYD3
SETMAR
SETD4
SETD6
KMT5A
EZH2
SUV39H2
SMYD2

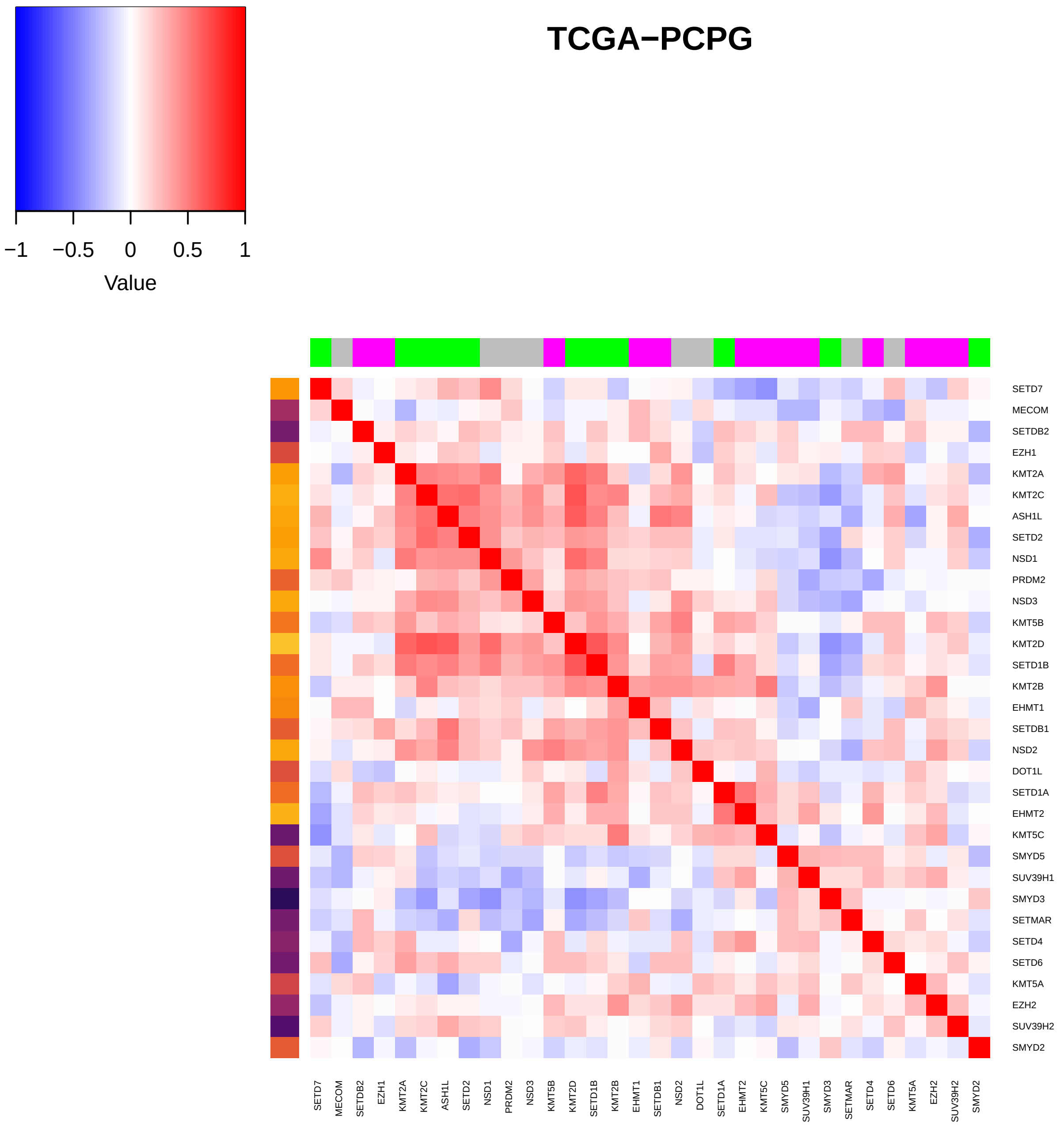
TCGA−PCPG
−1
−0.5
0
0.5
1
Value
SETD7
MECOM
SETDB2
EZH1
KMT2A
KMT2C
ASH1L
SETD2
NSD1
PRDM2
NSD3
KMT5B
KMT2D
SETD1B
KMT2B
EHMT1
SETDB1
NSD2
DOT1L
SETD1A
EHMT2
KMT5C
SMYD5
SUV39H1
SMYD3
SETMAR
SETD4
SETD6
KMT5A
EZH2
SUV39H2
SMYD2

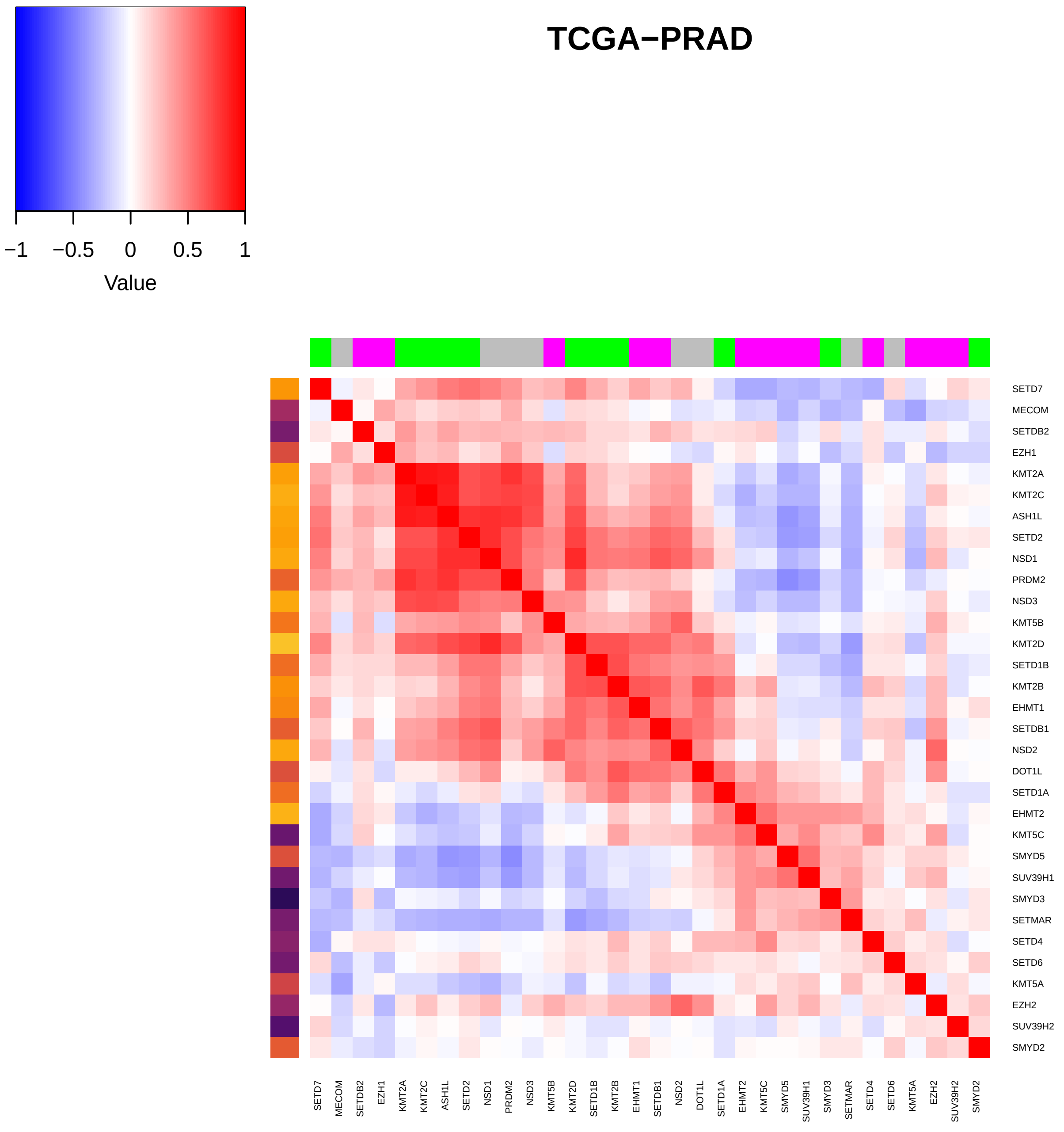

TCGA−PRAD
−1
−0.5
0
0.5
1
Value
SETD7
MECOM
SETDB2
EZH1
KMT2A
KMT2C
ASH1L
SETD2
NSD1
PRDM2
NSD3
KMT5B
KMT2D
SETD1B
KMT2B
EHMT1
SETDB1
NSD2
DOT1L
SETD1A
EHMT2
KMT5C
SMYD5
SUV39H1
SMYD3
SETMAR
SETD4
SETD6
KMT5A
EZH2
SUV39H2
SMYD2

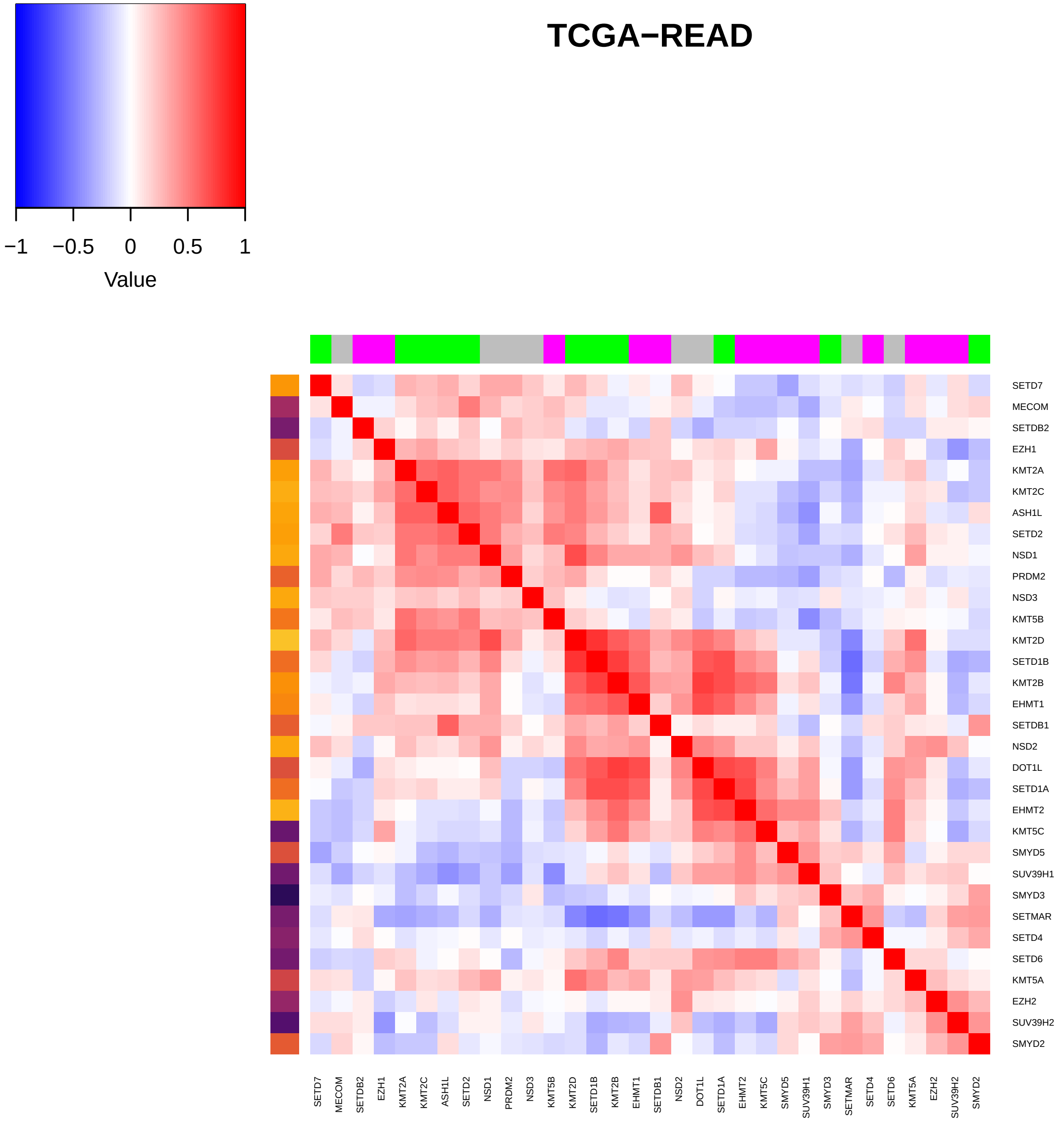

TCGA−READ
−1
−0.5
0
0.5
1
Value
SETD7
MECOM
SETDB2
EZH1
KMT2A
KMT2C
ASH1L
SETD2
NSD1
PRDM2
NSD3
KMT5B
KMT2D
SETD1B
KMT2B
EHMT1
SETDB1
NSD2
DOT1L
SETD1A
EHMT2
KMT5C
SMYD5
SUV39H1
SMYD3
SETMAR
SETD4
SETD6
KMT5A
EZH2
SUV39H2
SMYD2

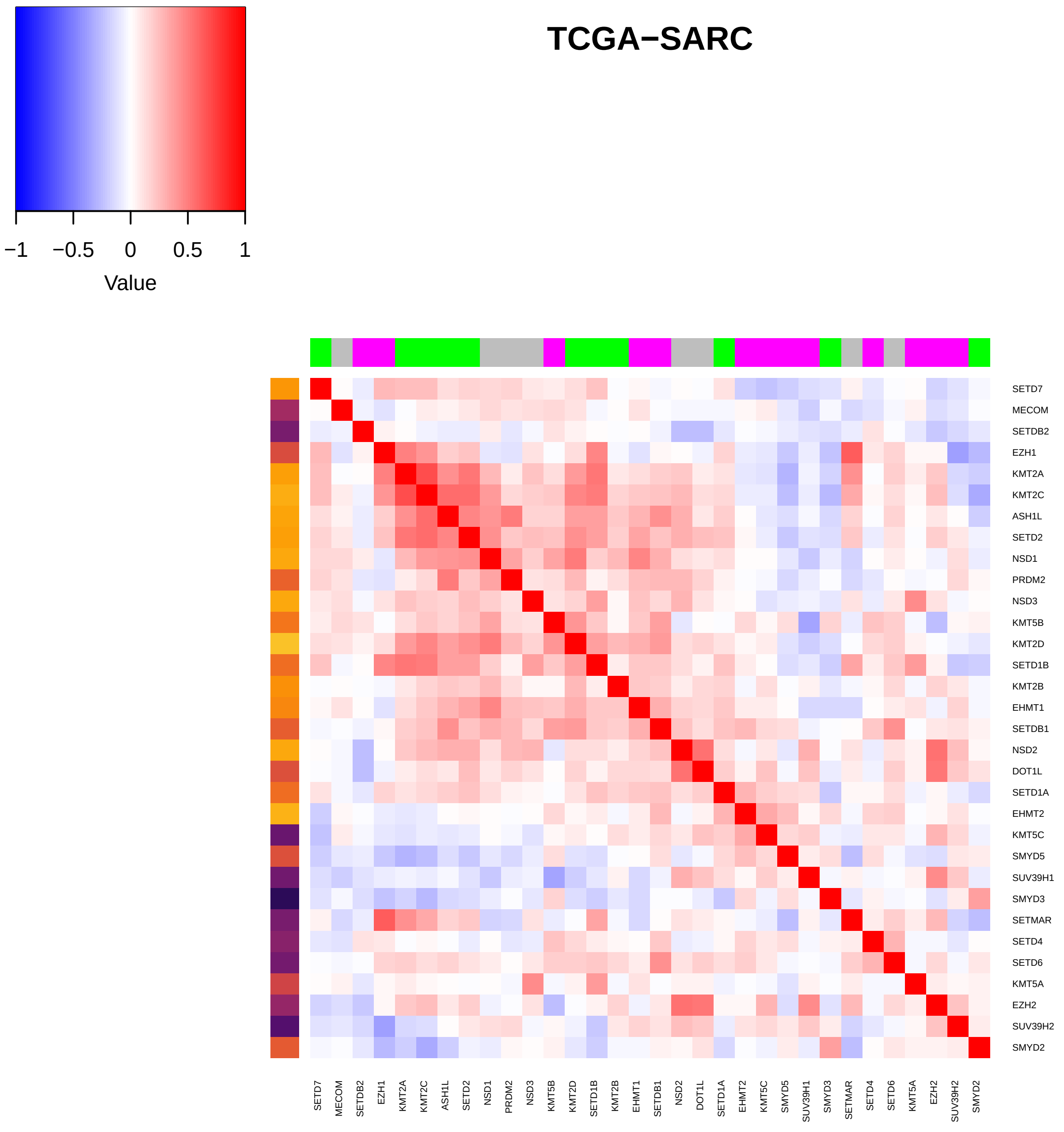

TCGA−SARC
−1
−0.5
0
0.5
1
Value
SETD7
MECOM
SETDB2
EZH1
KMT2A
KMT2C
ASH1L
SETD2
NSD1
PRDM2
NSD3
KMT5B
KMT2D
SETD1B
KMT2B
EHMT1
SETDB1
NSD2
DOT1L
SETD1A
EHMT2
KMT5C
SMYD5
SUV39H1
SMYD3
SETMAR
SETD4
SETD6
KMT5A
EZH2
SUV39H2
SMYD2

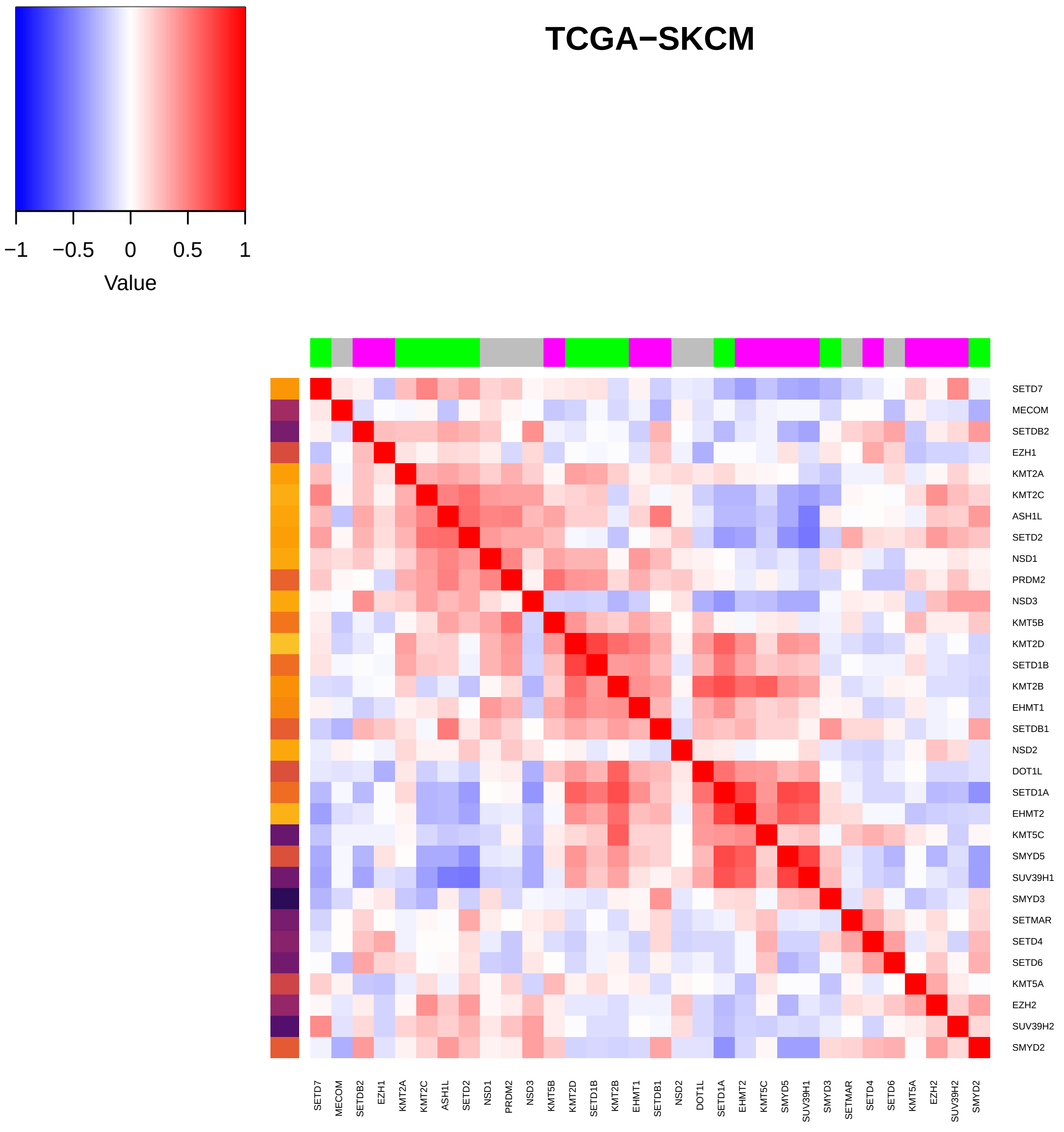

TCGA−SKCM
−1
−0.5
0
0.5
1
Value
SETD7
MECOM
SETDB2
EZH1
KMT2A
KMT2C
ASH1L
SETD2
NSD1
PRDM2
NSD3
KMT5B
KMT2D
SETD1B
KMT2B
EHMT1
SETDB1
NSD2
DOT1L
SETD1A
EHMT2
KMT5C
SMYD5
SUV39H1
SMYD3
SETMAR
SETD4
SETD6
KMT5A
EZH2
SUV39H2
SMYD2

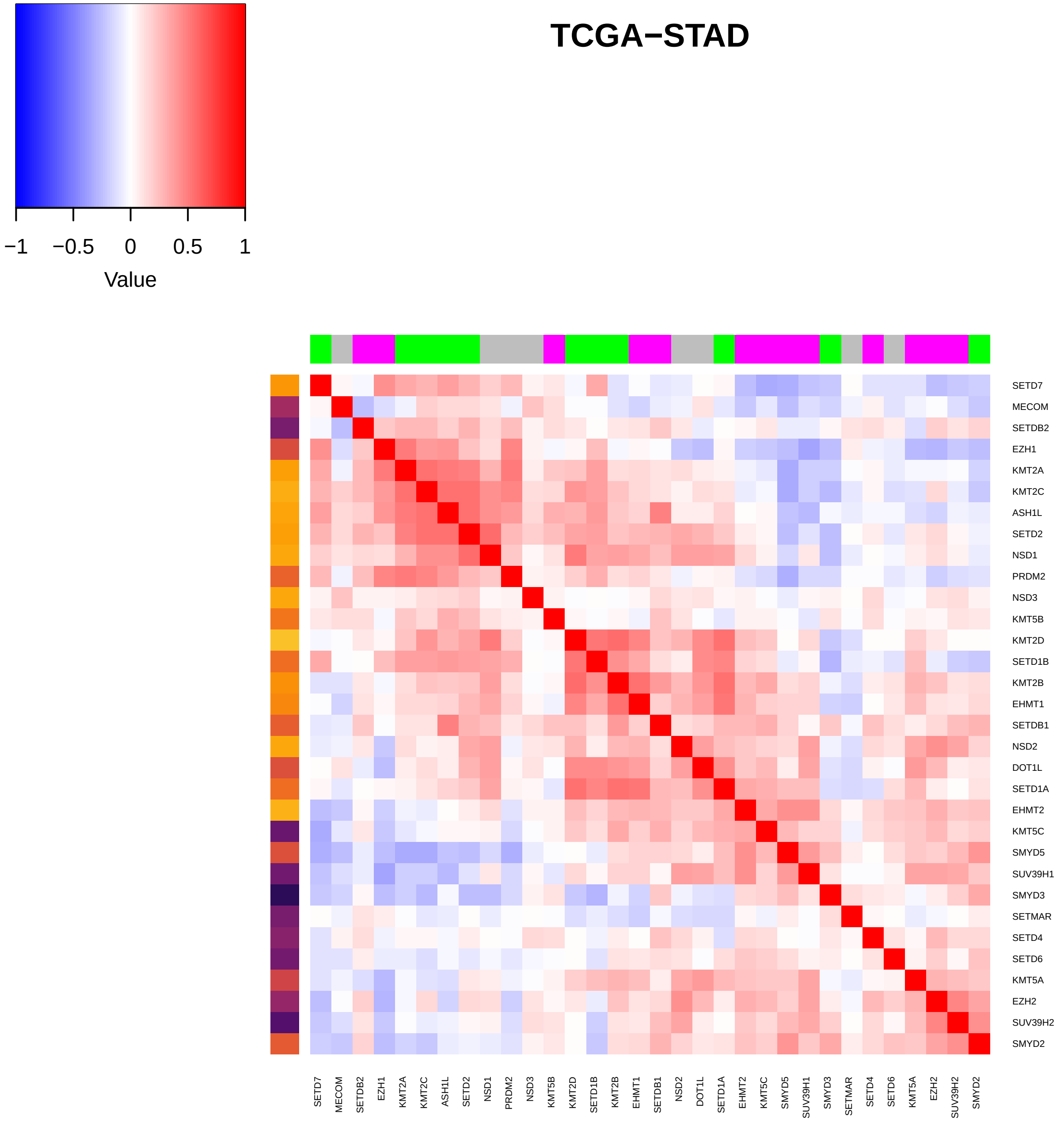
TCGA−STAD
−1
−0.5
0
0.5
1
Value
SETD7
MECOM
SETDB2
EZH1
KMT2A
KMT2C
ASH1L
SETD2
NSD1
PRDM2
NSD3
KMT5B
KMT2D
SETD1B
KMT2B
EHMT1
SETDB1
NSD2
DOT1L
SETD1A
EHMT2
KMT5C
SMYD5
SUV39H1
SMYD3
SETMAR
SETD4
SETD6
KMT5A
EZH2
SUV39H2
SMYD2

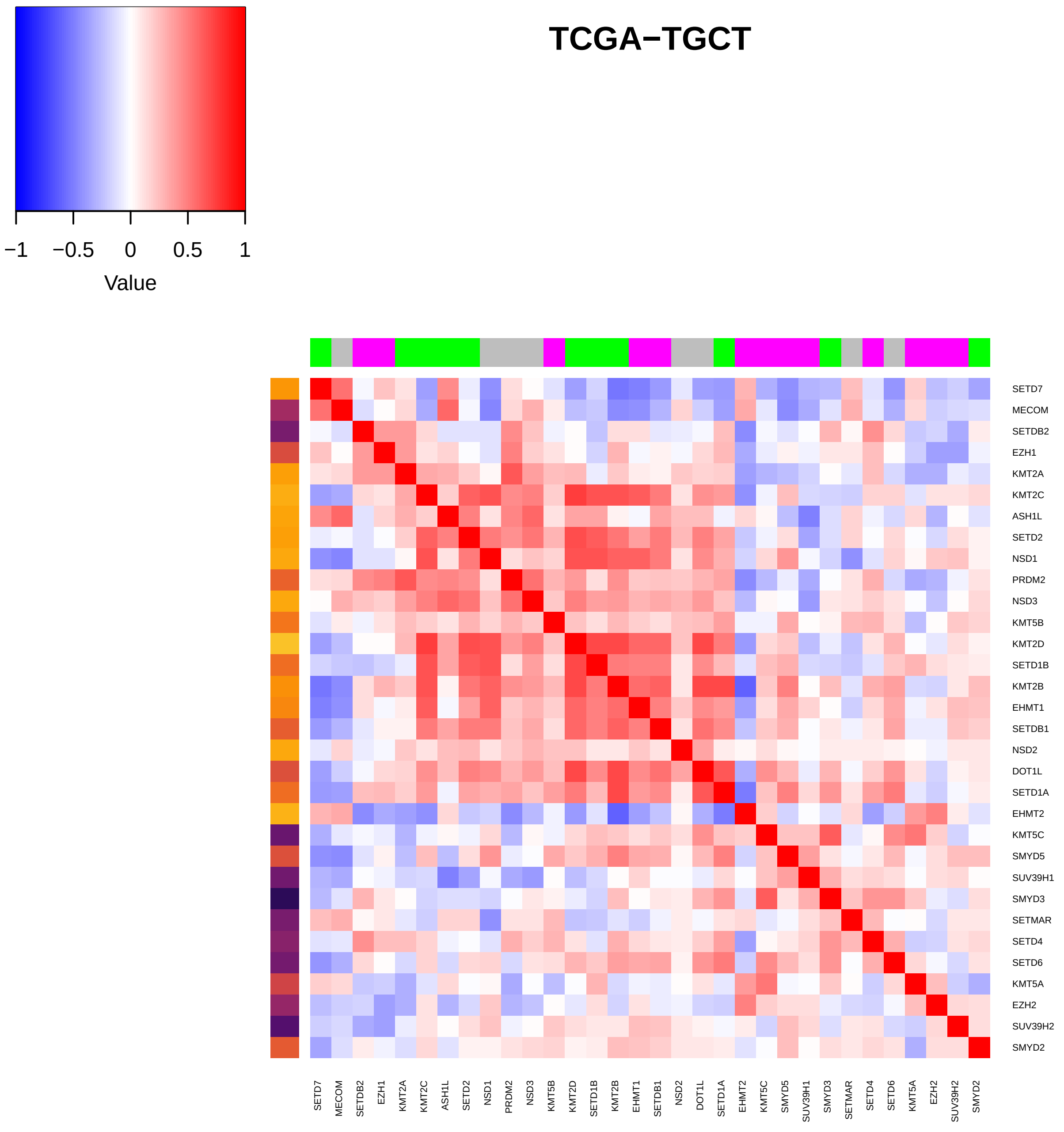

TCGA−TGCT
−1
−0.5
0
0.5
1
Value
SETD7
MECOM
SETDB2
EZH1
KMT2A
KMT2C
ASH1L
SETD2
NSD1
PRDM2
NSD3
KMT5B
KMT2D
SETD1B
KMT2B
EHMT1
SETDB1
NSD2
DOT1L
SETD1A
EHMT2
KMT5C
SMYD5
SUV39H1
SMYD3
SETMAR
SETD4
SETD6
KMT5A
EZH2
SUV39H2
SMYD2

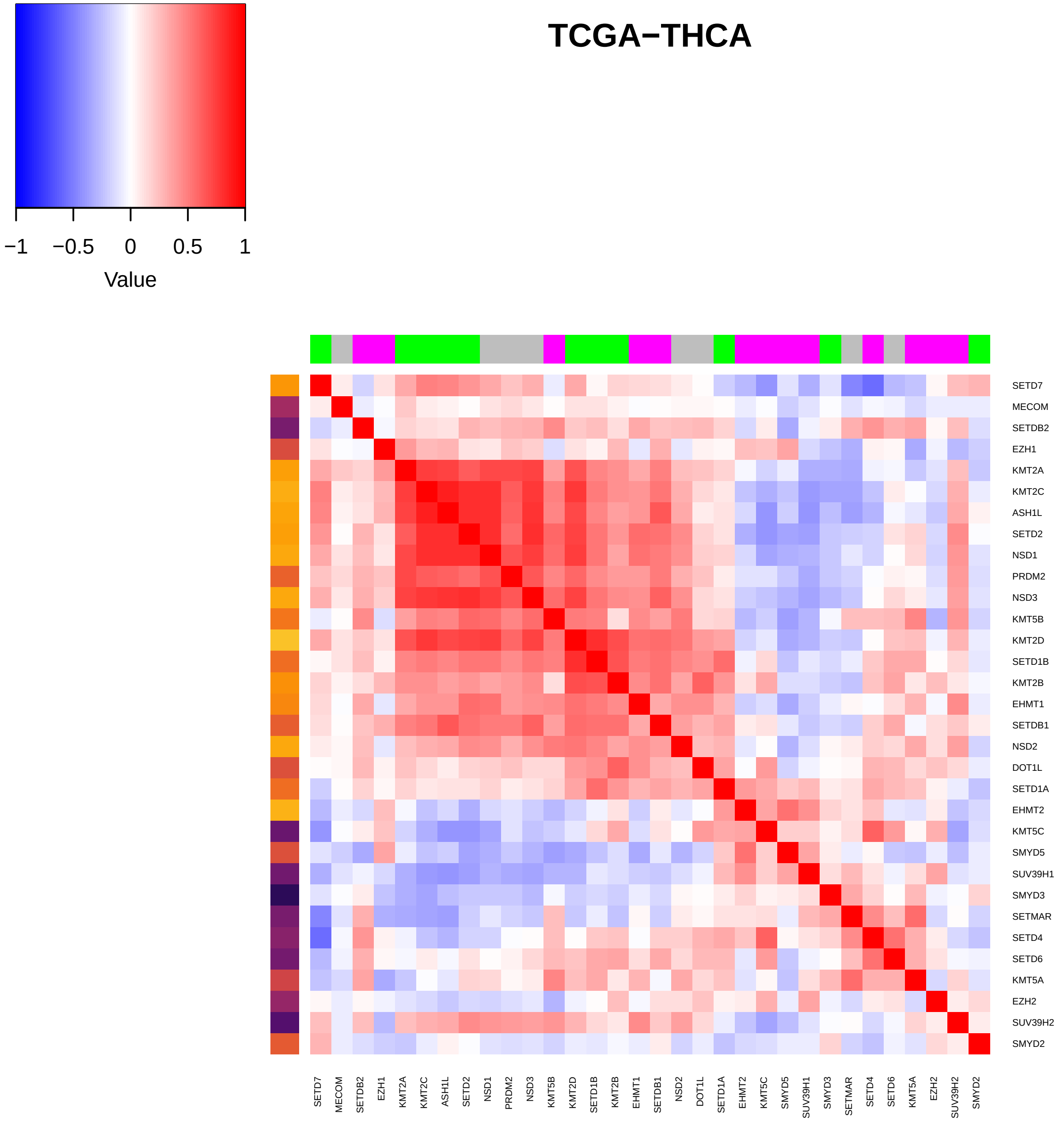

TCGA−THCA
−1
−0.5
0
0.5
1
Value
SETD7
MECOM
SETDB2
EZH1
KMT2A
KMT2C
ASH1L
SETD2
NSD1
PRDM2
NSD3
KMT5B
KMT2D
SETD1B
KMT2B
EHMT1
SETDB1
NSD2
DOT1L
SETD1A
EHMT2
KMT5C
SMYD5
SUV39H1
SMYD3
SETMAR
SETD4
SETD6
KMT5A
EZH2
SUV39H2
SMYD2

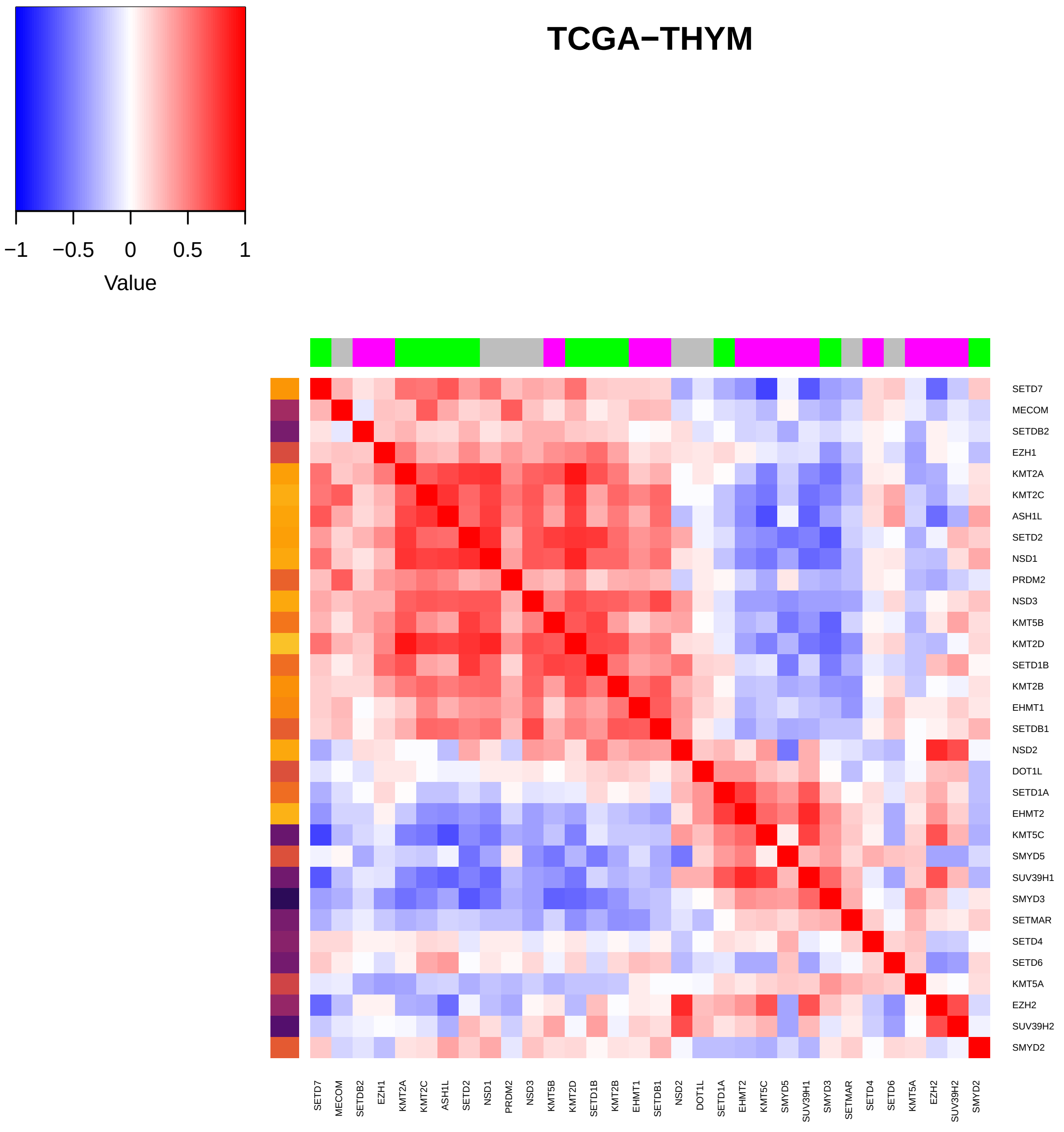
TCGA−THYM
−1
−0.5
0
0.5
1
Value
SETD7
MECOM
SETDB2
EZH1
KMT2A
KMT2C
ASH1L
SETD2
NSD1
PRDM2
NSD3
KMT5B
KMT2D
SETD1B
KMT2B
EHMT1
SETDB1
NSD2
DOT1L
SETD1A
EHMT2
KMT5C
SMYD5
SUV39H1
SMYD3
SETMAR
SETD4
SETD6
KMT5A
EZH2
SUV39H2
SMYD2

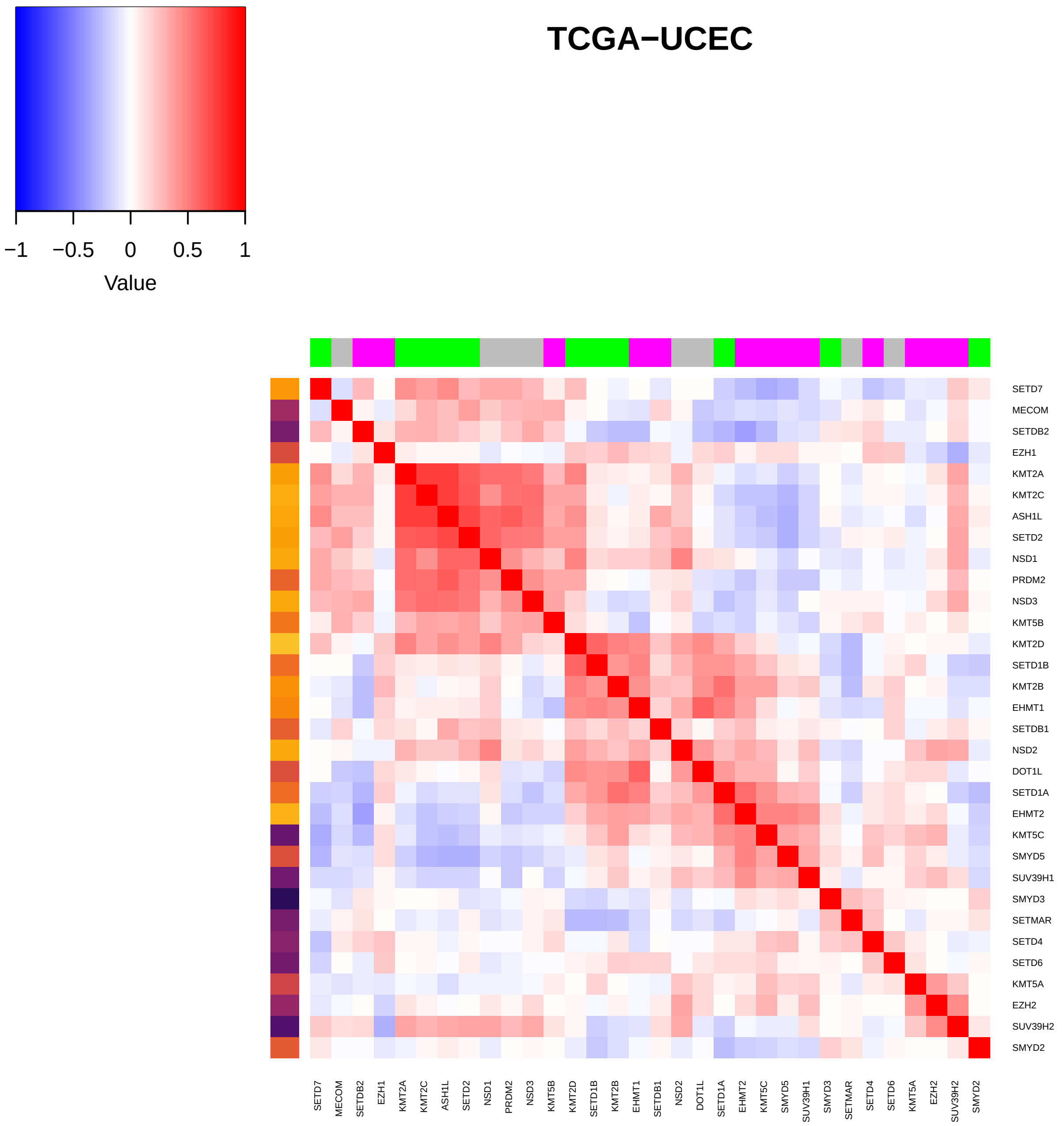

−1
−0.5
0
0.5
1
Value
TCGA−UCEC
SETD7
MECOM
SETDB2
EZH1
KMT2A
KMT2C
ASH1L
SETD2
NSD1
PRDM2
NSD3
KMT5B
KMT2D
SETD1B
KMT2B
EHMT1
SETDB1
NSD2
DOT1L
SETD1A
EHMT2
KMT5C
SMYD5
SUV39H1
SMYD3
SETMAR
SETD4
SETD6
KMT5A
EZH2
SUV39H2
SMYD2

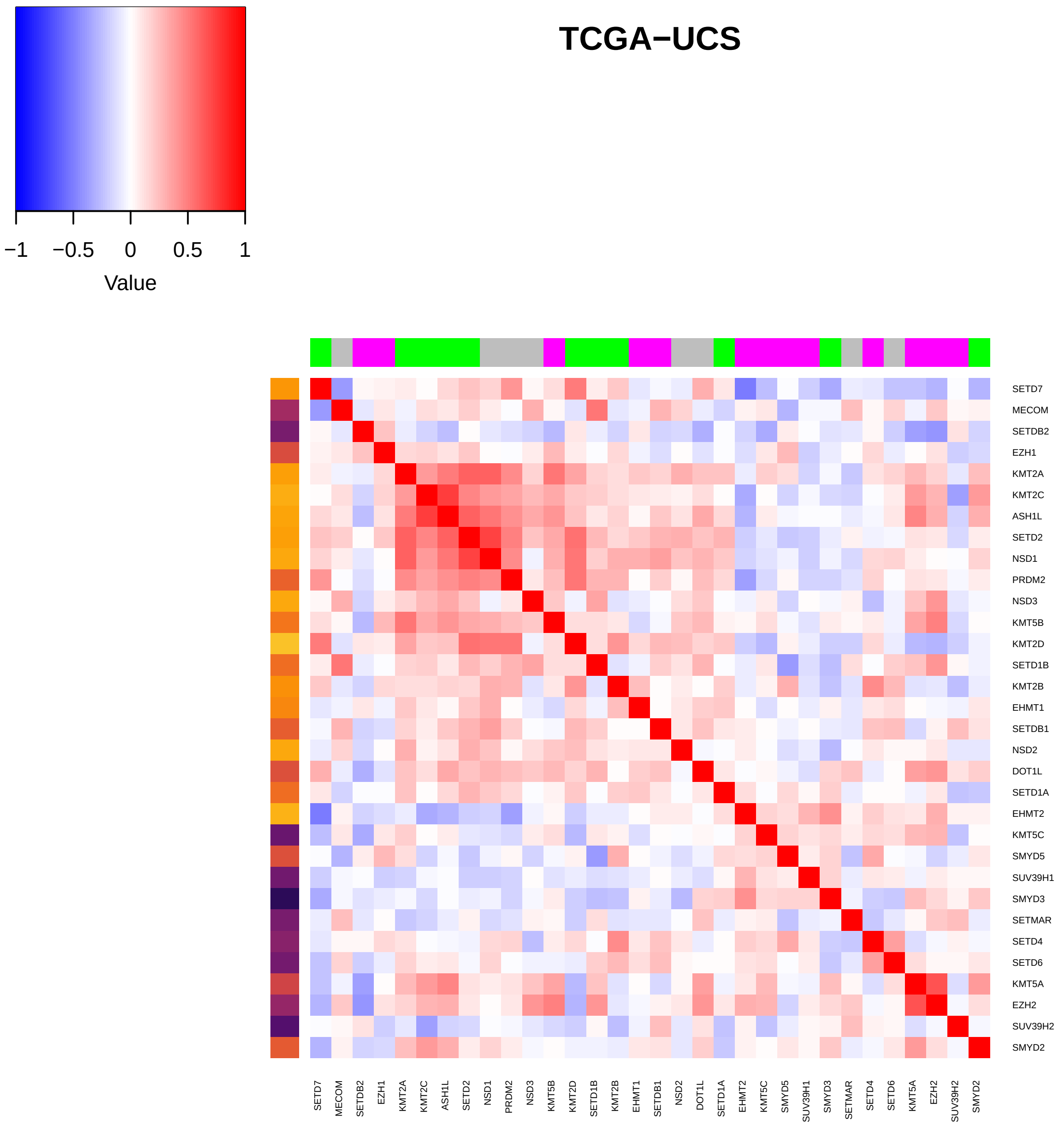

TCGA−UCS
−1
−0.5
0
0.5
1
Value
SETD7
MECOM
SETDB2
EZH1
KMT2A
KMT2C
ASH1L
SETD2
NSD1
PRDM2
NSD3
KMT5B
KMT2D
SETD1B
KMT2B
EHMT1
SETDB1
NSD2
DOT1L
SETD1A
EHMT2
KMT5C
SMYD5
SUV39H1
SMYD3
SETMAR
SETD4
SETD6
KMT5A
EZH2
SUV39H2
SMYD2

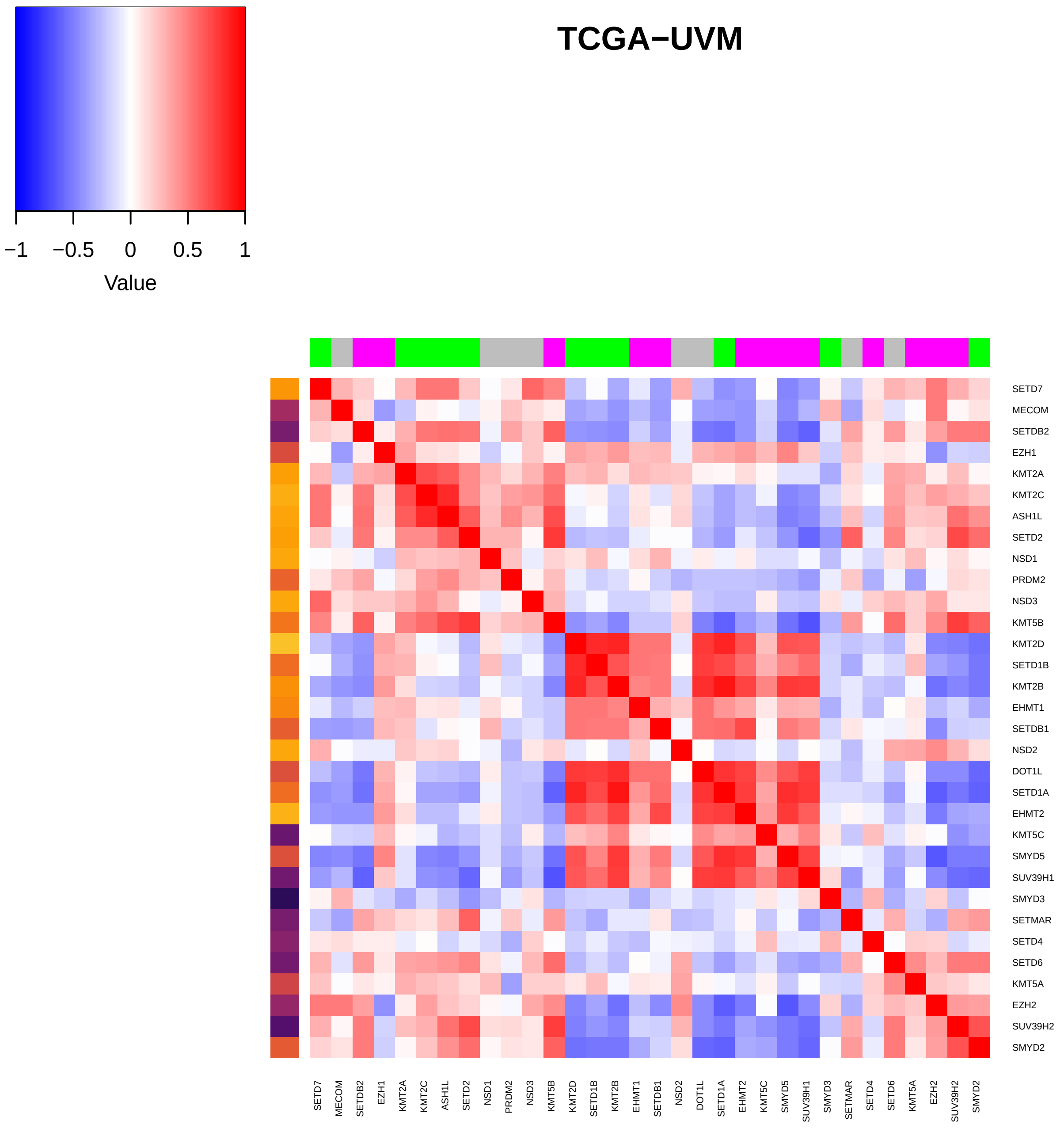
TCGA−UVM
−1
−0.5
0
0.5
1
Value
SETD7
MECOM
SETDB2
EZH1
KMT2A
KMT2C
ASH1L
SETD2
NSD1
PRDM2
NSD3
KMT5B
KMT2D
SETD1B
KMT2B
EHMT1
SETDB1
NSD2
DOT1L
SETD1A
EHMT2
KMT5C
SMYD5
SUV39H1
SMYD3
SETMAR
SETD4
SETD6
KMT5A
EZH2
SUV39H2
SMYD2
