## Supplemental Figures and Tables for "Histone methylation has a direct metabolic role in human cells": Supp File 7. Single HMT RB mutant boxplots.pdf

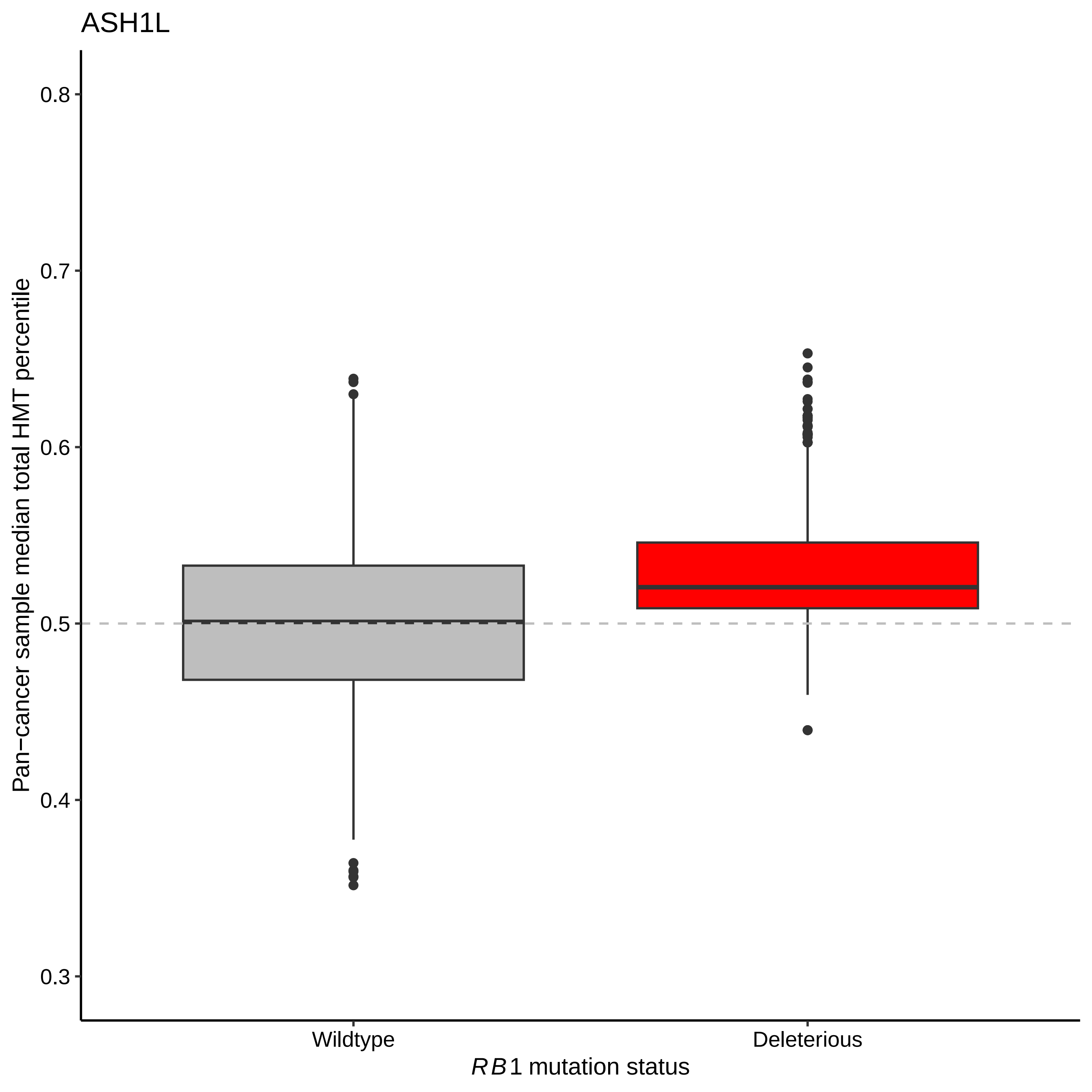

ASH1L
0.8
0.7
0.6
0.5
0.4
0.3
Pan−cancer sample median total HMT percentile
Wildtype
Deleterious
RB1 mutation status

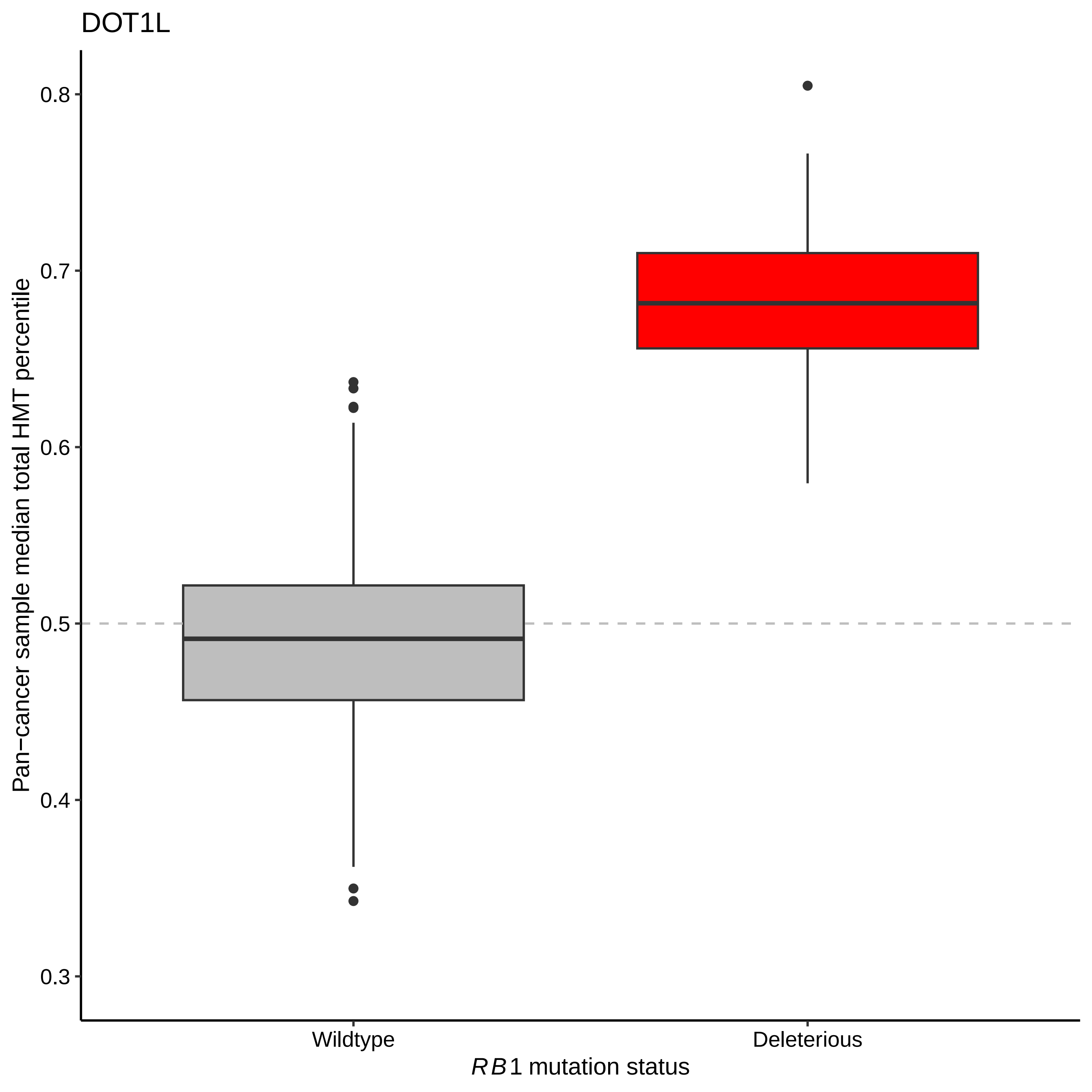
DOT1L
Pan−cancer sample median total HMT percentile
0.8
0.7
0.6
0.5
0.4
0.3
Wildtype
Deleterious
RB1 mutation status

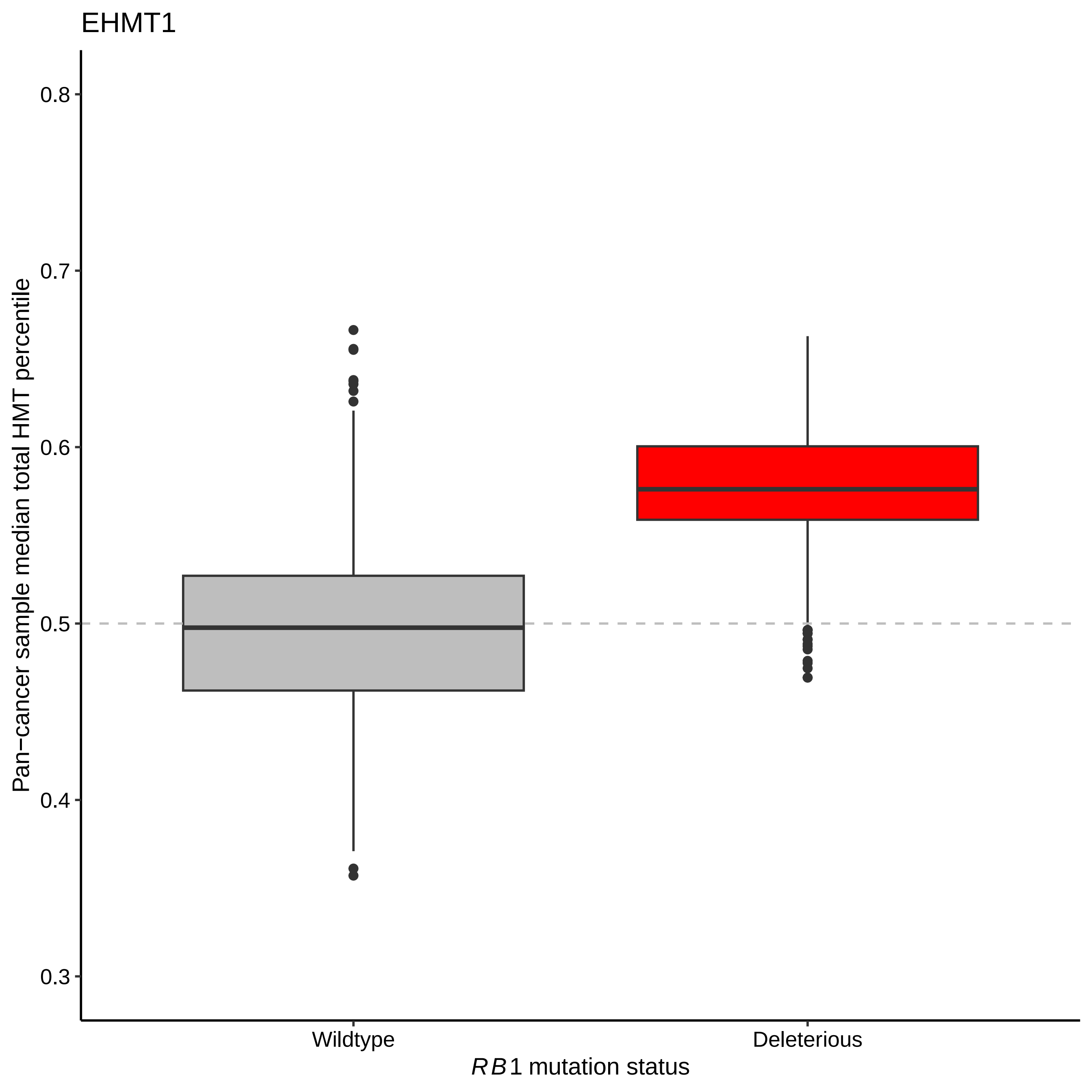
EHMT1
Pan−cancer sample median total HMT percentile
0.8
0.7
0.6
0.5
0.4
0.3
Wildtype
Deleterious
RB1 mutation status

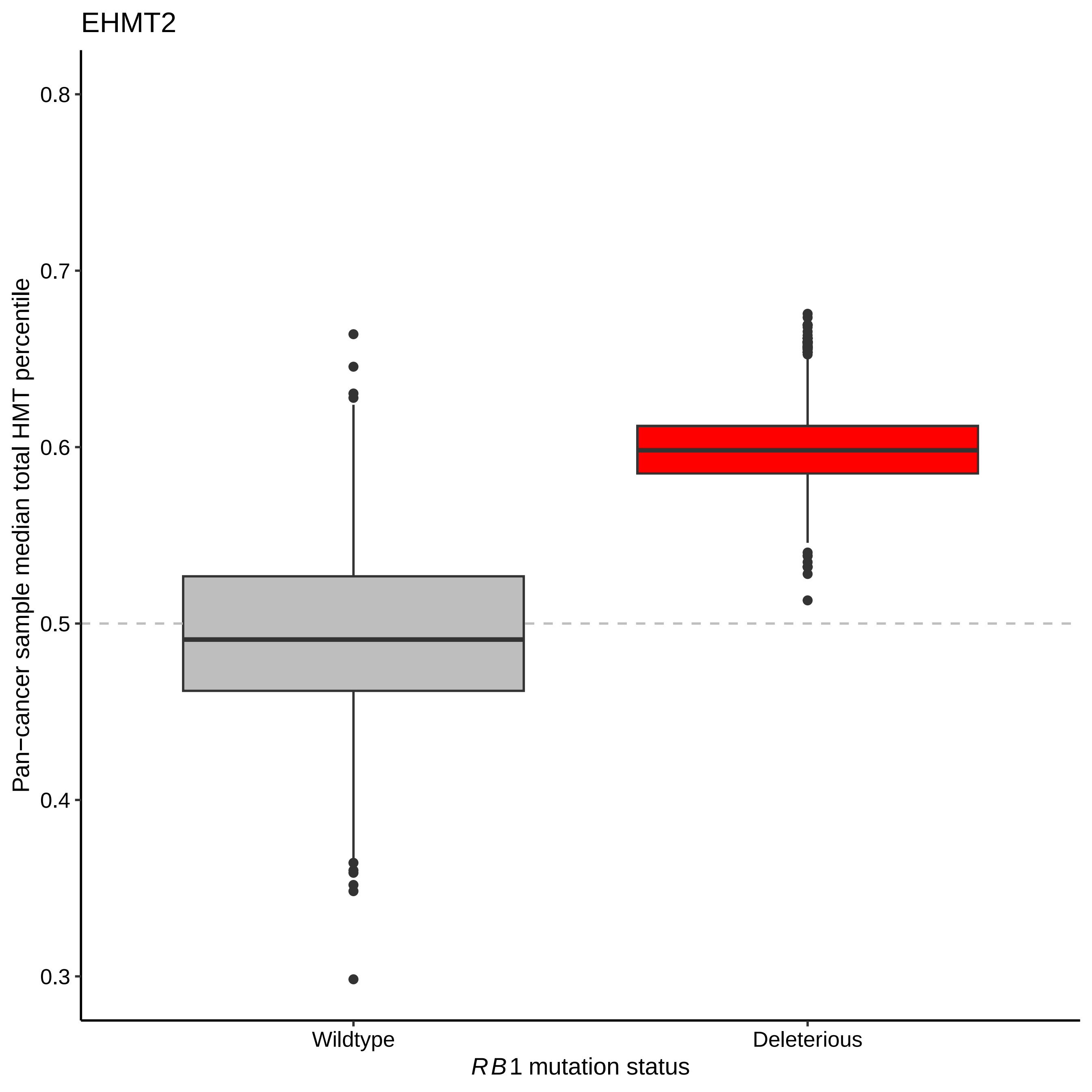
EHMT2
Pan−cancer sample median total HMT percentile
0.8
0.7
0.6
0.5
0.4
0.3
Wildtype
Deleterious
RB1 mutation status

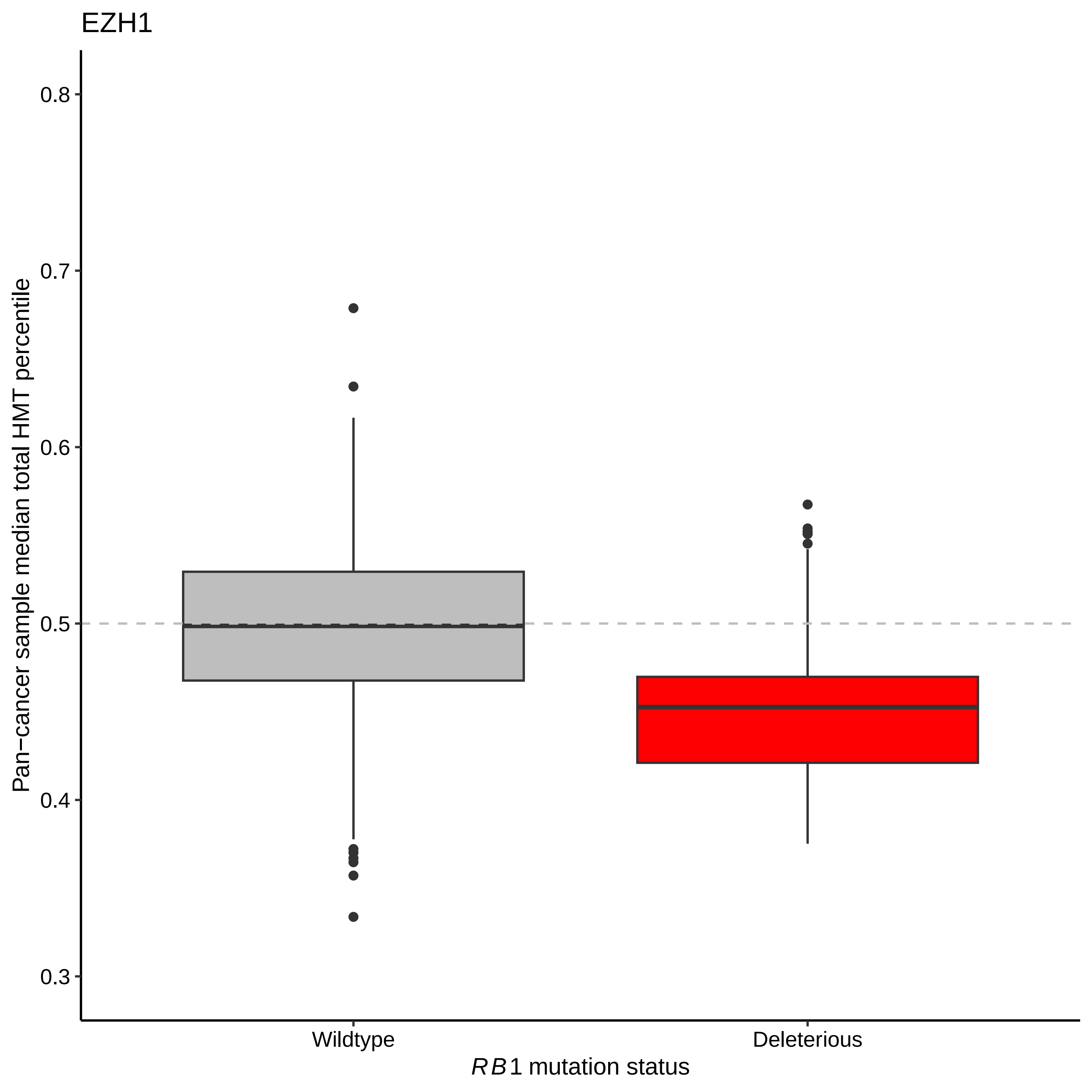
EZH1
Pan−cancer sample median total HMT percentile
0.8
0.7
0.6
0.5
0.4
0.3
Wildtype
Deleterious
RB1 mutation status

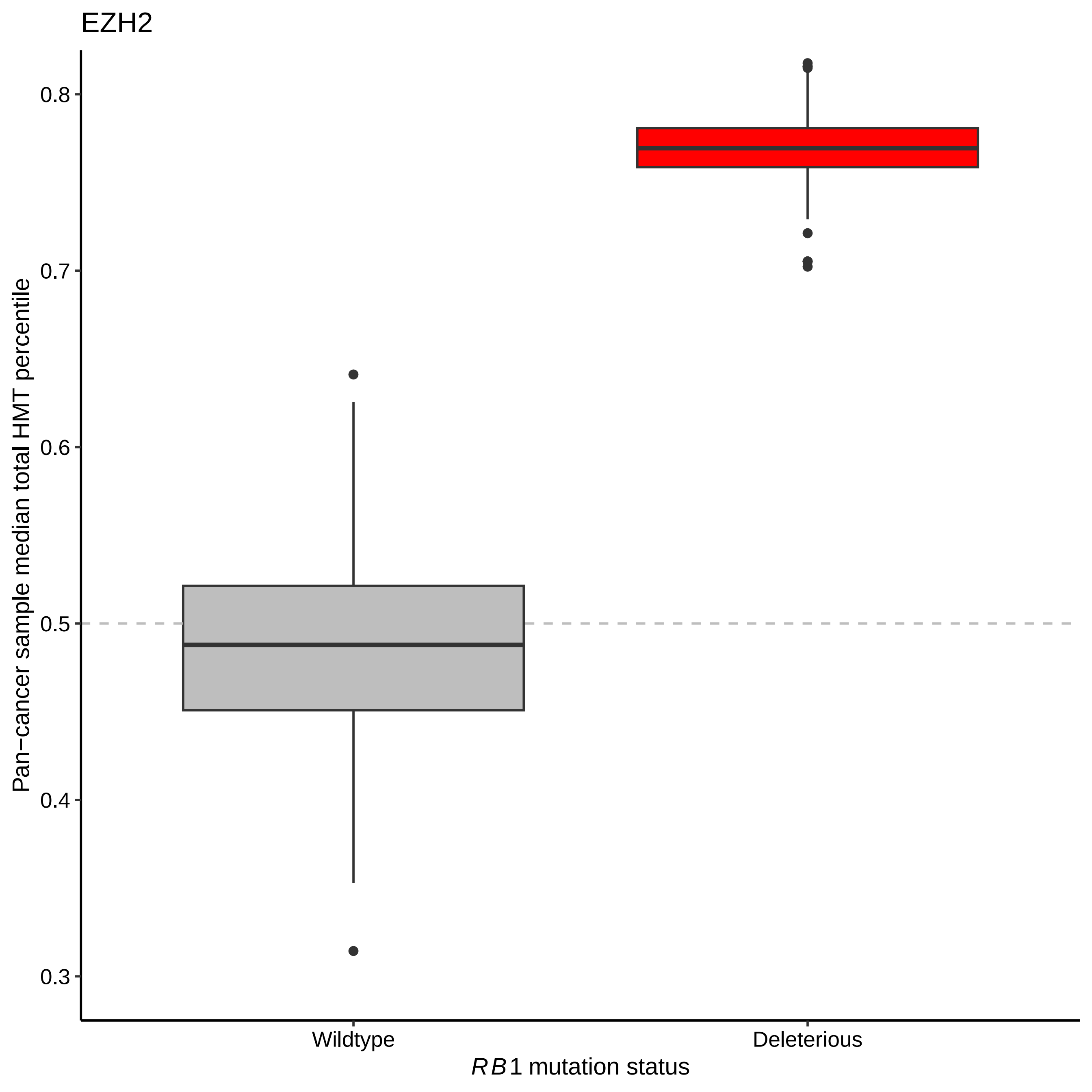

EZH2
Pan−cancer sample median total HMT percentile
0.8
0.7
0.6
0.5
0.4
0.3
Wildtype
Deleterious
RB1 mutation status

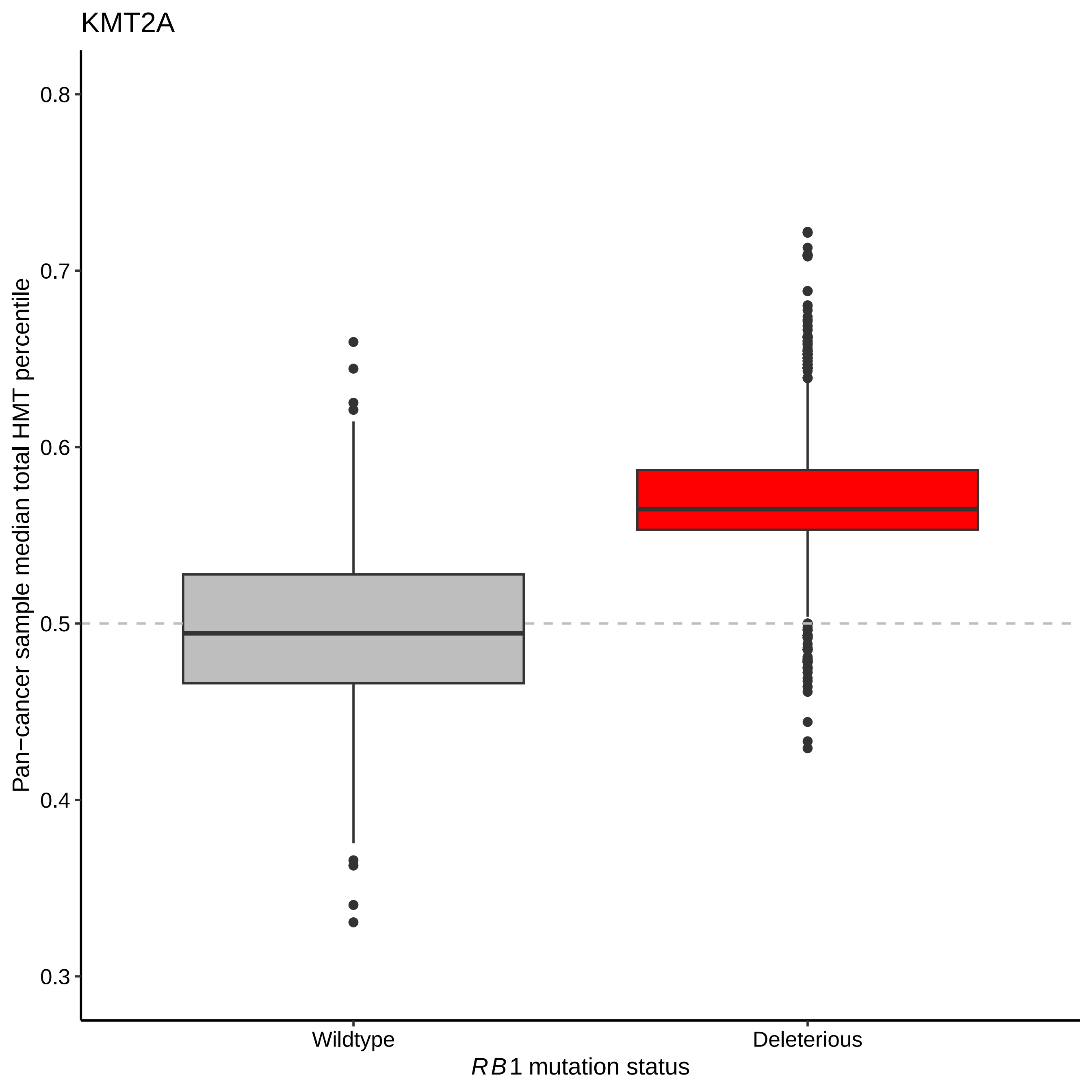
KMT2A
Pan−cancer sample median total HMT percentile
0.8
0.7
0.6
0.5
0.4
0.3
Wildtype
Deleterious
RB1 mutation status

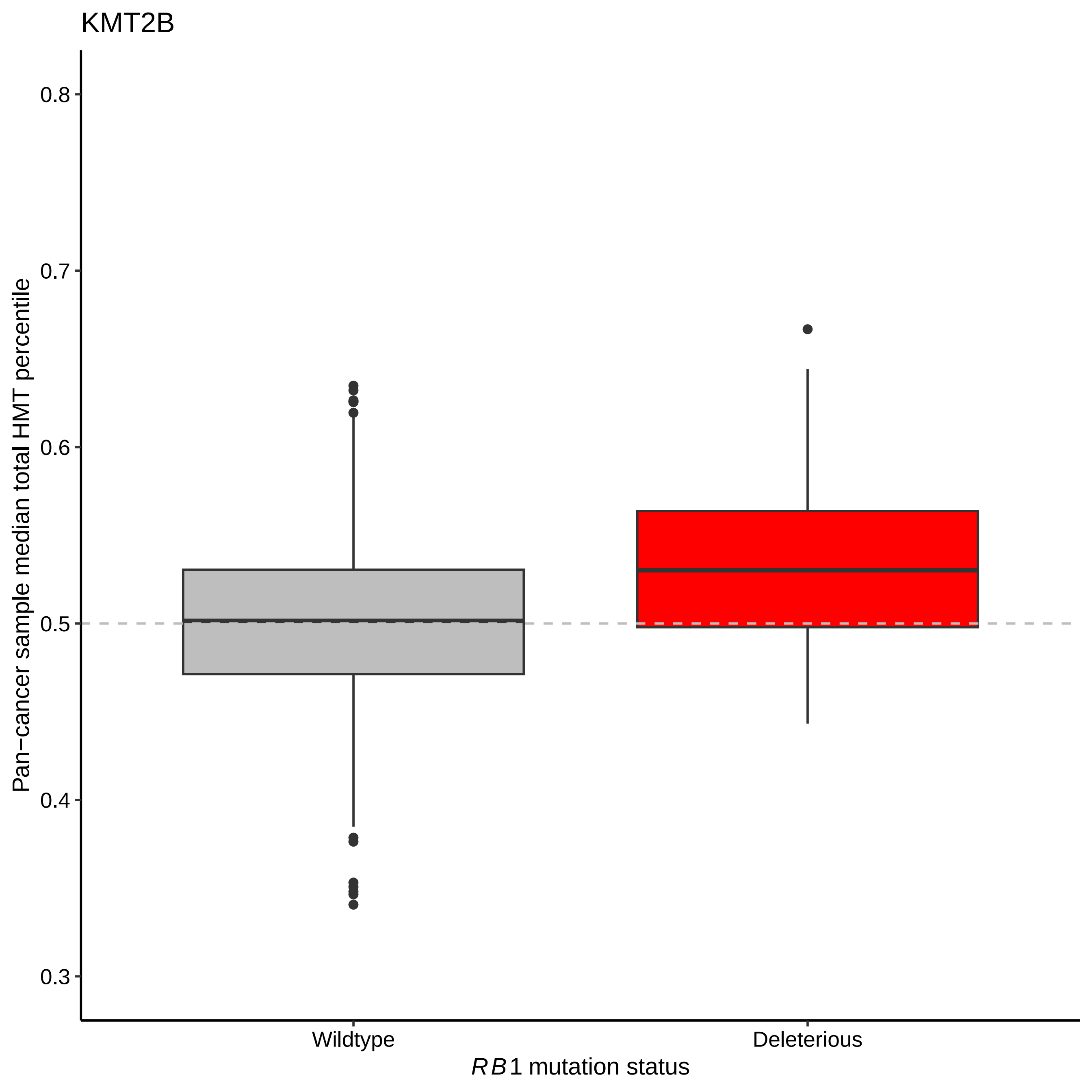

KMT2B
Pan−cancer sample median total HMT percentile
0.8
0.7
0.6
0.5
0.4
0.3
Wildtype
Deleterious
RB1 mutation status

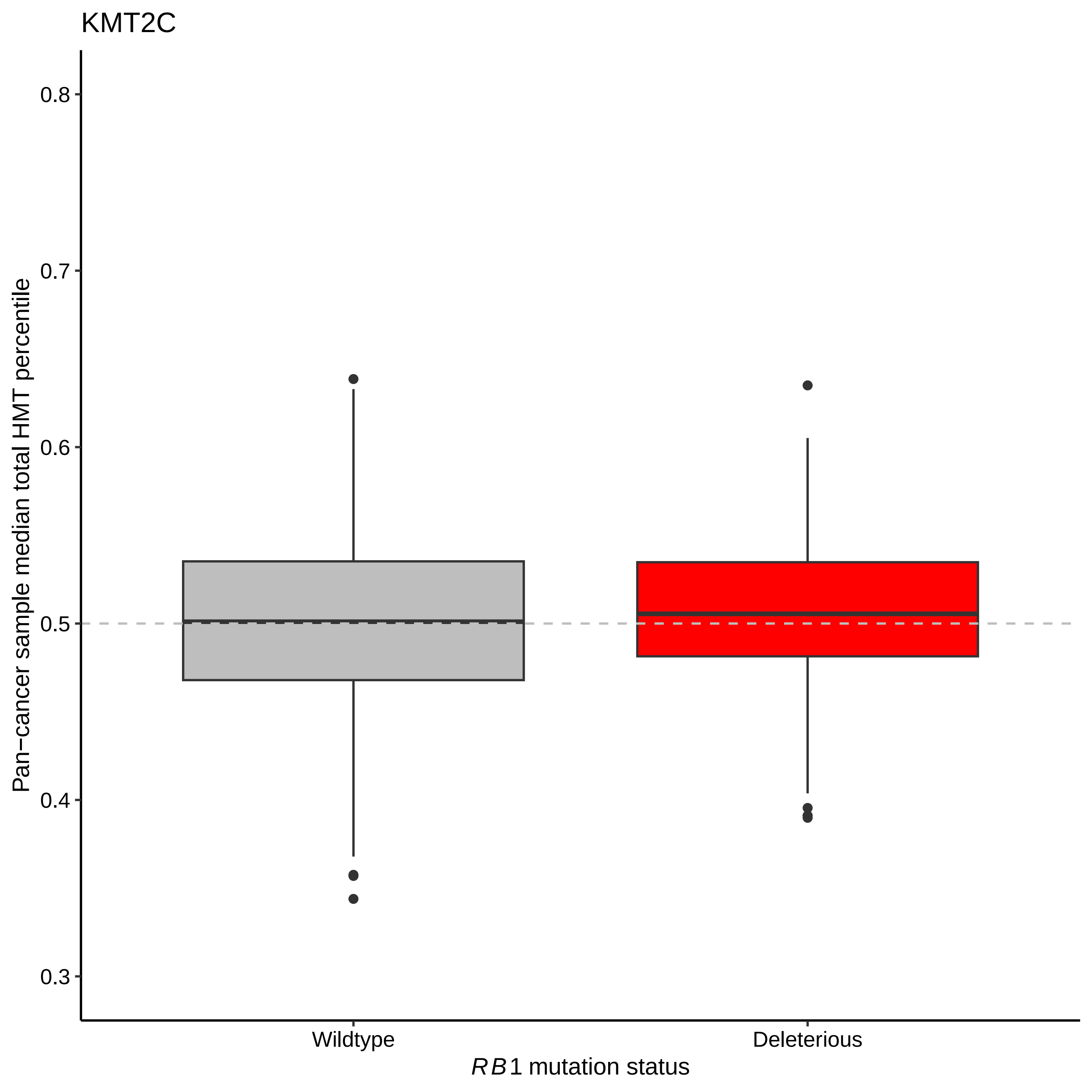
KMT2C
Pan−cancer sample median total HMT percentile
0.8
0.7
0.6
0.5
0.4
0.3
Wildtype
Deleterious
RB1 mutation status

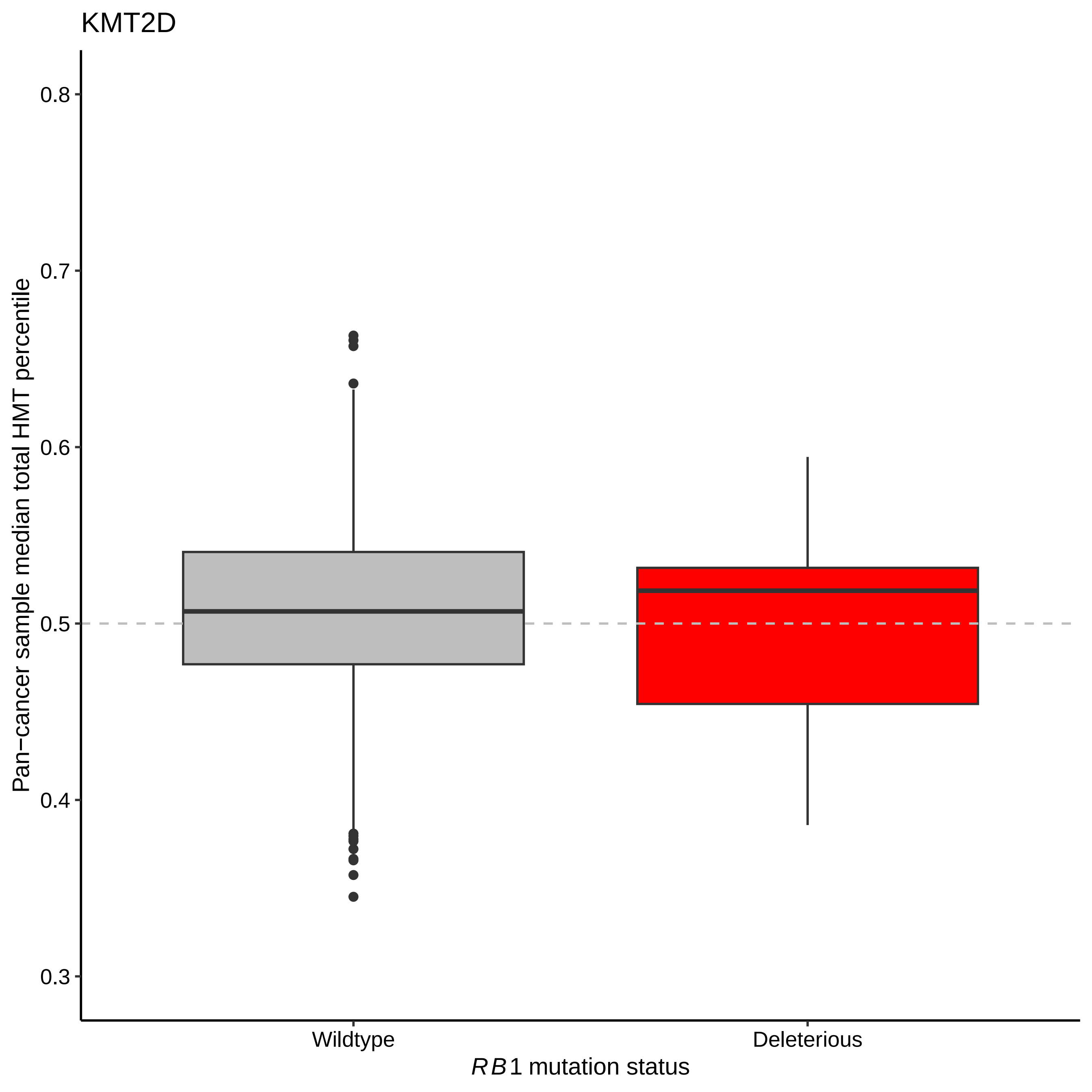

KMT2D
Pan−cancer sample median total HMT percentile
0.8
0.7
0.6
0.5
0.4
0.3
Wildtype
Deleterious
RB1 mutation status

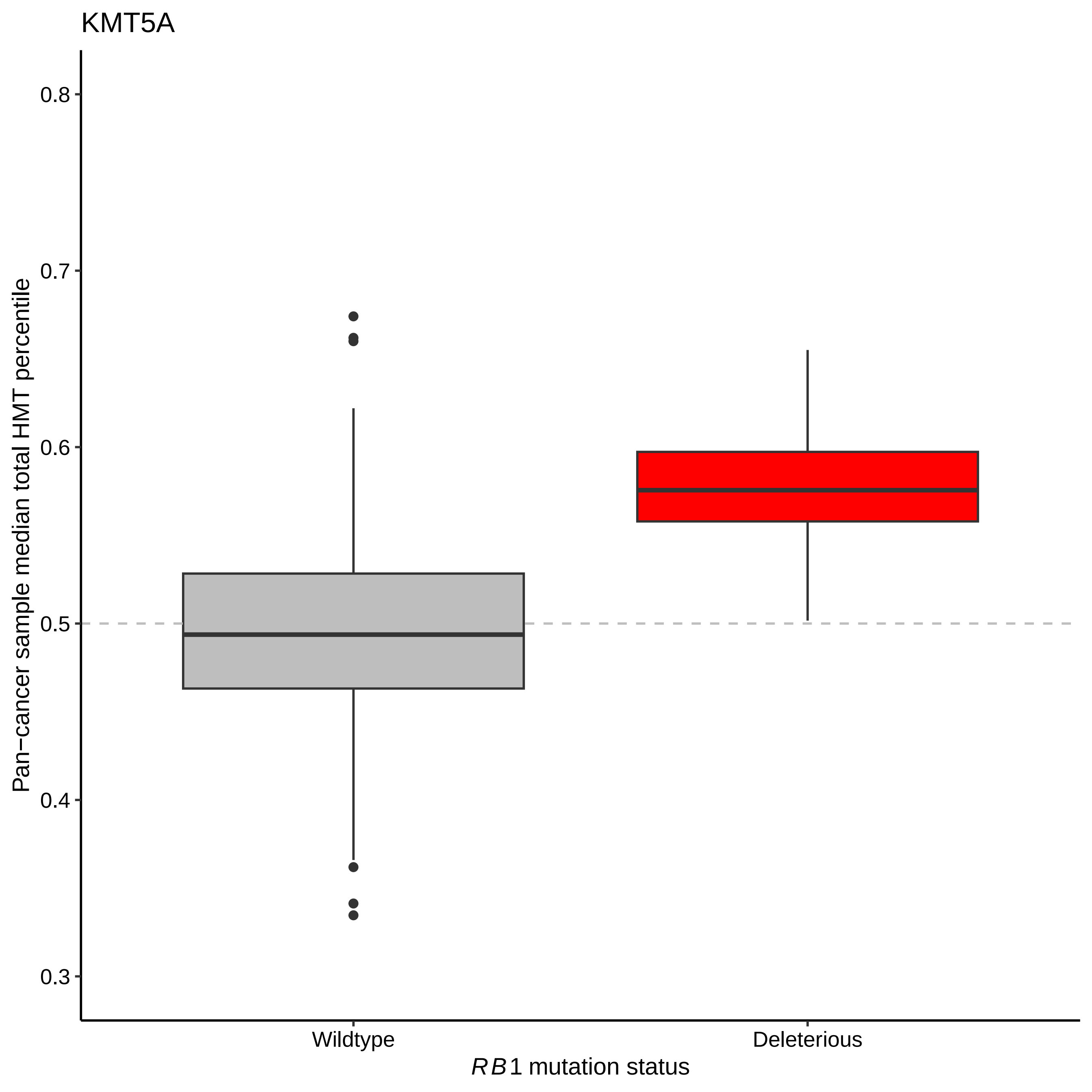
KMT5A
Pan−cancer sample median total HMT percentile
0.8
0.7
0.6
0.5
0.4
0.3
Wildtype
Deleterious
RB1 mutation status

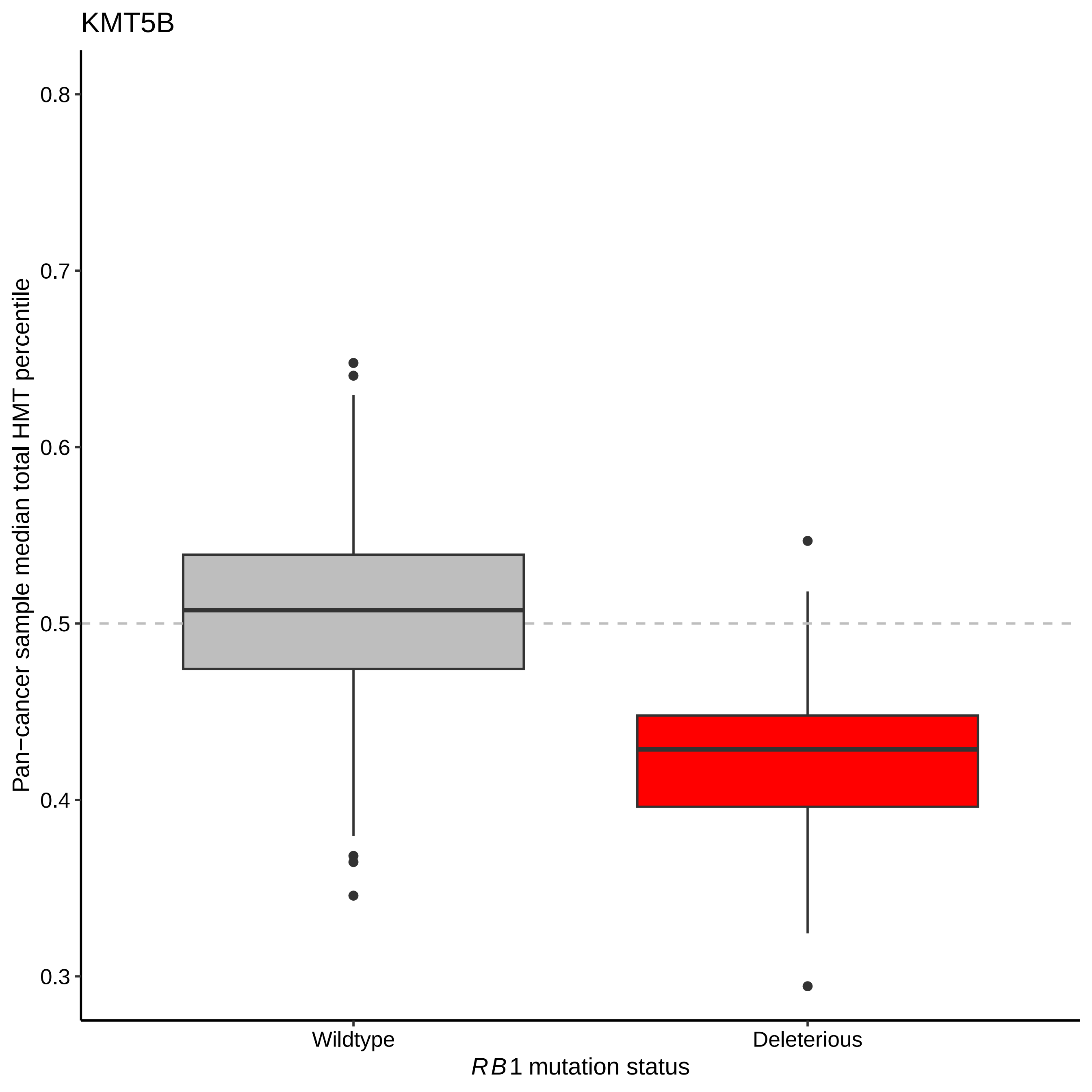

KMT5B
0.8
0.7
0.6
0.5
0.4
0.3
Pan−cancer sample median total HMT percentile
Wildtype
Deleterious
*RB1* mutation status

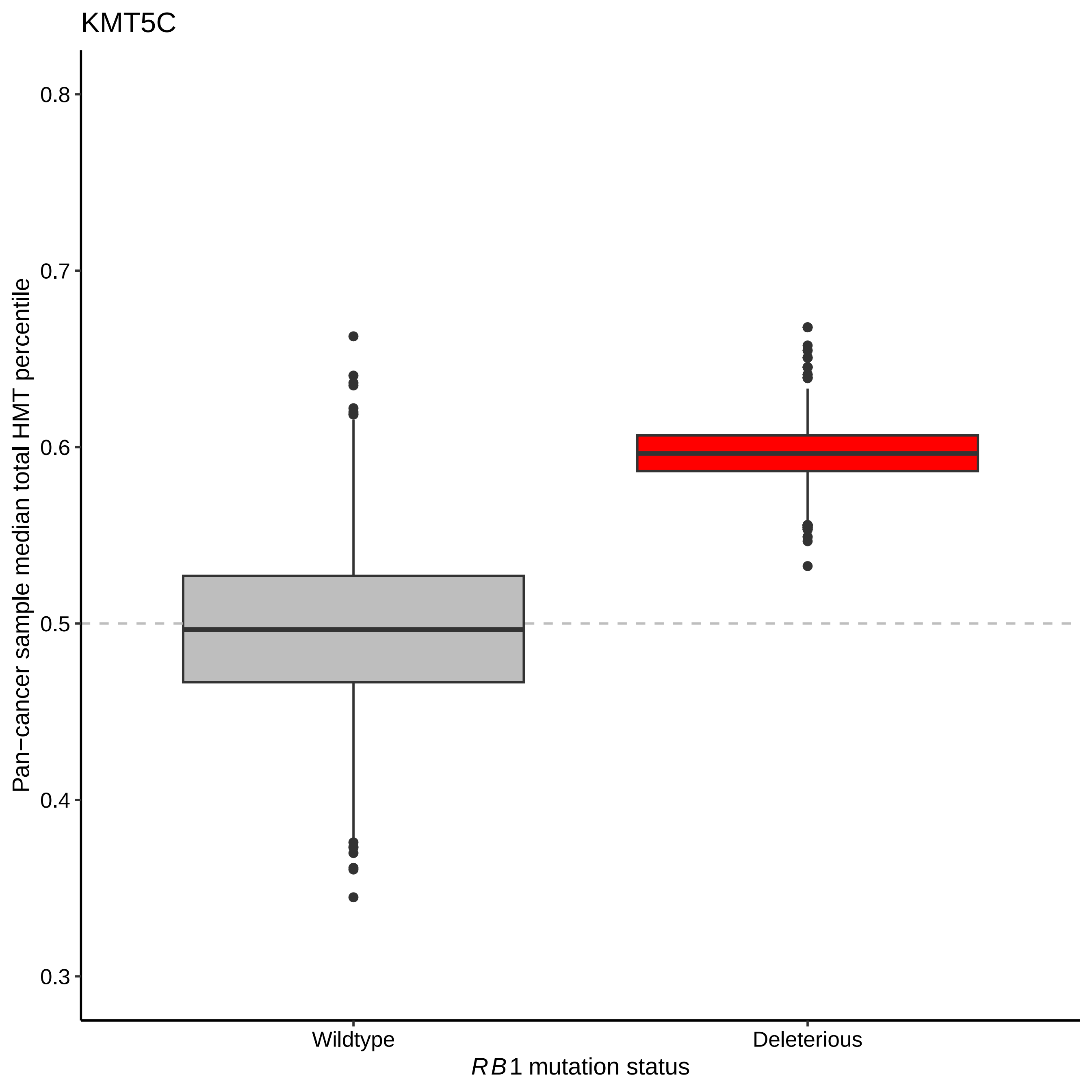
KMT5C
Pan−cancer sample median total HMT percentile
0.8
0.7
0.6
0.5
0.4
0.3
Wildtype
Deleterious
*RB1* mutation status

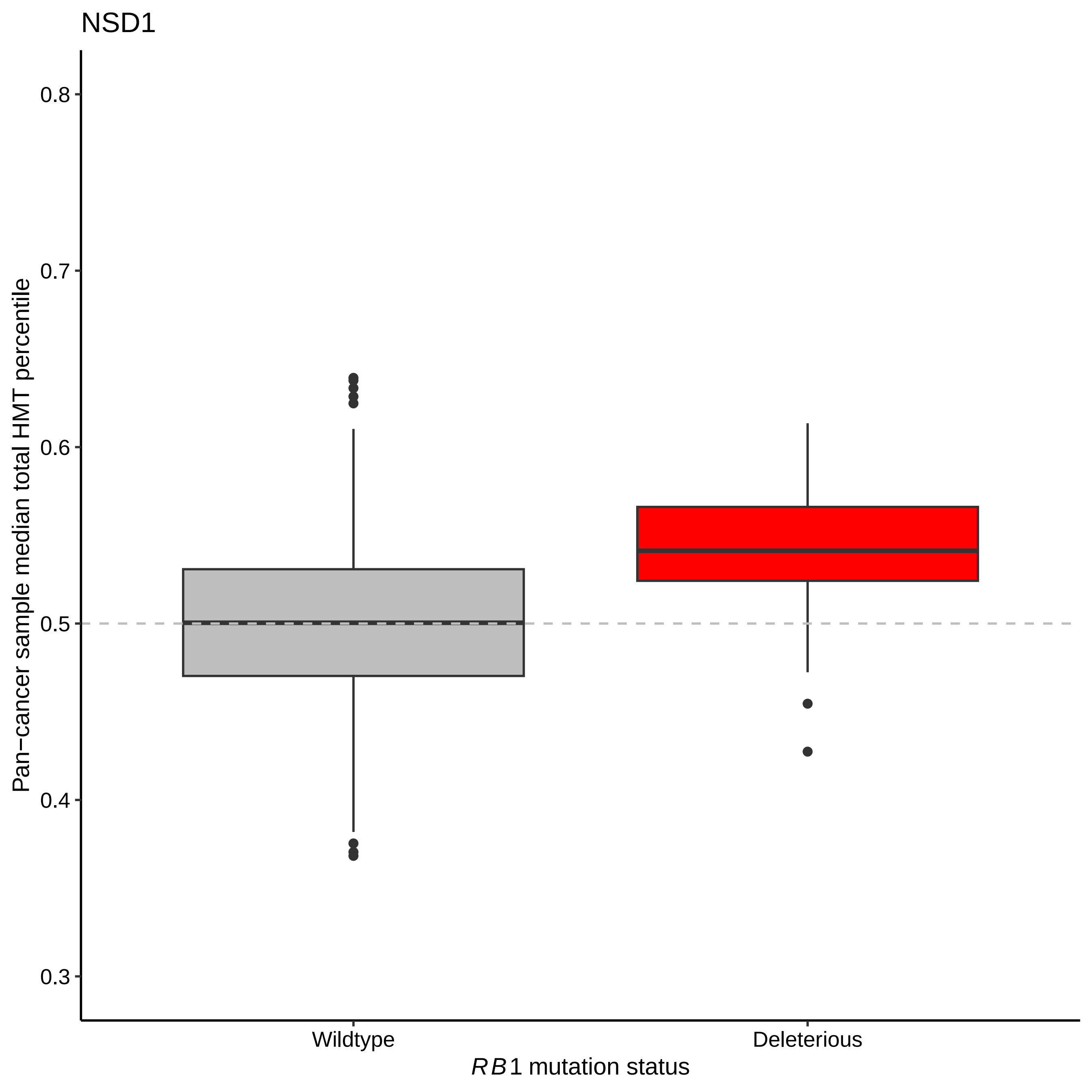

NSD1
Pan−cancer sample median total HMT percentile
0.8
0.7
0.6
0.5
0.4
0.3
Wildtype
Deleterious
RB1 mutation status

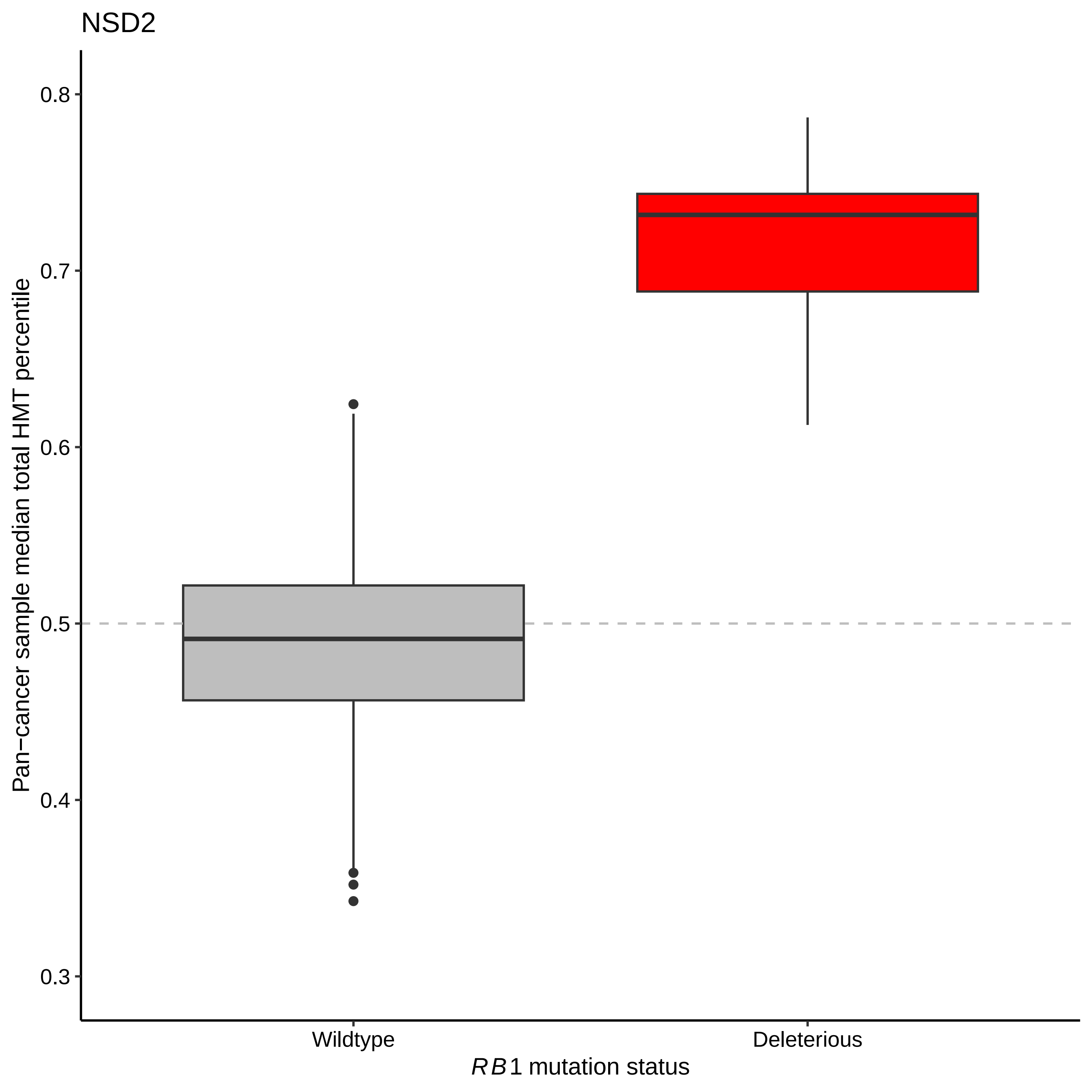
NSD2
Pan−cancer sample median total HMT percentile
0.8
0.7
0.6
0.5
0.4
0.3
Wildtype
Deleterious
RB1 mutation status

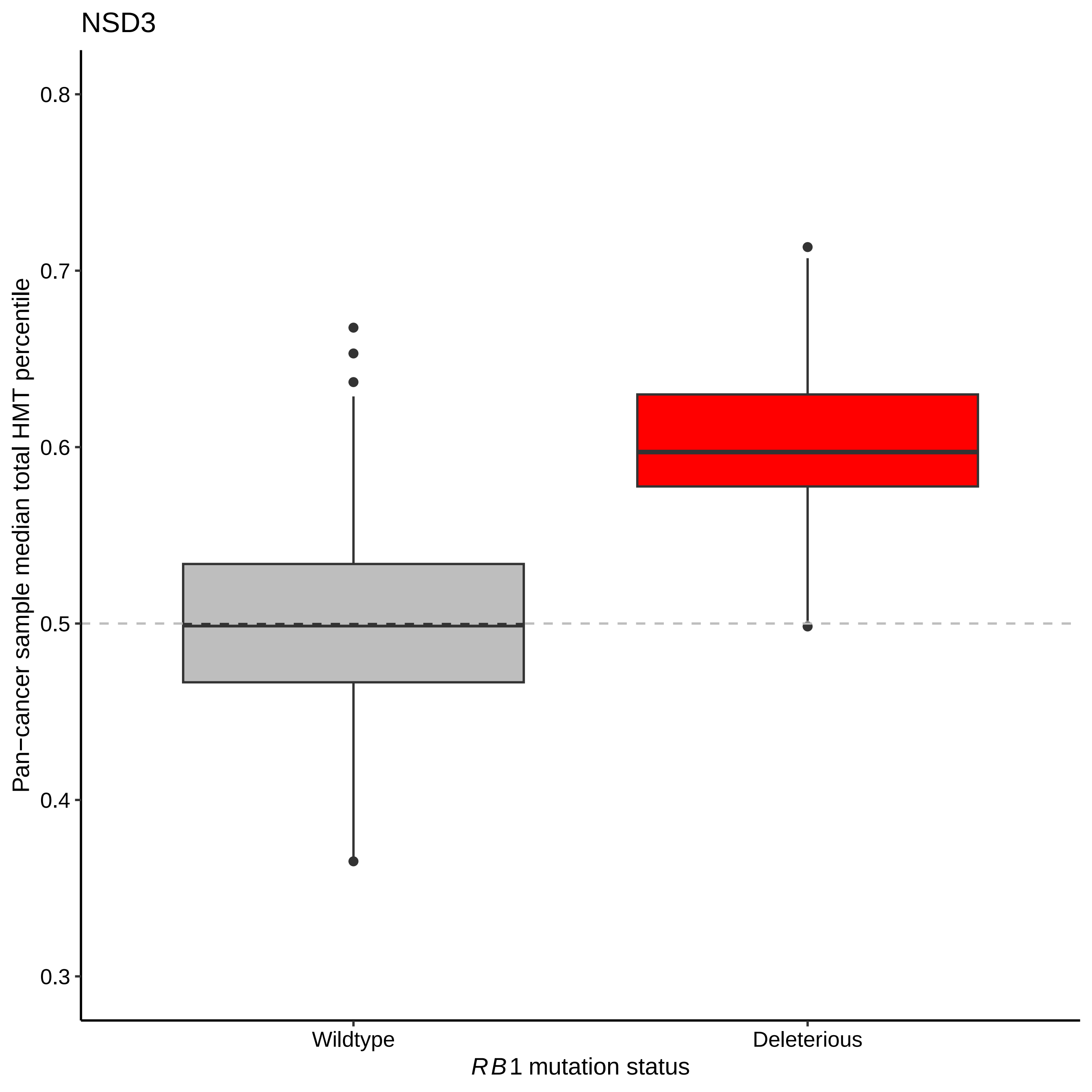

NSD3
Pan−cancer sample median total HMT percentile
0.8
0.7
0.6
0.5
0.4
0.3
Wildtype
Deleterious
RB1 mutation status

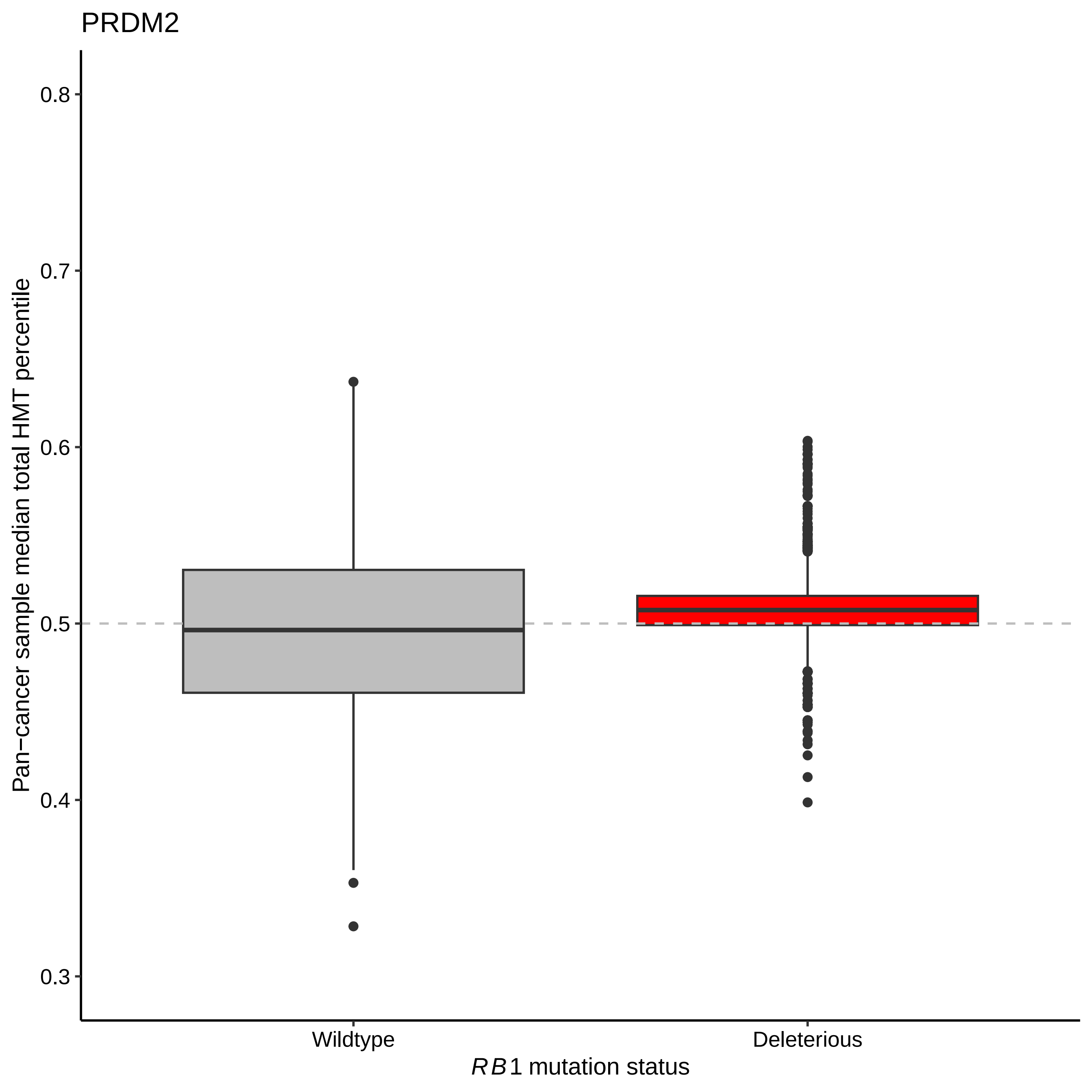

PRDM2
Pan−cancer sample median total HMT percentile
0.8
0.7
0.6
0.5
0.4
0.3
Wildtype
Deleterious
RB1 mutation status

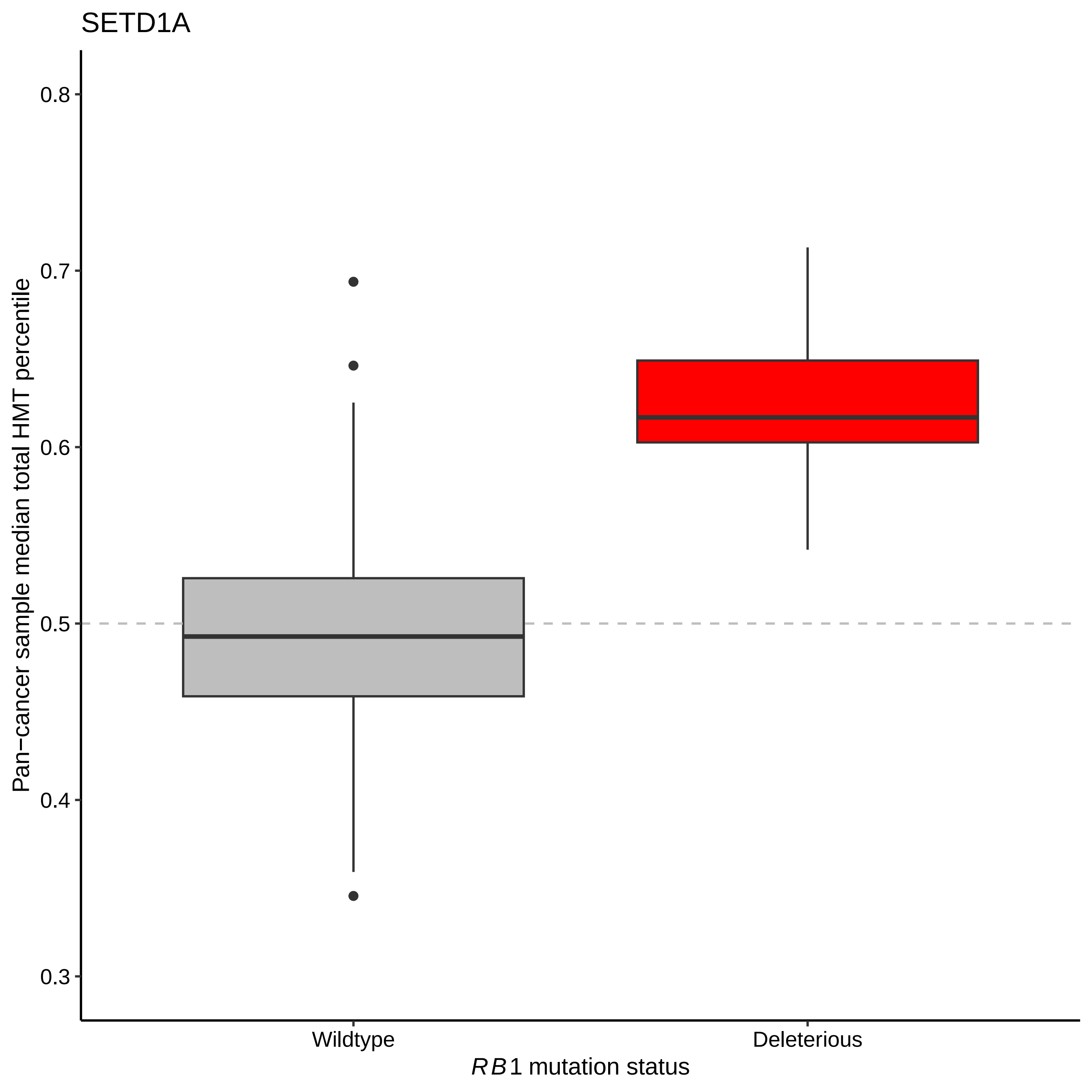

SETD1A
0.8
0.7
0.6
0.5
0.4
0.3
Pan−cancer sample median total HMT percentile
Wildtype
Deleterious
*RB1* mutation status

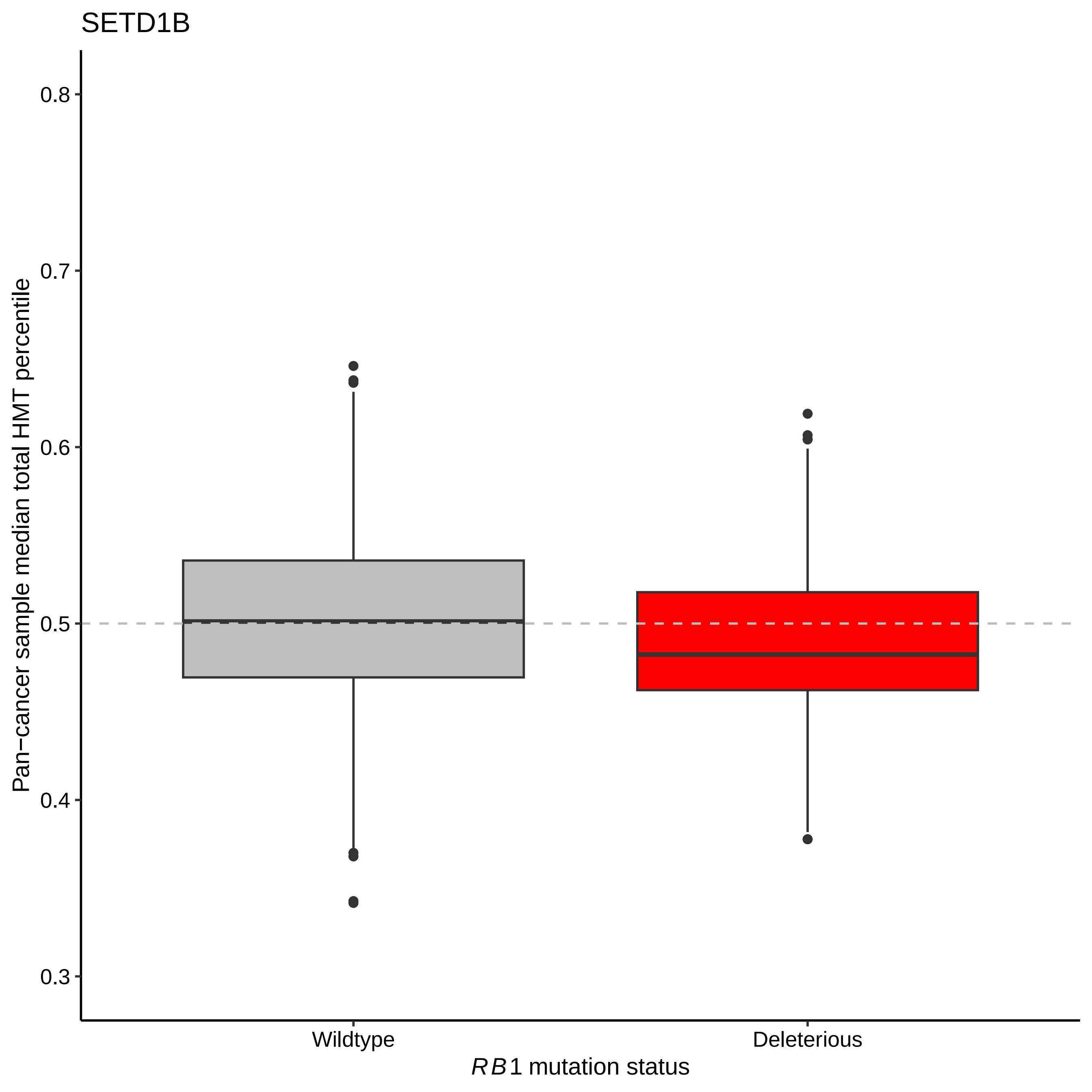

SETD1B
Pan−cancer sample median total HMT percentile
0.8
0.7
0.6
0.5
0.4
0.3
Wildtype
Deleterious
RB1 mutation status

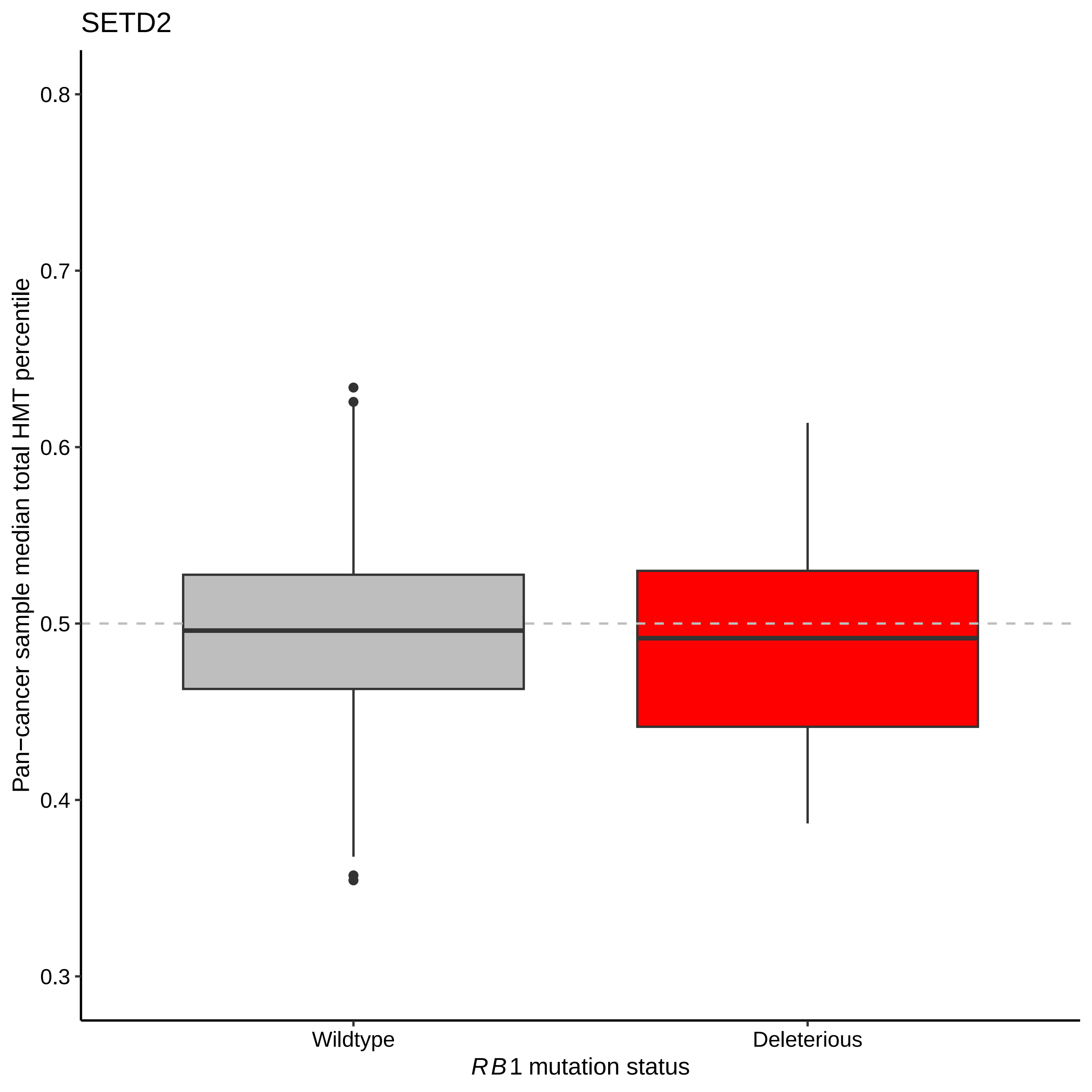
SETD2
Pan−cancer sample median total HMT percentile
0.8
0.7
0.6
0.5
0.4
0.3
Wildtype
Deleterious
RB1 mutation status

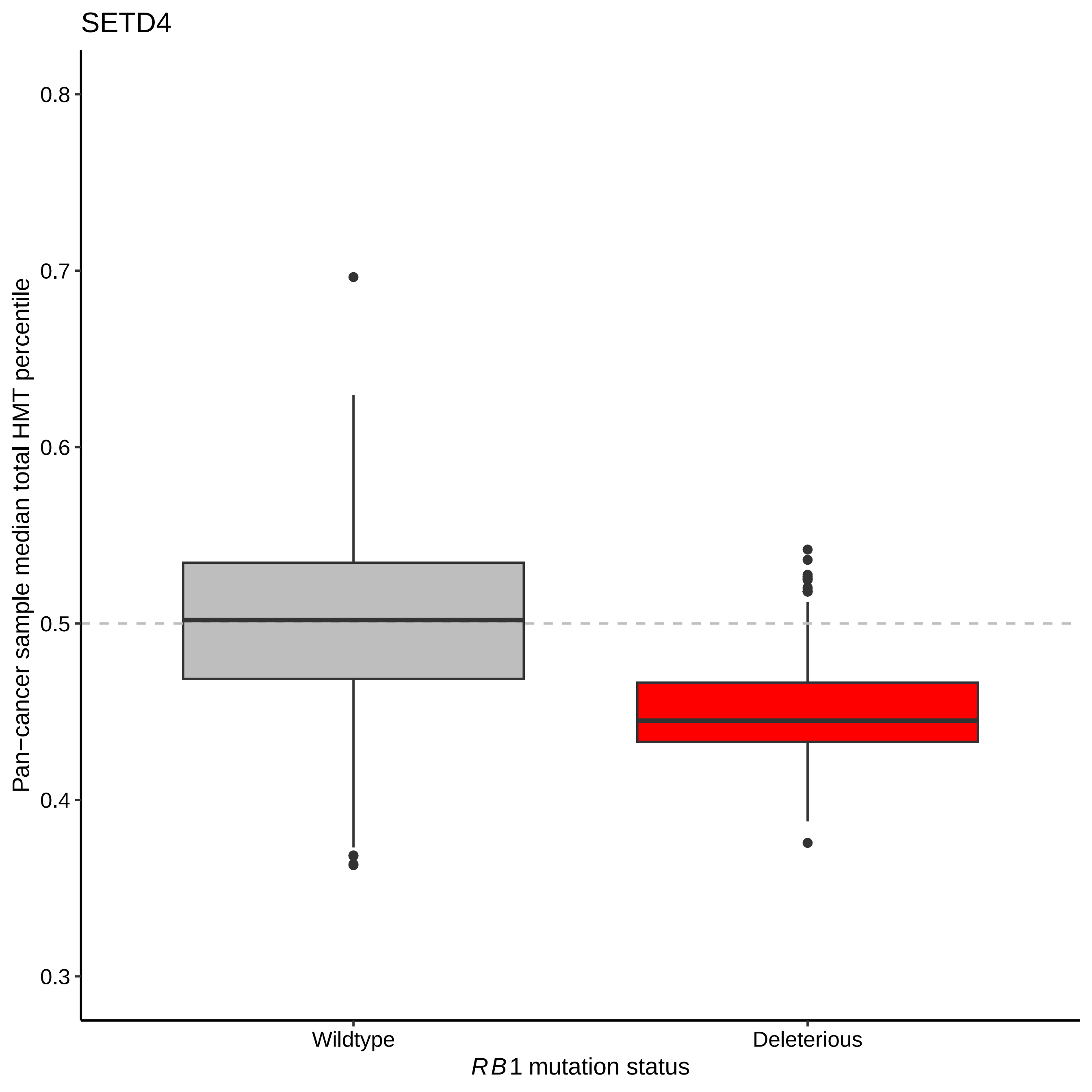

SETD4
Pan−cancer sample median total HMT percentile
0.8
0.7
0.6
0.5
0.4
0.3
Wildtype
Deleterious
RB1 mutation status

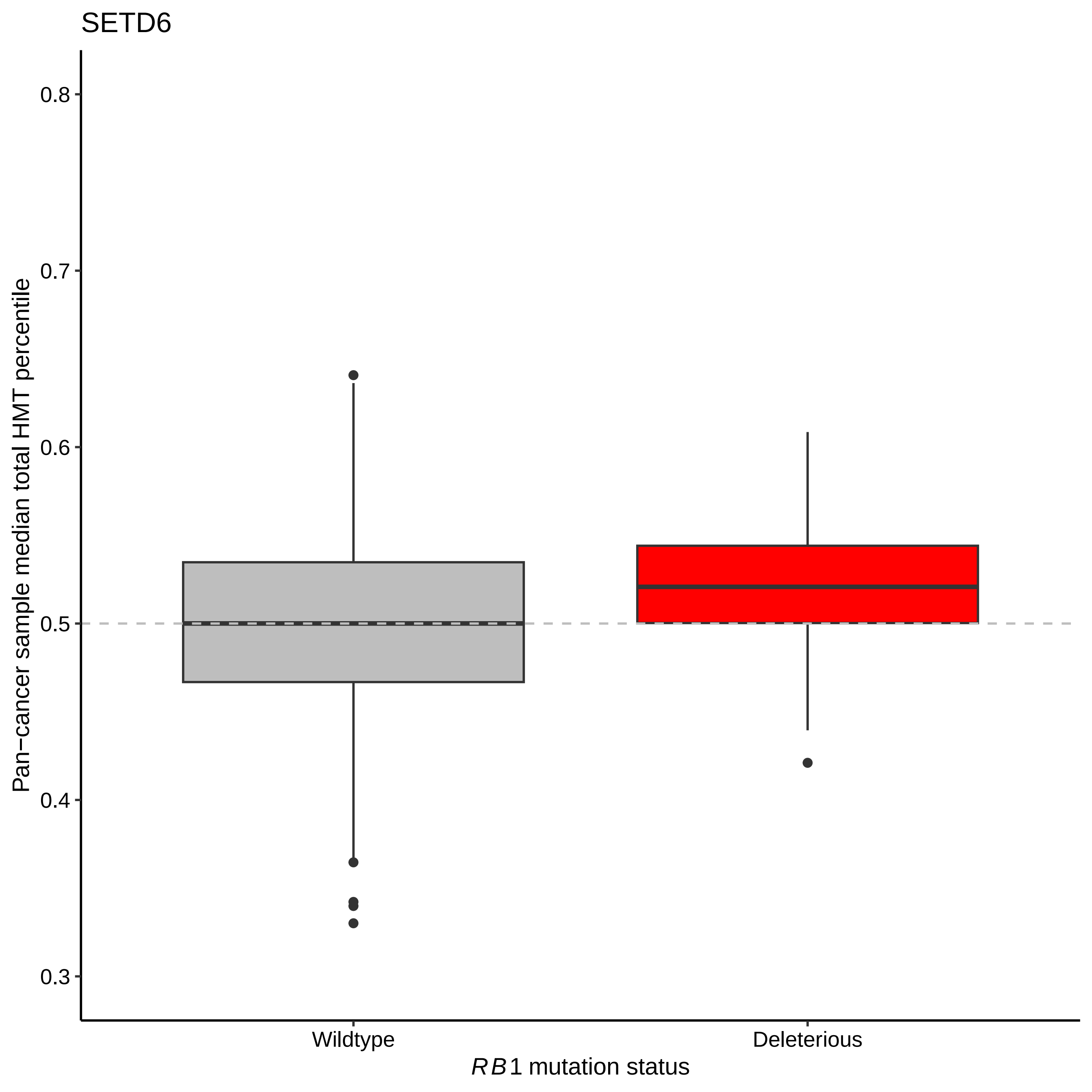
SETD6
0.8
0.7
0.6
0.5
0.4
0.3
Pan−cancer sample median total HMT percentile
Wildtype
Deleterious
RB1 mutation status

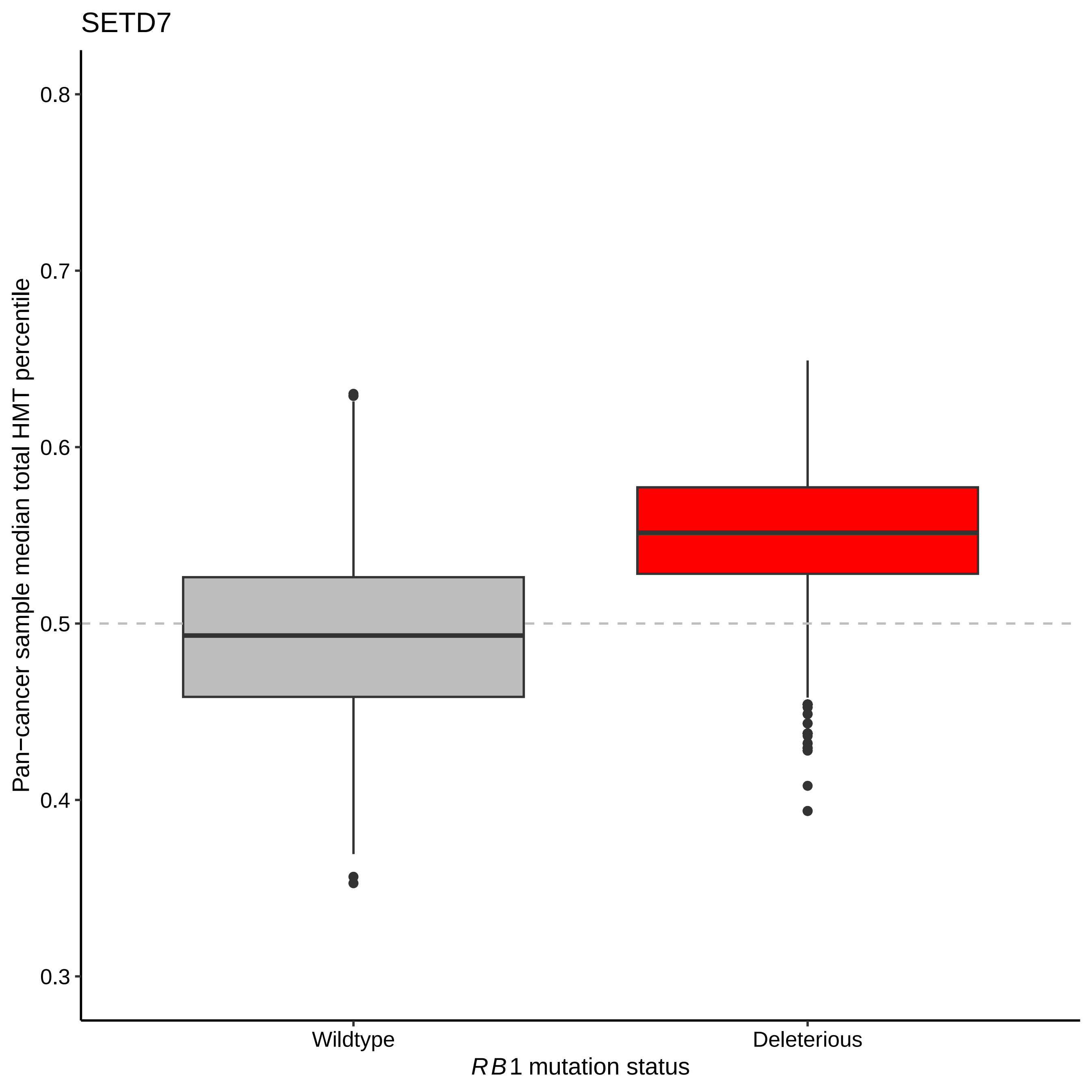
SETD7
Pan−cancer sample median total HMT percentile
0.8
0.7
0.6
0.5
0.4
0.3
Wildtype
Deleterious
RB1 mutation status

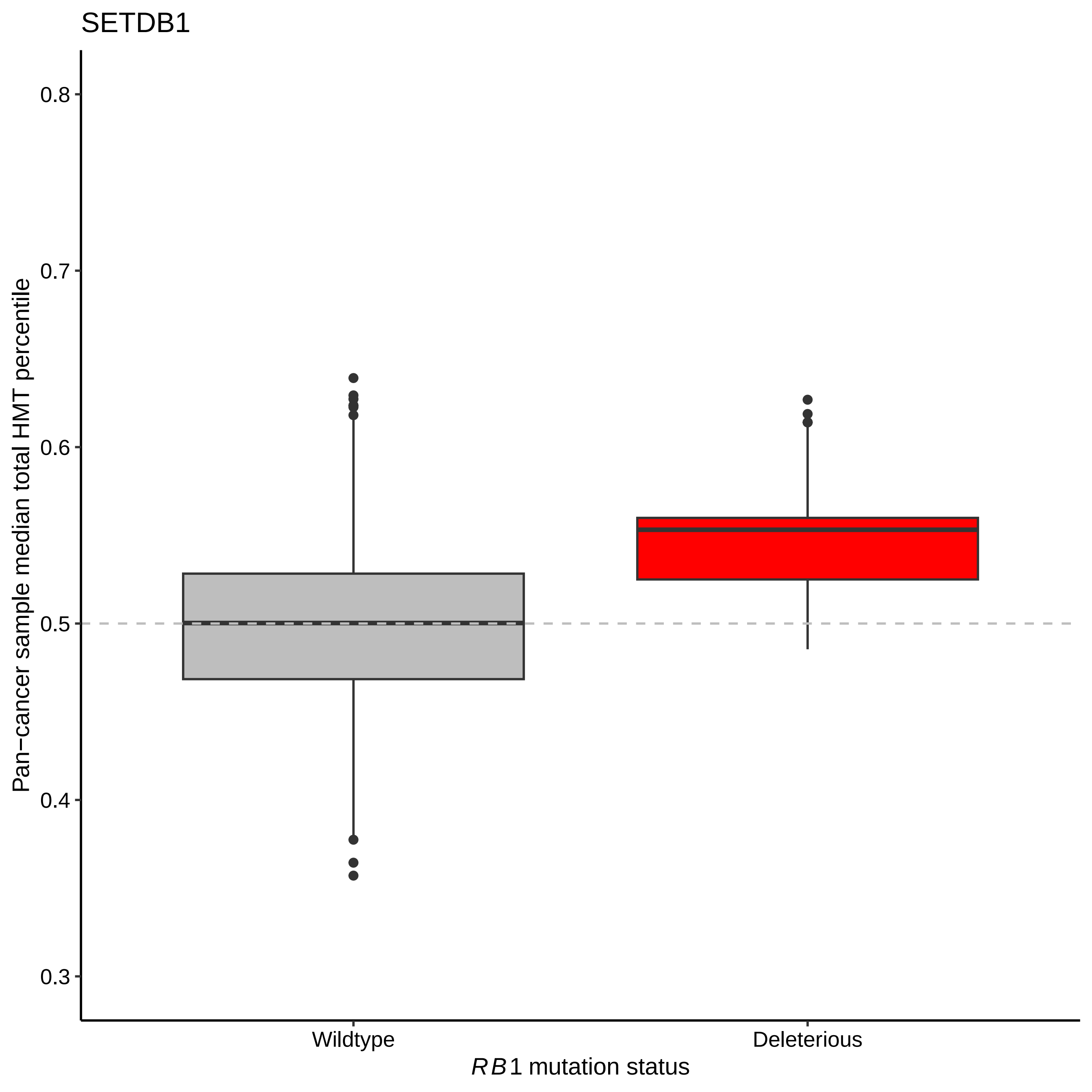

SETDB1
Pan−cancer sample median total HMT percentile
0.8
0.7
0.6
0.5
0.4
0.3
Wildtype
Deleterious
RB1 mutation status

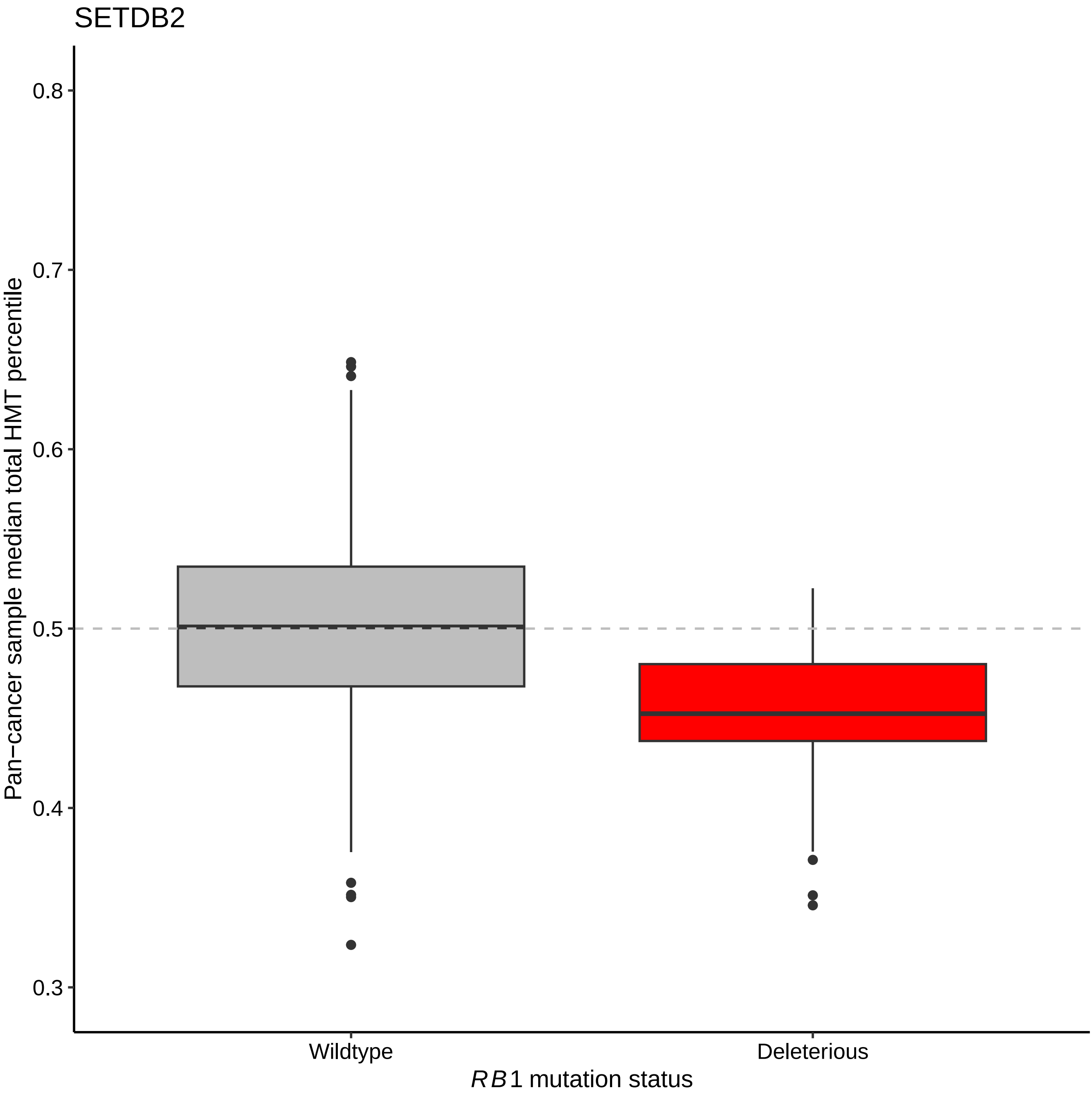

SETDB2
Pan−cancer sample median total HMT percentile
0.8
0.7
0.6
0.5
0.4
0.3
Wildtype
Deleterious
RB1 mutation status

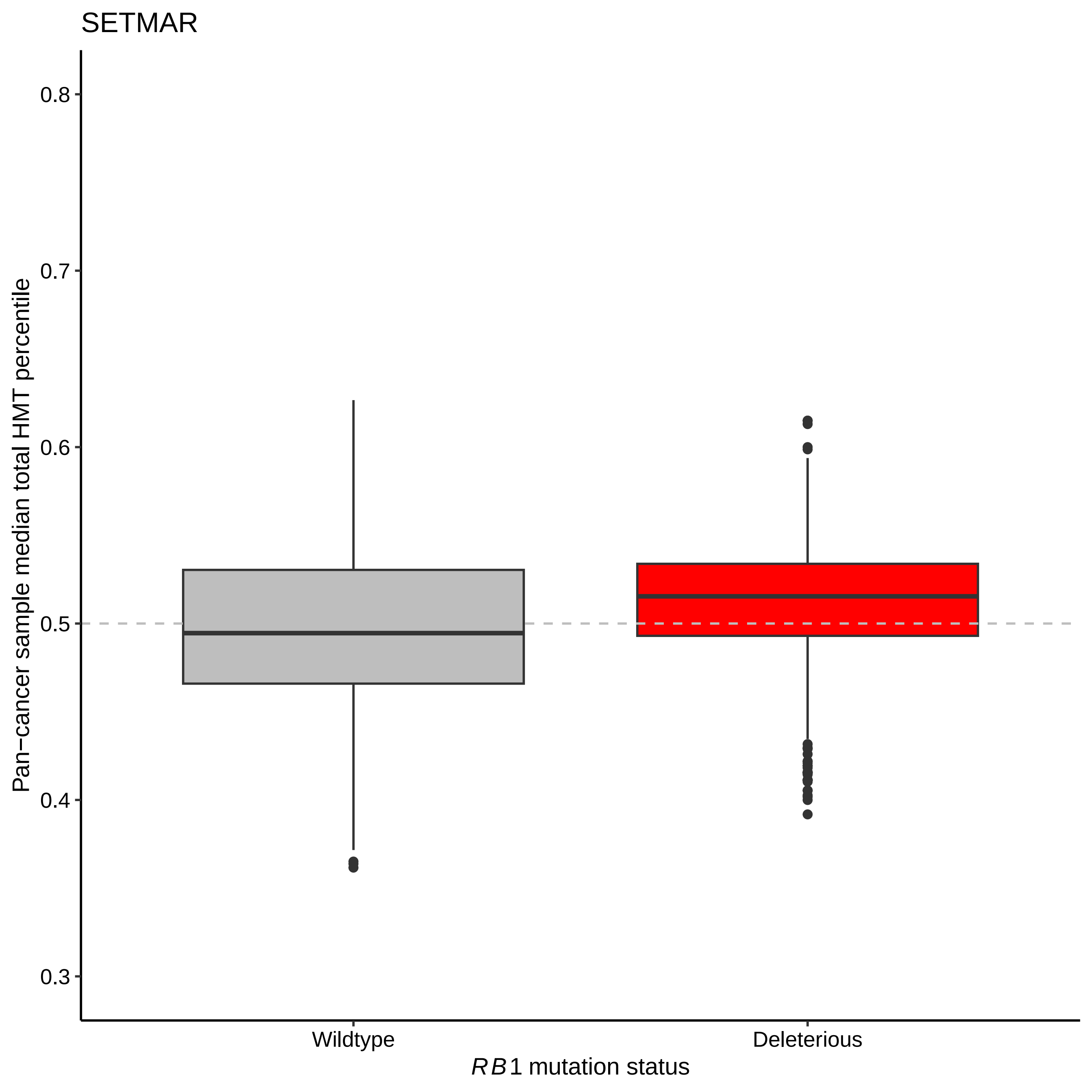
SETMAR
Pan−cancer sample median total HMT percentile
0.8
0.7
0.6
0.5
0.4
0.3
Wildtype
Deleterious
RB1 mutation status

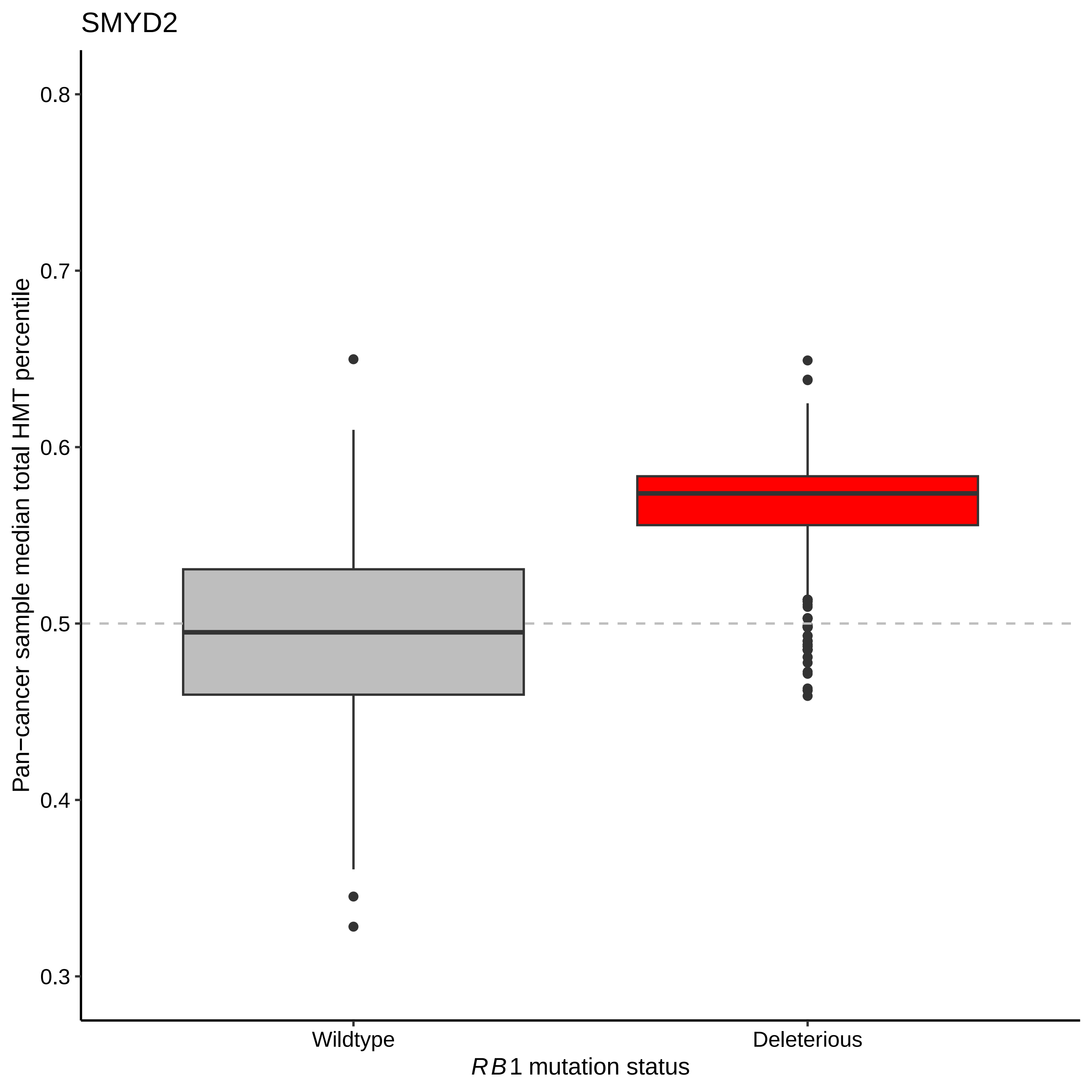
SMYD2
Pan−cancer sample median total HMT percentile
0.8
0.7
0.6
0.5
0.4
0.3
Wildtype
Deleterious
RB1 mutation status

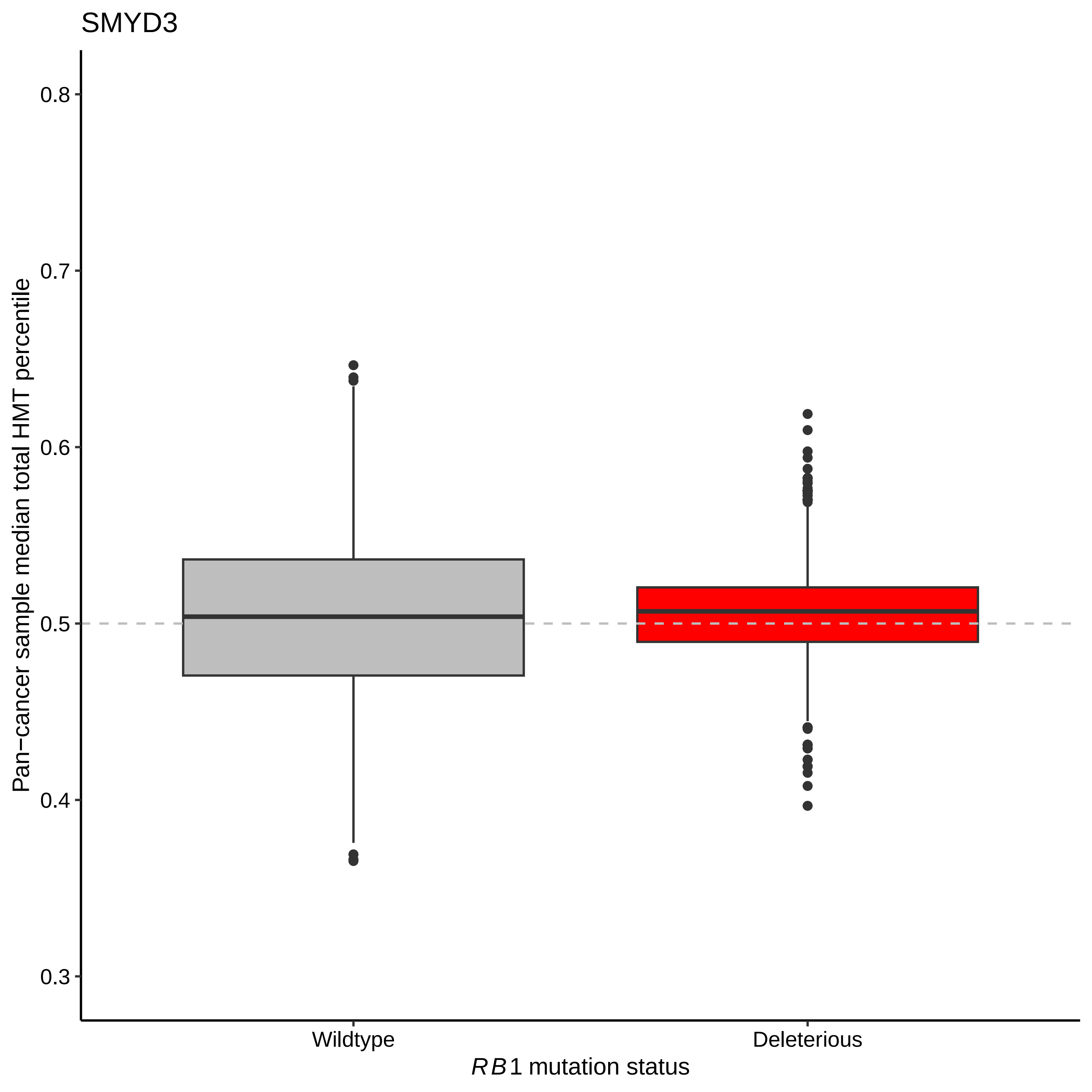

SMYD3
0.8
0.7
0.6
0.5
0.4
0.3
Pan−cancer sample median total HMT percentile
Wildtype
Deleterious
*RB1* mutation status

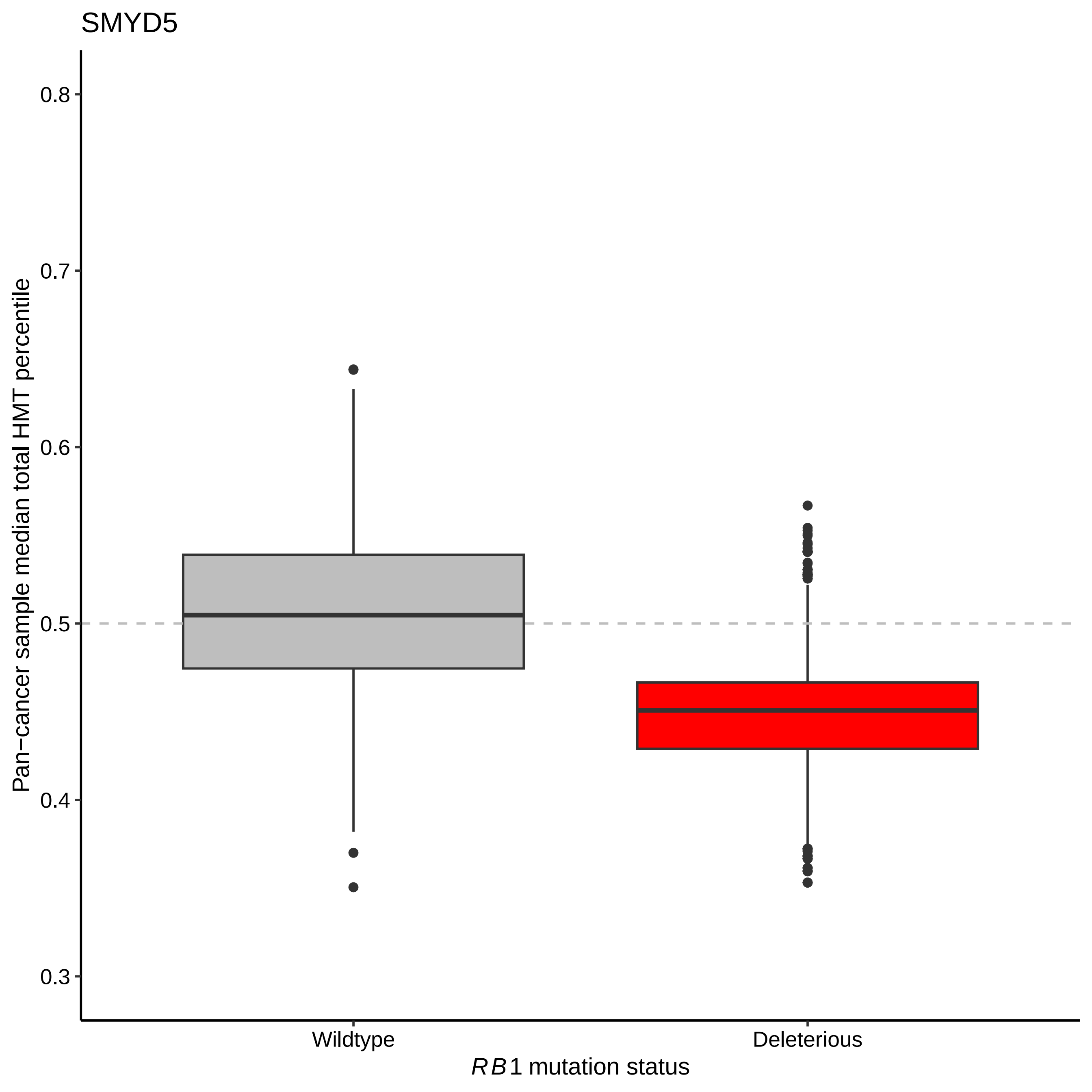

SMYD5
Pan−cancer sample median total HMT percentile
0.8
0.7
0.6
0.5
0.4
0.3
Wildtype
Deleterious
RB1 mutation status

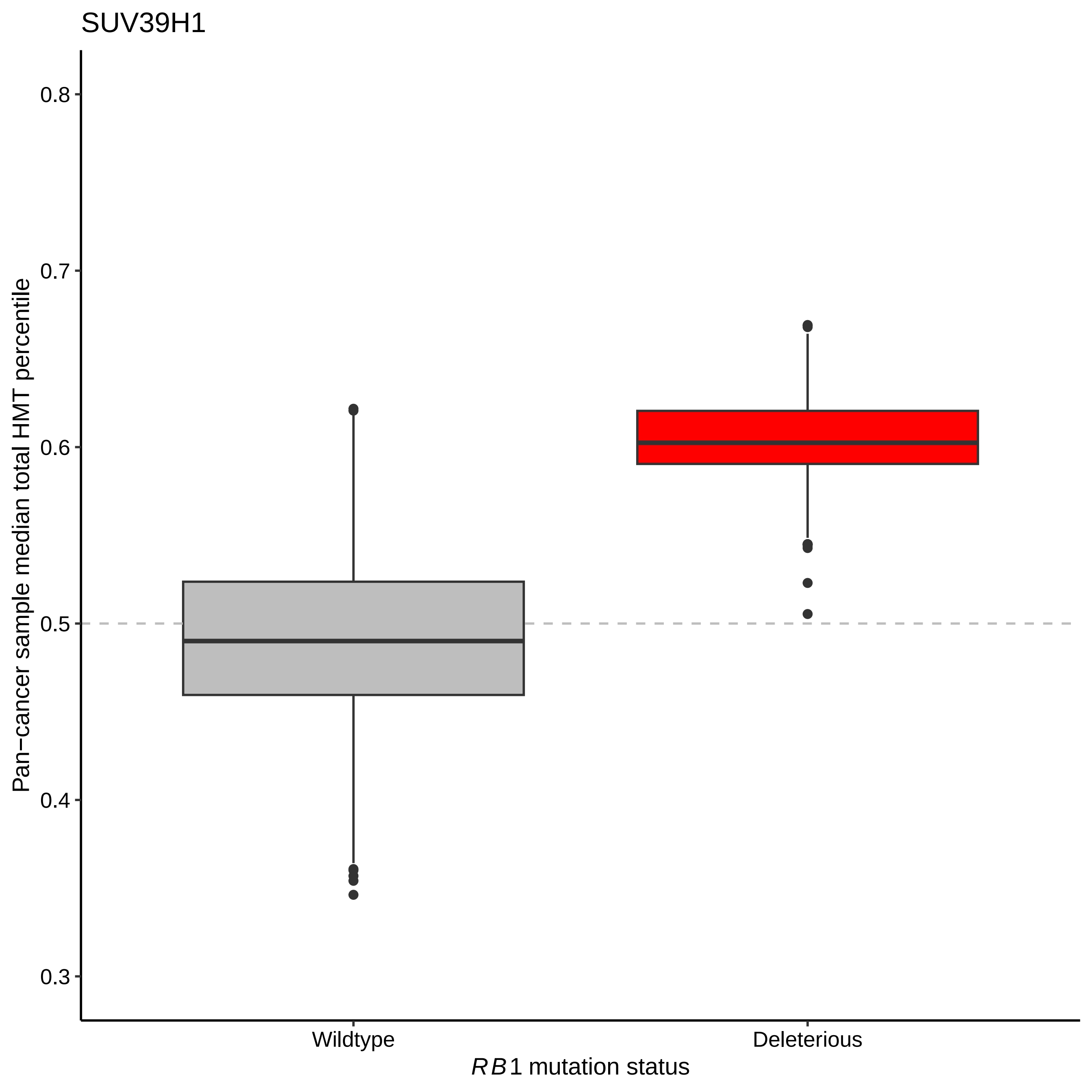

SUV39H1
Pan−cancer sample median total HMT percentile
0.8
0.7
0.6
0.5
0.4
0.3
Wildtype
Deleterious
RB1 mutation status

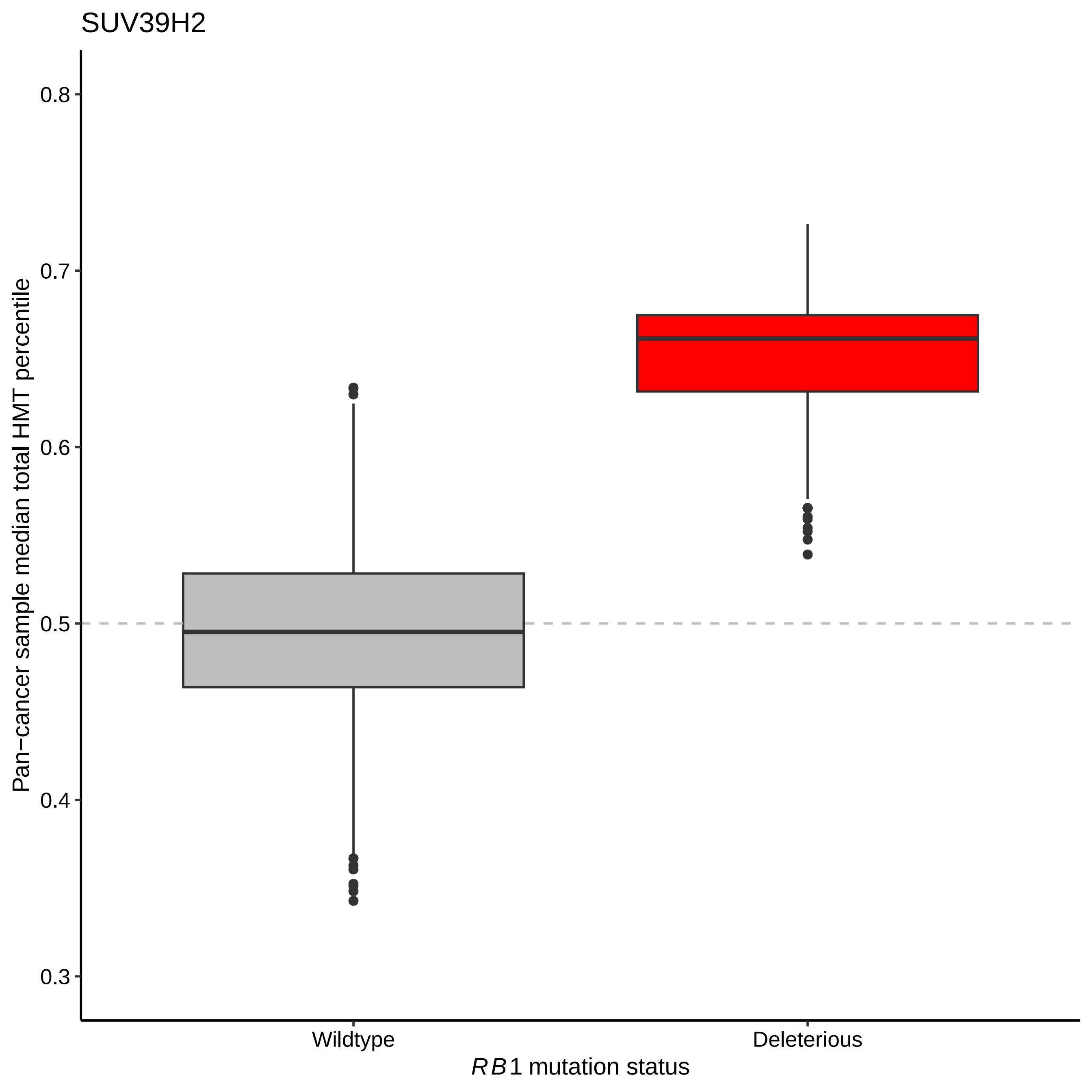

SUV39H2
Pan−cancer sample median total HMT percentile
0.8
0.7
0.6
0.5
0.4
0.3
Wildtype
Deleterious
RB1 mutation status
